# Translation Inhibiting Antibiotics Induce an asRNA Regulating Purine Metabolism in *S. pneumoniae* TIGR4 via an ISL3 Insertion Derived Hybrid Promoter

**DOI:** 10.64898/2026.04.28.721496

**Authors:** İrem Özkan, Tim vanOpijnen, Michelle M. Meyer

## Abstract

Translation inhibiting antibiotics (TIA) are an important class of antibiotics that elicit a wide variety of transcriptional responses. In this work we characterize the transcriptional responses of *Streptococcus pneumoniae* TIGR4, via RNA-seq, 5’-, and 3’-end mapping, to sub-MIC levels of three TIA, chloramphenicol, tetracycline, and kasugamycin. We find that kasugamycin (initiation inhibitor) displays a distinct transcriptomic profile compared to chloramphenicol and tetracycline (elongation inhibitors). However, genes in nucleotide metabolism were consistently downregulated. We also detected a TIA induced antisense transcript complementary to downregulated genes for purine metabolism, which we term SP_0835as. Mutating the promoter for SP_0835as eliminates both induction of SP_0835as and repression of the complementary genes in response to chloramphenicol. The mutated strain also displays slower growth rates than control strains in conditions that strongly induce SP_0835as, implying biologically relevant activity. Furthermore, the SP_0835as promoter straddles the boundary between SP_0835 and an insertion sequence (ISL3) immediately downstream. A survey of *S. pneumoniae* genomes shows the promoter is present in several phylogenetically distinct strain clusters. However, other common laboratory strains of *S. pneumoniae* lack the promoter, SP_0835as expression, and TIA repression of the complementary genes. Thus, SP_0835as represents a nascent RNA regulator that significantly remodels the metabolic response to TIA.

## INTRODUCTION

Bacteria utilize a variety of different RNA regulatory mechanisms to control gene expression in response to environmental changes (1), with mechanisms such as early termination of transcripts, inhibition of translation, or altered degradation rate, allowing fine tuning of gene expression (2-4). Such non-coding RNA (ncRNA) regulators play a particularly important role as they execute their regulatory functions without undergoing translation and thus may allow swifter more agile adaptive responses (1,5). Furthermore, there is growing evidence that regulatory RNAs play a crucial role in bacterial responses to environmental fluctuation, including mediation of antibiotic resistance (1,6), and such RNAs are considered drug targets (6,7). Despite these roles, many regulatory ncRNAs are narrowly distributed to only a few species (8-11), and our understanding of how such elements arise and are ultimately integrated into a host’s metabolic response network is still exceedingly limited.

<u>T</u>ranslation Inhibiting <u>A</u>ntibiotics (TIA) have been a mainstay for antibacterial therapeutics since the golden age of antibiotics (12) and remain a target for antibiotics in clinical development (13). Yet, transcriptomic changes in response to TIA appear quite diverse across different species (14-16). The high antibiotic concentrations used in clinical settings act as a strong selective force to accelerate the emergence of resistant bacterial strains. However, variable drug diffusion and metabolism also cause differences in drug concentration throughout a patient (17), and in natural environments bacteria often encounter antibiotic concentrations lower than the <u>M</u>inimum <u>I</u>nhibitory <u>C</u>oncentrations (MIC) of the drug (18,19). Antibiotics function as stressors at sub-MIC levels, inducing adaptive responses that facilitate the selection of resistant mutants (18-21).

In this work, we assessed how sub-MIC levels of TIA alter gene expression and transcript boundaries in the opportunistic pathogen *S. pneumoniae* TIGR4 using RNA-seq and 3’- and 5’-end sequencing. We show that elongation (chloramphenicol and tetracycline) and initiation (kasugamycin) inhibitors elicited responses that increase in similarity with both longer exposure time and higher drug concentration, with some metabolic processes, notably nucleotide metabolism, downregulated in all three antibiotics. Our analysis of transcript boundaries suggests that most high confidence transcription start and termination sites are maintained in the presence of sub-MIC TIA. However, we also observed several antisense transcripts induced by sub-MIC TIA, the most prominent of which is complementary to genes in purine salvage and precursor biosynthesis (SP_0828-SP_0835).

Assessment of gene expression at SP_0828-SP_0835 indicates that transcription of the asRNA, which we term SP_0835as, is anti-correlated with levels of the coding genes on the complementary strand. Furthermore, a strain carrying mutations to the promoter for SP_0835as eliminates both transcription of the asRNA and the downregulation of purine precursor biosynthesis and salvage genes. This strain also displays a significant growth defect in the presence of sub-MIC TIA antibiotics, suggesting biologically relevant regulatory functionality. However, the promoter driving SP_0835as transcription straddles the end of SP_0835 and an insertion sequence (ISL3) that is present at this locus. An analysis of complete *S. pneumoniae* genomes identified the promoter in a handful phylogenetically distinct strain clusters. Thus, in addition to differences in gene content, differences in regulatory elements such as asRNAs, may play a substantial role in the strain-to-strain differences in responses to antibiotics observed between strains within a given bacterial species. These results support the growing view that RNA regulators constitute a key layer in bacterial stress responses, enabling coordinated modulation of gene expression within core metabolic networks, with impacts on bacterial fitness (22). SP_0835as also provides a glimpse into how insertion elements remodel the genome to generate new regulatory features that integrate with existing metabolic responses to subtly alter phenotype.

## METHODS

### Strains

*S. pneumoniae* strain TIGR4 was used for the primary study. A derivative strain lacking expression of SP_0835as was created via replacement of the native promoter with mutations to the RpoD recognition sequence accompanied by a kanamycin cassette. To create this strain the genomic region proximal to the mutation was amplified with ∼1kb flanking regions from *S. pneumoniae* TIGR4 genomic DNA (Supplementary Figure S1). Mutation to the RpoD recognition site of the promoter was introduced via PCR (Supplementary Table S1). The two flanking regions were assembled with a kanamycin resistance cassette using HiFi DNA assembly (New England Biolabs), and transformed into *S. pneumoniae* TIGR4 to replace the native locus creating mP_SP_0835as. Transformants was screened for kanamycin resistance and were further confirmed via Sanger sequencing. A control strain was created with just the addition of kanamycin cassette but no mutation to the promoter sequence (Wt^Kan^).

### Minimum inhibitory concentration (MIC) Determination and Growth Rate Calculation

Minimum inhibitory concentrations (MIC) of chloramphenicol, kasugamycin and tetracycline were determined in the presence of increasing concentrations of antibiotics for *S. pneumoniae* TIGR4 strain by growth curves (Agilent BioTek BioSpa). Bacteria were grown to mid-exponential phase (OD_600_ ∼0.5) and diluted to OD_600_ ∼0.003 in Semi-Defined Minimal Media (SDMM) supplemented with 2-fold increasing concentrations of either antibiotic. OD_600_ was measured every 30 minutes for 16 hours to create growth curves. 96 well plate MIC determination was followed by MIC determination in tall culture tubes to ensure accuracy for collection conditions. MIC values for chloramphenicol, kasugamycin and tetracycline was determined to be 4 µg/ml, 500 µg/ml and 0.5 µg/ml, respectively (Supplementary Figure S2). To determine growth rates for mP_0835as, Wt^kan^, and TIGR4 in the presence of sub-MIC chloramphenicol, tetracycline, levofloxacin, and ciprofloxacin, similar growth curves were performed with two-fold decreasing concentrations of antibiotic starting at 2x the MIC. MICs for levofloxacin and ciprofloxacin were 2.5 µg/mL and 2 µg/mL respectively. Growth curves were analyzed with GrowthCurver to determine the growth rates (23).

### Growth Conditions and RNA preparation

For RNA-seq, 3’-end Seq, and 5’-end Seq library preparations, *S. pneumoniae* TIGR4 was grown in SDMM until initial OD_600_ of 0.5. Samples were diluted to OD of 0.1 in fresh SDMM, and then split into 3 different conditions: <u>N</u>o <u>d</u>rug <u>c</u>ontrol (NDC), ¼ x MIC, and ¾ x MIC of antibiotic treatment with 3 technical replicates for each condition. Samples were then incubated at 37°C, 5% CO_2,_ and 10 mL of each culture harvested by centrifugation (5000 rpm, 10 min at 4°C) at 60 and 90 minutes. Samples were flash frozen in a dry ice ethanol bath and stored at -80°C until RNA extraction. Total RNA isolation was performed via RNeasy Mini kit (Qiagen). Total RNA integrity of each sample was measured (all RINe scores >8.0). TIGR4 (Wt), Wt^Kan^ and mP_SP_0835as were grown in the presence or absence of ¾ xMIC chloramphenicol, and RNA isolated as described above.

### RNA-seq Library Preparation

For TIGR4 RNA-seq library preparation, RNAtag-seq protocol was followed as described (24,25). Briefly, 400 ng of total RNA was fragmented in FastAP buffer and treated with Turbo DNase. RNA adaptors with unique barcodes for each sample were ligated at the 3’-end of RNA samples followed by pooling. RNAseH based rRNA depletion was performed (26). Each RNA depletion reaction included up to 500 ng total RNA in 40 mM Tris-HCl (pH 7.5), 500 mM NaCl, and 6.7 mM of each depletion probe (Supplementary Table S2). Hybridization was performed with a hold at 95 °C for 2 min, ramp at −0.1 °C/s to 45 °C, and a hold at 45 °C for 5 min. A pre-warmed (42°C) RNaseH mastermix (3 µL Hybridase Thermostable RNaseH (Lucigen), 0.5 µL of 1 M Tris-HCl (pH 7.5), 0.2 µL of 5 M NaCl, 0.4 µL of 1 M MgCl_2_, and 0.9 µL of DEPC-treated water) was added after hybridization, and the solution mixed and incubated at 45 °C for 30 minutes. Reactions were adjusted to 40 µL using nuclease-free water prior to two sequential cleanups using Agencourt RNAClean XP magnetic beads (Beckman Coulter) according to the manufacturer’s instructions. After elution, 25 µL of the eluate was subjected to a second 2× SPRI cleanup, followed by elution in 15 µL water. First strand cDNA synthesis was performed with AffinityScript reaction kit and universal Illumina index primers were attached to each cDNA library pool by PCR for 17 cycles. Final size distribution and the sample concentration was retrieved by Qubit dsDNA HS Assay kit and dsDNA HS D1000 Tapestation kit. Samples were sequenced on an Illumina NextSeq500 using the v2 reagent kit. For TIGR4 (Wt), Wt^Kan^ and mP_SP_0835as samples, strand specific RNA-seq library preparation and sequencing were conducted by SeqCenter (Pittburg, PA).

### 5’- and 3’-end Seq Library Preparation

1 µg of the total RNA was divided into Tobacco Acid Pyrophosphates (TAP) treated (processed) and untreated (unprocessed) samples. All the samples were then treated with DNase, cleaned with magnetic SPRI beads and eluted with nuclease free water. Barcoded RNA adaptors were ligated to 5’-ends of each RNA sample and subsequent library preparation performed as described previously (27). Briefly, treated and untreated samples were fragmented in the FastAP buffer and pooled separately. rRNA depletion was performed as described above. First strand cDNA synthesis was performed with AffinityScript reaction kit and universal Illumina index primers were attached to each cDNA library pool via 17 cycles of PCR. Final size distribution and the sample concentration determined and samples sequenced as above.

3’-end Seq libraries were prepared as described previously (28). In brief, 2 µg of total RNA was DNase treated with TURBO DNase buffer and ligated to 3’-end RNA adaptors. Samples were then fragmented by using Ambion RNA fragmentation buffer and pooled. rRNA depletion was performed as described above. First strand cDNA synthesis was performed with AffinityScript reaction kit and universal Illumina index primers were added to each cDNA library pool by PCR for 17 cycles. Final size distribution and the sample concentration determined and samples sequenced as above.

### RNA-seq read processing

The single-end sequencing reads were processed and mapped to *S. pneumoniae* TIGR4 (NC_003028.3) via an internal pipeline (https://github.com/jsa-aerial/aerobio). Briefly, sequencing data was mapped to the genome via Bowtie2 (29) to generate a sorted and indexed BAM files for each sample. Raw reads generated in a previous study of *S. pneumoniae* TIGR4 and Taiwan 19F (15) was obtained from (BioProject PRJNA542628). Specific accession numbers are in Supplementary Table S3.

### Differential Expression and Pathway Enrichment

DESeq2 and pathway enrichment analyses were performed via featureCounts (-s 1 and -s 2 for stranded) for RNA-seq samples [26]. Genes that have log2fold change > 1 (or < -1) and padj < 0.05 were considered significant for RNA-seq. Differential expression results from DESeq2 were analyzed for pathway enrichment analysis custom scripts (available at https://github.com/iremozkanRNA). Genes with |log_2_FoldChange| > 1 and adjusted p-value (padj) < 0.05 were classified as significantly regulated and categorized based gene category. Missing annotations were assigned the value “Unknown.” Enrichment of significant genes within each functional category was assessed using a hypergeometric test in R, with the total gene set as background. p-values were corrected for multiple testing using the Benjamini–Hochberg method, and categories were ranked by adjusted p-value. Results including category name, overlap count, category size, and associated statistics were compiled into a single output file (Supplementary Table S4).

### Identification of 3’ -ends and High Confidence Transcription Termination Sites

3’-end peaks with signals significantly above background were called via PIPETS (30) from combined 3’-end sequencing data using default parameters with mapped .bed files (converted from bam files using bedtools (31)). The 3’-end annotation used a hierarchical approach to categorize 3’-ends with priority given to 3’-UTR designation (Supplementary Figure S3A). This approach yielded one to two thousand 3’-ends for each condition, with a large proportion (30-50%) identified within coding sequence (Supplementary Figure S4A, Supplementary Table S5). *S. pneumoniae* lacks the Rho termination factor, and thus relies heavily on intrinsic termination (32,33). To develop a set of high confidence termination sites we assessed each peak to determine whether it was associated with features of intrinsic termination (strong stem followed by poly uridine sequence). Briefly, 50 nt immediately upstream of each 3’ peak coordinate was extracted and secondary structure analysis performed using RNAfold (Vienna RNA package version 2.5.1) (33) and polyuridine sequences (5 of 7 consecutive unpaired bases following the stem) identified. Peaks whose upstream sequence possessed a free-folding energy <= 10 kcal/mol and were accompanied by an unpaired polyuridine sequence were considered associated with features of intrinsic termination (custom scripts at https://github.com/iremozkanRNA). The resulting peaks were much more likely to be in the 3’-UTR (Supplementary Figure S4B), and further assessment of RNA-seq coverage in regions before and after the 3’-end coordinate shows that these sites are also strongly associated with a drop in RNA-seq coverage, further suggesting they are high confidence termination sites (Supplementary Figure S4C,D).

### Identification of Transcription Start Sites

To differentiate between the primary (5-PPP) and processed or cleaved (5’-P) transcripts (34-36) we generated 5’-end seq libraries using both Tobacco Acid Pyrophosphatase (TAP) treated (converts triphosphate into monophosphate) and untreated (monophosphorylated) RNA samples and used the TSSpredator pipeline to identify TSS (37,38). 5’-ends reads from each condition and NCBI derived annotation file (.gff) was used to classify each transcription start site. Primary start point sites were identified via the ratio of 5’-phoshorylated/ all reads (e(i)processed / e(i)unprocessed) (39). Transcription start sites were categorized based on their position relative to coding sequences (Supplementary Figure S3B, custom scripts at https://github.com/iremozkanRNA). High confidence transcription start sites were identified based on the presence of RpoD consensus promoter motifs at the -10 and -35 elements (Supplementary Table S6) as *S. pneumoniae* has only two sigma factors, SigA (homolog of RpoD) and ComW (40), where ComW is only utilized when competence is induced (41). To call TSS for asRNA, PIPETS was used to identify significant peaks in both processed and unprocessed 5’-end sequencing data [41]. Significantly called peaks were then used to retrieve ratios (processed/unprocessed) from the raw 5’-end sequencing data (raw counts > 10 was used). Sites with ratios over 2 were counted as significant TSS (27) (Supplemental Table S7).

### Identification of asRNAs and strand-specific RT-PCR

To compare transcript abundance quantifications for sense vs. antisense transcription, raw counts from RNA-seq data were generated using DESeq2 featureCounts under two settings: strand-specific counting (-s 1, reads mapping to anti-sense transcript or -s 2, reads mapping to sense transcript) and strand-unsplit counting (no -s flag, total counts) (42,43). To avoid false identification of spurious anti-sense transcripts derived from library preparation protocols (cDNA synthesis & amplifications), we employed binomial testing methods to determine whether observed read distribution deviated significantly from an expected random allocation (42,43). Specifically, for each gene, the number of antisense mapping reads (featureCounts -s 2, successes) was tested against the number of sense mapping reads (featureCounts -s 1, trials) using a one-tailed binomial distribution test in R with the probability of success under the null hypothesis set to (<0.01). The resulting p-values for each gene across three replicates were combined using Fischer’s method. Extreme p-values (< 1e-100) were capped to prevent numerical overflow in the logarithmic calculations. To account for multiple hypothesis testing across all genes, the combined p-values were subsequently adjusted using the Benjamini-Hochberg method to control the false discovery rate (Supplementary Table S8).

To confirm that SP_0835as exists as a single transcript, strand specific RT-PCR was conducted on RNA samples collected from cells grown at ¾ x MIC tetracycline and chloramphenicol as above. RNA samples were treated with TURBO DNAse and reverse transcription performed using Induro Reverse transcriptase (NEB) with a primer complementary to the 5’ UTR of SP_0828 containing a 5’-tag sequence (Supplementary Table S1) alongside a no RT control lacking the enzyme. These reactions were treated with Hybridase Thermostable RNAseH (Lucigen) and amplified using Q5 polymerase (NEB) with a forward primer corresponding to the introduced tag sequence and reverse primers corresponding to the 3’-end of SP_0835 or the region just upstream of the predicted promoter (within SP_0836). For size comparison, TIGR4 genomic DNA was amplified with the reverse transcription primer and the same two reverse primers.

### Comparative Genomics of *S. pneumoniae* Genomes

A 10 kbp sequence corresponding to the locus encompassing SP_0825-SP_0837 NC_003028.3/777000-787000 was blasted against each genome in a collection of *S. pneumoniae* strains sequenced via long-read sequencing in BioProject PRJNA514780 (44). Contiguous sequences encompassing the matched regions were extracted and aligned with MUSCLE. The region that failed to align with most other genomes within this contiguous segment and an additional 50 nt of flanking sequence (NC_003028.3/784329-785849) was blasted back to the *S. pneumoniae* TIGR4 genome and the results aligned as above to confirm insertion sequence boundaries.

To extend comparative genomics, all *S. pneumoniae* genome assemblies with assembly level “complete” or “chromosome” were downloaded from NCBI GenBank (accessed May 2026), yielding 366 assemblies. To identify the genomes carrying the ISL3 insertion creating SP_0835as promoter, the 36-bp query sequence spanning the SP_0835/IS element (NC_003028.3/ 784362 - 784398) was blasted (BLASTn) against all 366 assemblies. Hits with >90% identity and >80% query coverage were retained as junction-positive (Supplementary Table S9). Each junction positive genome was further verified to carry insertion at the purine metabolism locus (SP_0825-SP_0837) by confirming junction for the 10kb overlap from NC_003028.3/777000-787000.

Global Pneumococcal Sequencing Clusters (GPSCs) were assigned to all 366 genomes using PopPUNK v2.7.8 with the GPS_v10 reference database (45,46). To create a phylogenetic tree, all 15 junction-positive genomes and one random representative genome from each cluster was chosen. Genomes were annotated with Prokka v1.14.6 and core gene alignments were generated with Roary v3.13.0 using a 95% core gene threshold (47,48). Maximum-likelihood phylogenies were inferred with IQ-TREE v2.2.0 with a general time-reversible nucleotide substitution model and gamma-distributed rate heterogeneity (GTR-G4), with branch support assessed by 1,000 ultrafast bootstrap replicates (49). Extracted IS element sequences from the four confirmed, junction-positive GPSCs (GPSC31, GPSC32, GPSC23, GPSC70) were aligned using MAFFT v7.525 and pairwise nucleotide identities calculated from the alignment (50). Custom scripts were deposited to GitHub (https://github.com/iremozkanRNA/SP_0835as)

## RESULTS

### TIAs with diverse mechanisms elicit a similar downregulation of metabolic processes at higher sub-MIC concentration and longer time points

To assess transcriptome profile changes in response to sub-MIC translation inhibiting antibiotics (TIA), we performed RNA-seq with *S. pneumoniae* TIGR4 grown in the absence or presence of ¼ and ¾ x MIC for 3 different TIA, elongation inhibitors tetracycline (51) and chloramphenicol (52), and initiation inhibitor kasugamycin (53). To appraise the quality and consistency of our RNA-seq datasets, we performed principal component analysis (PCA) on the variance-stabilized transformed (VST) read counts at each time point (Figure 1A). We see that individual samples subjected to the same treatment cluster together with self-consistent replicates. Furthermore, the No Drug Control (NDC) samples collected alongside each individual antibiotic also cluster indicating that observed changes are directly attributable to antibiotic treatment. The kasugamycin samples systematically cluster separately from the other two antibiotics, indicating expression changes in response to inhibition of translation initiation may differ from those induced by inhibition of translation elongation.

**Figure 1:**
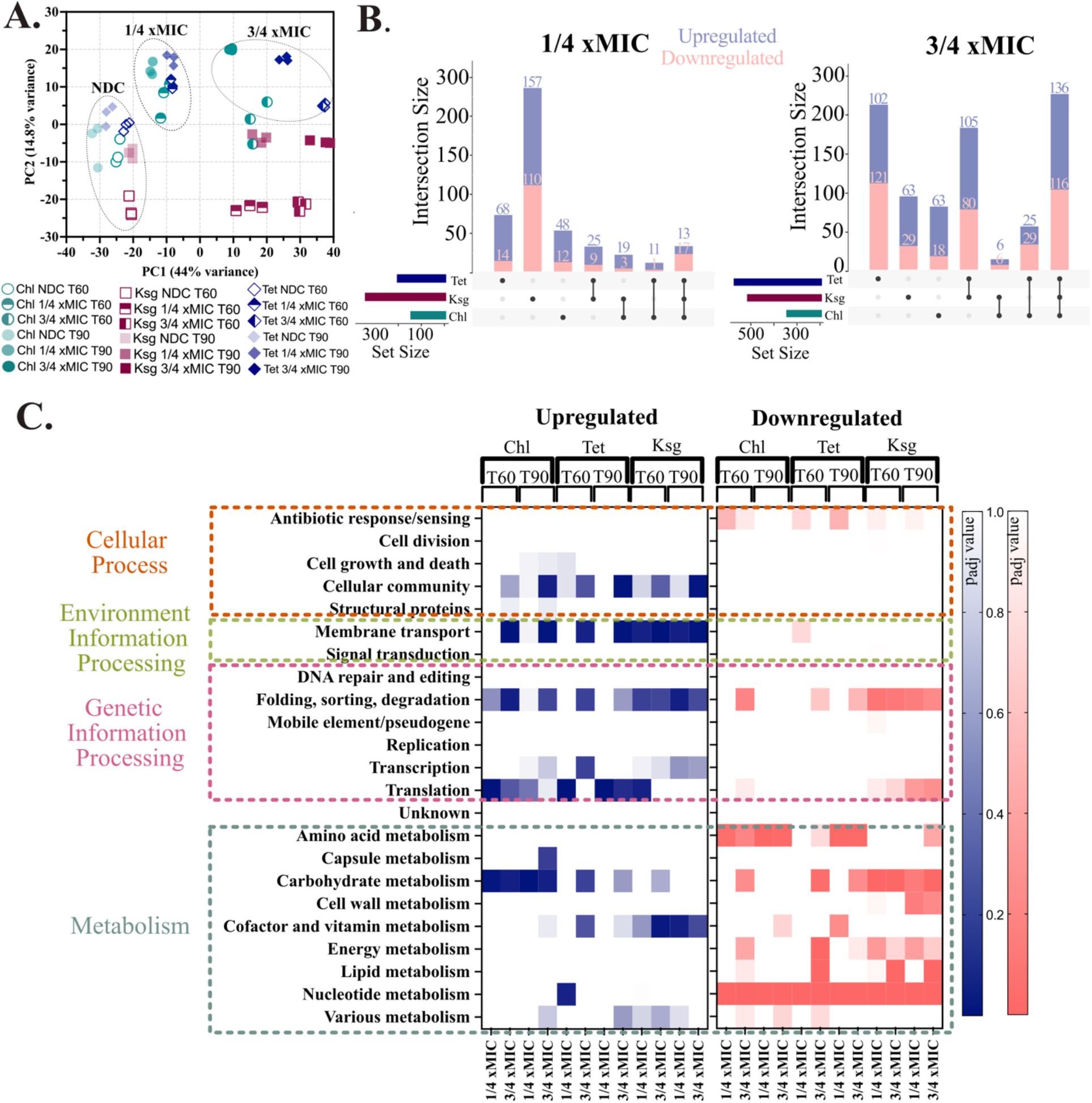
RNA-seq analysis of sub-MIC TIA treatments reveals overlapping transcriptional responses. **(A)** Principal Component Analysis (PCA) of RNA-seq data sets from *S. pneumoniae* TIGR4 strain grown in the presence of translation inhibiting antibiotics (TIA) treated with ¼ and ¾ x MIC for 60 (T60) and 90 (T90) minutes show clustering of replicates and differentiation of antibiotic treatments. PCA plot generated from variance-stabilized transformed (VST) read counts generated via DEseq2 (featureCounts). Clustering of untreated (NDC) samples, ¼ and ¾ x MIC treated samples for chloramphenicol and tetracycline is indicated. **(B)** Overlap of the differentially expressed genes in each antibiotic after 60 minutes exposure for both ¼ x and ¾ x MIC, blue bars represent upregulated and pink bars down regulated genes. (**C**) Pathway enrichment analysis for functional *S. pneumoniae* TIGR4 highlights metabolic down-regulation. Pathway enrichment for gene category panels was calculated from RNA-seq fold changes. Genes were considered significant if |log2FoldChange| > 1 and adjusted p-value < 0.05.

To determine general transcriptomic trends affected by antibiotic treatment, we analyzed each of our RNA-seq datasets relative to its matched NDC using DESeq2 (54). While ¾ xMIC samples typically show a greater response than ¼ xMIC samples, the responses are strongly correlated, especially for kasugamycin (R^2^=0.81 between ¼ xMIC and ¾ xMIC, Supplementary Figure S5, Table S4). In addition, for both chloramphenicol and tetracycline, longer exposure to antibiotics (60 vs. 90 minutes) increased the correlation between the two concentrations and the proportion of differentially expressed genes shared (Supplementary Figure S5). In comparing the responses to each antibiotic to each other we observe that ¼ xMIC kasugamycin elicits a strong response that is somewhat distinct, but at ¾ xMIC this difference dissipates (Figure 1B). At ¾ xMIC 60-65% of DE genes are shared amongst 2 or more conditions with ∼32% are shared by all three. Thus, we find that at ¾ xMIC, similar responses are elicited by all 3 TIA. Furthermore, chloramphenicol and kasugamycin share fewer DE genes than any other 2 antibiotic comparisons (Figure 1B).

To assess which pathways are affected by the antibiotic treatments, we performed pathway enrichment analysis separately with the up- and down-regulated gene sets (Figure 1C). We see that the majority of metabolic pathways were significantly downregulated upon sub-MIC TIA exposure. Most strikingly, nucleotide metabolism was downregulated in all the treatments regardless of the treatment time or concentration. Additionally, carbohydrate and lipid metabolisms were downregulated, especially for kasugamycin treatment (Figure 1C). Modification of nucleotide metabolism has been previously linked to changes in antibiotic susceptibility in several bacterial species (55-57). However, the consistent wide range of metabolic downregulation suggests a transcriptional response to conserve resources.

In addition to common metabolic downregulation, responses to individual antibiotics varied. For both chloramphenicol and tetracycline, genes involved in cellular community, membrane transport, translation and carbohydrate metabolism are significantly enriched in the upregulated category (Figure 1C) (58). For kasugamycin we observed significant enrichment of downregulated genes in folding, sorting and degradation pathways, which produce proteins responsible for the essential processes of quality control, delivery, and removal of other proteins within a cell. In comparing responses to different antibiotics, we observe that although tetracycline shares more individual DE genes with kasugamycin (Figure 1B), the gene categorization pattern displays more similarity with chloramphenicol (Figure 1C). The interplay of responses to the 3 antibiotics can also be observed from the PCA plot, where kasugamycin clearly separates from the other two antibiotics (Figure 1A). Together these results indicate that the response to kasugamycin, the initiation inhibitor, diverges somewhat from that of chloramphenicol and tetracycline, elongation inhibitors. Based on this analysis of the bulk RNA-seq data, we decided to assess transcript boundaries from two of the three antibiotics: chloramphenicol and kasugamycin.

### Transcript boundaries are largely stable in the presence of translation inhibiting antibiotics

Environmentally induced changes to transcription termination sites are relatively common in gram positive species, with cis-regulatory mechanisms such as riboswitches, t-boxes, and classic attenuators altering termination sites in response to specific metabolites, stresses, or antibiotics (59-61). Furthermore, the decoupling of transcription and translation in the presence of a translation inhibitor might be expected to alter sites of transcription termination (62). To characterize changes to transcription termination sites (TTS) under sub-MIC TIAs, we collected 3’-end sequencing data under chloramphenicol and kasugamycin treatments at ¼ xMIC and ¾ xMIC for 60 minutes alongside a No Drug Control condition for each antibiotic (28). We find that the high confidence termination sites (see Methods) largely overlap across the NDC and antibiotic conditions (Figure 2A). The overlap is comparable to that we observe for the two NDC samples collected alongside each antibiotic (Supplementary Figure S4E), and is consistent with the variability typically observed in 3’-end sequencing (63). Furthermore, manual inspection of sites that do appear uniquely in the antibiotic conditions indicated that most of them are present in the NDC, but fall below the limit of detection when using PIPETs with its default parameters. Thus, there do not appear to be substantial changes to transcription termination sites that impact gene expression in the presence of TIA (Figure 2A).

**Figure 2:**
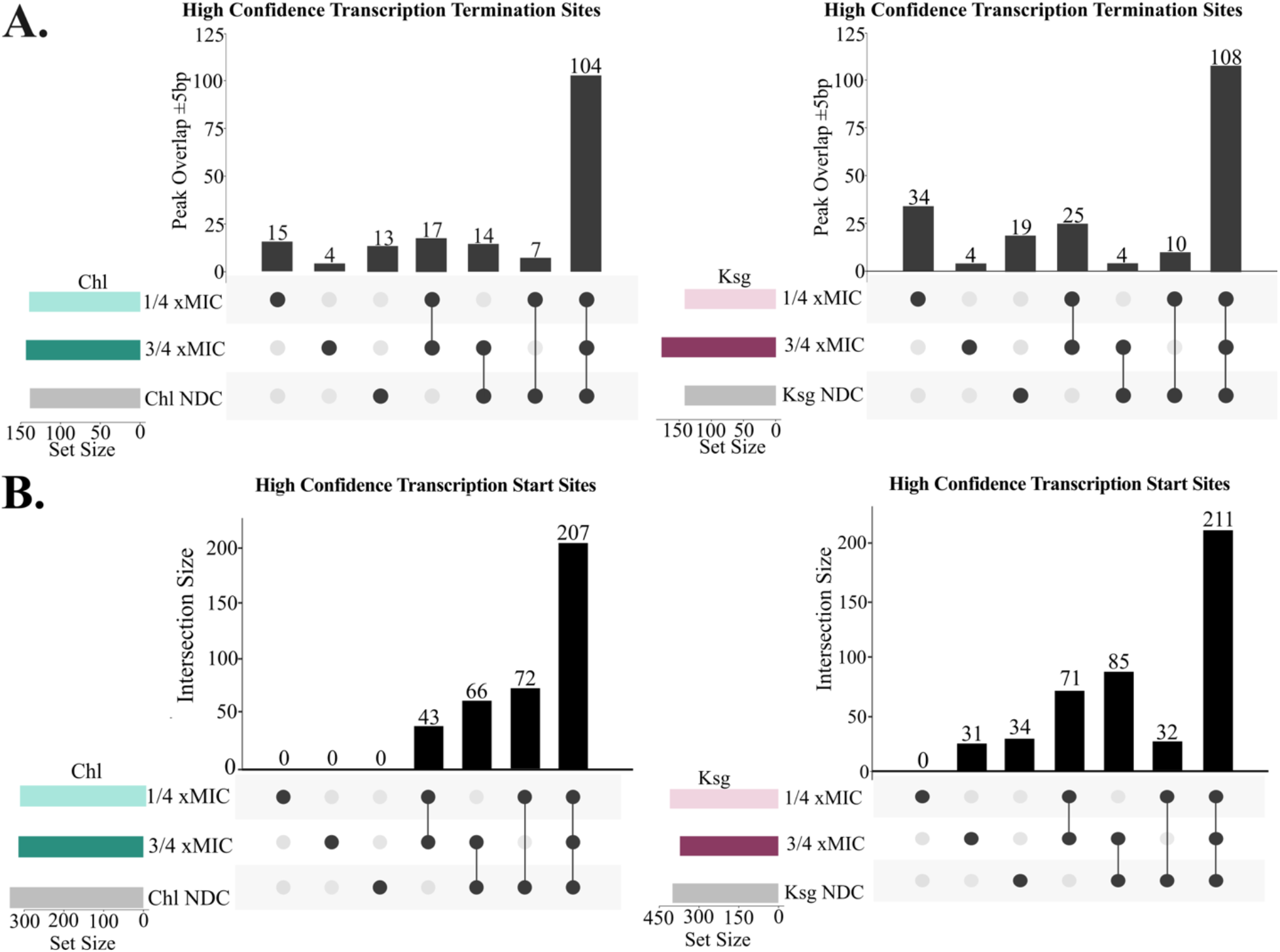
5’- and 3’-end sequencing of *S. pneumoniae* transcriptome under sub-MIC TIA treatment reflect only modest changes to high confidence transcription boundaries. **(A)** Overlap of high confidence *S. pneumoniae* transcription termination sites between low (¼ x MIC) and high (¾ x MIC) sub-MIC antibiotic TIA treatments at 60 minutes. **(B)** Overlap of *S. pneumoniae* high confidence transcription start sites between low (¼ x MIC) and high (¾ x MIC) sub-MIC antibiotic TIA treatments at 60 minutes.

Alongside our 3’-end sequencing data, we also collected 5’-end data to fully assess how transcript start sites (TSS) might change under antibiotic treatment. To assess TSS changes upon sub-MIC antibiotic treatment, we similarly analyzed TSS profiles for both antibiotic treatments. For both antibiotics, ∼80-90% of high confidence sites are shared by the NDC and at least one antibiotic (Figure 2B). Again, this is similar the rate at which sites are shared by the two NDC samples (Supplementary Figure S4F). These results show that high confidence TSS positions are largely stable in the presence of sub-MIC TIA treatments, suggesting that the sites of transcription initiation are broadly maintained under antibiotic stress, even as gene expression levels may change substantially

### Translation inhibition antibiotics induce antisense RNA expression

Antisense RNAs (asRNAs) originate from the opposite strand of protein-coding genes and may directly base-pair with mRNA sequences (7) to modulate gene expression by affecting mRNA stability, processing, and translation, or by altering transcription termination dynamics (64,65). However, past characterization of lactobacilli typically identified few antisense transcripts [32]. To detect whether *S. pneumoniae* possesses any antisense RNAs, we determined the ratio of antisense reads to total reads for each annotated gene. For each gene, we further assessed the likelihood that artifacts introduced by sequencing protocols or background transcription would result in spurious antisense reads using a binomial distribution test (42,43) (see Methods). We further considered that only genes where >50% of the total reads were antisense (ratios of antisense/total reads greater than 0.5) to be subject to asRNA-mediated modulations in gene expression (Figure 3A). In the absence of antibiotic (NDC) we find that between 22 and 33 genes show significant antisense transcription. However, in the presence of ¾ xMIC antibiotic, this number increases substantially to 73, 103, and 133 genes for chloramphenicol, kasugamycin and tetracycline, respectively (Figure 3A, Supplementary Table S8). Thus, under TIA treatment, *S. pneumoniae* induces antisense transcription over many genes.

**Figure 3:**
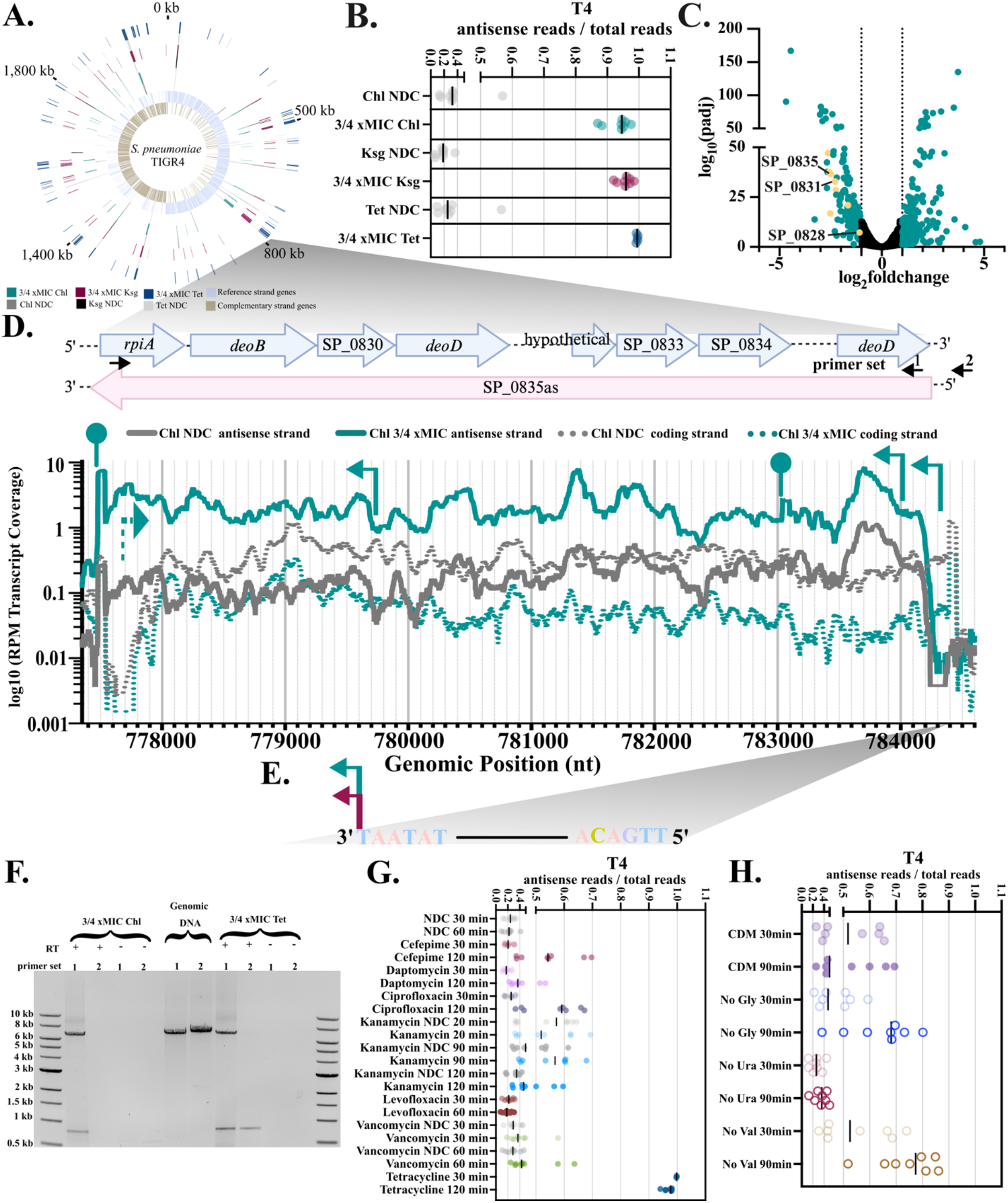
Combined RNA-seq, 5’- and 3’-end data support a TIA induced anti-sense RNA. **(A)** Assessment of RNA-seq data across all genes for antisense transcription. Binomial testing was used to identify significant antisense transcription over transcriptional noise for each gene assuming baseline success probability of 1% (42). Genes with antisense/total read ratios > 0.5 in each condition are displayed over the *S. pneumoniae* TIGR4 genome. **(B)** RNA-seq data antisense/total read ratios for SP_0828-SP_0835 gene cluster in presence of antibiotic (chloramphenicol, tetracycline and kasugamycin ¾ MIC) and no drug control treatments (NDC) collected in parallel. (**C**) Volcano plots of ¾ x MIC chloramphenicol vs. NDC showing significant changes in yellow for all the genes in SP_0828-0835. Similar data for kasugamycin and tetracycline shown in Supplementary Figure S7. (**D**) Genomic region of *S. pneumoniae* spanning SP_0828-SP_0835, encoding purine nucleotide metabolism genes. Colored arrows indicate the closest transcription start sites; dots indicate 3’-end sites. Coverage (average of triplicate) of both sense and antisense stands in RNA-seq is shown for chloramphenicol. Similar data for kasugamycin, and tetracycline are in Supplementary Figure S7B. Small black arrows indicate annealing positions for primers used in strand specific reverse-transcription PCR (RT-PCR) **(E)** The transcription start site identified for the antisense transcript is associated with a canonical RpoD (SigA) recognition site. **(F)** Strand-specific RT-PCR shows that SP_0835as is a single transcript with the dominant promoter in the region identified. Primer set 1 includes a forward primer corresponding to the 5’-tag sequence introduced by SP_0835as-specific RT and a reverse primer complementary to the 3’-end of SP_0835. Primer set 2 includes the same forward primer and a reverse primer in SP_0836. Genomic DNA is amplified for size comparison with the RT primer and each reverse primer. (**G**) RNA-seq data collected in a previous study from *S. pneumoniae* TIGR4 in the presence of diverse classes of antibiotics (15) shows significant SP_0835as transcription in the presence of several antibiotics, but levels similar to that observed in response to TIA appear only for tetracycline. (**H**) RNA-seq data collected from *S. pneumoniae* TIGR4 in several nutrient drop-outs during a previous study (15) shows a higher SP_0835as/total reads ratio in absence of both glycine and valine, but not in the absence of uracil.

If the observed antisense transcription is regulatory and negatively impacts sense strand transcription, we would expect to see an inverse correlation between these values. To assess this for the 162 genes with significant anti-sense transcription under any condition, we compared the change in gene expression of the coding (sense) strand with change in gene expression of the anti-sense strand in the presence of each antibiotic (Supplementary Figure S6). For genes that display significant differential expression (p_adj._ <0.05) of both strands, we observe a negative correlation in all three antibiotics (Spearmans’s correlation coefficient -0.41 to -0.61). Of particular note was a cluster of genes co-localized in the genome displaying substantial repression of gene expression and induction of antisense transcription in the presence of all three antibiotics compared to the no drug control, resulting in a very high ratio of anti-sense to total reads at this locus (Supplementary Figure S6, Figure 3B). These genes encode proteins involved in biosynthesis of purine precursors and the purine salvage pathway. All of the genes in this cluster display significant downregulation in coding strand expression for chloramphenicol and kasugamycin treatments, and coding transcripts are nearly undetectable in the presence of tetracycline (Figure 3C, Supplementary Figure S7A). These genes are responsible for the synthesis of ribose-5-P (*rpiA*/SP_0828, *deoB/*SP_0829) as well as the interconversion of purine bases (*deoD*/SP_0831/SP_0835) with their nucleoside and deoxynucleoside forms as part of the purine salvage pathway.

To corroborate the antisense transcription observed associated with purine salvage/biosynthesis genes, we extracted the transcript coverages for SP_0828-SP_0835 region for both strands and overlaid the previously determined TSS and TTS (Figure 3D, Supplementary Figure S7B, Supplementary Table S5-S7). TSSpredator does not annotate transcription start sites for features absent from the annotation file. Thus, to assign transcription start sites for the asRNA, we modified annotation agnostic PIPETS to identify potential TSS within this operon for both antibiotic treatments (30) (see Methods). Upstream of the higher confidence TSS, we found a clear consensus RpoD recognition sequence consisting of TATAAT -10 region and TTGACA in the -35 region (Figure 3E) that overlaps with the stop codon for SP_0835. To verify that the observed RNA-seq coverage corresponds to a single transcript we conducted strand-specific RT-PCR with RNA collected in both ¾ x MIC tetracycline and chloramphenicol (Figure 3F). We find that the entire antisense transcript from the region complementary to the 5’-UTR of SP_0828 through the end of SP_0835 (primer pair 1) is successfully amplified from strand-specific reverse transcription (see Methods). Furthermore, failure of a second primer placed just upstream of the proposed promoter (primer pair 2) to amplify a product from cDNA corroborates the proposed promoter site. Using this combined data, we identified a long asRNA, that we call SP_0835as, that spans purine nucleoside metabolism and salvage pathway (SP_0828-SP_0835), and is induced by all three TIA.

To determine whether this asRNA is present under other conditions, we turned to existing RNA-seq data for *S. pneumoniae* in the literature (15). The largest compendium of such data for *S. pneumoniae* TIGR4 was collected by Zhu & Surujon and co-workers and included multiple datasets in the presence of inhibitory concentrations of a range of different antibiotic classes including: vancomycin, levofloxacin, kanamycin, cefepime, ciprofloxacin, daptomycin and tetracycline (15). We assessed all of these data sets for asRNA transcription on SP_0828-SP_0835. We find that across all these samples there is a baseline level of asRNA expression that ranges from 10 to 60% of total reads. Although several antibiotics increased antisense reads modestly, tetracycline exposure (30- and 120-minutes) was the only condition in the published dataset that matched the high level of antisense transcription (>90%) we observed for SP_0828:SP_0835 (Figure 3G). We further assessed the nutrient dropout datasets collected with this study, and observed that the ratio of antisense/total transcription is elevated by amino acid dropouts (valine or glycine), but repressed by uracil dropout (Figure 3H). Together these data suggest that SP_0835as expression is strongly induced in response to translation inhibiting antibiotics, but also exists at a basal level in many conditions but may be elevated slightly by either nutrient or antibiotic stress.

### Purine salvage genes are regulated via asRNA transcription in response to TIA

To further investigate the relationship between transcription of the asRNA and the coding transcript (SP_0828-SP_0835) of *S. pneumoniae* TIGR4, we created an *S. pneumoniae* strain where the RpoD (SigA) consensus sequence of the native promoter for the asRNA is mutated (mP_SP_0835as) to prevent transcription (Figure 4A, Supplementary Figure S1. We also created a matched control strain containing only a Kan^r^ cassette (Wt^Kan^) inserted at an equivalent position). We then collected RNA-seq data with both strains in the absence or presence of ¾ x MIC (3 ug/mL) chloramphenicol for 60-minutes. In this data we observe that SP_0835as was significantly expressed in Wt^Kan^ in the presence of antibiotic, but antisense transcription was lost in mP_SP_0835as (Figure 4C), indicating that mutating the RpoD binding site upstream of the SP_0835as TSS significantly reduces its expression.

**Figure 4:**
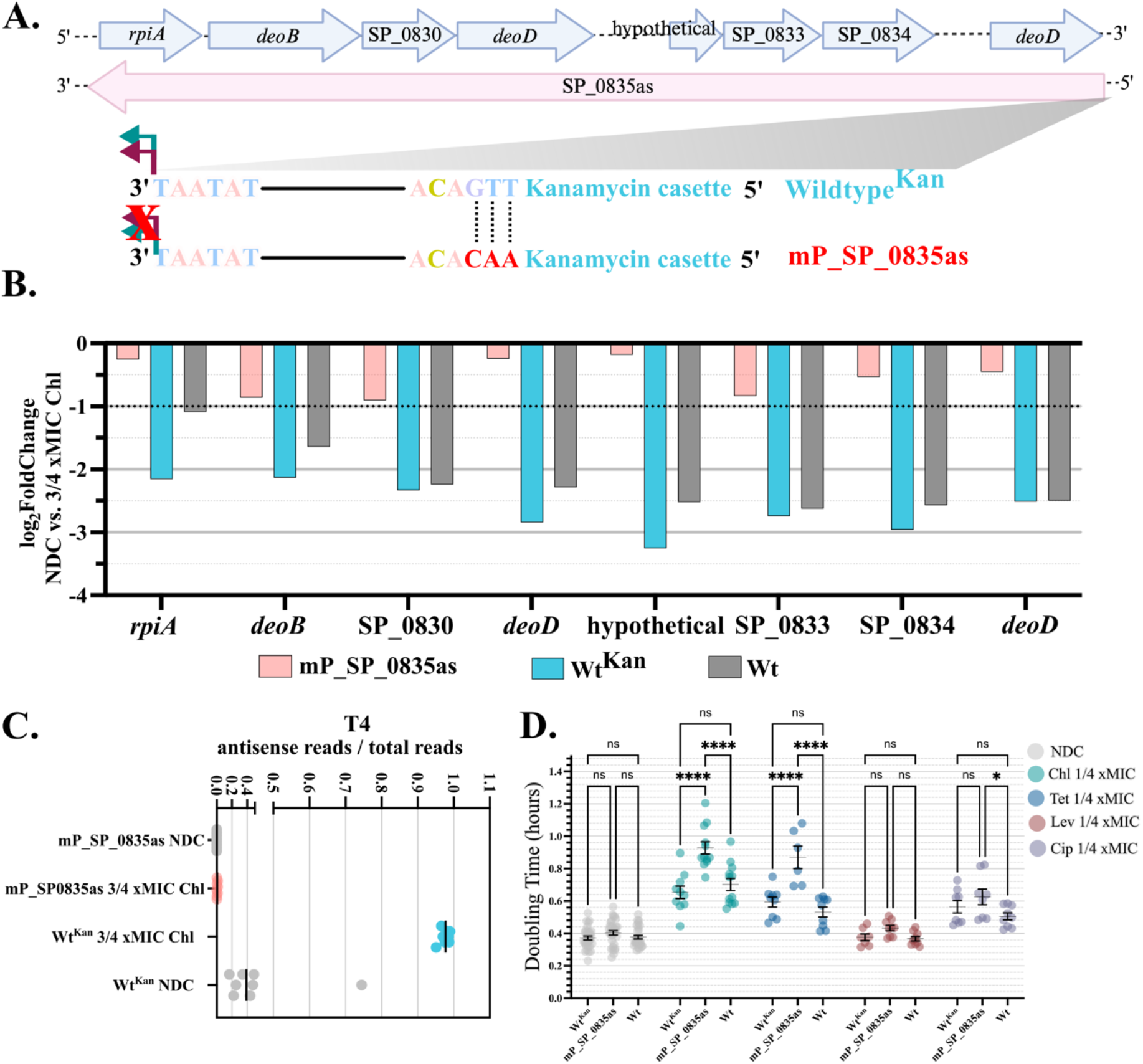
Anti-sense RNA SP_0835as negatively regulates SP_0828-SP_0835 to impact growth rate in the presence of TIA. **(A)** Diagram illustrating *S. pneumoniae* TIGR4 strains constructed to determine the relationship of SP_0835as transcription and expression of SP_0828-SP_0835. mP_SP0835as introduces three mutations into the -35 sequence of the anticipated promoter alongside the kanamycin resistance marker, and the Wildtype^Kan^ (Wt^Kan^) control strain with the kanamycin marker and no mutation. **(B)** Change in gene expression for SP_0828-SP_0835 in the presence (¾ x MIC) of chloramphenicol versus no drug control (NDC) for the promoter mutant mP_SP0835as, the Wt^Kan^ control strain, and unmodified *S. pneumoniae* TIGR4 (Wt) shows that mutating the promoter of SP_0835as eliminates the change in transcript levels observed for both the Wt and Wt^Kan^ strains. (**C**) Ratio of antisense/total transcription over SP_0828-SP_0835 shows that transcription of SP_0835as is eliminated in mP_SP0835as, but not in the Wt^Kan^ control strain. (**D**) The mutant strain lacking SP_0835as transcription (mP_SP0835as) displays an increased doubling time in the presence of ¼ MIC antibiotics that is more pronounced in conditions where SP_0835as is strongly induced (chloramphenicol (Chl) and tetracycline (Tet)) compared to the unmodified and control strains. In conditions where SP_0835as/total transcription is less perturbed from baseline (Levofloxacin (Lev), and Ciprofloxacin (Cip)), this increase is attenuated. Significance was determined via ordinary two-way ANOVA tests with Tukey’s multiple comparisons tests using a single pooled variance. Significant differences between datasets are indicated by asterisks (NDC n=33, Chl n = 12, Tet n = 9, Lev n=9, Cip n= 9, ns: not significant, ^*^ p < 0.0196, ^***^ p < 0.002, ^****^ p < 0.0001). Similar data collected at 1/8 x MIC and growth curves are available in Supplementary Figure S8.

To assess whether the loss of asRNA expression affects the transcript levels of genes on the coding strand in response to chloramphenicol treatment, we examined the relative expression of these genes for each strain (Wt, Wt^Kan^, mP_SP_0835as) in the presence of chloramphenicol compared to the matched NDC (54). These results show that for the mP_SP_0835as strain, gene expression levels for SP_0828-SP_0835 did not change significantly in the presence of antibiotic, suggesting that loss of asRNA transcription prevents the repression of these genes in response to antibiotic (Figure 4B). Additionally, we observed the Wt^Kan^ strain yields similar changes in gene expression of the coding genes as the unmodified strain suggesting that insertion of Kan^r^ alone does not substantially affect gene expression levels at this locus (Figure 4B). These results cumulatively indicate that in the presence of sub-MIC antibiotics, the asRNA SP_0835as is induced complementary to the purine biosynthesis and salvage genes (SP_0828-SP_0835), resulting in a reduction in expression of these genes.

Repression of purine biosynthesis has been linked to antibiotic tolerance and resistance for a number of different bacteria, although mechanisms leading to the effect are not always clear (66). Furthermore, the purine salvage pathway plays an important role in bacterial nucleotide metabolism by recycling purine bases and nucleosides from the environment into nucleotides, which can then be used for DNA and RNA synthesis in an energetically favorable manner compared to *de novo* purine biosynthesis (66). Given the extent to which nucleotide metabolism is downregulated in response to TIA (Figure 1C), we hypothesized that SP_0835as might provide a phenotypic benefit under the conditions in which it is highly expressed. To test this hypothesis, we assessed the growth rate of TIGR4 (Wt), Wt^Kan^, and mP_SP_0835as strains under sub-MIC antibiotic treatment. We observe that mP_SP_0835as grows significantly slower (∼40% increase in doubling time) than both unmodified *S. pneumoniae* TIGR4 and Wt^Kan^ at both 1/8 x and ¼ x MIC of chloramphenicol and tetracycline (Figure 4D, Supplementary Figure S8A,B). We also assessed growth at sub-MIC levels of antibiotics where the ratio of sense/antisense reads changes less compared to basal levels (ciprofloxacin and levofloxacin) and we find that although there is a slight increase in the doubling time compared to Wt in the presence of ciprofloxacin, this difference is far lower magnitude (∼10%). Thus, eliminating SP_0835as transcription not only forestalls the downregulation of SP_0828-SP_0835, but also results in substantially decreased growth rates under conditions where the antisense transcript dominates, suggesting that SP_0835as has biologically relevant functionality.

### The promoter for SP_0835as originates from an ISL3 insertion event

*S. pneumoniae* has a large and diverse pangenome (67) However, characterized RNA-regulators are typically within the core genome occurring in most clinical isolates (68,69). During assessment of existing *S. pneumoniae* transcriptomic data from other strains, we discovered that the promoter for SP_0835as is not present in other common *S. pneumoniae* laboratory strains including D39, or Taiwan 19F (Figure 5A). In TIGR4 immediately following SP_0835 is an insertion sequence (ISL3 variant) and SP_0836 encodes a predicted transposase. The RpoD recognition site for SP_0835as is formed from bases at the terminus of SP_0835 and part of the ISL3 sequence (Figure 5A). In D39 and 19F, the entire operon SP_0828-SP_0835 is present (SPD_0723-SPD_0730), but is directly followed by a DNA topology modulation protein FlaR (SPD_0731), which is encoded in TIGR4 by SP_0837 (Figure 5B). In *S. pneumoniae* Taiwan 19F this lack of promoter sequence is also associated with a lack of significant asRNA transcript representation in the presence of tetracycline (Figure 5D), as well as minimal changes in gene expression for the complementary coding genes (Figure 5C). This behavior stands in contrast to data collected by the same study for *S. pneumoniae* TIGR4 (Figure 3G), and in our data for TIGR4 (Figure 3B) (15). This finding further corroborates the relationship between SP_0835as transcription and regulation of the complementary purine biosynthesis and salvage genes.

**Figure 5:**
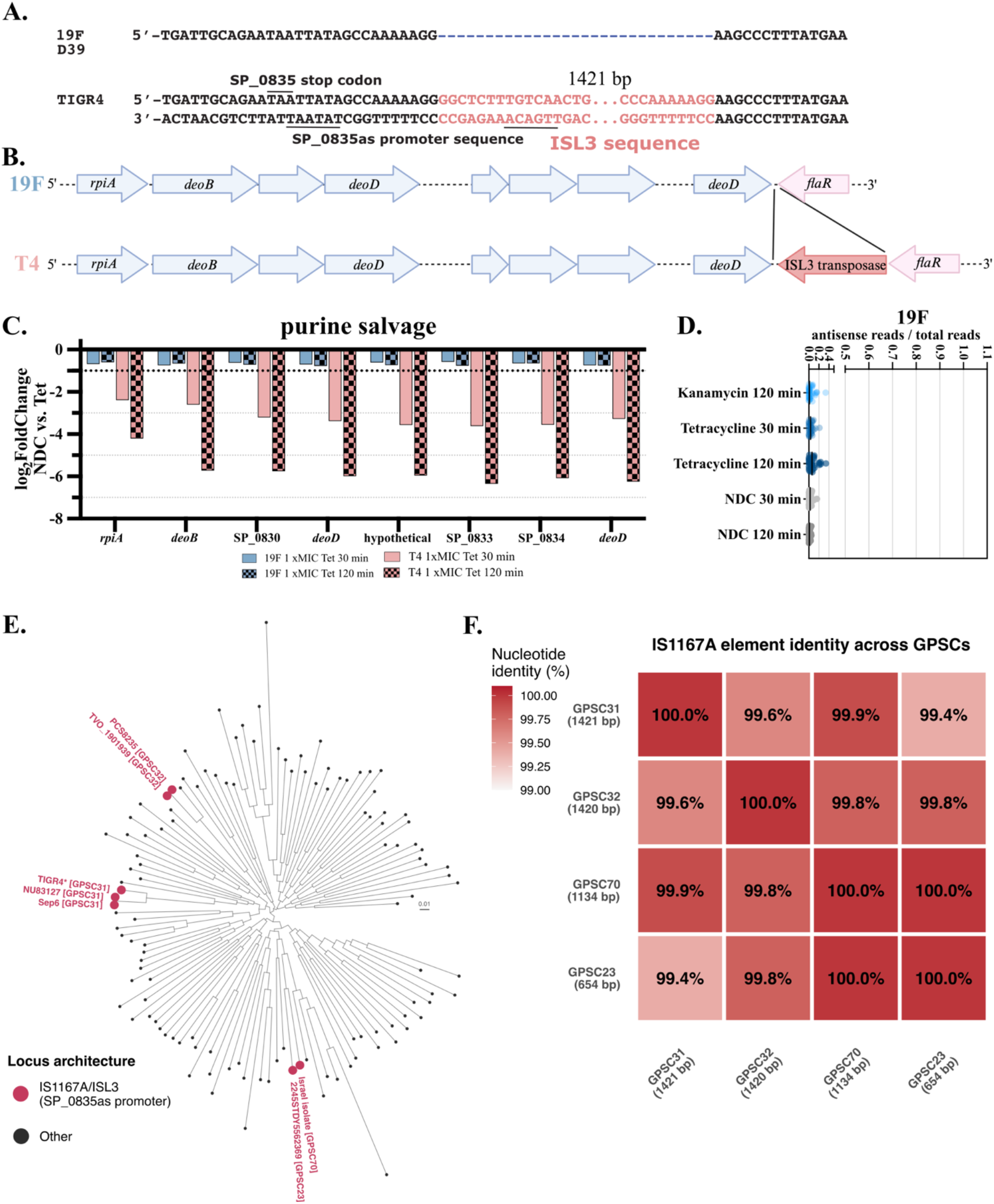
Promoter for SP_0835as formed via ISL3 insertion. **(A):** Sequences of *S. pneumoniae* strain TIGR4, and 19F (D39 is identical to 19F in this region) aligned to show promoter sequence spanning the end of SP_0835 and the ISL3 sequence is not present at this location in 19F or D39. **(B)** Overall genome architecture of *S. pneumoniae* strains Taiwan 19F (19F) and TIGR4 (T4) for the locus SP_0828-SP_0837. Complete alignment of the entire locus for 33 genomes including TIGR4 and 19F is in Supplementary Figure S9. **(C)** Comparing the change in gene expression upon treatment with tetracycline for 30 and 120 minutes for Taiwan 19F and TIGR4 shows that genes in this locus are not down regulated in the presence of tetracycline in strain Taiwan 19F, but are show substantial downregulation in TIGR4. Data collected as part of a pervious study and reanalyzed here from raw reads (15). **(D)** Analysis of antisense transcription over the equivalent locus shows that there is no significant antisense transcription in strain Taiwan 19F. **(E)** Core-genome maximum-likelihood phylogeny of 130 representative *S. pneumoniae* genomes. Tips are colored to indicate the presence of the ISL3 insertion creating the SP_0835as promoter at the purine metabolism locus. Junction-positive strains are distributed across four phylogenetically independent pneumococcal clusters: GPSC31, GPCS32, GPCS23, and GPCS70. Scale bar: 0.01 substitutions per site. **(F)** Pairwise nucleotide identity of ISL3 elements extracted from one representative genome per junction-positive pneumococcal cluster. The RpoD recognition site driving SP_0835as transcription is 100% identical across all four clusters. Elements were defined by BLASTn against the TIGR4 IS1167A reference (1,421 bp) and include both the transposase coding sequence and non-coding terminal sequences containing the inverted repeats and outward-directed promoter.

To assess the frequency of the SP_0835as promoter across the *S. pneumoniae* strain collection, we first assessed a small set of representative *S. pneumoniae* genomes sequenced via long-read sequencing, which are less likely to miss an insertion sequence due to assembly defects (44). We find that of these 33 genomes, in addition to TIGR4, the insertion sequence and asRNA promoter element are in only one of the genomes, that from TVO_1901939, a serotype 1 strain isolated in 2004 that is not expected to be exceedingly similar to TIGR4 (46). This is also the only strain among the long-read sequences that shares the region upstream of the start codon for SP_0828 with TIGR4, suggesting that the promoter regions for the coding transcript are also potentially distinct (Supplementary Figure S9).

To further examine the prevalence of the promoter for SP_0835as across the *S. pneumoniae* pangenome, we expanded our survey to the 366-chromosome level, complete *S. pneumoniae* genomes from NCBI GenBank. BLASTn search identified 15 genomes containing the specific junction creating the SP_0835as promoter at >90% identity, 7 of which are not TIGR4-derivative strains (Supplementary Table S9). To determine whether these strains are phylogenetically distinct, we constructed a core genome maximum-likelihood phylogeny from 130 representative genomes (see Methods). Our analysis revealed the promoter-creating junction is identified in strains assigned to four distinct global pneumococcus sequencing clusters (GPSC) that appear to be phylogenetically disparate (Figure 5E, Supplementary Table S9). To further explore the origins of the insertions, we extracted and aligned the IS element sequence from one representative genome per cluster (see Methods). This alignment shows that all the clusters share a very similar IS1167A elements, although that from GPSC23 has been truncated (Figure 5F). Thus, while SP_0835as is not part of the core *S. pneumoniae* genome, there is evidence that the element exists in clinically relevant strains.

## DISCUSSION

In this work we used RNA-sequencing, as well as 5’- and 3’-end sequencing to characterize the transcriptome of *S. pneumoniae* TIGR4 in the presence of sub-MIC translation inhibiting antibiotics (TIA). We find that metabolic genes are broadly downregulated, with a noticeable downregulation in nucleotide metabolism common to all three antibiotics. Our 5’ and 3’-end sequencing found that the majority of high confidence transcription start and termination sites are preserved. However, there are antisense RNAs induced in response to bacteriostatic TIA, most notably SP_0835as. Antisense RNAs have been reported in several Streptococcus species, but these reports are typically part of larger sequencing studies, and often refer to shorter transcripts (∼150-200 bp) rather than the long antisense transcript running complementary to many genes that we observe here (70,71). We are not aware of any work to date that explores the functionality of these transcripts in any detail, although several asRNA examples in Listeria have been characterized more comprehensively (35,72,73).

We show that expression of SP_0835as is anti-correlated with expression of the coding genes on the complementary strand. However, there are a variety of different mechanisms by which antisense transcription may modulate expression of the opposing genes that are consistent with our data (74). These mechanisms include transcriptional interference where two transcripts are transcribed in a mutually exclusive manner due to collision of the polymerases (75) or inhibition of a weaker promoter as RNAP initiated in the opposite direction elongates through it (76). However, selective degradation of double stranded RNA produced by the combination of sense and antisense transcripts (77), or transcript attenuation due to the formation of an intrinsic terminator mediated via the anti-sense strand (78) are also potential mechanisms given our observed changes in transcript levels. Furthermore, a combination of mechanisms may also occur, making deconvolution challenging, especially given the large expected sizes of both the sense and antisense transcripts (∼6-7kb), and their nearly complete overlap.

It has been argued that many antisense transcripts represent spurious transcription events rather than components of adaptive responses (26,79). However, there are several arguments for the biological relevance of SP_0835as. First, levels of the antisense transcript vary across conditions, with the antisense transcript highly expressed in relatively few of the conditions for which data are available, suggesting a specific trigger for strong expression. Second, our phenotypic data indicate that strains expressing SP_0835as (TIGR4, WT^Kan^) display a benefit relative to those lacking the RNA (mP_SP_0835as) that is more pronounced in conditions where the asRNA is highly expressed. Third, we identify the promoter element in several phylogenetically distinct strain clusters, suggesting that while SP_0835as expression is not common across the pangenome, it is either horizontally transferred or has arisen multiple times.

In addition, SP_0835as expression influences not only the complementary coding genes, but has affects across the transcriptome (Figure 1C). We find that SP_0835as expression amplifies the repression of purine metabolism, including purine *de novo* synthesis and transport genes that are not directly complimentary to the antisense RNA (Figure 6A). For *S. pneumoniae* strain Taiwan 19F a smattering of purine biosynthesis genes display significantly reduced expression in the presence of tetracycline (15). In comparison, the same study found that under similar conditions, strain TIGR4 strongly represses nearly all genes in purine metabolism (Figure 6B). Similarly, we observe that in our strain lacking SP_0835as expression (mP_SP0835as), purine metabolic genes show an attenuated response to ¾ x MIC chloramphenicol compared to the control strain (Wt^Kan^) (Figure 6C). While this is likely be the indirect impact of reducing purine precursors via reduction in levels of *rpiA* and *deoB*, it also shows that SP_0835as transcription impacts gene expression globally, amplifying a response that exists even in its absence.

**Figure 6:**
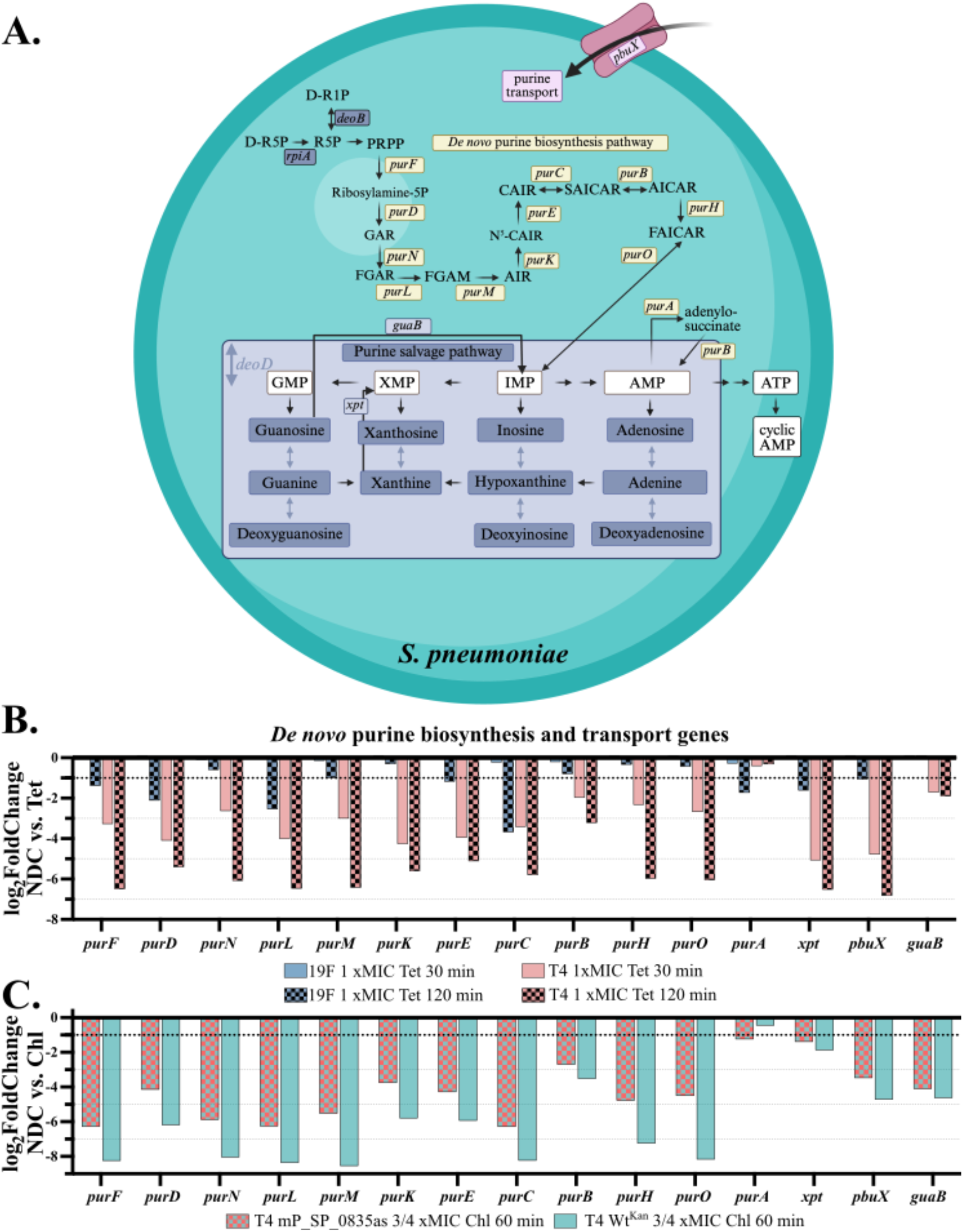
SP_0835as amplifies negative regulation of *de novo* purine synthesis and transport genes **(A)** Schematic showing genes involved in purine metabolism. **(B)** Change in expression for genes in purine biosynthesis and transport in the presence of 1 x MIC tetracycline for strain Taiwan 19F (no SP_0835as expression and no repression of purine metabolic genes *rpiA, deoB*, and *deoD*), and for strain TIGR4 (SP_0835as expression and repression of *rpiA, depB*, and *deoD*) shows that 19F significantly downregulates some purine biosynthesis genes, but this response is much stronger in TIGR4 (15). **(C)** Change in expression for genes in purine biosynthesis and transport in the presence of ¾ x MIC chloramphenicol for TIGR4 strain lacking SP_0835as (T4 mP_SP_0835as), and control strain (T4 Wt^Kan^) shows that similarly the response of purine biosynthesis genes is attenuated the strain lacking SP_0835as.

Regulatory RNAs are often narrowly distributed to subsets of bacterial species, and are extraordinarily diverse across the bacterial kingdom (8,11,80). There has been a great deal of speculation regarding the mechanisms that allow such regulatory elements to arise, and ultimately persist (10,26,81). Despite this interest, and the explosion in microbial genome data now available, there are few documented examples where nascent regulatory elements may be convincingly demonstrated (9,82). Although much remains to be determined regarding the requirements that balance the sense and antisense transcription, and the mechanism(s) by which regulation occurs, SP_0835as represents an important addition to this small group. Furthermore, SP_0835as also adds to a growing number of examples where insertion sequences remodel bacterial genomes, not purely via gene disruption (83-86), but also by introducing novel promoters that dramatically change gene expression patterns (87).

## Acknowledgements

We would like to thank Indu Warrier and Jon Anthony for helpful discussions regarding data collection and analysis. Some of the scientific sketches were created using BioRender.

## Author contributions

Conceptualization: M.M.M., T.vO. I.O., Formal Analysis: I.O., Funding acquisition: M.M.M., T.vO., Investigation: I.O., M.M.M., Methodology: I.O., Project administration: M.M.M., I.O., Resources: M.M.M., Visualization: I.O., Writing - original draft: I.O., Writing - review draft & editing: I.O. T.vO. M.M.M.

## Code availability

RNA-seq, 5’-end seq and 3’-end sequencing reads have been processed using the Aerobio platform (https://github.com/jsa-aerial/aerobio). Downstream custom scripts for RNA-seq analysis, 3’-end & 5’-end classification, intrinsic transcription termination filter, binomial testing for antisense RNA discovery have been deposited to SP_0835as GitHub platform (https://github.com/iremozkanRNA/SP_0835as or https://doi.org/10.5281/zenodo.20518592)

## Supplementary Data

Figures S1-S9 are a single pdf.

Tables S1-S9 are each excel spreadsheets

