## Supplementary material for "Translation Inhibiting Antibiotics Induce an asRNA Regulating Purine Metabolism in *S. pneumoniae* TIGR4 via an ISL3 Insertion Derived Hybrid Promoter": Figure S1

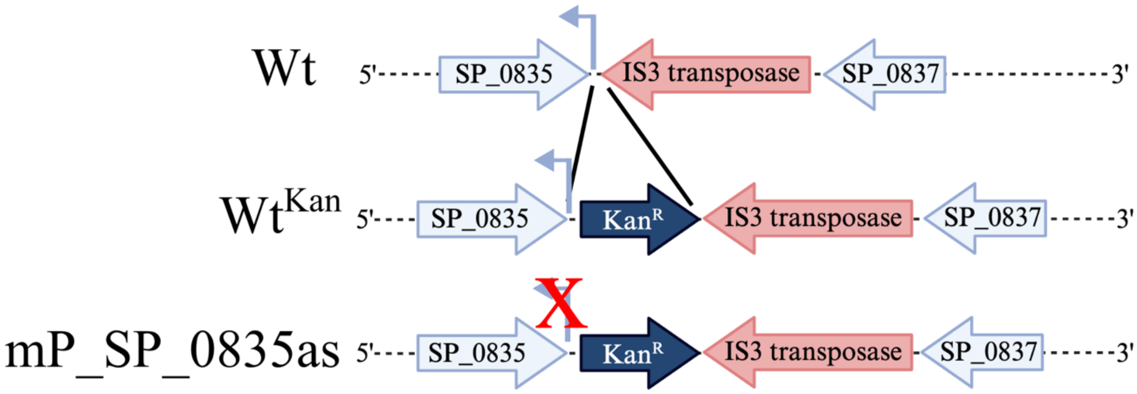


**Supplementary Figure S1**: Homologous recombination strategy to replace native promoter with a version that removes RpoD recognition sequence (Figure 4). Primers in Supplementary Table S1.
