## Supplementary material for "Translation Inhibiting Antibiotics Induce an asRNA Regulating Purine Metabolism in *S. pneumoniae* TIGR4 via an ISL3 Insertion Derived Hybrid Promoter": Figure S2

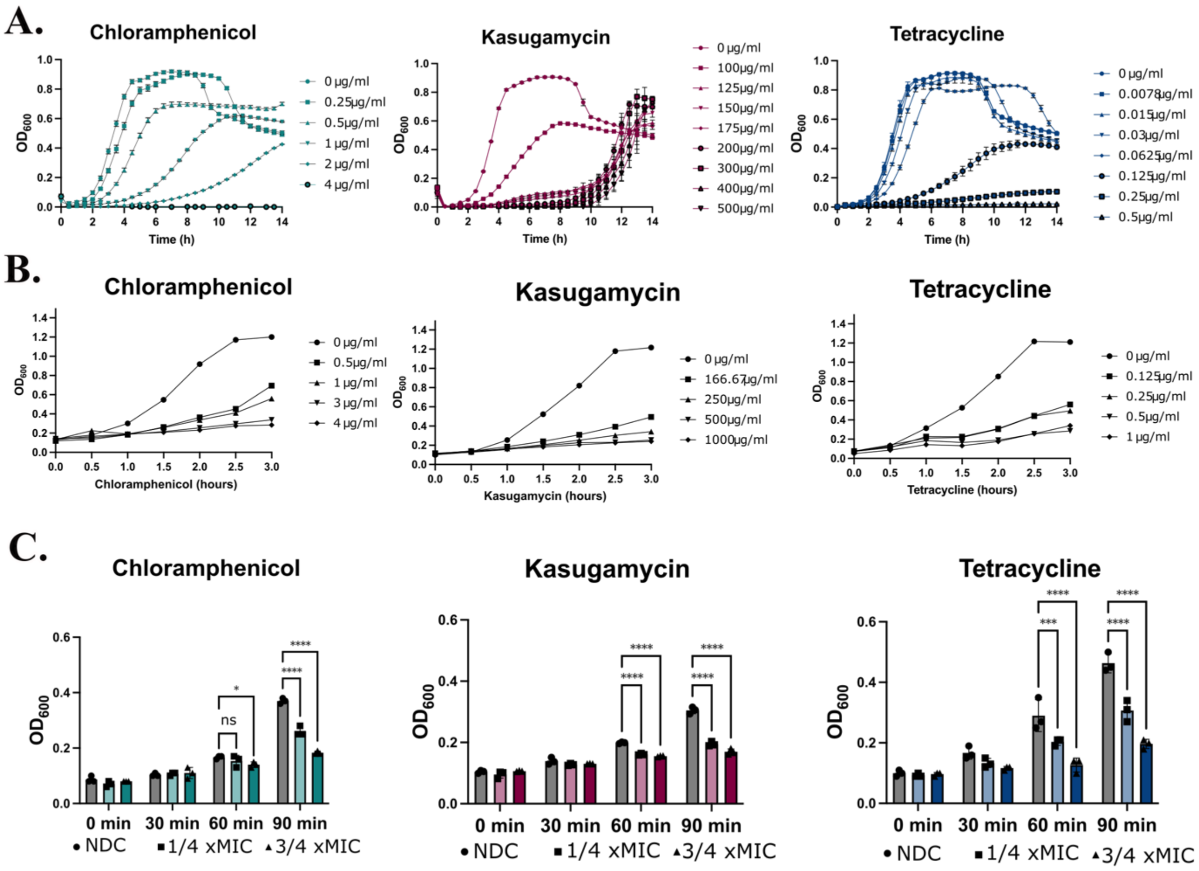


**Supplementary Figure 2:** Determination of MIC for each TIA for *S. pneumoniae* TIGR4 strain. **(A)** Growth curves performed in 96 well plates for chloramphenicol, tetracycline and kasugamycin treatments with increasing concentrations of antibiotic. Error bars correspond to standard deviation on 3 biological replicates. Highest concentrations shown in the figures was determined to be the MIC for each TIA. **(B)** $OD_{600}$ of each culture in tall culture tubes tested before sample collection (0-5 hour) and at different MIC for each antibiotic (n=2 replicate). **(C)** $OD_{600}$ of each culture during sample collection at different time points (0-, 30-, 60- and 90-minutes) and at different MIC for each antibiotic. Individual data points and the standard error are indicated (n=4). Statistical significance determined by two-way repeated measures ANOVA with Dunnett's post-hoc test. Significant differences between growth conditions are indicated by asterisks (**** $p < 0.0001$, *** $p < 0.003$, * $p < 0.021$, ns: not significant).
