## Supplementary material for "Translation Inhibiting Antibiotics Induce an asRNA Regulating Purine Metabolism in *S. pneumoniae* TIGR4 via an ISL3 Insertion Derived Hybrid Promoter": Figure S3

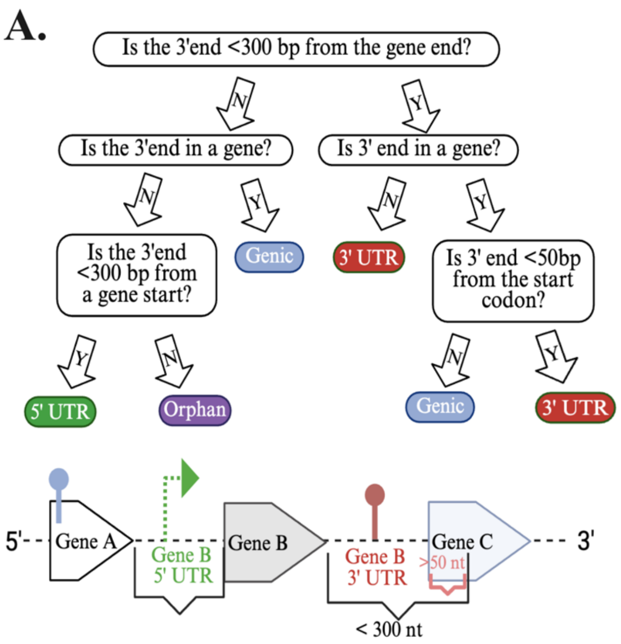


**Supplementary Figure S3: (A)** Assignment strategy for annotation of 3'-end sites (TTS) in the *S. pneumoniae* genome that integrates 3'-end data analyzed annotation agnostically (1) with gene annotation information. This strategy prioritizes assignment of 3'-UTR over other locations**. (B)** Assignment strategy for annotation of 5'-end sites (TSS) that integrates 5'-end data with gene annotation information. This strategy prioritizes assignment of 5'-UTR over other locations.
