## Supplementary material for "Translation Inhibiting Antibiotics Induce an asRNA Regulating Purine Metabolism in *S. pneumoniae* TIGR4 via an ISL3 Insertion Derived Hybrid Promoter": Figure S4

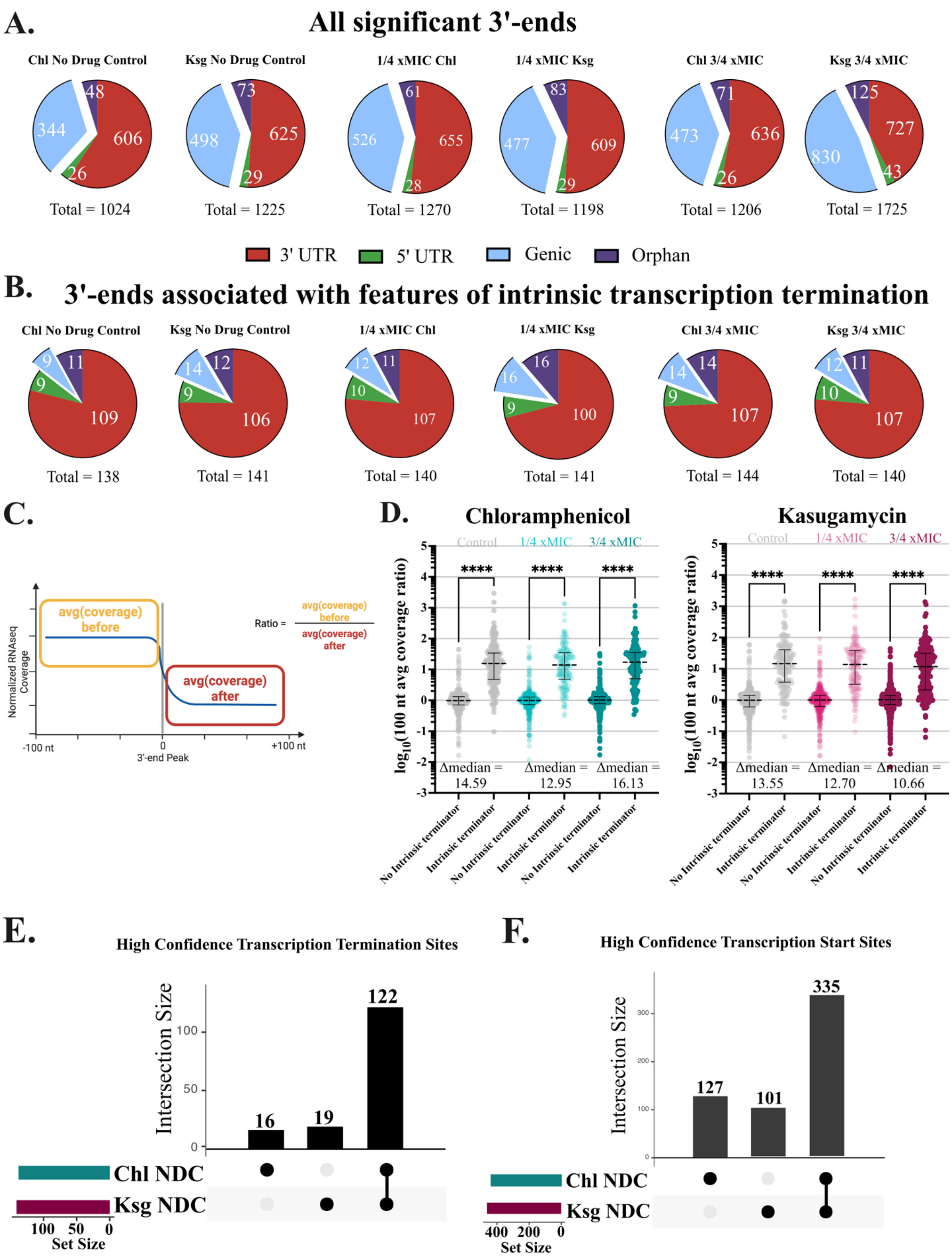


**Supplementary Figure S4: (A)** Distribution of position relative to genic features of significant 3'-ends identified with PIPETS from 3'-end sequencing data (1) (Supplementary Table S5)

according to hierarchy in Supplementary Figure S3. **(B)** Distribution of high confidence 3'-ends that are associated with features of intrinsic termination (strong stem followed by poly-uridine track). **(C)** Schematic to explain coverage ratio, calculated based on RNA-seq data collected under the same condition under which the 3'-end was considered significant. **(D**) 3'-ends associated with features of intrinsic termination are more likely to display a significant drop in RNA-seq coverage following the coordinate of termination (high value of coverage ratio) than sites lacking these features. Significance determined via Wilcoxon nonparametric tests for pairwise comparisons due to deviations of the population from a normal distribution. Significant differences between datasets are indicated by asterisks (**** $p < 0.0001$). **(E)** Overlap in high confidence transcription termination sites identified in two distinct no drug control (NDC) replicates collected alongside each antibiotic condition. **(F)** Overlap of high confidence transcription start sites identified in two distinct no drug control (NDC) replicates collected alongside each antibiotic condition.
