## Supplementary material for "Translation Inhibiting Antibiotics Induce an asRNA Regulating Purine Metabolism in *S. pneumoniae* TIGR4 via an ISL3 Insertion Derived Hybrid Promoter": Figure S5

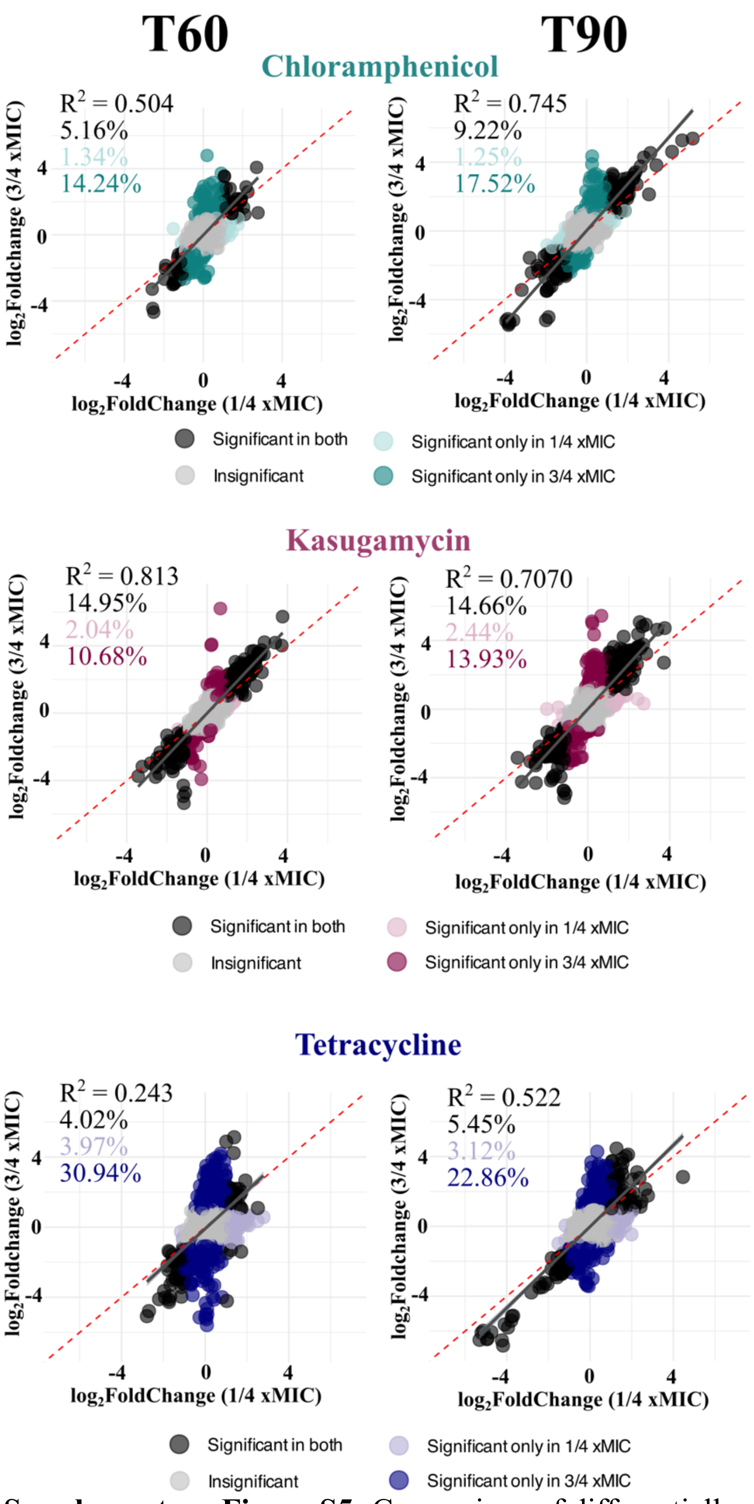


**Supplementary Figure S5:** Comparison of differentially expressed genes compared to NDC for ¼ (x-axis) and ¾ x MIC (y-axis) antibiotic treatment shows that responses at different antibiotic concentrations are largely correlated. Numbers in top left of graph correspond to proportion of genes significantly differentially expressed in both samples (black), only the ¼ x MIC sample (light color), and ¾ x MIC (dark color).
