## Supplementary material for "Translation Inhibiting Antibiotics Induce an asRNA Regulating Purine Metabolism in *S. pneumoniae* TIGR4 via an ISL3 Insertion Derived Hybrid Promoter": Figure S6

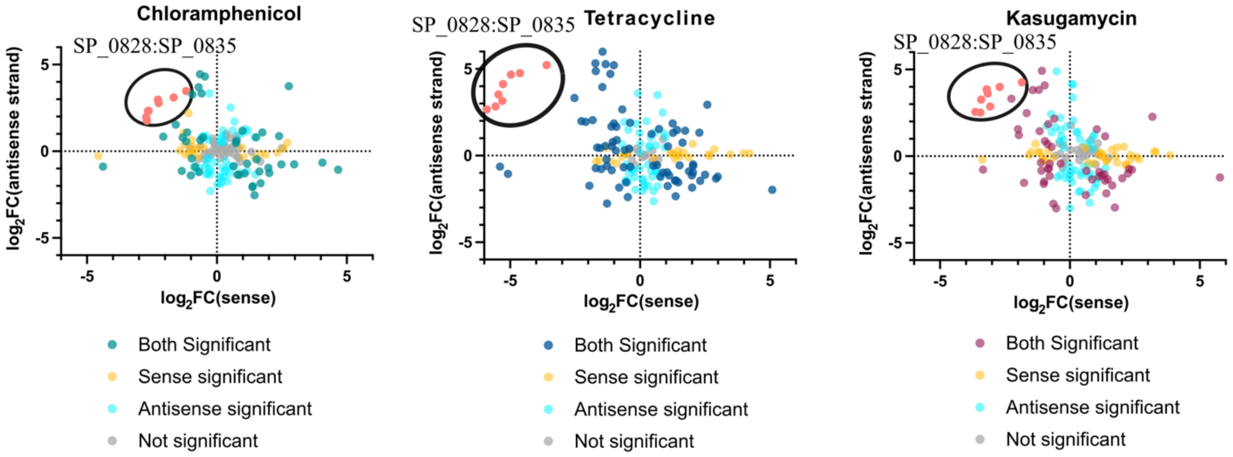


**Supplementary Figure S6:** Comparative analysis of sense and antisense strand expression changes in three antibiotic treatments (DESeq2). Scatter plots showing $log_2$FoldChanges in sense (x-axis) versus antisense (y-axis) strand expression for genes displaying significant antisense expression under any condition in response to chloramphenicol, tetracycline, and kasugamycin treatments (3/4 xMIC vs. NDC). Points are colored by statistical significance. light yellow indicates sense strand only significant ($p_{adj} < 0.05$); light blue indicates antisense strand only significant; and gray indicates neither strand significant. Significance of both strands are shown in teal for chloramphenicol, in dark blue for tetracycline, and purple for kasugamycin. For genes with significant change in expression of both the sense and antisense strand expression, these values are negatively correlated. For chloramphenicol Spearman's correlation constant = -0.416 (p=.0027, n= 55). For tetracycline Spearman's correlation constant = -0.521 (p<0.001, n= 80). For kasugamycin Spearman's correlation constant =-0.607 (p<0.001, n=50). The cluster of genes in red (SP_0828:SP_0835) shows anticorrelated antisense strand upregulation and sense strand downregulation across all three antibiotic conditions, suggesting stress-responsive antisense RNA regulation.
