## Supplementary material for "Translation Inhibiting Antibiotics Induce an asRNA Regulating Purine Metabolism in *S. pneumoniae* TIGR4 via an ISL3 Insertion Derived Hybrid Promoter": Figure S7

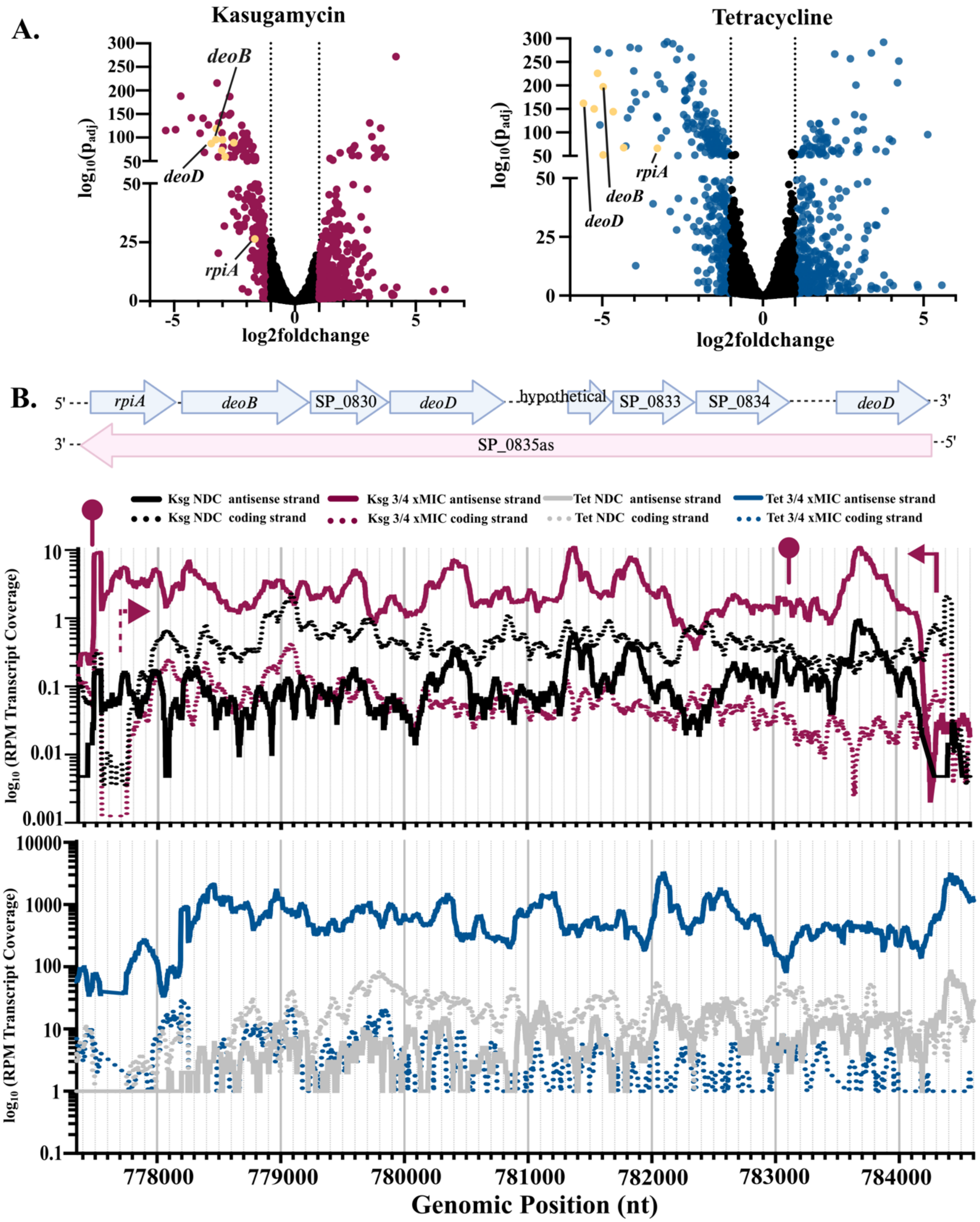


**Supplementary Figure S7:** (**A**) Volcano plots of ¾ x MIC kasugamycin vs. NDC and ¾ x MIC tetracycline vs NDC showing significant changes in yellow for all the genes in SP_0828-0835 including *rpiA*, *deoD*, and *deoB*. (**B**) Genomic region of *S. pneumoniae* spanning SP_0828-SP_0835, encoding purine nucleotide metabolism genes. RPM coverages (average of triplicate) of both sense and antisense stands in RNA-seq is shown for kasugamycin and tetracycline. Transcription start sites (TSS) and termination sites (TTS) shown for kasugamycin based on 3'- and 5'-end data.
