## Supplementary material for "Translation Inhibiting Antibiotics Induce an asRNA Regulating Purine Metabolism in *S. pneumoniae* TIGR4 via an ISL3 Insertion Derived Hybrid Promoter": Figure S8

A.

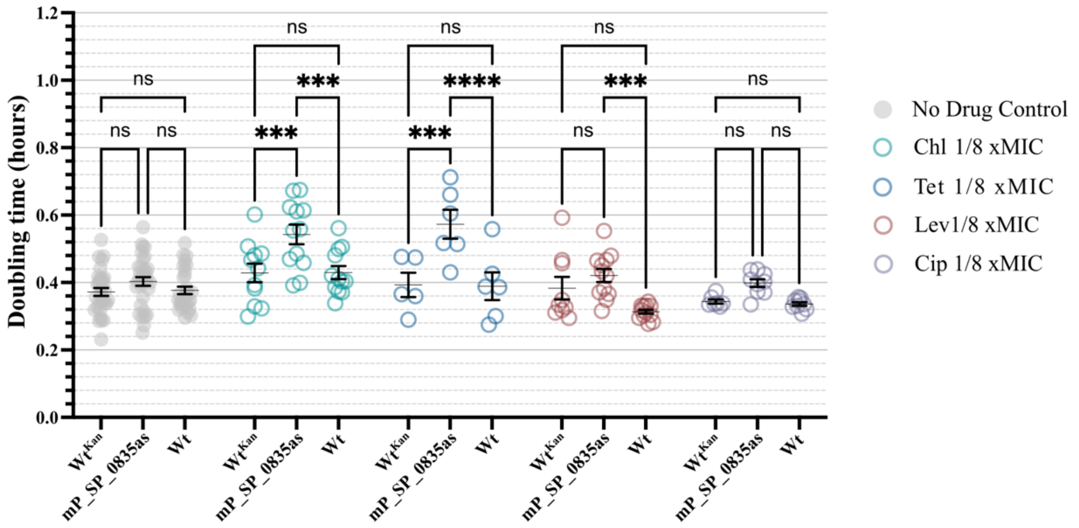


B.

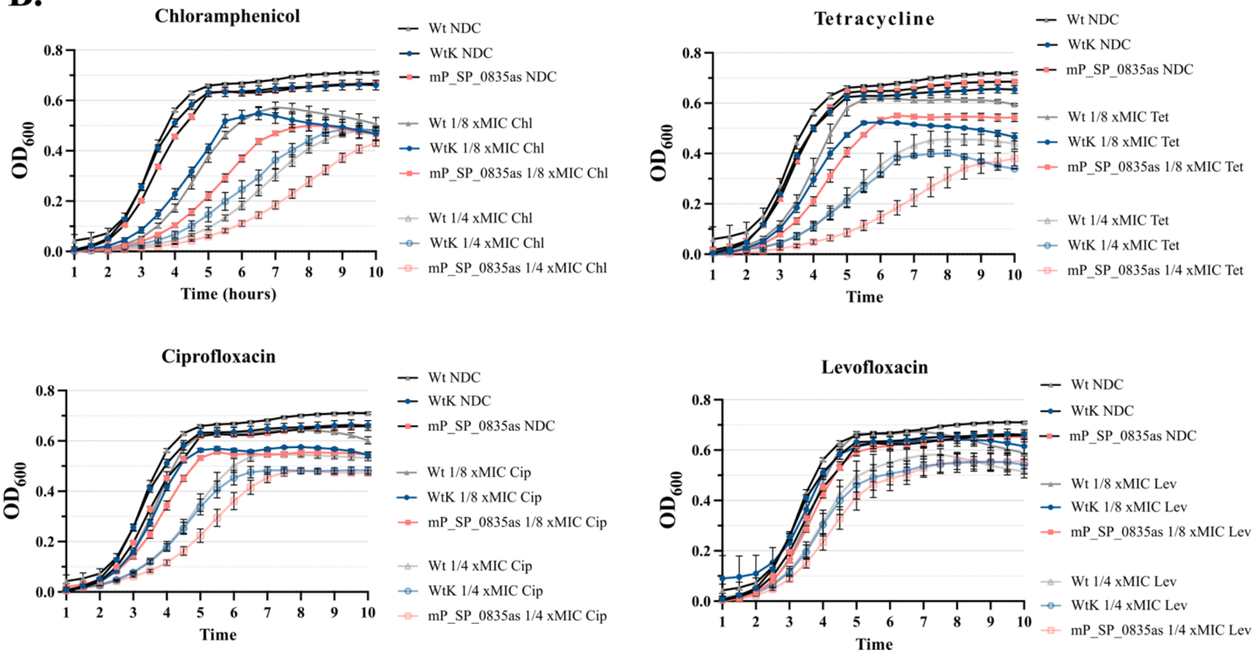


**Supplementary Figure S8: (A)** Doubling time for $Wt^{Kan}$, SP_0835as promoter mutant (mP_SP_0835as), and unmodified TIGR4 (Wt) in the presence of no antibiotic (No Drug Control), and 1/8 x MIC of: chloramphenicol (0.5 ug/mL), tetracycline (0.0625 ug/mL), levofloxacin (0.3125 ug/mL), and ciprofloxacin (0.25 ug/mL). **(B)** Growth curves used to calculate the doubling times presented above and in Figure 4. Error bars correspond to standard error of the mean. Significance was determined via ordinary two-way ANOVA tests with Tukey's multiple comparisons tests using a single pooled variance. Significant differences between datasets are indicated by asterisks (NDC n=33, Chl n = 12, Tet n = 6, Lev n=6, Cip n= 9, ns: not significant, *** $p < 0.004$, **** $p < 0.0001$).
