## Supplementary material for "Translation Inhibiting Antibiotics Induce an asRNA Regulating Purine Metabolism in *S. pneumoniae* TIGR4 via an ISL3 Insertion Derived Hybrid Promoter": Figure S9

### Ozkan et al. Supplementary Figures

```
NC_003028.3:777000-787000      1      AGGACAAGTTTGTTGTTTCCATCTAACCAGCTTATCTTAACTCCCCACCAAAAAGAATGGGAAAAACTGTCTGGTATTGCTATTGAAAAG
NZ_CP035248.1:765099-775111    1      AGGACAAGTTTGTTGTTTCCATCTAACCAGCTTATCTTAACTCCCCACCAAAAAGAATGGGAAAAACTGTCTGGTATTGCTATTGAAAAG
NZ_CP035235.1 c1326834-1317894 0      ------------------------------------------------------------------------------------------
NZ_CP035234.1:756284-765224    0      ------------------------------------------------------------------------------------------
NZ_CP038251.1:750745-759686    0      ------------------------------------------------------------------------------------------
NZ_CP035244.1:747023-755950    0      ------------------------------------------------------------------------------------------
NZ_CP035264.1:800836-809336    0      ------------------------------------------------------------------------------------------
NZ_CP035265.1:766128-774627    0      ------------------------------------------------------------------------------------------
NZ_CP035245.1 745261-753757    0      ------------------------------------------------------------------------------------------
NZ_CP035259.1:745700-753548    0      ------------------------------------------------------------------------------------------
NZ_CP035258.1:750874-758722    0      ------------------------------------------------------------------------------------------
NZ_CP035260.1:726156-733994    0      ------------------------------------------------------------------------------------------
NZ_CP035241.1:708296-716144    0      ------------------------------------------------------------------------------------------
NZ_CP035261.1:779086-786938    0      ------------------------------------------------------------------------------------------
NZ_CP035242.1:c1532236-1524385 0      ------------------------------------------------------------------------------------------
NZ_CP035243.1:777729-785580    0      ------------------------------------------------------------------------------------------
NZ_CP035237.1:c1311322-1303456 0      ------------------------------------------------------------------------------------------
NZ_CP035246.1:c1302892-1295026 0      ------------------------------------------------------------------------------------------
NZ_CP035247.1:c1301115-1293263 0      ------------------------------------------------------------------------------------------
NZ_CP035238.1:738547-746413    0      ------------------------------------------------------------------------------------------
NZ_CP035255.1:776866-784718    0      ------------------------------------------------------------------------------------------
NZ_CP035256.1:807236-815102    0      ------------------------------------------------------------------------------------------
NZ_CP038252.1:1151812-1159673  0      ------------------------------------------------------------------------------------------
NZ_CP035254.1:793365-801213    0      ------------------------------------------------------------------------------------------
NZ_CP035251.1:748733-756585    0      ------------------------------------------------------------------------------------------
NZ_CP035252.1:745670-753536    0      ------------------------------------------------------------------------------------------
NZ_CP035257.1:776317-784169    0      ------------------------------------------------------------------------------------------
NZ_CP038253.1:758105-765957    0      ------------------------------------------------------------------------------------------
NZ_CP035263.1:743050-750901    0      ------------------------------------------------------------------------------------------
NZ_CP035262.1:741595-749460    0      ------------------------------------------------------------------------------------------
NZ_CP035236.1:764915-772766    0      ------------------------------------------------------------------------------------------
NZ_CP035253.1 767286-775134    0      ------------------------------------------------------------------------------------------
NZ_CP035249.1:763080-770921    0      ------------------------------------------------------------------------------------------

NC_003028.3:777000-787000      91     CAAAACGAAGGTACAACATCTAGTGCCCTGACTTCTTTCCCTCAAGGAACAATTTTGGTAGAGAAAGGTCCAGCTACTCGTATTTGGCAA
NZ_CP035248.1:765099-775111    91     CAAAACGAAGGTACAACATCTAGTGCCCTGACTTCTTTCCCTCAAGGAACAATTTTGGTAGAGAAAGGTCCAGCTACTCGTATTTGGCAA
NZ_CP035235.1 c1326834-1317894 0      ------------------------------------------------------------------------------------------
NZ_CP035234.1:756284-765224    0      ------------------------------------------------------------------------------------------
NZ_CP038251.1:750745-759686    0      ------------------------------------------------------------------------------------------
NZ_CP035244.1:747023-755950    0      ------------------------------------------------------------------------------------------
NZ_CP035264.1:800836-809336    0      ------------------------------------------------------------------------------------------
NZ_CP035265.1:766128-774627    0      ------------------------------------------------------------------------------------------
NZ_CP035245.1 745261-753757    0      ------------------------------------------------------------------------------------------
NZ_CP035259.1:745700-753548    0      ------------------------------------------------------------------------------------------
NZ_CP035258.1:750874-758722    0      ------------------------------------------------------------------------------------------
NZ_CP035260.1:726156-733994    0      ------------------------------------------------------------------------------------------
NZ_CP035241.1:708296-716144    0      ------------------------------------------------------------------------------------------
NZ_CP035261.1:779086-786938    0      ------------------------------------------------------------------------------------------
NZ_CP035242.1:c1532236-1524385 0      ------------------------------------------------------------------------------------------
NZ_CP035243.1:777729-785580    0      ------------------------------------------------------------------------------------------
NZ_CP035237.1:c1311322-1303456 0      ------------------------------------------------------------------------------------------
NZ_CP035246.1:c1302892-1295026 0      ------------------------------------------------------------------------------------------
NZ_CP035247.1:c1301115-1293263 0      ------------------------------------------------------------------------------------------
NZ_CP035238.1:738547-746413    0      ------------------------------------------------------------------------------------------
NZ_CP035255.1:776866-784718    0      ------------------------------------------------------------------------------------------
NZ_CP035256.1:807236-815102    0      ------------------------------------------------------------------------------------------
NZ_CP038252.1:1151812-1159673  0      ------------------------------------------------------------------------------------------
NZ_CP035254.1:793365-801213    0      ------------------------------------------------------------------------------------------
NZ_CP035251.1:748733-756585    0      ------------------------------------------------------------------------------------------
NZ_CP035252.1:745670-753536    0      ------------------------------------------------------------------------------------------
NZ_CP035257.1:776317-784169    0      ------------------------------------------------------------------------------------------
NZ_CP038253.1:758105-765957    0      ------------------------------------------------------------------------------------------
NZ_CP035263.1:743050-750901    0      ------------------------------------------------------------------------------------------
NZ_CP035262.1:741595-749460    0      ------------------------------------------------------------------------------------------
NZ_CP035236.1:764915-772766    0      ------------------------------------------------------------------------------------------
NZ_CP035253.1 767286-775134    0      ------------------------------------------------------------------------------------------
NZ_CP035249.1:763080-770921    0      ------------------------------------------------------------------------------------------

NC_003028.3:777000-787000      181    GTTGGCCAGTCTGATTATTACCAGTTAAAGGTTGGCGGTCCCTATCAGGCGACTGGTGGTATGGGTGATACACTGGCTGGAATGATTGCA
NZ_CP035248.1:765099-775111    181    GTTGGCCAGTCTGATTATTACCAGTTAAAGGTTGGCGGTCCCTATCAGGCGACTGGTGGTATGGGTGATACACTGGCTGGAATGATTGCA
NZ_CP035235.1 c1326834-1317894 0      ------------------------------------------------------------------------------------------
NZ_CP035234.1:756284-765224    0      ------------------------------------------------------------------------------------------
NZ_CP038251.1:750745-759686    0      ------------------------------------------------------------------------------------------
NZ_CP035244.1:747023-755950    0      ------------------------------------------------------------------------------------------
NZ_CP035264.1:800836-809336    0      ------------------------------------------------------------------------------------------
NZ_CP035265.1:766128-774627    0      ------------------------------------------------------------------------------------------
NZ_CP035245.1 745261-753757    0      ------------------------------------------------------------------------------------------
NZ_CP035259.1:745700-753548    0      ------------------------------------------------------------------------------------------
NZ_CP035258.1:750874-758722    0      ------------------------------------------------------------------------------------------
NZ_CP035260.1:726156-733994    0      ------------------------------------------------------------------------------------------
NZ_CP035241.1:708296-716144    0      ------------------------------------------------------------------------------------------
NZ_CP035261.1:779086-786938    0      ------------------------------------------------------------------------------------------
NZ_CP035242.1:c1532236-1524385 0      ------------------------------------------------------------------------------------------
NZ_CP035243.1:777729-785580    0      ------------------------------------------------------------------------------------------
NZ_CP035237.1:c1311322-1303456 0      ------------------------------------------------------------------------------------------
NZ_CP035246.1:c1302892-1295026 0      ------------------------------------------------------------------------------------------
NZ_CP035247.1:c1301115-1293263 0      ------------------------------------------------------------------------------------------
NZ_CP035238.1:738547-746413    0      ------------------------------------------------------------------------------------------
NZ_CP035255.1:776866-784718    0      ------------------------------------------------------------------------------------------
NZ_CP035256.1:807236-815102    0      ------------------------------------------------------------------------------------------
NZ_CP038252.1:1151812-1159673  0      ------------------------------------------------------------------------------------------
NZ_CP035254.1:793365-801213    0      ------------------------------------------------------------------------------------------
NZ_CP035251.1:748733-756585    0      ------------------------------------------------------------------------------------------
NZ_CP035252.1:745670-753536    0      ------------------------------------------------------------------------------------------
NZ_CP035257.1:776317-784169    0      ------------------------------------------------------------------------------------------
```

```
NZ_CP038253.1:758105-765957    0      ------------------------------------------------------------------------------------------
NZ_CP035263.1:743050-750901    0      ------------------------------------------------------------------------------------------
NZ_CP035262.1:741595-749460    0      ------------------------------------------------------------------------------------------
NZ_CP035236.1:764915-772766    0      ------------------------------------------------------------------------------------------
NZ_CP035253.1 767286-775134    0      ------------------------------------------------------------------------------------------
NZ_CP035249.1:763080-770921    0      ------------------------------------------------------------------------------------------

NC_003028.3:777000-787000      271    GGATTTGCAGGCCAATTTCGACAGGCCAGTCTCTACGAACGTGTGGCAGTAGCAACCCATCTTCATTCAGCCATAGCCCAAGAACTATCT
NZ_CP035248.1:765099-775111    271    GGATTTGCAGGCCAATTTCGACAGGCCAGTCTCTACGAACGTGTGGCAGTAGCAACCCATCTTCATTCAGCCATAGCCCAAGAACTATCT
NZ_CP035235.1 c1326834-1317894 0      ------------------------------------------------------------------------------------------
NZ_CP035234.1:756284-765224    0      ------------------------------------------------------------------------------------------
NZ_CP038251.1:750745-759686    0      ------------------------------------------------------------------------------------------
NZ_CP035244.1:747023-755950    0      ------------------------------------------------------------------------------------------
NZ_CP035264.1:800836-809336    0      ------------------------------------------------------------------------------------------
NZ_CP035265.1:766128-774627    0      ------------------------------------------------------------------------------------------
NZ_CP035245.1 745261-753757    0      ------------------------------------------------------------------------------------------
NZ_CP035259.1:745700-753548    0      ------------------------------------------------------------------------------------------
NZ_CP035258.1:750874-758722    0      ------------------------------------------------------------------------------------------
NZ_CP035260.1:726156-733994    0      ------------------------------------------------------------------------------------------
NZ_CP035241.1:708296-716144    0      ------------------------------------------------------------------------------------------
NZ_CP035261.1:779086-786938    0      ------------------------------------------------------------------------------------------
NZ_CP035242.1:c1532236-1524385 0      ------------------------------------------------------------------------------------------
NZ_CP035243.1:777729-785580    0      ------------------------------------------------------------------------------------------
NZ_CP035237.1:c1311322-1303456 0      ------------------------------------------------------------------------------------------
NZ_CP035246.1:c1302892-1295026 0      ------------------------------------------------------------------------------------------
NZ_CP035247.1:c1301115-1293263 0      ------------------------------------------------------------------------------------------
NZ_CP035238.1:738547-746413    0      ------------------------------------------------------------------------------------------
NZ_CP035255.1:776866-784718    0      ------------------------------------------------------------------------------------------
NZ_CP035256.1:807236-815102    0      ------------------------------------------------------------------------------------------
NZ_CP038252.1:1151812-1159673  0      ------------------------------------------------------------------------------------------
NZ_CP035254.1:793365-801213    0      ------------------------------------------------------------------------------------------
NZ_CP035251.1:748733-756585    0      ------------------------------------------------------------------------------------------
NZ_CP035252.1:745670-753536    0      ------------------------------------------------------------------------------------------
NZ_CP035257.1:776317-784169    0      ------------------------------------------------------------------------------------------
NZ_CP038253.1:758105-765957    0      ------------------------------------------------------------------------------------------
NZ_CP035263.1:743050-750901    0      ------------------------------------------------------------------------------------------
NZ_CP035262.1:741595-749460    0      ------------------------------------------------------------------------------------------
NZ_CP035236.1:764915-772766    0      ------------------------------------------------------------------------------------------
NZ_CP035253.1 767286-775134    0      ------------------------------------------------------------------------------------------
NZ_CP035249.1:763080-770921    0      ------------------------------------------------------------------------------------------

NC_003028.3:777000-787000      361    CAAGAAAATTATGTGGTCTTGCCGACGGAAATTAGTAATTGTCTTCCTAAAGTAATGAAAAGATATGTCTAAAATAGTTAGACAAAAAAT
NZ_CP035248.1:765099-775111    361    CAAGAAAATTATGTGGTCTTGCCGACGGAAATTAGTAATTGTCTTCCTAAAGTAATGAAAAGATATGTCTAAAATAGTTAGACAAAAAAT
NZ_CP035235.1 c1326834-1317894 0      ------------------------------------------------------------------------------------------
NZ_CP035234.1:756284-765224    0      ------------------------------------------------------------------------------------------
NZ_CP038251.1:750745-759686    0      ------------------------------------------------------------------------------------------
NZ_CP035244.1:747023-755950    0      ------------------------------------------------------------------------------------------
NZ_CP035264.1:800836-809336    0      ------------------------------------------------------------------------------------------
NZ_CP035265.1:766128-774627    0      ------------------------------------------------------------------------------------------
NZ_CP035245.1 745261-753757    0      ------------------------------------------------------------------------------------------
NZ_CP035259.1:745700-753548    0      ------------------------------------------------------------------------------------------
NZ_CP035258.1:750874-758722    0      ------------------------------------------------------------------------------------------
NZ_CP035260.1:726156-733994    0      ------------------------------------------------------------------------------------------
NZ_CP035241.1:708296-716144    0      ------------------------------------------------------------------------------------------
NZ_CP035261.1:779086-786938    0      ------------------------------------------------------------------------------------------
NZ_CP035242.1:c1532236-1524385 0      ------------------------------------------------------------------------------------------
NZ_CP035243.1:777729-785580    0      ------------------------------------------------------------------------------------------
NZ_CP035237.1:c1311322-1303456 0      ------------------------------------------------------------------------------------------
NZ_CP035246.1:c1302892-1295026 0      ------------------------------------------------------------------------------------------
NZ_CP035247.1:c1301115-1293263 0      ------------------------------------------------------------------------------------------
NZ_CP035238.1:738547-746413    0      ------------------------------------------------------------------------------------------
NZ_CP035255.1:776866-784718    0      ------------------------------------------------------------------------------------------
NZ_CP035256.1:807236-815102    0      ------------------------------------------------------------------------------------------
NZ_CP038252.1:1151812-1159673  0      ------------------------------------------------------------------------------------------
NZ_CP035254.1:793365-801213    0      ------------------------------------------------------------------------------------------
NZ_CP035251.1:748733-756585    0      ------------------------------------------------------------------------------------------
NZ_CP035252.1:745670-753536    0      ------------------------------------------------------------------------------------------
NZ_CP035257.1:776317-784169    0      ------------------------------------------------------------------------------------------
NZ_CP038253.1:758105-765957    0      ------------------------------------------------------------------------------------------
NZ_CP035263.1:743050-750901    0      ------------------------------------------------------------------------------------------
NZ_CP035262.1:741595-749460    0      ------------------------------------------------------------------------------------------
NZ_CP035236.1:764915-772766    0      ------------------------------------------------------------------------------------------
NZ_CP035253.1 767286-775134    0      ------------------------------------------------------------------------------------------
NZ_CP035249.1:763080-770921    0      ------------------------------------------------------------------------------------------

NC_003028.3:777000-787000      451    GTTGATAATTTGTATCATTATTCTTAATTCACAAAAAACGAACGTTTAGTATTCTTCTTGCTAAGAAACTAAATTTGTTCGTTTTTTTAC
NZ_CP035248.1:765099-775111    451    GTTGATAATTTGTATCATTATTCTTAATTCACAAAAAACGAACGTTTAGTATTCTTCTTGCTAAGAAACTAAATTTGTTCGTTTTTTTAC
NZ_CP035235.1 c1326834-1317894 0      ------------------------------------------------------------------------------------------
NZ_CP035234.1:756284-765224    0      ------------------------------------------------------------------------------------------
NZ_CP038251.1:750745-759686    0      ------------------------------------------------------------------------------------------
NZ_CP035244.1:747023-755950    0      ------------------------------------------------------------------------------------------
NZ_CP035264.1:800836-809336    0      ------------------------------------------------------------------------------------------
NZ_CP035265.1:766128-774627    0      ------------------------------------------------------------------------------------------
NZ_CP035245.1 745261-753757    0      ------------------------------------------------------------------------------------------
NZ_CP035259.1:745700-753548    0      ------------------------------------------------------------------------------------------
NZ_CP035258.1:750874-758722    0      ------------------------------------------------------------------------------------------
NZ_CP035260.1:726156-733994    0      ------------------------------------------------------------------------------------------
NZ_CP035241.1:708296-716144    0      ------------------------------------------------------------------------------------------
NZ_CP035261.1:779086-786938    0      ------------------------------------------------------------------------------------------
NZ_CP035242.1:c1532236-1524385 0      ------------------------------------------------------------------------------------------
NZ_CP035243.1:777729-785580    0      ------------------------------------------------------------------------------------------
NZ_CP035237.1:c1311322-1303456 0      ------------------------------------------------------------------------------------------
NZ_CP035246.1:c1302892-1295026 0      ------------------------------------------------------------------------------------------
NZ_CP035247.1:c1301115-1293263 0      ------------------------------------------------------------------------------------------
NZ_CP035238.1:738547-746413    0      ------------------------------------------------------------------------------------------
```

### Ozkan et al. Supplementary Figures

```
NZ_CP035255.1:776866-784718    0      ------------------------------------------------------------------------------------------
NZ_CP035256.1:807236-815102    0      ------------------------------------------------------------------------------------------
NZ_CP038252.1:1151812-1159673  0      ------------------------------------------------------------------------------------------
NZ_CP035254.1:793365-801213    0      ------------------------------------------------------------------------------------------
NZ_CP035251.1:748733-756585    0      ------------------------------------------------------------------------------------------
NZ_CP035252.1:745670-753536    0      ------------------------------------------------------------------------------------------
NZ_CP035257.1:776317-784169    0      ------------------------------------------------------------------------------------------
NZ_CP038253.1:758105-765957    0      ------------------------------------------------------------------------------------------
NZ_CP035263.1:743050-750901    0      ------------------------------------------------------------------------------------------
NZ_CP035262.1:741595-749460    0      ------------------------------------------------------------------------------------------
NZ_CP035236.1:764915-772766    0      ------------------------------------------------------------------------------------------
NZ_CP035253.1 767286-775134    0      ------------------------------------------------------------------------------------------
NZ_CP035249.1:763080-770921    0      ------------------------------------------------------------------------------------------

NC_003028.3:777000-787000      541    TCTTGTAAATCTATTTTTGTTAGAGTTGATTTGGTTTACATCCGTACTTAAATTGATTTGTTAGAGCTCTACTTTTATTAAAAAAATTCA
NZ_CP035248.1:765099-775111    541    TCTTGTAAATCTATTTTTGTTAGAGTTGATTTGGTTTACATCCGTACTTAAATTGATTTGTTAGAGCTCTACTTTTATTAAAAAAATTCA
NZ_CP035235.1 c1326834-1317894 0      ------------------------------------------------------------------------------------------
NZ_CP035234.1:756284-765224    0      ------------------------------------------------------------------------------------------
NZ_CP038251.1:750745-759686    0      ------------------------------------------------------------------------------------------
NZ_CP035244.1:747023-755950    0      ------------------------------------------------------------------------------------------
NZ_CP035264.1:800836-809336    0      ------------------------------------------------------------------------------------------
NZ_CP035265.1:766128-774627    0      ------------------------------------------------------------------------------------------
NZ_CP035245.1 745261-753757    0      ------------------------------------------------------------------------------------------
NZ_CP035259.1:745700-753548    0      ------------------------------------------------------------------------------------------
NZ_CP035258.1:750874-758722    0      ------------------------------------------------------------------------------------------
NZ_CP035260.1:726156-733994    0      ------------------------------------------------------------------------------------------
NZ_CP035241.1:708296-716144    0      ------------------------------------------------------------------------------------------
NZ_CP035261.1:779086-786938    0      ------------------------------------------------------------------------------------------
NZ_CP035242.1:c1532236-1524385 0      ------------------------------------------------------------------------------------------
NZ_CP035243.1:777729-785580    0      ------------------------------------------------------------------------------------------
NZ_CP035237.1:c1311322-1303456 0      ------------------------------------------------------------------------------------------
NZ_CP035246.1:c1302892-1295026 0      ------------------------------------------------------------------------------------------
NZ_CP035247.1:c1301115-1293263 0      ------------------------------------------------------------------------------------------
NZ_CP035238.1:738547-746413    0      ------------------------------------------------------------------------------------------
NZ_CP035255.1:776866-784718    0      ------------------------------------------------------------------------------------------
NZ_CP035256.1:807236-815102    0      ------------------------------------------------------------------------------------------
NZ_CP038252.1:1151812-1159673  0      ------------------------------------------------------------------------------------------
NZ_CP035254.1:793365-801213    0      ------------------------------------------------------------------------------------------
NZ_CP035251.1:748733-756585    0      ------------------------------------------------------------------------------------------
NZ_CP035252.1:745670-753536    0      ------------------------------------------------------------------------------------------
NZ_CP035257.1:776317-784169    0      ------------------------------------------------------------------------------------------
NZ_CP038253.1:758105-765957    0      ------------------------------------------------------------------------------------------
NZ_CP035263.1:743050-750901    0      ------------------------------------------------------------------------------------------
NZ_CP035262.1:741595-749460    0      ------------------------------------------------------------------------------------------
NZ_CP035236.1:764915-772766    0      ------------------------------------------------------------------------------------------
NZ_CP035253.1 767286-775134    0      ------------------------------------------------------------------------------------------
NZ_CP035249.1:763080-770921    0      ------------------------------------------------------------------------------------------

NC_003028.3:777000-787000      631    ATTTCAAGGATAAATAAGCAGTATTCTAAAGGTACTTTTAGATGAAATAAAAGCCTTTACATGGTATAATAGAGGTAGCTCTTTAATGGA
NZ_CP035248.1:765099-775111    631    ATTTCAAGGATAAATAAGCAGTATTCTAAAGGTACTTTTAGATGAAATAAAAGCCTTTACATGGTATAATAGAGGTAGCTCTTTAATGGA
NZ_CP035235.1 c1326834-1317894 1      -------------------------------------------------------------------------------------ATGGA
NZ_CP035234.1:756284-765224    1      -------------------------------------------------------------------------------------ATGGA
NZ_CP038251.1:750745-759686    1      -------------------------------------------------------------------------------------ATGGA
NZ_CP035244.1:747023-755950    0      ------------------------------------------------------------------------------------------
NZ_CP035264.1:800836-809336    0      ------------------------------------------------------------------------------------------
NZ_CP035265.1:766128-774627    0      ------------------------------------------------------------------------------------------
NZ_CP035245.1 745261-753757    0      ------------------------------------------------------------------------------------------
NZ_CP035259.1:745700-753548    0      ------------------------------------------------------------------------------------------
NZ_CP035258.1:750874-758722    0      ------------------------------------------------------------------------------------------
NZ_CP035260.1:726156-733994    0      ------------------------------------------------------------------------------------------
NZ_CP035241.1:708296-716144    0      ------------------------------------------------------------------------------------------
NZ_CP035261.1:779086-786938    0      ------------------------------------------------------------------------------------------
NZ_CP035242.1:c1532236-1524385 0      ------------------------------------------------------------------------------------------
NZ_CP035243.1:777729-785580    0      ------------------------------------------------------------------------------------------
NZ_CP035237.1:c1311322-1303456 1      -------------------------------------------------------------------------------------ATGGA
NZ_CP035246.1:c1302892-1295026 1      -------------------------------------------------------------------------------------ATGGA
NZ_CP035247.1:c1301115-1293263 0      ------------------------------------------------------------------------------------------
NZ_CP035238.1:738547-746413    1      -------------------------------------------------------------------------------------ATGGA
NZ_CP035255.1:776866-784718    0      ------------------------------------------------------------------------------------------
NZ_CP035256.1:807236-815102    1      -------------------------------------------------------------------------------------ATGGA
NZ_CP038252.1:1151812-1159673  0      ------------------------------------------------------------------------------------------
NZ_CP035254.1:793365-801213    0      ------------------------------------------------------------------------------------------
NZ_CP035251.1:748733-756585    0      ------------------------------------------------------------------------------------------
NZ_CP035252.1:745670-753536    1      -------------------------------------------------------------------------------------ATGGA
NZ_CP035257.1:776317-784169    0      ------------------------------------------------------------------------------------------
NZ_CP038253.1:758105-765957    0      ------------------------------------------------------------------------------------------
NZ_CP035263.1:743050-750901    0      ------------------------------------------------------------------------------------------
NZ_CP035262.1:741595-749460    1      -------------------------------------------------------------------------------------ATGGA
NZ_CP035236.1:764915-772766    0      ------------------------------------------------------------------------------------------
NZ_CP035253.1 767286-775134    0      ------------------------------------------------------------------------------------------
NZ_CP035249.1:763080-770921    0      ------------------------------------------------------------------------------------------

NC_003028.3:777000-787000      721    GGTGTTTGAGTGGAAAATCTGAAGAAAATGGCAGGTATCACGGCTGCTGAATTTATCAAGGATGGGATGGTTGTAGGGCTAGGAACAGGT
NZ_CP035248.1:765099-775111    721    GGTGTTTGAGTGGAAAATCTGAAGAAAATGGCAGGTATCACGGCTGCTGAATTTATCAAGGATGGGATGGTTGTAGGGCTAGGAACAGGT
NZ_CP035235.1 c1326834-1317894 6      GGTAAATCGGTGGAAAATCTGAAGAAAATGGCAGGTATCAAGGCTGCTGAGTTCGTGAGCGATGGAATGGTCGTTGGACTTGGAACAGGC
NZ_CP035234.1:756284-765224    6      GGTAAATCGGTGGAAAATCTGAAGAAAATGGCAGGTATCAAGGCTGCTGAGTTCGTGAGCGATGGAATGGTCGTTGGACTTGGAACAGGC
NZ_CP038251.1:750745-759686    6      GGTAAATCGGTGGAAAATCTGAAGAAAATGGCAGGTATCAAGGCTGCTGAGTTCGTGAGCGATGGAATGGTCGTTGGACTTGGAACAGGC
NZ_CP035244.1:747023-755950    1      ---------GTGGAAAATCTGAAGAAAATGGCAGGTATCAAGGCTGCTGAGTTCGTGAGCGATGGAATGGTCGTTGGACTTGGAACAGGC
NZ_CP035264.1:800836-809336    1      ---------GTGGAAAATCTGAAGAAAATGGCAGGTATCAAGGCTGCTGAGTTCGTGAGCGATGGAATGGTCGTTGGACTTGGAACAGGC
NZ_CP035265.1:766128-774627    1      ---------GTGGAAAATCTGAAGAAAATGGCAGGTATCAAGGCTGCTGAGTTCGTGAGCGATGGAATGGTCGTTGGACTTGGAACAGGC
NZ_CP035245.1 745261-753757    1      ---------GTGGAAAATCTGAAGAAAATGGCAGGTATCAAGGCTGCTGAGTTCGTGAGCGATGGAATGGTCGTTGGACTTGGAACAGGC
NZ_CP035259.1:745700-753548    1      ---------GTGGAAAATCTGAAGAAAATGGCAGGTATCAAGGCTGCTGAGTTCGTGAGCGATGGAATGGTCGTTGGACTTGGAACGGGC
NZ_CP035258.1:750874-758722    1      ---------GTGGAAAATCTGAAGAAAATGGCAGGTATCAAGGCTGCTGAGTTCGTGAGCGATGGAATGGTCGTTGGACTTGGAACGGGC
NZ_CP035260.1:726156-733994    1      ---------GTGGAAAATCTGAAGAAAATGGCAGGTATCAAGGCTGCTGAGTTCGTGAGCGATGGAATGGTCGTTGGACTTGGAACAGGC
NZ_CP035241.1:708296-716144    1      ---------GTGGAAAATCTGAAGAAAATGGCAGGTATCAAGGCTGCTGAGTTCGTGAGCGATGGAATGGTCGTTGGACTTGGAACAGGC
```

### Ozkan et al. Supplementary Figures

```
NZ_CP035261.1:779086-786938    1      ---------GTGGAAAATCTGAAGAAAATGGCAGGTATCAAGGCTGCTGAGTTCGTGAGCGATGGAATGGTCGTTGGACTTGGAACAGGC
NZ_CP035242.1:c1532236-1524385 1      ---------GTGGAAAATCTGAAGAAAATGGCAGGTATCAAGGCTGCTGAGTTCGTGAGCGATGGAATGGTCGTTGGACTTGGAACAGGC
NZ_CP035243.1:777729-785580    1      ---------GTGGAAAATCTGAAGAAAATGGCAGGTATCAAGGCTGCTGAGTTCGTGAGCGATGGAATGGTCGTTGGACTTGGAACAGGC
NZ_CP035237.1:c1311322-1303456 6      GGTAAATCGGTGGAAAATCTGAAGAAAATGGCAGGTATCAAGGCTGCTGAGTTCGTGAGCGATGGAATGGTCGTTGGACTTGGAACAGGC
NZ_CP035246.1:c1302892-1295026 6      GGTAAATCGGTGGAAAATCTGAAGAAAATGGCAGGTATCAAGGCTGCTGAGTTCGTGAGCGATGGAATGGTCGTTGGACTTGGAACAGGC
NZ_CP035247.1:c1301115-1293263 1      ---------GTGGAAAATCTGAAGAAAATGGCAGGTATCAAGGCTGCTGAGTTCGTGAGCGATGGAATGGTCGTTGGACTTGGAACAGGC
NZ_CP035238.1:738547-746413    6      GGTAAATCGGTGGAAAATCTGAAGAAAATGGCAGGTATCAAGGCTGCTGAGTTCGTGAGCGATGGAATGGTCGTTGGACTTGGAACAGGC
NZ_CP035255.1:776866-784718    1      ---------GTGGAAAATCTGAAGAAAATGGCAGGTATCAAGGCTGCTGAGTTCGTGAGCGATGGAATGGTCGTTGGACTTGGAACAGGC
NZ_CP035256.1:807236-815102    6      GGTAAATCGGTGGAAAATCTGAAGAAAATGGCAGGTATCAAGGCTGCTGAGTTCGTGAGCGATGGAATGGTCGTTGGACTTGGAACAGGC
NZ_CP038252.1:1151812-1159673  1      -----------------------------------------------------------------AATGGTCGTTGGACTTGGAACAGGC
NZ_CP035254.1:793365-801213    1      ---------GTGGAAAATCTGAAGAAAATGGCAGGTATCAAGGCTGCTGAGTTCGTGAGCGATGGAATGGTCGTTGGACTTGGAACAGGC
NZ_CP035251.1:748733-756585    1      ---------GTGGAAAATCTGAAGAAAATGGCAGGTATCAAGGCTGCTGAGTTCGTGAGCGATGGAATGGTCGTTGGACTTGGAACAGGC
NZ_CP035252.1:745670-753536    6      GGTAAATCGGTGGAAAATCTGAAGAAAATGGCAGGTATCAAGGCTGCTGAGTTCGTGAGCGATGGAATGGTCGTTGGACTTGGAACAGGC
NZ_CP035257.1:776317-784169    1      ---------GTGGAAAATCTGAAGAAAATGGCAGGTATCAAGGCTGCTGAGTTCGTGAGCGATGGAATGGTCGTTGGACTTGGAACAGGC
NZ_CP038253.1:758105-765957    1      ---------GTGGAAAATCTGAAGAAAATGGCAGGTATCAAGGCTGCTGAGTTCGTGAGCGATGGAATGGTCGTTGGACTTGGAACAGGC
NZ_CP035263.1:743050-750901    1      ---------GTGGAAAATCTGAAGAAAATGGCAGGTATCAAGGCTGCTGAGTTCGTGAGCGATGGAATGGTCGTTGGACTTGGAACAGGC
NZ_CP035262.1:741595-749460    6      GGTAAATCGGTGGAAAATCTGAAGAAAATGGCAGGTATCAAGGCTGCTGAGTTCGTGAGCGATGGAATGGTCGTTGGACTTGGAACAGGC
NZ_CP035236.1:764915-772766    1      ---------GTGGAAAATCTGAAGAAAATGGCAGGTATCAAGGCTGCTGAGTTCGTGAGCGATGGAATGGTCGTTGGACTTGGAACAGGC
NZ_CP035253.1 767286-775134    1      ---------GTGGAAAATCTGAAGAAAATGGCAGGTATCAAGGCTGCTGAGTTCGTGAGCGATGGAATGGTCGTTGGACTTGGAACAGGC
NZ_CP035249.1:763080-770921    1      ---------GTGGAAAATCTGAAGAAAATGGCAGGTATCAAGGCTGCTGAGTTCGTGAGCGATGGAATGGTCGTTGGACTTGGAACAGGC

NC_003028.3:777000-787000      811    TCTACTGCCTATTATTTTGTCGAAGAAATCGGTCGTCGAATCAAGGAAGAAGGCTTGCAGATTACAGCTGTGACGACTTCTAGTGTGACC
NZ_CP035248.1:765099-775111    811    TCTACTGCCTATTATTTTGTCGAAGAAATCGGTCGTCGAATCAAGGAAGAAGGCTTGCAGATTACAGCTGTGACGACTTCTAGTGTGACC
NZ_CP035235.1 c1326834-1317894 96     TCGACTGCCTATTATTTTGTCGAAGAAATCGGTCGTCGTATCAAGGAAGAAGGATTGCAGATTACAGCTGTAACAACTTCTAGTGTGACC
NZ_CP035234.1:756284-765224    96     TCGACTGCCTATTATTTTGTCGAAGAAATCGGTCGTCGTATCAAGGAAGAAGGATTGCAGATTACAGCTGTAACAACTTCTAGTGTGACC
NZ_CP038251.1:750745-759686    96     TCGACTGCCTATTATTTTGTCGAAGAAATCGGTCGTCGTATCAAGGAAGAAGGATTGCAGATTACAGCTGTAACAACTTCTAGTGTGACC
NZ_CP035244.1:747023-755950    82     TCGACTGCCTATTATTTTGTCGAAGAAATCGGTCGTCGTATCAAGGAAGAAGGATTGCAGATTACAGCTGTAACAACTTCTAGTGTGACC
NZ_CP035264.1:800836-809336    82     TCGACTGCCTATTATTTTGTCGAAGAAATCGGTCGTCGTATCAAGGAAGAAGGATTACAGATTACAGCTGTAACAACTTCTAGTGTGACC
NZ_CP035265.1:766128-774627    82     TCGACTGCCTATTATTTTGTCGAAGAAATCGGTCGTCGTATCAAGGAAGAAGGATTACAGATTACAGCTGTAACAACTTCTAGTGTGACC
NZ_CP035245.1 745261-753757    82     TCGACTGCCTATTATTTTGTCGAAGAAATCGGTCGTCGTATCAAGGAAGAAGGATTGCAGATTACAGCTGTAACAACTTCTAGTGTGACC
NZ_CP035259.1:745700-753548    82     TCGACTGCCTATTATTTTGTCGAAGAAATCGGTCGTCGTATCAAGGAAGAAGGATTGCAGATTACAGCTGTAACAACTTCTAGTGTGACC
NZ_CP035258.1:750874-758722    82     TCGACTGCCTATTATTTTGTCGAAGAAATCGGTCGTCGTATCAAGGAAGAAGGATTGCAGATTACAGCTGTAACAACTTCTAGTGTGACC
NZ_CP035260.1:726156-733994    82     TCGACTGCCTATTATTTTGTCGAAGAAATCGGTCGTCGTATCAAGGAAGAAGGATTACAGATTACAGCTGTAACAACTTCTAGTGTGACC
NZ_CP035241.1:708296-716144    82     TCGACTGCCTATTATTTTGTCGAAGAAATCGGTCGTCGTATCAAGGAAGAAGGATTGCAGATTACAGCTGTAACAACTTCTAGTGTGACC
NZ_CP035261.1:779086-786938    82     TCGACTGCCTATTATTTTGTCGAAGAAATCGGTCGTCGTATCAAGGAAGAAGGATTGCAGATTACAGCTGTAACAACTTCTAGTGTGACC
NZ_CP035242.1:c1532236-1524385 82     TCGACTGCCTATTATTTTGTCGAAGAAATCGGTCGTCGTATCAAGGAAGAAGGATTACAGATTACAGCTGTAACAACTTCTAGTGTGACC
NZ_CP035243.1:777729-785580    82     TCGACTGCCTATTATTTTGTCGAAGAAATCGGTCGTCGTATCAAGGAAGAAGGATTACAGATTACAGCTGTAACAACTTCTAGTGTGACC
NZ_CP035237.1:c1311322-1303456 96     TCGACTGCCTATTATTTTGTCGAAGAAATCGGTCGTCGTATCAAGGAAGAAGGATTGCAGATTACAGCTGTAACAACTTCTAGTGTGACC
NZ_CP035246.1:c1302892-1295026 96     TCGACTGCCTATTATTTTGTCGAAGAAATCGGTCGTCGTATCAAGGAAGAAGGATTGCAGATTACAGCTGTAACAACTTCTAGTGTGACC
NZ_CP035247.1:c1301115-1293263 82     TCGACTGCCTATTATTTTGTCGAAGAAATCGGTCGTCGTATCAAGGAAGAAGGATTGCAGATTACAGCTGTAACAACTTCTAGTGTGACC
NZ_CP035238.1:738547-746413    96     TCGACTGCCTATTATTTTGTCGAAGAAATCGGTCGTCGTATCAAGGAAGAAGGATTGCAGATTACAGCTGTAACAACTTCTAGTGTGACC
NZ_CP035255.1:776866-784718    82     TCGACTGCCTATTATTTTGTCGAAGAAATCGGTCGTCGTATCAAGGAAGAAGGATTACAGATTACAGCTGTAACAACTTCTAGTGTGACC
NZ_CP035256.1:807236-815102    96     TCGACTGCCTATTATTTTGTCGAAGAAATCGGTCGTCGTATCAAGGAAGAAGGATTACAGATTACAGCTGTAACAACTTCTAGTGTGACC
NZ_CP038252.1:1151812-1159673  26     TCGACTGCCTATTATTTTGTCGAAGAAATCGGTCGTCGTATCAAGGAAGAAGGATTGCAGATTACAGCTGTAACAACTTCTAGTGTGACC
NZ_CP035254.1:793365-801213    82     TCGACTGCCTATTATTTTGTCGAAGAAATCGGTCGTCGTATCAAGGAAGAAGGATTGCAGATTACAGCTGTAACAACTTCTAGTGTGACC
NZ_CP035251.1:748733-756585    82     TCGACTGCCTATTATTTTGTCGAAGAAATCGGTCGTCGTATCAAGGAAGAAGGATTGCAGATTACAGCTGTAACAACTTCTAGTGTGACC
NZ_CP035252.1:745670-753536    96     TCGACTGCCTATTATTTTGTCGAAGAAATCGGTCGTCGTATCAAGGAAGAAGGATTGCAGATTACAGCTGTAACAACTTCTAGTGTGACC
NZ_CP035257.1:776317-784169    82     TCGACTGCCTATTATTTTGTCGAAGAAATCGGTCGTCGTATCAAGGAAGAAGGATTGCAGATTACAGCTGTAACAACTTCTAGTGTGACC
NZ_CP038253.1:758105-765957    82     TCGACTGCCTATTATTTTGTCGAAGAAATCGGTCGTCGTATCAAGGAAGAAGGATTGCAGATTACAGCTGTAACAACTTCTAGTGTGACC
NZ_CP035263.1:743050-750901    82     TCGACTGCCTATTATTTTGTCGAAGAAATCGGTCGTCGTATCAAGGAAGAAGGATTGCAGATTACAGCTGTAACAACTTCTAGTGTGACC
NZ_CP035262.1:741595-749460    96     TCGACTGCCTATTATTTTGTCGAAGAAATCGGTCGTCGTATCAAGGAAGAAGGATTGCAGATTACAGCTGTAACAACTTCTAGTGTGACC
NZ_CP035236.1:764915-772766    82     TCGACTGCCTATTATTTTGTCGAAGAAATCGGTCGTCGTATCAAGGAAGAAGGATTGCAGATTACAGCTGTAACAACTTCTAGTGTGACC
NZ_CP035253.1 767286-775134    82     TCGACTGCCTATTATTTTGTCGAAGAAATCGGTCGTCGTATCAAGGAAGAAGGATTGCAGATTACAGCTGTAACAACTTCTAGTGTGACC
NZ_CP035249.1:763080-770921    82     TCGACTGCCTATTATTTTGTCGAAGAAATCGGTCGTCGTATCAAGGAAGAAGGATTGCAGATTACAGCTGTAACAACTTCTAGTGTGACC

NC_003028.3:777000-787000      901    AGTAAACAGGCTGAAGGGCTCAATATCCCGCTCAAGTCTATTGACCAAGTAGACTTTGTCGATGTGACAGTCGACGGGGCGGATGAAGTG
NZ_CP035248.1:765099-775111    901    AGTAAACAGGCTGAAGGGCTCAATATCCCGCTCAAGTCTATTGACCAAGTAGACTTTGTCGATGTGACAGTCGACGGGGCGGATGAAGTG
NZ_CP035235.1 c1326834-1317894 186    ACTAAACAGGCAGAAGGCCTTAATATCCCGCTCAAGTCTATTGACCAAGTAGACTTTGTCGATGTGACGGTTGATGGCGCGGATGAAGTG
NZ_CP035234.1:756284-765224    186    ACTAAACAGGCAGAAGGCCTTAATATCCCGCTCAAGTCTATTGACCAAGTAGACTTTGTCGATGTGACGGTTGATGGCGCGGATGAAGTG
NZ_CP038251.1:750745-759686    186    ACTAAACAGGCAGAAGGCCTTAATATCCCGCTCAAGTCTATTGACCAAGTAGACTTTGTCGATGTGACGGTTGATGGCGCGGATGAAGTG
NZ_CP035244.1:747023-755950    172    ACTAAACAGGCAGAAGGCCTTAATATCCCGCTCAAGTCTATTGACCAAGTAGACTTTGTCGATGTGACGGTTGATGGCGCGGATGAAGTG
NZ_CP035264.1:800836-809336    172    ACTAAACAGGCAGAAGGCCTTAATATCCCGCTCAAGTCTATTGACCAAGTAGACTTTGTCGATGTGACGGTTGATGGCGCGGATGAAGTG
NZ_CP035265.1:766128-774627    172    ACTAAACAGGCAGAAGGCCTTAATATCCCGCTCAAGTCTATTGACCAAGTAGACTTTGTCGATGTGACGGTTGATGGCGCGGATGAAGTG
NZ_CP035245.1 745261-753757    172    ACTAAACAGGCAGAAGGCCTTAATATCCCGCTCAAGTCTATTGACCAAGTAGACTTTGTCGATGTGACGGTTGATGGCGCGGATGAAGTG
NZ_CP035259.1:745700-753548    172    ACTAAACAGGCAGAAGGCCTTAATATCCCGCTCAAGTCTATTGACCAAGTAGACTTTGTCGATGTGACGGTTGATGGCGCGGATGAAGTG
NZ_CP035258.1:750874-758722    172    ACTAAACAGGCAGAAGGCCTTAATATCCCGCTCAAGTCTATTGACCAAGTAGACTTTGTCGATGTGACGGTTGATGGCGCGGATGAAGTG
NZ_CP035260.1:726156-733994    172    ACTAAACAGGCAGAAGGCCTTAATATCCCGCTCAAGTCTATTGACCAAGTAGACTTTGTCGATGTGACGGTTGATGGCGCGGATGAAGTG
NZ_CP035241.1:708296-716144    172    ACTAAACAGGCAGAAGGCCTTAATATCCCGCTCAAGTCTATTGACCAAGTAGACTTTGTCGATGTGACGGTTGATGGCGCAGATGAAGTG
NZ_CP035261.1:779086-786938    172    ACTAAACAGGCAGAAGGCCTTAATATCCCGCTCAAGTCTATTGACCAAGTAGACTTTGTCGATGTGACGGTTGATGGCGCGGATGAAGTG
NZ_CP035242.1:c1532236-1524385 172    ACTAAACAGGCAGAAGGCCTTAATATCCCGCTCAAGTCTATTGACCAAGTAGACTTTGTCGATGTGACGGTTGATGGCGCGGATGAAGTG
NZ_CP035243.1:777729-785580    172    ACTAAACAGGCAGAAGGCCTTAATATCCCGCTCAAGTCTATTGACCAAGTAGACTTTGTCGATGTGACGGTTGATGGCGCGGATGAAGTG
NZ_CP035237.1:c1311322-1303456 186    ACTAAACAGGCAGAAGGCCTTAATATCCCGCTCAAGTCTATTGACCAAGTAGACTTTGTCGATGTGACGGTTGATGGCGCGGATGAAGTG
NZ_CP035246.1:c1302892-1295026 186    ACTAAACAGGCAGAAGGCCTTAATATCCCGCTCAAGTCTATTGACCAAGTAGATTTTGTCGATGTGACGGTTGATGGCGCGGATGAAGTG
NZ_CP035247.1:c1301115-1293263 172    ACTAAACAGGCAGAAGGCCTTAATATCCCGCTCAAGTCTATTGACCAAGTAGATTTTGTCGATGTGACGGTTGATGGCGCGGATGAAGTG
NZ_CP035238.1:738547-746413    186    ACTAAACAGGCAGAAGGCCTTAATATCCCGCTCAAGTCTATTGACCAAGTAGACTTTGTCGATGTGACGGTTGATGGCGCGGATGAAGTG
NZ_CP035255.1:776866-784718    172    ACTAAACAGGCAGAAGGCCTTAATATCCCGCTCAAGTCTATTGACCAAGTAGACTTTGTCGATGTGACGGTTGATGGCGCGGATGAAGTG
NZ_CP035256.1:807236-815102    186    ACTAAACAGGCAGAAGGCCTTAATATCCCGCTCAAGTCTATTGACCAAGTAGACTTTGTCGATGTGACGGTTGATGGCGCGGATGAAGTG
NZ_CP038252.1:1151812-1159673  116    ACTAAACAGGCAGAAGGCCTTAATATCCCGCTCAAGTCTATTGACCAAGTAGATTTTGTCGATGTGACGGTTGATGGCGCGGATGAAGTG
NZ_CP035254.1:793365-801213    172    ACTAAACAGGCAGAAGGCCTTAATATCCCGCTCAAGTCTATTGACCAAGTAGACTTTGTCGATGTGACGGTTGATGGCGCGGATGAAGTG
NZ_CP035251.1:748733-756585    172    ACTAAACAGGCAGAAGGCCTTAATATCCCGCTCAAGTCTATTGACCAAGTAGACTTTGTCGATGTGACGGTTGATGGCGCGGATGAAGTG
NZ_CP035252.1:745670-753536    186    ACTAAACAGGCAGAAGGCCTTAATATCCCGCTCAAGTCTATTGACCAAGTAGACTTTGTCGATGTGACGGTTGATGGCGCGGATGAAGTG
NZ_CP035257.1:776317-784169    172    ACTAAACAGGCAGAAGGCCTTAATATCCCGCTCAAGTCTATTGACCAAGTAGACTTTGTCGATGTGACGGTTGATGGCGCGGATGAAGTG
NZ_CP038253.1:758105-765957    172    ACTAAACAGGCAGAAGGCCTTAATATCCCGCTCAAGTCTATTGACCAAGTAGACTTTGTCGATGTGACGGTTGATGGCGCGGATGAAGTG
NZ_CP035263.1:743050-750901    172    ACTAAACAGGCAGAAGGCCTTAATATCCCGCTCAAGTCTATTGACCAAGTAGACTTTGTCGATGTGACGGTTGATGGCGCGGATGAAGTG
NZ_CP035262.1:741595-749460    186    ACTAAACAGGCAGAAGGCCTTAATATCCCGCTCAAGTCTATTGACCAAGTAGACTTTGTCGATGTGACGGTTGATGGCGCGGATGAAGTG
NZ_CP035236.1:764915-772766    172    ACTAAACAGGCAGAAGGCCTTAATATCCCGCTCAAGTCTATTGACCAAGTAGACTTTGTCGATGTGACGGTTGATGGCGCGGATGAAGTG
NZ_CP035253.1 767286-775134    172    ACTAAACAGGCAGAAGGCCTTAATATCCCGCTCAAGTCTATTGACCAAGTAGACTTTGTCGATGTGACGGTTGATGGCGCGGATGAAGTG
NZ_CP035249.1:763080-770921    172    ACTAAACAGGCAGAAGGCCTTAATATCCCGCTCAAGTCTATTGACCAAGTAGACTTTGTCGATGTGACGGTTGATGGCGCGGATGAAGTG

NC_003028.3:777000-787000      991    GATAGTCAGTTTAATGGAATCAAAGGCGGTGGTGGTGCCCTTCTCATGGAAAAGGTGGTCGCAACACCATCAAAAGAATACATTTGGGTG
NZ_CP035248.1:765099-775111    991    GATAGTCAGTTTAACGGAATCAAAGGCGGTGGTGGTGCCCTTCTGATGGAGAAGGTTGTGGCAACGCCCTCAAAAGAATACATTTGGGTG
NZ_CP035235.1 c1326834-1317894 276    GATAGTCAGTTTAATGGAATCAAAGGCGGTGGTGGTGCCCTTCTCATGGAAAAGGTGGTCGCAACACCATCAAAAGAATACATTTGGGTG
NZ_CP035234.1:756284-765224    276    GATAGTCAGTTTAATGGAATCAAAGGCGGTGGTGGTGCCCTTCTCATGGAAAAGGTGGTCGCAACACCATCAAAAGAATACATTTGGGTG
NZ_CP038251.1:750745-759686    276    GATAGTCAGTTTAATGGAATCAAAGGCGGTGGTGGTGCCCTTCTCATGGAAAAGGTGGTCGCAACACCATCAAAAGAATACATTTGGGTG
NZ_CP035244.1:747023-755950    262    GATAGTCAGTTTAATGGAATCAAAGGCGGTGGTGGTGCCCTTCTCATGGAAAAGGTGGTCGCAACACCATCAAAAGAATACATTTGGGTG
```

```
NZ_CP035264.1:800836-809336    262    GATAGTCAGTTTAATGGAATTAAAGGCGGTGGTGGTGCCCTTCTCATGGAAAAGGTGGTCGCAACACCATCAAAAGAATACATTTGGGTG
NZ_CP035265.1:766128-774627    262    GATAGTCAGTTTAATGGAATTAAAGGCGGTGGTGGTGCCCTTCTCATGGAAAAGGTGGTCGCAACACCATCAAAAGAATACATTTGGGTG
NZ_CP035245.1 745261-753757    262    GATAGTCAGTTTAATGGAATCAAAGGCGGTGGTGGTGCCCTTCTCATGGAAAAGGTGGTCGCAACACCATCAAAAGAATACATTTGGGTG
NZ_CP035259.1:745700-753548    262    GATAGTCAGTTTAATGGAATTAAAGGCGGTGGTGGTGCCCTTCTCATGGAAAAGGTGGTCGCAACACCATCAAAAGAATACATTTGGGTG
NZ_CP035258.1:750874-758722    262    GATAGTCAGTTTAATGGAATTAAAGGCGGTGGTGGTGCCCTTCTCATGGAAAAGGTGGTCGCAACACCATCAAAAGAATACATTTGGGTG
NZ_CP035260.1:726156-733994    262    GATAGTCAGTTTAATGGAATCAAAGGCGGTGGTGGTGCCCTTCTCATGGAAAAGGTGGTCGCAACACCATCAAAAGAATACATTTGGGTG
NZ_CP035241.1:708296-716144    262    GATAGTCAGTTTAATGGAATCAAAGGCGGTGGTGGTGCCCTTCTCATGGAAAAGGTGGTCGCAACACCATCAAAAGAATACATTTGGGTG
NZ_CP035261.1:779086-786938    262    GATAGTCAGTTTAATGGAATCAAAGGCGGTGGTGGTGCCCTTCTCATGGAAAAGGTGGTCGCAACACCATCAAAAGAATACATTTGGGTG
NZ_CP035242.1:c1532236-1524385 262    GATAGTCAGTTTAATGGAATCAAAGGCGGTGGTGGTGCCCTTCTCATGGAAAAGGTGGTCGCAACACCATCAAAAGAATACATTTGGGTG
NZ_CP035243.1:777729-785580    262    GATAGTCAGTTTAATGGAATCAAAGGCGGTGGTGGTGCCCTTCTCATGGAAAAGGTGGTCGCAACACCATCAAAAGAATACATTTGGGTG
NZ_CP035237.1:c1311322-1303456 276    GATAGTCAGTTTAATGGAATCAAAGGCGGTGGTGGTGCCCTTCTCATGGAAAAGGTGGTCGCAACACCATCAAAAGAATACATTTGGGTG
NZ_CP035246.1:c1302892-1295026 276    GATAGTCAGTTTAATGGAATCAAAGGCGGTGGTGGTGCCCTTCTCATGGAAAAGGTGGTCGCAACACCATCAAAAGAATACATTTGGGTG
NZ_CP035247.1:c1301115-1293263 262    GATAGTCAGTTTAATGGAATCAAAGGCGGTGGTGGTGCCCTTCTCATGGAAAAGGTGGTCGCAACACCATCAAAAGAATACATTTGGGTG
NZ_CP035238.1:738547-746413    276    GATAGTCAGTTTAATGGAATCAAAGGCGGTGGTGGTGCCCTTCTCATGGAAAAGGTGGTCGCAACACCATCAAAAGAATACATTTGGGTG
NZ_CP035255.1:776866-784718    262    GATAGTCAGTTTAATGGAATCAAAGGCGGTGGTGGTGCCCTTCTCATGGAAAAGGTGGTCGCAACACCATCAAAAGAATACATTTGGGTG
NZ_CP035256.1:807236-815102    276    GATAGTCAGTTTAATGGAATCAAAGGCGGTGGTGGTGCCCTTCTCATGGAAAAGGTGGTCGCAACACCATCAAAAGAATACATTTGGGTG
NZ_CP038252.1:1151812-1159673  206    GATAGTCAGTTTAATGGAATCAAAGGCGGTGGTGGTGCCCTTCTCATGGAAAAGGTGGTCGCAACACCATCAAAAGAATACATTTGGGTG
NZ_CP035254.1:793365-801213    262    GATAGTCAGTTTAATGGAATCAAAGGCGGTGGTGGTGCCCTTCTCATGGAAAAGGTGGTCGCAACACCATCAAAAGAATACATTTGGGTG
NZ_CP035251.1:748733-756585    262    GATAGTCAGTTTAATGGAATCAAAGGCGGTGGTGGTGCCCTTCTCATGGAAAAGGTGGTCGCAACACCATCAAAAGAATACATTTGGGTG
NZ_CP035252.1:745670-753536    276    GATAGTCAGTTTAATGGAATCAAAGGCGGTGGTGGTGCCCTTCTCATGGAAAAGGTGGTCGCAACACCATCAAAAGAATACATTTGGGTG
NZ_CP035257.1:776317-784169    262    GATAGTCAGTTTAATGGAATCAAAGGCGGTGGTGGTGCCCTTCTCATGGAAAAGGTGGTCGCAACACCATCAAAAGAATACATTTGGGTG
NZ_CP038253.1:758105-765957    262    GATAGTCAGTTTAATGGAATCAAAGGCGGTGGTGGTGCCCTTCTCATGGAAAAGGTGGTCGCAACACCATCAAAAGAATACATTTGGGTG
NZ_CP035263.1:743050-750901    262    GATAGTCAGTTTAATGGAATCAAAGGCGGTGGTGGTGCCCTTCTCATGGAAAAGGTGGTCGCAACACCATCAAAAGAATACATTTGGGTG
NZ_CP035262.1:741595-749460    276    GATAGTCAGTTTAATGGAATCAAAGGCGGTGGTGGTGCCCTTCTCATGGAAAAGGTGGTCGCAACACCATCAAAAGAATACATTTGGGTG
NZ_CP035236.1:764915-772766    262    GATAGTCAGTTTAATGGAATCAAAGGCGGTGGTGGTGCCCTTCTCATGGAAAAGGTGGTCGCAACACCATCAAAAGAATACATTTGGGTG
NZ_CP035253.1 767286-775134    262    GATAGTCAGTTTAATGGAATCAAAGGCGGTGGTGGTGCCCTTCTCATGGAAAAGGTGGTCGCAACACCATCAAAAGAATACATTTGGGTG
NZ_CP035249.1:763080-770921    262    GATAGTCAGTTTAATGGAATCAAAGGCGGTGGTGGTGCCCTTCTCATGGAAAAGGTGGTCGCAACACCATCAAAAGAATACATTTGGGTG

NC_003028.3:777000-787000      1081   GTGGATGAAAGCAAGCTGGTCGAAAAACTAGGTGCTTTTAAATTGCCAGTAGAAGTGGTTCAGTATGGTGCAGAGCAGGTCTTTCGTCAT
NZ_CP035248.1:765099-775111    1081   GTGGATGAAAGCAAGCTAGTCGAGAAACTGGGTGCTTTTAAATTGCCAGTAGAAGTGGTTCAGTATGGCGCAGAACAGGTCTTTCGTCGT
NZ_CP035235.1 c1326834-1317894 366    GTGGATGAAAGCAAGCTGGTCGAAAAACTAGGTGCTTTTAAATTGCCAGTAGAAGTGGTTCAGTATGGTGCAGAGCAGGTCTTTCGTCAT
NZ_CP035234.1:756284-765224    366    GTGGATGAAAGCAAGCTGGTCGAAAAACTAGGTGCTTTTAAATTGCCAGTAGAAGTGGTTCAGTATGGTGCAGAGCAGGTCTTTCGTCAT
NZ_CP038251.1:750745-759686    366    GTGGATGAAAGCAAGCTGGTCGAAAAACTAGGTGCTTTTAAATTGCCAGTAGAAGTGGTTCAGTATGGTGCAGAGCAGGTCTTTCGTCAT
NZ_CP035244.1:747023-755950    352    GTGGATGAAAGCAAGCTGGTCGAAAAACTAGGTGCTTTTAAATTGCCAGTAGAAGTGGTTCAGTATGGTGCAGAGCAGGTCTTTCGTCAT
NZ_CP035264.1:800836-809336    352    GTGGATGAAAGCAAGCTGGTCGAAAAACTAGGTGCTTTTAAATTGCCAGTAGAAGTGGTTCAGTATGGTGCAGAGCAGGTCTTTCGTCAT
NZ_CP035265.1:766128-774627    352    GTGGATGAAAGCAAGCTGGTCGAAAAACTAGGTGCTTTTAAATTGCCAGTAGAAGTGGTTCAGTATGGTGCAGAGCAGGTCTTTCGTCAT
NZ_CP035245.1 745261-753757    352    GTGGATGAAAGCAAGCTGGTCGAAAAACTAGGTGCTTTTAAATTGCCAGTAGAAGTGGTTCAGTATGGTGCAGAGCAGGTCTTTCGTCAT
NZ_CP035259.1:745700-753548    352    GTGGATGAAAGCAAGCTGGTCGAAAAACTAGGTGCTTTTAAATTGCCAGTAGAAGTGGTTCAGTATGGTGCAGAGCAGGTCTTTCGTCAT
NZ_CP035258.1:750874-758722    352    GTGGATGAAAGCAAGCTGGTCGAAAAACTAGGTGCTTTTAAATTGCCAGTAGAAGTGGTTCAGTATGGTGCAGAGCAGGTCTTTCGTCAT
NZ_CP035260.1:726156-733994    352    GTGGATGAAAGCAAGCTGGTCGAAAAACTAGGTGCTTTTAAATTGCCAGTAGAAGTGGTTCAGTATGGTGCAGAGCAGGTCTTTCGTCAT
NZ_CP035241.1:708296-716144    352    GTGGATGAAAGCAAGCTGGTCGAAAAACTAGGTGCTTTTAAATTGCCAGTAGAAGTGGTTCAGTATGGTGCAGAGCAGGTCTTTCGTCAT
NZ_CP035261.1:779086-786938    352    GTGGATGAAAGCAAGCTGGTCGAAAAACTAGGTGCTTTTAAATTGCCAGTAGAAGTGGTTCAGTATGGTGCAGAGCAGGTCTTTCGTCAT
NZ_CP035242.1:c1532236-1524385 352    GTGGATGAAAGCAAGCTGGTCGAAAAACTAGGTGCTTTTAAATTGCCAGTAGAAGTGGTTCAGTATGGTGCAGAGCAGGTCTTTCGTCAT
NZ_CP035243.1:777729-785580    352    GTGGATGAAAGCAAGCTGGTCGAAAAACTAGGTGCTTTTAAATTGCCAGTAGAAGTGGTTCAGTATGGTGCAGAGCAGGTCTTTCGTCAT
NZ_CP035237.1:c1311322-1303456 366    GTGGATGAAAGCAAGCTGGTCGAAAAACTAGGTGCTTTTAAATTGCCAGTAGAAGTGGTTCAGTATGGTGCAGAGCAGGTCTTTCGTCAT
NZ_CP035246.1:c1302892-1295026 366    GTGGATGAAAGCAAGCTGGTCGAAAAACTAGGTGCTTTTAAATTGCCAGTAGAAGTGGTTCAGTATGGTGCAGAGCAGGTCTTTCGTCAT
NZ_CP035247.1:c1301115-1293263 352    GTGGATGAAAGCAAGCTGGTCGAAAAACTAGGTGCTTTTAAATTGCCAGTAGAAGTGGTTCAGTATGGTGCAGAGCAGGTCTTTCGTCAT
NZ_CP035238.1:738547-746413    366    GTGGATGAAAGCAAGCTGGTCGAAAAACTAGGTGCTTTTAAATTGCCAGTAGAAGTGGTTCAGTATGGTGCAGAGCAGGTCTTTCGTCAT
NZ_CP035255.1:776866-784718    352    GTGGATGAAAGCAAGCTGGTCGAAAAACTAGGTGCTTTTAAATTGCCAGTAGAAGTGGTTCAGTATGGTGCAGAGCAGGTCTTTCGTCAT
NZ_CP035256.1:807236-815102    366    GTGGATGAAAGCAAGCTGGTCGAAAAACTAGGTGCTTTTAAATTGCCAGTAGAAGTGGTTCAGTATGGTGCAGAGCAGGTCTTTCGTCAT
NZ_CP038252.1:1151812-1159673  296    GTGGATGAAAGCAAGCTGGTCGAAAAACTAGGTGCTTTTAAATTGCCAGTAGAAGTGGTTCAGTATGGTGCAGAGCAGGTCTTTCGTCAT
NZ_CP035254.1:793365-801213    352    GTGGATGAAAGCAAGCTGGTCGAAAAACTAGGTGCTTTTAAATTGCCAGTAGAAGTGGTTCAGTATGGTGCAGAGCAGGTCTTTCGTCAT
NZ_CP035251.1:748733-756585    352    GTGGATGAAAGCAAGCTGGTCGAAAAACTAGGTGCTTTTAAATTGCCAGTAGAAGTGGTTCAGTATGGTGCAGAGCAGGTCTTTCGTCAT
NZ_CP035252.1:745670-753536    366    GTGGATGAAAGCAAGCTGGTCGAAAAACTAGGTGCTTTTAAATTGCCAGTAGAAGTGGTTCAGTATGGTGCAGAGCAGGTCTTTCGTCAT
NZ_CP035257.1:776317-784169    352    GTGGATGAAAGCAAGCTGGTCGAAAAACTAGGTGCTTTTAAATTGCCAGTAGAAGTGGTTCAGTATGGTGCAGAGCAGGTCTTTCGTCAT
NZ_CP038253.1:758105-765957    352    GTGGATGAAAGCAAGCTGGTCGAAAAACTAGGTGCTTTTAAATTGCCAGTAGAAGTGGTTCAGTATGGTGCAGAGCAGGTCTTTCGTCAT
NZ_CP035263.1:743050-750901    352    GTGGATGAAAGCAAGCTGGTCGAAAAACTAGGTGCTTTTAAATTGCCAGTAGAAGTGGTTCAGTATGGTGCAGAGCAGGTCTTTCGTCAT
NZ_CP035262.1:741595-749460    366    GTGGATGAAAGCAAGCTGGTCGAAAAACTAGGTGCTTTTAAATTGCCAGTAGAAGTGGTTCAGTATGGTGCAGAGCAGGTCTTTCGTCAT
NZ_CP035236.1:764915-772766    352    GTGGATGAAAGCAAGCTGGTCGAAAAACTAGGTGCTTTTAAATTGCCAGTAGAAGTGGTTCAGTATGGTGCAGAGCAGGTCTTTCGTCAT
NZ_CP035253.1 767286-775134    352    GTGGATGAAAGCAAGCTGGTCGAAAAACTAGGTGCTTTTAAATTGCCAGTAGAAGTGGTTCAGTATGGTGCAGAGCAGGTCTTTCGTCAT
NZ_CP035249.1:763080-770921    352    GTGGATGAAAGCAAGCTGGTCGAAAAACTAGGTGCTTTTAAATTGCCAGTAGAAGTGGTTCAGTATGGTGCAGAGCAGGTCTTTCGTCAT

NC_003028.3:777000-787000      1171   TTTGAACGAGCTGGCTACAAACCAAGTTTCCGTGAAAAAGACGGCCAACGTTTTGTGACCGATATGCAGAATTTTATCATTGACCTCGCC
NZ_CP035248.1:765099-775111    1171   TTTGAGCGAGCTGGCTACAAACCAAGTTTCCGTGAAAAAGACGGCCAACGTTTTGTGACCGATATGCAGAATTTTATCATTGACCTTGCT
NZ_CP035235.1 c1326834-1317894 456    TTTGAACGAGCTGGCTACAAACCAAGTTTCCGTGAAAAAGACGGCCAACGTTTTGTGACGGATATGCAGAATTTTATCATTGACCTCGCC
NZ_CP035234.1:756284-765224    456    TTTGAACGAGCTGGCTACAAACCAAGTTTCCGTGAAAAAGACGGCCAACGTTTTGTGACGGATATGCAGAATTTTATCATTGACCTCGCC
NZ_CP038251.1:750745-759686    456    TTTGAACGAGCTGGCTACAAACCAAGTTTCCGTGAAAAAGACGGCCAACGTTTTGTGACGGATATGCAGAATTTTATCATTGACCTCGCC
NZ_CP035244.1:747023-755950    442    TTTGAACGAGCTGGCTACAAACCAAGTTTCCGTGAAAAAGACGGCCAACGTTTTGTGACGGATATGCAGAATTTTATCATTGACCTCGCC
NZ_CP035264.1:800836-809336    442    TTTGAACGAGCTGGCTACAAACCAAGTTTTCGTGAAAAAGACGGCCAACGTTTTGTGACGGATATGCAGAATTTTATCATTGACCTCGCC
NZ_CP035265.1:766128-774627    442    TTTGAACGAGCTGGCTACAAACCAAGTTTCCGTGAAAAAGACGGCCAACGTTTTGTGACGGATATGCAGAATTTTATCATTGACCTCGCC
NZ_CP035245.1 745261-753757    442    TTTGAACGAGCTGGCTACAAACCAAGTTTCCGTGAAAAAGACGGCCAACGTTTTGTGACGGATATGCAGAATTTTATCATTGACCTCGCC
NZ_CP035259.1:745700-753548    442    TTTGAACGAGCTGGCTACAAACCAAGTTTCCGTGAAAAAGACGGCCAACGTTTTGTGACGGATATGCAGAATTTTATCATTGACCTCGCC
NZ_CP035258.1:750874-758722    442    TTTGAACGAGCTGGCTACAAACCAAGTTTCCGTGAAAAAGACGGCCAACGTTTTGTGACGGATATGCAGAATTTTATCATTGACCTCGCC
NZ_CP035260.1:726156-733994    442    TTTGAACGAGCTGGCTACAAACCAAGTTTCCGTGAAAAAGACGGCCAACGTTTTGTGACGGATATGCAGAATTTTATCATTGACCTCGCC
NZ_CP035241.1:708296-716144    442    TTTGAACGAGCTGGCTACAAACCAAGTTTCCGTGAAAAAGACGGCCAACGTTTTGTGACGGATATGCAGAATTTTATCATTGACCTCGCC
NZ_CP035261.1:779086-786938    442    TTTGAACGAGCTGGCTACAAACCAAGTTTCCGTGAAAAAGACGGTCAACGTTTTGTGACGGATATGCAGAATTTTATCATTGACCTCGCC
NZ_CP035242.1:c1532236-1524385 442    TTTGAACGAGCTGGCTACAAACCAAGTTTCCGTGAAAAAGACGGCCAACGTTTTGTGACGGATATGCAGAATTTTATCATTGACCTCGCC
NZ_CP035243.1:777729-785580    442    TTTGAACGAGCTGGCTACAAACCAAGTTTCCGTGAAAAAGACGGCCAACGTTTTGTGACGGATATGCAGAATTTTATCATTGACCTCGCC
NZ_CP035237.1:c1311322-1303456 456    TTTGAACGAGCTGGCTACAAACCAAGTTTCCGTGAAAAAGACGGCCAACGTTTTGTGACGGATATGCAGAATTTTATCATTGACCTCGCC
NZ_CP035246.1:c1302892-1295026 456    TTTGAACGAGCTGGCTACAAACCAAGTTTCCGTGAAAAAGACGGCCAACGTTTTGTGACGGATATGCAGAATTTTATCATTGACCTCGCC
NZ_CP035247.1:c1301115-1293263 442    TTTGAACGAGCTGGCTACAAACCAAGTTTCCGTGAAAAAGACGGCCAACGTTTTGTGACGGATATGCAGAATTTTATCATTGACCTCGCC
NZ_CP035238.1:738547-746413    456    TTTGAACGAGCTGGCTACAAACCAAGTTTCCGTGAAAAAGACGGCCAACGTTTTGTGACGGATATGCAGAATTTTATCATTGACCTCGCC
NZ_CP035255.1:776866-784718    442    TTTGAACGAGCTGGCTACAAACCAAGTTTCCGTGAAAAAGACGGCCAACGTTTTGTGACGGATATGCAGAATTTTATCATTGACCTCGCC
NZ_CP035256.1:807236-815102    456    TTTGAACGAGCTGGCTACAAACCAAGTTTCCGTGAAAAAGACGGCCAACGTTTTGTGACGGATATGCAGAATTTTATCATTGACCTCGCC
NZ_CP038252.1:1151812-1159673  386    TTTGAACGAGCTGGCTACAAACCAAGTTTCCGTGAAAAAGACGGCCAACGTTTTGTGACGGATATGCAGAATTTTATCATTGACCTCGCC
NZ_CP035254.1:793365-801213    442    TTTGAACGAGCTGGCTACAAACCAAGTTTCCGTGAAAAAGACGGCCAACGTTTTGTGACGGATATGCAGAATTTTATCATTGACCTCGCC
NZ_CP035251.1:748733-756585    442    TTTGAACGAGCTGGCTACAAACCAAGTTTCCGTGAAAAAGACGGCCAACGTTTTGTGACGGATATGCAGAATTTTATCATTGACCTCGCC
NZ_CP035252.1:745670-753536    456    TTTGAACGAGCTGGCTACAAACCAAGTTTCCGTGAAAAAGACGGCCAACGTTTTGTGACGGATATGCAGAATTTTATCATTGACCTCGCC
NZ_CP035257.1:776317-784169    442    TTTGAACGAGCTGGCTACAAACCAAGTTTCCGTGAAAAAGACGGCCAACGTTTTGTGACGGATATGCAGAATTTTATCATTGACCTCGCC
NZ_CP038253.1:758105-765957    442    TTTGAACGAGCTGGCTACAAACCAAGTTTCCGTGAAAAAGACGGCCAACGTTTTGTGACGGATATGCAGAATTTTATCATTGACCTCGCC
NZ_CP035263.1:743050-750901    442    TTTGAACGAGCTGGCTACAAACCAAGTTTCCGTGAAAAAGACGGCCAACGTTTTGTGACGGATATGCAGAATTTTATCATTGACCTCGCC
NZ_CP035262.1:741595-749460    456    TTTGAACGAGCTGGCTACAAACCAAGTTTCCGTGAAAAAGACGGCCAACGTTTTGTGACGGATATGCAGAATTTTATCATTGACCTCGCC
NZ_CP035236.1:764915-772766    442    TTTGAACGAGCTGGCTACAAACCAAGTTTCCGTGAAAAAGACGGCCAACGTTTTGTGACGGATATGCAGAATTTTATCATTGACCTCGCC
NZ_CP035253.1 767286-775134    442    TTTGAACGAGCTGGCTACAAACCAAGTTTCCGTGAAAAAGACGGCCAACGTTTTGTGACGGATATGCAGAATTTTATCATTGACCTCGCC
NZ_CP035249.1:763080-770921    442    TTTGAACGAGCTGGCTACAAACCAAGTTTCCGTGAAAAAGACGGTCAACGTTTTGTGACGGATATGCAGAATTTTATCATTGACCTCGCC
```

```
NC_003028.3:777000-787000       1261   TTGGATGTCATTGAAAATCCAATTGCTTTTGGACAAGAATTGGACCATGTCGTTGGTGTTGTGGAGCATGGTTTATTCAACCAAATGGTG
NZ_CP035248.1:765099-775111     1261   TTGGATGTCATTGAAGATCCAATTGCCTTTGGGCAAGAATTGGACCATGTCGTTGGTGTTGTAGAGCACGGTTTATTCAACCAAATGGTG
NZ_CP035235.1 c1326834-1317894 546    TTGGATGTCATTGAAAATCCAATTGCTTTTGGACAAGAATTGGACCATGTCGTTGGTGTTGTGGAGCATGGTTTATTCAACCAAATGGTG
NZ_CP035234.1:756284-765224     546    TTGGATGTCATTGAAAATCCAATTGCTTTTGGACAAGAATTGGACCATGTCGTTGGTGTTGTGGAGCATGGTTTATTCAACCAAATGGTG
NZ_CP038251.1:750745-759686     546    TTGGATGTCATTGAAAATCCAATTGCTTTTGGACAAGAATTGGACCATGTCGTTGGTGTTGTGGAGCATGGTTTATTCAACCAAATGGTG
NZ_CP035244.1:747023-755950     532    TTGGATGTCATTGAAAATCCAATTGCTTTTGGACAAGAATTGGACCATGTCGTTGGTGTTGTGGAGCATGGTTTATTCAACCAAATGGTG
NZ_CP035264.1:800836-809336     532    TTGGATGTCATTGAAAATCCAATTGCTTTTGGACAAGAATTGGACCATGTCGTTGGTGTTGTGGAGCATGGTTTATTCAACCAAATGGTG
NZ_CP035265.1:766128-774627     532    TTGGATGTCATTGAAAATCCAATTGCTTTTGGACAAGAATTGGACCATGTCGTTGGTGTTGTGGAGCATGGTTTATTCAACCAAATGGTG
NZ_CP035245.1 745261-753757     532    TTGGATGTCATTGAAAATCCAATTGCTTTTGGACAAGAATTGGACCATGTCGTTGGTGTTGTGGAGCATGGTTTATTCAACCAAATGGTG
NZ_CP035259.1:745700-753548     532    TTGGATGTCATTGAAAATCCAATTGCTTTTGGACAAGAATTGGACCATGTCGTTGGTGTTGTGGAGCATGGTTTATTCAACCAAATGGTG
NZ_CP035258.1:750874-758722     532    TTGGATGTCATTGAAAATCCAATTGCTTTTGGACAAGAATTGGACCATGTCGTTGGTGTTGTGGAGCATGGTTTATTCAACCAAATGGTG
NZ_CP035260.1:726156-733994     532    TTGGATGTCATTGAAAATCCAATTGCTTTTGGACAAGAATTGGACCATGTCGTTGGTGTTGTGGAGCATGGTTTATTCAACCAAATGGTG
NZ_CP035241.1:708296-716144     532    TTGGATGTCATTGAAAATCCAATTGCTTTTGGACAAGAATTGGACCATGTCGTTGGTGTTGTGGAGCATGGTTTATTCAACCAAATGGTG
NZ_CP035261.1:779086-786938     532    TTGGATGTCATTGAAAATCCAATTGCTTTTGGACAAGAATTGGACCATGTCGTTGGTGTTGTGGAGCATGGTTTATTCAACCAAATGGTG
NZ_CP035242.1:c1532236-1524385 532    TTGGATGTCATTGAAAATCCAATTGCTTTTGGACAAGAATTGGACCATGTCGTTGGTGTTGTGGAGCATGGTTTATTCAACCAAATGGTG
NZ_CP035243.1:777729-785580     532    TTGGATGTCATTGAAAATCCAATTGCTTTTGGACAAGAATTGGACCATGTCGTTGGTGTTGTGGAGCATGGTTTATTCAACCAAATGGTG
NZ_CP035237.1:c1311322-1303456 546    TTGGATGTCATTGAAAATCCAATTGCTTTTGGACAAGAATTGGATCATGTCGTTGGTGTTGTGGAGCATGGTTTATTCAACCAAATGGTG
NZ_CP035246.1:c1302892-1295026 546    TTGGATGTCATTGAAAATCCAATTGCTTTTGGACAAGAATTGGACCATGTCGTTGGTGTTGTGGAGCATGGTTTATTCAACCAAATGGTG
NZ_CP035247.1:c1301115-1293263 532    TTGGATGTCATTGAAAATCCAATTGCTTTTGGACAAGAATTGGACCATGTCGTTGGTGTTGTGGAGCATGGTTTATTCAACCAAATGGTG
NZ_CP035238.1:738547-746413     546    TTGGATGTCATTGAAAATCCAATTGCTTTTGGACAAGAATTGGACCATGTCGTTGGTGTTGTGGAGCATGGTTTATTCAACCAAATGGTG
NZ_CP035255.1:776866-784718     532    TTGGATGTCATTGAAAATCCAATTGCTTTTGGACAAGAATTGGACCATGTCGTTGGTGTTGTGGAGCATGGTTTATTCAACCAAATGGTG
NZ_CP035256.1:807236-815102     546    TTGGATGTCATTGAAAATCCAATTGCTTTTGGACAAGAATTGGACCATGTCGTTGGTGTTGTGGAGCATGGTTTATTCAACCAAATGGTG
NZ_CP038252.1:1151812-1159673  476    TTGGATGTCATTGAAAATCCAATTGCTTTTGGACAAGAATTGGACCATGTCGTTGGTGTTGTGGAGCATGGTTTATTCAACCAAATGGTG
NZ_CP035254.1:793365-801213     532    TTGGATGTCATTGAAAATCCAATTGCTTTTGGACAAGAATTGGACCATGTCGTTGGTGTTGTGGAGCATGGTTTATTCAACCAAATGGTG
NZ_CP035251.1:748733-756585     532    TTGGATGTCATTGAAAATCCAATTGCTTTTGGACAAGAATTGGACCATGTCGTTGGTGTTGTGGAGCATGGTTTATTCAACCAAATGGTG
NZ_CP035252.1:745670-753536     546    TTGGATGTCATTGAAAATCCAATTGCTTTTGGACAAGAATTGGACCATGTCGTTGGTGTTGTGGAGCATGGTTTATTCAACCAAATGGTG
NZ_CP035257.1:776317-784169     532    TTGGATGTCATTGAAAATCCAATTGCTTTTGGACAAGAATTGGACCATGTCGTTGGTGTTGTGGAGCATGGTTTATTCAACCAAATGGTG
NZ_CP038253.1:758105-765957     532    TTGGATGTCATTGAAAATCCAATTGCTTTTGGACAAGAATTGGACCATGTCGTTGGTGTTGTGGAGCATGGTTTATTCAACCAAATGGTG
NZ_CP035263.1:743050-750901     532    TTGGATGTCATTGAAAATCCAATTGCTTTTGGACAAGAATTGGACCATGTCGTTGGTGTTGTGGAGCATGGTTTATTCAACCAAATGGTG
NZ_CP035262.1:741595-749460     546    TTGGATGTCATTGAAAATCCAATTGCTTTTGGACAAGAATTGGACCATGTCGTTGGTGTTGTGGAGCATGGTTTATTCAACCAAATGGTG
NZ_CP035236.1:764915-772766     532    TTGGATGTCATTGAAAATCCAATTGCTTTTGGACAAGAATTGGACCATGTCGTTGGTGTTGTGGAGCATGGTTTATTCAACCAAATGGTG
NZ_CP035253.1 767286-775134     532    TTGGATGTCATTGAAAATCCAATTGCTTTTGGACAAGAATTGGACCATGTCGTTGGTGTTGTGGAGCATGGTTTATTCAACCAAATGGTG
NZ_CP035249.1:763080-770921     532    TTGGATGTCATTGAAAATCCAATTGCTTTTGGACAAGAATTGGACCATGTCGTTGGTGTTGTGGAGCATGGTTTATTCAACCAAATGGTG

NC_003028.3:777000-787000       1351   GATAAGGTAATCGTTGCTGGACGAGATGGAGTTCAGATTTCAACTTCAAAAAAAGGAAAATAG----------AAGGGGGCATAAGATGT
NZ_CP035248.1:765099-775111     1351   GATAAGGTCATTGTTGCTGGACGAGATGGGGTTCAGATTTTAACTTCAACAAAAGCAAACTAATCAACATAATAAGGAGATACACTATGT
NZ_CP035235.1 c1326834-1317894 636    GATAAGGTAATCGTTGCTGGACGAGATGGAGTTCAGATTTCAACTTCAAAAAAAGGAAAATAG----------AAGGGGGCATAAGATGT
NZ_CP035234.1:756284-765224     636    GATAAGGTAATCGTTGCTGGACGAGATGGAGTTCAGATTTCAACTTCAAAAAAAGGAAAATAG----------AAGGGGGCATAAGATGT
NZ_CP038251.1:750745-759686     636    GATAAGGTAATCGTTGCTGGACGAGATGGAGTTCAGATTTCAACTTCAAAAAAAGGAAAATAG----------AAGGGGGCATAAGATGT
NZ_CP035244.1:747023-755950     622    GATAAGGTAATCGTTGCTGGACGAGATGGAGTTCAGATTTCAACTTCAAAAAAAGGAAAATAG----------AAGGGGGCATAAGATGT
NZ_CP035264.1:800836-809336     622    GATAAGGTCATCGTTGCTGGACGAGATGGAGTTCAGATTTCAACTTCAAAAAAAGGAAAATAG----------AAGGGGGCATAAGATGT
NZ_CP035265.1:766128-774627     622    GATAAGGTCATCGTTGCTGGACGAGATGGAGTTCAGATTTCAACTTCAAAAAAAGGAAAATAG----------AAGGGGGCATAAGATGT
NZ_CP035245.1 745261-753757     622    GATAAGGTAATCGTTGCTGGACGAGATGGAGTTCAGATTTCAACTTCAAAAAAAGGAAAATAG----------AAGGGGGCATAAGATGT
NZ_CP035259.1:745700-753548     622    GATAAGGTCATCGTTGCTGGACGAGATGGAGTTCAGATTTCAACTTCAAAAAAAGGAAAATAG----------AAGGGGGCATAAGATGT
NZ_CP035258.1:750874-758722     622    GATAAGGTCATCGTTGCTGGACGAGATGGAGTTCAGATTTCAACTTCAAAAAAAGGAAAATAG----------AAGGGGGCATAAGATGT
NZ_CP035260.1:726156-733994     622    GATAAGGTAATCGTTGCTGGACGAGATGGAGTTCAGATTTCAACTTCAAAAAAAGGAAAATAG----------AAGGGGGCATAAGATGT
NZ_CP035241.1:708296-716144     622    GATAAGGTAATCGTTGCTGGACGAGATGGAGTTCAGATTTCAACTTCAAAAAAAGGAAAATAG----------AAGGGGGCATAAGATGT
NZ_CP035261.1:779086-786938     622    GATAAGGTAATCGTTGCTGGACGAGATGGAGTTCAGATTTCAACTTCAAAAAAAGGAAAATAG----------AAGGGGGCATAAGATGT
NZ_CP035242.1:c1532236-1524385 622    GATAAGGTAATCGTTGCTGGACGAGATGGAGTTCAGATTTCAACTTCAAAAAAAGGAAAATAG----------AAGGGGGCATAAGATGT
NZ_CP035243.1:777729-785580     622    GATAAGGTAATCGTTGCTGGACGAGATGGAGTTCAGATTTCAACTTCAAAAAAAGGAAAATAG----------AAGGGGGCATAAGATGT
NZ_CP035237.1:c1311322-1303456 636    GATAAGGTAATCGTTGCTGGACGGGATGGAGTTCAGATTTCAACTTCAAAAAAAGGAAAATAG----------AAGGGGGCATAAGATGT
NZ_CP035246.1:c1302892-1295026 636    GATAAGGTAATCGTTGCTGGACGAGATGGAGTTCAGATTTCAACTTCAAAAAAAGGAAAATAG----------AAGGGGGCATAAGATGT
NZ_CP035247.1:c1301115-1293263 622    GATAAGGTAATCGTTGCTGGACGAGATGGAGTTCAGATTTCAACTTCAAAAAAAGGAAAATAG----------AAGGGGGCATAAGATGT
NZ_CP035238.1:738547-746413     636    GATAAGGTAATCGTTGCTGGACGAGATGGAGTTCAGATTTCAACTTCAAAAAAAGGAAAATAG----------AAGGGGGCATAAGATGT
NZ_CP035255.1:776866-784718     622    GATAAGGTAATCGTTGCTGGACGAGATGGAGTTCAGATTTCAACTTCAAAAAAAGGAAAATAG----------AAGGGGGCATAAGATGT
NZ_CP035256.1:807236-815102     636    GATAAGGTAATCGTTGCTGGACGAGATGGAGTTCAGATTTCAACTTCAAAAAAAGGAAAATAG----------AAGGGGGCATAAGATGT
NZ_CP038252.1:1151812-1159673  566    GATAAGGTAATCGTTGCTGGACGAGATGGAGTTCAGATTTCAACTTCAAAAAAAGGAAAATAG----------AAGGGGGCATAAGATGT
NZ_CP035254.1:793365-801213     622    GATAAGGTAATCGTTGCTGGACGAGATGGAGTTCAGATTTCAACTTCAAAAAAAGGAAAATAG----------AAGGGGGCATAAGATGT
NZ_CP035251.1:748733-756585     622    GATAAGGTAATCGTTGCTGGACGAGATGGAGTTCAGATTTCAACTTCAAAAAAAGGAAAATAG----------AAGGGGGCATAAGATGT
NZ_CP035252.1:745670-753536     636    GATAAGGTAATCGTTGCTGGACGAGATGGAGTTCAGATTTCAACTTCAAAAAAAGGAAAATAG----------AAGGGGGCATAAGATGT
NZ_CP035257.1:776317-784169     622    GATAAGGTAATCGTTGCTGGACGAGATGGAGTTCAGATTTCAACTTCAAAAAAAGGAAAATAG----------AAGGGGGCATAAGATGT
NZ_CP038253.1:758105-765957     622    GATAAGGTAATCGTTGCTGGACGAGATGGAGTTCAGATTTCAACTTCAAAAAAAGGAAAATAG----------AAGGGGGCATAAGATGT
NZ_CP035263.1:743050-750901     622    GATAAGGTAATCGTTGCTGGACGAGATGGAGTTCAGATTTCAACTTCAAAAAAAGGAAAATAG----------AAGGGGGCATAAGATGT
NZ_CP035262.1:741595-749460     636    GATAAGGTAATCGTTGCTGGACGAGATGGAGTTCAGATTTCAACTTCAAAAAAAGGAAAATAG----------AAGGGGGCATAAGATGT
NZ_CP035236.1:764915-772766     622    GATAAGGTAATCGTTGCTGGACGAGATGGAGTTCAGATTTCAACTTCAAAAAAAGGAAAATAG----------AAGGGGGCATAAGATGT
NZ_CP035253.1 767286-775134     622    GATAAGGTAATCGTTGCTGGACGAGATGGAGTTCAGATTTCAACTTCAAAAAAAGGAAAATAG----------AAGGGGGCATAAGATGT
NZ_CP035249.1:763080-770921     622    GATAAGGTAATCGTTGCTGGACGAGATGGAGTTCAGATTTCAACTTCAAAAAAAGGAAAATAG----------AAGGGGGCATAAGATGT

NC_003028.3:777000-787000       1431   CTAAATTTAATCGTATTCATTTGGTGGTACTGGATTCTGTAGGAATCGGTGCAGCACCAGATGCTAATAACTTTGTCAATGCAGGGGTTC
NZ_CP035248.1:765099-775111     1441   CAAAATTTAATCGTATTCACTTGGTGGTACTAGATTCTGTAGGAATCGGTGCTGCACCAGATGCTAATAACTTTGTCAATGCAGGGGTTC
NZ_CP035235.1 c1326834-1317894 716    CTAAATTTAATCGTATTCATTTGGTGGTACTGGATTCTGTAGGAATCGGTGCAGCACCAGATGCTAATAACTTTGTCAATGCAGGGGTTC
NZ_CP035234.1:756284-765224     716    CTAAATTTAATCGTATTCATTTGGTGGTACTGGATTCTGTAGGAATCGGTGCAGCACCAGATGCTAATAACTTTGTCAATGCAGGGGTTC
NZ_CP038251.1:750745-759686     716    CTAAATTTAATCGTATTCATTTGGTGGTACTGGATTCTGTAGGAATCGGTGCAGCACCAGATGCTAATAACTTTGTCAATGCAGGGGTTC
NZ_CP035244.1:747023-755950     702    CTAAATTTAATCGTATTCATTTGGTGGTACTGGATTCTGTAGGAATCGGTGCAGCACCAGATGCTAATAACTTTGTCAATGCAGGGGTTC
NZ_CP035264.1:800836-809336     702    CTAAATTTAATCGTATTCATTTGGTGGTACTGGATTCTGTAGGAATCGGTGCAGCACCAGATGCTAATAACTTTGTCAATGCAGGGGTTC
NZ_CP035265.1:766128-774627     702    CTAAATTTAATCGTATTCATTTGGTGGTACTGGATTCTGTAGGAATCGGTGCAGCACCAGATGCTAATAACTTTGTCAATGCAGGGGTTC
NZ_CP035245.1 745261-753757     702    CTAAATTTAATCGTATTCATTTGGTGGTACTGGATTCTGTAGGAATCGGTGCAGCACCAGATGCTAATAACTTTGTCAATGCAGGGGTTC
NZ_CP035259.1:745700-753548     702    CTAAATTTAATCGTATTCATTTGGTGGTACTGGATTCTGTAGGAATCGGTGCAGCACCAGATGCTAATAACTTTGTCAATGCAGGGGTTC
NZ_CP035258.1:750874-758722     702    CTAAATTTAATCGTATTCATTTGGTGGTACTGGATTCTGTAGGAATCGGTGCAGCACCAGATGCTAATAACTTTGTCAATGCAGGGGTTC
NZ_CP035260.1:726156-733994     702    CTAAATTTAATCGTATTCATTTGGTGGTACTGGATTCTGTAGGAATCGGTGCAGCACCAGATGCTAATAACTTTGTCAATGCAGGGGTTC
NZ_CP035241.1:708296-716144     702    CTAAATTTAATCGTATTCATTTGGTGGTACTGGATTCTGTAGGAATCGGTGCAGCACCAGATGCTAATAACTTTGTCAATGCAGGGGTTC
NZ_CP035261.1:779086-786938     702    CTAAATTTAATCGTATTCATTTGGTGGTACTGGATTCTGTAGGAATCGGTGCAGCACCAGATGCTAATAACTTTGTCAATGCAGGGGTTC
NZ_CP035242.1:c1532236-1524385 702    CTAAATTTAATCGTATTCATTTGGTGGTACTGGATTCTGTAGGAATCGGTGCAGCACCAGATGCTAATAACTTTGTCAATGCAGGGGTTC
NZ_CP035243.1:777729-785580     702    CTAAATTTAATCGTATTCATTTGGTGGTACTGGATTCTGTAGGAATCGGTGCAGCACCAGATGCTAATAACTTTGTCAATGCAGGGGTTC
NZ_CP035237.1:c1311322-1303456 716    CTAAATTTAATCGTATTCATTTGGTGGTACTGGATTCTGTAGGAATCGGTGCAGCACCAGATGCTAATAACTTTGTCAATGCAGGGGTTC
NZ_CP035246.1:c1302892-1295026 716    CTAAATTTAATCGTATTCATTTGGTGGTACTGGATTCTGTAGGAATCGGTGCAGCACCAGATGCTAATAACTTTGTCAATGCAGGGGTTC
NZ_CP035247.1:c1301115-1293263 702    CTAAATTTAATCGTATTCATTTGGTGGTACTGGATTCTGTAGGAATCGGTGCAGCACCAGATGCTAATAACTTTGTCAATGCAGGGGTTC
NZ_CP035238.1:738547-746413     716    CTAAATTTAATCGTATTCATTTGGTGGTACTGGATTCTGTAGGAATCGGTGCAGCACCAGATGCTAATAACTTTGTCAATGCAGGGGTTC
NZ_CP035255.1:776866-784718     702    CTAAATTTAATCGTATTCATTTGGTGGTACTGGATTCTGTAGGAATCGGTGCAGCACCAGATGCTAATAACTTTGTCAATGCAGGGGTTC
NZ_CP035256.1:807236-815102     716    CTAAATTTAATCGTATTCATTTGGTGGTACTGGATTCTGTAGGAATCGGTGCAGCACCAGATGCTAATAACTTTGTCAATGCAGGGGTTC
NZ_CP038252.1:1151812-1159673  646    CTAAATTTAATCGTATTCATTTGGTGGTACTGGATTCTGTAGGAATCGGTGCAGCACCAGATGCTAATAACTTTGTCAATGCAGGGGTTC
NZ_CP035254.1:793365-801213     702    CTAAATTTAATCGTATTCATTTGGTGGTACTGGATTCTGTAGGAATCGGTGCAGCACCAGATGCTAATAACTTTGTCAATGCAGGGGTTC
NZ_CP035251.1:748733-756585     702    CTAAATTTAATCGTATTCATTTGGTGGTACTGGATTCTGTAGGAATCGGTGCAGCACCAGATGCTAATAACTTTGTCAATGCAGGGGTTC
NZ_CP035252.1:745670-753536     716    CTAAATTTAATCGTATTCATTTGGTGGTACTGGATTCTGTAGGAATCGGTGCAGCACCAGATGCTAATAACTTTGTCAATGCAGGGGTTC
```

```
NZ_CP035257.1:776317-784169    702    CTAAATTTAATCGTATTCATTTGGTGGTACTGGATTCTGTAGGAATCGGTGCAGCACCAGATGCTAATAACTTTGTCAATGCAGGGGTTC
NZ_CP038253.1:758105-765957    702    CTAAATTTAATCGTATTCATTTGGTGGTACTGGATTCTGTAGGAATCGGTGCAGCACCAGATGCTAATAACTTTGTCAATGCAGGGGTTC
NZ_CP035263.1:743050-750901    702    CTAAATTTAATCGTATTCATTTGGTGGTACTGGATTCTGTAGGAATCGGTGCAGCACCAGATGCTAATAACTTTGTCAATGCAGGGGTTC
NZ_CP035262.1:741595-749460    716    CTAAATTTAATCGTATTCATTTGGTGGTACTGGATTCTGTAGGAATCGGTGCAGCACCAGATGCTAATAACTTTGTCAATGCAGGGGTTC
NZ_CP035236.1:764915-772766    702    CTAAATTTAATCGTATTCATTTGGTGGTACTGGATTCTGTAGGAATCGGTGCAGCACCAGATGCTAATAACTTTGTCAATGCAGGGGTTC
NZ_CP035253.1 767286-775134    702    CTAAATTTAATCGTATTCATTTGGTGGTACTGGATTCTGTAGGAATCGGTGCAGCACCAGATGCTAATAACTTTGTCAATGCAGGGGTTC
NZ_CP035249.1:763080-770921    702    CTAAATTTAATCGTATTCATTTGGTGGTACTGGATTCTGTAGGAATCGGTGCAGCACCAGATGCTAATAACTTTGTCAATGCAGGGGTTC

NC_003028.3:777000-787000      1521   CAGATGGAGCTTCTGACACACTGGGACACATTTCAAAAACAGTTGGTTTGAATGTCCCAAACATGGCTAAAATAGGTCTTGGAAATATTC
NZ_CP035248.1:765099-775111    1531   CAGATGGAGCTTCTGACACACTGGGACACATTTCCAAATCAGTTGGTTTGAATGTGCCAAACATGGCTAAAATCGGTCTGGGAAATATTC
NZ_CP035235.1 c1326834-1317894 806    CAGATGGAGCTTCTGACACACTGGGACACATTTCAAAAACAGTTGGTTTGAATGTCCCAAACATGGCTAAAATAGGTCTTGGAAATATTC
NZ_CP035234.1:756284-765224    806    CAGATGGAGCTTCTGACACACTGGGACACATTTCAAAAACAGTTGGTTTGAATGTCCCAAACATGGCTAAAATAGGTCTTGGAAATATTC
NZ_CP038251.1:750745-759686    806    CAGATGGAGCTTCTGACACACTGGGACACATTTCAAAAACAGTTGGTTTGAATGTCCCAAACATGGCTAAAATAGGTCTTGGAAATATTC
NZ_CP035244.1:747023-755950    792    CAGATGGAGCTTCTGACACACTGGGACACATTTCAAAAACAGTTGGTTTGAATGTCCCAAACATGGCTAAAATAGGTCTTGGAAATATTC
NZ_CP035264.1:800836-809336    792    CAGATGGAGCTTCTGACACACTGGGACACATTTCAAAAACAGTTGGTTTGAATGTCCCAAACATGGCTAAAATAGGTCTTGGAAATATTC
NZ_CP035265.1:766128-774627    792    CAGATGGAGCTTCTGACACACTGGGACACATTTCAAAAACAGTTGGTTTGAATGTCCCAAACATGGCTAAAATAGGTCTTGGAAATATTC
NZ_CP035245.1 745261-753757    792    CAGATGGAGCTTCTGACACACTGGGACACATTTCAAAAACAGTTGGTTTGAATGTCCCAAACATGGCTAAAATAGGTCTTGGAAATATTC
NZ_CP035259.1:745700-753548    792    CAGATGGAGCTTCTGACACACTGGGACACATTTCAAAAACAGTTGGTTTGAATGTCCCAAACATGGCTAAAATAGGTCTTGGAAATATTC
NZ_CP035258.1:750874-758722    792    CAGATGGAGCTTCTGACACACTGGGACACATTTCAAAAACAGTTGGTTTGAATGTCCCAAACATGGCTAAAATAGGTCTTGGAAATATTC
NZ_CP035260.1:726156-733994    792    CAGATGGAGCTTCTGACACACTGGGACACATTTCAAAAACAGTTGGTTTGAATGTCCCAAACATGGCTAAAATAGGTCTTGGAAATATTC
NZ_CP035241.1:708296-716144    792    CAGATGGAGCTTCTGACACACTGGGACACATTTCAAAAACAGTTGGTTTGAATGTCCCAAACATGGCTAAAATAGGTCTTGGAAATATTC
NZ_CP035261.1:779086-786938    792    CAGATGGAGCTTCTGACACACTGGGACACATTTCAAAAACAGTTGGTTTGAATGTCCCAAACATGGCTAAAATAGGTCTTGGAAATATTC
NZ_CP035242.1:c1532236-1524385 792    CAGATGGAGCTTCTGACACACTGGGACACATTTCAAAAACAGTTGGTTTGAATGTCCCAAACATGGCTAAAATAGGTCTTGGAAATATTC
NZ_CP035243.1:777729-785580    792    CAGATGGAGCTTCTGACACACTGGGACACATTTCAAAAACAGTTGGTTTGAATGTCCCAAACATGGCTAAAATAGGTCTTGGAAATATTC
NZ_CP035237.1:c1311322-1303456 806    CAGATGGAGCTTCTGACACACTGGGACATATTTCAAAAACAGTTGGTTTGAATGTCCCAAACATGGCTAAAATAGGTCTTGGAAATATTC
NZ_CP035246.1:c1302892-1295026 806    CAGATGGAGCTTCTGACACACTGGGACACATTTCAAAAACAGTTGGTTTGAATGTCCCAAACATGGCTAAAATAGGTCTTGGAAATATTC
NZ_CP035247.1:c1301115-1293263 792    CAGATGGAGCTTCTGACACACTGGGACACATTTCAAAAACAGTTGGTTTGAATGTCCCAAACATGGCTAAAATAGGTCTTGGAAATATTC
NZ_CP035238.1:738547-746413    806    CAGATGGAGCTTCTGACACACTGGGACACATTTCAAAAACAGTTGGTTTGAATGTCCCAAACATGGCTAAAATAGGTCTTGGAAATATTC
NZ_CP035255.1:776866-784718    792    CAGATGGAGCTTCTGACACACTGGGACACATTTCAAAAACAGTTGGTTTGAATGTCCCAAACATGGCTAAAATAGGTCTTGGAAATATTC
NZ_CP035256.1:807236-815102    806    CAGATGGAGCTTCTGACACACTGGGACACATTTCAAAAACAGTTGGTTTGAATGTCCCAAACATGGCTAAAATAGGTCTTGGAAATATTC
NZ_CP038252.1:1151812-1159673  736    CAGATGGAGCTTCTGACACACTGGGACACATTTCAAAAACAGTTGGTTTGAATGTCCCAAACATGGCTAAAATAGGTCTTGGAAATATTC
NZ_CP035254.1:793365-801213    792    CAGATGGAGCTTCTGACACACTGGGACACATTTCAAAAACAGTTGGTTTGAATGTCCCAAACATGGCTAAAATAGGTCTTGGAAATATTC
NZ_CP035251.1:748733-756585    792    CAGATGGAGCTTCTGACACACTGGGACACATTTCAAAAACAGTTGGTTTGAATGTCCCAAACATGGCTAAAATAGGTCTTGGAAATATTC
NZ_CP035252.1:745670-753536    806    CAGATGGAGCTTCTGACACACTGGGACACATTTCAAAAACAGTTGGTTTGAATGTCCCAAACATGGCTAAAATAGGTCTTGGAAATATTC
NZ_CP035257.1:776317-784169    792    CAGATGGAGCTTCTGACACACTGGGACACATTTCAAAAACAGTTGGTTTGAATGTCCCAAACATGGCTAAAATAGGTCTTGGAAATATTC
NZ_CP038253.1:758105-765957    792    CAGATGGAGCTTCTGACACACTGGGACACATTTCAAAAACAGTTGGTTTGAATGTCCCAAACATGGCTAAAATAGGTCTTGGAAATATTC
NZ_CP035263.1:743050-750901    792    CAGATGGAGCTTCTGACACACTGGGACACATTTCAAAAACAGTTGGTTTGAATGTCCCAAACATGGCTAAAATAGGTCTTGGAAATATTC
NZ_CP035262.1:741595-749460    806    CAGATGGAGCTTCTGACACACTGGGACACATTTCAAAAACAGTTGGTTTGAATGTCCCAAACATGGCTAAAATAGGTCTTGGAAATATTC
NZ_CP035236.1:764915-772766    792    CAGATGGAGCTTCTGACACACTGGGACACATTTCAAAAACAGTTGGTTTGAATGTCCCAAACATGGCTAAAATAGGTCTTGGAAATATTC
NZ_CP035253.1 767286-775134    792    CAGATGGAGCTTCTGACACACTGGGACACATTTCAAAAACAGTTGGTTTGAATGTCCCAAACATGGCTAAAATAGGTCTTGGAAATATTC
NZ_CP035249.1:763080-770921    792    CAGATGGAGCTTCTGACACACTGGGACACATTTCAAAAACAGTTGGTTTGAATGTCCCAAACATGGCTAAAATAGGTCTTGGAAATATTC

NC_003028.3:777000-787000      1611   CTCGTGAAACTCCTCTTAAGACTGTAGCAGCTGAAAGCAATCCAACTGGATATGCAACAAAATTAGAGGAAGTATCTCTTGGTAAGGATA
NZ_CP035248.1:765099-775111    1621   CTCGCGAAACGCCTTTGAAGACTGTACCGGCGGAAAGCAATCCAACAGGATATGCAACAAAATTGGAAGAAGTATCTCTTGGTAAGGATA
NZ_CP035235.1 c1326834-1317894 896    CTCGTGAAACTCCTCTTAAGACTGTAGCAGCTGAAAGCAATCCAACTGGATATGCAACAAAATTAGAGGAAGTATCTCTTGGTAAGGATA
NZ_CP035234.1:756284-765224    896    CTCGTGAAACTCCTCTTAAGACTGTAGCAGCTGAAAGCAATCCAACTGGATATGCAACAAAATTAGAGGAAGTATCTCTTGGTAAGGATA
NZ_CP038251.1:750745-759686    896    CTCGTGAAACTCCTCTTAAGACTGTAGCAGCTGAAAGCAATCCAACTGGATATGCAACAAAATTAGAGGAAGTATCTCTTGGTAAGGATA
NZ_CP035244.1:747023-755950    882    CTCGTGAAACTCCTCTTAAGACTGTAGCAGCTGAAAGCAATCCAACTGGATATGCAACAAAATTAGAGGAAGTATCTCTTGGTAAGGATA
NZ_CP035264.1:800836-809336    882    CTCGTGAAACTCCTCTTAAGACTGTAGCAGCTGAAAGCAATCCAACTGGATATGCAACAAAATTAGAGGAAGTATCTCTTGGTAAGGATA
NZ_CP035265.1:766128-774627    882    CTCGTGAAACTCCTCTTAAGACTGTAGCAGCTGAAAGCAATCCAACTGGATATGCAACAAAATTAGAGGAAGTATCTCTTGGTAAGGATA
NZ_CP035245.1 745261-753757    882    CTCGTGAAACTCCTCTTAAGACTGTAGCAGCTGAAAGCAATCCAACTGGATATGCAACAAAATTAGAGGAAGTATCTCTTGGTAAGGATA
NZ_CP035259.1:745700-753548    882    CTCGTGAAACTCCTCTTAAGACTGTAGCAGCTGAAAGCAATCCAACTGGATATGCAACAAAATTAGAGGAAGTATCTCTTGGTAAGGATA
NZ_CP035258.1:750874-758722    882    CTCGTGAAACTCCTCTTAAGACTGTAGCAGCTGAAAGCAATCCAACTGGATATGCAACAAAATTAGAGGAAGTATCTCTTGGTAAGGATA
NZ_CP035260.1:726156-733994    882    CTCGTGAAACTCCTCTTAAGACTGTAGCAGCTGAAAGCAATCCAACTGGATATGCAACAAAATTAGAGGAAGTATCTCTTGGTAAGGATA
NZ_CP035241.1:708296-716144    882    CTCGTGAAACTCCTCTTAAGACTGTAGCAGCTGAAATCAATCCAACTGGATATGCAACAAAATTAGAGGAAGTATCTCTTGGTAAGGATA
NZ_CP035261.1:779086-786938    882    CTCGTGAAACTCCTCTTAAGACTGTAGCAGCTGAAAGCAATCCAACTGGATATGCAACAAAATTAGAGGAAGTATCTCTTGGTAAGGATA
NZ_CP035242.1:c1532236-1524385 882    CTCGTGAAACTCCTCTTAAGACCGTAGCAGCTGAAATCAATCCAACTGGATATGCAACAAAATTAGAGGAAGTATCTCTTGGTAAGGATA
NZ_CP035243.1:777729-785580    882    CTCGTGAAACTCCTCTTAAGACCGTAGCAGCTGAAATCAATCCAACTGGATATGCAACAAAATTAGAGGAAGTATCTCTTGGTAAGGATA
NZ_CP035237.1:c1311322-1303456 896    CTCGTGAAACTCCTCTTAAGACTGTAGCAGCTGAAAGCAATCCAACTGGATATGCAACAAAATTAGAGGAAGTATCTCTTGGTAAGGATA
NZ_CP035246.1:c1302892-1295026 896    CTCGTGAAACTCCTCTTAAGACTGTAGCAGCTGAAAGCAATCCAACTGGATATGCAACAAAATTAGAGGAAGTATCTCTTGGTAAGGATA
NZ_CP035247.1:c1301115-1293263 882    CTCGTGAAACTCCTCTTAAGACTGTAGCAGCTGAAAGCAATCCAACTGGATATGCAACAAAATTAGAGGAAGTATCTCTTGGTAAGGATA
NZ_CP035238.1:738547-746413    896    CTCGTGAAACTCCTCTTAAGACTGTAGCAGCTGAAAGCAATCCAACTGGATATGCAACAAAATTAGAGGAAGTATCTCTTGGTAAGGATA
NZ_CP035255.1:776866-784718    882    CTCGTGAAACTCCTCTTAAGACTGTAGCAGCTGAAAGCAATCCAACTGGATATGCAACAAAATTAGAGGAAGTATCTCTTGGTAAGGATA
NZ_CP035256.1:807236-815102    896    CTCGTGAAACTCCTCTTAAGACTGTAGCAGCTGAAAGCAATCCAACTGGATATGCAACAAAATTAGAGGAAGTATCTCTTGGTAAGGATA
NZ_CP038252.1:1151812-1159673  826    CTCGTGAAACTCCTCTTAAGACTGTAGCAGCTGAAAGCAATCCAACTGGATATGCAACAAAATTAGAGGAAGTATCTCTTGGTAAGGATA
NZ_CP035254.1:793365-801213    882    CTCGTGAAACTCCTCTTAAGACTGTAGCAGCTGAAATCAATCCAACTGGATATGCAACAAAATTAGAGGAAGTATCTCTTGGTAAGGATA
NZ_CP035251.1:748733-756585    882    CTCGTGAAACTCCTCTTAAGACTGTAGCAGCTGAAATCAATCCAACTGGATATGCAACAAAATTAGAGGAAGTATCTCTTGGTAAGGATA
NZ_CP035252.1:745670-753536    896    CTCGTGAAACTCCTCTTAAGACTGTAGCAGCTGAAATCAATCCAACTGGATATGCAACAAAATTAGAGGAAGTATCTCTTGGTAAGGATA
NZ_CP035257.1:776317-784169    882    CTCGTGAAACTCCTCTTAAGACTGTAGCAGCTGAAATCAATCCAACTGGATATGCAACAAAATTAGAGGAAGTATCTCTTGGTAAGGATA
NZ_CP038253.1:758105-765957    882    CTCGTGAAACTCCTCTTAAGACTGTAGCAGCTGAAATCAATCCAACTGGATATGCAACAAAATTAGAGGAAGTATCTCTTGGTAAGGATA
NZ_CP035263.1:743050-750901    882    CTCGTGAAACTCCTCTTAAGACTGTAGCAGCTGAAATCAATCCAACTGGATATGCAACAAAATTAGAGGAAGTATCTCTTGGTAAGGATA
NZ_CP035262.1:741595-749460    896    CTCGTGAAACTCCTCTTAAGACTGTAGCAGCTGAAATCAATCCAACTGGATATGCAACAAAATTAGAGGAAGTATCTCTTGGTAAGGATA
NZ_CP035236.1:764915-772766    882    CTCGTGAAACTCCTCTTAAGACTGTAGCAGCTGAAATCAATCCAACTGGATATGCAACAAAATTAGAGGAAGTATCTCTTGGTAAGGATA
NZ_CP035253.1 767286-775134    882    CTCGTGAAACTCCTCTTAAGACTGTAGCAGCTGAAATCAATCCAACTGGATATGCAACAAAATTAGAGGAAGTATCTCTTGGTAAGGATA
NZ_CP035249.1:763080-770921    882    CTCGTGAAACTCCTCTTAAGACTGTAGCAGCTGAAAGCAATCCAACTGGATATGCAACAAAATTAGAGGAAGTATCTCTTGGTAAGGATA

NC_003028.3:777000-787000      1701   CTATGACTGGACACTGGGAAATCATGGGACTCAACATTACTGAGCCTTTCGATACTTTCTGGAACGGATTCCCAGAAGAAATCCTGACAA
NZ_CP035248.1:765099-775111    1711   CTATGACTGGACACTGGGAAATCATGGGACTCAACATTACTGAGCCTTTCGATACTTTCTGGAACGGATTCCCAGAAGAAATCTTGACAA
NZ_CP035235.1 c1326834-1317894 986    CTATGACTGGACACTGGGAAATCATGGGACTCAACATTACTGAGCCTTTCGATACTTTCTGGAACGGATTCCCAGAAGAAATCCTGACAA
NZ_CP035234.1:756284-765224    986    CTATGACTGGACACTGGGAAATCATGGGACTCAACATTACTGAGCCTTTCGATACTTTCTGGAACGGATTCCCAGAAGAAATCCTGACAA
NZ_CP038251.1:750745-759686    986    CTATGACTGGACACTGGGAAATCATGGGACTCAACATTACTGAGCCTTTCGATACTTTCTGGAACGGATTCCCAGAAGAAATCCTGACAA
NZ_CP035244.1:747023-755950    972    CTATGACTGGACACTGGGAAATCATGGGACTCAACATTACTGAGCCTTTCGATACTTTCTGGAACGGATTCCCAGAAGAAATCCTGACAA
NZ_CP035264.1:800836-809336    972    CTATGACTGGACACTGGGAAATCATGGGACTCAACATTACTGAGCCTTTCGATACTTTCTGGAACGGATTCCCAGAAGAAATCCTGACAA
NZ_CP035265.1:766128-774627    972    CTATGACTGGACACTGGGAAATCATGGGACTCAACATTACTGAGCCTTTCGATACTTTCTGGAACGGATTCCCAGAAGAAATCCTGACAA
NZ_CP035245.1 745261-753757    972    CTATGACTGGACACTGGGAAATCATGGGACTCAACATTACTGAGCCTTTCGATACTTTCTGGAACGGATTCCCAGAAGAAATCCTGACAA
NZ_CP035259.1:745700-753548    972    CTATGACTGGACACTGGGAAATCATGGGACTCAACATTACTGAGCCTTTCGATACTTTCTGGAACGGATTCCCAGAAGAAATCCTGACAA
NZ_CP035258.1:750874-758722    972    CTATGACTGGACACTGGGAAATCATGGGACTCAACATTACTGAGCCTTTCGATACTTTCTGGAACGGATTCCCAGAAGAAATCCTGACAA
NZ_CP035260.1:726156-733994    972    CTATGACTGGACACTGGGAAATCATGGGACTCAACATTACTGAGCCTTTCGATACTTTCTGGAACGGATTCCCAGAAGAAATCCTGACAA
NZ_CP035241.1:708296-716144    972    CTATGACTGGACACTGGGAAATCATGGGACTCAACATTACTGAGCCTTTCGATACTTTCTGGAACGGATTCCCAGAAGAAATCCTGACAA
NZ_CP035261.1:779086-786938    972    CTATGACTGGACACTGGGAAATCATGGGACTCAACATTACTGAGCCTTTCGATACTTTCTGGAACGGATTCCCAGAAGAAATCCTGACAA
NZ_CP035242.1:c1532236-1524385 972    CTATGACTGGACACTGGGAAATCATGGGACTCAACATTACTGAGCCTTTCGATACTTTCTGGAACGGATTCCCAGAAGAAATCCTGACAA
NZ_CP035243.1:777729-785580    972    CTATGACTGGACACTGGGAAATCATGGGACTCAACATTACTGAGCCTTTCGATACTTTCTGGAACGGATTCCCAGAAGAAATCCTGACAA
NZ_CP035237.1:c1311322-1303456 986    CTATGACTGGACACTGGGAAATCATGGGACTCAACATTACTGAGCCTTTCGATACTTTCTGGAACGGATTCCCAGAAGAAATCCTGACAA
NZ_CP035246.1:c1302892-1295026 986    CTATGACTGGACACTGGGAAATCATGGGACTCAACATTACTGAGCCTTTCGATACTTTCTGGAACGGATTCCCAGAAGAAATCCTGACAA
NZ_CP035247.1:c1301115-1293263 972    CTATGACTGGACACTGGGAAATCATGGGACTCAACATTACTGAGCCTTTCGATACTTTCTGGAACGGATTCCCAGAAGAAATCCTGACAA
```

```
NZ_CP035238.1:738547-746413    986    CTATGACTGGACACTGGGAAATCATGGGACTCAACATTACTGAGCCTTTCGATACTTTCTGGAACGGATTCCCAGAAGAAATCCTGACAA
NZ_CP035255.1:776866-784718    972    CTATGACTGGACACTGGGAAATCATGGGACTCAACATTACTGAGCCTTTCGATACTTTCTGGAACGGATTCCCAGAAGAAATCCTGACAA
NZ_CP035256.1:807236-815102    986    CTATGACTGGACACTGGGAAATCATGGGACTCAACATTACTGAGCCTTTCGATACTTTCTGGAACGGATTCCCAGAAGAAATCCTGACAA
NZ_CP038252.1:1151812-1159673  916    CTATGACTGGACACTGGGAAATCATGGGACTCAACATTACTGAGCCTTTCGATACTTTCTGGAACGGATTCCCAGAAGAAATCCTGACAA
NZ_CP035254.1:793365-801213    972    CTATGACTGGACACTGGGAAATCATGGGACTCAACATTACTGAGCCTTTCGATACTTTCTGGAACGGATTCCCAGAAGAAATCCTGACAA
NZ_CP035251.1:748733-756585    972    CTATGACTGGACACTGGGAAATCATGGGACTCAACATTACTGAGCCTTTCGATACTTTCTGGAACGGATTCCCAGAAGAAATCCTGACAA
NZ_CP035252.1:745670-753536    986    CTATGACTGGACACTGGGAAATCATGGGACTCAACATTACTGAGCCTTTCGATACTTTCTGGAACGGATTCCCAGAAGAAATCCTGACAA
NZ_CP035257.1:776317-784169    972    CTATGACTGGACACTGGGAAATCATGGGACTCAACATTACTGAGCCTTTCGATACTTTCTGGAACGGATTCCCAGAAGAAATCCTGACAA
NZ_CP038253.1:758105-765957    972    CTATGACTGGACACTGGGAAATCATGGGACTCAACATTACTGAGCCTTTCGATACTTTCTGGAACGGATTCCCAGAAGAAATCCTGACAA
NZ_CP035263.1:743050-750901    972    CTATGACTGGACACTGGGAAATCATGGGACTCAACATTACTGAGCCTTTCGATACTTTCTGGAACGGATTCCCAGAAGAAATCCTGACAA
NZ_CP035262.1:741595-749460    986    CTATGACTGGACACTGGGAAATCATGGGACTCAACATTACTGAGCCTTTCGATACTTTCTGGAACGGATTCCCAGAAGAAATCCTGACAA
NZ_CP035236.1:764915-772766    972    CTATGACTGGACACTGGGAAATCATGGGACTCAACATTACTGAGCCTTTCGATACTTTCTGGAACGGATTCCCAGAAGAAATCCTGACAA
NZ_CP035253.1 767286-775134    972    CTATGACTGGACACTGGGAAATCATGGGACTCAACATTACTGAGCCTTTCGATACTTTCTGGAACGGATTCCCAGAAGAAATCCTGACAA
NZ_CP035249.1:763080-770921    972    CTATGACTGGACACTGGGAAATCATGGGACTCAACATTACTGAGCCTTTCGATACTTTCTGGAACGGATTCCCAGAAGAAATCCTGACAA

NC_003028.3:777000-787000      1791   AAATCGAAGAATTCTCAGGACGCAAGGTTATTCGTGAAGCCAACAAACCTTATTCAGGAACGGCTGTTATCTATGATTTTGGACCACGTC
NZ_CP035248.1:765099-775111    1801   AAATTGAAGAATTCTCAGGACGCAAGATCATTCGTGAAGCTAACAAACCTTACTCAGGTACAGCTGTTATCGATGATTTTGGACCACGTC
NZ_CP035235.1 c1326834-1317894 1076   AAATCGAAGAATTCTCAGGACGCAAGGTTATTCGTGAAGCCAACAAACCTTATTCAGGAACGGCTGTTATCGATGATTTTGGACCACGTC
NZ_CP035234.1:756284-765224    1076   AAATCGAAGAATTCTCAGGACGCAAGGTTATTCGTGAAGCCAACAAACCTTATTCAGGAACGGCTGTTATCGATGATTTTGGACCACGTC
NZ_CP038251.1:750745-759686    1076   AAATCGAAGAATTCTCAGGACGCAAGGTTATTCGTGAAGCCAACAAACCTTATTCAGGAACGGCTGTTATCGATGATTTTGGACCACGTC
NZ_CP035244.1:747023-755950    1062   AAATCGAAGAATTCTCAGGACGCAAGGTTATTCGTGAAGCCAACAAACCTTATTCAGGAACGGCTGTTATCGATGATTTTGGACCACGTC
NZ_CP035264.1:800836-809336    1062   AAATCGAAGAATTCTCAGGACGCAAGGTTATTCGTGAAGCCAACAAACCTTATTCAGGAACGGCTGTTATCGATGATTTTGGACCACGTC
NZ_CP035265.1:766128-774627    1062   AAATCGAAGAATTCTCAGGACGCAAGGTTATTCGTGAAGCCAACAAACCTTATTCAGGAACGGCTGTTATCGATGATTTTGGACCACGTC
NZ_CP035245.1 745261-753757    1062   AAATCGAAGAATTCTCAGGACGCAAGGTTATTCGTGAAGCCAACAAACCTTATTCAGGAACGGCTGTTATCGATGATTTTGGACCACGTC
NZ_CP035259.1:745700-753548    1062   AAATCGAAGAATTCTCAGGACGCAAGGTTATTCGTGAAGCCAACAAACCTTATTCAGGAACGGCTGTTATCGATGATTTTGGACCACGTC
NZ_CP035258.1:750874-758722    1062   AAATCGAAGAATTCTCAGGACGCAAGGTTATTCGTGAAGCCAACAAACCTTATTCAGGAACGGCTGTTATCGATGATTTTGGACCACGTC
NZ_CP035260.1:726156-733994    1062   AAATTGAAGAATTCTCAGGACGCAAGGTTATTCGTGAAGCCAACAAACCTTATTCAGGAACGGCTGTTATCGATGATTTTGGACCACGTC
NZ_CP035241.1:708296-716144    1062   AAATTGAAGAATTCTCAGGACGCAAGGTTATTCGTGAAGCCAACAAACCTTATTCAGGAACGGCTGTTATCGATGATTTTGGACCACGTC
NZ_CP035261.1:779086-786938    1062   AAATCGAAGAATTCTCAGGACGCAAGGTTATTCGTGAAGCCAACAAACCTTATTCAGGAACGGCTGTTATCGATGATTTTGGACCACGTC
NZ_CP035242.1:c1532236-1524385 1062   AAATCGAAGAATTCTCAGGACGCAAGGTTATTCGTGAAGCCAACAAACCTTATTCAGGAACGGCTGTTATCGATGATTTTGGACCACGTC
NZ_CP035243.1:777729-785580    1062   AAATCGAAGAATTCTCAGGACGCAAGGTTATTCGTGAAGCCAACAAACCTTATTCAGGAACGGCTGTTATCGATGATTTTGGACCACGTC
NZ_CP035237.1:c1311322-1303456 1076   AAATCGAAGAATTCTCAGGACGCAAGGTTATTCGTGAAGCCAACAAACCTTATTCAGGAACGGCTGTTATCGATGATTTTGGACCACGTC
NZ_CP035246.1:c1302892-1295026 1076   AAATCGAAGAATTCTCAGGACGCAAGGTTATTCGTGAAGCCAACAAACCTTATTCAGGAACGGCTGTTATCGATGATTTTGGACCACGTC
NZ_CP035247.1:c1301115-1293263 1062   AAATCGAAGAATTCTCAGGACGCAAGGTTATTCGTGAAGCCAACAAACCTTATTCAGGAACGGCTGTTATCGATGATTTTGGACCACGTC
NZ_CP035238.1:738547-746413    1076   AAATCGAAGAATTCTCAGGACGCAAGGTTATTCGTGAAACCAACAAACCTTATTCAGGAACGGCTGTTATCTATGATTTTGGACCACGTC
NZ_CP035255.1:776866-784718    1062   AAATCGAAGAATTCTCAGGACGCAAGGTTATTCGTGAAGCCAACAAACCTTATTCAGGAACGGCTGTTATCGATGATTTTGGACCACGTC
NZ_CP035256.1:807236-815102    1076   AAATCGAAGAATTCTCAGGACGCAAGGTTATTCGTGAAGCCAACAAACCTTATTCAGGAACGGCTGTTATCGATGATTTTGGACCACGTC
NZ_CP038252.1:1151812-1159673  1006   AAATCGAAGAATTCTCAGGACGCAAGGTTATTCGTGAAGCCAACAAACCTTATTCAGGAACGGCTGTTATCGATGATTTTGGACCACGTC
NZ_CP035254.1:793365-801213    1062   AAATCGAAGAATTCTCAGGACGCAAGGTTATTCGTGAAGCCAACAAACCTTATTCAGGAACGGCTGTTATCGATGATTTTGGACCACGTC
NZ_CP035251.1:748733-756585    1062   AAATCGAAGAATTCTCAGGACGCAAGGTTATTCGTGAAGCCAACAAACCTTATTCAGGAACGGCTGTTATCGATGATTTTGGACCACGTC
NZ_CP035252.1:745670-753536    1076   AAATCGAAGAATTCTCAGGACGCAAGGTTATTCGTGAAGCCAACAAACCTTATTCAGGAACGGCTGTTATCGATGATTTTGGACCACGTC
NZ_CP035257.1:776317-784169    1062   AAATCGAAGAATTCTCAGGACGCAAGGTTATTCGTGAAGCCAACAAACCTTATTCAGGAACGGCTGTTATCGATGATTTTGGACCACGTC
NZ_CP038253.1:758105-765957    1062   AAATCGAAGAATTCTCAGGACGCAAGGTTATTCGTGAAGCCAACAAACCTTATTCAGGAACGGCTGTTATCGATGATTTTGGACCACGTC
NZ_CP035263.1:743050-750901    1062   AAATCGAAGAATTCTCAGGACGCAAGGTTATTCGTGAAGCCAACAAACCTTATTCAGGAACGGCTGTTATCGATGATTTTGGACCACGTC
NZ_CP035262.1:741595-749460    1076   AAATCGAAGAATTCTCAGGACGCAAGGTTATTCGTGAAGCCAACAAACCTTATTCAGGAACGGCTGTTATCGATGATTTTGGACCACGTC
NZ_CP035236.1:764915-772766    1062   AAATCGAAGAATTCTCAGGACGCAAGGTTATTCGTGAAGCCAACAAACCTTATTCAGGAACGGCTGTTATCGATGATTTTGGACCACGTC
NZ_CP035253.1 767286-775134    1062   AAATCGAAGAATTCTCAGGACGCAAGGTTATTCGTGAAGCCAACAAACCTTATTCAGGAACGGCTGTTATCGATGATTTTGGACCACGTC
NZ_CP035249.1:763080-770921    1062   AAATCGAAGAATTCTCAGGACGCAAGGTTATTCGTGAAGCCAACAAACCTTATTCAGGAACGGCTGTTATCGATGATTTTGGACCACGTC

NC_003028.3:777000-787000      1881   AGATGGAAACTGGAGAGTTGATTATCTATACTTCAGCTGACCCTGTTTTGCAGATTGCTGCCCACGAAGACATTATTCCTTTGGATGAAT
NZ_CP035248.1:765099-775111    1891   AAATGGAAACTGGGGAGTTGATTATCTATACTTCAGCTGATCCTGTTTTGCAAATTGCTGCCCACGAAGACATTATTCCTTTGGACGAAT
NZ_CP035235.1 c1326834-1317894 1166   AGATGGAAACTGGAGAGTTGATTATCTATACTTCAGCTGACCCTGTTTTGCAGATTGCTGCCCACGAAGACATTATTCCTTTGGATGAAT
NZ_CP035234.1:756284-765224    1166   AGATGGAAACTGGAGAGTTGATTATCTATACTTCAGCTGACCCTGTTTTGCAGATTGCTGCCCACGAAGACATTATTCCTTTGGATGAAT
NZ_CP038251.1:750745-759686    1166   AGATGGAAACTGGAGAGTTGATTATCTATACTTCAGCTGACCCTGTTTTGCAGATTGCTGCCCACGAAGACATTATTCCTTTGGATGAAT
NZ_CP035244.1:747023-755950    1152   AGATGGAAACTGGAGAGTTGATTATCTATACTTCAGCTGACCCTGTTTTGCAGATTGCTGCCCACGAAGACATTATTCCTTTGGATGAAT
NZ_CP035264.1:800836-809336    1152   AGATGGAAACTGGAGAGTTGATTATCTATACTTCAGCTGACCCTGTTTTGCAGATTGCTGCCCACGAAGACATTATTCCTTTGGATGAAT
NZ_CP035265.1:766128-774627    1152   AGATGGAAACTGGAGAGTTGATTATCTATACTTCAGCTGACCCTGTTTTGCAGATTGCTGCCCACGAAGACATTATTCCTTTGGATGAAT
NZ_CP035245.1 745261-753757    1152   AGATGGAAACTGGAGAGTTGATTATCTATACTTCAGCTGACCCTGTTTTGCAGATTGCTGCCCACGAAGACATTATTCCTTTGGATGAAT
NZ_CP035259.1:745700-753548    1152   AGATGGAAACTGGAGAGTTGATTATCTATACTTCAGCTGACCCTGTTTTGCAGATTGCTGCCCACGAAGACATTATTCCTTTGGATGAAT
NZ_CP035258.1:750874-758722    1152   AGATGGAAACTGGAGAGTTGATTATCTATACTTCAGCTGACCCTGTTTTGCAGATTGCTGCCCACGAAGACATTATTCCTTTGGATGAAT
NZ_CP035260.1:726156-733994    1152   AGATGGAAACTGGAGAGTTGATTATCTATACTTCAGCTGACCCTGTTTTGCAGATTGCTGCCCACGAAGATATTATTCCTTTGGATGAAT
NZ_CP035241.1:708296-716144    1152   AGATGGAAACTGGAGAGTTGATTATCTATACTTCAGCTGACCCTGTTTTGCAGATTGCTGCCCACGAAGACATTATTCCTTTGGATGAAT
NZ_CP035261.1:779086-786938    1152   AGATGGAAACTGGAGAGTTGATTATCTATACTTCAGCTGACCCTGTTTTGCAGATTGCTGCCCACGAAGACATTATTCCTTTGGATGAAT
NZ_CP035242.1:c1532236-1524385 1152   AGATGGAAACTGGAGAGTTGATTATCTATACTTCAGCTGACCCTGTTTTGCAGATTGCTGCCCACGAAGACATTATTCCTTTGGATGAAT
NZ_CP035243.1:777729-785580    1152   AGATGGAAACTGGAGAGTTGATTATCTATACTTCAGCTGACCCTGTTTTGCAGATTGCTGCCCACGAAGACATTATTCCTTTGGATGAAT
NZ_CP035237.1:c1311322-1303456 1166   AGATGGAAACTGGAGAGTTGATTATCTATACTTCAGCTGACCCTGTTTTGCAGATTGCTGCCCACGAAGACATTATTCCTTTGGATGAAT
NZ_CP035246.1:c1302892-1295026 1166   AGATGGAAACTGGAGAGTTGATTATCTATACTTCAGCTGACCCTGTTTTGCAGATTGCTGCCCACGAAGACATTATTCCTTTGGATGAAT
NZ_CP035247.1:c1301115-1293263 1152   AGATGGAAACTGGAGAGTTGATTATCTATACTTCAGCTGACCCTGTTTTGCAGATTGCTGCCCACGAAGACATTATTCCTTTGGATGAAT
NZ_CP035238.1:738547-746413    1166   AGATGGAAACTGGAGAGTTGATTATCTATACTTCAGCTGACCCTGTTTTGCAGATTGCTGCCCACGAAGACATTATTCCTTTGGATGAAT
NZ_CP035255.1:776866-784718    1152   AGATGGAAACTGGAGAGTTGATTATCTATACTTCAGCTGACCCTGTTTTGCAGATTGCTGCCCACGAAGACATTATTCCTTTGGATGAAT
NZ_CP035256.1:807236-815102    1166   AGATGGAAACTGGAGAGTTGATTATCTATACTTCAGCTGACCCTGTTTTGCAGATTGCTGCCCACGAAGACATTATTCCTTTGGATGAAT
NZ_CP038252.1:1151812-1159673  1096   AGATGGAAACTGGAGAGTTGATTATCTATACTTCAGCTGACCCTGTTTTGCAGATTGCTGCCCACGAAGACATTATTCCTTTGGATGAAT
NZ_CP035254.1:793365-801213    1152   AGATGGAAACTGGAGAGTTGATTATCTATACTTCAGCTGACCCTGTTTTGCAGATTGCTGCCCACGAAGACATTATTCCTTTGGATGAAT
NZ_CP035251.1:748733-756585    1152   AGATGGAAACTGGAGAGTTGATTATCTATACTTCAGCTGACCCTGTTTTGCAGATTGCTGCCCACGAAGACATTATTCCTTTGGATGAAT
NZ_CP035252.1:745670-753536    1166   AGATGGAAACTGGAGAGTTGATTATCTATACTTCAGCTGACCCTGTTTTGCAGATTGCTGCCCACGAAGACATTATTCCTTTGGATGAAT
NZ_CP035257.1:776317-784169    1152   AGATGGAAACTGGAGAGTTGATTATCTATACTTCAGCTGACCCTGTTTTGCAGATTGCTGCCCACGAAGACATTATTCCTTTGGATGAAT
NZ_CP038253.1:758105-765957    1152   AGATGGAAACTGGAGAGTTGATTATCTATACTTCAGCTGACCCTGTTTTGCAGATTGCTGCCCACGAAGACATTATTCCTTTGGATGAAT
NZ_CP035263.1:743050-750901    1152   AGATGGAAACTGGAGAGTTGATTATCTATACTTCAGCTGACCCTGTTTTGCAGATTGCTGCCCACGAAGACATTATTCCTTTGGATGAAT
NZ_CP035262.1:741595-749460    1166   AGATGGAAACTGGAGAGTTGATTATCTATACTTCAGCTGACCCTGTTTTGCAGATTGCTGCCCACGAAGACATTATTCCTTTGGATGAAT
NZ_CP035236.1:764915-772766    1152   AGATGGAAACTGGAGAGTTGATTATCTATACTTCAGCTGACCCTGTTTTGCAGATTGCTGCCCACGAAGACATTATTCCTTTGGATGAAT
NZ_CP035253.1 767286-775134    1152   AGATGGAAACTGGAGAGTTGATTATCTATACTTCAGCTGACCCTGTTTTGCAGATTGCTGCCCACGAAGACATTATTCCTTTGGATGAAT
NZ_CP035249.1:763080-770921    1152   AGATGGAAACTGGAGAGTTGATTATCTATACTTCAGCTGACCCTGTTTTGCAGATTGCTGCCCACGAAGACATTATTCCTTTGGATGAAT

NC_003028.3:777000-787000      1971   TGTACCGTATCTGTGAATACGCTCGTTCGATTACCCTTGAGCGTCCTGCCCTTCTTGGTCGCATCATTGCTCGCCCTTATGTAGGTGAAC
NZ_CP035248.1:765099-775111    1981   TGTACCGTATCTGTGAATACGCTCGTTCAATTACACTTGAGCGCCCTGCCCTTCTAGGTCGTATCATTGCTCGCCCTTATGTAGGTGAAC
NZ_CP035235.1 c1326834-1317894 1256   TGTACCGTATCTGTGAATACGCTCGTTCGATTACCCTTGAGCGTCCTGCCCTTCTTGGTCGCATCATTGCTCGCCCTTATGTAGGTGAAC
NZ_CP035234.1:756284-765224    1256   TGTACCGTATCTGTGAATACGCTCGTTCGATTACCCTTGAGCGTCCTGCCCTTCTTGGTCGCATCATTGCTCGCCCTTATGTAGGTGAAC
NZ_CP038251.1:750745-759686    1256   TGTACCGTATCTGTGAATACGCTCGTTCGATTACCCTTGAGCGTCCTGCCCTTCTTGGTCGCATCATTGCTCGCCCTTATGTAGGTGAAC
NZ_CP035244.1:747023-755950    1242   TGTACCGTATCTGTGAATACGCTCGTTCGATTACCCTTGAGCGTCCTGCCCTTCTTGGTCGCATCATTGCTCGCCCTTATGTAGGTGAAC
NZ_CP035264.1:800836-809336    1242   TGTACCGTATCTGTGAATACGCTCGTTCGATTACCCTTGAGCGTCCTGCCCTTCTTGGTCGCATCATTGCTCGCCCTTATGTAGGTGAAC
NZ_CP035265.1:766128-774627    1242   TGTACCGTATCTGTGAATACGCTCGTTCGATTACCCTTGAGCGTCCTGCCCTTCTTGGTCGCATCATTGCTCGCCCTTATGTAGGTGAAC
NZ_CP035245.1 745261-753757    1242   TGTACCGTATCTGTGAATACGCTCGTTCGATTACCCTTGAGCGTCCTGCCCTTCTTGGTCGCATCATTGCTCGCCCTTATGTAGGTGAAC
NZ_CP035259.1:745700-753548    1242   TGTACCGTATCTGTGAATACGCTCGTTCGATTACCCTTGAGCGTCCTGCCCTTCTTGGTCGCATCATTGCTCGCCCTTATGTAGGTGAAC
NZ_CP035258.1:750874-758722    1242   TGTACCGTATCTGTGAATACGCTCGTTCGATTACCCTTGAGCGTCCTGCCCTTCTTGGTCGCATCATTGCTCGCCCTTATGTAGGTGAAC
NZ_CP035260.1:726156-733994    1242   TGTACCGTATCTGTGAATACGCTCGTTCGATTACCCTTGAGCGTCCTGCCCTTCTTGGCCGCATCATTGCTCGCCCTTATGTAGGTGAAC
```

```
NZ_CP035241.1:708296-716144    1242   TGTACCGTATCTGTGAATACGCTCGTTCGATTACCCTTGAGCGTCCTGCCCTTCTTGGTCGCATCATTGCTCGCCCTTATGTAGGTGAAC
NZ_CP035261.1:779086-786938    1242   TGTACCGTATCTGTGAATACGCTCGTTCGATTACCCTTGAGCGTCCTGCCCTTCTTGGTCGCATCATTGCTCGCCCTTATGTAGGTGAAC
NZ_CP035242.1:c1532236-1524385 1242   TGTACCGTATCTGTGAATACGCTCGTTCGATTACCCTTGAGCGTCCTGCCCTTCTTGGTCGCATCATTGCTCGCCCTTATGTAGGTGAAC
NZ_CP035243.1:777729-785580    1242   TGTACCGTATCTGTGAATACGCTCGTTCGATTACCCTTGAGCGTCCTGCCCTTCTTGGTCGCATCATTGCTCGCCCTTATGTAGGTGAAC
NZ_CP035237.1:c1311322-1303456 1256   TGTACCGTATCTGTGAATACGCTCGTTCGATTACCCTTGAGCGTCCTGCCCTTCTTGGTCGCATCATTGCTCGCCCTTATGTAGGTGAAC
NZ_CP035246.1:c1302892-1295026 1256   TGTACCGTATCTGTGAATACGCTCGTTCGATTACCCTTGAGCGTCCTGCCCTTCTTGGTCGCATCATTGCTCGCCCTTATGTAGGTGAAC
NZ_CP035247.1:c1301115-1293263 1242   TGTACCGTATCTGTGAATACGCTCGTTCGATTACCCTTGAGCGTCCTGCCCTTCTTGGTCGCATCATTGCTCGCCCTTATGTAGGTGAAC
NZ_CP035238.1:738547-746413    1256   TGTACCGTATCTGTGAATACGCTCGTTCGATTACCCTTGAGCGTCCTGCCCTTCTTGGTCGCATCATTGCTCGCCCTTATGTAGGTGAAC
NZ_CP035255.1:776866-784718    1242   TGTACCGTATCTGTGAATACGCTCGTTCGATTACCCTTGAGCGTCCTGCCCTTCTTGGTCGCATCATTGCTCGCCCTTATGTAGGTGAAC
NZ_CP035256.1:807236-815102    1256   TGTACCGTATCTGTGAATACGCTCGTTCGATTACCCTTGAGCGTCCTGCCCTTCTTGGTCGCATCATTGCTCGCCCTTATGTAGGTGAAC
NZ_CP038252.1:1151812-1159673  1186   TGTACCGTATCTGTGAATACGCTCGTTCGATTACCCTTGAGCGTCCTGCCCTTCTTGGTCGCATCATTGCTCGCCCTTATGTAGGTGAAC
NZ_CP035254.1:793365-801213    1242   TGTACCGTATCTGTGAATACGCTCGTTCGATTACCCTTGAGCGTCCTGCCCTTCTTGGTCGCATCATTGCTCGCCCTTATGTAGGTGAAC
NZ_CP035251.1:748733-756585    1242   TGTACCGTATCTGTGAATACGCTCGTTCGATTACCCTTGAGCGTCCTGCCCTTCTTGGTCGCATCATTGCTCGCCCTTATGTAGGTGAAC
NZ_CP035252.1:745670-753536    1256   TGTACCGTATCTGTGAATACGCTCGTTCGATTACCCTTGAGCGTCCTGCCCTTCTTGGTCGCATCATTGCTCGCCCTTATGTAGGTGAAC
NZ_CP035257.1:776317-784169    1242   TGTACCGTATCTGTGAATACGCTCGTTCGATTACCCTTGAGCGTCCTGCCCTTCTTGGTCGCATCATTGCTCGCCCTTATGTAGGTGAAC
NZ_CP038253.1:758105-765957    1242   TGTACCGTATCTGTGAATACGCTCGTTCGATTACCCTTGAGCGTCCTGCCCTTCTTGGTCGCATCATTGCTCGCCCTTATGTAGGTGAAC
NZ_CP035263.1:743050-750901    1242   TGTACCGTATCTGTGAATACGCTCGTTCGATTACCCTTGAGCGTCCTGCCCTTCTTGGTCGCATCATTGCTCGCCCTTATGTAGGTGAAC
NZ_CP035262.1:741595-749460    1256   TGTACCGTATCTGTGAATACGCTCGTTCGATTACCCTTGAGCGTCCTGCCCTTCTTGGTCGCATCATTGCTCGCCCTTATGTAGGTGAAC
NZ_CP035236.1:764915-772766    1242   TGTACCGTATCTGTGAATACGCTCGTTCGATTACCCTTGAGCGTCCTGCCCTTCTTGGTCGCATCATTGCTCGCCCTTATGTAGGTGAAC
NZ_CP035253.1 767286-775134    1242   TGTACCGTATCTGTGAATACGCTCGTTCGATTACCCTTGAGCGTCCTGCCCTTCTTGGTCGCATCATTGCTCGCCCTTATGTAGGTGAAC
NZ_CP035249.1:763080-770921    1242   TGTACCGTATCTGTGAATACGCTCGTTCGATTACCCTTGAGCGTCCTGCCCTTCTTGGTCGCATCATTGCTCGCCCTTATGTAGGTGAAC

NC_003028.3:777000-787000      2061   CAGGTAACTTCACTCGTACGGCAAACCGTCGTGACTTGGCTGTATCTCCATTTTTCCCAACTGTTTTGGATAAATTGAATGAGGCTGGTA
NZ_CP035248.1:765099-775111    2071   CAGGTAACTTCACTCGTACAGCAAACCGTCGTGACTTAGCCGTTTCTCCATTTGCTCCAACCGTTCTCGATAAACTCAATGAAGCTGGTA
NZ_CP035235.1 c1326834-1317894 1346   CAGGTAACTTCACTCGTACGGCAAACCGTCGTGACTTGGCTGTATCTCCATTTTCCCCAACTGTTTTGGATAAATTGAATGAGGCTGGTA
NZ_CP035234.1:756284-765224    1346   CAGGTAACTTCACTCGTACGGCAAACCGTCGTGACTTGGCTGTATCTCCATTTTCCCCAACTGTTTTGGATAAATTGAATGAGGCTGGTA
NZ_CP038251.1:750745-759686    1346   CAGGTAACTTCACTCGTACGGCAAACCGTCGTGACTTGGCTGTATCTCCATTTTCCCCAACTGTTTTGGATAAATTGAATGAGGCTGGTA
NZ_CP035244.1:747023-755950    1332   CAGGTAACTTCACTCGTACGGCAAACCGTCGTGACTTGGCTGTATCTCCATTTTCCCCAACTGTTTTGGATAAATTGAATGAGGCTGGTA
NZ_CP035264.1:800836-809336    1332   CAGGTAACTTCACTCGTACGGCAAACCGTCGTGACTTGGCTGTATCTCCATTTTCCCCAACTGTTTTGGATAAATTGAATGAGGCTGGTA
NZ_CP035265.1:766128-774627    1332   CAGGTAACTTCACTCGTACGGCAAACCGTCGTGACTTGGCTGTATCTCCATTTTCCCCAACTGTTTTGGATAAATTGAATGAGGCTGGTA
NZ_CP035245.1 745261-753757    1332   CAGGTAACTTCACTCGTACGGCAAACCGTCGTGACTTGGCTGTATCTCCATTTTCCCCAACTGTTTTGGATAAATTGAATGAGGCTGGTA
NZ_CP035259.1:745700-753548    1332   CAGGTAACTTCACTCGTACGGCAAACCGTCGTGACTTGGCTGTATCTCCATTTTCCCCAACTGTTTTGGATAAATTGAATGAGGCTGGTA
NZ_CP035258.1:750874-758722    1332   CAGGTAACTTCACTCGTACGGCAAACCGTCGTGACTTGGCTGTATCTCCATTTTCCCCAACTGTTTTGGATAAATTGAATGAGGCTGGTA
NZ_CP035260.1:726156-733994    1332   CAGGTAACTTCACTCGTACGGCAAACCGTCGTGACTTGGCTGTATCTCCATTTTCCCCAACTGTTTTGGATAAATTGAATGAGGCTGATA
NZ_CP035241.1:708296-716144    1332   CAGGTAACTTCACTCGTACGGCAAACCGTCGTGACTTGGCTGTATCTCCATTTTCCCCAACTGTTTTGGATAAATTGAATGAGGCTGATA
NZ_CP035261.1:779086-786938    1332   CAGGTAACTTCACTCGTACGGCAAACCGTCGTGACTTGGCTGTATCTCCATTTTCCCCAACTGTTTTGGATAAATTGAATGAGGCTGGTA
NZ_CP035242.1:c1532236-1524385 1332   CAGGTAACTTCACTCGTACGGCAAACCGTCGTGACTTGGCTGTATCTCCATTTTCCCCAACTGTTTTGGATAAATTGAATGAGGCTGGTA
NZ_CP035243.1:777729-785580    1332   CAGGTAACTTCACTCGTACGGCAAACCGTCGTGACTTGGCTGTATCTCCATTTTCCCCAACTGTTTTGGATAAATTGAATGAGGCTGGTA
NZ_CP035237.1:c1311322-1303456 1346   CAGGTAACTTCACTCGTACGGCAAACCGTCGTGACTTGGCTGTATCTCCATTTTCCCCAACTGTTTTGGATAAATTGAATGAGGCTGGTA
NZ_CP035246.1:c1302892-1295026 1346   CAGGTAACTTCACTCGTACGGCAAACCGTCGTGACTTGGCTGTATCTCCATTTTCCCCAACTGTTTTGGATAAATTGAATGAGGCTGGTA
NZ_CP035247.1:c1301115-1293263 1332   CAGGTAACTTCACTCGTACGGCAAACCGTCGTGACTTGGCTGTATCTCCATTTTCCCCAACTGTTTTGGATAAATTGAATGAGGCTGGTA
NZ_CP035238.1:738547-746413    1346   CAGGTAACTTCACTCGTACGGCAAACCGTCGTGACTTGGCTGTATCTCCATTTTCCCCAACTGTTTTGGATAAATTGAATGAGGCTGGTA
NZ_CP035255.1:776866-784718    1332   CAGGTAACTTCACTCGTACGGCAAACCGTCGTGACTTGGCTGTATCTCCATTTTCCCCAACTGTTTTGGATAAATTGAATGAGGCTGGTA
NZ_CP035256.1:807236-815102    1346   CAGGTAACTTCACTCGTACGGCAAACCGTCGTGACTTGGCTGTATCTCCATTTTCCCCAACTGTTTTGGATAAATTGAATGAGGCTGGTA
NZ_CP038252.1:1151812-1159673  1276   CAGGTAACTTCACTCGTACGGCAAACCGTCGTGACTTGGCTGTATCTCCATTTTCCCCAACTGTTTTGGATAAATTGAATGAGGCTGGTA
NZ_CP035254.1:793365-801213    1332   CAGGTAACTTCACTCGTACAGCAAACCGTCGTGACTTGGCTGTATCTCCATTTTCCCCAACTGTTTTGGATAAATTGAATGAGGCTGGTA
NZ_CP035251.1:748733-756585    1332   CAGGTAACTTCACTCGTACGGCAAACCGTCGTGACTTGGCTGTATCTCCATTTTCCCCAACTGTTTTGGATAAATTGAATGAGGCTGGTA
NZ_CP035252.1:745670-753536    1346   CAGGTAACTTCACTCGTACGGCAAACCGTCGTGACTTGGCTGTATCTCCATTTTCCCCAACTGTTTTGGATAAATTGAATGAGGCTGGTA
NZ_CP035257.1:776317-784169    1332   CAGGTAACTTCACTCGTACGGCAAACCGTCGTGACTTGGCTGTATCTCCATTTTCCCCAACTGTTTTGGATAAATTGAATGAGGCTGGTA
NZ_CP038253.1:758105-765957    1332   CAGGTAACTTCACTCGTACGGCAAACCGTCGTGACTTGGCTGTATCTCCATTTTCCCCAACTGTTTTGGATAAATTGAATGAGGCTGGTA
NZ_CP035263.1:743050-750901    1332   CAGGTAACTTCACTCGTACGGCAAACCGTCGTGACTTGGCTGTATCTCCATTTTCCCCAACTGTTTTGGATAAATTGAATGAGGCTGGTA
NZ_CP035262.1:741595-749460    1346   CAGGTAACTTCACTCGTACGGCAAACCGTCGTGACTTGGCTGTATCTCCATTTTCCCCAACTGTTTTGGATAAATTGAATGAGGCTGGTA
NZ_CP035236.1:764915-772766    1332   CAGGTAACTTCACTCGTACGGCAAACCGTCGTGACTTGGCTGTATCTCCATTTTCCCCAACTGTTTTGGATAAATTGAATGAGGCTGGTA
NZ_CP035253.1 767286-775134    1332   CAGGTAACTTCACTCGTACAGCAAACCGTCGTGACTTGGCTGTATCTCCATTTTCCCCAACTGTTTTGGATAAATTGAATGAGGCTGGTA
NZ_CP035249.1:763080-770921    1332   CAGGTAACTTCACTCGTACGGCAAACCGTCGTGACTTGGCTGTATCTCCATTTTCCCCAACTGTTTTGGATAAATTGAATGAGGCTGGTA

NC_003028.3:777000-787000      2151   TCGATACTTATGCTGTGGGTAAAATCAACGATATCTTTAACGGTGCTGGTATCAACCATGACATGGGTCACAACAAGTCAAATAGTCATG
NZ_CP035248.1:765099-775111    2161   TCGATACTTATGCTGTTGGTAAAATCAATGATATCTTTAACGGTGCTGGTATCAACCATGACATGGGCCACAACAAGTCAAACAGCCATG
NZ_CP035235.1 c1326834-1317894 1436   TCGATACTTATGCTGTGGGTAAAATCAACGATATCTTTAACGGTGCTGGTATCAACCATGACATGGGTCACAACAAGTCAAATAGTCATG
NZ_CP035234.1:756284-765224    1436   TCGATACTTATGCTGTGGGTAAAATCAACGATATCTTTAACGGTGCTGGTATCAACCATGACATGGGTCACAACAAGTCAAATAGTCATG
NZ_CP038251.1:750745-759686    1436   TCGATACTTATGCTGTGGGTAAAATCAACGATATCTTTAACGGTGCTGGTATCAACCATGACATGGGTCACAACAAGTCAAATAGTCATG
NZ_CP035244.1:747023-755950    1422   TCGATACTTATGCTGTGGGTAAAATCAACGATATCTTTAACGGTGCTGGTATCAACCATGACATGGGTCACAACAAGTCAAATAGTCATG
NZ_CP035264.1:800836-809336    1422   TCGATACTTATGCTGTGGGTAAAATCAACGATATCTTTAACGGTGCTGGTATCAACCATGACATGGGTCACAACAAGTCAAATAGTCATG
NZ_CP035265.1:766128-774627    1422   TCGATACTTATGCTGTGGGTAAAATCAACGATATCTTTAACGGTGCTGGTATCAACCATGACATGGGTCACAACAAGTCAAATAGTCATG
NZ_CP035245.1 745261-753757    1422   TCGATACTTATGCTGTGGGTAAAATCAACGATATCTTTAACGGTGCTGGTATCAACCATGACATGGGTCACAACAAGTCAAATAGTCATG
NZ_CP035259.1:745700-753548    1422   TCGATACTTATGCTGTGGGTAAAATCAACGATATCTTTAACGGTGCTGGTATCAACCATGACATGGGTCACAACAAGTCAAATAGTCATG
NZ_CP035258.1:750874-758722    1422   TCGATACTTATGCTGTGGGTAAAATCAACGATATCTTTAACGGTGCTGGTATCAACCATGACATGGGTCACAACAAGTCAAATAGTCATG
NZ_CP035260.1:726156-733994    1422   TCGATACTTATGCTGTGGGTAAAATCAACGATATCTTTAACGGTGCTGGTATCAACCATGACATGGGTCACAACAAGTCAAATAGTCATG
NZ_CP035241.1:708296-716144    1422   TCGATACTTATGCTGTGGGTAAAATCAACGATATCTTTAACGGTGCTGGTATCAACCATGACATGGGTCACAACAAGTCAAATAGTCATG
NZ_CP035261.1:779086-786938    1422   TCGATACTTATGCTGTGGGTAAAATCAACGATATCTTTAACGGTGCTGGTATCAACCATGACATGGGTCACAACAAGTCAAATAGTCATG
NZ_CP035242.1:c1532236-1524385 1422   TCGATACTTATGCTGTGGGTAAAATCAACGATATCTTTAACGGTGCTGGTATCAACCATGACATGGGTCACAACAAGTCAAATAGTCATG
NZ_CP035243.1:777729-785580    1422   TCGATACTTATGCTGTGGGTAAAATCAACGATATCTTTAACGGTGCTGGTATCAACCATGACATGGGTCACAACAAGTCAAATAGTCATG
NZ_CP035237.1:c1311322-1303456 1436   TCGATACTTATGCTGTGGGTAAAATCAACGATATCTTTAACGGTGCTGGTATCAACCATGACATGGGTCACAACAAGTCAAATAGTCATG
NZ_CP035246.1:c1302892-1295026 1436   TCGATACTTATGCTGTGGGTAAAATCAACGATATCTTTAACGGTGCTGGTATCAACCATGACATGGGTCACAACAAGTCAAATAGTCATG
NZ_CP035247.1:c1301115-1293263 1422   TCGATACTTATGCTGTGGGTAAAATCAACGATATCTTTAACGGTGCTGGTATCAACCATGACATGGGTCACAACAAGTCAAATAGTCATG
NZ_CP035238.1:738547-746413    1436   TCGATACTTATGCTGTGGGTAAAATCAACGATATCTTTAACGGTGCTGGTATCAACCATGACATGGGTCACAACAAGTCAAATAGTCATG
NZ_CP035255.1:776866-784718    1422   TCGATACTTATGCTGTGGGTAAAATCAACGATATCTTTAACGGCGCTGGTATCAACCATGACATGGGTCACAACAAGTCAAATAGTCATG
NZ_CP035256.1:807236-815102    1436   TCGATACTTATGCTGTGGGTAAAATCAACGATATCTTTAACGGCGCTGGTATCAACCATGACATGGGTCACAACAAGTCAAATAGTCATG
NZ_CP038252.1:1151812-1159673  1366   TCGATACTTATGCTGTGGGTAAAATCAACGATATCTTTAACGGTGCTGGTATCAACCATGACATGGGTCACAACAAGTCAAATAGTCATG
NZ_CP035254.1:793365-801213    1422   TCGATACTTATGCTGTGGGTAAAATCAACGATATCTTTAACGGCGCTGGTATCAACCATGACATGGGTCACAACAAGTCAAATAGTCATG
NZ_CP035251.1:748733-756585    1422   TCGATACTTATGCTGTGGGTAAAATCAACGATATCTTTAACGGTGCTGGTATCAACCATGACATGGGTCACAACAAGTCAAATAGTCATG
NZ_CP035252.1:745670-753536    1436   TCGATACTTATGCTGTGGGTAAAATCAACGATATCTTTAACGGTGCTGGTATCAACCATGACATGGGTCACAACAAGTCAAATAGTCATG
NZ_CP035257.1:776317-784169    1422   TCGATACTTATGCTGTGGGTAAAATCAACGATATCTTTAACGGTGCTGGTATCAACCATGACATGGGTCACAACAAGTCAAATAGTCATG
NZ_CP038253.1:758105-765957    1422   TCGATACTTATGCTGTGGGTAAAATCAACGATATCTTTAACGGTGCTGGTATCAACCATGACATGGGTCACAACAAGTCAAATAGTCATG
NZ_CP035263.1:743050-750901    1422   TCGATACTTATGCTGTGGGTAAAATCAACGATATCTTTAACGGTGCTGGTATCAACCATGACATGGGTCACAACAAGTCAAATAGTCATG
NZ_CP035262.1:741595-749460    1436   TCGATACTTATGCTGTGGGTAAAATCAACGATATCTTTAACGGTGCTGGTATCAACCATGACATGGGTCACAACAAGTCAAATAGTCATG
NZ_CP035236.1:764915-772766    1422   TCGATACTTATGCTGTGGGTAAAATCAACGATATCTTTAACGGTGCTGGTATCAACCATGACATGGGTCACAACAAGTCAAATAGTCATG
NZ_CP035253.1 767286-775134    1422   TCGATACTTATGCTGTGGGTAAAATCAACGATATCTTTAACGGCGCTGGTATCAACCATGACATGGGTCACAACAAGTCAAATAGTCATG
NZ_CP035249.1:763080-770921    1422   TCGATACTTATGCTGTGGGTAAAATCAACGATATCTTTAACGGTGCTGGTATCAACCATGACATGGGTCACAACAAGTCAAATAGTCATG

NC_003028.3:777000-787000      2241   GAATTGATACACTATTGAAGACTATGGGACTTGCTGAGTTTGAAAAAGGATTCTCATTCACAAACCTAGTTGACTTTGATGCCCTTTACG
NZ_CP035248.1:765099-775111    2251   GAATTGATACACTATTGAAGACCATGGGCCTTGCTGAGTTTGAAAAAGGATTCTCCTTCACAAACTTGGTTGACTTTGATGCCCTTTACG
NZ_CP035235.1 c1326834-1317894 1526   GAATTGATACACTATTGAAGACTATGGGACTTGCTGAGTTTGAAAAAGGATTCTCATTCACAAACCTAGTTGACTTTGATGCCCTTTACG
NZ_CP035234.1:756284-765224    1526   GAATTGATACACTATTGAAGACTATGGGACTTGCTGAGTTTGAAAAAGGATTCTCATTCACAAACCTAGTTGACTTTGATGCCCTTTACG
NZ_CP038251.1:750745-759686    1526   GAATTGATACACTATTGAAGACTATGGGACTTGCTGAGTTTGAAAAAGGATTCTCATTCACAAACCTAGTTGACTTTGATGCCCTTTACG
```

```
NZ_CP035244.1:747023-755950    1512   GAATTGATACACTATTGAAGACTATGGGACTTGCTGAGTTTGAAAAAGGATTCTCATTCACAAACCTAGTTGACTTTGATGCCCTTTACG
NZ_CP035264.1:800836-809336    1512   GAATTGATACACTATTGAAGACTATGGGACTTGCTGAGTTTGAAAAAGGATTCTCATTCACAAACCTAGTTGACTTTGATGCCCTTTACG
NZ_CP035265.1:766128-774627    1512   GAATTGATACACTATTGAAGACTATGGGACTTGCTGAGTTTGAAAAAGGATTCTCATTCACAAACCTAGTTGACTTTGATGCCCTTTACG
NZ_CP035245.1 745261-753757    1512   GAATTGATACACTATTGAAGACTATGGGACTTGCTGAGTTTGAAAAAGGATTCTCATTCACAAACCTAGTTGACTTTGATGCCCTTTACG
NZ_CP035259.1:745700-753548    1512   GAATTGATACACTATTGAAGACTATGGGACTTGCTGAGTTTGAAAAAGGATTCTCATTCACAAACCTAGTTGACTTTGATGCCCTTTACG
NZ_CP035258.1:750874-758722    1512   GAATTGATACACTATTGAAGACTATGGGACTTGCTGAGTTTGAAAAAGGATTCTCATTCACAAACCTAGTTGACTTTGATGCCCTTTACG
NZ_CP035260.1:726156-733994    1512   GAATTGATACACTATTGAAGACTATGGGACTTGCTGAGTTTGAAAAAGGATTCTCATTCACAAACCTAGTTGACTTTGATGCCCTTTACG
NZ_CP035241.1:708296-716144    1512   GAATTGATACACTATTGAAGACTATGGGACTTGCTGAGTTTGAAAAAGGATTCTCATTCACAAACCTAGTTGACTTTGATGCCCTTTACG
NZ_CP035261.1:779086-786938    1512   GAATTGATACACTATTGAAGACTATGGGACTTGCTGAGTTTGAAAAAGGATTCTCATTCACAAACCTAGTTGACTTTGATGCCCTTTACG
NZ_CP035242.1:c1532236-1524385 1512   GAATTGATACACTATTGAAGACTATGGGACTTGCTGAGTTTGAAAAAGGATTCTCATTTACAAACCTAGTTGACTTTGATGCCCTTTACG
NZ_CP035243.1:777729-785580    1512   GAATTGATACACTATTGAAGACTATGGGACTTGCTGAGTTTGAAAAAGGATTCTCATTTACAAACCTAGTTGACTTTGATGCCCTTTACG
NZ_CP035237.1:c1311322-1303456 1526   GAATTGATACACTATTGAAGACTATGGGACTTGCTGAGTTTGAAAAAGGATTCTCATTCACAAACCTAGTTGACTTTGATGCCCTTTACG
NZ_CP035246.1:c1302892-1295026 1526   GAATTGATACACTATTGAAGACTATGGGACTTGCTGAGTTTGAAAAAGGATTCTCATTCACAAACCTAGTTGACTTTGATGCCCTTTACG
NZ_CP035247.1:c1301115-1293263 1512   GAATTGATACACTATTGAAGACTATGGGACTTGCTGAGTTTGAAAAAGGATTCTCATTCACAAACCTAGTTGACTTTGATGCCCTTTACG
NZ_CP035238.1:738547-746413    1526   GAATTGATACACTATTGAAGACTATGGGACTTGCTGAGTTTGAAAAAGGATTCTCATTCACAAACCTAGTTGACTTTGATGCCCTTTACG
NZ_CP035255.1:776866-784718    1512   GAATTGATACACTATTGAAGACTATGGGACTTGCTGAGTTTGAAAAAGGATTCTCATTCACAAACCTAGTTGACTTTGATGCCCTTTACG
NZ_CP035256.1:807236-815102    1526   GAATTGATACACTATTGAAGACTATGGGACTTGCTGAGTTTGAAAAAGGATTCTCATTCACAAACCTAGTTGACTTTGATGCCCTTTACG
NZ_CP038252.1:1151812-1159673  1456   GAATTGATACACTATTGAAGACTATGGGACTTGCTGAGTTTGAAAAAGGATTCTCATTCACAAACCTAGTTGACTTTGATGCCCTTTACG
NZ_CP035254.1:793365-801213    1512   GAATTGATACACTATTGAAGACTATGGGACTTGCTGAGTTTGAAAAAGGATTCTCATTCACAAACCTAGTTGACTTTGATGCCCTTTACG
NZ_CP035251.1:748733-756585    1512   GAATTGATACACTATTGAAGACTATGGGACTTGCTGAGTTTGAAAAAGGATTCTCATTCACAAACCTAGTTGACTTTGATGCCCTTTACG
NZ_CP035252.1:745670-753536    1526   GAATTGATACACTATTGAAGACTATGGGACTTGCTGAGTTTGAAAAAGGATTCTCATTCACAAACCTAGTTGACTTTGATGCCCTTTACG
NZ_CP035257.1:776317-784169    1512   GAATTGATACACTATTGAAGACTATGGGACTTGCTGAGTTTGAAAAAGGATTCTCATTCACAAACCTAGTTGACTTTGATGCCCTTTACG
NZ_CP038253.1:758105-765957    1512   GAATTGATACACTATTGAAGACTATGGGACTTGCTGAGTTTGAAAAAGGATTCTCATTCACAAACCTAGTTGACTTTGATGCCCTTTACG
NZ_CP035263.1:743050-750901    1512   GAATTGATACACTATTGAAGACTATGGGACTTGCTGAGTTTGAAAAAGGATTCTCATTCACAAACCTAGTTGACTTTGATGCCCTTTACG
NZ_CP035262.1:741595-749460    1526   GAATTGATACACTATTGAAGACTATGGGACTTGCTGAGTTTGAAAAAGGATTCTCATTCACAAACCTAGTTGACTTTGATGCCCTTTACG
NZ_CP035236.1:764915-772766    1512   GAATTGATACACTATTGAAGACTATGGGACTTGCTGAGTTTGAAAAAGGATTCTCATTCACAAACCTAGTTGACTTTGATGCCCTTTACG
NZ_CP035253.1 767286-775134    1512   GAATTGATACACTATTGAAGACTATGGGACTTGCTGAGTTTGAAAAAGGATTCTCATTCACAAACCTAGTTGACTTTGATGCCCTTTACG
NZ_CP035249.1:763080-770921    1512   GAATTGATACACTATTGAAGACTATGGGACTTGCTGAGTTTGAAAAAGGATTCTCATTCACAAACCTAGTTGACTTTGATGCCCTTTACG

NC_003028.3:777000-787000      2331   GCCATCGTCGTAATGCTCACGGTTACCGTGATTGCTTGCATGAGTTTGATGAACGCTTACCTGAAATTATCGCAGCTATGAGAGAGAATG
NZ_CP035248.1:765099-775111    2341   GCCACCGTCGCAATGCTCATGGTTACCGTGATTGCTTGCATGAGTTTGATGAACGCTTGCCTGAAATTATCGCAGCCATGAGAGAGAATG
NZ_CP035235.1 c1326834-1317894 1616   GCCATCGTCGTAATGCTCACGGTTACCGTGATTGCTTGCATGAGTTTGATGAACGCTTACCTGAAATTATCGCAGCTATGAGAGAGAATG
NZ_CP035234.1:756284-765224    1616   GCCATCGTCGTAATGCTCACGGTTACCGTGATTGCTTGCATGAGTTTGATGAACGCTTACCTGAAATTATCGCAGCTATGAGAGAGAATG
NZ_CP038251.1:750745-759686    1616   GCCATCGTCGTAATGCTCACGGTTACCGTGATTGCTTGCATGAGTTTGATGAACGCTTACCTGAAATTATCGCAGCTATGAGAGAGAATG
NZ_CP035244.1:747023-755950    1602   GCCATCGTCGTAATGCTCACGGTTACCGTGATTGCTTGCATGAGTTTGATGAACGCTTACCTGAAATTATCGCAGCTATGAGAGAGAATG
NZ_CP035264.1:800836-809336    1602   GCCATCGTCGTAATGCTCACGGTTACCGTGATTGCTTGCATGAGTTTGATGAACGCTTACCTGAAATTATCGCAGCTATGAGAGAGAATG
NZ_CP035265.1:766128-774627    1602   GCCATCGTCGTAATGCTCACGGTTACCGTGATTGCTTGCATGAGTTTGATGAACGCTTACCTGAAATTATCGCAGCTATGAGAGAGAATG
NZ_CP035245.1 745261-753757    1602   GCCATCGTCGTAATGCTCACGGTTACCGTGATTGCTTGCATGAGTTTGATGAACGCTTACCTGAAATTATCGCAGCTATGAGAGAGAATG
NZ_CP035259.1:745700-753548    1602   GCCATCGTCGTAATGCTCACGGTTACCGTGATTGCTTGCATGAGTTTGATGAACGCTTACCTGAAATTATCGCAGCTATGAGAGAGAATG
NZ_CP035258.1:750874-758722    1602   GCCATCGTCGTAATGCTCACGGTTACCGTGATTGCTTGCATGAGTTTGATGAACGCTTACCTGAAATTATCGCAGCTATGAGAGAGAATG
NZ_CP035260.1:726156-733994    1602   GCCATCGTCGTAATGCTCACGGTTACCGTGATTGCTTGCATGAGTTTGATGAACGCTTACCTGAAATTATCGCAGCTATGAGAGAGAATG
NZ_CP035241.1:708296-716144    1602   GCCATCGTCGTAATGCTCACGGTTACCGTGATTGCTTGCATGAGTTTGATGAACGCTTACCTGAAATTATCGCAGCTATGAGAGAGAATG
NZ_CP035261.1:779086-786938    1602   GCCATCGTCGTAATGCTCACGGTTACCGTGATTGCTTGCATGAGTTTGATGAACGCTTACCTGAAATTATCGCAGCTATGAGAGAGAATG
NZ_CP035242.1:c1532236-1524385 1602   GCCATCGTCGTAATGCTCACGGTTACCGTGATTGCTTGCATGAGTTTGATGAACGCTTACCTGAAATTATCGCAGCTATGAGAGAGAATG
NZ_CP035243.1:777729-785580    1602   GCCATCGTCGTAATGCTCACGGTTACCGTGATTGCTTGCATGAGTTTGATGAACGCTTACCTGAAATTATCGCAGCTATGAGAGAGAATG
NZ_CP035237.1:c1311322-1303456 1616   GCCATCGTCGTAATGCTCACGGTTACCGTGATTGCTTGCATGAGTTTGATGAACGCTTACCTGAAATTATCGCAGCTATGAGAGAGAATG
NZ_CP035246.1:c1302892-1295026 1616   GCCATCGTCGTAATGCTCACGGTTACCGTGATTGCTTGCATGAGTTTGATGAACGCTTACCTGAAATTATCGCAGCTATGAGAGAGAATG
NZ_CP035247.1:c1301115-1293263 1602   GCCATCGTCGTAATGCTCACGGTTACCGTGATTGCTTGCATGAGTTTGATGAACGCTTACCTGAAATTATCGCAGCTATGAGAGAGAATG
NZ_CP035238.1:738547-746413    1616   GCCATCGTCGTAATGCTCACGGTTACCGTGATTGCTTGCATGAGTTTGATGAACGCTTACCTGAAATTATCGCAGCTATGAGAGAGAATG
NZ_CP035255.1:776866-784718    1602   GCCATCGTCGTAATGCTCACGGTTACCGTGATTGCTTGCATGAGTTTGATGAACGCTTACCTGAAATTATCGCAGCTATGAGAGAGAATG
NZ_CP035256.1:807236-815102    1616   GCCATCGTCGTAATGCTCACGGTTACCGTGATTGCTTGCATGAGTTTGATGAACGCTTACCTGAAATTATCGCAGCTATGAGAGAGAATG
NZ_CP038252.1:1151812-1159673  1546   GCCATCGTCGTAATGCTCACGGTTACCGTGATTGCTTGCATGAGTTTGATGAACGCTTACCTGAAATTATCGCAGCTATGAGAGAGAATG
NZ_CP035254.1:793365-801213    1602   GCCATCGTCGTAATGCTCACGGTTACCGTGATTGCTTGCATGAGTTTGATGAACGCTTACCTGAAATTATCGCAGCTATGAGAGAGAATG
NZ_CP035251.1:748733-756585    1602   GCCATCGTCGTAATGCTCACGGTTACCGTGATTGTTTGCATGAGTTTGATGAACGCTTACCTGAAATTATCGCAGCTATGAGAGAGAATG
NZ_CP035252.1:745670-753536    1616   GCCATCGTCGTAATGCTCACGGTTACCGTGATTGTTTGCATGAGTTTGATGAACGCTTACCTGAAATTATCGCAGCTATGAGAGAGAATG
NZ_CP035257.1:776317-784169    1602   GCCATCGTCGTAATGCTCACGGTTACCGTGATTGTTTGCATGAGTTTGATGAACGCTTACCTGAAATTATCGCAGCTATGAGAGAGAATG
NZ_CP038253.1:758105-765957    1602   GCCATCGTCGTAATGCTCACGGTTACCGTGATTGTTTGCATGAGTTTGATGAACGCTTACCTGAAATTATCGCAGCTATGAGAGAGAATG
NZ_CP035263.1:743050-750901    1602   GCCATCGTCGTAATGCTCACGGTTACCGTGATTGTTTGCATGAGTTTGATGAACGCTTACCTGAAATTATCGCAGCTATGAGAGAGAATG
NZ_CP035262.1:741595-749460    1616   GCCATCGTCGTAATGCTCACGGTTACCGTGATTGTTTGCATGAGTTTGATGAACGCTTACCTGAAATTATCGCAGCTATGAGAGAGAATG
NZ_CP035236.1:764915-772766    1602   GCCATCGTCGTAATGCTCACGGTTACCGTGATTGTTTGCATGAGTTTGATGAACGCTTACCTGAAATTATCGCAGCTATGAGAGAGAATG
NZ_CP035253.1 767286-775134    1602   GCCATCGTCGTAATGCTCACGGTTACCGTGATTGCTTGCATGAGTTTGATGAACGCTTACCTGAAATTATCGCAGCTATGAGAGAGAATG
NZ_CP035249.1:763080-770921    1602   GCCATCGTCGTAATGCTCACGGTTACCGTGATTGCTTGCATGAGTTTGATGAACGCTTACCTGAAATTATCGCAGCTATGAGAGAGAATG

NC_003028.3:777000-787000      2421   ACCTTCTCTTGATTACTGCGGACCATGGAAATGACCCAACGTATGCAGGAACGGATCACACTCGGGAATATATTCCATTGTTGGCCTATA
NZ_CP035248.1:765099-775111    2431   ATCTTCTTTTAATTACTGCGGACCATGGAAATGACCCAACCTATGCAGGAACAGACCACACTCGTGAATATATCCCACTTTTGGCTTACA
NZ_CP035235.1 c1326834-1317894 1706   ACCTTCTCTTGATTACTGCGGACCATGGAAATGACCCAACGTATGCAGGAACGGATCACACTCGGGAATATATTCCATTGTTGGCCTATA
NZ_CP035234.1:756284-765224    1706   ACCTTCTCTTGATTACTGCGGACCATGGAAATGACCCAACGTATGCAGGAACGGATCACACTCGGGAATATATTCCATTGTTGGCCTATA
NZ_CP038251.1:750745-759686    1706   ACCTTCTCTTGATTACTGCGGACCATGGAAATGACCCAACGTATGCAGGAACGGATCACACTCGGGAATATATTCCATTGTTGGCCTATA
NZ_CP035244.1:747023-755950    1692   ACCTTCTCTTGATTACTGCGGACCATGGAAATGACCCAACGTATGCAGGAACGGATCACACTCGGGAATATATTCCATTGTTGGCCTATA
NZ_CP035264.1:800836-809336    1692   ACCTTCTCTTGATTACTGCGGACCATGGAAATGACCCAACGTATGCAGGAACGGATCACACTCGGGAATATATTCCATTGTTGGCCTATA
NZ_CP035265.1:766128-774627    1692   ACCTTCTCTTGATTACTGCGGACCATGGAAATGACCCAACGTATGCAGGAACGGATCACACTCGGGAATATATTCCATTGTTGGCCTATA
NZ_CP035245.1 745261-753757    1692   ACCTTCTCTTGATTACTGCGGACCATGGAAATGACCCAACGTATGCAGGAACGGATCACACTCGGGAATATATTCCATTGTTGGCCTATA
NZ_CP035259.1:745700-753548    1692   ACCTTCTCTTGATTACTGCGGACCATGGAAATGACCCAACGTATGCAGGAACGGATCACACTCGGGAATATATTCCATTGTTGGCCTATA
NZ_CP035258.1:750874-758722    1692   ACCTTCTCTTGATTACTGCGGACCATGGAAATGACCCAACGTATGCAGGAACGGATCACACTCGGGAATATATTCCATTGTTGGCCTATA
NZ_CP035260.1:726156-733994    1692   ACCTTCTCTTGATTACTGCGGACCATGGAAATGACCCAACGTATGCAGGAACGGATCACACTCGGGAATATATTCCATTGTTGGCCTATA
NZ_CP035241.1:708296-716144    1692   ACCTTCTCTTGATTACTGCGGACCATGGAAATGACCCAACGTATGCAGGAACGGATCACACTCGGGAATATATTCCATTGTTGGCCTATA
NZ_CP035261.1:779086-786938    1692   ACCTTCTCTTGATTACTGCGGACCATGGAAATGACCCAACGTATGCAGGAACGGATCACACTCGGGAATATATTCCATTGTTGGCCTATA
NZ_CP035242.1:c1532236-1524385 1692   ACCTTCTCTTGATTACTGCGGACCATGGAAATGACCCAACGTATGCAGGAACGGATCACACTCGGGAATATATTCCATTGTTGGCCTATA
NZ_CP035243.1:777729-785580    1692   ACCTTCTCTTGATTACTGCGGACCATGGAAATGACCCAACGTATGCAGGAACGGATCACACTCGGGAATATATTCCATTGTTGGCCTATA
NZ_CP035237.1:c1311322-1303456 1706   ACCTTCTCTTGATTACTGCGGACCATGGAAATGACCCAACGTATGCAGGAACGGATCACACTCGGGAATATATTCCATTGTTGGCATATA
NZ_CP035246.1:c1302892-1295026 1706   ACCTTCTCTTGATTACTGCGGACCATGGAAATGACCCAACGTATGCAGGAACGGATCACACTCGGGAATATATTCCATTGTTGGCCTATA
NZ_CP035247.1:c1301115-1293263 1692   ACCTTCTCTTGATTACTGCGGACCATGGAAATGACCCAACGTATGCAGGAACGGATCACACTCGGGAATATATTCCATTGTTGGCCTATA
NZ_CP035238.1:738547-746413    1706   ACCTTCTCTTGATTACTGCGGATCATGGAAATGACCCAACGTATGCAGGAACGGATCACACTCGGGAATATATTCCATTGTTGGCCTATA
NZ_CP035255.1:776866-784718    1692   ACCTTCTCTTGATTACTGCGGACCATGGAAATGACCCAACGTATGCAGGAACGGATCACACTCGGGAATATATTCCATTGTTGGCCTATA
NZ_CP035256.1:807236-815102    1706   ACCTTCTCTTGATTACTGCGGACCATGGAAATGACCCAACGTATGCAGGAACGGATCACACTCGGGAATATATTCCATTGTTGGCCTATA
NZ_CP038252.1:1151812-1159673  1636   ACCTTCTCTTGATTACTGCGGACCATGGAAATGACCCAACGTATGCAGGAACGGATCACACTCGGGAATATATTCCATTGTTGGCCTATA
NZ_CP035254.1:793365-801213    1692   ACCTTCTCTTGATTACTGCGGACCATGGAAATGACCCAACGTATGCAGGAACGGATCACACTCGGGAATATATTCCATTGTTGGCCTATA
NZ_CP035251.1:748733-756585    1692   ACCTTCTCTTGATTACTGCGGACCATGGAAATGACCCAACGTATGCAGGGACGGATCACACTCGGGAATATATTCCATTGTTGGCATATA
NZ_CP035252.1:745670-753536    1706   ACCTTCTCTTGATTACTGCGGACCATGGAAATGACCCAACGTATGCAGGGACGGATCACACTCGGGAATATATTCCATTGTTGGCATATA
NZ_CP035257.1:776317-784169    1692   ACCTTCTCTTGATTACTGCGGACCATGGAAATGACCCAACGTATGCAGGGACGGATCACACTCGGGAATATATTCCATTGTTGGCATATA
NZ_CP038253.1:758105-765957    1692   ACCTTCTCTTGATTACTGCGGACCATGGAAATGACCCAACGTATGCAGGGACGGATCACACTCGGGAATATATTCCATTGTTGGCATATA
NZ_CP035263.1:743050-750901    1692   ACCTTCTCTTGATTACTGCGGACCATGGAAATGACCCAACGTATGCAGGGACGGATCACACTCGGGAATATATTCCATTGTTGGCATATA
NZ_CP035262.1:741595-749460    1706   ACCTTCTCTTGATTACTGCGGACCATGGAAATGACCCAACGTATGCAGGGACGGATCACACTCGGGAATATATTCCATTGTTGGCATATA
NZ_CP035236.1:764915-772766    1692   ACCTTCTCTTGATTACTGCGGACCATGGAAATGACCCAACGTATGCAGGGACGGATCACACTCGGGAATATATTCCATTGTTGGCATATA
NZ_CP035253.1 767286-775134    1692   ACCTTCTCTTGATTACTGCGGACCATGGAAATGACCCAACGTATGCAGGAACGGATCACACTCGGGAATATATTCCATTGTTGGCCTATA
```

```
NZ_CP035249.1:763080-770921    1692   ACCTTCTCTTGATTACTGCGGACCATGGAAATGACCCAACGTATGCAGGAACGGATCACACTCGGGAATATATTCCATTGTTGGCCTATA

NC_003028.3:777000-787000      2511   GCCCTGCCTTTAAAGGAAATGGTCTCATTCCAGTAGGACATTTTGCAGATATTTCAGCGACTGTTGCCGATAACTTTGGTGTGGAAACTG
NZ_CP035248.1:765099-775111    2521   GTCCTTCCTTTAAAGGAAATGGTGTCATTCCAGTTGGACATTTTGCAGATATCTCAGCGACAGTTGCGGATAACTTTGGTGTTGAAACTG
NZ_CP035235.1 c1326834-1317894 1796   GCCCTGCCTTTAAAGGAAATGGTCTCATTCCAGTAGGACATTTTGCAGATATTTCAGCGACTGTTGCCGATAACTTTGGTGTGGAAACTG
NZ_CP035234.1:756284-765224    1796   GCCCTGCCTTTAAAGGAAATGGTCTCATTCCAGTAGGACATTTTGCAGATATTTCAGCGACTGTTGCCGATAACTTTGGTGTGGAAACTG
NZ_CP038251.1:750745-759686    1796   GCCCTGCCTTTAAAGGAAATGGTCTCATTCCAGTAGGACATTTTGCAGATATTTCAGCGACTGTTGCCGATAACTTTGGTGTGGAAACTG
NZ_CP035244.1:747023-755950    1782   GCCCTGCCTTTAAAGGAAATGGTCTCATTCCAGTAGGACATTTTGCAGATATTTCAGCGACTGTTGCCGATAACTTTGGTGTGGAAACTG
NZ_CP035264.1:800836-809336    1782   GCCCTGCCTTTAAAGGAAATGGTCTCATTCCAGTAGGACATTTTGCGGATATTTCAGCGACTGTTGCCGATAACTTTGGTGTGGAAACTG
NZ_CP035265.1:766128-774627    1782   GCCCTGCCTTTAAAGGAAATGGTCTCATTCCAGTAGGACATTTTGCGGATATTTCAGCGACTGTTGCCGATAACTTTGGTGTGGAAACTG
NZ_CP035245.1 745261-753757    1782   GCCCTGCCTTTAAAGGAAATGGTCTCATTCCAGTAGGACATTTTGCAGATATTTCAGCGACTGTTGCCGATAACTTTGGTGTGGAAACTG
NZ_CP035259.1:745700-753548    1782   GCCCTGCCTTTAAAGGAAATGGTCTCATTCCAGTAGGACATTTTGCAGATATTTCAGCGACTGTTGCCGATAACTTTGGTGTGGAAACTG
NZ_CP035258.1:750874-758722    1782   GCCCTGCCTTTAAAGGAAATGGTCTCATTCCAGTAGGACATTTTGCAGATATTTCAGCGACTGTTGCCGATAACTTTGGTGTGGAAACTG
NZ_CP035260.1:726156-733994    1782   GCCCTGCCTTTAAAGGAAATGGTCTCATTCCAGTAGGACATTTTGCAGATATTTCAGCGACTGTTGCCGATAACTTTGGTGTGGAAACTG
NZ_CP035241.1:708296-716144    1782   GCCCTGCCTTTAAAGGAAATGGTCTCATTCCAGTAGGACATTTTGCAGATATTTCAGCGACTGTTGCCGATAACTTTGGTGTGGAAACTG
NZ_CP035261.1:779086-786938    1782   GCCCTGCCTTTAAAGGAAATGGTCTCATTCCAGTAGGACATTTTGCAGATATTTCAGCGACTGTTGCCGATAACTTTGGTGTGGAAACTG
NZ_CP035242.1:c1532236-1524385 1782   GCCCTGCCTTTAAAGGAAATGGTCTCATTCCAGTAGGACATTTTGCAGATATTTCAGCGACTGTTGCCGATAACTTTGGTGTGGAAACTG
NZ_CP035243.1:777729-785580    1782   GCCCTGCCTTTAAAGGAAATGGTCTCATTCCAGTAGGACATTTTGCAGATATTTCAGCGACTGTTGCCGATAACTTTGGTGTGGAAACTG
NZ_CP035237.1:c1311322-1303456 1796   GCCCTGCCTTTAAAGGAAATGGTCTCATTCCAGTAGGACATTTTGCAGATATTTCAGCGACTGTTGCCGATAACTTTGGTGTGGAAACTG
NZ_CP035246.1:c1302892-1295026 1796   GCCCTGCCTTTAAAGGAAATGGTCTCATTCCAGTAGGACATTTTGCAGATATTTCAGCGACTGTTGCCGATAACTTTGGTGTGGAAACTG
NZ_CP035247.1:c1301115-1293263 1782   GCCCTGCCTTTAAAGGAAATGGTCTCATTCCAGTAGGACATTTTGCAGATATTTCAGCGACTGTTGCCGATAACTTTGGTGTGGAAACTG
NZ_CP035238.1:738547-746413    1796   GCCCTGCCTTTAAAGGAAATGGTCTCATTCCAGTAGGACATTTTGCAGATATTTCAGCGACTGTTGCCGATAACTTTGGTGTGGAAACTG
NZ_CP035255.1:776866-784718    1782   GTCCTGCCTTTAAAGGAAATGGTCTCATTCCAGTAGGACATTTTGCAGATATTTCAGCGACTGTTGCCGATAACTTTGGTGTGGAAACTG
NZ_CP035256.1:807236-815102    1796   GTCCTGCCTTTAAAGGAAATGGTCTCATTCCAGTAGGACATTTTGCAGATATTTCAGCGACTGTTGCCGATAACTTTGGTGTGGAAACTG
NZ_CP038252.1:1151812-1159673  1726   GCCCTGCCTTTAAAGGAAATGGTCTCATTCCAGTAGGACATTTTGCAGATATTTCAGCGACTGTTGCCGATAACTTTGGTGTGGAAACTG
NZ_CP035254.1:793365-801213    1782   GTCCTGCCTTTAAAGGAAATGGTCTCATTCCAGTAGGACATTTTGCAGATATTTCAGCGACTGTTGCCGATAACTTTGGTGTGGAAACTG
NZ_CP035251.1:748733-756585    1782   GCCCTGCCTTTAAAGGAAATGGTCTCATTCCAGTAGGACATTTTGCAGATATTTCAGCGACTGTTGCCGATAACTTTGGTGTGGAAACTG
NZ_CP035252.1:745670-753536    1796   GCCCTGCCTTTAAAGGAAATGGTCTCATTCCAGTAGGACATTTTGCAGATATTTCAGCGACTGTTGCCGATAACTTTGGTGTGGAAACTG
NZ_CP035257.1:776317-784169    1782   GCCCTGCCTTTAAAGGAAATGGTCTCATTCCAGTAGGACATTTTGCAGATATTTCAGCGACTGTTGCCGATAACTTTGGTGTGGAAACTG
NZ_CP038253.1:758105-765957    1782   GCCCTGCCTTTAAAGGAAATGGTCTCATTCCAGTAGGACATTTTGCAGATATTTCAGCGACTGTTGCCGATAACTTTGGTGTGGAAACTG
NZ_CP035263.1:743050-750901    1782   GCCCTGCCTTTAAAGGAAATGGTCTCATTCCAGTAGGACATTTTGCAGATATTTCAGCGACTGTTGCCGATAACTTTGGTGTGGAAACTG
NZ_CP035262.1:741595-749460    1796   GCCCTGCCTTTAAAGGAAATGGTCTCATTCCAGTAGGACATTTTGCAGATATTTCAGCGACTGTTGCCGATAACTTTGGTGTGGAAACTG
NZ_CP035236.1:764915-772766    1782   GCCCTGCCTTTAAAGGAAATGGTCTCATTCCAGTAGGACATTTTGCAGATATTTCAGCGACTGTTGCCGATAACTTTGGTGTGGAAACTG
NZ_CP035253.1 767286-775134    1782   GTCCTGCCTTTAAAGGAAATGGTCTCATTCCAGTAGGACATTTTGCAGATATTTCAGCGACTGTTGCCGATAACTTTGGTGTGGAAACTG
NZ_CP035249.1:763080-770921    1782   GCCCTGCCTTTAAAGGAAATGGTCTCATTCCAGTAGGACATTTTGCAGATATTTCAGCGACTGTTGCCGATAACTTTGGTGTGGAAACTG

NC_003028.3:777000-787000      2601   CTATGATTGGGGAAAGTTTCTTAGATAAATTGGTATAAGATGACGCGCTATGCTTTGCTGGTGAGAGGTATCAATGTTGGTGGTAAGAAT
NZ_CP035248.1:765099-775111    2611   CCATGATTGGGGAAAGTTTCTTAGATAAATTGGTATAAGATGACGCGATATGCTTTACTTGTTCGGGGCATTAATGTAGGTGGGAAAAAT
NZ_CP035235.1 c1326834-1317894 1886   CTATGATTGGGGAAAGTTTCTTAGATAAATTGGTATAAGATGACGCGCTATTCTTTGCTGGTGAGAGGTATCAATGTTGGTGGTAAGAAT
NZ_CP035234.1:756284-765224    1886   CTATGATTGGGGAAAGTTTCTTAGATAAATTGGTATAAGATGACGCGCTATTCTTTGCTGGTGAGAGGTATCAATGTTGGTGGTAAGAAT
NZ_CP038251.1:750745-759686    1886   CTATGATTGGGGAAAGTTTCTTAGATAAATTGGTATAAGATGACGCGCTATTCTTTGCTGGTGAGAGGTATCAATGTTGGTGGTAAGAAT
NZ_CP035244.1:747023-755950    1872   CTATGATTGGGGAAAGTTTCTTAGATAAATTGGTATAAGATGACGCGCTATTCTTTGCTGGTGAGAGGTATCAATGTTGGTGGTAAGAAT
NZ_CP035264.1:800836-809336    1872   CTATGATTGGGGAAAGTTTCTTAGATAAATTGGTATAAGATGACGCGCTATGCTTTGCTGGTGAGAGGTATCAATGTTGGTGGTAAGAAT
NZ_CP035265.1:766128-774627    1872   CTATGATTGGGGAAAGTTTCTTAGATAAATTGGTATAAGATGACGCGCTATGCTTTGCTGGTGAGAGGTATCAATGTTGGTGGTAAGAAT
NZ_CP035245.1 745261-753757    1872   CTATGATTGGGGAAAGTTTCTTAGATAAATTGGTATAAGATGACGCGCTATGCTTTGCTGGTGAGAGGTATCAATGTTGGTGGTAAGAAT
NZ_CP035259.1:745700-753548    1872   CTATGATTGGGGAAAGTTTCTTAGATAAATTGGTATAAGATGACGCGCTATGCTTTGCTGGTGAGAGGTATCAATGTTGGTGGTAAGAAT
NZ_CP035258.1:750874-758722    1872   CTATGATTGGGGAAAGTTTCTTAGATAAATTGGTATAAGATGACGCGCTATGCTTTGCTGGTGAGAGGTATCAATGTTGGTGGTAAGAAT
NZ_CP035260.1:726156-733994    1872   CTATGATTGGGGAAAGTTTCTTAGATAAATTGGTATAAGATGACGCGCTATGCTTTGCTGGTGAGAGGTATCAATGTTGGTAGTAAGAAT
NZ_CP035241.1:708296-716144    1872   CTATGATTGGGGAAAGTTTCTTAGATAAATTGGTATAAGATGACGCGCTATGCTTTGCTGGTGAGAGGTATCAATGTTGGTAGTAAGAAT
NZ_CP035261.1:779086-786938    1872   CTATGATTGGGGAAAGTTTCTTAGATAAATTGGTATAAGATGACGCGCTATGCTTTGCTGGTGAGAGGTATCAATGTTGGTGGTAAGAAT
NZ_CP035242.1:c1532236-1524385 1872   CTATGATTGGGGAAAGTTTCTTAGATAAATTGGTATAAGATGACGCGCTATGCTTTGCTGGTGAGAGGTATCAATGTTGGTGGTAAGAAT
NZ_CP035243.1:777729-785580    1872   CTATGATTGGGGAAAGTTTCTTAGATAAATTGGTATAAGATGACGCGCTATGCTTTGCTGGTGAGAGGTATCAATGTTGGTGGTAAGAAT
NZ_CP035237.1:c1311322-1303456 1886   CTATGATTGGGGAAAGTTTCTTAGATAAATTGGTATAATATGACGCGCTATGCTTTGCTGGTGAGAGGTATCAATGTTGGTGGTAAGAAT
NZ_CP035246.1:c1302892-1295026 1886   CTATGATTGGGGAAAGTTTCTTAGATAAATTGGTATAAGATGACGCGCTATGCTTTGCTGGTGAGAGGTATCAATGTTGGTGGTAAGAAT
NZ_CP035247.1:c1301115-1293263 1872   CTATGATTGGGGAAAGTTTCTTAGATAAATTGGTATAAGATGACGCGCTATGCTTTGCTGGTGAGAGGTATCAATGTTGGTGGTAAGAAT
NZ_CP035238.1:738547-746413    1886   CTATGATTGGGGAAAGTTTCTTAGATAAATTGGTATAAGATGACGCGCTATGCTTTGCTGGTGAGAGGTATCAATGTTGGTGGTAAGAAT
NZ_CP035255.1:776866-784718    1872   CTATGATTGGGGAAAGTTTCTTAGATAAATTGGTATAAGATGACGCGCTATGCTTTGCTGGTGAGAGGTATCAATGTTGGTGGTAAGAAT
NZ_CP035256.1:807236-815102    1886   CTATGATTGGGGAAAGTTTCTTAGATAAATTGGTATAAGATGACGCGCTATGCTTTGCTGGTGAGAGGTATCAATGTTGGTGGTAAGAAT
NZ_CP038252.1:1151812-1159673  1816   CTATGATTGGGGAAAGTTTCTTAGATAAATTGGTATAAGATGACGCGCTATGCTTTGCTGGTGAGAGGTATCAATGTTGGTGGTAAGAAT
NZ_CP035254.1:793365-801213    1872   CTATGATTGGGGAAAGTTTCTTAGATAAATTGGTATAAGATGACGCGCTATGCTTTGCTGGTGAGAGGTATCAATGTTGGTGGTAAGAAT
NZ_CP035251.1:748733-756585    1872   CTATGATTGGGGAAAGTTTCTTAGATAAATTGGTATAAGATGACGCGCTATGCTTTGCTGGTGAGAGGTATCAATGTTGGTAGTAAGAAT
NZ_CP035252.1:745670-753536    1886   CTATGATTGGGGAAAGTTTCTTAGATAAATTGGTATAAGATGACGCGCTATGCTTTGCTGGTGAGAGGTATCAATGTTGGTAGTAAGAAT
NZ_CP035257.1:776317-784169    1872   CTATGATTGGGGAAAGTTTCTTAGATAAATTGGTATAAGATGACGCGCTATGCTTTGCTGGTGAGAGGTATCAATGTTGGTAGTAAGAAT
NZ_CP038253.1:758105-765957    1872   CTATGATTGGGGAAAGTTTCTTAGATAAATTGGTATAAGATGACGCGCTATGCTTTGCTGGTGAGAGGTATCAATGTTGGTAGTAAGAAT
NZ_CP035263.1:743050-750901    1872   CTATGATTGGGGAAAGTTTCTTAGATAAATTGGTATAAGATGACGCGCTATGCTTTGCTGGTGAGAGGTATCAATGTTGGTAGTAAGAAT
NZ_CP035262.1:741595-749460    1886   CTATGATTGGGGAAAGTTTCTTAGATAAATTGGTATAAGATGACGCGCTATGCTTTGCTGGTGAGAGGTATCAATGTTGGTAGTAAGAAT
NZ_CP035236.1:764915-772766    1872   CTATGATTGGGGAAAGTTTCTTAGATAAATTGGTATAAGATGACGCGCTATGCTTTGCTGGTGAGAGGTATCAATGTTGGTAGTAAGAAT
NZ_CP035253.1 767286-775134    1872   CTATGATTGGGGAAAGTTTCTTAGATAAATTGGTATAAGATGACGCGCTATGCTTTGCTGGTGAGAGGTATCAATGTTGGTGGTAAGAAT
NZ_CP035249.1:763080-770921    1872   CTATGATTGGGGAAAGTTTCTTAGATAAATTGGTATAAGATGACGCGCTATGCTTTGCTGGTGAGAGGTATCAATGTTGGTGGTAAGAAT

NC_003028.3:777000-787000      2691   AAGGTCGTCATGGCGGAGCTTCGTCAAGAATTGACAAACTTGGGACTGGAAAAGGTTGAGAGCTACATCAATAGTGGCAATATTTTCTTT
NZ_CP035248.1:765099-775111    2701   AAGGTTGTCATGGCGCAACTTCGTCAAGAACTGACAGAGTTGGGACTGGGAAAGGTTGAGACCTACATCAACAGTGGCAATATTTTCTTT
NZ_CP035235.1 c1326834-1317894 1976   AAGGTCGTCATGGCGGAGCTTCGTCAAGAATTGACAAACTTGGGACTGGAAAAGGTTGAGAGCTACATCAACAGTGGCAATATTTTCTTT
NZ_CP035234.1:756284-765224    1976   AAGGTCGTCATGGCGGAGCTTCGTCAAGAATTGACAAACTTGGGACTGGAAAAGGTTGAGAGCTACATCAACAGTGGCAATATTTTCTTT
NZ_CP038251.1:750745-759686    1976   AAGGTCGTCATGGCGGAGCTTCGTCAAGAATTGACAAACTTGGGACTGGAAAAGGTTGAGAGCTACATCAACAGTGGCAATATTTTCTTT
NZ_CP035244.1:747023-755950    1962   AAGGTCGTCATGGCGGAGCTTCGTCAAGAATTGACAAACTTGGGACTGGAAAAGGTTGAGAGCTACATCAACAGTGGCAATATTTTCTTT
NZ_CP035264.1:800836-809336    1962   AAGGTCGTCATGGCGGAGCTTCGTCAAGAATTGACAAACTTGGGACTGGAAAAGTTTGAGAGCTACATCAACAGTGGCAATATTTTCTTT
NZ_CP035265.1:766128-774627    1962   AAGGTCGTCATGGCGGAGCTTCGTCAAGAATTGACAAACTTGGGACTGGAAAAGTTTGAGAGCTACATCAACAGTGGCAATATTTTCTTT
NZ_CP035245.1 745261-753757    1962   AAGGTCGTCATGGCGGAGCTTCGTCAAGAATTGACAAACTTGGGACTGGAAAAGGTTGAGAGCTACATCAATAGTGGCAATATTTTCTTT
NZ_CP035259.1:745700-753548    1962   AAGGTCGTCATGGCGGAGCTTCGTCAAGAATTGACAAACTTGGGACTGGAAAAGGTTGAGAGCTACATCAACAGTGGCAATATTTTCTTT
NZ_CP035258.1:750874-758722    1962   AAGGTCGTCATGGCGGAGCTTCGTCAAGAATTGACAAACTTGGGACTGGAAAAGGTTGAGAGCTACATCAACAGTGGCAATATTTTCTTT
NZ_CP035260.1:726156-733994    1962   AAGGTCGTCATGGCGGAGCTTCATCAAGAATTGACAAACTTGGGACTGGAAAAGGTTGAGAGCTACATCAACAGTGGCAATATTTTCTTT
NZ_CP035241.1:708296-716144    1962   AAGGTCGTCATGGCGGAGCTTCATCAAGAATTGACAAACTTGGGACTGGAAAAGGTTGAGAGCTACATCAACAGTGGCAATATTTTCTTT
NZ_CP035261.1:779086-786938    1962   AAGGTCGTCATGGCGGAGCTTCGTCAAGAATTGACAAACTTGGGACTGGAAAAGGTTGAGAGCTACATCAATAGTGGCAATATTTTCTTT
NZ_CP035242.1:c1532236-1524385 1962   AAGGTCGTCATGGCGGAGCTTCGTCAAGAATTGACAAACTTGGGACTGGAAAAGGTTGAGAGCTACATCAATAGTGGCAATATTTTCTTT
NZ_CP035243.1:777729-785580    1962   AAGGTCGTCATGGCGGAGCTTCGTCAAGAATTGACAAACTTGGGACTGGAAAAGGTTGAGAGCTACATCAATAGTGGCAATATTTTCTTT
NZ_CP035237.1:c1311322-1303456 1976   AAGGTCGTCATGGCGGAGCTTCGTCAAGAATTGACAAACTTGGGACTGGAAAAGGTTGAGAGCTACATCAACAGTGGCAATATTTTCTTT
NZ_CP035246.1:c1302892-1295026 1976   AAGGTCGTCATGGCGGAGCTTCGTCAAGAATTGACAAACTTGGGACTGGAAAAGGTTGAGAGCTACATCAATAGTGGCAATATTTTCTTT
NZ_CP035247.1:c1301115-1293263 1962   AAGGTCGTCATGGCGGAGCTTCGTCAAGAATTGACAAACTTGGGACTGGAAAAGGTTGAGAGCTACATCAATAGTGGCAATATTTTCTTT
NZ_CP035238.1:738547-746413    1976   AAGGTCGTCATGGCGGAACTTCGTCAAGAATTGACAAACTTGGGACTGGAAAAGGTTGAGAGCTACATCAACAGTGGCAATATTTTCTTT
NZ_CP035255.1:776866-784718    1962   AAGGTCGTCATGGCGGAGCTTCGTCAAGAATTGACAAACTTGGGACTGGAAAAGGTTGAGAGCTACATCAACAGTGGCAATATTTTCTTT
NZ_CP035256.1:807236-815102    1976   AAGGTCGTCATGGCGGAGCTTCGTCAAGAATTGACAAACTTGGGACTGGAAAAGGTTGAGAGCTACATCAACAGTGGCAATATTTTCTTT
NZ_CP038252.1:1151812-1159673  1906   AAGGTCGTCATGGCGGAGCTTCGTCAAGAATTGACAAACTTGGGACTGGAAAAGGTTGAGAGCTACATCAATAGTGGCAATATTTTCTTT
NZ_CP035254.1:793365-801213    1962   AAGGTCGTCATGGCGGAGCTTCGTCAAGAATTGACAAACTTGGGACTGGAAAAGGTTGAGAGCTACATCAACAGTGGCAATATTTTCTTT
NZ_CP035251.1:748733-756585    1962   AAGGTCGTTATGGCGGAGCTTCGTCAAGAATTGACAAACTTGGGACTGGAAAAGGTTGAGAGCTACATCAACAGTGGCAATATTTTCTTT
```

### Ozkan et al. Supplementary Figures

```
NZ_CP035252.1:745670-753536    1976    AAGGTCGTTATGGCGGAGCTTCGTCAAGAATTGACAAACTTGGGACTGGAAAAGGTTGAGAGCTACATCAACAGTGGCAATATTTTCTTT
NZ_CP035257.1:776317-784169    1962    AAGGTCGTTATGGCGGAGCTTCGTCAAGAATTGACAAACTTTGGACTGGAAAAGGTTGAGAGCTACATCAACAGTGGCAATATTTTCTTT
NZ_CP038253.1:758105-765957    1962    AAGGTCGTTATGGCGGAGCTTCGTCAAGAATTGACAAACTTGGGACTGGAAAAGGTTGAGAGCTACATCAACAGTGGCAATATTTTCTTT
NZ_CP035263.1:743050-750901    1962    AAGGTCGTTATGGCGGAGCTTCGTCAAGAATTGACAAACTTGGGACTGGAAAAGGTTGAGAGCTACATCAACAGTGGCAATATTTTCTTT
NZ_CP035262.1:741595-749460    1976    AAGGTCGTTATGGCGGAGCTTCGTCAAGAATTGACAAACTTGGGACTGGAAAAGGTTGAGAGCTACATCAACAGTGGCAATATTTTCTTT
NZ_CP035236.1:764915-772766    1962    AAGGTCGTTATGGCGGAGCTTCGTCAAGAATTGACAAACTTGGGACTGGAAAAGGTTGAGAGCTACATCAACAGTGGCAATATTTTCTTT
NZ_CP035253.1 767286-775134    1962    AAGGTCGTCATGGCGGAGCTTCGTCAAGAATTGACAAACTTGGGACTGGAAAAGGTTGAGAGCTACATCAACAGTGGCAATATTTTCTTT
NZ_CP035249.1:763080-770921    1962    AAGGTCGTCATGGCGGAGCTTCGTCAAGAATTGACAAACTTGGGACTGGAAAAGGTTGAGAGCTACATCAATAGTGGCAATATTTTCTTT

NC_003028.3:777000-787000      2781    ACTTCGATAGATTCCAAAGCCCAATTGGTTGAAAAGCTAGAGACTTTCTTTGCAGTCCATTATCCATTATTCAGAGCTTTTCTTTACTG
NZ_CP035248.1:765099-775111    2791    ACTTCGACAGATCCCAAAGCCCAATTGGTTGAAAAGTTAGAGTCTTTCTTTGAAATCCATTATCCATTATTCAGAGCTTTTCGTTGCTG
NZ_CP035235.1 c1326834-1317894 2066    ACTTCGATAGATTCCAAAGCCCAATTGGTTGAAAAGCTAGAGACTTTCTTTGCAGTCCATTATCCATTATTCAGAGCTTTTCTTTACTG
NZ_CP035234.1:756284-765224    2066    ACTTCGATAGATTCCAAAGCCCAATTGGTTGAAAAGCTAGAGACTTTCTTTGCAGTCCATTATCCATTATTCAGAGCTTTTCTTTACTG
NZ_CP038251.1:750745-759686    2066    ACTTCGATAGATTCCAAAGCCCAATTGGTTGAAAAGCTAGAGACTTTCTTTGCAGTCCATTATCCATTATTCAGAGCTTTTCTTTACTG
NZ_CP035244.1:747023-755950    2052    ACTTCGATAGATTCCAAAGCCCAATTGGTTGAAAAGCTAGAGACTTTCTTTGCAGTCCATTATCCATTATTCAGAGCTTTTCTTTACTG
NZ_CP035264.1:800836-809336    2052    ACTTCGATAGATTCCAAAGCCCAATTGGTTGAAAAGCTAGAGACTTTCTTTGCAGTCCATTATCCATTATTCAGAGCTTTTCTTTACTG
NZ_CP035265.1:766128-774627    2052    ACTTCGATAGATTCCAAAGCCCAATTGGTTGAAAAGCTAGAGACTTTCTTTGCAGTCCATTATCCATTATTCAGAGCTTTTCTTTACTG
NZ_CP035245.1 745261-753757    2052    ACTTCGATAGATTCCAAAGCCCAATTGGTTGAAAAGCTAGAGACTTTCTTTGCAGTCCATTATCCATTATTCAGAGCTTTTCTTTACTG
NZ_CP035259.1:745700-753548    2052    GCTTCGATAGATTCCAAAGCCCAATTGGTTGAAAAGCTAGAGACTTTCTTTGCAGTCCATTATCCATTATTCAGAGCTTTTCTTTACTG
NZ_CP035258.1:750874-758722    2052    GCTTCGATAGATTCCAAAGCCCAATTGGTTGAAAAGCTAGAGACTTTCTTTGCAGTCCATTATCCATTATTCAGAGCTTTTCTTTACTG
NZ_CP035260.1:726156-733994    2052    GCTTCGATAGATTCCAAAGCCCAATTGGTTGAAAAGCTAGAGACTTTCTTTGCAGTCCATTATCCATTATTCAGAGCTTTTCTTTACTG
NZ_CP035241.1:708296-716144    2052    GCTTCGATAGATTCCAAAGCCCAATTGGTTGAAAAGCTAGAGACTTTCTTTGCAGTCCATTATCCATTATTCAGAGCTTTTCTTTACTG
NZ_CP035261.1:779086-786938    2052    ACTTCGATAGATTCCAAAGCCCAATTGGTTGAAAAGCTAGAGACTTTCTTTGCAGTCCATTATCCATTATTCAGAGCTTTTCTTTACTG
NZ_CP035242.1:c1532236-1524385 2052    ACTTCGATAGATTCCAAAGCCCAATTGGTTGAAAAGCTAGAGACTTTCTTTGCAGTCCATTATCCATTATTCAGAGCTTTTCTTTACTG
NZ_CP035243.1:777729-785580    2052    ACTTCGATAGATTCCAAAGCCCAATTGGTTGAAAAGCTAGAGACTTTCTTTGCAGTCCATTATCCATTATTCAGAGCTTTTCTTTACTG
NZ_CP035237.1:c1311322-1303456 2066    ACTTCGATAGATTCCAAAGCCCAATTGGTTGAAAAGCTAGGGACTTTCTTTGCAGTCCATTATCCATTATTCAGAGCTTTTCTTTACTG
NZ_CP035246.1:c1302892-1295026 2066    ACTTCGATAGATTCCAAAGCCCAATTGGTTGAAAAGCTAGAGACTTTCTTTGCAGTCCATTATCCATTATTCAGAGCTTTTCTTTACTG
NZ_CP035247.1:c1301115-1293263 2052    ACTTCGATAGATTCCAAAGCCCAATTGGTTGAAAAGCTAGAGACTTTCTTTGCAGTCCATTATCCATTATTCAGAGCTTTTCTTTACTG
NZ_CP035238.1:738547-746413    2066    ACTTCGATAGATTCCAAAGCCCAATTGGTTGAAAAGCTAGAGACTTTCTTTGCAGTCCATTATCCATTATTCAGAGCTTTTCTTTACTG
NZ_CP035255.1:776866-784718    2052    ACTTCGATAGATTCCAAAGCCCAATTGGTTGAAAAGCTAGAGACTTTCTTTGCAGTCCATTATCCATTATTCAGAGCTTTTCTTTACTG
NZ_CP035256.1:807236-815102    2066    ACTTCGATAGATTCCAAAGCCCAATTGGTTGAAAAGCTAGAGACTTTCTTTGCAGTCCATTATCCATTATTCAGAGCTTTTCTTTACTG
NZ_CP038252.1:1151812-1159673  1996    ACTTCGATAGATTCCAAAGCCCAATTGGTTGAAAAGCTAGAGACTTTCTTTGCAGTCCATTATCCATTATTCAGAGCTTTTCTTTACTG
NZ_CP035254.1:793365-801213    2052    ACTTCGATAGATTCCAAAGCCCAATTGGTTGAAAAGCTAGAGACTTTCTTTGCAGTCCATTATCCATTATTCAGAGCTTTTCTTTACTG
NZ_CP035251.1:748733-756585    2052    ACTTCGATAGATTCCAAAGCCCAATTGGTTTAAAAGCTAGAGACTTTCTTTGCAGTCCATTATCCATTATTCAGAGCTTTTCTTTACTG
NZ_CP035252.1:745670-753536    2066    ACTTCGATAGATTCCAAAGCCCAATTGGTTTAAAAGCTAGAGACTTTCTTTGCAGTCCATTATCCATTATTCAGAGCTTTTCTTTACTG
NZ_CP035257.1:776317-784169    2052    ACTTCGATAGATTCCAAAGCCCAATTGGTTTAAAAGCTAGAGACTTTCTTTGCAGTCCATTATCCATTATTCAGAGCTTTTCTTTACTG
NZ_CP038253.1:758105-765957    2052    ACTTCGATAGATTCCAAAGCCCAATTGGTTTAAAAGCTAGAGACTTTCTTTGCAGTCCATTATCCATTATTCAGAGCTTTTCTTTACTG
NZ_CP035263.1:743050-750901    2052    ACTTCGATAGATTCCAAAGCCCAATTGGTTTAAAAGCTAGAGACTTTCTTTGCAGTCCATTATCCATTATTCAGAGCTTTTCTTTACTG
NZ_CP035262.1:741595-749460    2066    ACTTCGATAGATTCCAAAGCCCAATTGGTTTAAAAGCTAGAGACTTTCTTTGCAGTCCATTATCCATTATTCAGAGCTTTTCTTTACTG
NZ_CP035236.1:764915-772766    2052    ACTTCGATAGATTCCAAAGCCCAATTGGTTTAAAAGCTAGAGACTTTCTTTGCAGTCCATTATCCATTATTCAGAGCTTTTCTTTACTG
NZ_CP035253.1 767286-775134    2052    ACTTCGATAGATTCCAAAGCCCAATTGGTTGAAAAGCTAGAGACTTTCTTTGCAGTCCATTATCCATTATTCAGAGCTTTTCTTTACTG
NZ_CP035249.1:763080-770921    2052    ACTTCGATAGATTCCAAAGCCCAATTGGTTGAAAAGCTAGAGACTTTCTTTGCAGTCCATTATCCATTATTCAGAGCTTTTCTTTACTG

NC_003028.3:777000-787000      2871    AGTCTAGAGGACTTTGAGGCGGAACTTGAAAATCTACCAGCTTGGTGGAGCAGAGACTTGGCACGAAAAGATTTTCTCTTTTACACTGAG
NZ_CP035248.1:765099-775111    2881    AGTCATGAAGACTATGAAGCGGAACTTGAAAATCTACCTAAATGGTGGACCAAAGATTTGGCACGAAAGGATGTCCTCTTTTACACGGAG
NZ_CP035235.1 c1326834-1317894 2156    AGCCTAGAGGACTTTGAGGCGGAACTTGAAAATCTACCAGCTTGGTGGAGCAGAGACTTGGCACGAAAAGATTTTCTCTTTTACACTGAG
NZ_CP035234.1:756284-765224    2156    AGCCTAGAGGACTTTGAGGCGGAACTTGAAAATCTACCAGCTTGGTGGAGCAGAGACTTGGCACGAAAAGATTTTCTCTTTTACACTGAG
NZ_CP038251.1:750745-759686    2156    AGCCTAGAGGACTTTGAGGCGGAACTTGAAAATCTACCAGCTTGGTGGAGCAGAGACTTGGCACGAAAAGATTTTCTCTTTTACACTGAG
NZ_CP035244.1:747023-755950    2142    AGCCTAGAGGACTTTGAGGCGGAACTTGAAAATCTACCAGCTTGGTGGAGCAGAGACTTGGCACGAAAAGATTTTCTCTTTTACACTGAG
NZ_CP035264.1:800836-809336    2142    AGTCTAGAGGACTTTGAGGCGGAACTTGAAAATCTACCAGCTTGGTGGAGCAGAGACTTGGCACGAAAAGATTTTCTCTTTTACACTGAG
NZ_CP035265.1:766128-774627    2142    AGTCTAGAGGACTTTGAGGCGGAACTTGAAAATCTACCAGCTTGGTGGAGCAGAGACTTGGCACGAAAAGATTTTCTCTTTTACACTGAG
NZ_CP035245.1 745261-753757    2142    AGTCTAGAGGACTTTGAGGCGGAACTTGAAAATCTACCAGCTTGGTGGAGCAGAGACTTGGCACGAAAAGATTTTCTCTTTTACACTGAG
NZ_CP035259.1:745700-753548    2142    AGTCTAGAGGACTTTGAGGCGGAACTTGAAAATCTACCAGCTTGGTGGAGCAGAGACTTGGCACGAAAAGATTTTCTCTTTTACACTGAG
NZ_CP035258.1:750874-758722    2142    AGTCTAGAGGACTTTGAGGCGGAACTTGAAAATCTACCAGCTTGGTGGAGCAGAGACTTGGCACGAAAAGATTTTCTCTTTTACACTGAG
NZ_CP035260.1:726156-733994    2142    AGTCTAGAGGACTTTGAGGCGGAACTTGAAAATCTACCAGCTTGGTGGAGCAGAGACTTGGCACGAAAAGATTTTCTCTTTTACACTGAG
NZ_CP035241.1:708296-716144    2142    AGTCTAGAGGACTTTGAGGCGGAACTTGAAAATCTACCAGCTTGGTGGAGCAGAGACTTGGCACGAAAAGATTTTCTCTTTTACACTGAG
NZ_CP035261.1:779086-786938    2142    AGTCTAGTGGACTTTGAGGCGGAACTTGAAAATCTACCAGCTTGGTGGAGCAGAGACTTGGCACGAAAAGATTTTCTCTTTTACACTGAG
NZ_CP035242.1:c1532236-1524385 2142    AGTCTAGAGGACTTTGAGGCGGAACTTGAAAATCTACCAGCTTGGTGGAGCAGAGACTTGGCACGAAAAGATTTTCTCTTTTACACTGAG
NZ_CP035243.1:777729-785580    2142    AGTCTAGAGGACTTTGAGGCGGAACTTGAAAATCTACCAGCTTGGTGGAGCAGAGACTTGGCACGAAAAGATTTTCTCTTTTACACTGAG
NZ_CP035237.1:c1311322-1303456 2156    AGTCTAGAGGACTTTGAGGCGGAACTTGAAAATCTACCAGCTTGGTGGAGCAGAGACTTGGCACGAAAAGATTTTCTCTTTTACACTGAG
NZ_CP035246.1:c1302892-1295026 2156    AGTCTAGAGGACTTTGAGGCGGAACTTGAAAATCTACCAGCTTGGTGGAGCAGAGACTTGGCACGAAAAGATTTTCTCTTTTACACTGAG
NZ_CP035247.1:c1301115-1293263 2142    AGTCTAGAGGACTTTGAGGCGGAACTTGAAAATCTACCAGCTTGGTGGAGCAGAGACTTGGCACGAAAAGATTTTCTCTTTTACACTGAG
NZ_CP035238.1:738547-746413    2156    AGTCTAGAGGACTTTGAGGCGGAACTTGAAAATCTACCAGCTTGGTGGAGCAGAGACTTGGCACGAAAAGATTTTCTCTTTTACACTGAG
NZ_CP035255.1:776866-784718    2142    AGTCTAGAGGACTTTGAGGCGGAACTTGAAAATCTACCAGCTTGGTGGAGCAGAGACTTGGCACGAAAAGATTTTCTCTTTTACACTGAG
NZ_CP035256.1:807236-815102    2156    AGTCTAGAGGACTTTGAGGCGGAACTTGAAAATCTACCAGCTTGGTGGAGCAGAGACTTGGCACGAAAAGATTTTCTCTTTTACACTGAG
NZ_CP038252.1:1151812-1159673  2086    AGTCTAGAGGACTTTGAGGCGGAACTTGAAAATCTACCAGCTTGGTGGAGCAGAGACTTGGCACGAAAAGATTTTCTCTTTTACACTGAG
NZ_CP035254.1:793365-801213    2142    AGTCTAGAGGACTTTGAGGCGGAACTTGAAAATCTACCAGCTTGGTGGAGCAGAGACTTGGCACGAAAAGATTTTCTCTTTTACACTGAG
NZ_CP035251.1:748733-756585    2142    AGTCTAGAGGACTTTGAGGCGGAACTTGAAAATCTACCAGCTTGGTGGAGCAGAGACTTGGCACGAAAAGATTTTCTCTTTTACACTGAG
NZ_CP035252.1:745670-753536    2156    AGTCTAGAGGACTTTGAGGCGGAACTTGAAAATCTACCAGCTTGGTGGAGCAGAGACTTGGCACGAAAAGATTTTCTCTTTTACACTGAG
NZ_CP035257.1:776317-784169    2142    AGTCTAGAGGACTTTGAGGCGGAACTTGAAAATCTACCAGCTTGGTGGAGCAGAGACTTGGCACGAAAAGATTTTCTCTTTTACACTGAG
NZ_CP038253.1:758105-765957    2142    AGTCTAGAGGACTTTGAGGCGGAACTTGAAAATCTACCAGCTTGGTGGAGCAGAGACTTGGCACGAAAAGATTTTCTCTTTTACACTGAG
NZ_CP035263.1:743050-750901    2142    AGTCTAGAGGACTTTGAGGCGGAACTTGAAAATCTACCAGCTTGGTGGAGCAGAGACTTGGCACGAAAAGATTTTCTCTTTTACACTGAG
NZ_CP035262.1:741595-749460    2156    AGTCTAGAGGACTTTGAGGCGGAACTTGAAAATCTACCAGCTTGGTGGAGCAGAGACTTGGCACGAAAAGATTTTCTCTTTTACACTGAG
NZ_CP035236.1:764915-772766    2142    AGTCTAGAGGACTTTGAGGCGGAACTTGAAAATCTACCAGCTTGGTGGAGCAGAGACTTGGCACGAAAAGATTTTCTCTTTTACACTGAG
NZ_CP035253.1 767286-775134    2142    AGTCTAGAGGACTTTGAGGCGGAACTTGAAAATCTACCAGCTTGGTGGAGCAGAGACTTGGCACGAAAAGATTTTCTCTTTTACACTGAG
NZ_CP035249.1:763080-770921    2142    AGTCTAGAGGACTTTGAGGCGGAACTTGAAAATCTACCAGCTTGGTGGAGCAGAGACTTGGCACGAAAAGATTTTCTCTTTTACACTGAG

NC_003028.3:777000-787000      2961    GGTTTGGATGTGGACCAAGTCATCGCGACAGTTGAAAGTTTAGAGCTGAAAGATGAAGTGCTTTATTTTGGAAAACTTGGGATTTTCTGG
NZ_CP035248.1:765099-775111    2971    AGCTTGGATGTGGATCAAGTGATCGAGAAAGTGAACAGTTTGGAATTGGAAGATGAAGTGGTTCATTTTGGAAGACTTGGGATTTTCTGG
NZ_CP035235.1 c1326834-1317894 2246    GGTTTGGATGTGGACCAAGTCATCGCGACAGTTGAAAGTTTAGAGCTGAAAGATGAAGTGCTTTATTTTGGAAAACTTGGGATTTTCTGG
NZ_CP035234.1:756284-765224    2246    GGTTTGGATGTGGACCAAGTCATCGCGACAGTTGAAAGTTTAGAGCTGAAAGATGAAGTGCTTTATTTTGGAAAACTTGGGATTTTCTGG
NZ_CP038251.1:750745-759686    2246    GGTTTGGATGTGGACCAAGTCATCGCGACAGTTGAAAGTTTAGAGCTGAAAGATGAAGTGCTTTATTTTGGAAAACTTGGGATTTTCTGG
NZ_CP035244.1:747023-755950    2232    GGTTTGGATGTGGACCAAGTCATCGCGACAGTTGAAAGTTTAGAGCTGAAAGATGAAGTGCTTTATTTTGGAAAACTTGGGATTTTCTGG
NZ_CP035264.1:800836-809336    2232    GGTTTGGATGTGGACCAAGTCATCGCGACAGTTGAAAGTTTAGAGCTGAAAGATGAAGTGCTTTATTTTGGAAAACTTGGGATTTTCTGG
NZ_CP035265.1:766128-774627    2232    GGTTTGGATGTGGACCAAGTCATCGCGACAGTTGAAAGTTTAGAGCTGAAAGATGAAGTGCTTTATTTTGGAAAACTTGGGATTTTCTGG
NZ_CP035245.1 745261-753757    2232    GGTTTGGATGTGGACCAAGTCATCGCGACAGTTGAAAGTTTAGAGCTGAAAGATGAAGTGCTTTATTTTGGAAAACTTGGGATTTTCTGG
NZ_CP035259.1:745700-753548    2232    GGTTTGGATGTGGACCAAGTCATCGCGACAGTTGAAAGTTTAGAGCTGAAAGATGAAGTGCTTTATTTTGGAAAACTTGGGATTTTCTGG
NZ_CP035258.1:750874-758722    2232    GGTTTGGATGTGGACCAAGTCATCGCGACAGTTGAAAGTTTAGAGCTGAAAGATGAAGTGCTTTATTTTGGAAAACTTGGGATTTTCTGG
NZ_CP035260.1:726156-733994    2232    GGTTTGGATGTGGACCAAGTCATCGCGACAGTTGAAAGTTTAGAGCTGAAAGATGAAGTGCTTTATTTTGGAAAACTTGGGATTTTCTGG
NZ_CP035241.1:708296-716144    2232    GGTTTGGATGTGGACCAAGTCATCGCGACAGTTGAAAGTTTAGAGCTGAAAGATGAAGTGCTTTATTTTGGAAAACTTGGGATTTTCTGG
NZ_CP035261.1:779086-786938    2232    GGTTTGGATGTGGACCAAGTCATCGCGACAGTTGAAAGTTTAGAGCTGAAAGATGAAGTGCTTTATTTTGGAAAACTTGGGATTTTCTGG
NZ_CP035242.1:c1532236-1524385 2232    GGTTTGGATGTGGACCAAGTCATCGCGACAGTTGAAAGTTTAGAGCTGAAAGATGAAGTGCTTTATTTTGGAAAACTTGGGATTTTCTGG
NZ_CP035243.1:777729-785580    2232    GGTTTGGATGTGGACCAAGTCATCGCGACAGTTGAAAGTTTAGAGCTGAAAGATGAAGTGCTTTATTTTGGAAAACTTGGGATTTTCTGG
NZ_CP035237.1:c1311322-1303456 2246    GGTTTGGATGTGGACCAAGTCATCGCGACAGTTGAAAGTTTAGAGCTGAAAGATGAAGTGCTTTATTTTGGAAAACTTGGGATTTTCTGG
NZ_CP035246.1:c1302892-1295026 2246    GGTTTGGATGTGGACCAAGTCATCGCGACAGTTGAAAGTTTAGAGCTGAAAGATGAAGTGCTTTATTTTGGAAAACTTGGGATTTTCTGG
```

### Ozkan et al. Supplementary Figures

```
NZ_CP035247.1:c1301115-1293263 2232   GGTTTGGATGTGGACCAAGTCATCGCGACAGTTGAAAGTTTAGAGCTGAAAGATGAAGTGCTTTATTTTGGAAAACTTGGGATTTTCTGG
NZ_CP035238.1:738547-746413    2246   GGTTTGGATGTGGACCAAGTCATCGCGACAGTTGAAAGTTTAGAGCTGAAAGATGAAGTGCTTTATTTTGGAAAACTTGGGATTTTCTAG
NZ_CP035255.1:776866-784718    2232   GGTTTGGATGTGGACCAAGTCATCGCGACAGTTGAAAGTTTAGAGCTGAAAGATGAAGTGCTTTATTTTGGAAAACTTGGGATTTTCTGG
NZ_CP035256.1:807236-815102    2246   GGTTTGGATGTGGACCAAGTCATCGCGACAGTTGAAAGTTTAGAGCTGAAAGATGAAGTGCTTTATTTTGGAAAACTTGGGATTTTCTGG
NZ_CP038252.1:1151812-1159673  2176   GGTTTGGATGTGGACCAAGTCATCGCGACAGTTGAAAGTTTAGAGCTGAAAGATGAAGTGCTTTATTTTGGAAAACTTGGGATTTTCTGG
NZ_CP035254.1:793365-801213    2232   GGTTTGGATGTGGACCAAGTCATCGCGACAGTTGAAAGTTTAGAGCTGAAAGATGAAGTGCTTTATTTTGGAAAACTTGGGATTTTCTGG
NZ_CP035251.1:748733-756585    2232   GGTTTGGATGTGGACCAAGTCATCGCGACAGTTGAAAGTTTAGAGCTGAAAGATGAAGTGCTTTATTTTAGAAAACTTGGGATTTTCTGG
NZ_CP035252.1:745670-753536    2246   GGTTTGGATGTGGACCAAGTCATCGCGACAGTTGAAAGTTTAGAGCTGAAAGATGAAGTGCTTTATTTTAGAAAACTTGGGATTTTCTGG
NZ_CP035257.1:776317-784169    2232   GGTTTGGATGTGGACCAAGTCATCGCGACAGTTGAAAGTTTAGAGCTGAAAGATGAAGTGCTTTATTTTAGAAAACTTGGGATTTTCTGG
NZ_CP038253.1:758105-765957    2232   GGTTTGGATGTGGACCAAGTCATCGCGACAGTTGAAAGTTTAGAGCTGAAAGATGAAGTGCTTTATTTTAGAAAACTTGGGATTTTCTGG
NZ_CP035263.1:743050-750901    2232   GGTTTGGATGTGGACCAAGTCATCGCGACAGTTGAAAGTTTAGAGCTGAAAGATGAAGTGCTTTATTTTAGAAAACTTGGGATTTTCTGG
NZ_CP035262.1:741595-749460    2246   GGTTTGGATGTGGACCAAGTCATCGCGACAGTTGAAAGTTTAGAGCTGAAAGATGAAGTGCTTTATTTTAGAAAACTTGGGATTTTCTGG
NZ_CP035236.1:764915-772766    2232   GGTTTGGATGTGGACCAAGTCATCGCGACAGTTGAAAGTTTAGAGCTGAAAGATGAAGTGCTTTATTTTAGAAAACTTGGGATTTTCTGG
NZ_CP035253.1 767286-775134    2232   GGTTTGGATGTGGACCAAGTCATCGCGACAGTTGAAAGTTTAGAGCTGAAAGATGAAGTGCTTTATTTTGGAAAACTTGGGATTTTCTGG
NZ_CP035249.1:763080-770921    2232   GGTTTGGATGTGGACCAAGTCATCGCGACAGTTGAAAGTTTAGAGCTGAAAGATGAAGTGCTTTATTTTGGAAAACTTGGGATTTTCTGG

NC_003028.3:777000-787000      3051   GGGAAATTTTCTGAAGAATCCTATTCTAAGACTGCCTATCATAAGTACTTGCTGAAGGTGCCTTTCTACCGCCACATTACTATTCGTAAT
NZ_CP035248.1:765099-775111    3061   GGGAAATTCTCTGAGGAATCCTACTATGCAACTGCCTATCATAAGTACTTGCTCAAGATGCCTTTTTATCGCAATATCACCATTCGCAAT
NZ_CP035235.1 c1326834-1317894 2336   GGGAAATTTTCTGAAGAATCCTATTCTAAGACTGCCTATCATAAGTACTTGCTGAAGGTGCCTTTCTACCGCCACATTACTATTCGTAAT
NZ_CP035234.1:756284-765224    2336   GGGAAATTTTCTGAAGAATCCTATTCTAAGACTGCCTATCATAAGTACTTGCTGAAGGTGCCTTTCTACCGCCACATTACTATTCGTAAT
NZ_CP038251.1:750745-759686    2336   GGGAAATTTTCTGAAGAATCCTATTCTAAGACTGCCTATCATAAGTACTTGCTGAAGGTGCCTTTCTACCGCCACATTACTATTCGTAAT
NZ_CP035244.1:747023-755950    2322   GGGAAATTTTCTGAAGAATCCTATTCTAAGACTGCCTATCATAAGTACTTGCTGAAGGTGCCTTTCTACCGCCACATTACTATTCGTAAT
NZ_CP035264.1:800836-809336    2322   GGGAAATTTTCTGAAGAATCCTATTCTAAGACTGCCTATCATAAGTACTTGCTGAAGGTGCCTTTCTACCGCCACATTACTATTCGTAAT
NZ_CP035265.1:766128-774627    2322   GGGAAATTTTCTGAAGAATCCTATTCTAAGACTGCCTATCATAAGTACTTGCTGAAGGTGCCTTTCTACCGCCACATTACTATTCGTAAT
NZ_CP035245.1 745261-753757    2322   GGGAAATTTTCTGAAGAATCCTATTCTAAGACTGCCTATCAGAAGTACTTGCTGAAGGTGCCTTTCTACCGCCACATTACTATTCGTAAT
NZ_CP035259.1:745700-753548    2322   GGGAAATTTTCTGAAGAATCCTATTCTAAGACTGCCTATCATAAGTACTTGCTGAAGGTGCCTTTCTACCGCCACATTACTATTCGTAAT
NZ_CP035258.1:750874-758722    2322   GGGAAATTTTCTGAAGAATCCTATTCTAAGACTGCCTATCATAAGTACTTGCTGAAGGTGCCTTTCTACCGCCACATTACTATTCGTAAT
NZ_CP035260.1:726156-733994    2322   GGGAAATTTTCTGAAGAATCCTATTCTAAGACTGCCTATCAGAAGTACTTGCTGAAGGTGCCTTTCTACCGCCACATTACTATTCGTAAT
NZ_CP035241.1:708296-716144    2322   GGGAAATTTTCTGAAGAATCCTATTCTAAGACTGCCTATCAGAAGTATTTGCTGAAGGTGCCTTTCTACCGCCACATTACTATTCGTAAT
NZ_CP035261.1:779086-786938    2322   GGGAAATTTTCTGAAGAATCCTATTCTAAGACTGCCTATCATAAGTACTTGCTGAAGGTGCCTTTCTACCGCCACATTACTATTCGTAAT
NZ_CP035242.1:c1532236-1524385 2322   GGGAAATTTTCTGAAGAATCCTATTCTAAGACTGCCTATCATAAGTACTTGCTGAAGGTGCCTTTCTACCGCCACATTACTATTCGTAAT
NZ_CP035243.1:777729-785580    2322   GGGAAATTTTCTGAAGAATCCTATTCTAAGACTGCCTATCATAAGTACTTGCTGAAGGTGCCTTTCTACCGCCACATTACTATTCGTAAT
NZ_CP035237.1:c1311322-1303456 2336   GGGAAATTTTCTGAAGAATCCTATTCTAAGACTGCCTATCAGAAGTACTTGCTGAAGGTGCCTTTCTACCGCCACATTACTATTCGTAAT
NZ_CP035246.1:c1302892-1295026 2336   GGGAAATTTTCTGAAGAATCCTATTCTAAGACTGCCTATCATAAGTACTTGCTGAAGGTGCCTTTCTACCGCCACATTACTATTCGTAAT
NZ_CP035247.1:c1301115-1293263 2322   GGGAAATTTTCTGAAGAATCCTATTCTAAGACTGCCTATCATAAGTACTTGCTGAAGGTGCCTTTCTACCGCCACATTACTATTCGTAAT
NZ_CP035238.1:738547-746413    2336   GGGAAATTTTCTGAAGAATCCTATTCTAAGACTGCCTATCATAAGTACTTGCTGAAGGTGCCTTTCTACCGCCACATTACTATTCGTAAT
NZ_CP035255.1:776866-784718    2322   GGGAAATTTTCTGAAGAATCCTATTCTAAGACTGCCTATCAGAAGTACTTGCTGAAGGTGCCTTTCTACCGCCACATTACTATTCGTAAT
NZ_CP035256.1:807236-815102    2336   GGGAAATTTTCTGAAGAATCCTATTCTAAGACTGCCTATCAGAAGTACTTGCTGAAGGTGCCTTTCTACCGCCACATTACTATTCGTAAT
NZ_CP038252.1:1151812-1159673  2266   GGGAAATTTTCTGAAGAATCCTATTCTAAGACTGCCTATCATAAGTACTTGCTGAAGGTGCCTTTCTACCGCCACATTACTATTCGTAAT
NZ_CP035254.1:793365-801213    2322   GGGAAATTTTCTGAAGAATCCTATTCTAAGACTGCCTATCAGAAGTACTTGCTGAAGGTGCCTTTCTACCGCCACATTACTATTCGTAAT
NZ_CP035251.1:748733-756585    2322   GGGAAATTTTCTGAAGAATCCTATTCTAAGACTGCCTATCATAAGTACTTGCTGAAGGTGCCTTTCTACCGCCACATTACTATTCGTAAT
NZ_CP035252.1:745670-753536    2336   GGGAAATTTTCTGAAGAATCCTATTCTAAGACTGCCTATCATAAGTACTTGCTGAAGGTGCCTTTCTACCGCCACATTACTATTCGTAAT
NZ_CP035257.1:776317-784169    2322   GGGAAATTTTCTGAAGAATCCTATTCTAAGACTGCCTATCATAAGTACTTGCTGAAGGTGCCTTTCTACCGCCACATTACTATTCGTAAT
NZ_CP038253.1:758105-765957    2322   GGGAAATTTTCTGAAGAATCCTATTCTAAGACTGCCTATCATAAGTACTTGCTGAAGGTGCCTTTCTACCGCCACATTACTATTCGTAAT
NZ_CP035263.1:743050-750901    2322   GGGAAATTTTCTGAAGAATCCTATTCTAAGACTGCCTATCATAAGTACTTGCTGAAGGTGCCTTTCTACCGCCACATTACTATTCGTAAT
NZ_CP035262.1:741595-749460    2336   GGGAAATTTTCTGAAGAATCCTATTCTAAGACTGCCTATCATAAGTACTTGCTGAAGGTGCCTTTCTACCGCCACATTACTATTCGTAAT
NZ_CP035236.1:764915-772766    2322   GGGAAATTTTCTGAAGAATCCTATTCTAAGACTGCCTATCATAAGTACTTGCTGAAGGTGCCTTTCTACCGCCACATTACTATTCGTAAT
NZ_CP035253.1 767286-775134    2322   GGGAAATTTTCTGAAGAATCCTATTCTAAGACTGCCTATCAGAAGTACTTGCTGAAGGTGCCTTTCTACCGCCACATTACTATTCGTAAT
NZ_CP035249.1:763080-770921    2322   GGGAAATTTTCTGAAGAATCCTATTCTAAGACTGCCTATCATAAGTACTTGCTGAAGGTGCCTTTCTACCGCCACATTACTATTCGTAAT

NC_003028.3:777000-787000      3141   GCTAAAACCTTTGACAAAATTGGTCAAATGCTAAAAAAATAATAAAGGAGACACACAATGACATTTTTAAACAAAATCCATGAAACTGCT
NZ_CP035248.1:765099-775111    3151   GCTAAAACTTTTGACAAAATCGGTCAAATGCT-AAAAACTAAAAAAGGAGACACACAATGACATTTTTAGATAAAATCAAGGAAACAGCT
NZ_CP035235.1 c1326834-1317894 2426   GCTAAAACCTTTGACAAAATTGGTCAAATGCTAAAAAAATAATAAAGGAGACACACAATGACATTTTTAAACAAAATCCATGAAACTGCT
NZ_CP035234.1:756284-765224    2426   GCTAAAACCTTTGACAAAATTGGTCAAATGCTAAAAAAATAATAAAGGAGACACACAATGACATTTTTAAACAAAATCCATGAAACTGCT
NZ_CP038251.1:750745-759686    2426   GCTAAAACCTTTGACAAAATTGGTCAAATGCTAAAAAAATAATAAAGGAGACACACAATGACATTTTTAAACAAAATCCATGAAACTGCT
NZ_CP035244.1:747023-755950    2412   GCTAAAACCTTTGACAAAATTGGTCAAATGCTAAAAAAATAATAAAGGAGACACACAATGACATTTTTAAACAAAATCCATGAAACTGCT
NZ_CP035264.1:800836-809336    2412   GCTAAAACCTTTGACAAAATTGGTCAAATGCTAAAAAAATAATAAAGGAGACACACAATGACATTTTTAAACAAAATCCATGAAACTGCT
NZ_CP035265.1:766128-774627    2412   GCTAAAACCTTTGACAAAATTGGTCAAATGCTAAAAAAATAATAAAGGAGACACACAATGACATTTTTAAACAAAATCCATGAAACTGCT
NZ_CP035245.1 745261-753757    2412   GCTAAAACCTTTGACAAAATTGGTCAAATGCTAAAAAAATAATAAAGGAGACACACAATGACATTTTTAAACAAAATCCATGAAACTGCT
NZ_CP035259.1:745700-753548    2412   GCTAAAACCTTTGACAAAATTGGTCAAATGCTAAAAAAATAATAAAGGAGACACACAATGACATTTTTAAACAAAATCCATGAAACTGCT
NZ_CP035258.1:750874-758722    2412   GCTAAAACCTTTGACAAAATTGGTCAAATGCTAAAAAAATAATAAAGGAGACACACAATGACATTTTTAAACAAAATCCATGAAACTGCT
NZ_CP035260.1:726156-733994    2412   GCTAAAACCTTTGACAAAATTGGTCAAATGCTAAAAAAATAATAAAGGAGACACACAATGACATTTTTAAACAAAATCCATGAAACTGCT
NZ_CP035241.1:708296-716144    2412   GCTAAAACCTTTGACAAAATTGGTCAAATGCTAAAAAAATAATAAAGGAGACACACAATGACATTTTTAAACAAAATCCATGAAACTGCT
NZ_CP035261.1:779086-786938    2412   GCTAAAACCTTTGACAAAATTGGTCAAATGCTAAAAAAATAATAAAGGAGACACACAATGACATTTTTAAACAAAATCCATGAAACTGCT
NZ_CP035242.1:c1532236-1524385 2412   GCTAAAACCTTTGACAAAATTGGTCAAATGCT-AAAAAATAATAAAGGAGACACACAATGACATTTTTAAACAAAATCCATGAAACTGCT
NZ_CP035243.1:777729-785580    2412   GCTAAAACCTTTGACAAAATTGGTCAAATGCT-AAAAAATAATAAAGGAGACACACAATGACATTTTTAAACAAAATCCATGAAACTGCT
NZ_CP035237.1:c1311322-1303456 2426   GCTAAAACCTTTGACAAAATTGGTCAAATGCTAAAAAAATAATAAAGGAGACACACAATGACATTTTTAAACAAAATCCATGAAACTGCT
NZ_CP035246.1:c1302892-1295026 2426   GCTAAAACCTTTGACAAAATTGGTCAAATGCTAAAAAAATAATAAAGGAGACACACAATGACATTTTTAAACAAAATCCATGAAACTGCT
NZ_CP035247.1:c1301115-1293263 2412   GCTAAAACCTTTGACAAAATTGGTCAAATGCTAAAAAAATAATAAAGGAGACACACAATGACATTTTTAAACAAAATCCATGAAACTGCT
NZ_CP035238.1:738547-746413    2426   GCTAAAACCTTTGACAAAATTGGTCAAATGCTAAAAAAATAATAAAGGAGACACACAATGACATTTTTAAACAAAATCCATGAAACTGCT
NZ_CP035255.1:776866-784718    2412   GCTAAAACCTTTGACAAAATTGGTCAAATGCTAAAAAAATAATAAAGGAGACACACAATGACATTTTTAAACAAAATCCATGAAACTGCT
NZ_CP035256.1:807236-815102    2426   GCTAAAACCTTTGACAAAATTGGTCAAATGCTAAAAAAATAATAAAGGAGACACACAATGACATTTTTAAACAAAATCCATGAAACTGCT
NZ_CP038252.1:1151812-1159673  2356   GCTAAAACCTTTGACAAAATTGGTCAAATGCT-AAAAAATAATAAAGGAGACACACAATGACATTTTTAAACAAAATCCATGAAACTGCT
NZ_CP035254.1:793365-801213    2412   GCTAAAACCTTTGACAAAATTGGTCAAATGCTAAAAAAATAATAAAGGAGACACACAATGACATTTTTAAACAAAATCCATGAAACTGCT
NZ_CP035251.1:748733-756585    2412   GCTAAAACCTTTGACAAAATTGGTCAAATGCT-AAAAAATAATAAAGGAGACACATAATGACATTTTTAAACAAAATCCATGAAACTGCT
NZ_CP035252.1:745670-753536    2426   GCTAAAACCTTTGACAAAATTGGTCAAATGCT-AAAAAATAATAAAGGAGACACATAATGACATTTTTAAACAAAATCCATGAAACTGCT
NZ_CP035257.1:776317-784169    2412   GCTAAAACCTTTGACAAAATTGGTCAAATGCT-AAAAAATAATAAAGGAGACACATAATGACATTTTTAAACAAAATCCATGAAACTGCT
NZ_CP038253.1:758105-765957    2412   GCTAAAACCTTTGACAAAATTGGTCAAATGCT-AAAAAATAATAAAGGAGACACATAATGACATTTTTAAACAAAATCCATGAAACTGCT
NZ_CP035263.1:743050-750901    2412   GCTAAAACCTTTGACAAAATTGGTCAAATGCT-AAAAAATAATAAAGGAGACACATAATGACATTTTTAAACAAAATCCATGAAACTGCT
NZ_CP035262.1:741595-749460    2426   GCTAAAACCTTTGACAAAATTGGTCAAATGCT-AAAAAATAATAAAGGAGACACATAATGACATTTTTAAACAAAATCCATGAAACTGCT
NZ_CP035236.1:764915-772766    2412   GCTAAAACCTTTGACAAAATTGGTCAAATGCT-AAAAAATAATAAAGGAGACACATAATGACATTTTTAAACAAAATCCATGAAACTGCT
NZ_CP035253.1 767286-775134    2412   GCTAAAACCTTTGACAAAATTGGTCAAATGCTAAAAAAATAATAAAGGAGACACACAATGACATTTTTAAACAAAATCCATGAAACTGCT
NZ_CP035249.1:763080-770921    2412   GCTAAAACCTTTGACAAAATTGGTCAAATGCTAAAAAAATAATAAAGGAGACACACAATGACATTTTTAAACAAAATCCATGAAACTGCT

NC_003028.3:777000-787000      3231   ACTTTCCTGAAAGAAAAGGGAATTGCAGCCCCTGAGTTCGGTCTAATCCTTGGATCAGGACTTGGAGAATTGGCAGAAGAAATCGAAAAT
NZ_CP035248.1:765099-775111    3240   ACTTTTCTGAAAGATAAAGGAATCCAAGCGCCTGAGTTCGGTCTAATCCTTGGATCAGGACTTGGAGAATTGGCAGAAGAAATCGAAAAT
NZ_CP035235.1 c1326834-1317894 2516   ACTTTCCTGAAAGAAAAGGGAATTGCAGCCCCTGAGTTCGGTCTAATCCTTGGATCAGGACTTGGAGAATTGGCAGAAGAAATCGAAAAT
NZ_CP035234.1:756284-765224    2516   ACTTTCCTGAAAGAAAAGGGAATTGCAGCCCCTGAGTTCGGTCTAATCCTTGGATCAGGACTTGGAGAATTGGCAGAAGAAATCGAAAAT
NZ_CP038251.1:750745-759686    2516   ACTTTCCTGAAAGAAAAGGGAATTGCAGCCCCTGAGTTCGGTCTAATCCTTGGATCAGGACTTGGAGAATTGGCAGAAGAAATCGAAAAT
NZ_CP035244.1:747023-755950    2502   ACTTTCCTGAAAGAAAAGGGAATTGCAGCCCCTGAGTTCGGTCTAATCCTTGGATCAGGACTTGGAGAATTGGCAGAAGAAATCGAAAAT
NZ_CP035264.1:800836-809336    2502   ACTTTCCTGAAAGAAAAGGGAATTGCAGCCCCTGAGTTCGGTCTAATCCTTGGGTCAGGACTTGGAGAATTGGCAGAAGAAATCGAAAAT
NZ_CP035265.1:766128-774627    2502   ACTTTCCTGAAAGAAAAGGGAATTGCAGCCCCTGAGTTCGGTCTAATCCTTGGGTCAGGACTTGGAGAATTGGCAGAAGAAATCGAAAAT
NZ_CP035245.1 745261-753757    2502   ACTTTCCTGAAAGAAAAGGGAATTGCAGCCCCTGAGTTCGGTCTAATCCTTGGGTCAGGACTTGGAGAATTGGCAGAAGAAATCGAAAAT
NZ_CP035259.1:745700-753548    2502   ACTTTCCTGAAAGAAAAGGGAATTGCAGCCCCTGAGTTCGGTCTAATCCTTGGGTCAGGACTTGGAGAATTGGCAGAAGAAATCGAAAAT
NZ_CP035258.1:750874-758722    2502   ACTTTCCTGAAAGAAAAGGGAATTGCAGCCCCTGAGTTCGGTCTAATCCTTGGGTCAGGACTTGGAGAATTGGCAGAAGAAATCGAAAAT
```

```
NZ_CP035260.1:726156-733994    2502   ACTTTCCTGAAAGAAAAGGGAATTGCAGCCCCTGAGTTCGGTCTAATCCTTGGGTCAGGACTTGGAGAATTGGCAGAAGAAATCGAAAAT
NZ_CP035241.1:708296-716144    2502   ACTTTCCTGAAAGAAAAGGGAATTGCAGCCCCTGAGTTCGGTCTAATCCTTGGGTCAGGACTTGGAGAATTGGCAGAAGAAATCGAAAAT
NZ_CP035261.1:779086-786938    2502   ACTTTCCTGAAAGAAAAGGGAATTGCAGCCCCTGAGTTCGGTCTAATCCTTGGATCAGGACTTGGAGAATTGGCAGAAGAAATCGAAAAT
NZ_CP035242.1:c1532236-1524385 2501   ACTTTCCTGAAAGAAAAGGGAATTGCAGCCCCTGAGTTCGGTCTAATCCTTGGATCAGGACTTGGAGAATTGGCAGAAGAAATCGAAAAT
NZ_CP035243.1:777729-785580    2501   ACTTTCCTGAAAGAAAAGGGAATTGCAGCCCCTGAGTTCGGTCTAATCCTTGGATCAGGACTTGGAGAATTGGCAGAAGAAATCGAAAAT
NZ_CP035237.1:c1311322-1303456 2516   ACTTTCCTGAAAGAAAAGGGAATTGCAGCCCCTGAGTTCGGTCTAATCCTTGGATCAGGACTTGGAGAATTGGCAGAAGAAATCGAAAAT
NZ_CP035246.1:c1302892-1295026 2516   ACTTTCCTGAAAGAAAAGGGAATTGCAGCCCCTGAGTTCGGTCTAATCCTTGGATCAGGACTTGGAGAATTGGCAGAAGAAATCGAAAAT
NZ_CP035247.1:c1301115-1293263 2502   ACTTTCCTGAAAGAAAAGGGAATTGCAGCCCCTGAGTTCGGTCTAATCCTTGGATCAGGACTTGGAGAATTGGCAGAAGAAATCGAAAAT
NZ_CP035238.1:738547-746413    2516   ACTTTCCTGAAAGAAAAGGGAATTGCAGCCCCTGAGTTCGGTCTAATCCTTGGATCAGGACTTGGAGAATTGGCAGAAGAAATCGAAAAT
NZ_CP035255.1:776866-784718    2502   ACTTTCCTGAAAGAAAAGGGAATTGCAGCCCCTGAGTTCGGTCTAATCCTTGGGTCAGGACTTGGAGAATTGGCAGAAGAAATCGAAAAT
NZ_CP035256.1:807236-815102    2516   ACTTTCCTGAAAGAAAAGGGAATTGCAGCCCCTGAGTTCGGTCTAATCCTTGGGTCAGGACTTGGAGAATTGGCAGAAGAAATCGAAAAT
NZ_CP038252.1:1151812-1159673  2445   ACTTTCCTGAAAGAAAAGGGAATTGCAGCCCCTGAGTTCGGTCTAATCCTTGGATCAGGACTTGGAGAATTGGCAGAAGAAATCGAAAAT
NZ_CP035254.1:793365-801213    2502   ACTTTCCTGAAAGAAAAGGGAATTGCAGCCCCTGAGTTCGGTCTAATCCTTGGATCAGGACTTGGAGAATTGGCAGAAGAAATCGAAAAT
NZ_CP035251.1:748733-756585    2501   ACTTTCCTGAAAGAAAAGGGAATTGCAGCCCCTGAGTTCGGTCTAATCCTTGGGTCAGGACTTGGAGAATTGGCAGAAGAAATCGAAAAT
NZ_CP035252.1:745670-753536    2515   ACTTTCCTGAAAGAAAAGGGAATTGCAGCCCCTGAGTTCGGTCTAATCCTTGGGTCAGGACTTGGAGAATTGGCAGAAGAAATCGAAAAT
NZ_CP035257.1:776317-784169    2501   ACTTTCCTGAAAGAAAAGGGAATTGCAGCCCCTGAGTTCGGTCTAATCCTTGGGTCAGGACTTGGAGAATTGGCAGAAGAAATCGAAAAT
NZ_CP038253.1:758105-765957    2501   ACTTTCCTGAAAGAAAAGGGAATTGCAGCCCCTGAGTTCGGTCTAATCCTTGGGTCAGGACTTGGAGAATTGGCAGAAGAAATCGAAAAT
NZ_CP035263.1:743050-750901    2501   ACTTTCCTGAAAGAAAAGGGAATTGCAGCCCCTGAGTTCGGTCTAATCCTTGGGTCAGGACTTGGAGAATTGGCAGAAGAAATCGAAAAT
NZ_CP035262.1:741595-749460    2515   ACTTTCCTGAAAGAAAAGGGAATTGCAGCCCCTGAGTTCGGTCTAATCCTTGGGTCAGGACTTGGAGAATTGGCAGAAGAAATCGAAAAT
NZ_CP035236.1:764915-772766    2501   ACTTTCCTGAAAGAAAAGGGAATTGCAGCCCCTGAGTTCGGTCTAATCCTTGGGTCAGGACTTGGAGAATTGGCAGAAGAAATCGAAAAT
NZ_CP035253.1 767286-775134    2502   ACTTTCCTGAAAGAAAAGGGAATTGCAGCCCCTGAGTTCGGTCTAATCCTTGGATCAGGACTTGGAGAATTGGCAGAAGAAATCGAAAAT
NZ_CP035249.1:763080-770921    2502   ACTTTCCTGAAAGAAAAGGGAATTGCAGCCCCTGAGTTCGGTCTAATCCTTGGATCAGGACTTGGAGAATTGGCAGAAGAAATCGAAAAT

NC_003028.3:777000-787000      3321   CCAGTTGTAGTAGACTATGCTGAGATTCCAAACTGGGGCCGTTCAACAGTAGTCGGTCATGCTGGTAAATTGGTATATGGTGAACTGGCA
NZ_CP035248.1:765099-775111    3330   CCAGTTGTAGTAGACTATGCAGATATTCCAAACTGGGGTCGTTCGACAGTAGTCGGTCACGCTGGTAAATTGGTATATGGTGACCTTGCA
NZ_CP035235.1 c1326834-1317894 2606   CCAGTTGTAGTAGACTATGCTGAGATTCCAAACTGGGGCCGTTCAACAGTAGTCGGTCATGCTGGTAAATTGGTATATGGTGAACTGGCA
NZ_CP035234.1:756284-765224    2606   CCAGTTGTAGTAGACTATGCTGAGATTCCAAACTGGGGCCGTTCAACAGTAGTCGGTCATGCTGGTAAATTGGTATATGGTGAACTGGCA
NZ_CP038251.1:750745-759686    2606   CCAGTTGTAGTAGACTATGCTGAGATTCCAAACTGGGGCCGTTCAACAGTAGTCGGTCATGCTGGTAAATTGGTATATGGTGAACTGGCA
NZ_CP035244.1:747023-755950    2592   CCAGTTGTAGTAGACTATGCTGAGATTCCAAACTGGGGCCGTTCAACAGTAGTCGGTCATGCTGGTAAATTGGTATATGGTGAACTGGCA
NZ_CP035264.1:800836-809336    2592   CCAGTTGTAGTAGACTATGCTGAGATTCCAAACTGGGGCCGTTCAACAGTAGTCGGTCATGCTGGTAAATTGGTATATGGTGAACTGGCA
NZ_CP035265.1:766128-774627    2592   CCAGTTGTAGTAGACTATGCTGAGATTCCAAACTGGGGCCGTTCAACAGTAGTCGGTCATGCTGGTAAATTGGTATATGGTGAACTGGCA
NZ_CP035245.1 745261-753757    2592   CCAGTTGTAGTAGACTATGCTGAGATTCCAAACTGGGGCCGTTCAACAGTAGTCGGTCATGCTGGTAAATTGGTATATGGTGAACTGGCA
NZ_CP035259.1:745700-753548    2592   CCAGTTGTAGTAGACTATGCTGAGATTCCAAACTGGGGCCGTTCAACAGTAGTCGGTCATGCTGGTAAATTGGTATATGGTGAACTGGCA
NZ_CP035258.1:750874-758722    2592   CCAGTTGTAGTAGACTATGCTGAGATTCCAAACTGGGGCCGTTCAACAGTAGTCGGTCATGCTGGTAAATTGGTATATGGTGAACTGGCA
NZ_CP035260.1:726156-733994    2592   CCAGTTGTAGTAGACTATGCTGAGATTCCAAACTGGGGCCGTTCAACAGTAGTCGGTCATGCTGGTAAATTGGTATATGGTGAACTGGCA
NZ_CP035241.1:708296-716144    2592   CCAGTTGTAGTAGACTATGCTGAGATTCCAAACTGGGGCCGTTCAACAGTAGTCGGTCATGCTGGTAAATTGGTATATGGTGAACTGGCA
NZ_CP035261.1:779086-786938    2592   CCAGTTGTAGTAGACTATGCTGAGATTCCAAACTGGGGCCGTTCAACAGTAGTCGGTCATGCTGGTAAATTGGTATATGGTGAACTGGCA
NZ_CP035242.1:c1532236-1524385 2591   CCAGTTGTAGTAGACTATGCTGAGATTCCAAACTGGGGCCGTTCAACAGTAGTCGGTCATGCTGGTAAATTGGTATATGGTGAACTGGCA
NZ_CP035243.1:777729-785580    2591   CCAGTTGTAGTAGACTATGCTGAGATTCCAAACTGGGGCCGTTCAACAGTAGTCGGTCATGCTGGTAAATTGGTATATGGTGAACTGGCA
NZ_CP035237.1:c1311322-1303456 2606   CCAGTTGTAGTAGACTATGCTGAGATTCCAAACTGGGGCCGTTCAACAGTAGTCGGTCATGCTGGTAAATTGGTATATGGTGAACTGGCA
NZ_CP035246.1:c1302892-1295026 2606   CCAGTTGTAGTAGACTATGCTGAGATTCCAAACTGGGGCCGTTCAACAGTAGTCGGTCATGCTGGTAAATTGGTATATGGTGAACTGGCA
NZ_CP035247.1:c1301115-1293263 2592   CCAGTTGTAGTAGACTATGCTGAGATTCCAAACTGGGGCCGTTCAACAGTAGTCGGTCATGCTGGTAAATTGGTATATGGTGAACTGGCA
NZ_CP035238.1:738547-746413    2606   CCAGTTGTAGTAGACTATGCTGAGATTCCAAACTGGGGCCGTTCAACAGTAGTCGGTCATGCTGGTAAATTGGTATATGGTGAACTGGCA
NZ_CP035255.1:776866-784718    2592   CCAGTTGTAGTAGACTATGCTGAGATTCCAAACTGGGGCCGTTCAACAGTAGTCGGTCATGCTGGTAAATTGGTATATGGTGAACTGGCA
NZ_CP035256.1:807236-815102    2606   CCAGTTGTAGTAGACTATGCTGAGATTCCAAACTGGGGCCGTTCAACAGTAGTCGGTCATGCTGGTAAATTGGTATATGGTGAACTGGCA
NZ_CP038252.1:1151812-1159673  2535   CCAGTTGTAGTAGACTATGCTGAGATTCCAAACTGGGGCCGTTCAACAGTAGTCGGTCATGCTGGTAAATTGGTATATGGTGAACTGGCA
NZ_CP035254.1:793365-801213    2592   CCAGTTGTAGTAGACTATGCTGAGATTCCAAACTGGGGCCGTTCAACAGTAGTCGGTCATGCTGGTAAATTGGTATATGGTGAACTGGCA
NZ_CP035251.1:748733-756585    2591   CCAGTTGTAGTAGACTATGCTGAGATTCCAAACTGGGGCCGTTCAACAGTAGTCGGTCATGCTGGTAAATTGGTATATGGTGAACTGGCA
NZ_CP035252.1:745670-753536    2605   CCAGTTGTAGTAGACTATGCTGAGATTCCAAACTGGGGCCGTTCAACAGTAGTCGGTCATGCTGGTAAATTGGTATATGGTGAACTGGCA
NZ_CP035257.1:776317-784169    2591   CCAGTTGTAGTAGACTATGCTGAGATTCCAAACTGGGGCCGTTCAACAGTAGTCGGTCATGCTGGTAAATTGGTATATGGTGAACTGGCA
NZ_CP038253.1:758105-765957    2591   CCAGTTGTAGTAGACTATGCTGAGATTCCAAACTGGGGCCGTTCAACAGTAGTCGGTCATGCTGGTAAATTGGTATATGGTGAACTGGCA
NZ_CP035263.1:743050-750901    2591   CCAGTTGTAGTAGACTATGCTGAGATTCCAAACTGGGGCCGTTCAACAGTAGTCGGTCATGCTGGTAAATTGGTATATGGTGAACTGGCA
NZ_CP035262.1:741595-749460    2605   CCAGTTGTAGTAGACTATGCTGAGATTCCAAACTGGGGCCGTTCAACAGTAGTCGGTCATGCTGGTAAATTGGTATATGGTGAACTGGCA
NZ_CP035236.1:764915-772766    2591   CCAGTTGTAGTAGACTATGCTGAGATTCCAAACTGGGGCCGTTCAACAGTAGTCGGTCATGCTGGTAAATTGGTATATGGTGAACTGGCA
NZ_CP035253.1 767286-775134    2592   CCAGTTGTAGTAGACTATGCTGAGATTCCAAACTGGGGCCGTTCAACAGTAGTCGGTCATGCTGGTAAATTGGTATATGGTGAACTGGCA
NZ_CP035249.1:763080-770921    2592   CCAGTTGTAGTAGACTATGCTGAGATTCCAAACTGGGGCCGTTCAACAGTAGTCGGTCATGCTGGTAAATTGGTATATGGTGAACTGGCA

NC_003028.3:777000-787000      3411   GGTCGCAAGGTCTTGGCTCTTCAAGGGCGTTTCCATTTCTATGAAGGGAATCCTCTGGAAGTGGTGACTTTCCCAGTTCGTGTGATGAAA
NZ_CP035248.1:765099-775111    3420   GGTCGCAAGGTCTTGGCTCTTCAAGGTCGTTTCCATTTCTACGAAGGAAATCCTCTGGAAGTAGTTACCTTCCCAGTACGTGTCATGAAA
NZ_CP035235.1 c1326834-1317894 2696   GGTCGCAAGGTCTTGGCTCTTCAAGGGCGTTTCCATTTCTATGAAGGGAATCCTCTGGAAGTGGTGACTTTCCCAGTTCGTGTGATGAAA
NZ_CP035234.1:756284-765224    2696   GGTCGCAAGGTCTTGGCTCTTCAAGGGCGTTTCCATTTCTATGAAGGGAATCCTCTGGAAGTGGTGACTTTCCCAGTTCGTGTGATGAAA
NZ_CP038251.1:750745-759686    2696   GGTCGCAAGGTCTTGGCTCTTCAAGGGCGTTTCCATTTCTATGAAGGGAATCCTCTGGAAGTGGTGACTTTCCCAGTTCGTGTGATGAAA
NZ_CP035244.1:747023-755950    2682   GGTCGCAAGGTCTTGGCTCTTCAAGGGCGTTTCCATTTCTATGAAGGGAATCCTCTGGAAGTGGTGACTTTCCCAGTTCGTGTGATGAAA
NZ_CP035264.1:800836-809336    2682   GGTCGCAAGGTCTTGGCTCTTCAAGGTCGTTTCCATTTCTATGAAGGGAATCCTCTGGAAGTGGTGACTTTCCCAGTTCGTGTGATGAAA
NZ_CP035265.1:766128-774627    2682   GGTCGCAAGGTCTTGGCTCTTCAAGGTCGTTTCCATTTCTATGAAGGGAATCCTCTGGAAGTGGTGACTTTCCCAGTTCGTGTGATGAAA
NZ_CP035245.1 745261-753757    2682   GGTCGCAAGGTCTTGGCTCTTCAAGGTCGTTTCCATTTCTATGAAGGGAATCCTCTGGAAGTGGTGACTTTCCCAGTTCGTGTGATGAAA
NZ_CP035259.1:745700-753548    2682   GGTCGCAAGGTCTTGGCTCTTCAAGGTCGTTTCCATTTCTATGAAGGGAATCCTCTGGAAGTGGTGACTTTCCCAGTTCGTGTGATGAAA
NZ_CP035258.1:750874-758722    2682   GGTCGCAAGGTCTTGGCTCTTCAAGGTCGTTTCCATTTCTATGAAGGGAATCCTCTGGAAGTGGTGACTTTCCCAGTTCGTGTGATGAAA
NZ_CP035260.1:726156-733994    2682   GGTCGCAAGGTCTTGGCTCTTCAAGGGCGTTTCCATTTCTATGAAGGGAATCCTCTGGAAGTGGTGACTTTCCCAGTTCGTGTGATGAAA
NZ_CP035241.1:708296-716144    2682   GGTCGCAAGGTCTTGGCTCTTCAAGGGCGTTTCCATTTCTATGAAGGGAATCCTCTGGAAGTGGTGACTTTCCCAGTTCGTGTGATGAAA
NZ_CP035261.1:779086-786938    2682   GGTCGCAAGGTCTTGGCTCTTCAAGGGCGTTTCCATTTCTATGAAGGGAATCCTCTGGAAGTGGTGACTTTCCCAGTTCGTGTGATGAAA
NZ_CP035242.1:c1532236-1524385 2681   GGTCGCAAGGTCTTGGCTCTTCAAGGGCGTTTCCATTTCTATGAAGGGAATCCTCTGGAAGTGGTGACTTTCCCAGTTCGTGTGATGAAA
NZ_CP035243.1:777729-785580    2681   GGTCGCAAGGTCTTGGCTCTTCAAGGGCGTTTCCATTTCTATGAAGGGAATCCTCTGGAAGTGGTGACTTTCCCAGTTCGTGTGATGAAA
NZ_CP035237.1:c1311322-1303456 2696   GGTCGCAAGGTCTTGGCTCTTCAAGGTCGTTTCCATTTCTATGAAGGGAATCCTCTGGAAGTGGTGACTTTCCCAGTTCGTGTGATGAAA
NZ_CP035246.1:c1302892-1295026 2696   GGTCGCAAGGTCTTGGCTCTTCAAGGGCGTTTCCATTTCTATGAAGGGAATCCTCTAGAAGTGGTGACTTTCCCAGTTCGTGTGATGAAA
NZ_CP035247.1:c1301115-1293263 2682   GGTCGCAAGGTCTTGGCTCTTCAAGGGCGTTTCCATTTCTATGAAGGGAATCCTCTAGAAGTGGTGACTTTCCCAGTTCGTGTGATGAAA
NZ_CP035238.1:738547-746413    2696   GGTCGCAAGGTCTTGGCTCTTCAAGGTCGTTTCCATTTCTATGAAGGGAATCCTCTGGAAGTGGTGACTTTCCCAGTTCGTGTGATGAAA
NZ_CP035255.1:776866-784718    2682   GGTCGCAAGGTCTTGGCTCTTCAAGGTCGTTTCCATTTCTATGAAGGGAATCCTCTGGAAGTGGTGACTTTCCCAGTTCGTGTGATGAAA
NZ_CP035256.1:807236-815102    2696   GGTCGCAAGGTCTTGGCTCTTCAAGGTCGTTTCCATTTCTATGAAGGGAATCCTCTGGAAGTGGTGACTTTCCCAGTTCGTGTGATGAAA
NZ_CP038252.1:1151812-1159673  2625   GGTCGCAAGGTCTTGGCTCTTCAAGGGCGTTTCCATTTCTATGAAGGGAATCCTCTGGAAGTGGTGACTTTCCCAGTTCGTGTGATGAAA
NZ_CP035254.1:793365-801213    2682   GGTCGCAAGGTCTTGGCTCTTCAAGGGCGTTTCCATTTCTATGAAGGGAATCCTCTGGAAGTGGTGACTTTCCCAGTTCGTGTGATGAAA
NZ_CP035251.1:748733-756585    2681   GGTCGCAAGGTCTTGGCTCTTCAAGGGCGTTTCCATTTCTATGAAGGGAATCCTCTGGAAGTGGTGACTTTCCCAGTTCGTGTGATGAAA
NZ_CP035252.1:745670-753536    2695   GGTCGCAAGGTCTTGGCTCTTCAAGGGCGTTTCCATTTCTATGAAGGGAATCCTCTGGAAGTGGTGACTTTCCCAGTTCGTGTGATGAAA
NZ_CP035257.1:776317-784169    2681   GGTCGCAAGGTCTTGGCTCTTCAAGGTCGTTTCCATTTCTATGAAGGGAATCCTCTGGAAGTGGTGACTTTCCCAGTTCGTGTGATGAAA
NZ_CP038253.1:758105-765957    2681   GGTCGCAAGGTCTTGGCTCTTCAAGGTCGTTTCCATTTCTATGAAGGGAATCCTCTGGAAGTGGTGACTTTCCCAGTTCGTGTGATGAAA
NZ_CP035263.1:743050-750901    2681   GGTCGCAAGGTCTTGGCTCTTCAAGGGCGTTTCCATTTCTATGAAGGGAATCCTCTGGAAGTGGTGACTTTCCCAGTTCGTGTGATGAAA
NZ_CP035262.1:741595-749460    2695   GGTCGCAAGGTCTTGGCTCTTCAAGGGCGTTTCCATTTCTATGAAGGGAATCCTCTGGAAGTGGTGACTTTCCCAGTTCGTGTGATGAAA
NZ_CP035236.1:764915-772766    2681   GGTCGCAAGGTCTTGGCTCTTCAAGGGCGTTTCCATTTCTATGAAGGGAATCCTCTGGAAGTGGTGACTTTCCCAGTTCGTGTGATGAAA
NZ_CP035253.1 767286-775134    2682   GGTCGCAAGGTCTTGGCTCTTCAAGGGCGTTTCCATTTCTATGAAGGGAATCCTCTGGAAGTGGTGACTTTCCCAGTTCGTGTGATGAAA
NZ_CP035249.1:763080-770921    2682   GGTCGCAAGGTCTTGGCTCTTCAAGGGCGTTTCCATTTCTATGAAGGGAATCCTCTGGAAGTGGTGACTTTCCCAGTTCGTGTGATGAAA

NC_003028.3:777000-787000      3501   GTTCTTGGATGTGAAGGTGTTATTGTAACCAATGCAGCTGGCGGTATCGGATTTGGTCCTGGTACCTTGATGGCTATCTCAGACCATATC
NZ_CP035248.1:765099-775111    3510   GTTCTCGGATGTGAAGGTGTCATTGTAACAAATGCAGCTGGTGGTATTGGATATGGTCCTGGTACTTTGATGGCTATCTCAGACCATATC
NZ_CP035235.1 c1326834-1317894 2786   GTTCTTGGATGTGAAGGTGTTATTGTAACCAATGCAGCTGGCGGTATCGGATTTGGTCCTGGTACCTTGATGGCTATCTCAGACCATATC
NZ_CP035234.1:756284-765224    2786   GTTCTTGGATGTGAAGGTGTTATTGTAACCAATGCAGCTGGCGGTATCGGATTTGGTCCTGGTACCTTGATGGCTATCTCAGACCATATC
```

```
NZ_CP038251.1:750745-759686    2786   GTTCTTGGATGTGAAGGTGTTATTGTAACCAATGCAGCTGGCGGTATCGGATTTGGTCCTGGTACCTTGATGGCTATCTCAGACCATATC
NZ_CP035244.1:747023-755950    2772   GTTCTTGGATGTGAAGGTGTTATTGTAACCAATGCAGCTGGCGGTATCGGATTTGGTCCTGGTACCTTGATGGCTATCTCAGACCATATC
NZ_CP035264.1:800836-809336    2772   GTTCTTGGATGTGAAGGTGTTATTGTAACCAATGCAGCTGGCGGTATCGGATTTGGTCCTGGTACCTTGATGGCTATCTCAGACCATATC
NZ_CP035265.1:766128-774627    2772   GTTCTTGGATGTGAAGGTGTTATTGTAACCAATGCAGCTGGCGGTATCGGATTTGGTCCTGGTACCTTGATGGCTATCTCAGACCATATC
NZ_CP035245.1 745261-753757    2772   GTTCTTGGATGTGAAGGTGTTATTGTAACCAATGCAGCTGGCGGTATCGGATTTGGTCCTGGTACCTTGATGGCTATCTCAGACCATATC
NZ_CP035259.1:745700-753548    2772   GTTATTGGATGTGAAGGTGTTATTGTAACCAATGCAGCTGGCGGTATCGGATTTGGTCCTGGTACCTTGATGGCTATCTCAGACCATATC
NZ_CP035258.1:750874-758722    2772   GTTATTGGATGTGAAGGTGTTATTGTAACCAATGCAGCTGGCGGTATCGGATTTGGTCCTGGTACCTTGATGGCTATCTCAGACCATATC
NZ_CP035260.1:726156-733994    2772   GTTCTTGGATGTGAAGGTGTTATTGTAACCAATGCAGCTGGCGGTATCAGATTTGGTCCTGGTACCTTGATGGCTATCTCAGACCATATC
NZ_CP035241.1:708296-716144    2772   GTTCTTGGATGTGAAGGTGTTATTGTAACCAATGCAGCTGGCGGTATCAGATTTGGTCCTGGTACCTTGATGGCTATCTCAGACCATATC
NZ_CP035261.1:779086-786938    2772   GTTCTTGGATGTGAAGGTGTTATTGTAACCAATGCAGCTGGCGGTATCGGATTTGGTCCTGGTACCTTGATGGCTATCTCAGACCATATC
NZ_CP035242.1:c1532236-1524385 2771   GTTCTTGGATGTGAAGGTGTTATTGTAACCAATGCAGCTGGCGGTATCGGATTTGGTCCTGGTACCTTGATGGCTATCTCAGACCATATC
NZ_CP035243.1:777729-785580    2771   GTTCTTGGATGTGAAGGTGTTATTGTAACCAATGCAGCTGGCGGTATCGGATTTGGTCCTGGTACCTTGATGGCTATCTCAGACCATATC
NZ_CP035237.1:c1311322-1303456 2786   GTTCTTGGATGTGAAGGTGTTATTGTAACCAATGCAGCTGGCGGTATCAGATTTGGTCCTGGTACCTTGATGGCTATCTCAGACCATATC
NZ_CP035246.1:c1302892-1295026 2786   GTTCTTGGATGTGAAGGTGTTATTGTAACCAATGCAGCTGGCGGTATCGGATTTGGTCCTGGTACCTTGATGGCTATCTCAGACCATATC
NZ_CP035247.1:c1301115-1293263 2772   GTTCTTGGATGTGAAGGTGTTATTGTAACCAATGCAGCTGGCGGTATCGGATTTGGTCCTGGTACCTTGATGGCTATCTCAGACCATATC
NZ_CP035238.1:738547-746413    2786   GTTCTTGGATGTGAAGGTGTTATTGTAACCAATGCAGCTGGCGGTATCGGATTTGGTCCTGGTACCTTGATGGCTATCTCAGACCATATC
NZ_CP035255.1:776866-784718    2772   GTTCTTGGATGTGAAGGTGTTATTGTAACCAATGCAGCTGGCGGTATCGGATTTGGTCCTGGTACCTTGATGGCTATCTCAGACCATATC
NZ_CP035256.1:807236-815102    2786   GTTCTTGGATGTGAAGGTGTTATTGTAACCAATGCAGCTGGCGGTATCGGATTTGGTCCTGGTACCTTGATGGCTATCTCAGACCATATC
NZ_CP038252.1:1151812-1159673  2715   GTTCTTGGATGTGAAGGTGTTATTGTAACCAATGCAGCTGGCGGTATCGGATTTGGTCCTGGTACCTTGATGGCTATCTCAGACCATATC
NZ_CP035254.1:793365-801213    2772   GTTCTTGGATGTGAAGGTGTTATTGTAACCAATGCAGCTGGCGGTATCGGATTTGGTCCTGGTACCTTGATGGCTATCTCAGACCATATC
NZ_CP035251.1:748733-756585    2771   GTTCTTGGATGTGAAGGTGTTATTGTAACCAATGCAGCTGGCGGTATCGGATTTGGTCCTGGTACCTTGATGGCTATCTCAGACCATATC
NZ_CP035252.1:745670-753536    2785   GTTCTTGGATGTGAAGGTGTTATTGTAACCAATGCAGCTGGCGGTATCGGATTTGGTCCTGGTACCTTGATGGCTATCTCAGACCATATC
NZ_CP035257.1:776317-784169    2771   GTTCTTGGATGTGAAGGTGTTATTGTAACCAATGCAGCTGGCGGTATCGGATTTGGTCCTGGTACCTTGATGGCTATCTCAGACCATATC
NZ_CP038253.1:758105-765957    2771   GTTCTTGGATGTGAAGGTGTTATTGTAACCAATGCAGCTGGCGGTATCGGATTTGGTCCTGGTACCTTGATGGCTATCTCAGACCATATC
NZ_CP035263.1:743050-750901    2771   GTTCTTGGATGTGAAGGTGTTATTGTAACCAATGCAGCTGGCGGTATCGGATTTGGTCCTGGTACCTTGATGGCTATCTCAGACCATATC
NZ_CP035262.1:741595-749460    2785   GTTCTTGGATGTGAAGGTGTTATTGTAACCAATGCAGCTGGCGGTATCGGATTTGGTCCTGGTACCTTGATGGCTATCTCAGACCATATC
NZ_CP035236.1:764915-772766    2771   GTTCTTGGATGTGAAGGTGTTATTGTAACCAATGCAGCTGGCGGTATCGGATTTGGTCCTGGTACCTTGATGGCTATCTCAGACCATATC
NZ_CP035253.1 767286-775134    2772   GTTCTTGGATGTGAAGGTGTTATTGTAACCAATGCAGCTGGCGGTATCGGATTTGGTCCTGGTACCTTGATGGCTATCTCAGACCATATC
NZ_CP035249.1:763080-770921    2772   GTTCTTGGATGTGAAGGTGTTATTGTAACCAATGCAGCTGGCGGTATCGGATTTGGTCCTGGTACCTTGATGGCTATCTCAGACCATATC

NC_003028.3:777000-787000      3591   AACATGACGGGGCAAAATCCATTGATGGGTGAAAACTTGGATGACTTTGGCCCACGTTTCCCAGATATGTCTAGGGCCTACACACCAGAA
NZ_CP035248.1:765099-775111    3600   AACATGACGGGGCAGAACCCATTGATGGGTGAAAACTTGGATGACTTTGGTCCACGTTTCCCAGATATGTCTAAATCCTACACTCCAGAA
NZ_CP035235.1 c1326834-1317894 2876   AACATGACGGGGCAAAATCCATTGATGGGTGAAAACTTGGATGACTTTGGCCCACGTTTCCCAGATATGTCTAGGGCTTACACACCAGAA
NZ_CP035234.1:756284-765224    2876   AACATGACGGGGCAAAATCCATTGATGGGTGAAAACTTGGATGACTTTGGCCCACGTTTCCCAGATATGTCTAGGGCTTACACACCAGAA
NZ_CP038251.1:750745-759686    2876   AACATGACGGGGCAAAATCCATTGATGGGTGAAAACTTGGATGACTTTGGCCCACGTTTCCCAGATATGTCTAGGGCTTACACACCAGAA
NZ_CP035244.1:747023-755950    2862   AACATGACGGGGCAAAATCCATTGATGGGTGAAAACTTGGATGACTTTGGCCCACGTTTCCCAGATATGTCTAGGGCTTACACACCAGAA
NZ_CP035264.1:800836-809336    2862   AACATGACGGGGCAAAATCCATTGATGGGTGAAAACTTGGATGACTTTGGCCCACGTTTCCCAGATATGTCTAGGGCCTACACACCAGAA
NZ_CP035265.1:766128-774627    2862   AACATGACGGGGCAAAATCCATTGATGGGTGAAAACTTGGATGACTTTGGCCCACGTTTCCCAGATATGTCTAGGGCCTACACACCAGAA
NZ_CP035245.1 745261-753757    2862   AACATGACGGGGCAAAATCCATTGATGGGTGAAAACTTGGATGACTTTGGCCCACGTTTCCCAGATATGTCTAGGACCTACACACCAGAA
NZ_CP035259.1:745700-753548    2862   AACATGACGGGGCAAAATCCATTGATGGGTGAAAACTTGGATGACTTTGGCCCACGTTTCCCAGATATGTCTAGGGCCTACACACCAGAA
NZ_CP035258.1:750874-758722    2862   AACATGACGGGGCAAAATCCATTGATGGGTGAAAACTTGGATGACTTTGGCCCACGTTTCCCAGATATGTCTAGGGCCTACACACCAGAA
NZ_CP035260.1:726156-733994    2862   AACATGACGGGGCAAAATCCATTGATGGGTGAAAACTTGGATGACTTTGGCCCACGTTTCCCAGATATGTCTAGGGCCTACACACCAGAA
NZ_CP035241.1:708296-716144    2862   AACATGACGGGGCAAAATCCATTGATGGGGGAAAACTTGGATGACTTTGGCCCACGTTTCCCAGATATGTCTAGGGCCTACACACCAGAA
NZ_CP035261.1:779086-786938    2862   AACATGACGGGGCAAAATCCATTGATGGGTGAAAACTTGGATGACTTTGGCCCACGTTTCCCAGATATGTCTAGGGCCTACACACCAGAA
NZ_CP035242.1:c1532236-1524385 2861   AACATGACGGGGCAAAATCCATTGATGGGTGAAAACTTGGATGACTTTGGCCCACGTTTCCCAGATATGTCTAGGGCCTACACACCAGAA
NZ_CP035243.1:777729-785580    2861   AACATGACGGGGCAAAATCCATTGATGGGTGAAAACTTGGATGACTTTGGCCCACGTTTCCCAGATATGTCTAGGGCCTACACACCAGAA
NZ_CP035237.1:c1311322-1303456 2876   AACATGACGGGGCAAAATCCATTGATGGGTGAAAACTTGGATGACTTTGGCCCACGTTTCCCAGATATGTCTAGGGCCTACACACCAGAA
NZ_CP035246.1:c1302892-1295026 2876   AACATGACGGGGCAAAATCCATTGATGGGTGAAAACTTGGATGACTTTGGCCCACGTTTCCCAGATATGTCTAGGGCCTACACACCAGAA
NZ_CP035247.1:c1301115-1293263 2862   AACATGACGGGGCAAAATCCATTGATGGGTGAAAACTTGGATGACTTTGGCCCACGTTTCCCAGATATGTCTAGGGCCTACACACCAGAA
NZ_CP035238.1:738547-746413    2876   AACATGACGGGGCAAAATCCATTGATGGGTGAAAACTTGGATGACTTTGGCCCACGTTTCCCAGATATGTCTAGGGCCTACACACCAGAA
NZ_CP035255.1:776866-784718    2862   AACATGACGGGGCAAAATCCATTGATGGGTGAAAACTTGGATGACTTTGGCCCACGTTTCCCAGATATGTCTAGGGCCTACACACCAGAA
NZ_CP035256.1:807236-815102    2876   AACATGACGGGGCAAAATCCATTGATGGGTGAAAACTTGGATGACTTTGGCCCACGTTTCCCAGATATGTCTAGGGCCTACACACCAGAA
NZ_CP038252.1:1151812-1159673  2805   AACATGACGGGGCAAAATCCATTGATGGGTGAAAACTTGAATGACTTTGGCCCACGTTTCCCAGATATGTCTAGGACCTACACACCAGAA
NZ_CP035254.1:793365-801213    2862   AACATGACGGGGCAAAATCCATTGATGGGTGAAAACTTGGATGACTTTGGCCCACGTTTCCCAGATATGTCTAGGGCCTACACACCAGAA
NZ_CP035251.1:748733-756585    2861   AACATGACGGGGCAAAATCCATTGATGGGTGAAAACTTGGATGACTTTGGCCCACGTTTCCCAGATATGTCTAGGGCCTACACACCAGAA
NZ_CP035252.1:745670-753536    2875   AACATGACGGGGCAAAATCCATTGATGGGTGAAAACTTGGATGACTTTGGCCCACGTTTCCCAGATATGTCTAGGGCCTACACACCAGAA
NZ_CP035257.1:776317-784169    2861   AACATGACGGGGCAAAATCCATTGATGGGTGAAAACTTGGATGACTTTGGCCCACGTTTCCCAGATATGTCTAGGGCCTACACACCAGAA
NZ_CP038253.1:758105-765957    2861   AACATGACGGGGCAAAATCCATTGATGGGTGAAAACTTGGATGACTTTGGCCCACGTTTCCCAGATATGTCTAGGGCCTACACACCAGAA
NZ_CP035263.1:743050-750901    2861   AACATGACGGGGCAAAATCCATTGATGGGTGAAAACTTGGATGACTTTGGCCCACGTTTCCCAGATATGTCTAGGGCCTACACACCAGAA
NZ_CP035262.1:741595-749460    2875   AACATGACGGGGCAAAATCCATTGATGGGTGAAAACTTGGATGACTTTGGCCCACGTTTCCCAGATATGTCTAGGGCCTACACACCAGAA
NZ_CP035236.1:764915-772766    2861   AACATGACGGGGCAAAATCCATTGATGGGTGAAAACTTGGATGACTTTGGCCCACGTTTCCCAGATATGTCTAGGGCCTACACACCAGAA
NZ_CP035253.1 767286-775134    2862   AACATGACGGGGCAAAATCCATTGATGGGTGAAAACTTGGATGACTTTGGCCCACGTTTCCCAGATATGTCTAGGGCCTACACACCAGAA
NZ_CP035249.1:763080-770921    2862   AACATGACGGGGCAAAATCCATTGATGGGTGAAAACTTGGATGACTTTGGCCCACGTTTCCCAGATATGTCTAGGGCCTACACACCAGAA

NC_003028.3:777000-787000      3681   TACCGTGCCACTGCCCATGAAGTGGCTAAAAAACTTAATATCAAGCTTGATGAAGGTGTCTATATCGGAGTTACTGGTCCGACTTATGAA
NZ_CP035248.1:765099-775111    3690   TATCGTGCTACTGCCCATGAAGTTGCTAAGAAACTTGGTATCAAGCTTGATGAAGGTGTCTATATCGGTGTTACTGGTCCAACTTATGAA
NZ_CP035235.1 c1326834-1317894 2966   TACCGTGCCACTGCCCATGAAGTGGCTAAAAAACTTAATATCAAGCTTGATGAAGGTGTCTATATCGGAGTTACTGGTCCGACTTATGAA
NZ_CP035234.1:756284-765224    2966   TACCGTGCCACTGCCCATGAAGTGGCTAAAAAACTTAATATCAAGCTTGATGAAGGTGTCTATATCGGAGTTACTGGTCCGACTTATGAA
NZ_CP038251.1:750745-759686    2966   TACCGTGCCACTGCCCATGAAGTGGCTAAAAAACTTAATATCAAGCTTGATGAAGGTGTCTATATCGGAGTTACTGGTCCGACTTATGAA
NZ_CP035244.1:747023-755950    2952   TACCGTGCCACTGCCCATGAAGTGGCTAAAAAACTTAATATCAAGCTTGATGAAGGTGTCTATATCGGAGTTACTGGTCCGACTTATGAA
NZ_CP035264.1:800836-809336    2952   TACCGTGCCACTGCCCATGAAGTGGCTAAAAAACTTAATATCAAGCTTGATGAAGGTGTCTATATCGGAGTTACTGGTCCGACTTATGAA
NZ_CP035265.1:766128-774627    2952   TACCGTGCCACTGCCCATGAAGTGGCTAAAAAACTTAATATCAAGCTTGATGAAGGTGTCTATATCGGAGTTACTGGTCCGACTTATGAA
NZ_CP035245.1 745261-753757    2952   TACCGTGCCACTGCCCATGAAGTAGCTAAAAAACTTAATATCAAGCTTGATGAAGGTGTCTATATCGGAGTTACTGGTCCGACTTATGAA
NZ_CP035259.1:745700-753548    2952   TACCGTGCTACTGCCCATGAAGTGGCTAAAAAACTTAATATCAAGCTTGATGAAGGTGTCTATATCGGAGTTACTGGTCCGACTTATGAA
NZ_CP035258.1:750874-758722    2952   TACCGTGCTACTGCCCATGAAGTGGCTAAAAAACTTAATATCAAGCTTGATGAAGGTGTCTATATCGGAGTTACTGGTCCGACTTATGAA
NZ_CP035260.1:726156-733994    2952   TACCGTGCCACTGCCCATGAAGTGGCTAAAAAACTTAATATCAAGATTGATGAAGGTGTCTATATCGGAGTTACTGGTCCGACTTATGAA
NZ_CP035241.1:708296-716144    2952   TACCGTGCCACTGCCCATGAAGTGGCTAAAAAACTTAATATCAAGATTGATGAAGGTGTCTATATCGGAGTTACTGGTCCGACTTATGAA
NZ_CP035261.1:779086-786938    2952   TACCGTGCCACTGCCCATGAAGTGGCTAAAAAACTTAATATCAAGCTTGATGAAGGTGTCTATATCGGAGTTACTGGTCCGACTTATGAA
NZ_CP035242.1:c1532236-1524385 2951   TACCGTGCCACTGCCCATGAAGTGGCTAAAAAACTTAATATCAAGCTTGATGAAGGTGTCTATATCGGAGTTACTGGTCCGACTTATGAA
NZ_CP035243.1:777729-785580    2951   TACCGTGCCACTGCCCATGAAGTGGCTAAAAAACTTAATATCAAGCTTGATGAAGGTGTCTATATCGGAGTTACTGGTCCGACTTATGAA
NZ_CP035237.1:c1311322-1303456 2966   TACCGTGCCACTGCCCATGAAGTGGCTAAAAAACTTAATATCAAGCTTGATGAAGGTGTCTATATCGGAGTTACTGGTCCGACTTATGAA
NZ_CP035246.1:c1302892-1295026 2966   TACCGTGCCACTGCCCATGAAGTGGCTAAAAAACTTAATATCAAGCTTGATGAAGGTGTCTATATCGGAGTTACTGGTCCGACTTATGAA
NZ_CP035247.1:c1301115-1293263 2952   TACCGTGCCACTGCCCATGAAGTGGCTAAAAAACTTAATATCAAGCTTGATGAAGGTGTCTATATCGGAGTTACTGGTCCGACTTATGAA
NZ_CP035238.1:738547-746413    2966   TACCGTGCCACTGCCCATGAAGTGGCTAAAAAACTTAATATCAAGCTTGATGAAGGTGTCTATATCGGAGTTACTGGTCCGACTTATGAA
NZ_CP035255.1:776866-784718    2952   TACCGTGCCACTGCCCATGAAGTGGCTAAAAAACTTAATATCAAGCTTGATGAAGGTGTCTATATCGGAGTTACTGGTCCGACTTATGAA
NZ_CP035256.1:807236-815102    2966   TACCGTGCCACTGCCCATGAAGTGGCTAAAAAACTTAATATCAAGCTTGATGAAGGTGTCTATATCGGAGTTACTGGTCCGACTTATGAA
NZ_CP038252.1:1151812-1159673  2895   TACCGTGCCACTGCCCATGAAGTGGCTAAAAAACTTAATATCAAGCTTGATGAAGGTGTCTATATCGGAGTTACTGGTCCGACTTATGAA
NZ_CP035254.1:793365-801213    2952   TACCGTGCCACTGCCCATGAAGTGGCTAAAAAACTTAATATCAAGCTTGATGAAGGTGTCTATATCGGAGTTACTGGTCCGACTTATGAA
NZ_CP035251.1:748733-756585    2951   TACCGTGCCACTGCCCATGAAGTGGCTAAAAAACTTAATATCAAGCTTGATGAAGGTGTCTATATCGGAGTTACTGGTCCGACTTATGAA
NZ_CP035252.1:745670-753536    2965   TACCGTGCCACTGCCCATGAAGTGGCTAAAAAACTTAATATCAAGCTTGATGAAGGTGTCTATATCGGAGTTACTGGTCCGACTTATGAA
NZ_CP035257.1:776317-784169    2951   TACCGTGCCACTGCCCATGAAGTGGCTAAAAAACTTAATATCAAGCTTGATGAAGGTGTCTATATCGGAGTTACTGGTCCGACTTATGAA
NZ_CP038253.1:758105-765957    2951   TACCGTGCCACTGCCCATGAAGTGGCTAAAAAACTTAATATCAAGCTTGATGAAGGTGTCTATATCGGAGTTACTGGTCCGACTTATGAA
NZ_CP035263.1:743050-750901    2951   TACCGTGCCACTGCCCATGAAGTGGCTAAAAAACTTAATATCAAGCTTGATGAAGGTGTCTATATCGGAGTTACTGGTCCGACTTATGAA
NZ_CP035262.1:741595-749460    2965   TACCGTGCCACTGCCCATGAAGTGGCTAAAAAACTTAATATCAAGCTTGATGAAGGTGTCTATATCGGAGTTACTGGTCCGACTTATGAA
NZ_CP035236.1:764915-772766    2951   TACCGTGCCACTGCCCATGAAGTGGCTAAAAAACTTAATATCAAGCTTGATGAAGGTGTCTATATCGGAGTTACTGGTCCGACTTATGAA
```

```
NZ_CP035253.1 767286-775134    2952   TACCGTGCCACTGCCCATGAAGTGGCTAAAAAACTTAATATCAAGCTTGATGAAGGTGTCTATATCGGAGTTACTGGTCCGACTTATGAA
NZ_CP035249.1:763080-770921    2952   TACCGTGCCACTGCCCATGAAGTGGCTAAAAAACTTAATATCAAGCTTGATGAAGGTGTCTATATCGGAGTTACTGGTCCGACTTATGAA

NC_003028.3:777000-787000      3771   ACACCAGCAGAAATTCGTTCCTATAAGACACTGGGAGCAGATGCAGTTGGTATGTCTACGGTTCCTGAAGTTATCGTGGCAGCCCACTCT
NZ_CP035248.1:765099-775111    3780   ACACCAGCAGAGATTCGCGCCTATAAGATACTTGGAGCAGATGCAGTTGGTATGTCTACAGTTCCTGAAGTGATTGTGGCAGCCCACTCA
NZ_CP035235.1 c1326834-1317894 3056   ACACCAGCAGAAATTCGTTCCTATAAGACACTGGGAGCAGATGCAGTTGGTATGTCTACGGTTCCTGAAGTTATCGTGGCAGCCCACTCT
NZ_CP035234.1:756284-765224    3056   ACACCAGCAGAAATTCGTTCCTATAAGACACTGGGAGCAGATGCAGTTGGTATGTCTACGGTTCCTGAAGTTATCGTGGCAGCCCACTCT
NZ_CP038251.1:750745-759686    3056   ACACCAGCAGAAATTCGTTCCTATAAGACACTGGGAGCAGATGCAGTTGGTATGTCTACGGTTCCTGAAGTTATCGTGGCAGCCCACTCT
NZ_CP035244.1:747023-755950    3042   ACACCAGCAGAAATTCGTTCCTATAAGACACTGGGAGCAGATGCAGTTGGTATGTCTACGGTTCCTGAAGTTATCGTGGCAGCCCACTCT
NZ_CP035264.1:800836-809336    3042   ACACCAGCAGAAATTCGTTCCTATAAGACACTGGGAGCAGATGCAGTTGGTATGTCTACGGTTCCTGAAGTTATCGTGGCAGCCCACTCT
NZ_CP035265.1:766128-774627    3042   ACACCAGCAGAAATTCGTTCCTATAAGACACTGGGAGCAGATGCAGTTGGTATGTCTACGGTTCCTGAAGTTATCGTGGCAGCCCACTCT
NZ_CP035245.1 745261-753757    3042   ACACCAGCAGAAATTCGTTCCTATAAGACACTGGGAGCAGATGCAGTTGGTATGTCTACGGTTCCTGAAGTTATCGTGGCAGCCCACTCT
NZ_CP035259.1:745700-753548    3042   ACACCAGCAGAAATTCGTTCCTATAAGACACTGGGAGCAGATGCAGTTGGTATGTCTACGGTTCCTGAAGTTATCGTGGCAGCCCACTCT
NZ_CP035258.1:750874-758722    3042   ACACCAGCAGAAATTCGTTCCTATAAGACACTGGGAGCAGATGCAGTTGGTATGTCTACGGTTCCTGAAGTTATCGTGGCAGCCCACTCT
NZ_CP035260.1:726156-733994    3042   ACACCAGCAGAAATTCGTTCCTATAAGACACTGGGAGCAGATGCAGTTGGTATGTCTACGGTTCCTGAAGTTATCGTGGCAGCCCACTCT
NZ_CP035241.1:708296-716144    3042   ACACCAGCAGAAATTCGTTCCTATAAGACACTGGGAGCAGATGCAGTTGGTATGTCTACGGTTCCTGAAGTTATCGTGGCAGCCCACTCT
NZ_CP035261.1:779086-786938    3042   ACACCAGCAGAAATTCGTTCCTATAAGACACTGGGAGCAGATGCAGTTGGTATGTCTACGGTTCCTGAAGTTATCGTGGCAGCCCACTCT
NZ_CP035242.1:c1532236-1524385 3041   ACACCAGCAGAAATTCGTTCCTATAAGACACTGGGAGCAGATGCAGTTGGTATGTCTACGGTTCCTGAAGTTATCGTGGCAGCCCACTCT
NZ_CP035243.1:777729-785580    3041   ACACCAGCAGAAATTCGTTCCTATAAGACACTGGGAGCAGATGCAGTTGGTATGTCTACGGTTCCTGAAGTTATCGTGGCAGCCCACTCT
NZ_CP035237.1:c1311322-1303456 3056   ACACCAGCAGAAATTCGTTCCTATAAGACACTGGGAGCAGATGCAGTTGGTATGTCTACGGTTCCTGAAGTTATCGTGGCAGCCCACTCT
NZ_CP035246.1:c1302892-1295026 3056   ACACCAGCAGAAATTCGTTCCTATAAGACACTGGGAGCAGATGCAGTTGGTATGTCTACGGTTCCTGAAGTTATCGTGGCAGCCCACTCT
NZ_CP035247.1:c1301115-1293263 3042   ACACCAGCAGAAATTCGTTCCTATAAGACACTGGGAGCAGATGCAGTTGGTATGTCTACGGTTCCTGAAGTTATCGTGGCAGCCCACTCT
NZ_CP035238.1:738547-746413    3056   ACACCAGCAGAAATTCGTTCCTATAAGACACTGGGAGCAGATGCAGTTGGTATGTCTACGGTTCCTGAAGTTATCGTGGCAGCCCACTCT
NZ_CP035255.1:776866-784718    3042   ACACCAGCAGAAATTCGTTCCTATAAGACACTGGGAGCAGATGCAGTTGGTATGTCTACGGTTCCTGAAGTTATCGTGGCAGCCCACTCT
NZ_CP035256.1:807236-815102    3056   ACACCAGCAGAAATTCGTTCCTATAAGACACTGGGAGCAGATGCAGTTGGTATGTCTACGGTTCCTGAAGTTATCGTGGCAGCCCACTCT
NZ_CP038252.1:1151812-1159673  2985   ACACCAGCAGAAATTCGTTCCTATAAGACACTGGGAGCAGATGCAGTTGGTATGTCTACGGTTCCTGAAGTTATCGTGGCAGCCCACTCT
NZ_CP035254.1:793365-801213    3042   ACACCAGCAGAAATTCGTTCCTATAAGACACTGGGAGCAGATGCAGTTGGTATGTCTACGGTTCCTGAAGTTATCGTGGCAGCCCACTCT
NZ_CP035251.1:748733-756585    3041   ACACCAGCAGAAATTCGTTCCTATAAGACACTGGGAGCAGATGCAGTTGGTATGTCTACGGTTCCTGAAGTTATCGTGGCGGCCCACTCT
NZ_CP035252.1:745670-753536    3055   ACACCAGCAGAAATTCGTTCCTATAAGACACTGGGAGCAGATGCAGTTGGTATGTCTACGGTTCCTGAAGTTATCGTGGCGGCCCACTCT
NZ_CP035257.1:776317-784169    3041   ACACCAGCAGAAATTCGTTCCTATAAGACACTGGGAGCAGATGCAGTTGGTATGTCTACGGTTCCTGAAGTTATCGTGGCAGCCCACTCT
NZ_CP038253.1:758105-765957    3041   ACACCAGCAGAAATTCGTTCCTATAAGACACTGGGAGCAGATGCAGTTGGTATGTCTACGGTTCCTGAAGTTATCGTGGCAGCCCACTCT
NZ_CP035263.1:743050-750901    3041   ACACCAGCAGAAATTCGTTCCTATAAGACACTGGGAGCAGATGCAGTTGGTATGTCTACGGTTCCTGAAGTTATCGTGGCGGCCCACTCT
NZ_CP035262.1:741595-749460    3055   ACACCAGCAGAAATTCGTTCCTATAAGACACTGGGAGCAGATGCAGTTGGTATGTCTACGGTTCCTGAAGTTATCGTGGCGGCCCACTCT
NZ_CP035236.1:764915-772766    3041   ACACCAGCAGAAATTCGTTCCTATAAGACACTGGGAGCAGATGCAGTTGGTATGTCTACGGTTCCTGAAGTTATCGTGGCGGCCCACTCT
NZ_CP035253.1 767286-775134    3042   ACACCAGCAGAAATTCGTTCCTATAAGACACTGGGAGCAGATGCAGTTGGTATGTCTACGGTTCCTGAAGTTATCGTGGCAGCCCACTCT
NZ_CP035249.1:763080-770921    3042   ACACCAGCAGAAATTCGTTCCTATAAGACACTGGGAGCAGATGCAGTTGGTATGTCTACGGTTCCTGAAGTTATCGTGGCAGCCCACTCT

NC_003028.3:777000-787000      3861   GGCTTGAAAGTTCTGGGAATTTCATGTATCACTAACTTTGCGGCCGGTTTCCAAGAAGAACTCAATCACGAAGAAGTTGTAGAAGTGACT
NZ_CP035248.1:765099-775111    3870   GGTTTGAAAGTTCTTGGTATTTCATGTATCACTAACTTTGCGGCTGGTTTCCAAGAAGAACTTAATCACGAAGAAGTTGTAGAAGTAACT
NZ_CP035235.1 c1326834-1317894 3146   GGCTTGAAAGTTCTGGGAATTTCATGTATCACTAACTTTGCGGCCGGTTTCCAAGAAGAACTCAATCACGAAGAAGTTGTAGAAGTGACT
NZ_CP035234.1:756284-765224    3146   GGCTTGAAAGTTCTGGGAATTTCATGTATCACTAACTTTGCGGCCGGTTTCCAAGAAGAACTCAATCACGAAGAAGTTGTAGAAGTGACT
NZ_CP038251.1:750745-759686    3146   GGCTTGAAAGTTCTGGGAATTTCATGTATCACTAACTTTGCGGCCGGTTTCCAAGAAGAACTCAATCACGAAGAAGTTGTAGAAGTGACT
NZ_CP035244.1:747023-755950    3132   GGCTTGAAAGTTCTGGGAATTTCATGTATCACTAACTTTGCGGCCGGTTTCCAAGAAGAACTCAATCACGAAGAAGTTGTAGAAGTGACT
NZ_CP035264.1:800836-809336    3132   GGCTTGAAAGTTCTGGGAATTTCATGTATCACTAACTTTGCGGCTGGTTTCCAAGAAGAACTCAATCACGAAGAAGTTGTAGAAGTGACT
NZ_CP035265.1:766128-774627    3132   GGCTTGAAAGTTCTGGGAATTTCATGTATCACTAACTTTGCGGCTGGTTTCCAAGAAGAACTCAATCACGAAGAAGTTGTAGAAGTGACT
NZ_CP035245.1 745261-753757    3132   GGCTTGAAAGTTCTGGGAATTTCATGTATCACTAACTTTGCGGCTGGTTTCCAAGAAGAACTCAATCACGAAGAAGTTGTAGAAGTGACT
NZ_CP035259.1:745700-753548    3132   GGCTTGAAAGTTCTGGGAATTTCATGTATCACTAACTTTGCGGCTGGTTTCCAAGAAGAACTCAATCACGAAGAAGTTGTAGAAGTGACT
NZ_CP035258.1:750874-758722    3132   GGCTTGAAAGTTCTGGGAATTTCATGTATCACTAACTTTGCGGCTGGTTTCCAAGAAGAACTCAATCACGAAGAAGTTGTAGAAGTGACT
NZ_CP035260.1:726156-733994    3132   GGCTTGAAAGTTCTGGGAATTTCATGTATCACTAACTTTGCGGCTGGTTTCCAAGAAGAACTCAATCACGAAGAAGTTGTAGAAGTGACT
NZ_CP035241.1:708296-716144    3132   GGCTTGAAAGTTCTGGGAATTTCATGTATCACTAACTTTGCGGCTGGTTTCCAAGAAGAACTCAATCACGAAGAAGTTGTAGAAGTGACT
NZ_CP035261.1:779086-786938    3132   GGCTTGAAAGTTCTGGGAATTTCATGTATCACTAACTTTGCGGCCGGTTTCCAAGAAGAACTCAATCACGAAGAAGTTGTAGAAGTGACT
NZ_CP035242.1:c1532236-1524385 3131   GGCTTGAAAGTTCTGGGAATTTCATGTATCACTAACTTTGCGGCCGGTTTCCAAGAAGAACTCAATCACGAAGAAGTTGTAGAAGTGACT
NZ_CP035243.1:777729-785580    3131   GGCTTGAAAGTTCTGGGAATTTCATGTATCACTAACTTTGCGGCCGGTTTCCAAGAAGAACTCAATCACGAAGAAGTTGTAGAAGTGACT
NZ_CP035237.1:c1311322-1303456 3146   GGCTTGAAAGTTCTGGGAATTTCATGTATCACTAACTTTGCGGCTGGTTTCCAAGAAGAACTCAATCACGAAGAAGTTGTAGAAGTGACT
NZ_CP035246.1:c1302892-1295026 3146   GGCTTGAAAGTTCTGGGAATTTCATGTATCACTAACTTTGCGGCTGGTTTCCAAGAAGAACTCAATCACGAAGAAGTTGTAGAAGTGACT
NZ_CP035247.1:c1301115-1293263 3132   GGCTTGAAAGTTCTGGGAATTTCATGTATCACTAACTTTGCGGCTGGTTTCCAAGAAGAACTCAATCACGAAGAAGTTGTAGAAGTGACT
NZ_CP035238.1:738547-746413    3146   GGCTTGAAAGTTCTGGGAATTTCATGTATCACTAACTTTGCGGCTGGTTTCCAAGAAGAACTCAATCACGAAGAAGTTGTAGAAGTGACT
NZ_CP035255.1:776866-784718    3132   GGCTTGAAAGTTCTGGGAATTTCATGTATCACTAACTTTGCGGCTGGTTTCCAAGAAGAACTCAATCACGAAGAAGTTGTAGAAGTGACT
NZ_CP035256.1:807236-815102    3146   GGCTTGAAAGTTCTGGGAATTTCATGTATCACTAACTTTGCGGCTGGTTTCCAAGAAGAACTCAATCACGAAGAAGTTGTAGAAGTGACT
NZ_CP038252.1:1151812-1159673  3075   GGCTTGAAAGTTCTGGGAATTTCATGTATCACTAACTTTGCGGCTGGTTTCCAAGAAGAACTCAATCACGAAGAAGTTGTAGAAGTGACT
NZ_CP035254.1:793365-801213    3132   GGCTTGAAAGTTCTGGGAATTTCATGTATCACTAACTTTGCGGCTGGTTTCCAAGAAGAACTCAATCACGAAGAAGTTGTAGAAGTGACT
NZ_CP035251.1:748733-756585    3131   GGCTTGAAAGTTCTGGGAATTTCATGTATCACTAACTTTGCGGCCGGTTTCCAAGAAGAACTCAATCACGAAGAAGTTGTAGAAGTGACT
NZ_CP035252.1:745670-753536    3145   GGCTTGAAAGTTCTGGGAATTTCATGTATCACTAACTTTGCGGCCGGTTTCCAAGAAGAACTCAATCACGAAGAAGTTGTAGAAGTGACT
NZ_CP035257.1:776317-784169    3131   GGCTTGAAAGTTCTAGGAATTTCATGTATCACTAACTTTGCGGCTGGTTTCCAAGAAGAACTCAATCACGAAGAAGTTGTAGAAGTGACT
NZ_CP038253.1:758105-765957    3131   GGCTTGAAAGTTCTAGGAATTTCATGTATCACTAACTTTGCGGCTGGTTTCCAAGAAGAACTCAATCACGAAGAAGTTGTAGAAGTGACT
NZ_CP035263.1:743050-750901    3131   GGCTTGAAAGTTCTGGGAATTTCATGTATCACTAACTTTGCGGCCGGTTTCCAAGAAGAACTCAATCACGAAGAAGTTGTAGAAGTGACT
NZ_CP035262.1:741595-749460    3145   GGCTTGAAAGTTCTGGGAATTTCATGTATCACTAACTTTGCGGCCGGTTTCCAAGAAGAACTCAATCACGAAGAAGTTGTAGAAGTGACT
NZ_CP035236.1:764915-772766    3131   GGCTTGAAAGTTCTGGGAATTTCATGTATCACTAACTTTGCGGCCGGTTTCCAAGAAGAACTCAATCACGAAGAAGTTGTAGAAGTGACT
NZ_CP035253.1 767286-775134    3132   GGCTTGAAAGTTCTGGGAATTTCATGTATCACTAACTTTGCGGCTGGTTTCCAAGAAGAACTCAATCACGAAGAAGTTGTAGAAGTGACT
NZ_CP035249.1:763080-770921    3132   GGCTTGAAAGTTCTGGGAATTTCATGTATCACTAACTTTGCGGCCGGTTTCCAAGAAGAACTCAATCACGAAGAAGTTGTAGAAGTGACT

NC_003028.3:777000-787000      3951   GAACGTGTTAAAGGTGATTTCAAAGGCTTGCTTAAAGCGATTCTTGCTGAATTGTAAGAAAAAAGATTTAAAAGGGGGAGTGCCTCTGTT
NZ_CP035248.1:765099-775111    3960   GAACGTGTTAAAGGTGATTTCAAAGGCTTGCTTAAAGCGATTCTTGCTGAATTGTAAGAAAAAAGATTTAAAAGGGGGAGTGCCTCTGTT
NZ_CP035235.1 c1326834-1317894 3236   GAACGTGTTAAAGGTGGTTTCAAAGGCTTGCTTAAAGCGATTCTTGCTGAATTGTAAGAAAAAAGATTTAAAAGGGGGAGTGCCTCTGTC
NZ_CP035234.1:756284-765224    3236   GAACGTGTTAAAGGTGGTTTCAAAGGCTTGCTTAAAGCGATTCTTGCTGAATTGTAAGAAAAAAGATTTAAAAGGGGGAGTGCCTCTGTC
NZ_CP038251.1:750745-759686    3236   GAACGTGTTAAAGGTGGTTTCAAAGGCTTGCTTAAAGCGATTCTTGCTGAATTGTAAGAAAAAAGATTTAAAAGGGGGAGTGCCTCTGTC
NZ_CP035244.1:747023-755950    3222   GAACGTGTTAAAGGTGGTTTCAAAGGCTTGCTTAAAGCGATTCTTGCTGAATTGTAAGAAAAAAGATTTAAAAGGGGGAGTGCCTCTGTC
NZ_CP035264.1:800836-809336    3222   GAACGTGTTAAAGGTGATTTCAAAGGCTTGCTTAAAGCGATTCTTGCTGAATTGTAAGAAAAAAGATTTAAAAGGGGAAGTGCCTCTGTT
NZ_CP035265.1:766128-774627    3222   GAACGTGTTAAAGGTGATTTCAAAGGCTTGCTTAAAGCGATTCTTGCTGAATTGTAAGAAAAAAGATTTAAAAGGGGAAGTGCCTCTGTT
NZ_CP035245.1 745261-753757    3222   GAACGTGTTAAAGGTGATTTCAAAGGCTTGCTTAAAGCGATTCTTGCTGAATTGTAAGAAAAAAGATTTAAAAGGGGGAGTGCCTCTGTC
NZ_CP035259.1:745700-753548    3222   GAACGTGTTAAAGGTGATTTTAAAGGCTTGCTTAAAGCGATTCTTGCTGAATTGTAAGAAAAAAGATTTAAAAGGGGGAGTGCCTCTGTC
NZ_CP035258.1:750874-758722    3222   GAACGTGTTAAAGGTGATTTTAAAGGCTTGCTTAAAGCGATTCTTGCTGAATTGTAAGAAAAAAGATTTAAAAGGGGGAGTGCCTCTGTC
NZ_CP035260.1:726156-733994    3222   GAACGTGTTAAAGGTGATTTCAAAGGCTTGCTTAAAGCGATTCTTGCTGAATTGTAAGAAAAAAGATTTAAAAGGGGGAGTGCCTCTGTC
NZ_CP035241.1:708296-716144    3222   GAACGTGTTAAAGGTGATTTCAAAGGCTTGCTTAAAGCGATTCTTGCTGAATTGTAAGAAAAAAGATTTAAAAGGGGGAGTGCCTCTGTC
NZ_CP035261.1:779086-786938    3222   GAACGTGTTAAAGGTGATTTCAAAGGCTTGCTTAAAGCGATTCTTGCTGAATTGTAAGAAAAAAGATTTAAAAGGGGGAGTGCCTCTGTT
NZ_CP035242.1:c1532236-1524385 3221   GAACGTGTTAAAGGTGATTTCAAAGGCTTGCTTAAAGCGATTCTTGCTGAATTGTAAGAAAAAAGATTTAAAAGGGGGAGTGCCTCTGTT
NZ_CP035243.1:777729-785580    3221   GAACGTGTTAAAGGTGATTTCAAAGGCTTGCTTAAAGCGATTCTTGCTGAATTGTAAGAAAAAAGATTTAAAAGGGGGAGTGCCTCTGTT
NZ_CP035237.1:c1311322-1303456 3236   GAACGTGTTAAAGGTGATTTCAAAGGCTTGCTTAAAGCGATTCTTGCTGAATTGTAAGAAAAAAGATTTAAAAGGGGTAGTGTCTCTGTC
NZ_CP035246.1:c1302892-1295026 3236   GAACGTGTTAAAGGTGATTTCAAAGGCTTGCTTAAAGCGATTCTTGCTGAATTGTAAGAAAAAAGATTTAAAAGGGGGAGTGCCTCTGTC
NZ_CP035247.1:c1301115-1293263 3222   GAACGTGTTAAAGGTGATTTCAAAGGCTTGCTTAAAGCGATTCTTGCTGAATTGTAAGAAAAAAGATTTAAAAGGGGGAGTGCCTCTGTC
NZ_CP035238.1:738547-746413    3236   GAACGTGTTAAAGGTGATTTCAAAGGCTTGCTTAAAGCGATTCTTGCTGAATTGTAAGAAAAAAGATTTAAAAGGGGGAGTGCCTCTGTC
NZ_CP035255.1:776866-784718    3222   GAACGTGTTAAAGGTGATTTCAAAGGCTTGCTTAAAGCGATTCTTGCTGAATTGTAAGAAAAAAGATTTAAAAGGGGGAGTGCCTCTGTC
NZ_CP035256.1:807236-815102    3236   GAACGTGTTAAAGGTGATTTCAAAGGCTTGCTTAAAGCGATTCTTGCTGAATTGTAAGAAAAAAGATTTAAAAGGGGGAGTGCCTCTGTC
NZ_CP038252.1:1151812-1159673  3165   GAACGTGTTAAAGGTGATTTCAAAGGCTTGCTTAAAGCGATTCTTGCTGAATTGTAAGAAAAAAGATTTAAAAGGGGGAGTGCCTCTGTC
NZ_CP035254.1:793365-801213    3222   GAACGTGTTAAAGGTGATTTCAAAGGCTTGCTTAAAGCGATTCTTGCTGAATTGTAAGAAAAAAGATTTAAAAGGGGGAGTGCCTCTGTC
```

### Ozkan et al. Supplementary Figures

```
NZ_CP035251.1:748733-756585    3221   GAACGTGTTAAAGGTGATTTCAAAGGCTTGCTTAAAGCGATTCTTGCTGAATTGTAAGAAAAAAGATTTAAAAGGGGAAGTACCTCTGTT
NZ_CP035252.1:745670-753536    3235   GAACGTGTTAAAGGTGATTTCAAAGGCTTGCTTAAAGCGATTCTTGCTGAATTGTAAGAAAAAAGATTTAAAAGGGGAAGTACCTCTGTT
NZ_CP035257.1:776317-784169    3221   GAACGTGTTAAAGGTGATTTCAAAGGCTTGCTTAAAGCGATTCTTGCTGAATTGTAAGAAAAAAGATTTAAAAGGGGAAGTACCTCTGTT
NZ_CP038253.1:758105-765957    3221   GAACGTGTTAAAGGTGATTTCAAAGGCTTGCTTAAAGCGATTCTTGCTGAATTGTAAGAAAAAAGATTTAAAAGGGGAAGTACCTCTGTT
NZ_CP035263.1:743050-750901    3221   GAACGTGTTAAAGGTGATTTCAAAGGCTTGCTTAAAGCGATTCTTGCTGAATTGTAAGAAAAAAGATTTAAAAGGGGGAGTGCCTCTGTT
NZ_CP035262.1:741595-749460    3235   GAACGTGTTAAAGGTGATTTCAAAGGCTTGCTTAAAGCGATTCTTGCTGAATTGTAAGAAAAAAGATTTAAAAGGGGGAGTGCCTCTGTT
NZ_CP035236.1:764915-772766    3221   GAACGTGTTAAAGGTGATTTCAAAGGCTTGCTTAAAGCGATTCTTGCTGAATTGTAAGAAAAAAGATTTAAAAGGGGGAGTGCCTCTGTT
NZ_CP035253.1 767286-775134    3222   GAACGTGTTAAAGGTGATTTCAAAGGCTTGCTTAAAGCGATTCTTGCTGAATTGTAAGAAAAAAGATTTAAAAGGGGGAGTGCCTCTGTC
NZ_CP035249.1:763080-770921    3222   GAACGTGTTAAAGGTGATTTCAAAGGCTTGCTTAAAGCGATTCTTGCTGAATTGTAAGAAAAAAGATTTAAAAGGGGGAGTGCCTCTGTT

NC_003028.3:777000-787000      4041   TTTTCAGGATTGACTGCCTATCCGGATTAAAGAAGAAACAGAGGAATACTATGAGCTTCTTCCTGCTCTTATAACTGAAAGAAGCGGAAG
NZ_CP035248.1:765099-775111    4050   TTTTCAGGATTGACTGCCTATCCGGATTAAAGAAGAAACAGAGGAATACTATGAGCTTCTTCCTGCTCTTATAACTGAAAGAAGCGGAAG
NZ_CP035235.1 c1326834-1317894 3326   TTTTCAGGATTGACTGCCTATCCGTATTAAAGAAGAAACAGAGGAATACTATGAGCTTCTTCCTGCTCTTATAACTGAAAGAAGCGGAAG
NZ_CP035234.1:756284-765224    3326   TTTTCAGGATTGACTGCCTATCCGTATTAAAGAAGAAACAGAGGAATACTATGAGCTTCTTCCTGCTCTTATAACTGAAAGAAGCGGAAG
NZ_CP038251.1:750745-759686    3326   TTTTCAGGATTGACTGCCTATCCGTATTAAAGAAGAAACAGAGGAATACTATGAGCTTCTTCCTGCTCTTATAACTGAAAGAAGCGGAAG
NZ_CP035244.1:747023-755950    3312   TTTTCAGGATTGACTGCCTATCCGTATTAAAGAAGAAACAGAGGAATACTATGAGCTTCTTCCTGCTCTTATAACTGAAAGAAGCGGAAG
NZ_CP035264.1:800836-809336    3312   TTTTCAGGATTGACTGCCTATCCGGATTAAAGAAGAAACAGAGGAATACTATGAGCTTCTTCCTGCTCTTATAACTGAAAGAAGCGGAAG
NZ_CP035265.1:766128-774627    3312   TTTTCAGGATTGACTGCCTATCCGGATTAAAGAAGAAACAGAGGAATACTATGAGCTTCTTCCTGCTCTTATAACTGAAAGAAGCGGAAG
NZ_CP035245.1 745261-753757    3312   TTTTCAGGATTGACTGCCTATCCGGATTAAAGAAGAAACAGAGGAATACTATGAGCTTCTTCCTGCTCTTATAACTGAAAGAAGTGGAAG
NZ_CP035259.1:745700-753548    3312   TTTTCAGGATTGACTGCCTATCCGTATTAAAGAAGAAACAGAGGAATACTATGAGCTTCTTCCTGCTCTTATAACTGAAAGAAGCGGAAG
NZ_CP035258.1:750874-758722    3312   TTTTCAGGATTGACTGCCTATCCGTATTAAAGAAGAAACAGAGGAATACTATGAGCTTCTTCCTGCTCTTATAACTGAAAGAAGCGGAAG
NZ_CP035260.1:726156-733994    3312   TTTTCAGGATTGACTGCCTATCCGTATTAAAGAAGAAACAGAGGAATACTATGAGCTTCTTCCTGCTCTTATAACTGAAAGAAGCGGAAG
NZ_CP035241.1:708296-716144    3312   TTTTCAGGATTGACTGCCTATCCGTATTAAAGAAGAAACAGAGGAATACTATGAGCTTCTTCCTGCTCTTATAACTGAAAGAAGCGGAAG
NZ_CP035261.1:779086-786938    3312   TTTTCAGGATTGACTGCCTATCCGGATTAAAGAAGAAACAGAGGAATACTATGAGCTTCTTCCTGCTCTTATAACTGAAAGAAGCGGAAG
NZ_CP035242.1:c1532236-1524385 3311   TTTTCAGGATTGACTGCCTATCCGGATTAAAGAAGAAACAGAGGAATACTATGAGCTTCTTCCTGCTCTTATAACTGAAAGAAGCGGAAG
NZ_CP035243.1:777729-785580    3311   TTTTCAGGATTGACTGCCTATCCGGATTAAAGAAGAAACAGAGGAATACTATGAGCTTCTTCCTGCTCTTATAACTGAAAGAAGCGGAAG
NZ_CP035237.1:c1311322-1303456 3326   TTTTCAGGATTGACTGCCTATCCGTATTAAAGAAGAAACAGAGGAATACTATGAGCTTCTTCCTGCTCTTATAACTGAAAGAAGCGGAAG
NZ_CP035246.1:c1302892-1295026 3326   TTTTCAGGATTGACTGCCTATCCGTATTAAAGAAGAAACAGAGGAATACTATGAGCTTCTTCCTGCTCTTATAACTGAAAGAAGCGGAAG
NZ_CP035247.1:c1301115-1293263 3312   TTTTCAGGATTGACTGCCTATCCGTATTAAAGAAGAAACAGAGGAATACTATGAGCTTCTTCCTGCTCTTATAACTGAAAGAAGCGGAAG
NZ_CP035238.1:738547-746413    3326   TTTTCAGGATTGACTGCCTATCCGTATTAAAGAAGAAACAGAGGAATACTATGAGCTTCTTCCTGCTCTTATAACTGAAAGAAGCGGAAG
NZ_CP035255.1:776866-784718    3312   TTTTCAGGATTGACTGCCTATCCGTATTAAAGAAGAAACAGAGGAATACTATGAGCTTCTTCCTGCTCTTATAACTGAAAGAAGCGGAAG
NZ_CP035256.1:807236-815102    3326   TTTTCAGGATTGACTGCCTATCCGTATTAAAGAAGAAACAGAGGAATACTATGAGCTTCTTCCTGCTCTTATAACTGAAAGAAGCGGAAG
NZ_CP038252.1:1151812-1159673  3255   TTTTCAGGATTGATTGCCTATCCGTATTAAAGAAGAAACAGAGGAATACTATGAGCTTCTTCCTGCTCTTATAACTGAAAGAAGCGGAAG
NZ_CP035254.1:793365-801213    3312   TTTTCAGGATTGACTGCCTATCCGTATTAAAGAAGAAACAGAGGAATACTATGAGCTTCTTCCTGCTCTTATAACTGAAAGAAGCGGAAG
NZ_CP035251.1:748733-756585    3311   TTTTCAGGATTGACTGCCTATCCGGATTAAAGAAGAAACAGAGGAATACTATGAGCTTCTTCCTGCTCTTATAACTGAAAGAAGCGGAAG
NZ_CP035252.1:745670-753536    3325   TTTTCAGGATTGACTGCCTATCCGGATTAAAGAAGAAACAGAGGAATACTATGAGCTTCTTCCTGCTCTTATAACTGAAAGAAGCGGAAG
NZ_CP035257.1:776317-784169    3311   TTTTCAGGATTGACTGCCTATCCGGATTAAAGAAGAAACAGAGGAATACTATGAGCTTCTTCCTGCTCTTATAACTGAAAGAAGCGGAAG
NZ_CP038253.1:758105-765957    3311   TTTTCAGGATTGACTGCCTATCCGGATTAAAGAAGAAACAGAGGAATACTATGAGCTTCTTCCTGCTCTTATAACTGAAAGAAGCGGAAG
NZ_CP035263.1:743050-750901    3311   TTTTCAGGATTGACTGCCTATCCGGATTAAAGAAGAAACAGAGGAATACTATGAGCTTCTTCCTGCTCTTATAACTGAAAGAAGCGGAAG
NZ_CP035262.1:741595-749460    3325   TTTTCAGGATTGACTGCCTATCCGGATTAAAGAAGAAACAGAGGAATACTATGAGCTTCTTCCTGCTCTTATAACTGAAAGAAGCGGAAG
NZ_CP035236.1:764915-772766    3311   TTTTCAGGATTGACTGCCTATCCGGATTAAAGAAGAAACAGAGGAATACTATGAGCTTCTTCCTGCTCTTATAACTGAAAGAAGCGGAAG
NZ_CP035253.1 767286-775134    3312   TTTTCAGGATTGACTGCCTATCCGTATTAAAGAAGAAACAGAGGAATACTATGAGCTTCTTCCTGCTCTTATAACTGAAAGAAGCGGAAG
NZ_CP035249.1:763080-770921    3312   TTTTCAGGATTGACTGCCTATCCGGATTAAAGAAGAAACAGAGGAATACTATGAGCTTCTTCCTGCTCTTATAACTGAAAGAAGCGGAAG

NC_003028.3:777000-787000      4131   AATAGGTATGTCTGATCTGATAGCCAGCATTGTGAAAGACAAGATTCTAGGATACTAGCATTAGCTTCCTAGCCAAGCAGACTAGTATGA
NZ_CP035248.1:765099-775111    4140   AATAGGTATGTCTGATCTGATAGCCAGCATTGTGAAAGACAAGATTCTAGAATACTAGCATTAGCTTCCTAGCCAAGCAGACTAGTATGA
NZ_CP035235.1 c1326834-1317894 3416   AATAGGTATGTCTGATCTGATAGCCAGCATTGTGAAAGACAAGATTCTAGGATACTAGCATTAGCTTCCTAGCCAAGCAGACTAGTATGA
NZ_CP035234.1:756284-765224    3416   AATAGGTATGTCTGATCTGATAGCCAGCATTGTGAAAGACAAGATTCTAGGATACTAGCATTAGCTTCCTAGCCAAGCAGACTAGTATGA
NZ_CP038251.1:750745-759686    3416   AATAGGTATGTCTGATCTGATAGCCAGCATTGTGAAAGACAAGATTCTAGGATACTAGCATTAGCTTCCTAGCCAAGCAGACTAGTATGA
NZ_CP035244.1:747023-755950    3402   AATAGGTATGTCTGATCTGATAGCCAGCATTGTGAAAGACAAGATTCTAGGATACTAGCATTAGCTTCCTAGCCAAGCAGACTAGTATGA
NZ_CP035264.1:800836-809336    3402   AATAGGTATGTCTGATCTGATAGCCAGCATTGTGAAAGACAAGATTCTAGGATACTAGCATTAGCTTTCTAGCCAAGCAGACTAGTATGA
NZ_CP035265.1:766128-774627    3402   AATAGGTATGTCTGATCTGATAGCCAGCATTGTGAAAGACAAGATTCTAGGATACTAGCATTAGCTTTCTAGCCAAGCAGACTAGTATGA
NZ_CP035245.1 745261-753757    3402   AATAGGTATGTCTGATCTGATAGCCAGCATTGTGAAAGACAAGATTCTAGGATACTAGCATTAGCTTCCTAGCCAAGTAGACTAGTATGA
NZ_CP035259.1:745700-753548    3402   AATAGGTGTGTCTGATCTGATAGCCAGCATTGTGAAAGACAAGATTCTAGGATACTAGCATTAGCTTCCTAGCCAAGCAGACTAGTATGA
NZ_CP035258.1:750874-758722    3402   AATAGGTGTGTCTGATCTGATAGCCAGCATTGTGAAAGACAAGATTCTAGGATACTAGCATTAGCTTCCTAGCCAAGCAGACTAGTATGA
NZ_CP035260.1:726156-733994    3402   AATAGGTGTGTCTGATCTGATAGCCAGCATTGTGAAAGACAAGATTCTAGGATACTAGCATTAGCTTCCTAGCCAAGCAGACTAGTATGA
NZ_CP035241.1:708296-716144    3402   AATAGGTGTGTCTGATCTGATAGCCAGCATTGTGAAAGACAAGATTCTAGGATACTAGCATTAGCTTCCTAGCCAAGCAGACTAGTATGA
NZ_CP035261.1:779086-786938    3402   AATAGGTATGTCTGATCTGATAGCCAGCATTGTGAAAGACAAGATTCTAGGATACTAGCATTAGCTTCCTAGCCAAGCAGACTAGTATGA
NZ_CP035242.1:c1532236-1524385 3401   AATAGGTATGTCTGATCTGATAGCCAGCATTGTGAAAGACAAGATTCTAGGATACTAGCATTAGCTTCCTAGCCAAGCAGACTAGTATGA
NZ_CP035243.1:777729-785580    3401   AATAGGTATGTCTGATCTGATAGCCAGCATTGTGAAAGACAAGATTCTAGGATACTAGCATTAGCTTCCTAGCCAAGCAGACTAGTATGA
NZ_CP035237.1:c1311322-1303456 3416   AATAGGTATGTCTGATCTGATAGCCAGCATTGTGAAAGACAAGATTCTAGGATACTAGCATTAGCTTTCTAGCCAAGCAGACTAGTATGA
NZ_CP035246.1:c1302892-1295026 3416   AATAGGTATGTCTGATCTGATAGCCAGCATTGTGAAAGACAAGATTCTAGGATACTAGCATTAGCTTCCTAGCCAAGCAGACTAGTATGA
NZ_CP035247.1:c1301115-1293263 3402   AATAGGTATGTCTGATCTGATAGCCAGCATTGTGAAAGACAAGATTCTAGGATACTAGCATTAGCTTCCTAGCCAAGCAGACTAGTATGA
NZ_CP035238.1:738547-746413    3416   AATAGGTATGTCTGATCTGATAGCCAGCATTGTGAAAGACAAGATTCTAGGATACTAGCATTAGCTTCCTAGCCAAGCAGACTAGTATGA
NZ_CP035255.1:776866-784718    3402   AATAGGTATGTCTGATCTGATAGCCAGCATTGTGAAAGACAAGATTCTAGGATACTAGCATTAGCTTCCTAGCCAAGCAGACTAGTATGA
NZ_CP035256.1:807236-815102    3416   AATAGGTATGTCTGATCTGATAGCCAGCATTGTGAAAGACAAGATTCTAGGATACTAGCATTAGCTTCCTAGCCAAGCAGACTAGTATGA
NZ_CP038252.1:1151812-1159673  3345   AATAGGTATGTCTGATCTGATAGCCAGCATTGTGAAAGACAAGATTCTAGGATACTAGCATTAGCTTCCTAGCCAAGCAGACTAGTATGA
NZ_CP035254.1:793365-801213    3402   AATAGGTATGTCTGATCTGATAGCCAGCATTGTGAAAGACAAGATTCTAGGATACTAGCATTAGCTTCCTAGCCAAGCAGACTAGTATGA
NZ_CP035251.1:748733-756585    3401   AATAGGTATGTCTGATCTGATAGCCAGCATTGTGAAAGACAAGATTCTAGGATACTAGCATTAGCTTCCTAGCCAAGCAGACTAGTATGA
NZ_CP035252.1:745670-753536    3415   AATAGGTATGTCTGATCTGATAGCCAGCATTGTGAAAGACAAGATTCTAGGATACTAGCATTAGCTTCCTAGCCAAGCAGACTAGTATGA
NZ_CP035257.1:776317-784169    3401   AATAGGTATGTCTGATCTGATAGCCAGCATTGTGAAAGACAAGATTCTAGGATACTAGCATTAGCTTCCTAGCCAAGCAGACTAGTATGA
NZ_CP038253.1:758105-765957    3401   AATAGGTATGTCTGATCTGATAGCCAGCATTGTGAAAGACAAGATTCTAGGATACTAGCATTAGCTTCCTAGCCAAGCAGACTAGTATGA
NZ_CP035263.1:743050-750901    3401   AATAGGTGTGTCTGATCTGATAGCCAGCATTGTGAAAGACAAGATTCTAGGATACTAGCATTAGCTTCCTAGCCAAGTAGACTAGTATGA
NZ_CP035262.1:741595-749460    3415   AATAGGTGTGTCTGATCTGATAGCCAGCATTGTGAAAGACAAGATTCTAGGATACTAGCATTAGCTTCCTAGCCAAGTAGACTAGTATGA
NZ_CP035236.1:764915-772766    3401   AATAGGTGTGTCTGATCTGATAGCCAGCATTGTGAAAGACAAGATTCTAGGATACTAGCATTAGCTTCCTAGCCAAGTAGACTAGTATGA
NZ_CP035253.1 767286-775134    3402   AATAGGTATGTCTGATCTGATAGCCAGCATTGTGAAAGACAAGATTCTAGGATACTAGCATTAGCTTCCTAGCCAAGCAGACTAGTATGA
NZ_CP035249.1:763080-770921    3402   AATAGGTATGTCTGATCTGATAGCCAGCATTGTGAAAGACAAGATTCTAGGATACTAGCATTAGCTTCCTAGCCAAGCAGACTAGTATGA

NC_003028.3:777000-787000      4221   TAAGGAGAGATGAGAATGAATTGACTTTCTGAATTTCTCAGTCTTATCATATATAGCACAATGAGATTTCGCTTGAGTCTGCTTGTAAAT
NZ_CP035248.1:765099-775111    4230   TAAGGAGAGATGAGAATGAATTGACTTTCTGAATTTCTCAGTCTTATCATATATAGCACAATGAGATTTCGCTTGAGTCTGCTTGTAAAT
NZ_CP035235.1 c1326834-1317894 3506   TAAGGAGAGATGAGAATGAATTGACTTTCTGAATTTCTCAGTCTTATCATATATAGCACAATGAGATTTCGCTTGAGTCTGCTTGTAAAT
NZ_CP035234.1:756284-765224    3506   TAAGGAGAGATGAGAATGAATTGACTTTCTGAATTTCTCAGTCTTATCATATATAGCACAATGAGATTTCGCTTGAGTCTGCTTGTAAAT
NZ_CP038251.1:750745-759686    3506   TAAGGAGAGATGAGAATGAATTGACTTTCTGAATTTCTCAGTCTTATCATATATAGCACAATGAGATTTCGCTTGAGTCTGCTTGTAAAT
NZ_CP035244.1:747023-755950    3492   TAAGGAGAGATGAGAATGAATTGACTTTCTGAATTTCTCAGTCTTATCATATATAGCACAATGAGATTTCGCTTGAGTCTGCTTGTAAAT
NZ_CP035264.1:800836-809336    3492   TAAGGAGAGATGAGAATGAATTGACTTTCTGAATTTCTCAGTCTTATCATATATAGCACAATGAGATTTCGCTTGAGTCTGCTTGTAAAT
NZ_CP035265.1:766128-774627    3492   TAAGGAGAGATGAGAATGAATTGACTTTCTGAATTTCTCAGTCTTATCATATATAGCACAATGAGATTTCGCTTGAGTCTGCTTGTAAAT
NZ_CP035245.1 745261-753757    3492   TAAGGAGAGATGAGAATGAATTGACTTTCTGAATTTCTTAGTCTTATCATATATAGCACAATGAGATTTCGCTTGAGTCTGCTTGTAAAT
NZ_CP035259.1:745700-753548    3492   TAAGGAGAGATGAGAATGAATTGACTTTCTGAATTTCTCAGTCTTATCATATATAGCACAATGAGATTTCGCTTGAGTCTGCTTGTAAAT
NZ_CP035258.1:750874-758722    3492   TAAGGAGAGATGAGAATGAATTGACTTTCTGAATTTCTCAGTCTTATCATATATAGCACAATGAGATTTCGCTTGAGTCTGCTTGTAAAT
NZ_CP035260.1:726156-733994    3492   TAAGGAGAGATGAGAATGAATTGACTTTCTGAATTTCTCAGTCTTATCATATATAGCACAATGAGATTTCGCTTGAGTCTGCTTGTAAAT
NZ_CP035241.1:708296-716144    3492   TAAGGAGAGATGAGAATGAATTGACTTTCTGAATTTCTCAGTCTTATCATATATAGCACAATGAGATTTCGCTTGAGTCTGCTTGTAAAT
NZ_CP035261.1:779086-786938    3492   TAAGGAGAGATGAGAATGAATTGACTTTCTGAATTTCTCAGTCTTATCATATATAGCACAATGAGATTTCGCTTGAGTCTGCTTGTAAAT
NZ_CP035242.1:c1532236-1524385 3491   TAAGGAGAGATGAGAATGAATTGACTTTCTGAATTTCTCAGTCTTATCATATATAGCACAATGAGATTTCGCTTGAGTCTGCTTGTAAAT
NZ_CP035243.1:777729-785580    3491   TAAGGAGAGATGAGAATGAATTGACTTTCTGAATTTCTCAGTCTTATCATATATAGCACAATGAGATTTCGCTTGAGTCTGCTTGTAAAT
NZ_CP035237.1:c1311322-1303456 3506   TAAGGAGAGATGAGAATGAATTGACTTTCTGAATTTCTCAGTCTTATCATATATAGCACAATGAGATTTCGCTTGAGTCTGCTTGTAAAT
```

### Ozkan et al. Supplementary Figures

```
NZ_CP035246.1:c1302892-1295026 3506   TAAGGAGAGATGAGAATGAATTGACTTTCTGAATTTCTCAGTCTTATCATATATAGCACAATGAGATTTCGCTTGAGTCTGCTTGTAAAT
NZ_CP035247.1:c1301115-1293263 3492   TAAGGAGAGATGAGAATGAATTGACTTTCTGAATTTCTCAGTCTTATCATATATAGCACAATGAGATTTCGCTTGAGTCTGCTTGTAAAT
NZ_CP035238.1:738547-746413    3506   TAAGGAGAGATGAGAATGAATTGACTTTCTGAATTTCTCAGTCTTATCATATATAGCACAATGAGATTTCGCTTGAGTCTGCTTGTAAAT
NZ_CP035255.1:776866-784718    3492   TAAGGAGAGATGAGAATGAATTGACTTTCTGAATTTCTCAGTCTTATCATATATAGCACAATGAGATTTCGCTTGAGTCTGCTTGTAAAT
NZ_CP035256.1:807236-815102    3506   TAAGGAGAGATGAGAATGAATTGACTTTCTGAATTTCTCAGTCTTATCATATATAGCACAATGAGATTTCGCTTGAGTCTGCTTGTAAAT
NZ_CP038252.1:1151812-1159673  3435   TAAGGAGAGATGAGAATGAATTGACTTTCTGAATTTCTCAGTCTTATCATATATAGCACAATGAGATTTCGCTTGAGTCTGCTTGTAAAT
NZ_CP035254.1:793365-801213    3492   TAAGGAGAGATGAGAATGAATTGACTTTCTGAATTTCTCAGTCTTATCATATATAGCACAATGAGATTTCGCTTGAGTCTGCTTGTAAAT
NZ_CP035251.1:748733-756585    3491   TAAGGAGAGATGAGAATGAATTGACTTTCTGAATTTCTCAGTCTTATCATATATAGCACAATGAGATTTCGCTTGAGTCTGCTTGTAAAT
NZ_CP035252.1:745670-753536    3505   TAAGGAGAGATGAGAATGAATTGACTTTCTGAATTTCTCAGTCTTATCATATATAGCACAATGAGATTTCGCTTGAGTCTGCTTGTAAAT
NZ_CP035257.1:776317-784169    3491   TAAGGAGAGATGAGAATGAATTGACTTTCTGAATTTCTCAGTCTTATCATATATAGCACAATGAGATTTCGCTTGAGTCTGCTTGTAAAT
NZ_CP038253.1:758105-765957    3491   TAAGGAGAGATGAGAATGAATTGACTTTCTGAATTTCTCAGTCTTATCATATATAGCACAATGAGATTTCGCTTGAGTCTGCTTGTAAAT
NZ_CP035263.1:743050-750901    3491   TAAGGAGAGATGAGAATGAATTGACTTTCTGAATTTCTCAGTCTTATCATATATAGCACAATGAGATTTCGCTTGAGTCTGCTTGTAAAT
NZ_CP035262.1:741595-749460    3505   TAAGGAGAGATGAGAATGAATTGACTTTCTGAATTTCTCAGTCTTATCATATATAGCACAATGAGATTTCGCTTGAGTCTGCTTGTAAAT
NZ_CP035236.1:764915-772766    3491   TAAGGAGAGATGAGAATGAATTGACTTTCTGAATTTCTCAGTCTTATCATATATAGCACAATGAGATTTCGCTTGAGTCTGCTTGTAAAT
NZ_CP035253.1 767286-775134    3492   TAAGGAGAGATGAGAATGAATTGACTTTCTGAATTTCTCAGTCTTATCATATATAGCACAATGAGATTTCGCTTGAGTCTGCTTGTAAAT
NZ_CP035249.1:763080-770921    3492   TAAGGAGAGATGAGAATGAATTGACTTTCTGAATTTCTCAGTCTTATCATATATAGCACAATGAGATTTCGCTTGAGTCTGCTTGTAAAT

NC_003028.3:777000-787000      4311   AAACGAAAAGAAAGATAAGAAATAATGAAAATTGGTCAACGAATTATGCGCTTTGGCATAAAAAATTAAGTATCGGAGTTGTATCTGTTG
NZ_CP035248.1:765099-775111    4320   AAACGAAAAGAAAGATAAGAAATAATGAAAATTGGTCAACGAATTATGCGCTTTGGCATAAAAAATTAAGTATCGGAGTTGTATCTGTTG
NZ_CP035235.1 c1326834-1317894 3596   AAACGAAAAGAAAGATAAGAAATAATGAAAATTGGTCAACGAATTATGCGCTTTGGCATAAAAAATTAAGTATCGGAGTTGTATCTGTTG
NZ_CP035234.1:756284-765224    3596   AAACGAAAAGAAAGATAAGAAATAATGAAAATTGGTCAACGAATTATGCGCTTTGGCATAAAAAATTAAGTATCGGAGTTGTATCTGTTG
NZ_CP038251.1:750745-759686    3596   AAACGAAAAGAAAGATAAGAAATAATGAAAATTGGTCAACGAATTATGCGCTTTGGCATAAAAAATTAAGTATCGGAGTTGTATCTGTTG
NZ_CP035244.1:747023-755950    3582   AAACGAAAAGAAAGATAAGAAATAATGAAAATTGGTCAACGAATTATGCGCTTTGGCATAAAAAATTAAGTATCGGAGTTGTATCTGTTG
NZ_CP035264.1:800836-809336    3582   AAACGAAAAGAAAGATAAGAAATAATGAAAATTGGTCAACGAATTATGCGCTTTGGCATAAAAAATTAAGTATCGGAGTTGTATCTGTTG
NZ_CP035265.1:766128-774627    3582   AAACGAAAAGAAAGATAAGAAATAATGAAAATTGGTCAACGAATTATGCGCTTTGGCATAAAAAATTAAGTATCGGAGTTGTATCTGTTG
NZ_CP035245.1 745261-753757    3582   AAACGAAAAGAAAGATAAGAAATAATGAAAATTGGTCAACGAATTATGCGCTTTGGCATAAAAAATTAAGTATCGGAGTTGTATCTGTTG
NZ_CP035259.1:745700-753548    3582   AAACGAAAAGAAAGATAAGAAATAATGAAAATTGGTCAACGAATTATGCGCTTTGGCATAAAAAATTAAGTATCGGAGTTGTATCTGTTG
NZ_CP035258.1:750874-758722    3582   AAACGAAAAGAAAGATAAGAAATAATGAAAATTGGTCAACGAATTATGCGCTTTGGCATAAAAAATTAAGTATCGGAGTTGTATCTGTTG
NZ_CP035260.1:726156-733994    3582   AAACGAAAAGAAAGATAAGAAATAATGAAAATTGGTCAACGAATTATGCGCTTTGGCATAAAAAATTAAGTATCGGAGTTGTATCTGTTG
NZ_CP035241.1:708296-716144    3582   AAACGAAAAGAAAGATAAGAAATAATGAAAATTGGTCAACGAATTATGCGCTTTGGCATAAAAAATTAAGTATCGGAGTTGTATCTGTTG
NZ_CP035261.1:779086-786938    3582   AAACGAAAAGAAAGATAAGAAATAATGAAAATTGGTCAACGAATTATGCGCTTTGGCATAAAAAATTAAGTATCGGAGTTGTATCTGTTG
NZ_CP035242.1:c1532236-1524385 3581   AAACGAAAAGAAAGATAAGAAATAATGAAAATTGGTCAACGAATTATGCGCTTTGGCATAAAAAATTAAGTATCGGAGTTGTATCTGTTG
NZ_CP035243.1:777729-785580    3581   AAACGAAAAGAAAGATAAGAAATAATGAAAATTGGTCAACGAATTATGCGCTTTGGCATAAAAAATTAAGTATCGGAGTTGTATCTGTTG
NZ_CP035237.1:c1311322-1303456 3596   AAACGAAAAGAAAGATAATAAATAATGAAAATTGGTCAACGAATTATGCGCTTTGGCATAAAAAATTAAGTATCGGAGTTGTATCTGTTG
NZ_CP035246.1:c1302892-1295026 3596   AAACGAAAAGAAAGATAAGAAATAATGAAAATTGGTCAACGAATTATGCGCTTTGGCATAAAAAATTAAGTATCGGAGTTGTATCTGTTG
NZ_CP035247.1:c1301115-1293263 3582   AAACGAAAAGAAAGATAAGAAATAATGAAAATTGGTCAACGAATTATGCGCTTTGGCATAAAAAATTAAGTATCGGAGTTGTATCTGTTG
NZ_CP035238.1:738547-746413    3596   AAACGAAAAGAAAGATAAGAAATAATGAAAATTGGTCAACGAATTATGCGCTTTGGCATAAAAAATTAAGTATCGGAGTTGTATCTGTTG
NZ_CP035255.1:776866-784718    3582   AAACGAAAAGAAAGATAAGAAATAATGAAAATTGGTCAACGAATTATGCGCTTTGGCATAAAAAATTAAGTATCGGAGTTGTATCTGTTG
NZ_CP035256.1:807236-815102    3596   AAACGAAAAGAAAGATAAGAAATAATGAAAATTGGTCAACGAATTATGCGCTTTGGCATAAAAAATTAAGTATCGGAGTTGTATCTGTTG
NZ_CP038252.1:1151812-1159673  3525   AAATGAAAAGAAAGATAAGAAATAATGAAAATTGGTCAACGAATTATGCGCTTTGGCATAAAAAATTAAGTATCGGAGTTGTATCTGTTG
NZ_CP035254.1:793365-801213    3582   AAACGAAAAGAAAGATAAGAAATAATGAAAATTGGTCAACGAATTATGCGCTTTGGCATAAAAAATTAAGTATCGGAGTTGTATCTGTTG
NZ_CP035251.1:748733-756585    3581   AAACGAAAAGAAAGATAAGAAATAATGAAAATTGGTCAACGAATTATGCGCTTTGGCATAAAAAATTAAGTATCGGAGTTGTATCTGTTG
NZ_CP035252.1:745670-753536    3595   AAACGAAAAGAAAGATAAGAAATAATGAAAATTGGTCAACGAATTATGCGCTTTGGCATAAAAAATTAAGTATCGGAGTTGTATCTGTTG
NZ_CP035257.1:776317-784169    3581   AAACGAAAAGAAAGATAAGAAATAATGAAAATTGGTCAACGAATTATGCGCTTTGGCATAAAAAATTAAGTATCGGAGTTGTATCTGTTG
NZ_CP038253.1:758105-765957    3581   AAACGAAAAGAAAGATAAGAAATAATGAAAATTGGTCAACGAATTATGCGCTTTGGCATAAAAAATTAAGTATCGGAGTTGTATCTGTTG
NZ_CP035263.1:743050-750901    3581   AAACGAAAAGAAAGATAAGAAATAATGAAAATTGGTCAACGAATTATGCGCTTTGGCATAAAAAATTAAGTATCGGAGTTGTATCTGTTG
NZ_CP035262.1:741595-749460    3595   AAACGAAAAGAAAGATAAGAAATAATGAAAATTGGTCAACGAATTATGCGCTTTGGCATAAAAAATTAAGTATCGGAGTTGTATCTGTTG
NZ_CP035236.1:764915-772766    3581   AAACGAAAAGAAAGATAAGAAATAATGAAAATTGGTCAACGAATTATGCGCTTTGGCATAAAAAATTAAGTATCGGAGTTGTATCTGTTG
NZ_CP035253.1 767286-775134    3582   AAACGAAAAGAAAGATAAGAAATAATGAAAATTGGTCAACGAATTATGCGCTTTGGCATAAAAAATTAAGTATCGGAGTTGTATCTGTTG
NZ_CP035249.1:763080-770921    3582   AAACGAAAAGAAAGATAAGAAATAATGAAAATTGGTCAACGAATTATGCGCTTTGGCATAAAAAATTAAGTATCGGAGTTGTATCTGTTG

NC_003028.3:777000-787000      4401   TAGTCGGCTTTGATTTCTAGCTCCAGCTGGAATTTCAGCCAATGAAGTAAAGCAAGATGTAACATCTGAAGTGGTAATAGGTGTGCTAGA
NZ_CP035248.1:765099-775111    4410   TAGTCGGCTTTGATTTCTAGCTCCAGCTGGAATTTCAGCCAATGAAGTAAAGCAAAATGTAACATCTGAAGTGGTAATAGGTGTGCTAGA
NZ_CP035235.1 c1326834-1317894 3686   TAGTCGGCTTTGATTTCTAGCTCCAGCTGGAATTTCAGCCAATGAAGTAAAGCAAGATGTAACATCTGAAGTGGTAATAGGTGTGCTAGA
NZ_CP035234.1:756284-765224    3686   TAGTCGGCTTTGATTTCTAGCTCCAGCTGGAATTTCAGCCAATGAAGTAAAGCAAGATGTAACATCTGAAGTGGTAATAGGTGTGCTAGA
NZ_CP038251.1:750745-759686    3686   TAGTCGGCTTTGATTTCTAGCTCCAGCTGGAATTTCAGCCAATGAAGTAAAGCAAGATGTAACATCTGAAGTGGTAATAGGTGTGCTAGA
NZ_CP035244.1:747023-755950    3672   TAGTCGGCTTTGATTTCTAGCTCCAGCTGGAATTTCAGCCAATGAAGTAAAGCAAGATGTAACATCTGAAGTGGTAATAGGTGTGCTAGA
NZ_CP035264.1:800836-809336    3672   TAGTCGGCTTTGATTTCTAGCTCCAGCTGGAATTTCAGCCAATGAAGTAAAGCAAGATGTAACATTTGAAGTGGTAATAGGTGTGCTAGA
NZ_CP035265.1:766128-774627    3672   TAGTCGGCTTTGATTTCTAGCTCCAGCTGGAATTTCAGCCAATGAAGTAAAGCAAGATGTAACATTTGAAGTGGTAATAGGTGTGCTAGA
NZ_CP035245.1 745261-753757    3672   TAGTCGGCTTTGATTTCTAGCTCCAGCTGGAATTTCAGCCAATGAAGTAAAGCAAGATGTAACATCTGAAGTGGTAATAGGTGTGCTAGA
NZ_CP035259.1:745700-753548    3672   TAGTCGGCTTTGATTTCTAGCTCCAGCTGGAATTTCAGCCAATGAAGTAAAGCAAGATGTAACATCTGAAGTGGTAATAGGTGTGCTAGA
NZ_CP035258.1:750874-758722    3672   TAGTCGGCTTTGATTTCTAGCTCCAGCTGGAATTTCAGCCAATGAAGTAAAGCAAGATGTAACATCTGAAGTGGTAATAGGTGTGCTAGA
NZ_CP035260.1:726156-733994    3672   TAGTCGGCTTTGATTTCTAGCTCCAGCTGGAATTTCAGCCAATGAAGTAAAGCAAGATGTAACATCTGAAGTGGTAATAGGTGTGCTAGA
NZ_CP035241.1:708296-716144    3672   TAGTCGGCTTTGATTTCTAGCTCCAGCTGGAATTTCAGCCAATGAAGTAAAGCAAGATGTAACATCTGAAGTGGTAATAGGTGTGCTAGA
NZ_CP035261.1:779086-786938    3672   TAGTCGGCTTTGATTTCTAGCTCCAGCTGGAATTTCAGCCAATGAAGTAAAGCAAGATGTAACATCTGAAGTGGTAATAGGTGTGCTAGA
NZ_CP035242.1:c1532236-1524385 3671   TAGTTGGCTTTGATTTCTAGCTCCAGCTGGAATTTCAGCCAATGAAGTAAAGCAAGATGTAACATCTGAAGTGGTAATAGGTGTGCTAGA
NZ_CP035243.1:777729-785580    3671   TAGTTGGCTTTGATTTCTAGCTCCAGCTGGAATTTCAGCCAATGAAGTAAAGCAAGATGTAACATCTGAAGTGGTAATAGGTGTGCTAGA
NZ_CP035237.1:c1311322-1303456 3686   TAGTCGGCTTTGATTTCTAGCTCCAGCTGGAATTTCAGCCAATGAAGTAAAGCAAGATGTAACATCTGAAGTGGTAATAGGTGTGCTAGA
NZ_CP035246.1:c1302892-1295026 3686   TAGTCGGCTTTGATTTCTAGCTCCAGCTGGAATTTCAGCCAATGAAGTAAAGCAAGATGTAACATCTGAAGTGGTAATAGGTGTGCTAGA
NZ_CP035247.1:c1301115-1293263 3672   TAGTCGGCTTTGATTTCTAGCTCCAGCTGGAATTTCAGCCAATGAAGTAAAGCAAGATGTAACATCTGAAGTGGTAATAGGTGTGCTAGA
NZ_CP035238.1:738547-746413    3686   TAGTCGGCTTTGATTTCTAGCTCCAGCTGGAATTTCAGCCAATGAAGTAAAGCAAGATGTAACATCTGAAGTGGTAATAGGTGTGCTAGA
NZ_CP035255.1:776866-784718    3672   TAGTCGGCTTTGATTTCTAGCTCCAGCTGGAATTTCAGCCAATGAAGTAAAGCAAGATGTAACATCTGAAGTGGTAATAGGTGTGCTAGA
NZ_CP035256.1:807236-815102    3686   TAGTCGGCTTTGATTTCTAGCTCCAGCTGGAATTTCAGCCAATGAAGTAAAGCAAGATGTAACATCTGAAGTGGTAATAGGTGTGCTAGA
NZ_CP038252.1:1151812-1159673  3615   TAGTCGGCTTTGATTTCTAGCTCCAGCTGGAATTTCAGCCAATGAAGTAAAGCAAGATGTAACATCTGAAGTGGTAATAGGTGTGCTAGA
NZ_CP035254.1:793365-801213    3672   TAGTCGGCTTTGATTTCTAGCTCCAGCTGGAATTTCAGCCAATGAAGTAAAGCAATATGTAACATCTGAAGTGGTAATAGGTGTGCTAGA
NZ_CP035251.1:748733-756585    3671   TAGTCGGCTTTGATTTCTAGCTCCAGCTGGAATTTCAGCCAATGAAGTAAAGTAAGATGTAACATCTGAAGTGGTAATAGGTGTGTTAGA
NZ_CP035252.1:745670-753536    3685   TAGTCGGCTTTGATTTCTAGCTCCAGCTGGAATTTCAGCCAATGAAGTAAAGTAAGATGTAACATCTGAAGTGGTAATAGGTGTGTTAGA
NZ_CP035257.1:776317-784169    3671   TAGTCGGCTTTGATTTCTAGCTCCAGCTGGAATTTCAGCCAATGAAGTAAAGTAAGATGTAACATCTGAAGTGGTAATAGGTGTGTTAGA
NZ_CP038253.1:758105-765957    3671   TAGTCGGCTTTGATTTCTAGCTCCAGCTGGAATTTCAGCCAATGAAGTAAAGTAAGATGTAACATCTGAAGTGGTAATAGGTGTGTTAGA
NZ_CP035263.1:743050-750901    3671   TAGTCGGCTTTGATTTCTAGCTCCAGCTGGAATTTCAGCCAATGAAGTAAAGCAAGATGTAACATCTGAAGTGGTAATAGGTGTGCTAGA
NZ_CP035262.1:741595-749460    3685   TAGTCGGCTTTGATTTCTAGCTCCAGCTGGAATTTCAGCCAATGAAGTAAAGCAAGATGTAACATCTGAAGTGGTAATAGGTGTGCTAGA
NZ_CP035236.1:764915-772766    3671   TAGTCGGCTTTGATTTCTAGCTCCAGCTGGAATTTCAGCCAATGAAGTAAAGCAAGATGTAACATCTGAAGTGGTAATAGGTGTGCTAGA
NZ_CP035253.1 767286-775134    3672   TAGTCGGCTTTGATTTCTAGCTCCAGCTGGAATTTCAGCCAATGAAGTAAAGCAATATGTAACATCTGAAGTGGTAATAGGTGTGCTAGA
NZ_CP035249.1:763080-770921    3672   TAGTCGGCTTTGATTTCTAGCTCCAGCTGGAATTTCAGCCAATGAAGTAAAGCAAGATGTAACATCTGAAGTGGTAATAGGTGTGCTAGA

NC_003028.3:777000-787000      4491   TTCTAAGGAGGAATTGAAAGAGTCAGAAAATGATGCTCCAAAACTAGAAACTCCTCTTAGAGAGGAGCCAAGACTAGCTCCTCAAACGCT
NZ_CP035248.1:765099-775111    4500   TTCTAAGGAGGAATTGAAAGAGTCAGAAAATGATGCTCCAAAACTAGAAACTCCTCTTAGAGAGGAGCCAAGACTAGCTCCTCAAACGCT
NZ_CP035235.1 c1326834-1317894 3776   TTCTAAGGAGGAATTGAAAGAGTCAGAAAATGATGCTCCAAAACTAGAAACTCCTCTTAGAGAGGAGCCAAGACTAGCTCCTCAAACGCT
NZ_CP035234.1:756284-765224    3776   TTCTAAGGAGGAATTGAAAGAGTCAGAAAATGATGCTCCAAAACTAGAAACTCCTCTTAGAGAGGAGCCAAGACTAGCTCCTCAAACGCT
NZ_CP038251.1:750745-759686    3776   TTCTAAGGAGGAATTGAAAGAGTCAGAAAATGATGCTCCAAAACTAGAAACTCCTCTTAGAGAGGAGCCAAGACTAGCTCCTCAAACGCT
NZ_CP035244.1:747023-755950    3762   TTCTAAGGAGGAATTGAAAGAGTCAGAAAATGATGCTCCAAAACTAGAAACTCCTCTTAGAGAGGAGCCAAGACTAGCTCCTCAAACGCT
NZ_CP035264.1:800836-809336    3762   TTCTAAGGAGGAATTGAAAGAGTCAGAAAATGATGCTCCAAAACTAGAAACTCCTCTTAGAGAGGAGCCAAGACTAGCTCCTCAAACGCT
NZ_CP035265.1:766128-774627    3762   TTCTAAGGAGGAATTGAAAGAGTCAGAAAATGATGCTCCAAAACTAGAAACTCCTCTTAGAGAGGAGCCAAGACTAGCTCCTCAAACGCT
NZ_CP035245.1 745261-753757    3762   TTCTAAGGAGGAATTGAAAGAGTCAGAAAATGATGCTCCAAAACTAGAAACTCCTCTTAGAGAGGAGCCAAGACTGGCTCCTCAAACGCT
NZ_CP035259.1:745700-753548    3762   TTCTAAGGAGGAATTGAAAGAGTCAGAAAATGATGCTCCAAAACTAGAAACTCCTCTTAGAGAGGAGCCAAGACTAGCTCCTCAAACGCT
```

```
NZ_CP035258.1:750874-758722    3762    TTCTAAGGAGGAATTGAAAGAGTCAGAAAATGATGCTCCAAAACTAGAAACTCCTCTTAGAGAGGAGCCAAGACTAGCTCCTCAAACGCT
NZ_CP035260.1:726156-733994    3762    TTCTAAGGAGGAATTGAAAGAGTCAGAAAATGATGCTCCAAAACTAGAAACTCCTCTTAGAGAGGAGCCAAGACTAGCTCCTCAAACGCT
NZ_CP035241.1:708296-716144    3762    TTCTAAGGAGGAATTGAAAGAGTCAGAAAATGATGCTCCAAAACTAGAAACTCCTCTTAGAGAGGAGCCAAGACTAGCTCCTCAAACGCT
NZ_CP035261.1:779086-786938    3762    TTCTAAGGAGGAATTGAAAGAGTCAGAAAATGATGCTCCAAAACTAGAAACTCCTCTTAGAGAGGAGCCAAGACTAGCTCCTCAAACGCT
NZ_CP035242.1:c1532236-1524385 3761    TTCTAAGGAGGAATTGAAAGAGTCAGAAAATGATGCTCCAAAACTAGAAACTCCTCTTAGAGAGGAGCCAAGACTAGCTCCTCAAACGCT
NZ_CP035243.1:777729-785580    3761    TTCTAAGGAGGAATTGAAAGAGTCAGAAAATGATGCTCCAAAACTAGAAACTCCTCTTAGAGAGGAGCCAAGACTAGCTCCTCAAACGCT
NZ_CP035237.1:c1311322-1303456 3776    TTCTAAGGAGGAATTGAAAGAGTCAGAAAATGATGCTCCAAAACTAGAAACTCCTCTTAGAGAGGAGCCAAGACTAGCTCCTCAAACGCT
NZ_CP035246.1:c1302892-1295026 3776    TTCTAAGGAGGAATTGAAAGAGTCAGAAAATGATGCTCCAAAACTAGAAACTCCTCTTAGAGAGGAGCCAAGACTAGCTCCTCAAACGCT
NZ_CP035247.1:c1301115-1293263 3762    TTCTAAGGAGGAATTGAAAGAGTCAGAAAATGATGCTCCAAAACTAGAAACTCCTCTTAGAGAGGAGCCAAGACTAGCTCCTCAAACGCT
NZ_CP035238.1:738547-746413    3776    TTCTAAGGAGGAATTGAAAGAGTCAGAAAATGATGCTCCAAAACTAGAAACTCCTCTTAGAGAGGAGCCAAGACTAGCTCCTCAAACGCT
NZ_CP035255.1:776866-784718    3762    TTCTAAGGAGGAATTGAAAGAGTCAGAAAATGATGCTCCAAAACTAGAAACTCCTCTTAGAGAGGAGCCAAGACTAGCTCCTCAAACGCT
NZ_CP035256.1:807236-815102    3776    TTCTAAGGAGGAATTGAAAGAGTCAGAAAATGATGCTCCAAAACTAGAAACTCCTCTTAGAGAGGAGCCAAGACTAGCTCCTCAAACGCT
NZ_CP038252.1:1151812-1159673  3705    TTCTAAGGAGGAATTGAAAGAGTCAGAAAATGATGCTCCAAAACTAGAAACTCCTCTTAGAGAGGAGCCAAGACTAGCTCCTCAAACGCT
NZ_CP035254.1:793365-801213    3762    TTCTAAGGAGGAATTGAAAGAGTCAGAAAATGATGCTCCAAAACTAGAAACTCCTCTTAGAGAGGAGCCAAGACTAGCTCCTCAAACGCT
NZ_CP035251.1:748733-756585    3761    TTCTAAGGAGGAATTGAAAGAGTCAGAAAATGATGCTCTAAAACTAGAAACTCCTCTTAGAGAGGAGCCAAGACTAGCTCCTCAAACGCT
NZ_CP035252.1:745670-753536    3775    TTCTAAGGAGGAATTGAAAGAGTCAGAAAATGATGCTCTAAAACTAGAAACTCCTCTTAGAGAGGAGCCAAGACTAGCTCCTCAAACGCT
NZ_CP035257.1:776317-784169    3761    TTCTAAGGAGGAATTGAAAGAGTCAGAAAATGATGCTCTAAAACTAGAAACTCCTCTTAGAGAGGAGCCAAGACTAGCTCCTCAAACGCT
NZ_CP038253.1:758105-765957    3761    TTCTAAGGAGGAATTGAAAGAGTCAGAAAATGATGCTCTAAAACTAGAAACTCCTCTTAGAGAGGAGCCAAGACTAGCTCCTCAAACGCT
NZ_CP035263.1:743050-750901    3761    TTCTAAGGAGGAATTGAAAGAGTCAGAAAATGATGCTCCAAAACTAGAAACTCCTCTTAGAGAGGAGCCAAGACTAGCTCCTCAAACGCT
NZ_CP035262.1:741595-749460    3775    TTCTAAGGAGGAATTGAAAGAGTCAGAAAATGATGCTCCAAAACTAGAAACTCCTCTTAGAGAGGAGCCAAGACTAGCTCCTCAAACGCT
NZ_CP035236.1:764915-772766    3761    TTCTAAGGAGGAATTGAAAGAGTCAGAAAATGATGCTCCAAAACTAGAAACTCCTCTTAGAGAGGAGCCAAGACTAGCTCCTCAAACGCT
NZ_CP035253.1 767286-775134    3762    TTCTAAGGAGGAATTGAAAGAGTCAGAAAATGATGCTCCAAAACTAGAAACTCCTCTTAGAGAGGAGCCAAGACTAGCTCCTCAAACGCT
NZ_CP035249.1:763080-770921    3762    TTCTAAGGAGGAATTGAAAGAGTCAGAAAATGATGCTCCAAAACTAGAAACTCCTCTTAGAGAGGAGCCAAGACTAGCTCCTCAAACGCT

NC_003028.3:777000-787000      4581    TCCGGAAGCAAGTGA----AGTTCTTGAAAACAAAAGGGAAGAGTCAAAAGTAGAGATAACATAACC-----------------------
NZ_CP035248.1:765099-775111    4590    TCCGGAAGCAAGTGA----AGTTCTTGAAAACAAAAGGGAAGAGTCAAAAGTAGAGATAACATAACC-----------------------
NZ_CP035235.1 c1326834-1317894 3866    TCCGGAAGCAAGTGA----AGTTCTTGAAAACAAAAGGGAAGAGTCAAAAGTAGAGATAACAGAGCC-----------------------
NZ_CP035234.1:756284-765224    3866    TCCGGAAGCAAGTGA----AGTTCTTGAAAACAAAAGGGAAGAGTCAAAAGTAGAGATAACAGAGCC-----------------------
NZ_CP038251.1:750745-759686    3866    TCCGGAAGCAAGTGA----AGTTCTTGAAAACAAAAGGGAAGAGTCAAAAGTAGAGATAACAGAGCC-----------------------
NZ_CP035244.1:747023-755950    3852    TCCGGAAGCAAGTGA----AGTTCTTGAAAACAAAAGGGAAGAGTCAAAAGTAGAGATAACAGAGCC-----------------------
NZ_CP035264.1:800836-809336    3852    TCCGGAAGCAAGTGA----AGTTCTTGAAAACAAAAGGGAAGAGTCAAAAGTAGAGATAACAGAGCCAGCTCAAGACTCACCTATTCTTG
NZ_CP035265.1:766128-774627    3852    TCCGGAAGCAAGTGA----AGTTCTTGAAAACAAAAGGGAAGAGTCAAAAGTAGAGATAACAGAGCCAGCTCAAGACTCACCTATTCTTG
NZ_CP035245.1 745261-753757    3852    TCTGGAAGCAAGTGA----AGTTCTTGAAAACAAAAGGGAAGAGTCAAAAGTAGGGATAACAGAGCCAGCTCAAGACTCACCTATTCTTG
NZ_CP035259.1:745700-753548    3852    TCCGGAAGCAAGTGA----AGTTCTTGAAAACAAAAGGGAAGAGTCAAAAGTAGAGATAACATAACC-----------------------
NZ_CP035258.1:750874-758722    3852    TCCGGAAGCAAGTGA----AGTTCTTGAAAACAAAAGGGAAGAGTCAAAAGTAGAGATAACATAACC-----------------------
NZ_CP035260.1:726156-733994    3852    TCCGGAAGCAAGTGA----AGTTCTTGAAAACAAAAGGGAAGAGTCAAAAGTAGAGATAACAGAGCC-----------------------
NZ_CP035241.1:708296-716144    3852    TCCGGAAGCAAGTGA----AGTTCTTGAAAACAAAAGGGAAGAGTCAAAAGTAGAGATAACAGAGCC-----------------------
NZ_CP035261.1:779086-786938    3852    TCCGGAAGCAAGTGA----AGTTCTTGAAAACAAAAGGGAAGAGTCAAAAGTAGAGATAACATAACC-----------------------
NZ_CP035242.1:c1532236-1524385 3851    TCCGGAAGCAAGTGA----AGTTCTTGAAAACAAAAGGGAAGAGTCAAAAGTAGAGATAACATAACC-----------------------
NZ_CP035243.1:777729-785580    3851    TCCGGAAGCAAGTGA----AGTTCTTGAAAACAAAAGGGAAGAGTCAAAAGTAGAGATAACATAACC-----------------------
NZ_CP035237.1:c1311322-1303456 3866    TCCGGAAGCAAGTGA----AGTTCTTGAAAACAAAAGGGAAGAGTCAAAAGTAGAGATAACAGAGCC-----------------------
NZ_CP035246.1:c1302892-1295026 3866    TCCGGAAGCAAGTGA----AGTTCTTGAAAACAAAAGGGAAGAGTCAAAAGTAGAGATAACAGAGCC-----------------------
NZ_CP035247.1:c1301115-1293263 3852    TCCGGAAGCAAGTGA----AGTTCTTGAAAACAAAAGGGAAGAGTCAAAAGTAGAGATAACAGAGCC-----------------------
NZ_CP035238.1:738547-746413    3866    TCCGGAAGCAAGTGA----AGTTCTTGAAAACAAAAGGGAAGAGTCAAAAGTAGAGATAACAGAGCC-----------------------
NZ_CP035255.1:776866-784718    3852    TCCGGAAGCAAGTGA----AGTTCTTGAAAACAAAAGGGAAGAGTCAAAAGTAGAGATAACAGAGCC-----------------------
NZ_CP035256.1:807236-815102    3866    TCCGGAAGCAAGTGA----AGTTCTTGAAAACAAAAGGGAAGAGTCAAAAGTAGAGATAACAGAGCC-----------------------
NZ_CP038252.1:1151812-1159673  3795    TCCGGAAGCAAGTGA----AGTTCTTGAAAACAAAAGGGAAGAGTCAAAAGTAGAGATAACAGAGCC-----------------------
NZ_CP035254.1:793365-801213    3852    TCCGGAAGCAAGTGA----AGTTCTTGAAAACAAAAGGGAAGAGTCAAAAGTAGAGATAACAGAGCC-----------------------
NZ_CP035251.1:748733-756585    3851    TCCGGAAGCAAGTGAAGTTAGTTCTTGAAAACAAAAGGGAAGAGTCAAAAGTAGAGATAACAGAACC-----------------------
NZ_CP035252.1:745670-753536    3865    TCCGGAAGCAAGTGAAGTTAGTTCTTGAAAACAAAAGGGAAGAGTCAAAAGTAGAGATAACAGAACC-----------------------
NZ_CP035257.1:776317-784169    3851    TCCGGAAGCAAGTGAAGTTAGTTCTTGAAAACAAAAGGGAAGAGTCAAAAGTAGAGATAACAGAACC-----------------------
NZ_CP038253.1:758105-765957    3851    TCCGGAAGCAAGTGAAGTTAGTTCTTGAAAACAAAAGGGAAGAGTCAAAAGTAGAGATAACAGAACC-----------------------
NZ_CP035263.1:743050-750901    3851    TCCGGAAGCAAGTGA----AGTTCTTGAAAACAAAAGGGAAGAGTCAAAAGTAGAGATAATAGAGCC-----------------------
NZ_CP035262.1:741595-749460    3865    TCCGGAAGCAAGTGA----AGTTCTTGAAAACAAAAGGGAAGAGTCAAAAGTAGAGATAATAGAGCC-----------------------
NZ_CP035236.1:764915-772766    3851    TCCGGAAGCAAGTGA----AGTTCTTGAAAACAAAAGGGAAGAGTCAAAAGTAGAGATAATAGAGCC-----------------------
NZ_CP035253.1 767286-775134    3852    TCCGGAAGCAAGTGA----AGTTCTTGAAAACAAAAGGGAAGAGTCAAAAGTAGAGATAACAGAGCC-----------------------
NZ_CP035249.1:763080-770921    3852    TCCGGAAGCAAGTGA----AGTTCTTGAAAACAAAAGGGAAGAGTCAAAAGTAGAGATAACATAACC-----------------------

NC_003028.3:777000-787000      4643    ------------------------------------------------------------------------------------------
NZ_CP035248.1:765099-775111    4652    ------------------------------------------------------------------------------------------
NZ_CP035235.1 c1326834-1317894 3928    ------------------------------------------------------------------------------------------
NZ_CP035234.1:756284-765224    3928    ------------------------------------------------------------------------------------------
NZ_CP038251.1:750745-759686    3928    ------------------------------------------------------------------------------------------
NZ_CP035244.1:747023-755950    3914    ------------------------------------------------------------------------------------------
NZ_CP035264.1:800836-809336    3938    CTCCTGTAGAAGAAACTAAAGAAGAAGCTGTGACAGAAAAACCAACAAATACTCGTTCTCTAACTGCAGAAGATTTGGTGAAGATTTCCA
NZ_CP035265.1:766128-774627    3938    CTCCTGTAGAAGAAACTAAAGAAGAAGCTGTGACAGAAAAACCAACAAATACTCGTTCTCTAACTGCAGAAGATTTGGTGAAGATTTCCA
NZ_CP035245.1 745261-753757    3938    CTCCTGTAGAAGAAACTAAAGAAGAAGCTGTGACAGAAAAACCAACAAATACTCGTTCTCTAACTGCAGAAGATTTGGTGAAGATTTCCA
NZ_CP035259.1:745700-753548    3914    ------------------------------------------------------------------------------------------
NZ_CP035258.1:750874-758722    3914    ------------------------------------------------------------------------------------------
NZ_CP035260.1:726156-733994    3914    ------------------------------------------------------------------------------------------
NZ_CP035241.1:708296-716144    3914    ------------------------------------------------------------------------------------------
NZ_CP035261.1:779086-786938    3914    ------------------------------------------------------------------------------------------
NZ_CP035242.1:c1532236-1524385 3913    ------------------------------------------------------------------------------------------
NZ_CP035243.1:777729-785580    3913    ------------------------------------------------------------------------------------------
NZ_CP035237.1:c1311322-1303456 3928    ------------------------------------------------------------------------------------------
NZ_CP035246.1:c1302892-1295026 3928    ------------------------------------------------------------------------------------------
NZ_CP035247.1:c1301115-1293263 3914    ------------------------------------------------------------------------------------------
NZ_CP035238.1:738547-746413    3928    ------------------------------------------------------------------------------------------
NZ_CP035255.1:776866-784718    3914    ------------------------------------------------------------------------------------------
NZ_CP035256.1:807236-815102    3928    ------------------------------------------------------------------------------------------
NZ_CP038252.1:1151812-1159673  3857    ------------------------------------------------------------------------------------------
NZ_CP035254.1:793365-801213    3914    ------------------------------------------------------------------------------------------
NZ_CP035251.1:748733-756585    3917    ------------------------------------------------------------------------------------------
NZ_CP035252.1:745670-753536    3931    ------------------------------------------------------------------------------------------
NZ_CP035257.1:776317-784169    3917    ------------------------------------------------------------------------------------------
NZ_CP038253.1:758105-765957    3917    ------------------------------------------------------------------------------------------
NZ_CP035263.1:743050-750901    3913    ------------------------------------------------------------------------------------------
NZ_CP035262.1:741595-749460    3927    ------------------------------------------------------------------------------------------
NZ_CP035236.1:764915-772766    3913    ------------------------------------------------------------------------------------------
NZ_CP035253.1 767286-775134    3914    ------------------------------------------------------------------------------------------
NZ_CP035249.1:763080-770921    3914    ------------------------------------------------------------------------------------------

NC_003028.3:777000-787000      4643    ------------------------------------------------------------------------------------------
NZ_CP035248.1:765099-775111    4652    ------------------------------------------------------------------------------------------
NZ_CP035235.1 c1326834-1317894 3928    ------------------------------------------------------------------------------------------
```

```
NZ_CP035234.1:756284-765224    3928   ------------------------------------------------------------------------------------------
NZ_CP038251.1:750745-759686    3928   ------------------------------------------------------------------------------------------
NZ_CP035244.1:747023-755950    3914   ------------------------------------------------------------------------------------------
NZ_CP035264.1:800836-809336    4028   AAGGGGAATTGCATTTAGAAAATGATTTGATTGATGAATCTTTCTATGGTGAAAAAGCTCTTGATTTGGAAGGGGATGATGACCAGGATG
NZ_CP035265.1:766128-774627    4028   AAGGGGAATTGCATTTAGAAAATGATTTGATTGATGAATCTTTCTATGGTGAAAAAGCTCTTGATTTGGAAGGGGATGATGACCAGGATG
NZ_CP035245.1 745261-753757    4028   AAGGGGAATTGCATTTAGAAAATGATTTGATTGATGAATCTTTCTATGGTGAAAAAGCTCTTGATTTGGAAGGGGATGATTACCAGGATG
NZ_CP035259.1:745700-753548    3914   ------------------------------------------------------------------------------------------
NZ_CP035258.1:750874-758722    3914   ------------------------------------------------------------------------------------------
NZ_CP035260.1:726156-733994    3914   ------------------------------------------------------------------------------------------
NZ_CP035241.1:708296-716144    3914   ------------------------------------------------------------------------------------------
NZ_CP035261.1:779086-786938    3914   ------------------------------------------------------------------------------------------
NZ_CP035242.1:c1532236-1524385 3913   ------------------------------------------------------------------------------------------
NZ_CP035243.1:777729-785580    3913   ------------------------------------------------------------------------------------------
NZ_CP035237.1:c1311322-1303456 3928   ------------------------------------------------------------------------------------------
NZ_CP035246.1:c1302892-1295026 3928   ------------------------------------------------------------------------------------------
NZ_CP035247.1:c1301115-1293263 3914   ------------------------------------------------------------------------------------------
NZ_CP035238.1:738547-746413    3928   ------------------------------------------------------------------------------------------
NZ_CP035255.1:776866-784718    3914   ------------------------------------------------------------------------------------------
NZ_CP035256.1:807236-815102    3928   ------------------------------------------------------------------------------------------
NZ_CP038252.1:1151812-1159673  3857   ------------------------------------------------------------------------------------------
NZ_CP035254.1:793365-801213    3914   ------------------------------------------------------------------------------------------
NZ_CP035251.1:748733-756585    3917   ------------------------------------------------------------------------------------------
NZ_CP035252.1:745670-753536    3931   ------------------------------------------------------------------------------------------
NZ_CP035257.1:776317-784169    3917   ------------------------------------------------------------------------------------------
NZ_CP038253.1:758105-765957    3917   ------------------------------------------------------------------------------------------
NZ_CP035263.1:743050-750901    3913   ------------------------------------------------------------------------------------------
NZ_CP035262.1:741595-749460    3927   ------------------------------------------------------------------------------------------
NZ_CP035236.1:764915-772766    3913   ------------------------------------------------------------------------------------------
NZ_CP035253.1 767286-775134    3914   ------------------------------------------------------------------------------------------
NZ_CP035249.1:763080-770921    3914   ------------------------------------------------------------------------------------------

NC_003028.3:777000-787000      4643   ------------------------------------------------------------------------------------------
NZ_CP035248.1:765099-775111    4652   ------------------------------------------------------------------------------------------
NZ_CP035235.1 c1326834-1317894 3928   ------------------------------------------------------------------------------------------
NZ_CP035234.1:756284-765224    3928   ------------------------------------------------------------------------------------------
NZ_CP038251.1:750745-759686    3928   ------------------------------------------------------------------------------------------
NZ_CP035244.1:747023-755950    3914   ------------------------------------------------------------------------------------------
NZ_CP035264.1:800836-809336    4118   GCATCAAAAACAAAGATGGTAAGGATTATCTAGGATATAACAGTCATCCCTTGCTAGCAGACAGTGATGGGGATGGTTTGGCAGATGGGG
NZ_CP035265.1:766128-774627    4118   GCATCAAAAACAAAGATGGTAAGGATTATCTAGGATATAACAGTCATCCCTTGCTAGCAGACAGTGATGGGGATGGTTTGGCAGATGGGG
NZ_CP035245.1 745261-753757    4118   GCATCAAAAACAAAGATGGTAAGGATTATCTAGGATATAACAGTCATCCCTTGCTAGCAGACAGTGATGGGGATGGTTTGGCAGATGGGG
NZ_CP035259.1:745700-753548    3914   ------------------------------------------------------------------------------------------
NZ_CP035258.1:750874-758722    3914   ------------------------------------------------------------------------------------------
NZ_CP035260.1:726156-733994    3914   ------------------------------------------------------------------------------------------
NZ_CP035241.1:708296-716144    3914   ------------------------------------------------------------------------------------------
NZ_CP035261.1:779086-786938    3914   ------------------------------------------------------------------------------------------
NZ_CP035242.1:c1532236-1524385 3913   ------------------------------------------------------------------------------------------
NZ_CP035243.1:777729-785580    3913   ------------------------------------------------------------------------------------------
NZ_CP035237.1:c1311322-1303456 3928   ------------------------------------------------------------------------------------------
NZ_CP035246.1:c1302892-1295026 3928   ------------------------------------------------------------------------------------------
NZ_CP035247.1:c1301115-1293263 3914   ------------------------------------------------------------------------------------------
NZ_CP035238.1:738547-746413    3928   ------------------------------------------------------------------------------------------
NZ_CP035255.1:776866-784718    3914   ------------------------------------------------------------------------------------------
NZ_CP035256.1:807236-815102    3928   ------------------------------------------------------------------------------------------
NZ_CP038252.1:1151812-1159673  3857   ------------------------------------------------------------------------------------------
NZ_CP035254.1:793365-801213    3914   ------------------------------------------------------------------------------------------
NZ_CP035251.1:748733-756585    3917   ------------------------------------------------------------------------------------------
NZ_CP035252.1:745670-753536    3931   ------------------------------------------------------------------------------------------
NZ_CP035257.1:776317-784169    3917   ------------------------------------------------------------------------------------------
NZ_CP038253.1:758105-765957    3917   ------------------------------------------------------------------------------------------
NZ_CP035263.1:743050-750901    3913   ------------------------------------------------------------------------------------------
NZ_CP035262.1:741595-749460    3927   ------------------------------------------------------------------------------------------
NZ_CP035236.1:764915-772766    3913   ------------------------------------------------------------------------------------------
NZ_CP035253.1 767286-775134    3914   ------------------------------------------------------------------------------------------
NZ_CP035249.1:763080-770921    3914   ------------------------------------------------------------------------------------------

NC_003028.3:777000-787000      4643   ------------------------------------------------------------------------------------------
NZ_CP035248.1:765099-775111    4652   ------------------------------------------------------------------------------------------
NZ_CP035235.1 c1326834-1317894 3928   ------------------------------------------------------------------------------------------
NZ_CP035234.1:756284-765224    3928   ------------------------------------------------------------------------------------------
NZ_CP038251.1:750745-759686    3928   ------------------------------------------------------------------------------------------
NZ_CP035244.1:747023-755950    3914   ------------------------------------------------------------------------------------------
NZ_CP035264.1:800836-809336    4208   AAGATGATAATAAGAAAGAATGGTATGTCACAGACCGTGATTCTCTTCTCTTTATGGAGTTAGCTTATCGAGACGATGATTATATTGAGA
NZ_CP035265.1:766128-774627    4208   AAGATGATAATAAGAAAGAATGGTATGTCACAGACCGTGATTCTCTTCTCTTTATGGAGTTAGCTTATCGAGACGATGATTATATTGAGA
NZ_CP035245.1 745261-753757    4208   AAGATGATAATAAGAAAGAATGGTATGTCACAGACCGTGATTCTCTTCTCTTTATGGAGTTAGCTTATCGAGACGATGATTATATTGAGA
NZ_CP035259.1:745700-753548    3914   ------------------------------------------------------------------------------------------
NZ_CP035258.1:750874-758722    3914   ------------------------------------------------------------------------------------------
NZ_CP035260.1:726156-733994    3914   ------------------------------------------------------------------------------------------
NZ_CP035241.1:708296-716144    3914   ------------------------------------------------------------------------------------------
NZ_CP035261.1:779086-786938    3914   ------------------------------------------------------------------------------------------
NZ_CP035242.1:c1532236-1524385 3913   ------------------------------------------------------------------------------------------
NZ_CP035243.1:777729-785580    3913   ------------------------------------------------------------------------------------------
NZ_CP035237.1:c1311322-1303456 3928   ------------------------------------------------------------------------------------------
NZ_CP035246.1:c1302892-1295026 3928   ------------------------------------------------------------------------------------------
NZ_CP035247.1:c1301115-1293263 3914   ------------------------------------------------------------------------------------------
NZ_CP035238.1:738547-746413    3928   ------------------------------------------------------------------------------------------
NZ_CP035255.1:776866-784718    3914   ------------------------------------------------------------------------------------------
NZ_CP035256.1:807236-815102    3928   ------------------------------------------------------------------------------------------
NZ_CP038252.1:1151812-1159673  3857   ------------------------------------------------------------------------------------------
NZ_CP035254.1:793365-801213    3914   ------------------------------------------------------------------------------------------
NZ_CP035251.1:748733-756585    3917   ------------------------------------------------------------------------------------------
NZ_CP035252.1:745670-753536    3931   ------------------------------------------------------------------------------------------
NZ_CP035257.1:776317-784169    3917   ------------------------------------------------------------------------------------------
NZ_CP038253.1:758105-765957    3917   ------------------------------------------------------------------------------------------
NZ_CP035263.1:743050-750901    3913   ------------------------------------------------------------------------------------------
NZ_CP035262.1:741595-749460    3927   ------------------------------------------------------------------------------------------
```

```
NZ_CP035236.1:764915-772766    3913    ------------------------------------------------------------------------------------------
NZ_CP035253.1 767286-775134    3914    ------------------------------------------------------------------------------------------
NZ_CP035249.1:763080-770921    3914    ------------------------------------------------------------------------------------------

NC_003028.3:777000-787000      4643    ------------------------------------------------------------------------------------------
NZ_CP035248.1:765099-775111    4652    ------------------------------------------------------------------------------------------
NZ_CP035235.1 c1326834-1317894 3928    ------------------------------------------------------------------------------------------
NZ_CP035234.1:756284-765224    3928    ------------------------------------------------------------------------------------------
NZ_CP038251.1:750745-759686    3928    ------------------------------------------------------------------------------------------
NZ_CP035244.1:747023-755950    3914    ------------------------------------------------------------------------------------------
NZ_CP035264.1:800836-809336    4298    AAATTTTATATCATAAGAATCTTTTCCCTAGTCTCTATCTTGACCGTCAAGAACACAAACTCATGCACAATGAATTGGCTCCTTTCTGGA
NZ_CP035265.1:766128-774627    4298    AAATTTTATATCATAAGAATCTTTTCCCTAGTCTCTATCTTGACCGTCAAGAACACAAACTCATGCACAATGAATTGGCTCCTTTCTGGA
NZ_CP035245.1 745261-753757    4298    AAATTTTAGATCATAAGAATCTTTTCCCTAGTCTCTATCTTGACCGTCAAGAACACAAACTCATGCACAATGAATTGGCTCCTTTCTGGA
NZ_CP035259.1:745700-753548    3914    ------------------------------------------------------------------------------------------
NZ_CP035258.1:750874-758722    3914    ------------------------------------------------------------------------------------------
NZ_CP035260.1:726156-733994    3914    ------------------------------------------------------------------------------------------
NZ_CP035241.1:708296-716144    3914    ------------------------------------------------------------------------------------------
NZ_CP035261.1:779086-786938    3914    ------------------------------------------------------------------------------------------
NZ_CP035242.1:c1532236-1524385 3913    ------------------------------------------------------------------------------------------
NZ_CP035243.1:777729-785580    3913    ------------------------------------------------------------------------------------------
NZ_CP035237.1:c1311322-1303456 3928    ------------------------------------------------------------------------------------------
NZ_CP035246.1:c1302892-1295026 3928    ------------------------------------------------------------------------------------------
NZ_CP035247.1:c1301115-1293263 3914    ------------------------------------------------------------------------------------------
NZ_CP035238.1:738547-746413    3928    ------------------------------------------------------------------------------------------
NZ_CP035255.1:776866-784718    3914    ------------------------------------------------------------------------------------------
NZ_CP035256.1:807236-815102    3928    ------------------------------------------------------------------------------------------
NZ_CP038252.1:1151812-1159673  3857    ------------------------------------------------------------------------------------------
NZ_CP035254.1:793365-801213    3914    ------------------------------------------------------------------------------------------
NZ_CP035251.1:748733-756585    3917    ------------------------------------------------------------------------------------------
NZ_CP035252.1:745670-753536    3931    ------------------------------------------------------------------------------------------
NZ_CP035257.1:776317-784169    3917    ------------------------------------------------------------------------------------------
NZ_CP038253.1:758105-765957    3917    ------------------------------------------------------------------------------------------
NZ_CP035263.1:743050-750901    3913    ------------------------------------------------------------------------------------------
NZ_CP035262.1:741595-749460    3927    ------------------------------------------------------------------------------------------
NZ_CP035236.1:764915-772766    3913    ------------------------------------------------------------------------------------------
NZ_CP035253.1 767286-775134    3914    ------------------------------------------------------------------------------------------
NZ_CP035249.1:763080-770921    3914    ------------------------------------------------------------------------------------------

NC_003028.3:777000-787000      4643    ------------------------------------------------------------------------------------------
NZ_CP035248.1:765099-775111    4652    ------------------------------------------------------------------------------------------
NZ_CP035235.1 c1326834-1317894 3928    ------------------------------------------------------------------------------------------
NZ_CP035234.1:756284-765224    3928    ------------------------------------------------------------------------------------------
NZ_CP038251.1:750745-759686    3928    ------------------------------------------------------------------------------------------
NZ_CP035244.1:747023-755950    3914    ------------------------------------------------------------------------------------------
NZ_CP035264.1:800836-809336    4388    AGATGAAAAAAGCCTACTATACAGATAGTGGCTTGGATGCTTTCTTATTTGAGACCAAGAGCGACCTTCCTTATCTCAAAGATGGAACGG
NZ_CP035265.1:766128-774627    4388    AGATGAAAAAAGCCTACTATACAGATAGTGGCTTGGATGCTTTCTTATTTGAGACCAAGAGCGACCTTCCTTATCTCAAAGATGGAACGG
NZ_CP035245.1 745261-753757    4388    AGATGAAAAAAGCCTACTATACAGATAGTGGCTTGGATGCTTTCTTATTTGAGACCAAGAGCGACCTTCCTTATCTCAAAGATGGAACGG
NZ_CP035259.1:745700-753548    3914    ------------------------------------------------------------------------------------------
NZ_CP035258.1:750874-758722    3914    ------------------------------------------------------------------------------------------
NZ_CP035260.1:726156-733994    3914    ------------------------------------------------------------------------------------------
NZ_CP035241.1:708296-716144    3914    ------------------------------------------------------------------------------------------
NZ_CP035261.1:779086-786938    3914    ------------------------------------------------------------------------------------------
NZ_CP035242.1:c1532236-1524385 3913    ------------------------------------------------------------------------------------------
NZ_CP035243.1:777729-785580    3913    ------------------------------------------------------------------------------------------
NZ_CP035237.1:c1311322-1303456 3928    ------------------------------------------------------------------------------------------
NZ_CP035246.1:c1302892-1295026 3928    ------------------------------------------------------------------------------------------
NZ_CP035247.1:c1301115-1293263 3914    ------------------------------------------------------------------------------------------
NZ_CP035238.1:738547-746413    3928    ------------------------------------------------------------------------------------------
NZ_CP035255.1:776866-784718    3914    ------------------------------------------------------------------------------------------
NZ_CP035256.1:807236-815102    3928    ------------------------------------------------------------------------------------------
NZ_CP038252.1:1151812-1159673  3857    ------------------------------------------------------------------------------------------
NZ_CP035254.1:793365-801213    3914    ------------------------------------------------------------------------------------------
NZ_CP035251.1:748733-756585    3917    ------------------------------------------------------------------------------------------
NZ_CP035252.1:745670-753536    3931    ------------------------------------------------------------------------------------------
NZ_CP035257.1:776317-784169    3917    ------------------------------------------------------------------------------------------
NZ_CP038253.1:758105-765957    3917    ------------------------------------------------------------------------------------------
NZ_CP035263.1:743050-750901    3913    ------------------------------------------------------------------------------------------
NZ_CP035262.1:741595-749460    3927    ------------------------------------------------------------------------------------------
NZ_CP035236.1:764915-772766    3913    ------------------------------------------------------------------------------------------
NZ_CP035253.1 767286-775134    3914    ------------------------------------------------------------------------------------------
NZ_CP035249.1:763080-770921    3914    ------------------------------------------------------------------------------------------

NC_003028.3:777000-787000      4643    -------------------------------------------------------------------------------------AGCTC
NZ_CP035248.1:765099-775111    4652    -------------------------------------------------------------------------------------AGCTC
NZ_CP035235.1 c1326834-1317894 3928    -------------------------------------------------------------------------------------AGCTC
NZ_CP035234.1:756284-765224    3928    -------------------------------------------------------------------------------------AGCTC
NZ_CP038251.1:750745-759686    3928    -------------------------------------------------------------------------------------AGCTC
NZ_CP035244.1:747023-755950    3914    -------------------------------------------------------------------------------------AGCTC
NZ_CP035264.1:800836-809336    4478    TGTACATGTTGGCTATTCGTGGAACGCGAGTTAATGACGCCAAGGACTTGAGTGCAGATTTGGTTTTATTAGGTGGAAATAAACCAGCTC
NZ_CP035265.1:766128-774627    4478    TGTACATGTTGGCTATTCGTGGAACGCGAGTTAATGACGCCAAGGACTTGAGTGCAGATTTGGTTTTATTAGGTGGAAATAAACCAGCTC
NZ_CP035245.1 745261-753757    4478    TGCACATGTTGGCTATTCGTGGAACGCGAGTTAATGACGCCAAGGACTTGAGTGCAGATTTTGTTTTATTAGGTGGAAATAAACTAGCTC
NZ_CP035259.1:745700-753548    3914    -------------------------------------------------------------------------------------AGCTC
NZ_CP035258.1:750874-758722    3914    -------------------------------------------------------------------------------------AGCTC
NZ_CP035260.1:726156-733994    3914    -------------------------------------------------------------------------------------AGCTC
NZ_CP035241.1:708296-716144    3914    -------------------------------------------------------------------------------------AGCTC
NZ_CP035261.1:779086-786938    3914    -------------------------------------------------------------------------------------AGCTC
NZ_CP035242.1:c1532236-1524385 3913    -------------------------------------------------------------------------------------AGCTC
NZ_CP035243.1:777729-785580    3913    -------------------------------------------------------------------------------------AGCTC
NZ_CP035237.1:c1311322-1303456 3928    -------------------------------------------------------------------------------------AGCTC
NZ_CP035246.1:c1302892-1295026 3928    -------------------------------------------------------------------------------------AGCTC
NZ_CP035247.1:c1301115-1293263 3914    -------------------------------------------------------------------------------------AGCTC
NZ_CP035238.1:738547-746413    3928    -------------------------------------------------------------------------------------AGCTC
NZ_CP035255.1:776866-784718    3914    -------------------------------------------------------------------------------------AGCTC
NZ_CP035256.1:807236-815102    3928    -------------------------------------------------------------------------------------AGCTC
NZ_CP038252.1:1151812-1159673  3857    -------------------------------------------------------------------------------------AGCTC
```

### Ozkan et al. Supplementary Figures

| | | |
|---|---|---|
| NZ_CP035254.1:793365-801213 | 3914 | -------------------------------------------------------------------------------------AGCTC |
| NZ_CP035251.1:748733-756585 | 3917 | -------------------------------------------------------------------------------------AGCTC |
| NZ_CP035252.1:745670-753536 | 3931 | -------------------------------------------------------------------------------------AGCTC |
| NZ_CP035257.1:776317-784169 | 3917 | -------------------------------------------------------------------------------------AGCTC |
| NZ_CP038253.1:758105-765957 | 3917 | -------------------------------------------------------------------------------------AGCTC |
| NZ_CP035263.1:743050-750901 | 3913 | -------------------------------------------------------------------------------------AGCTC |
| NZ_CP035262.1:741595-749460 | 3927 | -------------------------------------------------------------------------------------AGCTC |
| NZ_CP035236.1:764915-772766 | 3913 | -------------------------------------------------------------------------------------AGCTC |
| NZ_CP035253.1 767286-775134 | 3914 | -------------------------------------------------------------------------------------AGCTC |
| NZ_CP035249.1:763080-770921 | 3914 | -------------------------------------------------------------------------------------AGCTC |

| | | |
|---|---|---|
| NC_003028.3:777000-787000 | 4649 | AAGCGGATGATATCCGCAAGGTTGTTGGGGAATTAGCCAAGGATATAAGTATTACTAAGTTGTATATGACAGGTCATTCTCTTGGATGTT |
| NZ_CP035248.1:765099-775111 | 4658 | AAGCGGATGATATCCGCAAGGTTGTTGGGGAATTAGCCAAGGATATAAGTATTACTAAGTTGTATATGACAGGTCATTCTCTTGGATGTT |
| NZ_CP035235.1 c1326834-1317894 | 3934 | AAGCGGATGATATCCGCAAGGTTGTTGGGGAATTAGCCAAGGATATAAGTATTACTAAGTTGTATATGACAGGTCATTCTCTTGGAGGCT |
| NZ_CP035234.1:756284-765224 | 3934 | AAGCGGATGATATCCGCAAGGTTGTTGGGGAATTAGCCAAGGATATAAGTATTACTAAGTTGTATATGACAGGTCATTCTCTTGGAGGCT |
| NZ_CP038251.1:750745-759686 | 3934 | AAGCGGATGATATCCGCAAGGTTGTTGGGGAATTAGCCAAGGATATAAGTATTACTAAGTTGTATATGACAGGTCATTCTCTTGGAGGCT |
| NZ_CP035244.1:747023-755950 | 3920 | AAGCGGATGATATCCGCAAGGTTGTTGGGGAATTAGCCAAGGATATAAGTATTACTAAGTTGTATATGACAGGTCATTCTCTTGGAGGCT |
| NZ_CP035264.1:800836-809336 | 4568 | AAGCGGATGATATCCGCAAGGTTGTTGGGGAATTAGCCAAGGATATAAGTATTACTAAGTTGTATATGACAGGTCATTCTCTTGGAGGCT |
| NZ_CP035265.1:766128-774627 | 4568 | AAGCGGATGATATCCGCAAGGTTGTTGGGGAATTAGCCAAGGATATAAGTATTACTAAGTTGTATATGACAGGTCATTCTCTTGGAGGCT |
| NZ_CP035245.1 745261-753757 | 4568 | AAGCGGATGATATCCGCAAGGTTGTTGGGGAATTAGCCAAGGATATAAGTATTACTAAGTTGTATATGACAGGTCATTCTCTTGGAGGCT |
| NZ_CP035259.1:745700-753548 | 3920 | AAGCGGATGATATCCGCAAGGTTGTTGGGGAATTAGCCAAGGATATAAGTATTACTAAGTTGTATATGACAGGTCATTCTCTTGGATGTT |
| NZ_CP035258.1:750874-758722 | 3920 | AAGCGGATGATATCCGCAAGGTTGTTGGGGAATTAGCCAAGGATATAAGTATTACTAAGTTGTATATGACAGGTCATTCTCTTGGATGTT |
| NZ_CP035260.1:726156-733994 | 3920 | AAGCGGATGATATCCGCAAGGTTGTTGGGGAATTAGCCAAGGATA----------TAAGTTGTATATGACAGGTCATTCTCTTGGAGGCT |
| NZ_CP035241.1:708296-716144 | 3920 | AAGCGGATGATATCCGCAAGGTTGTTGGGGAATTAGCCAAGGATATAAGTATTACTAAGTTGTATATGACAGGTCATTCTCTTGGAGGCT |
| NZ_CP035261.1:779086-786938 | 3920 | AAGCGGATGATATCCGCAAGGTTGTTGGGGAATTAGCCAAGGATATAAGTATTACTAAGTTGTATATGACAGGTCATTCTCTTGGATGTT |
| NZ_CP035242.1:c1532236-1524385 | 3919 | AAGCGGATGATATCCGCAAGGTTGTTGGGGAATTAGCCAAGGATATAAGTATTACTAAGTTGTATATGACAGGTCATTCTCTTGGATGTT |
| NZ_CP035243.1:777729-785580 | 3919 | AAGCGGATGATATCCGCAAGGTTGTTGGGGAATTAGCCAAGGATATAAGTATTACTAAGTTGTATATGACAGGTCATTCTCTTGGATGTT |
| NZ_CP035237.1:c1311322-1303456 | 3934 | AAGCGGATGATATCCGCAAGGTTGTTGGGGAATTAGCCAAGGATATAAGTATTACTAAGTTGTATATGACAGGTCATTCTCTTGGAGGCT |
| NZ_CP035246.1:c1302892-1295026 | 3934 | AAGCGGATGATATCCGCAAGGTTGTTGGGGAATTAGCCAAGGATATAAGTATTACTAAGTTGTATATGACAGGTCATTCTCTTGGATGCT |
| NZ_CP035247.1:c1301115-1293263 | 3920 | AAGCGGATGATATCCGCAAGGTTGTTGGGGAATTAGCCAAGGATATAAGTATTACTAAGTTGTATATGACAGGTCATTCTCTTGGATGCT |
| NZ_CP035238.1:738547-746413 | 3934 | AAGCGGATGATATCCGCAAGGTTGTTGGGGAATTAGCCAAGGATATAAGTATTACTAAGTTGTATATGACAGGTCATTCTCTTGGATGCT |
| NZ_CP035255.1:776866-784718 | 3920 | AAGCGGATGATATCCGCAAGGTTGTTGGGGAATTAGCCAAGGATATAAGTATTACTAAGTTGTATATGACAGGTCATTCTCTTGGAGGCT |
| NZ_CP035256.1:807236-815102 | 3934 | AAGCGGATGATATCCGCAAGGTTGTTGGGGAATTAGCCAAGGATATAAGTATTACTAAGTTGTATATGACAGGTCATTCTCTTGGAGGCT |
| NZ_CP038252.1:1151812-1159673 | 3863 | AAGCGGATGATATCCGCAAGGTTGTTGGGGAATTAGCCAAGGATATAAGTATTACTAAGTTGTATATGACAGGTCATTCTCTTGGATGTT |
| NZ_CP035254.1:793365-801213 | 3920 | AAGCGGATGATATCCGCAAGGTTGTTGGGGAATTAGCCAAGGATATAAGTATTACTAAGTTGTATATGACAGGTCATTCTCTTGGAGGCT |
| NZ_CP035251.1:748733-756585 | 3923 | AAGCGGATGATATCCGCAAGGTTGTTGGGGAATTAGCCAAGGATATAAGTATTACTAAGTTGTATATGACAGGTCATTTTCTTGGATGTT |
| NZ_CP035252.1:745670-753536 | 3937 | AAGCGGATGATATCCGCAAGGTTGTTGGGGAATTAGCCAAGGATATAAGTATTACTAAGTTGTATATGACAGGTCATTTTCTTGGATGTT |
| NZ_CP035257.1:776317-784169 | 3923 | AAGCGGATGATATCCGCAAGGTTGTTGGGGAATTAGCCAAGGATATAAGTATTACTAAGTTGTATATGACAGGTCATTTTCTTGGATGTT |
| NZ_CP038253.1:758105-765957 | 3923 | AAGCGGATGATATCCGCAAGGTTGTTGGGGAATTAGCCAAGGATATAAGTATTACTAAGTTGTATATGACAGGTCATTTTCTTGGATGTT |
| NZ_CP035263.1:743050-750901 | 3919 | AAGCGGATGATATCCGCAAGGTTGTTGGGGAATTAGCCAAGGATATAAGTATTACTAAGTTGTATATGACAGGTCATTCTCTTGGATGTT |
| NZ_CP035262.1:741595-749460 | 3933 | AAGCGGATGATATCCGCAAGGTTGTTGGGGAATTAGCCAAGGATATAAGTATTACTAAGTTGTATATGACAGGTCATTCTCTTGGATGTT |
| NZ_CP035236.1:764915-772766 | 3919 | AAGCGGATGATATCCGCAAGGTTGTTGGGGAATTAGCCAAGGATATAAGTATTACTAAGTTGTATATGACAGGTCATTCTCTTGGATGTT |
| NZ_CP035253.1 767286-775134 | 3920 | AAGCGGATGATATCCGCAAGGTTGTTGGGGAATTAGCCAAGGATATAAGTATTACTAAGTTGTATATGACAGGTCATTCTCTTGGAGGCT |
| NZ_CP035249.1:763080-770921 | 3920 | AAGCGGATGATATCCGCAAGGTTGTTGGGGAATTAGCCAAGGATATAAGTATTACTAAGTTGTATATGACAGGTCATTCTCTTGGATGTT |

| | | |
|---|---|---|
| NC_003028.3:777000-787000 | 4739 | ACCTAGCTCAGATTGCAGCGGTTGAAGCTTACCAAAAATATCCTGATTTTTATAACCATGTATTGAGGAAAGTGACAACTTTCAGTGCTC |
| NZ_CP035248.1:765099-775111 | 4748 | ACCTAGCTCAGATTGCAGCGGTTGAAGCTTACCAAAAATATCCTGATTTTTATAACCATGTATTGAGGAAAGTGACAACTTTCAGTGCTC |
| NZ_CP035235.1 c1326834-1317894 | 4024 | ACCTAGCTCAGATTGCAGCGGTTGAAGATTACCAAAAATATCCTGATTTTTATAACCATGTATTGAGGAAAGTGACAACTTTCAGTGCTC |
| NZ_CP035234.1:756284-765224 | 4024 | ACCTAGCTCAGATTGCAGCGGTTGAAGATTACCAAAAATATCCTGATTTTTATAACCATGTATTGAGGAAAGTGACAACTTTCAGTGCTC |
| NZ_CP038251.1:750745-759686 | 4024 | ACCTAGCTCAGATTGCAGCGGTTGAAGATTACCAAAAATATCCTGATTTTTATAACCATGTATTGAGGAAAGTGACAACTTTCAGTGCTC |
| NZ_CP035244.1:747023-755950 | 4010 | ACCTAGCTCAGATTGCAGCGGTTGAAGATTACCAAAAATATCCTGATTTTTATAACCATGTATTGAGGAAAGTGACAACTTTCAGTGCTC |
| NZ_CP035264.1:800836-809336 | 4658 | ACCTAGCTCAGATTGCAGCGGTTGAAGATTACCAAAAATATCCTGATTTTTATAACCATGTATTGAGGAAAGTGACAACTTTCAGTGCTC |
| NZ_CP035265.1:766128-774627 | 4658 | ACCTAGCTCAGATTGCAGCGGTTGAAGATTACCAAAAATATCCTGATTTTTATAACCATGTATTGAGGAAAGTGACAACTTTCAGTGCTC |
| NZ_CP035245.1 745261-753757 | 4658 | ACCTAGCTCAGATTGCAGCGGTTGAAGATTACCAAAAATATCCTGATTTTTATAACCATGTATTGAGGAAAGTGACAACTTTCAGTGCTC |
| NZ_CP035259.1:745700-753548 | 4010 | ACCTAGCTCAGATTGCAGCGGTTGAAGCTTACCAAAAATATCCTGATTTTTATAACCATGTATTGAGGAAAGTGACAACTTTCAGTGCTC |
| NZ_CP035258.1:750874-758722 | 4010 | ACCTAGCTCAGATTGCAGCGGTTGAAGCTTACCAAAAATATCCTGATTTTTATAACCATGTATTGAGGAAAGTGACAACTTTCAGTGCTC |
| NZ_CP035260.1:726156-733994 | 4000 | ACCTAGCTCAGATTGCAGCGGTTGAAGATTACCAAAAATATCCTGATTTTTATAACCATGTATTGAGGAAAGTGACAACTTTCAGTGCTC |
| NZ_CP035241.1:708296-716144 | 4010 | ACCTAGCTCAGATTGCAGCGGTTGAAGATTACCAAAAATATCCTGATTTTTATAACCATGTATTGAGGAAAGTGACAACTTTCAGTGCTC |
| NZ_CP035261.1:779086-786938 | 4010 | ACCTAGCTCAGATTGCAGCGGTTGAAGCTTACCAAAAATATCCTGATTTTTATAACCATGTATTGAGGAAAGTGACAACTTTCAGTGCTC |
| NZ_CP035242.1:c1532236-1524385 | 4009 | ACCTAGCTCAGATTGCAGCGGTTGAAGCTTACCAAAAATATCCTGATTTTTATAACCATGTATTGAGGAAAGTGACAACTTTCAGTGCTC |
| NZ_CP035243.1:777729-785580 | 4009 | ACCTAGCTCAGATTGCAGCGGTTGAAGCTTACCAAAAATATCCTGATTTTTATAACCATGTATTGAGGAAAGTGACAACTTTCAGTGCTC |
| NZ_CP035237.1:c1311322-1303456 | 4024 | ACCTAGCTCAGATTGCAGCGGTTGAAGATTACCAAAAATATCCTGATTTTTATAACCATGTATTGAGGAAAGTGACAACTTTCAGTGCTC |
| NZ_CP035246.1:c1302892-1295026 | 4024 | ACCTAGCTCAGATTGCAGAGGTTGAAGCTTACCAAAAATATCCTGATTTTTATAACCATGTATTGAGGAAAGTGACAACTTTCAGTGCTC |
| NZ_CP035247.1:c1301115-1293263 | 4010 | ACCTAGCTCAGATTGCAGAGGTTGAAGCTTACCAAAAATATCCTGATTTTTATAACCATGTATTGAGGAAAGTGACAACTTTCAGTGCTC |
| NZ_CP035238.1:738547-746413 | 4024 | ACCTAGCTCAGATTGCAGAGGTTGAAGCTTACCAAAAATATCCTGATTTTTATAACCATGTATTGAGGAAAGTGACAACTTTCAGTGCTC |
| NZ_CP035255.1:776866-784718 | 4010 | ACCTAGCTCAGATTGCAGAGGTTGAAGCTTACCAAAAATATCCTGATTTTTATAACCATGTATTGAGGAAAGTGACAACTTTCAGTGCTC |
| NZ_CP035256.1:807236-815102 | 4024 | ACCTAGCTCAGATTGCAGAGGTTGAAGCTTACCAAAAATATCCTGATTTTTATAACCATGTATTGAGGAAAGTGACAACTTTCAGTGCTC |
| NZ_CP038252.1:1151812-1159673 | 3953 | ACCTAGCTCAGATTGCAGCGGTTGAAGCTTACCAAAAATATCCTGATTTTTATAACCATGTATTGAGGAAAGTGACAACTTTCAGTGCTC |
| NZ_CP035254.1:793365-801213 | 4010 | ACCTAGCTCAGATTGCAGCGGTTGAAGATTACCAAAAATATCCTGATTTTTATAACCATGTATTGAGGAAAGTGACAACTTTCAGTGCTC |
| NZ_CP035251.1:748733-756585 | 4013 | ACCTAGCTCAGATTGCAGCGGTTGAAGCTTACCAAAAATATCCTGATTTTTATAACCATGTATTGAGGAGAGTGACAACTTTCAGTGCTC |
| NZ_CP035252.1:745670-753536 | 4027 | ACCTAGCTCAGATTGCAGCGGTTGAAGCTTACCAAAAATATCCTGATTTTTATAACCATGTATTGAGGAGAGTGACAACTTTCAGTGCTC |
| NZ_CP035257.1:776317-784169 | 4013 | ACCTAGCTCAGATTGCAGCGGTTGAAGCTTACCAAAAATATCCTGATTTTTATAACCATGTATTGAGGAGAGTGACAACTTTCAGTGCTC |
| NZ_CP038253.1:758105-765957 | 4013 | ACCTAGCTCAGATTGCAGCGGTTGAAGCTTACCAAAAATATCCTGATTTTTATAACCATGTATTGAGGAGAGTGACAACTTTCAGTGCTC |
| NZ_CP035263.1:743050-750901 | 4009 | ACCTAGCTCAGATTGCAGCGGTTGAAGCTTACCAAAAATATCCTGATTTTTATAACCATGTATTGAGGAAAGTGACAACTTTCAGTGCTC |
| NZ_CP035262.1:741595-749460 | 4023 | ACCTAGCTCAGATTGCAGCGGTTGAAGCTTACCAAAAATATCCTGATTTTTATAACCATGTATTGAGGAAAGTGACAACTTTCAGTGCTC |
| NZ_CP035236.1:764915-772766 | 4009 | ACCTAGCTCAGATTGCAGCGGTTGAAGCTTACCAAAAATATCCTGATTTTTATAACCATGTATTGAGGAAAGTGACAACTTTCAGTGCTC |
| NZ_CP035253.1 767286-775134 | 4010 | ACCTAGCTCAGATTGCAGCGGTTGAAGATTACCAAAAATATCCTGATTTTTATAACCATGTATTGAGGAAAGTGACAACTTTCAGTGCTC |
| NZ_CP035249.1:763080-770921 | 4010 | ACCTAGCTCAGATTGCAGCGGTTGAAGCTTACCAAAAATATCCTGATTTTTATAACCATGTATTGAGGAAAGTGACAACTTTCAGTGCTC |

| | | |
|---|---|---|
| NC_003028.3:777000-787000 | 4829 | CTAAAGTGATTACTTCCAGAACTGTTTGGAATGCTAAGAATGGTTTCTGGGATGTTGGTTTGGAAAGTCGTAAATTAGCTGTTAGTGGAA |
| NZ_CP035248.1:765099-775111 | 4838 | CTAAAGTGATTACTTCCAGAACTGTTTGGAATGCTAAGAATGGTTTCTGGGATGTTGGTTTGGAAAGTCGTAAATTAGCTGTTAGTGGAA |
| NZ_CP035235.1 c1326834-1317894 | 4114 | CTAAAGTGATTACTTCCAGAACTGTTTGGGATGCTAAGAATGGTTTCTGAGATGTTGGTTTGGAAAGTCGTAAATTAGCTGTTAGTGGAA |
| NZ_CP035234.1:756284-765224 | 4114 | CTAAAGTGATTACTTCCAGAACTGTTTGGGATGCTAAGAATGGTTTCTGAGATGTTGGTTTGGAAAGTCGTAAATTAGCTGTTAGTGGAA |
| NZ_CP038251.1:750745-759686 | 4114 | CTAAAGTGATTACTTCCAGAACTGTTTGGGATGCTAAGAATGGTTTCTGAGATGTTGGTTTGGAAAGTCGTAAATTAGCTGTTAGTGGAA |
| NZ_CP035244.1:747023-755950 | 4100 | CTAAAGTGATTACTTCCAGAACTGTTTGGGATGCTAAGAATGGTTTCTGAGATGTTGGTTTGGAAAGTCGTAAATTAGCTGTTAGTGGAA |
| NZ_CP035264.1:800836-809336 | 4748 | CTAAAGTGATTACTTCCAGAACTGTTTGGGATGCTAAGAATGGTTTCTGAGATGTTGGTTTGGAAAGTCGTAAATTAGCTGTTAGTGGAA |
| NZ_CP035265.1:766128-774627 | 4748 | CTAAAGTGATTACTTCCAGAACTGTTTGGGATGCTAAGAATGGTTTCTGAGATGTTGGTTTGGAAAGTCGTAAATTAGCTGTTAGTGGAA |
| NZ_CP035245.1 745261-753757 | 4748 | CTAAAGTCATTACTTCCAGAACTGTTTGGGATGCTAAGAATGGTTTCTGAGATGTTGGTTTGGAAAGTCGTAAATTAGCTGTTAGTGGAA |
| NZ_CP035259.1:745700-753548 | 4100 | CTAAAGTGATTCCTTCCAGAACTGTTTGGAATGCTAAGAATGGTTTCTGGGATGTTGGTTTGGAAAGTCGTAAATTAGCTGTTAGTGGAA |
| NZ_CP035258.1:750874-758722 | 4100 | CTAAAGTGATTCCTTCCAGAACTGTTTGGAATGCTAAGAATGGTTTCTGGGATGTTGGTTTGGAAAGTCGTAAATTAGCTGTTAGTGGAA |
| NZ_CP035260.1:726156-733994 | 4090 | CTAAAGTGATTACTTCCAGAACTGTTTGGGATGCTAAGAATGGTTTCTGAGATGTTGGTTTGGAAAGTCGTAAATTAGCTGTTAGTGGAA |
| NZ_CP035241.1:708296-716144 | 4100 | CTAAAGTGATTACTTCCAGAACTGTTTGGGATGCTAAGAATGGTTTCTGAGATGTTGGTTTGGAAAGTCGTAAATTAGCTGTTAGTGGAA |
| NZ_CP035261.1:779086-786938 | 4100 | CTAAAGTGATTACTTCCAGAACTGTTTGGAATGCTAAGAATGGTTTCTGGGATGTTGGTTTGGAAAGTCGTAAATTAGCTGTTAGTGGAA |
| NZ_CP035242.1:c1532236-1524385 | 4099 | CTAAAGTGATTACTTCCAGAACTGTTTGGAATGCTAAGAATGGTTTCTGGGATGTTGGTTTGGAAAGTCGTAAATTAGCTGTTAGTGGAA |
| NZ_CP035243.1:777729-785580 | 4099 | CTAAAGTGATTACTTCCAGAACTGTTTGGAATGCTAAGAATGGTTTCTGGGATGTTGGTTTGGAAAGTCGTAAATTAGCTGTTAGTGGAA |

### Ozkan et al. Supplementary Figures

```
NZ_CP035237.1:c1311322-1303456 4114   CTAAAGTCATTACTTCCAGAACTGTTTGGGATGCTAAGAATGGTTTCTGAGATGTTGGTTTGGAAAGTCGTAAATTAGCTGTTAGTGGAA
NZ_CP035246.1:c1302892-1295026 4114   CTAAAGTGATTACTTCCAGAACTGTTTGGAATGCTAAGAATGGTTTCTGGGATGTTGGTTTGGAAAGTCGTAAATTAGCTGTTAGTGGAA
NZ_CP035247.1:c1301115-1293263 4100   CTAAAGTGATTACTTCCAGAACTGTTTGGAATGCTAAGAATGGTTTCTGGGATGTTGGTTTGGAAAGTCGTAAATTAGCTGTTAGTGGAA
NZ_CP035238.1:738547-746413    4114   CTAAAGTGATTACTTCCAGAACTGTTTGGAATGCTAAGAATGGTTTCTGGGATGTTGGTTTGGAAAGTCGTAAATTAGCTGTTAGTGGAA
NZ_CP035255.1:776866-784718    4100   CTAAAGTGATTACTTCCAGAACTGTTTGGAATGCTAAGAATGGTTTCTGGGATGTTGGTTTGGAAAGTCGTAAATTAGCTGTTAGTGGAA
NZ_CP035256.1:807236-815102    4114   CTAAAGTGATTACTTCCAGAACTGTTTGGAATGCTAAGAATGGTTTCTGGGATGTTGGTTTGGAAAGTCGTAAATTAGCTGTTAGTGGAA
NZ_CP038252.1:1151812-1159673  4043   CTAAAGTGATTACTTCCAGAACTGTTTGGAATGCTAAGAATGGTTTCTGGGATGTTGGTTTGGAAAGTCGTAAATTAGCTGTTAGTGGAA
NZ_CP035254.1:793365-801213    4100   CTAAAGTGATTACTTCCAGAACTGTTTGGGATGCTAAGAATGGTTTCTGAGATGTTGGTTTGGAAAGTCGTAAATTAGCTGTTAGTGGAA
NZ_CP035251.1:748733-756585    4103   CTAAAGTGATTACTTCCAGAACTGTTTGGAATGCTAAGAATGGTTTCTGGGATGTTGGTTTGGAAAGTCGTAAATTAGCTGTTAGTGGAA
NZ_CP035252.1:745670-753536    4117   CTAAAGTGATTACTTCCAGAACTGTTTGGAATGCTAAGAATGGTTTCTGGGATGTTGGTTTGGAAAGTCGTAAATTAGCTGTTAGTGGAA
NZ_CP035257.1:776317-784169    4103   CTAAAGTGATTACTTCCAGAACTGTTTGGAATGCTAAGAATGGTTTCTGGGATGTTGGTTTGGAAAGTCGTAAATTAGCTGTTAGTGGAA
NZ_CP038253.1:758105-765957    4103   CTAAAGTGATTACTTCCAGAACTGTTTGGAATGCTAAGAATGGTTTCTGGGATGTTGGTTTGGAAAGTCGTAAATTAGCTGTTAGTGGAA
NZ_CP035263.1:743050-750901    4099   CTAAAGTGATTACTTCCAGAACTGTTTGGGATGCTAAGAATGGTTTCTGAGATGTTGGTTTGGAAAGTCGTAAATTAGCTGTTAGTGGAA
NZ_CP035262.1:741595-749460    4113   CTAAAGTGATTACTTCCAGAACTGTTTGGGATGCTAAGAATGGTTTCTGAGATGTTGGTTTGGAAAGTCGTAAATTAGCTGTTAGTGGAA
NZ_CP035236.1:764915-772766    4099   CTAAAGTGATTACTTCCAGAACTGTTTGGGATGCTAAGAATGGTTTCTGAGATGTTGGTTTGGAAAGTCGTAAATTAGCTGTTAGTGGAA
NZ_CP035253.1 767286-775134    4100   CTAAAGTGATTACTTCCAGAACTGTTTGGGATGCTAAGAATGGTTTCTGAGATGTTGGTTTGGAAAGTCGTAAATTAGCTGTTAGTGGAA
NZ_CP035249.1:763080-770921    4100   CTAAAGTGATTACTTCCAGAACTGTTTGGAATGCTAAGAATGGTTTCTGGGATGTTGGTTTGGAAAGTCGTAAATTAGCTGTTAGTGGAA

NC_003028.3:777000-787000      4919   AAATTAAGCATTATGTGGTTGATAATGACAATGTTGTGACTCCCTTGATTCATAATAATCGTGATATTGTTACATTTACAGGTAATTCAC
NZ_CP035248.1:765099-775111    4928   AAATTAAGCATTATGTGGTTGATAATGACAATGTTGTGACTCCCTTGATTCATAATAATCGTGATATTGTTACATTTACAGGTAATTCAC
NZ_CP035235.1 c1326834-1317894 4204   AAATTAAGCATTATGTGGTTGATAATGACAATGTTGTGACTCCCTTGATTCATAATAATCGTGATATTGTTACATTTACAGGTAATTCAC
NZ_CP035234.1:756284-765224    4204   AAATTAAGCATTATGTGGTTGATAATGACAATGTTGTGACTCCCTTGATTCATAATAATCGTGATATTGTTACATTTACAGGTAATTCAC
NZ_CP038251.1:750745-759686    4204   AAATTAAGCATTATGTGGTTGATAATGACAATGTTGTGACTCCCTTGATTCATAATAATCGTGATATTGTTACATTTACAGGTAATTCAC
NZ_CP035244.1:747023-755950    4190   AAATTAAGCATTATGTGGTTGATAATGACAATGTTGTGACTCCCTTGATTCATAATAATCGTGATATTGTTACATTTACAGGTAATTCAC
NZ_CP035264.1:800836-809336    4838   AAATTAAGCATTATGTGGTTGATAATGACAATGTTGTGACTCCCTTGATTCATAATAATCGTGATATTGTTACATTTACAGGTAATTCAC
NZ_CP035265.1:766128-774627    4838   AAATTAAGCATTATGTGGTTGATAATGACAATGTTGTGACTCCCTTGATTCATAATAATCGTGATATTGTTACATTTACAGGTAATTCAC
NZ_CP035245.1 745261-753757    4838   AAATTAAGCATTATGTGGTTGATAATGACAATGTTGTGACTCCCTTGATTCATAATAATCGTGATATTGTTACATTTACAGGTAATTCAC
NZ_CP035259.1:745700-753548    4190   AAATTAAGCATTATGTGGTTGATAATGACAATGTTGTGACTCCCTTGATTCATAATGATCGTGATATTGTTACATTTACAGGTAATTCAC
NZ_CP035258.1:750874-758722    4190   AAATTAAGCATTATGTGGTTGATAATGACAATGTTGTGACTCCCTTGATTCATAATGATCGTGATATTGTTACATTTACAGGTAATTCAC
NZ_CP035260.1:726156-733994    4180   AAATTAAGCATTATGTGGTTGATAATGACAATGTTGTGACTCCCTTGATTCATAATAATCGTGATATTGTTACATTTACAGGTAATTCAC
NZ_CP035241.1:708296-716144    4190   AAATTAAGCATTATGTGGTTGATAATGACAATGTTGTGACTCCCTTGATTCATAATAATCGTGATATTGTTACATTTACAGGTAATTCAC
NZ_CP035261.1:779086-786938    4190   AAATTAAGCATTATGTGGTTGATAATGACAATGTTGTGACTCCCTTGATTCATAATAATCGTGATATTGTTACATTTACAGGTAATTCAC
NZ_CP035242.1:c1532236-1524385 4189   AAATTAAGCATTATGTGGTTGATAATGACAATGTTGTGACTCCCTTGATTCATAATAATCGTGATATTGTTACATTTACAGGTAATTCAC
NZ_CP035243.1:777729-785580    4189   AAATTAAGCATTATGTGGTTGATAATGACAATGTTGTGACTCCCTTGATTCATAATAATCGTGATATTGTTACATTTACAGGTAATTCAC
NZ_CP035237.1:c1311322-1303456 4204   AAATTAAGCATTATGTGGTTGATAATGACAATGTTGTGACTCCCTTGATTCATAATAATCGTGATATTGTTACATTTACAGGTAATTCAC
NZ_CP035246.1:c1302892-1295026 4204   AAATTAAGCATTATGTGGTTGATAATGACAATGTTGTGACTCCCTTGATTCATAATGATCGTGATATTGTTACATTTACAGGTAATTCAC
NZ_CP035247.1:c1301115-1293263 4190   AAATTAAGCATTATGTGGTTGATAATGACAATGTTGTGACTCCCTTGATTCATAATGATCGTGATATTGTTACATTTACAGGTAATTCAC
NZ_CP035238.1:738547-746413    4204   AAATTAAGCATTATGTGGTTGATAATGACAATGTTGTGACTCCCTTGATTCATAATGATCGTGATATTGTTACATTTACAGGTAATTCAC
NZ_CP035255.1:776866-784718    4190   AAATTAAGCATTATGTGGTTGATAATGACAATGTTGTGACTCCCTTGATTCATAATGATCGTGATATTGTTACATTTACAGGTAATTCAC
NZ_CP035256.1:807236-815102    4204   AAATTAAGCATTATGTGGTTGATAATGACAATGTTGTGACTCCCTTGATTCATAATGATCGTGATATTGTTACATTTACAGGTAATTCAC
NZ_CP038252.1:1151812-1159673  4133   AAATTAAGCATTATGTGGTTGATAATGACAATGTTGTGACTCCCTTGATTCATAATAATCGTGATATTGTTACATTTACAGGTAATTCAC
NZ_CP035254.1:793365-801213    4190   AAATTAAGCATTATGTGGTTGATAATGACAATGTTGTGACTCCCTTGATTCATAATAATCGTGATATTGTTACATTTACAGGTAATTCAC
NZ_CP035251.1:748733-756585    4193   AAATTAAGCATTATGTGGTTGATAATGACAATGTTGTGACTCCCTTGATTCATAATGATCGTGATATTGTTACATTTACAGGTAATTCAC
NZ_CP035252.1:745670-753536    4207   AAATTAAGCATTATGTGGTTGATAATGACAATGTTGTGACTCCCTTGATTCATAATGATCGTGATATTGTTACATTTACAGGTAATTCAC
NZ_CP035257.1:776317-784169    4193   AAATTAAGCATTATGTGGTTGATAATGACAATGTTGTGACTCCCTTGATTCATAATGATCGTGATATTGTTACATTTACAGGTAATTCAC
NZ_CP038253.1:758105-765957    4193   AAATTAAGCATTATGTGGTTGATAATGACAATGTTGTGACTCCCTTGATTCATAATGATCGTGATATTGTTACATTTACAGGTAATTCAC
NZ_CP035263.1:743050-750901    4189   AAATTAAGCATTATGTGGTTGATAATGACAATGTTGTGACTCCCTTGATTCATAATAATCGTGATATTGTTACATTTACAGGTAATTCAC
NZ_CP035262.1:741595-749460    4203   AAATTAAGCATTATGTGGTTGATAATGACAATGTTGTGACTCCCTTGATTCATAATAATCGTGATATTGTTACATTTACAGGTAATTCAC
NZ_CP035236.1:764915-772766    4189   AAATTAAGCATTATGTGGTTGATAATGACAATGTTGTGACTCCCTTGATTCATAATAATCGTGATATTGTTACATTTACAGGTAATTCAC
NZ_CP035253.1 767286-775134    4190   AAATTAAGCATTATGTGGTTGATAATGACAATGTTGTGACTCCCTTGATTCATAATAATCGTGATATTGTTACATTTACAGGTAATTCAC
NZ_CP035249.1:763080-770921    4190   AAATTAAGCATTATGTGGTTGATAATGACAATGTTGTGACTCCCTTGATTCATAATAATCGTGATATTGTTACATTTACAGGTAATTCAC

NC_003028.3:777000-787000      5009   GCTTTAAACACCGTTCTCGTGGCTATTTTGAAAGTCCAATGAATGATATTCCTAACTTTAATATTGGTAAACAAGCTACCTTGGATAAAC
NZ_CP035248.1:765099-775111    5018   GCTTTAAACACCGTTCTCGTGGCTATTTTGAAAGTCCAATGAATGATATTCCTAACTTTAATATTGGTAAACAAGCTACCTTGGATAAAC
NZ_CP035235.1 c1326834-1317894 4294   GCTTTAAACACCGTTCTCGTGGCTATTTTGAAAGTCCAATGAATGATATTCCTAACTTTAATATTGGTAAACAAGCTACCTTGGATAAAC
NZ_CP035234.1:756284-765224    4294   GCTTTAAACACCGTTCTCGTGGCTATTTTGAAAGTCCAATGAATGATATTCCTAACTTTAATATTGGTAAACAAGCTACCTTGGATAAAC
NZ_CP038251.1:750745-759686    4294   GCTTTAAACACCGTTCTCGTGGCTATTTTGAAAGTCCAATGAATGATATTCCTAACTTTAATATTGGTAAACAAGCTACCTTGGATAAAC
NZ_CP035244.1:747023-755950    4280   GCTTTAAACACCGTTCTCGTGGCTATTTTGAAAGTCCAATGAATGATATTCCTAACTTTAATATTGGTAAACAAGCTACCTTGGATAAAC
NZ_CP035264.1:800836-809336    4928   GCTTTAAACACCGTTCTCGTGGCTATTTTGAAAGTCCAATGAATGATATTCCTAACTTTAATATTGGTAAACAAGCTACCTTGGATAAAC
NZ_CP035265.1:766128-774627    4928   GCTTTAAACACCGTTCTCGTGGCTATTTTGAAAGTCCAATGAATGATATTCCTAACTTTAATATTGGTAAACAAGCTACCTTGGATAAAC
NZ_CP035245.1 745261-753757    4928   GCTTTAAACACCGTTCTCGTGGCTATTTTGAAAGTCCAATGAATGATATTCCTAACTTTAATATTGGTAAACAAGCTACCTTGGATAAAC
NZ_CP035259.1:745700-753548    4280   GCTTTAAACACCGTTCTCGTGGCTATTTTGAAAGTCCAATGAATGATATTCCTAACTTTAATATTGGTAAACAAGCTACCTTGGATAAAC
NZ_CP035258.1:750874-758722    4280   GCTTTAAACACCGTTCTCGTGGCTATTTTGAAAGTCCAATGAATGATATTCCTAACTTTAATATTGGTAAACAAGCTACCTTGGATAAAC
NZ_CP035260.1:726156-733994    4270   GCTTTAAACACCGTTCTCGTGGCTATTTTGAAAGTCCAATGAATGATATTCCTAACTTTAATATTGGTAAACAAGCTACCTTGGATAAAC
NZ_CP035241.1:708296-716144    4280   GCTTTAAACACCGTTCTCGTGGCTATTTTGAAAGTCCAATGAATGATATTCCTAACTTTAATATTGGTAAACAAGCTACCTTGGATAAAC
NZ_CP035261.1:779086-786938    4280   GCTTTAAACACCGTTCTCGTGGCTATTTTGAAAGTCCAATGAATGATATTCCTAACTTTAATATTGGTAAACAAGCTACCTTGGATAAAC
NZ_CP035242.1:c1532236-1524385 4279   GCTTTAAACACCGTTCTCGTGGCTATTTTGAAAGTCCAATGAATGATATTCCTAACTTTAATATTGGTAAACAAGCTACCTTGGATAAAC
NZ_CP035243.1:777729-785580    4279   GCTTTAAACACCGTTCTCGTGGCTATTTTGAAAGTCCAATGAATGATATTCCTAACTTTAATATTGGTAAACAAGCTACCTTGGATAAAC
NZ_CP035237.1:c1311322-1303456 4294   GCTTTAAACACCGTTCTCGTGGCTATTTTGAAAGTCCAATGAATGATATTCCTAACTTTAATATTGGTAAACAAGCTACCTTGGATAAAC
NZ_CP035246.1:c1302892-1295026 4294   GCTTTAAACACCGTTCTCGTGGCTATTTTGAAAGTCCAATGAATGATATTCCTAACTTTAATATTGGTAAACAAGCTACCTTGGATAAAC
NZ_CP035247.1:c1301115-1293263 4280   GCTTTAAACACCGTTCTCGTGGCTATTTTGAAAGTCCAATGAATGATATTCCTAACTTTAATATTGGTAAACAAGCTACCTTGGATAAAC
NZ_CP035238.1:738547-746413    4294   GCTTTAAACACCGTTCTCGTGGCTATTTTGAAAGTCCAATGAATGATATTCCTAACTTTAATATTGGTAAACAAGCTACCTTGGATAAAC
NZ_CP035255.1:776866-784718    4280   GCTTTAAACACCGTTCTCGTGGCTATTTTGAAAGTCCAATGAATGATATTCCTAACTTTAATATTGGTAAACAAGCTACCTTGGATAAAC
NZ_CP035256.1:807236-815102    4294   GCTTTAAACACCGTTCTCGTGGCTATTTTGAAAGTCCAATGAATGATATTCCTAACTTTAATATTGGTAAACAAGCTACCTTGGATAAAC
NZ_CP038252.1:1151812-1159673  4223   GCTTTAAACACCGTTCTCGTGGCTATTTTGAAAGTCCAATGAATGATATTCCTAACTTTAATATTGGTAAACAAGCTACCTTGGATAAAC
NZ_CP035254.1:793365-801213    4280   GCTTTAAACACCGTTCTCGTGGCTATTTTGAAAGTCCAATGAATGATATTCCTAACTTTAATATTGGTAAACAAGCTACCTTGGATAAAC
NZ_CP035251.1:748733-756585    4283   GCTTTAAACACCGTTCTCGTGGCTATTTTGAAAGTCCAATGAATGATATTCCTAACTTTAATATTGGTAAACAAGCTACCTTTGATAAAC
NZ_CP035252.1:745670-753536    4297   GCTTTAAACACCGTTCTCGTGGCTATTTTGAAAGTCCAATGAATGATATTCCTAACTTTAATATTGGTAAACAAGCTACCTTTGATAAAC
NZ_CP035257.1:776317-784169    4283   GCTTTAAACACCGTTCTCGTGGCTATTTTGAAAGTCCAATGAATGATATTCCTAACTTTAATATTGGTAAACAAGCTACCTTTGATAAAC
NZ_CP038253.1:758105-765957    4283   GCTTTAAACACCGTTCTCGTGGCTATTTTGAAAGTCCAATGAATGATATTCCTAACTTTAATATTGGTAAACAAGCTACCTTTGATAAAC
NZ_CP035263.1:743050-750901    4279   GCTTTAAACACCGTTCTCGTGGCTATTTTGAAAGTCCAATGAATGATATTCCTAACTTTAATATTGGTAAACAAGCTACCTTGGATAAAC
NZ_CP035262.1:741595-749460    4293   GCTTTAAACACCGTTCTCGTGGCTATTTTGAAAGTCCAATGAATGATATTCCTAACTTTAATATTGGTAAACAAGCTACCTTGGATAAAC
NZ_CP035236.1:764915-772766    4279   GCTTTAAACACCGTTCTCGTGGCTATTTTGAAAGTCCAATGAATGATATTCCTAACTTTAATATTGGTAAACAAGCTACCTTGGATAAAC
NZ_CP035253.1 767286-775134    4280   GCTTTAAACACCGTTCTCGTGGCTATTTTGAAAGTCCAATGAATGATATTCCTAACTTTAATATTGGTAAACAAGCTACCTTGGATAAAC
NZ_CP035249.1:763080-770921    4280   GCTTTAAACACCGTTCTCGTGGCTATTTTGAAAGTCCAATGAATGATATTCCTAACTTTAATATTGGTAAACAAGCTACCTTGGATAAAC

NC_003028.3:777000-787000      5099   ATGGTTATCGTGATCCGAAATTGGATAAAGTGCGATTCTTTAAGAAACAGGCTCTGCCTCGATCTTCTAGTCAACCAAGCGCTGAACCAA
NZ_CP035248.1:765099-775111    5108   ATGGTTATCGTGATCCGAAATTGGATAAAGTGCGATTCTTTAAGAAACAGGCTCTGCCTCGATCTTCTAGTCAACCAAGCGCTGAACCAA
NZ_CP035235.1 c1326834-1317894 4384   ATGGTTATCGTGATCCGAAATTGGATAAAGTGCGATTCTTTAAGAAACAGGCTCTACCTCAATCTTCTAGTCAACCAAGCGCTGAACCAA
NZ_CP035234.1:756284-765224    4384   ATGGTTATCGTGATCCGAAATTGGATAAAGTGCGATTCTTTAAGAAACAGGCTCTACCTCAATCTTCTAGTCAACCAAGCGCTGAACCAA
NZ_CP038251.1:750745-759686    4384   ATGGTTATCGTGATCCGAAATTGGATAAAGTGCGATTCTTTAAGAAACAGGCTCTACCTCAATCTTCTAGTCAACCAAGCGCTGAACCAA
NZ_CP035244.1:747023-755950    4370   ATGGTTATCGTGATCCGAAATTGGATAAAGTGCGATTCTTTAAGAAACAGGCTCTACCTCAATCTTCTAGTCAACCAAGCGCTGAACCAA
NZ_CP035264.1:800836-809336    5018   ATGGTTATCGTGATCCGAAATTGGATAAAGCGCGATTCTTTAAGAAACAGGCTCTACCTCAATCTTCTAGTCAACCAAGCGCTGAACCAA
NZ_CP035265.1:766128-774627    5018   ATGGTTATCGTGATCCGAAATTGGATAAAGCGCGATTCTTTAAGAAACAGGCTCTACCTCAATCTTCTAGTCAACCAAGCGCTGAACCAA
NZ_CP035245.1 745261-753757    5018   ATGGTTATCGTGATCCGAAATTGGATAAAGTGCGATTCTTTAAGAAACAGGCTCTACCTCAATCTTCTAGTCAACCAAGCGCTGAACCAA
```

```
NZ_CP035259.1:745700-753548    4370   ATGGTTATCGTGATCCGAAATTGGATAAAGTGCGATTCTTTAAGAAACAGGCTCTGCCTCGATCTTCTAGTCAACCAAGCGCTGAACCAA
NZ_CP035258.1:750874-758722    4370   ATGGTTATCGTGATCCGAAATTGGATAAAGTGCGATTCTTTAAGAAACAGGCTCTGCCTCGATCTTCTAGTCAACCAAGCGCTGAACCAA
NZ_CP035260.1:726156-733994    4360   ATGGTTATCGTGATCCGAAATTGGATAAAGCGCGATTCTTTAAGAAACAGGCTCTACCTCAATCTTCTAGTCAACCAAGCGCTGAACCAA
NZ_CP035241.1:708296-716144    4370   ATGGTTATCGTGATCCGAAATTGGATAAAGCGCGATTCTTTAAGAAACAGGCTCTACCTCAATCTTCTAGTCAACCAAGCGCTGAACCAA
NZ_CP035261.1:779086-786938    4370   ATGGTTATCGTGATCCGAAATTGGATAAAGTGCGATTCTTTAAGAAACAGGCTCTGCCTCGATCTTCTAGTCAACCAAGCGCTGAACCAA
NZ_CP035242.1:c1532236-1524385 4369   ATGGTTATCGTGATCCGAAATTGGATAAAGTGCGATTCTTTAAGAAACAGGCTCTGCCTCGATCTTCTAGTCAACCAAGCGCTGAACCAA
NZ_CP035243.1:777729-785580    4369   ATGGTTATCGTGATCCGAAATTGGATAAAGTGCGATTCTTTAAGAAACAGGCTCTGCCTCGATCTTCTAGTCAACCAAGCGCTGAACCAA
NZ_CP035237.1:c1311322-1303456 4384   ATGGTTATCGTGATCCGAAATTGGATAAAGTGCGATTCTTTAAGAAACAGGCTCTACCTCAATCTTCTAGTCAACCAAGCGCTGAACCAA
NZ_CP035246.1:c1302892-1295026 4384   ATGGTTATCGTGATCCGAAATTGGATAAAGTGCGATTCTTTAAGAAACAGGCTCTGCCTCGATCTTCTAGTCAACCAAGCGCTGAACCAA
NZ_CP035247.1:c1301115-1293263 4370   ATGGTTATCGTGATCCGAAATTGGATAAAGTGCGATTCTTTAAGAAACAGGCTCTGCCTCGATCTTCTAGTCAACCAAGCGCTGAACCAA
NZ_CP035238.1:738547-746413    4384   ATGGTTATCGTGATCCGAAATTGGATAAAGTGCGATTCTTTAAGAAACAGGCTCTGCCTCGATCTTCTAGTCAACCAAGCGCTGAACCAA
NZ_CP035255.1:776866-784718    4370   ATGGTTATCGTGATCCGAAATTGGATAAAGTGCGATTCTTTAAGAAACAGGCTCTGCCTCGATCTTCTAGTCAACCAAGCGCTGAACCAA
NZ_CP035256.1:807236-815102    4384   ATGGTTATCGTGATCCGAAATTGGATAAAGTGCGATTCTTTAAGAAACAGGCTCTGCCTCGATCTTCTAGTCAACCAAGCGCTGAACCAA
NZ_CP038252.1:1151812-1159673  4313   ATGGTTATCGTGATCCGAAATTGGATAAAGTGCGATTCTTTAAGAAACAGGCTCTGCCTCGATCTTCTAGTCAACCAAGCGCTGAACCAA
NZ_CP035254.1:793365-801213    4370   ATGGTTATCGTGATCCGAAATTGGATAAAGTGCGATTCTTTAAGAAACAGGCTCTACCTCAATCTTCTAGTCAACCAAGCGCTGAACCAA
NZ_CP035251.1:748733-756585    4373   ATGGTTATCGTGATCCGAAATTGGATAAAGTGCGATTCTTTAAGAAACAGGCTCTACCTCAATCTTCTAGTCAACCAAGCGCTGAACCAA
NZ_CP035252.1:745670-753536    4387   ATGGTTATCGTGATCCGAAATTGGATAAAGTGCGATTCTTTAAGAAACAGGCTCTACCTCAATCTTCTAGTCAACCAAGCGCTGAACCAA
NZ_CP035257.1:776317-784169    4373   ATGGTTATCGTGATCCGAAATTGGATAAAGTGCGATTCTTTAAGAAACAGGCTCTACCTCAATCTTCTAGTCAACCAAGCGCTGAACCAA
NZ_CP038253.1:758105-765957    4373   ATGGTTATCGTGATCCGAAATTGGATAAAGTGCGATTCTTTAAGAAACAGGCTCTACCTCAATCTTCTAGTCAACCAAGCGCTGAACCAA
NZ_CP035263.1:743050-750901    4369   ATGGTTATCGTGATCCGAAATTGGATAAAGTGCGATTCTTTAAGAAACAGGCTCTACCTCAATCTTCTAGTCAACCAAGCGCTGAACCAA
NZ_CP035262.1:741595-749460    4383   ATGGTTATCGTGATCCGAAATTGGATAAAGTGCGATTCTTTAAGAAACAGGCTCTACCTCAATCTTCTAGTCAACCAAGCGCTGAACCAA
NZ_CP035236.1:764915-772766    4369   ATGGTTATCGTGATCCGAAATTGGATAAAGTGCGATTCTTTAAGAAACAGGCTCTACCTCAATCTTCTAGTCAACCAAGCGCTGAACCAA
NZ_CP035253.1 767286-775134    4370   ATGGTTATCGTGATCCGAAATTGGATAAAGTGCGATTCTTTAAGAAACAGGCTCTACCTCAATCTTCTAGTCAACCAAGCGCTGAACCAA
NZ_CP035249.1:763080-770921    4370   ATGGTTATCGTGATCCGAAATTGGATAAAGTGCGATTCTTTAAGAAACAGGCTCTGCCTCGATCTTCTAGTCAACCAAGCGCTGAACCAA

NC_003028.3:777000-787000      5189   TGGAAAATATTGCCTCAGGAAAACAGGTTACTCAAAGTTCGACAGCTTTCGGAGGAGATGCTAGAAGAGCTGTGGATGGCAAAGTCGATG
NZ_CP035248.1:765099-775111    5198   TGGAAAATATTGCCTTAGGAAAACAGGTTACTCAAAGTTCGACAGTTTTCGGAGGAGATGCTAGAAGAGCTGTGGATGGCAAAGTCGATG
NZ_CP035235.1 c1326834-1317894 4474   TGGAAAATATTGCCTTAGGAAAACAGGTTACTCAAAGTTCGACAGCTTTCGGAGGAGATGCTAGAAGAGCTGTGGATGGCAAAGTCGATG
NZ_CP035234.1:756284-765224    4474   TGGAAAATATTGCCTTAGGAAAACAGGTTACTCAAAGTTCGACAGCTTTCGGAGGAGATGCTAGAAGAGCTGTGGATGGCAAAGTCGATG
NZ_CP038251.1:750745-759686    4474   TGGAAAATATTGCCTTAGGAAAACAGGTTACTCAAAGTTCGACAGCTTTCGGAGGAGATGCTAGAAGAGCTGTGGATGGCAAAGTCGATG
NZ_CP035244.1:747023-755950    4460   TGGAAAATATTGCCTTAGGAAAACAGGTTACTCAAAGTTCGACAGCTTTCGGAGGAGATGCTAGAAGAGCTGTGGATGGCAAAGTCGATG
NZ_CP035264.1:800836-809336    5108   TGGAAAATATTGCCTTAGGAAAACAGGTTACTCAAAGTTCGACAGCTTTCGGAGGAGATGCTAGAAGAGCTGTGGATGGCAAAGTCGATG
NZ_CP035265.1:766128-774627    5108   TGGAAAATATTGCCTTAGGAAAACAGGTTACTCAAAGTTCGACAGCTTTCGGAGGAGATGCTAGAAGAGCTGTGGATGGCAAAGTCGATG
NZ_CP035245.1 745261-753757    5108   TGGAAAATATTGCCTTAGGAAAACAGGTTACTCAAAGTTCGACAGCTTTCGGAGGAGATGCTAGAAGAGCTGTGGATGGCAAAGTCGATG
NZ_CP035259.1:745700-753548    4460   TGGAAAATATTGCCTCAGGAAAACAGGTTACTCAAAGTTCGACAGCTTTCGGAGGAGATGCTAGAAGAGCTGTGGATGGCAAAGTCGATG
NZ_CP035258.1:750874-758722    4460   TGGAAAATATTGCCTCAGGAAAACAGGTTACTCAAAGTTCGACAGCTTTCGGAGGAGATGCTAGAAGAGCTGTGGATGGCAAAGTCGATG
NZ_CP035260.1:726156-733994    4450   TGGAAAATATTGCCTTAGGAAAACAGGTTACTCAAAGTTCGACAGCTTTCGGAGGAGATGCTAGAAGAGCTGTGGATGGCAAAGTCGATG
NZ_CP035241.1:708296-716144    4460   TGGAAAATATTGCCTTAGGAAAACAGGTTACTCAAAGTTCGACAGCTTTCGGAGGAGATGCTAGAAGAGCTGTGGATGGCAAAGTCGATG
NZ_CP035261.1:779086-786938    4460   TGGAAAATATTGCCTTAGGAAAACAGGTTACTCAAAGTTCGACAGTTTTCGGAGGAGATGCTAGAAGAGCTGTGGATGGCAAAGTCGATG
NZ_CP035242.1:c1532236-1524385 4459   TGGAAAATATTGCCTTAGGAAAACAGGTTACTCAAAGTTCGACAGTTTTCGGAGGAGATGCTAGAAGAGCTGTGGATGGCAAAGTCGATG
NZ_CP035243.1:777729-785580    4459   TGGAAAATATTGCCTTAGGAAAACAGGTTACTCAAAGTTCGACAGTTTTCGGAGGAGATGCTAGAAGAGCTGTGGATGGCAAAGTCGATG
NZ_CP035237.1:c1311322-1303456 4474   TGGAAAATATTGCCTTAGGAAAACAGGTTACTCAAAGTTCGACAGCTTTCGGAGGAGATGCTAGAAGAGCTGTGGATGGCAAAGTCGATG
NZ_CP035246.1:c1302892-1295026 4474   TGGAAAATATTGCCTTAGGAAAACAGGTTACTCAAAGTTCGACAGCTTTCGGAGGAGATGCTAGAAGAGCTGTGGATGGCAAAGTCGATG
NZ_CP035247.1:c1301115-1293263 4460   TGGAAAATATTGCCTTAGGAAAACAGGTTACTCAAAGTTCGACAGCTTTCGGAGGAGATGCTAGAAGAGCTGTGGATGGCAAAGTCGATG
NZ_CP035238.1:738547-746413    4474   TGGAAAATATTGCCTTAGGAAAATAGGTTACTCAAAGTTCGACAGCTTTCGGAGGAGATGCTAGAAGAGCTGTGGATGGCAAAGTCGATG
NZ_CP035255.1:776866-784718    4460   TGGAAAATATTGCCTTAGGAAAACAGGTTACTCAAAGTTCGACAGCTTTCGGAGGAGATGCTAGAAGAGCTGTGGATGGCAAAGTCGATG
NZ_CP035256.1:807236-815102    4474   TGGAAAATATTGCCTTAGGAAAACAGGTTACTCAAAGTTCGACAGCTTTCGGAGGAGATGCTAGAAGAGCTGTGGATGGCAAAGTCGATG
NZ_CP038252.1:1151812-1159673  4403   TGGAAAATATTGCCTTAGGAAAACAGGTTACTCAAAGTTCGACAGTTTTCGGAGGAGATGCTAGAAGAGCTGTGGATGGCAAAGTCGATG
NZ_CP035254.1:793365-801213    4460   TGGAAAATATTGCCTTAGGAAAACAGGTTACTCAAAGTTCGACAGCTTTCGGAGGAGATGCTAGAAGAGCTGTGGATGGCAAAGTCGATG
NZ_CP035251.1:748733-756585    4463   TGGAAAATATTGCCTTAGGAAAACAGGTTACTCAAAGTTCGACAGCTTTCGGAGGAGATGCTAGAAGAGCTGTGGATGGCAAAGTCGATG
NZ_CP035252.1:745670-753536    4477   TGGAAAATATTGCCTTAGGAAAACAGGTTACTCAAAGTTCGACAGCTTTCGGAGGAGATGCTAGAAGAGCTGTGGATGGCAAAGTCGATG
NZ_CP035257.1:776317-784169    4463   TGGAAAATATTGCCTTAGGAAAACAGGTTACTCAAAGTTCGACAGCTTTCGGAGGAGATGCTAGAAGAGCTGTGGATGGCAAAGTCGATG
NZ_CP038253.1:758105-765957    4463   TGGAAAATATTGCCTTAGGAAAACAGGTTACTCAAAGTTCGACAGCTTTCGGAGGAGATGCTAGAAGAGCTGTGGATGGCAAAGTCGATG
NZ_CP035263.1:743050-750901    4459   TGGAAAATATTGCCTTAGGAAAACAGGTTACTCAAAGTTCGACAGCTTTCGGAGGAGATGCTAGAAGAGCTGTGGATGGCAAAGTCGATG
NZ_CP035262.1:741595-749460    4473   TGGAAAATATTGCCTTAGGAAAACAGGTTACTCAAAGTTCGACAGCTTTCGGAGGAGATGCTAGAAGAGCTGTGGATGGCAAAGTCGATG
NZ_CP035236.1:764915-772766    4459   TGGAAAATATTGCCTTAGGAAAACAGGTTACTCAAAGTTCGACAGCTTTCGGAGGAGATGCTAGAAGAGCTGTGGATGGCAAAGTCGATG
NZ_CP035253.1 767286-775134    4460   TGGAAAATATTGCCTTAGGAAAACAGGTTACTCAAAGTTCGACAGCTTTCGGAGGAGATGCTAGAAGAGCTGTGGATGGCAAAGTCGATG
NZ_CP035249.1:763080-770921    4460   TGGAAAATATTGCCTTAGGAAAACAGGTTACTCAAAGTTCGACAGCTTTCGGAGGAGATGCTAGAAGAGCTGTGGATGGCAAAGTCGATG

NC_003028.3:777000-787000      5279   GTAACTATGGTCACAATTCTGTCACTCATACAAACTTCCAATCTAAGCCTTGGTGGCAAGTAGATTTGGCTAAAGAAGAAACCATTCGCC
NZ_CP035248.1:765099-775111    5288   GTAACTATGGTCACAATTCTGTCACTCATACAAACTTCCAATCTAAGCCTTGGTGGCAAGTAGATTTGGCTAAAGAAGAAACCATTCGCC
NZ_CP035235.1 c1326834-1317894 4564   GTAACTATGGTCACAATTCTGTCACTCATACAAACTTCCAATCTAAGCCTTGGTGGCAAGTAGATTTGGCTAAAGAAGAAACCATTCGCC
NZ_CP035234.1:756284-765224    4564   GTAACTATGGTCACAATTCTGTCACTCATACAAACTTCCAATCTAAGCCTTGGTGGCAAGTAGATTTGGCTAAAGAAGAAACCATTCGCC
NZ_CP038251.1:750745-759686    4564   GTAACTATGGTCACAATTCTGTCACTCATACAAACTTCCAATCTAAGCCTTGGTGGCAAGTAGATTTGGCTAAAGAAGAAACCATTCGCC
NZ_CP035244.1:747023-755950    4550   GTAACTATGGTCACAATTCTGTCACTCATACAAACTTCCAATCTAAGCCTTGGTGGCAAGTAGATTTGGCTAAAGAAGAAACCATTCGCC
NZ_CP035264.1:800836-809336    5198   GTAACTATGGTCACAATTCTGTCACTCATACAAACTTCCAATCTAAGCCTTGGTGGCAAGTAGATTTGGCTAAAGAAGAAACCATTCGCC
NZ_CP035265.1:766128-774627    5198   GTAACTATGGTCACAATTCTGTCACTCATACAAACTTCCAATCTAAGCCTTGGTGGCAAGTAGATTTGGCTAAAGAAGAAACCATTCGCC
NZ_CP035245.1 745261-753757    5198   GTAACTATGGTCACAATTCTGTCACTCATACAAACTTCCAATCTAAGCCTTGGTGGCAAGTAGATTTGGCTAAAGAAGAAACCATTCGCC
NZ_CP035259.1:745700-753548    4550   GTAACTATGGTCACAATTCTGTCACTCATACAAACTTCCAATCTAAGCCTTGGTGGCAAGTAGATTTGGCTAAAGAAGAAACCATTCGCC
NZ_CP035258.1:750874-758722    4550   GTAACTATGGTCACAATTCTGTCACTCATACAAACTTCCAATCTAAGCCTTGGTGGCAAGTAGATTTGGCTAAAGAAGAAACCATTCGCC
NZ_CP035260.1:726156-733994    4540   GTAACTATGGTCACAATTCTGTCACTCATACAAACTTCCAATCTAAGCCTTGGTGGCAAGTAGATTTGGCTAAAGAAGAAACCATTCGCC
NZ_CP035241.1:708296-716144    4550   GTAACTATGGTCACAATTCTGTCACTCATACAAACTTCCAATCTAAGCCTTGGTGGCAAGTAGATTTGGCTAAAGAAGAAACCATTCGCC
NZ_CP035261.1:779086-786938    4550   GTAACTATGGTCACAATTCTGTCACTCATACAAACTTCCAATCTAAGCCTTGGTGGCAAGTAGATTTGGCTAAAGAAGAAACCATTCGCC
NZ_CP035242.1:c1532236-1524385 4549   GTAACTATGGTCACAATTCTGTCACTCATACAAACTTCCAATCTAAGCCTTGGTGGCAAGTAGATTTGGCTAAAGAAGAAACCATTCGCC
NZ_CP035243.1:777729-785580    4549   GTAACTATGGTCACAATTCTGTCACTCATACAAACTTCCAATCTAAGCCTTGGTGGCAAGTAGATTTGGCTAAAGAAGAAACCATTCGCC
NZ_CP035237.1:c1311322-1303456 4564   GTAACTATGGTCACAATTCTGTCACTCATACAAACTTCCAATCTAAGCCTTGGTGGCAAGTAGATTTGGCTAAAGAAGAAACCATTCGCC
NZ_CP035246.1:c1302892-1295026 4564   GTAACTATGGTCACAATTCTGTCACTCATACAAACTTCCAATCTAAGCCTTGGTGGCAAGTAGATTTGGCTAAAGAAGAAACCATTCGCC
NZ_CP035247.1:c1301115-1293263 4550   GTAACTATGGTCACAATTCTGTCACTCATACAAACTTCCAATCTAAGCCTTGGTGGCAAGTAGATTTGGCTAAAGAAGAAACCATTCGCC
NZ_CP035238.1:738547-746413    4564   GTAACTATGGTCACAATTCTGTCACTCATACAAACTTTCAATCTAAGCCTTGGTGGCAAGTAGATTTGGCTAAAGAAGAAACCATTCGCC
NZ_CP035255.1:776866-784718    4550   GTAACTATGGTCACAATTCTGTCACTCATACAAACTTTCAATCTAAGCCTTGGTGGCAAGTAGATTTGGCTAAAGAAGAAACCATTCGCC
NZ_CP035256.1:807236-815102    4564   GTAACTATGGTCACAATTCTGTCACTCATACAAACTTTCAATCTAAGCCTTGGTGGCAAGTAGATTTGGCTAAAGAAGAAACCATTCGCC
NZ_CP038252.1:1151812-1159673  4493   GTAACTATGGTCACAATTCTGTCACTCATACAAACTTCCAATCTAAGCCTTGGTGGCAAGTAGATTTGGCTAAAGAAGAAACCATTCGCC
NZ_CP035254.1:793365-801213    4550   GTAACTATGGTCACAATTCTGTCACTCATACAAACTTCCAATCTAAGCCTTGGTGGCAAGTAGATTTGGCTAAAGAAGAAACCATTCGCC
NZ_CP035251.1:748733-756585    4553   GTAACTATGGTCACAATTCTGTCACTCATACAAACTTCCAATCTAAGCCTTGGTGGCAAGTAGATTTGGCTAAAGAAGAAACCATTCGCC
NZ_CP035252.1:745670-753536    4567   GTAACTATGGTCACAATTCTGTCACTCATACAAACTTCCAATCTAAGCCTTGGTGGCAAGTAGATTTGGCTAAAGAAGAAACCATTCGCC
NZ_CP035257.1:776317-784169    4553   GTAACTATGGTCACAATTCTGTCACTCATACAAACTTCCAATCTAAGCCTTGGTGGCAAGTAGATTTGGCTAAAGAAGAAACCATTCGCC
NZ_CP038253.1:758105-765957    4553   GTAACTATGGTCACAATTCTGTCACTCATACAAACTTCCAATCTAAGCCTTGGTGGCAAGTAGATTTGGCTAAAGAAGAAACCATTCGCC
NZ_CP035263.1:743050-750901    4549   GTAACTATGGTCACAATTCTGTCACTCATACAAACTTCCAATCTAAGCCTTGGTGGCAAGTAGATTTGGCTAAAGAAGAAACCATTCGCC
NZ_CP035262.1:741595-749460    4563   GTAACTATGGTCACAATTCTGTCACTCATACAAACTTCCAATCTAAGCCTTGGTGGCAAGTAGATTTGGCTAAAGAAGAAACCATTCGCC
NZ_CP035236.1:764915-772766    4549   GTAACTATGGTCACAATTCTGTCACTCATACAAACTTCCAATCTAAGCCTTGGTGGCAAGTAGATTTGGCTAAAGAAGAAACCATTCGCC
NZ_CP035253.1 767286-775134    4550   GTAACTATGGTCACAATTCTGTCACTCATACAAACTTCCAATCTAAGCCTTGGTGGCAAGTAGATTTGGCTAAAGAAGAAACCATTCGCC
NZ_CP035249.1:763080-770921    4550   GTAACTATGGTCACAATTCTGTCACTCATACAAACTTCCAATCTAAGCCTTGGTGGCAAGTAGATTTGGCTAAAGAAGAAACCATTCGCC

NC_003028.3:777000-787000      5369   AAATCAATATTTACAACCGAACAGACACTGCCCAGGATAGATTGGCAAACTTTGATGTCATTCTTTTAGACAGTTCTGGTAAAGAAATTG
NZ_CP035248.1:765099-775111    5378   AAATCAATATTTACAACCGAACAGACACTGCCCAGGATAGATTGGCAAACTTTGATGTCATTCTTTTAGACAGTTCTGGTAAAGAAATTG
```

```
NZ_CP035235.1 c1326834-1317894 4654   AAATCAATATTTACAACCGAACAGACACTGCCCAGGATAGATTGGCAAACTTTGATGTCATTCTTTTAGACAGTTCTGGTAAAGAAATTG
NZ_CP035234.1:756284-765224    4654   AAATCAATATTTACAACCGAACAGACACTGCCCAGGATAGATTGGCAAACTTTGATGTCATTCTTTTAGACAGTTCTGGTAAAGAAATTG
NZ_CP038251.1:750745-759686    4654   AAATCAATATTTACAACCGAACAGACACTGCCCAGGATAGATTGGCAAACTTTGATGTCATTCTTTTAGACAGTTCTGGTAAAGAAATTG
NZ_CP035244.1:747023-755950    4640   AAATCAATATTTACAACCGAACAGACACTGCCCAGGATAGATTGGCAAACTTTGATGTCATTCTTTTAGACAGTTCTGGTAAAGAAATTG
NZ_CP035264.1:800836-809336    5288   AAATCAATATTTACAACCGAACAGACACTGCCCAGGATAGATTGGCAAACTTTGATGTCATTCTTTTAGACAGTTCTGGTAAAGAAATTG
NZ_CP035265.1:766128-774627    5288   AAATCAATATTTACAACCGAACAGACACTGCCCAGGATAGATTGGCAAACTTTGATGTCATTCTTTTAGACAGTTCTGGTAAAGAAATTG
NZ_CP035245.1 745261-753757    5288   AAATCAATATTTACAACCGAACAGACACTGCCCAGGATAGATTGGCAAACTTTGATGTCATTCTTTTAGACAGTTCTGGTAAAGAAATTG
NZ_CP035259.1:745700-753548    4640   AAATCAATATTTACAACCGAACAGACACTGCCCAGGATAGATTGGCAAACTTTGATGTCATTCTTTTAGACAGTTCTGGTAAAGAAATTG
NZ_CP035258.1:750874-758722    4640   AAATCAATATTTACAACCGAACAGACACTGCCCAGGATAGATTGGCAAACTTTGATGTCATTCTTTTAGACAGTTCTGGTAAAGAAATTG
NZ_CP035260.1:726156-733994    4630   AAATCAATATTTACAACCGAACAGACACTGCCCAGGATAGATTGGCAAACTTTGATGTCATTCTTTTAGACGGTTCTGGTAAAGAAATTG
NZ_CP035241.1:708296-716144    4640   AAATCAATATTTACAACCGAACAGACACTGCCCAGGATAGATTGGCAAACTTTGATGTCATTCTTTTAGACGGTTCTGGTAAAGAAATTG
NZ_CP035261.1:779086-786938    4640   AAATCAATATTTACAACCGAACAGACACTGCCCAGGATAGATTGGCAAACTTTGATGTCATTCTTTTAGACAGTTCTGGTAAAGAAATTG
NZ_CP035242.1:c1532236-1524385 4639   AAATCAATATTTACAACCGAACAGACACTGCCCAGGATAGATTGGCAAACTTTGATGTCATTCTTTTAGACAGTTCTGGTAAAGAAATTG
NZ_CP035243.1:777729-785580    4639   AAATCAATATTTACAACCGAACAGACACTGCCCAGGATAGATTGGCAAACTTTGATGTCATTCTTTTAGACAGTTCTGGTAAAGAAATTG
NZ_CP035237.1:c1311322-1303456 4654   AAATCAATATTTACAACCGAACAGACACTGCCCAGGATAGATTGGCAAACTTTGATGTCATTCTTTTAGACAGTTCTGGTAAAGAAATTG
NZ_CP035246.1:c1302892-1295026 4654   AAATCAATATTTACAACCGAACAGACACTGCCCAGGATAGATTGGCAAACTTTGATGTCATTCTTTTAGACAGTTCTGGTAAAGAAATTG
NZ_CP035247.1:c1301115-1293263 4640   AAATCAATATTTACAACCGAACAGACACTGCCCAGGATAGATTGGCAAACTTTGATGTCATTCTTTTAGACAGTTCTGGTAAAGAAATTG
NZ_CP035238.1:738547-746413    4654   AAATCAATATTTACAACCGAACAGACACTGCCCAGGATAGATTGGCAAACTTTGATGTCATTCTTTTAGACAGTTCTGGTAAAGAAATTG
NZ_CP035255.1:776866-784718    4640   AAATCAATATTTACAACCGAACAGACACTGCCCAGGATAGATTGGCAAACTTTGATGTCATTCTTTTAGACAGTTCTGGTAAAGAAATTG
NZ_CP035256.1:807236-815102    4654   AAATCAATATTTACAACCGAACAGACACTGCCCAGGATAGATTGGCAAACTTTGATGTCATTCTTTTAGACAGTTCTGGTAAAGAAATTG
NZ_CP038252.1:1151812-1159673  4583   AAATCAATATTTACAACCGAACAGACACTGCCCAGGATAGATTGGCAAACTTTGATGTCATTCTTTTAGACAGTTCTGGTAAAGAAATTG
NZ_CP035254.1:793365-801213    4640   AAATCAATATTTACAACCGAACAGACACTGCCCAGGATAGATTGGCAAACTTTGATGTCATTCTTTTAGACAGTTCTGGTAAAGAAATTG
NZ_CP035251.1:748733-756585    4643   AAATCAATATTTACAACCGAACAGACACTGCCCAGGATAGATTGGCAAACTTTGATGTCATTCTTTTAGACAGTTCTGGTAAAGAAATTG
NZ_CP035252.1:745670-753536    4657   AAATCAATATTTACAACCGAACAGACACTGCCCAGGATAGATTGGCAAACTTTGATGTCATTCTTTTAGACAGTTCTGGTAAAGAAATTG
NZ_CP035257.1:776317-784169    4643   AAATCAATATTTACAACCGAACAGACACTGCCCAGGATAGATTGGCAAACTTTGATGTCATTCTTTTAGACAGTTCTGGTAAAGAAATTG
NZ_CP038253.1:758105-765957    4643   AAATCAATATTTACAACCGAACAGACACTGCCCAGGATAGATTGGCAAACTTTGATGTCATTCTTTTAGACAGTTCTGGTAAAGAAATTG
NZ_CP035263.1:743050-750901    4639   AAATCAATATTTACAACCGAACAGACACTGTCCAGGATAGATTGGCAAACTTTGATGTCATTCTTTTAGACAGTTCTGGTAAAGAAATTG
NZ_CP035262.1:741595-749460    4653   AAATCAATATTTACAACCGAACAGACACTGCCCAGGATAGATTGGCAAACTTTGATGTCATTCTTTTAGACAGTTCTGGTAAAGAAATTG
NZ_CP035236.1:764915-772766    4639   AAATCAATATTTACAACCGAACAGACACTGCCCAGGATAGATTGGCAAACTTTGATGTCATTCTTTTAGACAGTTCTGGTAAAGAAATTG
NZ_CP035253.1 767286-775134    4640   AAATCAATATTTACAACCGAACAGACACTGCCCAGGATAGATTGGCAAACTTTGATGTCATTCTTTTAGACAGTTCTGGTAAAGAAATTG
NZ_CP035249.1:763080-770921    4640   AAATCAATATTTACAACCGAACAGACACTGCCCAGGATAGATTGGCAAACTTTGATGTCATTCTTTTAGACAGTTCTGGTAAAGAAATTG

NC_003028.3:777000-787000      5459   AGTGAAAACGTATAACATCTCCTAAAGATGTGTCAGCACAAATTACGATTAACCAT-AAAAAAGCGCGCTATGTTCGGATTGAGCTAGAA
NZ_CP035248.1:765099-775111    5468   AGTAAAAACGTATAACATCTCCTAAAGATGTGTCAGCACAAATTACGATTAACCAT-AAAAAAGTGCGCTATGTTCGGATTGAGCTAGAA
NZ_CP035235.1 c1326834-1317894 4744   AGTGAAAACGTATAATATCTCCTAAAGATGTGTCAGCACAAATTACGATTAACCAT--AAAAAACGCGCTATGTTCGGATTGAGCTAGAA
NZ_CP035234.1:756284-765224    4744   AGTGAAAACGTATAATATCTCCTAAAGATGTGTCAGCACAAATTACGATTAACCAT--AAAAAACGCGCTATGTTCGGATTGAGCTAGAA
NZ_CP038251.1:750745-759686    4744   AGTGAAAACGTATAATATCTCCTAAAGATGTGTCAGCACAAATTACGATTAACCAT--AAAAAACGCGCTATGTTCGGATTGAGCTAGAA
NZ_CP035244.1:747023-755950    4730   AGTGAAAACGTATAATATCTCCTAAAGATGTGTCAGCACAAATTACGATTAACCAT--AAAAAACGCGCTATGTTCGGATTGAGCTAGAA
NZ_CP035264.1:800836-809336    5378   AGTGAAAACGTATAACATCTCCTAAAGATGTGTCAGCACAAATTACGATTAACCAT-AAAAAAGTGCGCTATGTTCGGATTGAGCTAGAA
NZ_CP035265.1:766128-774627    5378   AGTGAAAACGTATAACATCTCCTAAAGATGTGTCAGCACAAATTACGATTAACCAT--AAAAAGTGCGCTATGTTCGGATTGAGCTAGAA
NZ_CP035245.1 745261-753757    5378   AGTGAAAACGTATAATATCTCCTAAAGATGTGTCAGCACAAATTACGATTAACCAT-AAAAAAGCGCGCTATTTTCGGATTGAGCTAGAA
NZ_CP035259.1:745700-753548    4730   AGTGAAAACGTATAACATCTCCTAAAGATGTGTCAGCACAAATTACGATTAACCAT-AAAAAAGCGCGCTATGTTCGGATTGAGCTAGAA
NZ_CP035258.1:750874-758722    4730   AGTGAAAACGTATAACATCTCCTAAAGATGTGTCAGCACAAATTACGATTAACCAT-AAAAAAGCGCGCTATGTTCGGATTGAGCTAGAA
NZ_CP035260.1:726156-733994    4720   AGTGAAAACGTATAATATCTCCTAAAGATGTGTCAGCACAAATTACGATTAACCAT-AAAAAAGCGCGCTATGTTCGGATTGAGCTAGAA
NZ_CP035241.1:708296-716144    4730   AGTGAAAACGTATAATATCTCCTAAAGATGTGTCAGCACAAATTACGATTAACCAT-AAAAAAGCGCGCTATGTTCGGATTGAGCTAGAA
NZ_CP035261.1:779086-786938    4730   AGTGAAAACGTATAACATCTCCTAAAGATGTGTCAGCACAAATTACGATTAACCAT-AAAAAAGTGCGCTATGTTCGGATTGAGCTAGAA
NZ_CP035242.1:c1532236-1524385 4729   AGTAAAAACGTATAACATCTCCTAAAGATGTGTCAGCACAAATTACGATTAACCAT-AAAAAAGTGCGCTATGTTCGGATTGAGCTAGAA
NZ_CP035243.1:777729-785580    4729   AGTAAAAACGTATAACATCTCCTAAAGATGTGTCAGCACAAATTACGATTAACCAT-AAAAAAGTGCGCTATGTTCGGATTGAGCTAGAA
NZ_CP035237.1:c1311322-1303456 4744   AGTGAAAACGTATAATATCTCCTAAAGATGTGTCAGCACAAATTACGATTAACCAT-AAAAAAGCGCGCTATGTTCGGATTGAGCTAGAA
NZ_CP035246.1:c1302892-1295026 4744   AGTGAAAACGTATAACATCTCCTAAAGATGTGTCAGCACAAATTACGATTAACCAT-AAAAAAGCGCGCTATGTTCGGATTGAGCTAGAA
NZ_CP035247.1:c1301115-1293263 4730   AGTGAAAACGTATAACATCTCCTAAAGATGTGTCAGCACAAATTACGATTAACCAT-AAAAAAGCGCGCTATGTTCGGATTGAGCTAGAA
NZ_CP035238.1:738547-746413    4744   AGTGAAAACGTATAACATCTCCTAAAGATGTGTCAGCACAAATTACGATTAACCAT-AAAAAAGCGCGCTATGTTCGGATTGAGCTAGAA
NZ_CP035255.1:776866-784718    4730   AGTGAAAACGTATAACATCTCCTAAAGATGTGTCAGCACAAATTACGATTAACCAT-AAAAAAGCGCGCTATGTTCGGATTGAGCTAGAA
NZ_CP035256.1:807236-815102    4744   AGTGAAAACGTATAACATCTCCTAAAGATGTGTCAGCACAAATTACGATTAACCAT-AAAAAAGCGCGCTATGTTCGGATTGAGCTAGAA
NZ_CP038252.1:1151812-1159673  4673   AGTGAAAACGTATAACATCTCCTAAAGATGTGTCAGCACAAATTACGATTAACCAT-AAAAAAGTGCGCTATGTTCGGATTGAGCTAGAA
NZ_CP035254.1:793365-801213    4730   AGTGAAAACGTATAATATCTCCTAAAGATGTGTCAGCACAAATTACGATTAACCAT-AAAAAAGTACGCTATGTTCGGATTGAGCTAGAA
NZ_CP035251.1:748733-756585    4733   AGTGAAAACGTATAACATCTCCTAAAGATGTGTCAGCACAAATTACGATTAACCATAAAAAAAGCGCGCTATGTTCGGATTGAGCTAGAA
NZ_CP035252.1:745670-753536    4747   AGTGAAAACGTATAACATCTCCTAAAGATGTGTCAGCACAAATTACGATTAACCATAAAAAAAGCGCGCTATGTTCGGATTGAGCTAGAA
NZ_CP035257.1:776317-784169    4733   AGTGAAAACGTATAACATCTCCTAAAGATGTGTCAGCACAAATTACGATTAACCATAAAAAAAGCGCGCTATGTTCGGATTGAGCTAGAA
NZ_CP038253.1:758105-765957    4733   AGTGAAAACGTATAACATCTCCTAAAGATGTGTCAGCACAAATTACGATTAACCATAAAAAAAGCGCGCTATGTTCGGATTGAGCTAGAA
NZ_CP035263.1:743050-750901    4729   AGTGAAAACGTATAATATCTCCTAAAGATGTGTCAGCACAAATTACGATTAACCAT-AAAAAAGCGCGCTATGTTCGGATTGGGCTAGAA
NZ_CP035262.1:741595-749460    4743   AGTGAAAACGTATAATATCTCCTAAAGATGTGTCAGCACAAATTACGATTAACCAT-AAAAAAGCGCGCTATGTTCGGATTGGGCTAGAA
NZ_CP035236.1:764915-772766    4729   AGTGAAAACGTATAATATCTCCTAAAGATGTGTCAGCACAAATTACGATTAACCAT-AAAAAAGCGCGCTATGTTCGGATTGGGCTAGAA
NZ_CP035253.1 767286-775134    4730   AGTGAAAACGTATAATATCTCCTAAAGATGTGTCAGCACAAATTACGATTAACCAT-AAAAAAGTACGCTATGTTCGGATTGAGCTAGAA
NZ_CP035249.1:763080-770921    4730   AGTGAAAACGTATAATATCTCCTAAAGATGTGTCAGCACAAATTACGATTAACCAT-AAAAAAGCGCGCTATGTTCGGATTGAGCTAGAA

NC_003028.3:777000-787000      5548   GGCTATAATGCCCTCAGTCTTGCAGAAGTTGAAGTTTTCTGCTTTATAGCTACGAATGCTGAAACGGCGACACAAGTTTCTAAGCCAGTT
NZ_CP035248.1:765099-775111    5557   GGCTATAATGCCCTCAGTCTTGCAGAAGTTGAAGTTTTCCGCTTTATAGCTACGAATGCTGAAACGGCGACACAAGTTTCTAAGCCAGTT
NZ_CP035235.1 c1326834-1317894 4832   GGCTATAATGCCCTCAGTCTTGCAGAAGTTGAAGTTTTCCGCTTTATAGCTACGAATGCTGAAACGGCGACACAAGTTTCTAAGCCAGTT
NZ_CP035234.1:756284-765224    4832   GGCTATAATGCCCTCAGTCTTGCAGAAGTTGAAGTTTTCCGCTTTATAGCTACGAATGCTGAAACGGCGACACAAGTTTCTAAGCCAGTT
NZ_CP038251.1:750745-759686    4832   GGCTATAATGCCCTCAGTCTTGCAGAAGTTGAAGTTTTCCGCTTTATAGCTACGAATGCTGAAACGGCGACACAAGTTTCTAAGCCAGTT
NZ_CP035244.1:747023-755950    4818   GGCTATAATGCCCTCAGTCTTGCAGAAGTTGAAGTTTTCCGCTTTATAGCTACGAATGCTGAAACGGCGACACAAGTTTCTAAGCCAGTT
NZ_CP035264.1:800836-809336    5467   GGCTATAATGCCCTCAGTCTTGCAGAAGTTGAAGTTTTCCGCTTTATAGCTACGAATGCTGAAACGGCGACACAAGTTTCTAAGCCAGTT
NZ_CP035265.1:766128-774627    5466   GGCTATAATGCCCTCAGTCTTGCAGAAGTTGAAGTTTTCCGCTTTATAGCTACGAATGCTGAAACGGCGACACAAGTTTCTAAGCCAGTT
NZ_CP035245.1 745261-753757    5467   GGCTATAATGCCCTCAGTCTTGCAGAAGTTGAAGTTTTCCGCTTTATAGCTACGAATGCTGAAACGGCGACACAAGTTTCTAAGCCAGTT
NZ_CP035259.1:745700-753548    4819   GGCTATAATGCCCTCAGTCTTGCAGAAGTTGAAGTTTTCTGCTTTATAGCTACGAATGCTGAAACGGCGACACAAGTTTCTAAGCCAGTT
NZ_CP035258.1:750874-758722    4819   GGCTATAATGCCCTCAGTCTTGCAGAAGTTGAAGTTTTCTGCTTTATAGCTACGAATGCTGAAACGGCGACACAAGTTTCTAAGCCAGTT
NZ_CP035260.1:726156-733994    4809   GGCTATAATGCCCTCAGTCTTGCAGAAGTTGAAGTTTTCCGCTTTATAGCTACGAATGGTGAAACGGCGATACAAGTTTCTAAGTCAGTT
NZ_CP035241.1:708296-716144    4819   GGCTATAATGCCCTCAGTCTTGCAGAAGTTGAAGTTTTCCGCTTTATAGCTACGAATGGTGAAACGGCGATACAAGTTTCTAAGTCAGTT
NZ_CP035261.1:779086-786938    4819   GGCTATAATGCCCTCAGTCTTGCAGAAGTTGAAGTTTTCCGCTTTATAGCTACGAATGCTGAAACGGCGACACAAGTTTCTAAGCCAGTT
NZ_CP035242.1:c1532236-1524385 4818   GGCTATAATGCCCTCAGTCTTGCAGAAGTTGAAGTTTTCCGCTTTATAGCTACGAATGCTGAAACGGCGACACAAGTTTCTAAGCCAGTT
NZ_CP035243.1:777729-785580    4818   GGCTATAATGCCCTCAGTCTTGCAGAAGTTGAAGTTTTCCGCTTTATAGCTACGAATGCTGAAACGGCGACACAAGTTTCTAAGCCAGTT
NZ_CP035237.1:c1311322-1303456 4833   GGCTATAATGCCCTCAGTCTTGCAGAAGTTGAAGTTTTCCGCTTTATAGCTACGAATGCTGAAACGGCGACACAAGTTTCTAAGCCAGTT
NZ_CP035246.1:c1302892-1295026 4833   GGCTATAATGCCCTCAGTCTTGCAGAAGTTGAAGTTTTCCGCTTTATAGCTACGAATGGTGAAACGGCGATACAAGTTTCTAAGTCAGTT
NZ_CP035247.1:c1301115-1293263 4819   GGCTATAATGCCCTCAGTCTTGCAGAAGTTGAAGTTTTCCGCTTTATAGCTACGAATGGTGAAACGGCGATACAAGTTTCTAAGTCAGTT
NZ_CP035238.1:738547-746413    4833   GGCTATAATGCCCTCAGTCTTGCAGAAGTTGAAGTTTTCCGCTTTATAGCTACGAATGGTGAAACGGCGATACAAGTTTCTAAGTCAGTT
NZ_CP035255.1:776866-784718    4819   GGCTATAATGCCCTCAGTCTTGCAGAAGTTGAAGTTTTCCGCTTTATAGCTACGAATGGTGAAACGGCGATACAAGTTTCTAAGTCAGTT
NZ_CP035256.1:807236-815102    4833   GGCTATAATGCCCTCAGTCTTGCAGAAGTTGAAGTTTTCCGCTTTATAGCTACGAATGGTGAAACGGCGATACAAGTTTCTAAGTCAGTT
NZ_CP038252.1:1151812-1159673  4762   GGCTATAATGCCCTCAGTCTTGCAGAAGTTGAAGTTTTCCGCTTTATAGCTACGAATGCTGAAACGGCGACACAAGTTTCTAAGCCAGTT
NZ_CP035254.1:793365-801213    4819   GGCTATAATGCCCTCAGTCTTGCAGAAGTTGAAGTTTTCCGCTTTATAGCTACGAATGCTGAAACGGCGACACAAGTTTCTAAGCCAGTT
NZ_CP035251.1:748733-756585    4823   GGCTATAATGCCCTCAGTCTTGCAGAAGTTGAAGTTTTCCGCTTTATAGCTACGAATGGTGAAACGGCGATACAAGTTTCTAAGTCAGTT
NZ_CP035252.1:745670-753536    4837   GGCTATAATGCCCTCAGTCTTGCAGAAGTTGAAGTTTTCCGCTTTATAGCTACGAATGGTGAAACGGCGATACAAGTTTCTAAGTCAGTT
NZ_CP035257.1:776317-784169    4823   GGCTATAATGCCCTCAGTCTTGCAGAAGTTGAAGTTTTCCGCTTTATAGCTACGAATGGTGAAACGGCGATACAAGTTTCTAAGTCAGTT
NZ_CP038253.1:758105-765957    4823   GGCTATAATGCCCTCAGTCTTGCAGAAGTTGAAGTTTTCCGCTTTATAGCTACGAATGGTGAAACGGCGATACAAGTTTCTAAGTCAGTT
NZ_CP035263.1:743050-750901    4818   GGCTATAATGCCCTCAGTCTTGCAGAAGTTGAAGTTTTCCGCTTTATAGCTACGAATGCTGAAACGGCGACACAAGTTTCTAAGCCAGTT
```

### Ozkan et al. Supplementary Figures

```
NZ_CP035262.1:741595-749460    4832   GGCTATAATGCCCTCAGTCTTGCAGAAGTTGAAGTTTTCCGCTTTATAGCTACGAATGCTGAAACGGCGACACAAGTTTCTAAGCCAGTT
NZ_CP035236.1:764915-772766    4818   GGCTATAATGCCCTCAGTCTTGCAGAAGTTGAAGTTTTCCGCTTTATAGCTACGAATGCTGAAACGGCGACACAAGTTTCTAAGCCAGTT
NZ_CP035253.1 767286-775134    4819   GGCTATAATGCCCTCAGTCTTGCAGAAGTTGAAGTTTTCCGCTTTATAGCTACGAATGCTGAAACGGCGACACAAGTTTCTAAGCCAGTT
NZ_CP035249.1:763080-770921    4819   GGCTATAATGCCCTCAGTCTTGCAGAAGTTGAAGTTTTCCGCTTTATAGCTACGAATGCTGAAACGGCGACACAAGTTTCTAAGCCAGTT

NC_003028.3:777000-787000      5638   CAACCAATCAGTCAGACTCCTGTGAAGGATAAAACATTGACAATTCAACACAGTGGAGCTTACATTGCCCGCTACTCCATAACTTGGGAA
NZ_CP035248.1:765099-775111    5647   CAACCAATCAGTCAGACTCCTGTGAAAGATAAAACATTGACAATTCAACACAGTGGAGCTTACATTGCCCGCTACTCCATAACTTGGGAA
NZ_CP035235.1 c1326834-1317894 4922   CAACCAATCAGTCAGACTCCTGTGAAAGATAAAACATTGACAATTCAACACAGTGGAGCTTACATTGCCCGCTACTCCATAACTTGGGAA
NZ_CP035234.1:756284-765224    4922   CAACCAATCAGTCAGACTCCTGTGAAAGATAAAACATTGACAATTCAACACAGTGGAGCTTACATTGCCCGCTACTCCATAACTTGGGAA
NZ_CP038251.1:750745-759686    4922   CAACCAATCAGTCAGACTCCTGTGAAAGATAAAACATTGACAATTCAACACAGTGGAGCTTACATTGCCCGCTACTCCATAACTTGGGAA
NZ_CP035244.1:747023-755950    4908   CAACCAATCAGTCAGACTCCTGTGAAAGATAAAACATTGACAATTCAACACAGTGGAGCTTACATTGCCCGCTACTCCATAACTTGGGAA
NZ_CP035264.1:800836-809336    5557   CAACCAATCAGTCAGACTCCTGTGAAAGATAAAACATTGACAATTCAACATAGTGGAGCTTACATTGCCCGCTACTCCATAACTTGGGAA
NZ_CP035265.1:766128-774627    5556   CAACCAATCAGTCAGACTCCTGTGAAAGATAAAACATTGACAATTCAACATAGTGGAGCTTACATTGCCCGCTACTCCATAACTTGGGAA
NZ_CP035245.1 745261-753757    5557   CAACCAATCAGTCAGACTCCTGTGAAAGATAAAACATTGATAATTCAACACAGTGGAGCTTACATTGCCCGCTACTCCATAACTTGGGAA
NZ_CP035259.1:745700-753548    4909   CAACCAATCAGTCAGACTCCTGTGAAGGATAAAACATTGACAATTCAACACAGTGGAGCTTACATTGCCCGCTACTCCATAACTTGGGAA
NZ_CP035258.1:750874-758722    4909   CAACCAATCAGTCAGACTCCTGTGAAGGATAAAACATTGACAATTCAACACAGTGGAGCTTACATTGCCCGCTACTCCATAACTTGGGAA
NZ_CP035260.1:726156-733994    4899   CAACCAATCAGTCAGACTCCTGTGAAAGATAAAACATTGACAATTCAACACAGTGGAGCTTACATTGCCCGCTACTCCATAACTTGGGAA
NZ_CP035241.1:708296-716144    4909   CAACCAATCAGTCAGACTCCTGTGAAAGATAAAACATTGACAATTCAACACAGTGGAGCTTACATTGCCCGCTACTCCATAACTTGGGAA
NZ_CP035261.1:779086-786938    4909   CAACCAATCAGTCAGACTCCTGTGAAAGATAAAACATTGACAATTCAATACAGTGGAGCTTACATTGCTCGCTACTCCATAACTTGGGAA
NZ_CP035242.1:c1532236-1524385 4908   CAACCAATCAGTCAGACTCCTGTGAAAGATAAAACATTGACAATTCAACACAGTGGAGCTTACATTGCCCGCTACTCCATAACTTGGGAA
NZ_CP035243.1:777729-785580    4908   CAACCAATCAGTCAGACTCCTGTGAAAGATAAAACATTGACAATTCAACACAGTGGAGCTTACATTGCCCGCTACTCCATAACTTGGGAA
NZ_CP035237.1:c1311322-1303456 4923   CAACCAATCAGTCAGACTCCTGTGAAAGATAAAACATTGACAATTCAACACAGTGGAGCTTACATTGCCCGCTACTCCATAACTTGGGAA
NZ_CP035246.1:c1302892-1295026 4923   CAACCAATCAGTCAGACTCCTGTGAAGGATAAAACATTGACAATTCAACACAGTGGAGCTTACATTGCCCGCTACTCCATAACTTGGGAA
NZ_CP035247.1:c1301115-1293263 4909   CAACCAATCAGTCAGACTCCTGTGAAGGATAAAACATTGACAATTCAACACAGTGGAGCTTACATTGCCCGCTACTCCATAACTTGGGAA
NZ_CP035238.1:738547-746413    4923   CAACCAATCAGTCAGACTCCTGTGAAGGATAAAACATTGACAATTCAACACAGTGGAGCTTACATTGCCCGCTACTCCATAACTTGGGAA
NZ_CP035255.1:776866-784718    4909   CAACCAATCAGTCAGACTCCTGTGAAGGATAAAACATTGACAATTCAACACAGTGGAGCTTACATTGCCCGCTACTCCATAACTTGGGAA
NZ_CP035256.1:807236-815102    4923   CAACCAATCAGTCAGACTCCTGTGAAGGATAAAACATTGACAATTCAACACAGTGGAGCTTACATTGCCCGCTACTCCATAACTTGGGAA
NZ_CP038252.1:1151812-1159673  4852   CAACCAATCAGTCAGACTCCTGTGAAAGATAAAACATTGACAATTCAACATAGTGGAGCTTACATTGCCCGCTACTCCATAACTTGGGAA
NZ_CP035254.1:793365-801213    4909   CAACCAATCAGTCAGACTCCTGTGAAAGATAAAACATTGACAATTCAACACAGTGGAGCTTACATTGCCCGCTACTCCATAACTTGGGAA
NZ_CP035251.1:748733-756585    4913   CAACCAATCAGTCAGACTCCTGTGAAAGATAAAACATTGACAATTCAACACAGTGGAGCTTACATTGCCCGCTACTCCATAACTTGGGAA
NZ_CP035252.1:745670-753536    4927   CAACCAATCAGTCAGACTCCTGTGAAAGATAAAACATTGACAATTCAACACAGTGGAGCTTACATTGCCCGCTACTCCATAACTTGGGAA
NZ_CP035257.1:776317-784169    4913   CAACCAATCAGTCAGACTCCTGTGAAAGATAAAACATTGACAATTCAACACAGTGGAGCTTACATTGCCCGCTACTCCATAACTTGGGAA
NZ_CP038253.1:758105-765957    4913   CAACCAATCAGTCAGACTCCTGTGAAAGATAAAACATTGACAATTCAACACAGTGGAGCTTACATTGCCCGCTACTCCATAACTTGGGAA
NZ_CP035263.1:743050-750901    4908   CAACCAATCAGTCAGACTCCTGTGAAAGATAAAACATTGACAATTCAACACAGTGGAGCTTACATTGCCCGCTACTCCATAACTTGGGAA
NZ_CP035262.1:741595-749460    4922   CAACCAATCAGTCAGACTCCTGTGAAAGATAAAACATTGACAATTCAACACAGTGGAGCTTACATTGCCCGCTACTCCATAACTTGGGAA
NZ_CP035236.1:764915-772766    4908   CAACCAATCAGTCAGACTCCTGTGAAAGATAAAACATTGACAATTCAACACAGTGGAGCTTACATTGCCCGCTACTCCATAACTTGGGAA
NZ_CP035253.1 767286-775134    4909   CAACCAATCAGTCAGACTCCTGTGAAAGATAAAACATTGACAATTCAACACAGTGGAGCTTACATTGCCCGCTACTCCATAACTTGGGAA
NZ_CP035249.1:763080-770921    4909   CAACCAATCAGTCAGACTCCTGTGAAAGATAAAACATTGACAATTCAACACAGTGGAGCTTACATTGCCCGCTACTCCATAACTTGGGAA

NC_003028.3:777000-787000      5728   GAAGTTCCAGTAGATAAAGATGGAAACCAAGTTGTTCGTAGTCATTCTTGGGAAGGAAGCGGTCGCAACCAGACTGCAGGTTTTGTCCTC
NZ_CP035248.1:765099-775111    5737   GAAGTTCCAGTAGATAAAGATGGAAACCAAGTTGTTCGTAGTCATTCTTGGGAAGGAAACGGTTGCAACCAGACTGCAGGTTTTGTCCTC
NZ_CP035235.1 c1326834-1317894 5012   GAAGTTCCAGTAGATAAAGATGGAAACCAAGTTGTTCGTAGTCATTCTTGGGAAGGAAACGGTTGCAACCAGACTGCAGGTTTTGTCCTC
NZ_CP035234.1:756284-765224    5012   GAAGTTCCAGTAGATAAAGATGGAAACCAAGTTGTTCGTAGTCATTCTTGGGAAGGAAACGGTTGCAACCAGACTGCAGGTTTTGTCCTC
NZ_CP038251.1:750745-759686    5012   GAAGTTCCAGTAGATAAAGATGGAAACCAAGTTGTTCGTAGTCATTCTTGGGAAGGAAACGGTTGCAACCAGACTGCAGGTTTTGTCCTC
NZ_CP035244.1:747023-755950    4998   GAAGTTCCAGTAGATAAAGATGGAAACCAAGTTGTTCGTAGTCATTCTTGGGAAGGAAACGGTTGCAACCAGACTGCAGGTTTTGTCCTC
NZ_CP035264.1:800836-809336    5647   GAAGTTCCAGTAGATAAAGATGGAAACCAAGTTGTTCGTAGTCATTCTTGGGAAGGAAACGGTTGCAACCAGACTGCAGGTTTTGTCCTC
NZ_CP035265.1:766128-774627    5646   GAAGTTCCAGTAGATAAAGATGGAAACCAAGTTGTTCGTAGTCATTCTTGGGAAGGAAACGGTTGCAACCAGACTGCAGGTTTTGTCCTC
NZ_CP035245.1 745261-753757    5647   GAAGTTCCAGTAGATAAAGATGGAAACCAAGTTGTTCGTAGTCATTCTTGGGAAGGAAACGGTCGCAACCAGACTGCAGGTTTTGTCCTC
NZ_CP035259.1:745700-753548    4999   GAAGTTCCAGTAGATAAAGATGGAAACCAAGTTGTTCGTAGTCATTCTTGGGAAGGAAGCGGTCGCAACCAGACTGCAGGTTTTGTCCTC
NZ_CP035258.1:750874-758722    4999   GAAGTTCCAGTAGATAAAGATGGAAACCAAGTTGTTCGTAGTCATTCTTGGGAAGGAAGCGGTCGCAACCAGACTGCAGGTTTTGTCCTC
NZ_CP035260.1:726156-733994    4989   GAAGTTACAGTAGATAAAGATGGAAACCAAGTTGTTCGTAGTCATTCTTGGGAAGGAAACGGTCGCAACCAGACTGCAGGTTTTGTCCTC
NZ_CP035241.1:708296-716144    4999   GAAGTTACAGTAGATAAAGATGGAAACCAAGTTGTTCGTAGTCATTCTTGGGAAGGAAACGGTCGCAACCAGACTGCAGGTTTTGTCCTC
NZ_CP035261.1:779086-786938    4999   GAAGTTCCAGTAGATAAAGATGGAAACCAAGTTGTTCGTAGTCATTCTTGGGAAGGAAACGGTTGCAACCAGACTGCAGGTTTTGTCCTC
NZ_CP035242.1:c1532236-1524385 4998   GAAGTTCCAGTAGATAAAGATGGAAACCAAGTTGTTCGTAGTCATTCTTGGGAAGGAAACGGTTGCAACCAGACTGCAGATTTTGTCCTC
NZ_CP035243.1:777729-785580    4998   GAAGTTCCAGTAGATAAAGATGGAAACCAAGTTGTTCGTAGTCATTCTTGGGAAGGAAACGGTTGCAACCAGACTGCAGATTTTGTCCTC
NZ_CP035237.1:c1311322-1303456 5013   GAAGTTCCAGTAGATAAAGATGGAAACCAAGTTGTTCGTAGTCATTCTTGGGAAGGAAACGGTTGCAACCAGACTGCAGGTTTTGTCCTC
NZ_CP035246.1:c1302892-1295026 5013   GAAGTTACAGTAGATAAAGATGGAAACCAAGTTGTTCGTAGTCATTCTTGGGAAGGAAACGGTCGCAACCAGACTGCAGGTTTTGTCCTC
NZ_CP035247.1:c1301115-1293263 4999   GAAGTTACAGTAGATAAAGATGGAAACCAAGTTGTTCGTAGTCATTCTTGGGAAGGAAACGGTCGCAACCAGACTGCAGGTTTTGTCCTC
NZ_CP035238.1:738547-746413    5013   GAAGTTACAGTAGATAAAGATGGAAACCAAGTTGTTCGTAGTCATTCTTGGGAAGGAAACGGTCGCAACCAGACTGCAGGTTTTGTCCTC
NZ_CP035255.1:776866-784718    4999   GAAGTTACAGTAGATAAAGATGGAAACCAAGTTGTTCGTAGTCATTCTTGGGAAGGAAACGGTCGCAACCAGACTGCAGGTTTTGTCCTC
NZ_CP035256.1:807236-815102    5013   GAAGTTACAGTAGATAAAGATGGAAACCAAGTTGTTCGTAGTCATTCTTGGGAAGGAAACGGTCGCAACCAGACTGCAGGTTTTGTCCTC
NZ_CP038252.1:1151812-1159673  4942   GAAGTTCCAGTAGATAAAGATGGAAACCAAGTTGTTCGTAGTCATTCTTGGGAAGGAAACGGTTGCAACCAGACTGCAGGTTTTGTCCTC
NZ_CP035254.1:793365-801213    4999   GAAGTTCCAGTAGATAAAGATGGAAACCAAGTTGTTCGTAGTCATTCTTGGGAAGGAAACGGTTGCAACCAGACTGCAGGTTTTGTCCTC
NZ_CP035251.1:748733-756585    5003   GAAGTTACAGTAGATAAAGATGGAAACCAAGTTGTTCGTAGTCATTCTTGGGAAGGAAACGGTCGCAACCAGACTGCAGGTTTTGTCCTC
NZ_CP035252.1:745670-753536    5017   GAAGTTACAGTAGATAAAGATGGAAACCAAGTTGTTCGTAGTCATTCTTGGGAAGGAAACGGTCGCAACCAGACTGCAGGTTTTGTCCTC
NZ_CP035257.1:776317-784169    5003   GAAGTTACAGTAGATAAAGATGGAAACCAAGTTGTTCGTAGTCATTCTTGGGAAGGAAACGGTCGCAACCAGACTGCAGGTTTTGTCCTC
NZ_CP038253.1:758105-765957    5003   GAAGTTACAGTAGATAAAGATGGAAACCAAGTTGTTCGTAGTCATTCTTGGGAAGGAAACGGTCGCAACCAGACTGCAGGTTTTGTCCTC
NZ_CP035263.1:743050-750901    4998   GAAGTTACAGTAGATAAAGATGGAAACCAAGTTGTTCTTAGTCATTCTTGGGAAGGAAACGGTCGCAACCAGACTGCAGGTTTTGTCCTC
NZ_CP035262.1:741595-749460    5012   GAAGTTACAGTAGATAAAGATGGAAACCAAGTTGTTCTTAGTCATTCTTGGGAAGGAAACGGTCGCAACCAGACTGCAGGTTTTGTCCTC
NZ_CP035236.1:764915-772766    4998   GAAGTTACAGTAGATAAAGATGGAAACCAAGTTGTTCGTAGTCATTCTTGGGAAGGAAACGGTCGCAACCAGACTGCAGGTTTTGTCCTC
NZ_CP035253.1 767286-775134    4999   GAAGTTCCAGTAGATAAAGATGGAAACCAAGTTGTTCGTAGTCATTCTTGGGAAGGAAACGGTTGCAACCAGACTGCAGGTTTTGTCCTC
NZ_CP035249.1:763080-770921    4999   GAAGTTCCAGTAGATAAAGATGGAAACCAAGTTGTTCGTAGTCATTCTTGGGAAGGAAACGGTCGCAACCAGACTGCGGGTTTTGTCCTC

NC_003028.3:777000-787000      5818   AACCTCCCAATCAAAGAAAATATGAGAAATCTGCGAGTTAAGATTGAGAAAAAGACGGGCCTACTATGGAATAGATGGCAAACAATCTAT
NZ_CP035248.1:765099-775111    5827   AACTTCCCAATCAAAGAAAATATGAGAAATCTGCGAGTTAAGATTGAGAAAAAGACGGGCCTACTATGGAATAGATGGCAAACAATCTAT
NZ_CP035235.1 c1326834-1317894 5102   AACCTCCCAATCAAAGAAAATATGAGAAATCTGCGAGTTAAGATTGAGAAAAAGACGGGCCTACTATGGAATAGATGGCAAACAATCTAT
NZ_CP035234.1:756284-765224    5102   AACCTCCCAATCAAAGAAAATATGAGAAATCTGCGAGTTAAGATTGAGAAAAAGACGGGCCTACTATGGAATAGATGGCAAACAATCTAT
NZ_CP038251.1:750745-759686    5102   AACCTCCCAATCAAAGAAAATATGAGAAATCTGCGAGTTAAGATTGAGAAAAAGACGGGCCTACTATGGAATAGATGGCAAACAATCTAT
NZ_CP035244.1:747023-755950    5088   AACCTCCCAATCAAAGAAAATATGAGAAATCTGCGAGTTAAGATTGAGAAAAAGACGGGCCTACTATGGAATAGATGGCAAACAATCTAT
NZ_CP035264.1:800836-809336    5737   AACCTCCCAATCAAAGAAAATATGAGAAATCTGCGAGTTAAGATTGAGAAAAAGACGGGCCTACTATGGAATAGATGGCAAACAATCTAT
NZ_CP035265.1:766128-774627    5736   AACCTCCCAATCAAAGAAAATATGAGAAATCTGCGAGTTAAGATTGAGAAAAAGACGGGCCTACTATGGAATAGATGGCAAACAATCTAT
NZ_CP035245.1 745261-753757    5737   AACCTCCCAATCAAAGAAAATATGAGAAATCTGCGAGTTAAGATTGAGAAAAAGACGGGCCTACTATGGAATAGATGGCAAACAATCTAT
NZ_CP035259.1:745700-753548    5089   AACCTCCCAATCAAAGAAAATATGAGAAATCTGCGAGTTAAGATTGAGAAAAAGACGGGCCTACTATGGAATAGATGGCAAACAATCTAT
NZ_CP035258.1:750874-758722    5089   AACCTCCCAATCAAAGAAAATATGAGAAATCTGCGAGTTAAGATTGAGAAAAAGACGGGCCTACTATGGAATAGATGGCAAACAATCTAT
NZ_CP035260.1:726156-733994    5079   AACCTCCCAATCAAAGAAAATATGAGAAATCTGCGAGTTAAGATTGAGAAAAAGACGGGCCTACTATGGAATAGATGGCAAACAATCTAT
NZ_CP035241.1:708296-716144    5089   AACCTCCCAATCAAAGAAAATATGAGAAATCTGCGAGTTAAGATTGAGAAAAAGACGGGCCTACTATGGAATAGATGGCAAACAATCTAT
NZ_CP035261.1:779086-786938    5089   AACCTCCCAATCAAAGAAAATATGAGAAATCTGCGAGTTAAGATTGAGAAAAAGACGGGCCTACTATGGAATAGATGGCAAACAATCTAT
NZ_CP035242.1:c1532236-1524385 5088   AACCTCCCAATCAAAGAAAATATGAGAAATCTGCGAGTTAAGATTGAGAAAAAGACGGGCCTACTATGGAATAGATGGCAAACAATCTAT
NZ_CP035243.1:777729-785580    5088   AACCTCCCAATCAAAGAAAATATGAGAAATCTGCGAGTTAAGATTGAGAAAAAGACGGGCCTACTATGGAATAGATGGCAAACAATCTAT
NZ_CP035237.1:c1311322-1303456 5103   AACCTCCCAATCAAAGAAAATATGAGAAATCTGCGAGTTAAGATTGAGAAAAAGACGGGCCTACTATGGAATAGATGGCAAACAATCTAT
NZ_CP035246.1:c1302892-1295026 5103   AACCTCCCAATCAAAGAAAATATGAGAAATCTGCGAGTTAAGATTGAGAAAAAGACGGGCCTACTATGGAATAGATGGCAAACAATCTAT
NZ_CP035247.1:c1301115-1293263 5089   AACCTCCCAATCAAAGAAAATATGAGAAATCTGCGAGTTAAGATTGAGAAAAAGACGGGCCTACTATGGAATAGATGGCAAACAATCTAT
NZ_CP035238.1:738547-746413    5103   AACCTCCCAATCAAAGAAAATATGAGAAATCTGCGAGTTAAGATTGAGAAAAAGACGGGCCTACTATGGAATAGATGGCAAACAATCTAT
NZ_CP035255.1:776866-784718    5089   AACCTCCCAATCAAAGAAAATATGAGAAATCTGCGAGTTAAGATTGAGAAAAAGACGGGCCTACTATGGAATAGATGGCAAACAATCTAT
NZ_CP035256.1:807236-815102    5103   AACCTCCCAATCAAAGAAAATATGAGAAATCTGCGAGTTAAGATTGAGAAAAAGACGGGCCTACTATGGAATAGATGGCAAACAATCTAT
```

```
NZ_CP038252.1:1151812-1159673  5032   AACCTCCCAATCAAAGAAAATATGAGAAATCTGCGAGTTAAGATTGAGAAAAAGACGGGCCTACTATGGAATAGATGGCAAACAATCTAT
NZ_CP035254.1:793365-801213    5089   AACCTCCCAATCAAAGAAAATATGAGAAATCTGCGAGTTAAGATTGAGAAAAAGACGGGCCTACTATGGAATAGATGGCAAACAATCTAT
NZ_CP035251.1:748733-756585    5093   AACCTCCCAATCAAAGAAAATATGAGAAATCTGCGAGTTAAGATTGAGAAAAAGACGGGCCTACTATGGAATAGATGGCAAACAATCTAT
NZ_CP035252.1:745670-753536    5107   AACCTCCCAATCAAAGAAAATATGAGAAATCTGCGAGTTAAGATTGAGAAAAAGACGGGCCTACTATGGAATAGATGGCAAACAATCTAT
NZ_CP035257.1:776317-784169    5093   AACCTCCCAATCAAAGAAAATATGAGAAATCTGCGAGTTAAGATTGAGAAAAAGACGGGCCTACTATGGAATAGATGGCAAACAATCTAT
NZ_CP038253.1:758105-765957    5093   AACCTCCCAATCAAAGAAAATATGAGAAATCTGCGAGTTAAGATTGAGAAAAAGACGGGCCTACTATGGAATAGATGGCAAACAATCTAT
NZ_CP035263.1:743050-750901    5088   AACCTCCCAATCAAAGAAAATATGAGAAATCTGCGAGTTAAGATTGAGAAAAAGACGGGCCTACTATGGAATAGATGGCAAACAATCTAT
NZ_CP035262.1:741595-749460    5102   AACCTCCCAATCAAAGAAAATATGAGAAATCTGCGAGTTAAGATTGAGAAAAAGACGGGCCTACTATGGAATAGATGGCAAACAATCTAT
NZ_CP035236.1:764915-772766    5088   AACCTCCCAATCAAAGAAAATATGAGAAATCTGCGAGTTAAGATTGAGAAAAAGACGGGCCTACTATGGAATAGATGGCAAACAATCTAT
NZ_CP035253.1 767286-775134    5089   AACCTCCCAATCAAAGAAAATATGAGAAATCTGCGAGTTAAGATTGAGAAAAAGACGGGCCTACTATGGAATAGATGGCAAACAATCTAT
NZ_CP035249.1:763080-770921    5089   AACCTCCCAATCAAAGAAAATATGAGAAATCTGCGAGTTAAGATTGAGAAAAAGACGGGCCTACTATGGAATAGATGGCAAACAATCTAT

NC_003028.3:777000-787000      5908   GAAAACAGACCAATTTTAGCTCAACCCCACCGTAAAATTACCCATTGGGGTACGACATTGAATTCCAAGGTGAGTGACGATGATGTCTTG
NZ_CP035248.1:765099-775111    5917   GAAAACAGACCAATTTTAGCTCAACCCCACCGTAAAATTACCCATTGGGGTACGACATTGAATTCCAAGGTGAGTGACGATGATGTCTTG
NZ_CP035235.1 c1326834-1317894 5192   GAAAACAGACCAATTTTAGCTCAACCCCACCGTAAAATTACCCATTGGGGTACGACATTGAATTCCAAGGTGAGTGACGATGATGTCTTG
NZ_CP035234.1:756284-765224    5192   GAAAACAGACCAATTTTAGCTCAACCCCACCGTAAAATTACCCATTGGGGTACGACATTGAATTCCAAGGTGAGTGACGATGATGTCTTG
NZ_CP038251.1:750745-759686    5192   GAAAACAGACCAATTTTAGCTCAACCCCACCGTAAAATTACCCATTGGGGTACGACATTGAATTCCAAGGTGAGTGACGATGATGTCTTG
NZ_CP035244.1:747023-755950    5178   GAAAACAGACCAATTTTAGCTCAACCCCACCGTAAAATTACCCATTGGGGTACGACATTGAATTCCAAGGTGAGTGACGATGATGTCTTG
NZ_CP035264.1:800836-809336    5827   GAAAACAGACCAATTTTAGCTCAACCCCACCGTAAAATTACCCATTGGGGTACGACATTGAATTCCAAGGTGAGTGACGATGATGTCTTG
NZ_CP035265.1:766128-774627    5826   GAAAACAGACCAATTTTAGCTCAACCCCACCGTAAAATTACCCATTGGGGTACGACATTGAATTCCAAGGTGAGTGACGATGATGTCTTG
NZ_CP035245.1 745261-753757    5827   GAAAACAGACCAATTTTAGCTCAACCCCACCGTAAAATTACCCATTGGGGTACGACATTGAATTCCAAGGTGAGTGACGATGATGTCTTG
NZ_CP035259.1:745700-753548    5179   GAAAACAGACCAATTTTAGCTCAACCCCACCGTAAAATTACCCATTGGGGTACGACATTGAATTCCAAGGTGAGTGACGATGATGTCTTG
NZ_CP035258.1:750874-758722    5179   GAAAACAGACCAATTTTAGCTCAACCCCACCGTAAAATTACCCATTGGGGTACGACATTGAATTCCAAGGTGAGTGACGATGATGTCTTG
NZ_CP035260.1:726156-733994    5169   GAAAACAGACCAATTTTAGCTCAACCCCACCGTAAAATTACCCATTGGGGTACGACATTGAATTCCAAGGTGAGTGACGATGATGTCTTG
NZ_CP035241.1:708296-716144    5179   GAAAACAGACCAATTTTAGCTCAACCCCACCGTAAAATTACCCATTGGGGTACGACATTGAATTCCAAGGTGAGTGACGATGATGTCTTG
NZ_CP035261.1:779086-786938    5179   GAAAACAGACCAATTTTAGCTCAACCCCACCGTAAAATTACCCATTGGGGTACGACATTGAATTCCAAGGTGAGTGACGATGATGTCTTG
NZ_CP035242.1:c1532236-1524385 5178   GAAAACAGACCAATTTTAGCTCAACCCCACCGTAAAATTACCCATTGGGGTACGACATTGAATTCCAAGGTGAGTGACGATGATGTCTTG
NZ_CP035243.1:777729-785580    5178   GAAAACAGACCAATTTTAGCTCAACCCCACCGTAAAATTACCCATTGGGGTACGACATTGAATTCCAAGGTGAGTGACGATGATGTCTTG
NZ_CP035237.1:c1311322-1303456 5193   GAAAACAGACCAATTTTAGCTCAACCCCACCGTAAAATTACCCATTGGGGTACGACATTGAATTCCAAGGTGAGTGACGATGATGTCTTG
NZ_CP035246.1:c1302892-1295026 5193   GAAAACAGACCAATTTTAGCTCAACCCCACCGTAAAATTACCCATTGGGGTACGACATTGAATTCCAAGGTGAGTGACGATGATGTCTTG
NZ_CP035247.1:c1301115-1293263 5179   GAAAACAGACCAATTTTAGCTCAACCCCACCGTAAAATTACCCATTGGGGTACGACATTGAATTCCAAGGTGAGTGACGATGATGTCTTG
NZ_CP035238.1:738547-746413    5193   GAAAACAGACCAATTTTAGCTCAACCCCACCGTAAAATTACCCATTGGGGTACGACATTGAATTCCAAGGTGAGTGACGATGATGTCTTG
NZ_CP035255.1:776866-784718    5179   GAAAACAGACCAATTTTAGCTCAACCCCACCGTAAAATTACCCATTGGGGTACGACATTGAATTCCAAGGTGAGTGACGATGATGTCTTG
NZ_CP035256.1:807236-815102    5193   GAAAACAGACCAATTTTAGCTCAACCCCACCGTAAAATTACCCATTGGGGTACGACATTGAATTCCAAGGTGAGTGACGATGATGTCTTG
NZ_CP038252.1:1151812-1159673  5122   GAAAACAGACCAATTTTAGCTCAACCCCACCGTAAAATTACCCATTGGGGTACGACATTGAATTCCAAGGTGAGTGACGATGATGTCTTG
NZ_CP035254.1:793365-801213    5179   GAAAACAGACCAATTTTAGCTCAACCCCACCGTAAAATTACCCATTGGGGTACGACATTGAATTCCAAGGTGAGTGACGATGATGTCTTG
NZ_CP035251.1:748733-756585    5183   GAAAACAGACCAATTTTAGCTCAACCCCACCGTAAAATTACCCATTGGGGTACGACATTGAATTCCAAGGTGAGTGACGATGATGTCTTG
NZ_CP035252.1:745670-753536    5197   GAAAACAGACCAATTTTAGCTCAACCCCACCGTAAAATTACCCATTGGGGTACGACATTGAATTCCAAGGTGAGTGACGATGATGTCTTG
NZ_CP035257.1:776317-784169    5183   GAAAACAGACCAATTTTAGCTCAACCCCACCGTAAAATTACCCATTGGGGTACGACATTGAATTCCAAGGTGAGTGACGATGATGTCTTG
NZ_CP038253.1:758105-765957    5183   GAAAACAGACCAATTTTAGCTCAACCCCACCGTAAAATTACCCATTGGGGTACGACATTGAATTCCAAGGTGAGTGACGATGATGTCTTG
NZ_CP035263.1:743050-750901    5178   GAAAACAGACCAATTTTAGCTCAACCCCACCGTAAAATTACCCATTGGGGTACGACATTGAATTCCAAGGTGAGTGACGATGATGTCTTG
NZ_CP035262.1:741595-749460    5192   GAAAACAGACCAATTTTAGCTCAACCCCACCGTAAAATTACCCATTGGGGTACGACATTGAATTCCAAGGTGAGTGACGATGATGTCTTG
NZ_CP035236.1:764915-772766    5178   GAAAACAGACCAATTTTAGCTCAACCCCACCGTAAAATTACCCATTGGGGTACGACATTGAATTCCAAGGTGAGTGACGATGATGTCTTG
NZ_CP035253.1 767286-775134    5179   GAAAACAGACCAATTTTAGCTCAACCCCACCGTAAAATTACCCATTGGGGTACGACATTGAATTCCAAGGTGAGTGACGATGATGTCTTG
NZ_CP035249.1:763080-770921    5179   GAAAACAGACCAATTTCAGCTCAACCCCACCGTAAAATTACCCATTGGGGTACGACATTGAATTCCAAGGTGAGTGACGATGATGTCTTG

NC_003028.3:777000-787000      5998   TAATCTGATGGTAGAATGACAGTTAGTTTGTCTAGTTTATAAGAAAGTACTACCTGAGCTTGAATAGGACTCAGGTAGCTCTCTATGAAA
NZ_CP035248.1:765099-775111    6007   TAATCTGATGGTAGAATGACAGTTAGTTTGTCTAGTTTATAAGAAAGTACTATCTGAGCTTGAATAGAACTCAGGTAGCTCTCTGTGAAA
NZ_CP035235.1 c1326834-1317894 5282   TAATCTGATGGTACAATGACAGTTAGTTTGTCTAGTTTATAAGAAAGTACTACCTGAGCTTGAATAGAACTCAGGTAGCTCTCTGTGAAA
NZ_CP035234.1:756284-765224    5282   TAATCTGATGGTACAATGACAGTTAGTTTGTCTAGTTTATAAGAAAGTACTACCTGAGCTTGAATAGAACTCAGGTAGCTCTCTGTGAAA
NZ_CP038251.1:750745-759686    5282   TAATCTGATGGTACAATGACAGTTAGTTTGTCTAGTTTATAAGAAAGTACTACCTGAGCTTGAATAGAACTCAGGTAGCTCTCTGTGAAA
NZ_CP035244.1:747023-755950    5268   TAATCTGATGGTACAATGACAGTTAGTTTGTCTAGTTTATAAGAAAGTACTACCTGAGCTTGAATAGAACTCAGGTAGCTCTCTGTGAAA
NZ_CP035264.1:800836-809336    5917   TAATCTGATGGTAGAATGACAGTTAGTTTGTCTAGTTTATAAGAAAGTACTACCTGAGCTTGAATAGAACTCAGGTAGCTCTCTGTGAAA
NZ_CP035265.1:766128-774627    5916   TAATCTGATGGTAGAATGACAGTTAGTTTGTCTAGTTTATAAGAAAGTACTACCTGAGCTTGAATAGAACTCAGGTAGCTCTCTGTGAAA
NZ_CP035245.1 745261-753757    5917   TAATCTGATGGTAGAATGACAGTTAGTTTGTCTAGTTTATAAGAAAGTACTACCTGAGCTTGAATAGAACTCAGGTAGCTCTCTATGAAA
NZ_CP035259.1:745700-753548    5269   TAATCTGATGGTAGAATGACAGTTAGTTTGTCTAGTTTATAAGAAAGTACTACCTGAGCTTGAATAGGACTCAGGTAGCTCTCTATGAAA
NZ_CP035258.1:750874-758722    5269   TAATCTGATGGTAGAATGACAGTTAGTTTGTCTAGTTTATAAGAAAGTACTACCTGAGCTTGAATAGGACTCAGGTAGCTCTCTATGAAA
NZ_CP035260.1:726156-733994    5259   TAATCTGATGGTACAATGACAGTTAGTTTGTCTAGTTTATAAGAAAGTACTACCTGAGCTTGAATAGAACTCAGGTAGCTCTCTGTGAAA
NZ_CP035241.1:708296-716144    5269   TAATCTGATGGTACAATGACAGTTAGTTTGTCTAGTTTATAAGAAAGTACTACCTGAGCTTGAATAGAACTCAGGTAGCTCTCTGTGAAA
NZ_CP035261.1:779086-786938    5269   TAATCTGATGGTACAATGACAGTTAGTTTGTCTAGTTTATAAGAAAGTACTACCTGAGCTTGAATAGGACTCAGGTAGTTCTCTGTGAAA
NZ_CP035242.1:c1532236-1524385 5268   TAATCTGATGGTACAATGACAGTTAGTTTGTCTAGTTTATAAGAAAGTACTACCTGAGCTTGAATAGAACTCAGGTAGCTCTCTGTGAAA
NZ_CP035243.1:777729-785580    5268   TAATCTGATGGTACAATGACAGTTAGTTTGTCTAGTTTATAAGAAAGTACTACCTGAGCTTGAATAGAACTCAGGTAGCTCTCTGTGAAA
NZ_CP035237.1:c1311322-1303456 5283   TAATCTGATGGTAGAATGACAGTTAGTTTGTCTAGTTTATAAGAAAGTACTACCTGAGCTTGAATAGAACTCAGGTAGCTCTCTATGAAA
NZ_CP035246.1:c1302892-1295026 5283   TAATCTGATGGTACAATGATAGTTAGTTTGTCTAGTTTATAAGAAAGTACTACCTGAGCTTGAATAGGACTCAGGTAGCTCTCTGTGAAA
NZ_CP035247.1:c1301115-1293263 5269   TAATCTGATGGTACAATGATAGTTAGTTTGTCTAGTTTATAAGAAAGTACTACCTGAGCTTGAATAGGACTCAGGTAGCTCTCTGTGAAA
NZ_CP035238.1:738547-746413    5283   TAATCTGATGGTACAATGATAGTTAGTTTGTCTAGTTTATAAGAAAGTACTACCTGAGCTTGAATAGGACTCAGGTAGCTCTCTGTGAAA
NZ_CP035255.1:776866-784718    5269   TAATCTGATGGTACAATGATAGTTAGTTTGTCTAGTTTATAAGAAAGTACTACCTGAGCTTGAATAGGACTCAGGTAGCTCTCTGTGAAA
NZ_CP035256.1:807236-815102    5283   TAATCTGATGGTACAATGATAGTTAGTTTGTCTAGTTTATAAGAAAGTACTACCTGAGCTTGAATAGGACTCAGGTAGCTCTCTGTGAAA
NZ_CP038252.1:1151812-1159673  5212   TAATCTGATGGTACAATGACAGTTAGTTTGTCTAGTTTATAAGAAAGTACTACCTGAGCTTGAATAGAACTCAGGTAGCTCTCTGTGAAA
NZ_CP035254.1:793365-801213    5269   TAATCTGATGGTACAATGACAGTTAGTTTGTCTAGTTTATAAGAAAGTACTACCTGAGCTTGAATAGAACTCAGGTAGCTCTCTATGAAA
NZ_CP035251.1:748733-756585    5273   TAATCTGATGGTAGAATGACAGTTAGTTTGTCTAGTTTATAAGAAAGTACTACCTGAGCTTGAATAGAACTCAGGTAGCTCTCTATGAAA
NZ_CP035252.1:745670-753536    5287   TAATCTGATGGTAGAATGACAGTTAGTTTGTCTAGTTTATAAGAAAGTACTACCTGAGCTTGAATAGAACTCAGGTAGCTCTCTATGAAA
NZ_CP035257.1:776317-784169    5273   TAATCTGATGGTAGAATGACAGTTAGTTTGTCTAGTTTATAAGAAAGTACTACCTGAGCTTGAATAGAACTCAGGTAGCTCTCTATGAAA
NZ_CP038253.1:758105-765957    5273   TAATCTGATGGTAGAATGACAGTTAGTTTGTCTAGTTTATAAGAAAGTACTACCTGAGCTTGAATAGAACTCAGGTAGCTCTCTATGAAA
NZ_CP035263.1:743050-750901    5268   TAATCTGATGGTACAATGACAGTTAGTTTGTCTAGTTTATAAGAAAGTACTACCTGAGCTTGAATAGAACTCAGGTAGCTCTCTGTGAAA
NZ_CP035262.1:741595-749460    5282   TAATCTGATGGTACAATGACAGTTAGTTTGTCTAGTTTATAAGAAAGTACTACCTGAGCTTGAATAGAACTCAGGTAGCTCTCTGTGAAA
NZ_CP035236.1:764915-772766    5268   TAATCTGATGGTACAATGACAGTTAGTTTGTCTAGTTTATAAGAAAGTACTACCTGAGCTTGAATAGAACTCAGGTAGCTCTCTGTGAAA
NZ_CP035253.1 767286-775134    5269   TAATCTGATGGTACAATGACAGTTAGTTTGTCTAGTTTATAAGAAAGTACTACCTGAGCTTGAATAGAACTCAGGTAGCTCTCTATGAAA
NZ_CP035249.1:763080-770921    5269   TAATCTGATGGTACAATGACAGTTAGTTTGTCTAGTTTATAAGAAAGTACTACCTGAGCTTGAATAGAACTCAGGTAGCTCTCTGTGAAA

NC_003028.3:777000-787000      6088   GAACAAAATTAATACTCAATGAAAATCAAAGAGCAAACTAAGAAACTAGCCGCAGGTTGCTCAAAGCACTGCTTTGAGGTTGTAGATAAG
NZ_CP035248.1:765099-775111    6097   GAACAAAATTAATACTCAATGAAAATCAAAGAGCAAACTAGGAAACTAGCCGCAGGTTGCTCAAAGCACTGCTTTGAGGTTGTAGATAAG
NZ_CP035235.1 c1326834-1317894 5372   GAACAAAATTAATACTCAATGAAAATCAAAGAGCAAACTAGGAAACTAGCCGCAGGTTGCTCAAAGCACTGCTTTGAGGTTGTAGATAAG
NZ_CP035234.1:756284-765224    5372   GAACAAAATTAATACTCAATGAAAATCAAAGAGCAAACTAGGAAACTAGCCGCAGGTTGCTCAAAGCACTGCTTTGAGGTTGTAGATAAG
NZ_CP038251.1:750745-759686    5372   GAACAAAATTAATACTCAATGAAAATCAAAGAGCAAACTAGGAAACTAGCCGCAGGTTGCTCAAAGCACTGCTTTGAGGTTGTAGATAAG
NZ_CP035244.1:747023-755950    5358   GAACAAAATTAATACTCAATGAAAATCAAAGAGCAAACTAGGAAACTAGCCGCAGGTTGCTCAAAGCACTGCTTTGAGGTTGTAGATAAG
NZ_CP035264.1:800836-809336    6007   GAACAAAATTAATACTCAATGAAAATCAAAGAGCAAACTAGGAAACTAGCCGCAGGTTGCTCAAAGCACTGCTTTGAGGTTGTAGATAAG
NZ_CP035265.1:766128-774627    6006   GAACAAAATTAATACTCAATGAAAATCAAAGAGCAAACTAGGAAACTAGCCGCAGGTTGCTCAAAGCACTGCTTTGAGGTTGTAGATAAG
NZ_CP035245.1 745261-753757    6007   GAACAAAATTAATACTCAATGAAAATCAAAGAGCAAACTAGGAAACTAGCCGCAGGTTGCTCAAAGCACTGCTTTGAGGTTGTAGATAAG
NZ_CP035259.1:745700-753548    5359   GAACAAAATTAATACTCAATGAAAATCAAAGAGCAAACTAAGAAACTAGCCGCAGGTTGCTCAAAGCACTGCTTTGAGGTTGTAGATAAG
NZ_CP035258.1:750874-758722    5359   GAACAAAATTAATACTCAATGAAAATCAAAGAGCAAACTAAGAAACTAGCCGCAGGTTGCTCAAAGCACTGCTTTGAGGTTGTAGATAAG
NZ_CP035260.1:726156-733994    5349   GAACAAAATTAATACTCAATGAAAATCAAAGAGCAAACTAAGAAACTAGCCGCAGGTTGCTCAAAGCACTGCTTTGAGGTTGTAGATAAG
NZ_CP035241.1:708296-716144    5359   GAACAAAATTAATACTCAATGAAAATCAAAGAGCAAACTAAGAAACTAGCCGCAGGTTGCTCAAAGCACTGCTTTGAGGTTGTAGATAAG
NZ_CP035261.1:779086-786938    5359   GAACAAAATTAATACTCAATGAAAATCAAAGAGCAAACTAGGAAACTAGCCGCAGGTTGCTCAAAGCACTGCTTTGAGGTTGTAGATAAG
NZ_CP035242.1:c1532236-1524385 5358   GAACAAAATTAATACTCAATGAAAATCAAAGAGCAAACTAGGAAACTAGCCGCAGGTTGCTCAAAGCACTGCTTTGAGGTTGTAGATAAG
```

```
NZ_CP035243.1:777729-785580    5358   GAACAAAATTAATACTCAATGAAAATCAAAGAGCAAACTAGGAAACTAGCCGCAGGTTGCTCAAAGCACTGCTTTGAGGTTGTAGATAAG
NZ_CP035237.1:c1311322-1303456 5373   GAACAAAATTAATACTCAATGAAAATCAAAGAGCAAACTAGGAAACTAGCCGCAGGTTGCTCAAAGCACTGCTTTGAGGTTGTAGATAAG
NZ_CP035246.1:c1302892-1295026 5373   GAACAAAATTAATACTCAATGAAAATCAAAGAGCAAACTAGGAAACTAGCCGCAGGTTGCTCAAAGCACTGCTTTGAGGTTGTAGATAAG
NZ_CP035247.1:c1301115-1293263 5359   GAACAAAATTAATACTCAATGAAAATCAAAGAGCAAACTAGGAAACTAGCCGCAGGTTGCTCAAAGCACTGCTTTGAGGTTGTAGATAAG
NZ_CP035238.1:738547-746413    5373   GAACAAAATTAATACTCAATGAAAATCAAAGAGCAAACTAGGAAACTAGCCGCAGGTTGCTCAAAGCACTGCTTTGAGGTTGTAGATAAG
NZ_CP035255.1:776866-784718    5359   GAACAAAATTAATACTCAATGAAAATCAAAGAGCAAACTAGGAAACTAGCCGCAGGTTGCTCAAAGCACTGCTTTGAGGTTGTAGATAAG
NZ_CP035256.1:807236-815102    5373   GAACAAAATTAATACTCAATGAAAATCAAAGAGCAAACTAGGAAACTAGCCGCAGGTTGCTCAAAGCACTGCTTTGAGGTTGTAGATAAG
NZ_CP038252.1:1151812-1159673  5302   GAACAAAATTAATACTCAATGAAAATCAAAGAGCAAACTAGGAAACTAGCCGCAGGTTGCTCAAAGCACTGCTTTGAGGTTGTAGATAAG
NZ_CP035254.1:793365-801213    5359   GAACAAAATTAATACTCAATGAAAATCAAAGAGCAAACTAAGAAACTAGCCGCAGGTTGCTCAAAGCACTGCTTTGAGGTTGTAGATAAG
NZ_CP035251.1:748733-756585    5363   GAACAAAATTAATACTCAATGAAAATCAAAGAGCAAACTAGGAAACTAGCCGCAGGTTGCTCAAAGCACTACTTTGAGGTTGTAGATAAG
NZ_CP035252.1:745670-753536    5377   GAACAAAATTAATACTCAATGAAAATCAAAGAGCAAACTAGGAAACTAGCCGCAGGTTGCTCAAAGCACTACTTTGAGGTTGTAGATAAG
NZ_CP035257.1:776317-784169    5363   GAACAAAATTAATACTCAATGAAAATCAAAGAGCAAACTAGGAAACTAGCCGCAGGTTGCTCAAAGCACTACTTTGAGGTTGTAGATAAG
NZ_CP038253.1:758105-765957    5363   GAACAAAATTAATACTCAATGAAAATCAAAGAGCAAACTAGGAAACTAGCCGCAGGTTGCTCAAAGCACTACTTTGAGGTTGTAGATAAG
NZ_CP035263.1:743050-750901    5358   GAACAAAATTAATACTCAATGAAAATCAAAGAGCAAACTAAGAAACTAGCCGCAGGTTGCTCAAAGCACTGCTTTGAGGTTGTAGATAAG
NZ_CP035262.1:741595-749460    5372   GAACAAAATTAATACTCAATGAAAATCAAAGAGCAAACTAAGAAACTAGCCGCAGGTTGCTCAAAGCACTGCTTTGAGGTTGTAGATAAG
NZ_CP035236.1:764915-772766    5358   GAACAAAATTAATACTCAATGAAAATCAAAGAGCAAACTAAGAAACTAGCCGCAGGTTGCTCAAAGCACTGCTTTGAGGTTGTAGATAAG
NZ_CP035253.1 767286-775134    5359   GAACAAAATTAATACTCAATGAAAATCAAAGAGCAAACTAAGAAACTAGCCGCAGGTTGCTCAAAGCACTGCTTTGAGGTTGTAGATAAG
NZ_CP035249.1:763080-770921    5359   GAACAAAATTAATACTCAATGAAAATCAAAGAGCAAACTAGGAAACTAGCCGCAGGTTGCTCAAAGCACTGCTTTGAGGTTGTAGATAAG

NC_003028.3:777000-787000      6178   ACTGACGAAGTCAGTCACATATATAATCCAAGGCGACGTTGACGTGGTTTGAAGAGATTTTCGAAGAGTATAAACAGAAAGGTAGAGCGC
NZ_CP035248.1:765099-775111    6187   ACTGACGAAGTCAGTCACATATATAATCCAAGGCGACGTTGACGTGGTTTGAAGAGATTTTCGAAGAGTATAAACAGAAAGGTAGAGCGC
NZ_CP035235.1 c1326834-1317894 5462   ACTGACGAAGTCAGTCACATATATAATCCAAGGCGACGTTGACGTGGTTTGAAGAGATTTTCGAAGAGTATAAACAGAAAGGTAGAGCGC
NZ_CP035234.1:756284-765224    5462   ACTGACGAAGTCAGTCACATATATAATCCAAGGCGACGTTGACGTGGTTTGAAGAGATTTTCGAAGAGTATAAACAGAAAGGTAGAGCGC
NZ_CP038251.1:750745-759686    5462   ACTGACGAAGTCAGTCACATATATAATCCAAGGCGACGTTGACGTGGTTTGAAGAGATTTTCGAAGAGTATAAACAGAAAGGTAGAGCGC
NZ_CP035244.1:747023-755950    5448   ACTGACGAAGTCAGTCACATATATAATCCAAGGCGACGTTGACGTGGTTTGAAGAGATTTTCGAAGAGTATAAACAGAAAGGTAGAGCGC
NZ_CP035264.1:800836-809336    6097   ACTGACGAAGTCAGTCACATATATAATCCAAGGCGACGTTGACGTGGTTTGAAGAGATTTTCGAAGAGTATAAACAGAAAGGTAGAGCGC
NZ_CP035265.1:766128-774627    6096   ACTGACGAAGTCAGTCACATATATAATCCAAGGCGACGTTGACGTGGTTTGAAGAGATTTTCGAAGAGTATAAACAGAAAGGTAGAGCGC
NZ_CP035245.1 745261-753757    6097   ACTGACGAAGTCAGTCACATATATAATCCAAGGCGACGTTGACGTGGTTTGAAGAGATTTTAGAAGAGTATAAACAGAAAGGTAGAGCGC
NZ_CP035259.1:745700-753548    5449   ACTGACGAAGTCAGTCACATATATAATCCAAGGCGACGTTGACGTGGTTTGAAGAGATTTTCGAAGAGTATAAACAGAAAGGTAGAGCGC
NZ_CP035258.1:750874-758722    5449   ACTGACGAAGTCAGTCACATATATAATCCAAGGCGACGTTGACGTGGTTTGAAGAGATTTTCGAAGAGTATAAACAGAAAGGTAGAGCGC
NZ_CP035260.1:726156-733994    5439   ACTGACGAAGTCAGTCACATATATAATCCAAGGCGACGTTGACGTGGTTTGAAGAGATTTTCGAAGAGTATAAACAGAAAGGTAGAGCGC
NZ_CP035241.1:708296-716144    5449   ACTGACGAAGTCAGTCACATATATAATCCAAGGCGACGTTGACGTGGTTTGAAGAGATTTTCGAAGAGTATAAACAGAAAGGTAGAGCGC
NZ_CP035261.1:779086-786938    5449   ACTGACAAAGTCAGTCACATATATAATCCAAGGCGACGTTGACGTGGTTTGAAGAGATTTTCGAAGAGTATAAACAGAAAGGTAGAGCGC
NZ_CP035242.1:c1532236-1524385 5448   ACTGACGAAGTCAGTCACATATATAATCCAAGGCGACGTTGACGTGGTTTGAAGAGATTTTCGAAGAGTATAAACAGAAAGGTAGAGCGC
NZ_CP035243.1:777729-785580    5448   ACTGACGAAGTCAGTCACATATATAATCCAAGGCGACGTTGACGTGGTTTGAAGAGATTTTCGAAGAGTATAAACAGAAAGGTAGAGCGC
NZ_CP035237.1:c1311322-1303456 5463   ACTGACGAAGTCAGTCACATATATAATCCAAGGCGACGTTGACGTGGTTTGAAGAGATTTTCGAAGAGTATAAACAGAAAGGTAGAGCGC
NZ_CP035246.1:c1302892-1295026 5463   ACTGACGAAGTCAGTCACATATATAATCCAAGGCGACGTTGACGTGGTTTGAAGAGATTTTCGAAGAGTATAAACAGAAAGGTAGAGCGC
NZ_CP035247.1:c1301115-1293263 5449   ACTGACGAAGTCAGTCACATATATAATCCAAGGCGACGTTGACGTGGTTTGAAGAGATTTTCGAAGAGTATAAACAGAAAGGTAGAGCGC
NZ_CP035238.1:738547-746413    5463   ACTGACGAAGTCAGTCACATATATAATCCAAGGCGACGTTGACGTGGTTTGAAGAGATTTTCGAAGAGTATAAACAGAAAGGTAGAGCGC
NZ_CP035255.1:776866-784718    5449   ACTGACGAAGTCAGTCACATATATAATCCAAGGCGACGTTGACGTGGTTTGAAGAGATTTTCGAAGAGTATAAACAGAAAGGTAGAGCGC
NZ_CP035256.1:807236-815102    5463   ACTGACGAAGTCAGTCACATATATAATCCAAGGCGACGTTGACGTGGTTTGAAGAGATTTTCGAAGAGTATAAACAGAAAGGTAGAGCGC
NZ_CP038252.1:1151812-1159673  5392   ACTGACGAAGTCAGTCACATATATAATCCAAGGCGACGTTGACGTGGTTTGAAGAGATTTTCGAAGAGTATAAACAGAAAGGTAGAGCGC
NZ_CP035254.1:793365-801213    5449   ACTGACGAAGTCAGTCACATATATAATCCAAGGCGACGTTGACGTGGTTTGAAGAGATTTTCGAAGAGTATAAACAGAAAGGTAGAGCGC
NZ_CP035251.1:748733-756585    5453   ACTGACGAAGTCAGTCACATATATAATCCAAGGCGACGTTGACGTGGTTTGAAGAGATTTTAGAAGAGTATAAACAGAAAGGTAGAGCGC
NZ_CP035252.1:745670-753536    5467   ACTGACGAAGTCAGTCACATATATAATCCAAGGCGACGTTGACGTGGTTTGAAGAGATTTTAGAAGAGTATAAACAGAAAGGTAGAGCGC
NZ_CP035257.1:776317-784169    5453   ACTGACGAAGTCAGTCACATATATAATCCAAGGCGACGTTGACGTGGTTTGAAGAGATTTTAGAAGAGTATAAACAGAAAGGTAGAGCGC
NZ_CP038253.1:758105-765957    5453   ACTGACGAAGTCAGTCACATATATAATCCAAGGCGACGTTGACGTGGTTTGAAGAGATTTTAGAAGAGTATAAACAGAAAGGTAGAGCGC
NZ_CP035263.1:743050-750901    5448   ACTGACGAAGTCAGTCACATATATAATCCAAGGCGACGTTGACGTGGTTTGAAGAGATTTTCGAAGAGTATAAACAGAAAGGTAGAGCGC
NZ_CP035262.1:741595-749460    5462   ACTGACGAAGTCAGTCACATATATAATCCAAGGCGACGTTGACGTGGTTTGAAGAGATTTTCGAAGAGTATAAACAGAAAGGTAGAGCGC
NZ_CP035236.1:764915-772766    5448   ACTGACGAAGTCAGTCACATATATAATCCAAGGCGACGTTGACGTGGTTTGAAGAGATTTTCGAAGAGTATAAACAGAAAGGTAGAGCGC
NZ_CP035253.1 767286-775134    5449   ACTGACGAAGTCAGTCACATATATAATCCAAGGCGACGTTGACGTGGTTTGAAGAGATTTTCGAAGAGTATAAACAGAAAGGTAGAGCGC
NZ_CP035249.1:763080-770921    5449   ACTGACGAAGTCAGTCACATATATAATCCAAGGCGACGTTGACGTGGTTTGAAGAGATTTTCGAAGAGTATAAACAGAAAGGTAGAGCGC

NC_003028.3:777000-787000      6268   GTGTTCTAATTTGAACACGAGTAGAAAACTTTTCTAAAAACAAAAACGAAAGGATGGGTAAACTGTATTCGCTGAACTGAATACGGGCGA
NZ_CP035248.1:765099-775111    6277   GTGTTCTAATTTGAACACGAGTAGAAAACTTTTCTAAAAACAAAAACGAAAGGATGGGTAAACTGTATTCGCTGAACTGAATACGGGCGA
NZ_CP035235.1 c1326834-1317894 5552   GTGTTCTAATTTGAACACGAGTAGAAAACTTTTCTAAAAACAAAAACGAAAGGATGGGTAAACTGTATTCGCTGAACTGAATACGGGCGA
NZ_CP035234.1:756284-765224    5552   GTGTTCTAATTTGAACACGAGTAGAAAACTTTTCTAAAAACAAAAACGAAAGGATGGGTAAACTGTATTCGCTGAACTGAATACGGGCGA
NZ_CP038251.1:750745-759686    5552   GTGTTCTAATTTGAACACGAGTAGAAAACTTTTCTAAAAACAAAAACGAAAGGATGGGTAAACTGTATTCGCTGAACTGAATACGGGCGA
NZ_CP035244.1:747023-755950    5538   GTGTTCTAATTTGAACACGAGTAGAAAACTTTTCTAAAAACAAAAACGAAAGGATGGGTAAACTGTATTCGCTGAACTGAATACGGGCGA
NZ_CP035264.1:800836-809336    6187   GTGTTCTAATTTGAACACGAGTAGAAAACTTTTCTAAAAACAAAAACGAAAGGATGGGTAAACTGTATTCGCTGAACTGAATACGGGCGA
NZ_CP035265.1:766128-774627    6186   GTGTTCTAATTTGAACACGAGTAGAAAACTTTTCTAAAAACAAAAACGAAAGGATGGGTAAACTGTATTCGCTGAACTGAATACGGGCGA
NZ_CP035245.1 745261-753757    6187   GTGTTCTAATTTGAACACGAGTAGAAAACTTTTCTAAAAGCAAAAACGAAAGGATGGGTAAACTGTATTCGCTGAACTGAATACGGGCGA
NZ_CP035259.1:745700-753548    5539   GTGTTCTAATTTGAACACGAGTAGAAAACTTTTCTAAAAACAAAAACGAAAGGATGGGTAAACTGTATTCGCTGAACTGAATACGGGCGA
NZ_CP035258.1:750874-758722    5539   GTGTTCTAATTTGAACACGAGTAGAAAACTTTTCTAAAAACAAAAACGAAAGGATGGGTAAACTGTATTCGCTGAACTGAATACGGGCGA
NZ_CP035260.1:726156-733994    5529   GTGTTCTAATTTGAACACGAGTAGAAAACTTTTCTAAAAACAAAAACGAAAGGATGGGTAAACTGTATTCGCTGAACTGAATACGGGCGA
NZ_CP035241.1:708296-716144    5539   GTGTTCTAATTTGAACACGAGTAGAAAACTTTTCTAAAAACAAAAACGAAAGGATGGGTAAACTGTATTCGCTGAACTGAATACGGGCGA
NZ_CP035261.1:779086-786938    5539   GTGTTCTAATTTGAACACGAGTAGAAAACTTTTCTAAAAACAAAAACGAAAGGATGGGTAAACTGTATTCGCTGAACTGAATACGGGCGA
NZ_CP035242.1:c1532236-1524385 5538   GTGTTCTAATTTGAACACGAGTAGAAAACTTTTCTAAAAACAAAAACGAAAGGATGGGTAAACTGTATTCGCTGAACTGAATACGGGCGA
NZ_CP035243.1:777729-785580    5538   GTGTTCTAATTTGAACACGAGTAGAAAACTTTTCTAAAAACAAAAACGAAAGGATGGGTAAACTGTATTCGCTGAACTGAATACGGGCGA
NZ_CP035237.1:c1311322-1303456 5553   GTGTTCTAATTTGAACACGAGTAGAAAACTTTTCTAAAAACAAAAACGAAAGGATGGGTAAACTGTATTCGCTGAACTGAATACGGGCGA
NZ_CP035246.1:c1302892-1295026 5553   GTGTTCTAATTTGAACACGAGTAGAAAACTTTTCTAAAAATAAAAACGAAAGGATGGGTAAACTGTATTCGCTGAACTGAATACGGGCGA
NZ_CP035247.1:c1301115-1293263 5539   GTGTTCTAATTTGAACACGAGTAGAAAACTTTTCTAAAAATAAAAACGAAAGGATGGGTAAACTGTATTCGCTGAACTGAATACGGGCGA
NZ_CP035238.1:738547-746413    5553   GTGTTCTAATTTGAACACGAGTAGAAAACTTTTCTAAAAATAAAAACGAAAGGATGGGTAAACTGTATTCGCTGAACTGAATACGGGCGA
NZ_CP035255.1:776866-784718    5539   GTGTTCTAATTTGAACACGAGTAGAAAACTTTTCTAAAAATAAAAACGAAAGGATGGGTAAACTGTATTCGCTGAACTGAATACGGGCGA
NZ_CP035256.1:807236-815102    5553   GTGTTCTAATTTGAACACGAGTAGAAAACTTTTCTAAAAATAAAAACGAAAGGATGGGTAAACTGTATTCGCTGAACTGAATACGGGCGA
NZ_CP038252.1:1151812-1159673  5482   GTGTTCTAATTTGAACACGAGTAGAAAACTTTTCTAAAAACAAAAACGAAAGGATGGGTAAACTGTATTCGCTGAACTGAATACGGGCGA
NZ_CP035254.1:793365-801213    5539   GTGTTCTAATTTGAACACGAGTAGAAAACTTTTCTAAAAACAAAAACGAAAGGATGGGTAAACTGTATTCGCTGAACTGAATACGGGCGA
NZ_CP035251.1:748733-756585    5543   GTGTTCTAATTTGAACACGAGTAGAAAACTTTTCTAAAAACAAAAACGAAAGGATGGGTAAACTGTATTCGCTGAACTGAATACGGGCGA
NZ_CP035252.1:745670-753536    5557   GTGTTCTAATTTGAACACGAGTAGAAAACTTTTCTAAAAACAAAAACGAAAGGATGGGTAAACTGTATTCGCTGAACTGAATACGGGCGA
NZ_CP035257.1:776317-784169    5543   GTGTTCTAATTTGAACACGAGTAGAAAACTTTTCTAAAAACAAAAACGAAAGGATGGGTAAACTGTATTCGCTGAACTGAATACGGGCGA
NZ_CP038253.1:758105-765957    5543   GTGTTCTAATTTGAACACGAGTAGAAAACTTTTCTAAAAACAAAAACGAAAGGATGGGTAAACTGTATTCGCTGAACTGAATACGGGCGA
NZ_CP035263.1:743050-750901    5538   GTGTTCTAATTTGAACACGAGTAGAAAACTTTTCTAAAAATAAAAACGAAAGGATGGGTAAACTGTATTCGCTGAACTGAATACGGGCGA
NZ_CP035262.1:741595-749460    5552   GTGTTCTAATTTGAACACGAGTAGAAAACTTTTCTAAAAATAAAAACGAAAGGATGGGTAAACTGTATTCGCTGAACTGAATACGGGCGA
NZ_CP035236.1:764915-772766    5538   GTGTTCTAATTTGAACACGAGTAGAAAACTTTTCTAAAAATAAAAACGAAAGGATGGGTAAACTGTATTCGCTGAACTGAATACGGGCGA
NZ_CP035253.1 767286-775134    5539   GTGTTCTAATTTGAACACGAGTAGAAAACTTTTCTAAAAACAAAAACGAAAGGATGGGTAAACTGTATTCGCTGAACTGAATACGGGCGA
NZ_CP035249.1:763080-770921    5539   GTGTTCTAATTTGAACACGAGTAGAAAACTTTTCTAAAAACAAAAACGAAAGGATGGGTAAACTGTATTCGCTGAACTGAATACGGGCGA

NC_003028.3:777000-787000      6358   CTCTCCTCTAAATCAAAATT----AAGAAAGGAATTGACCCCACCCTAAAAGTAGTGGGAAAAAGATAGTTGATCTAGCGAGCATCGCTC
NZ_CP035248.1:765099-775111    6367   CTCTCCTCTAAATCAAAATTAAGAAAGAAAGGAATTGACCCCACCCTAAAAGCAGTGGGAAAAAGATAGTTGGTCTAGCGAGCATCGCTC
NZ_CP035235.1 c1326834-1317894 5642   CTCTCCTCTAAATCAAAATTAAGAAAGAAAGGAATTGACCCCACCCTAAAAGCAGTGGGAAAAAGATAGTTGGTCTAGCGAGCATCGCTC
NZ_CP035234.1:756284-765224    5642   CTCTCCTCTAAATCAAAATTAAGAAAGAAAGGAATTGACCCCACCCTAAAAGCAGTGGGAAAAAGATAGTTGGTCTAGCGAGCATCGCTC
NZ_CP038251.1:750745-759686    5642   CTCTCCTCTAAATCAAAATTAAGAAAGAAAGGAATTGACCCCACCCTAAAAGCAGTGGGAAAAAGATAGTTGGTCTAGCGAGCATCGCTC
NZ_CP035244.1:747023-755950    5628   CTCTCCTCTAAATCAAAATTAAGAAAGAAAGGAATTGACCCCACCCTAAAAGCAGTGGGAAAAAGATAGTTGGTCTAGCGAGCATCGCTC
NZ_CP035264.1:800836-809336    6277   CTCTCCTCTAAATCAAAATTAAGAAAGAAAGGAATTGAACCCACCCTAAAAGCAGTGGGAAAAAGATAGTTGGTCTAGCGAGCATCGCTC
NZ_CP035265.1:766128-774627    6276   CTCTCCTCTAAATCAAAATTAAGAAAGAAAGGAATTGAACCCACCCTAAAAGCAGTGGGAAAAAGATAGTTGGTCTAGCGAGCATCGCTC
```

```
NZ_CP035245.1 745261-753757    6277   CTCTCCTCTAAATCAAAATT----AAGAAAGGAATTGACCCCACCCTAAAAGTAGTGGGAAAAAGATAGTTGGTCTAGCGAGCATCGCTC
NZ_CP035259.1:745700-753548    5629   CTCTCCTCTAAATCAAAATT----AAGAAAGGAATTGACCCCACCCTAAAAGTAGTGGGAAAAAGATAGTTGATCTAGCGAGCATCGCTC
NZ_CP035258.1:750874-758722    5629   CTCTCCTCTAAATCAAAATT----AAGAAAGGAATTGACCCCACCCTAAAAGTAGTGGGAAAAAGATAGTTGATCTAGCGAGCATCGCTC
NZ_CP035260.1:726156-733994    5619   CTCTCCTCTAAATCAAAATT----AAGAAAGGAATTGACCCCACCCTAAAAGTAGTGGGAAAAAGATAGTTGGTCTAGCGAGCATCGCTC
NZ_CP035241.1:708296-716144    5629   CTCTCCTCTAAATCAAAATT----AAGAAAGGAATTGACCCCACCCTAAAAGTAGTGGGAAAAAGATAGTTGGTCTAGCGAGCATCGCTC
NZ_CP035261.1:779086-786938    5629   CTCTCCTCTAAATCAAAATTAAGAAAGAAAGGAATTGAACCCACCCTAAAAGCAGTGGGAAAAAGATAGTTGGTCTAGCGAGCATCGCTC
NZ_CP035242.1:c1532236-1524385 5628   CTCTCCTCTAAATCAAAATTAAGAAAGAAAGGAATTGAACCCACCCTAAAAGCAGTGGGAAAAAGATAGTTGGTCTAGCGAGCATCGCTC
NZ_CP035243.1:777729-785580    5628   CTCTCCTCTAAATCAAAATTAAGAAAGAAAGGAATTGAACCCACCCTAAAAGCAGTGGGAAAAAGATAGTTGGTCTAGCGAGCATCGCTC
NZ_CP035237.1:c1311322-1303456 5643   CTCTCCTCTAAATCAAAATTAAGAAAGAAAGGAATTGCCCCCACCCTAAAAGCAGTGGGAAAAAGATAGTTGATCTAGCGAGCATCGCTC
NZ_CP035246.1:c1302892-1295026 5643   CTCTCCTCTAAATCAAAATTAAGAAAGAAAGGAATTGACCCCACCCTAAAAGCAGTGGGAAAAAGATAGTTGGTCTAGCGAGCATCGCTC
NZ_CP035247.1:c1301115-1293263 5629   CTCTCCTCTAAATCAAAATTAAGAAAGAAAGGAATTGACCCCACCCTAAAAGCAGTGGGAAAAAGATAGTTGGTCTAGCGAGCATCGCTC
NZ_CP035238.1:738547-746413    5643   CTCTCCTCTAAATCAAAATTAAGAAAGAAAGGAATTGACCCCACCCTAAAAGCAGTGGGAAAAAGATAGTTGGTCTAGCGAGCATCGCTC
NZ_CP035255.1:776866-784718    5629   CTCTCCTCTAAATCAAAATTAAGAAAGAAAGGAATTGACCCCACCCTAAAAGCAGTGGGAAAAAGATAGTTGGTCTAGCGAGCATCGCTC
NZ_CP035256.1:807236-815102    5643   CTCTCCTCTAAATCAAAATTAAGAAAGAAAGGAATTGACCCCACCCTAAAAGCAGTGGGAAAAAGATAGTTGGTCTAGCGAGCATCGCTC
NZ_CP038252.1:1151812-1159673  5572   CTCTCCTCTAAATCAAAATT----AAGAAAGGAATTGACCCCACCCTAAAAGTAGTGGGAAAAAGATAGTTGATCTAGCGAGCATCGCTC
NZ_CP035254.1:793365-801213    5629   CTCTCCTCTAAATCGAAATT----AAGAAAGGAATTGACCCCACCCTAAAAGTAGTGGGAAAAAGATAGTTGATCTAGCGAGCATCGCTC
NZ_CP035251.1:748733-756585    5633   CTCTCCTCTAAATCAAAATT----AAGAAAGGAATTGACCCCACCCTAAAAGTAGTGGGAAAAAGATAGTTGGTCTAGCGAGCATCGCTC
NZ_CP035252.1:745670-753536    5647   CTCTCCTCTAAATCAAAATT----AAGAAAGGAATTGACCCCACCCTAAAAGTAGTGGGAAAAAGATAGTTGGTCTAGCGAGCATCGCTC
NZ_CP035257.1:776317-784169    5633   CTCTCCTCTAAATCAAAATT----AAGAAAGGAATTGACCCCACCCTAAAAGTAGTGGGAAAAAGATAGTTGGTCTAGCGAGCATCGCTC
NZ_CP038253.1:758105-765957    5633   CTCTCCTCTAAATCAAAATT----AAGAAAGGAATTGACCCCACCCTAAAAGTAGTGGGAAAAAGATAGTTGGTCTAGCGAGCATCGCTC
NZ_CP035263.1:743050-750901    5628   CTCTCCTCTAAATCAAAATTAAGAAAGAAAGGAATTGACCCCACCCTAAAAGCAGTGGGAAAAAGATAGTTGGTCTAGCGAGCATCGCTC
NZ_CP035262.1:741595-749460    5642   CTCTCCTCTAAATCAAAATTAAGAAAGAAAGGAATTGCCCCCACCCTAAAAGCAGTGGGAAAAAGATAGTTGGTCTAGCGAGCATCGCTC
NZ_CP035236.1:764915-772766    5628   CTCTCCTCTAAATCAAAATTAAGAAAGAAAGGAATTGACCCCACCCTAAAAGCAGTGGGAAAAAGATAGTTGGTCTAGCGAGCATCGCTC
NZ_CP035253.1 767286-775134    5629   CTCTCCTCTAAATCGAAATT----AAGAAAGGAATTGACCCCACCCTAAAAGTAGTGGGAAAAAGATAGTTGATCTAGCGAGCATCGCTC
NZ_CP035249.1:763080-770921    5629   CTCTCCTCTAAATCAAAATT----AAGAAAGGAATTGACCCCACCCTAAAAGTAGTGGGAAAAAGATAGTTGGTCTAGCGAGCATCGCTC

NC_003028.3:777000-787000      6444   ACTGCGCCCAACTCCTATTTTCCCTTCGCTTTTTGATGGGTTTGGTATCTTTCTCAATATAAAATATAAAATAAAGAAAGGTAGAGCGTG
NZ_CP035248.1:765099-775111    6457   ACTGCGCCCAACTCCTATTTTCCCTTCGCTTTTTGACGGGTTTGGTATCTTTCTCAATATAAAATATAAAATAAAGAAAGGTAGAGCGTG
NZ_CP035235.1 c1326834-1317894 5732   ACTGCGCCCAACTCCTATTTTCCCTTCGCTTTTTGACGGGTTTGGTATCTTTCTCAATATAAAATATAAAATAAAGAAAGGTAGAGCGTG
NZ_CP035234.1:756284-765224    5732   ACTGCGCCCAACTCCTATTTTCCCTTCGCTTTTTGACGGGTTTGGTATCTTTCTCAATATAAAATATAAAATAAAGAAAGGTAGAGCGTG
NZ_CP038251.1:750745-759686    5732   ACTGCGCCCAACTCCTATTTTCCCTTCGCTTTTTGACGGGTTTGGTATCTTTCTCAATATAAAATATAAAATAAAGAAAGGTAGAGCGTG
NZ_CP035244.1:747023-755950    5718   ACTGCGCCCAACTCCTATTTTCCCTTCGCTTTTTGACGGGTTTGGTATCTTTCTCAATATAAAATATAAAATAAAGAAAGGTAGAGCGTG
NZ_CP035264.1:800836-809336    6367   ACTGCGCCCAACTCCTATTTTCCCTTCGCTTTTTGACGGGTTTGGTATCTTTCTCAATATAAAATATAAAATAAAGAAAGGTAGAGCGTG
NZ_CP035265.1:766128-774627    6366   ACTGCGCCCAACTCCTATTTTCCCTTCGCTTTTTGACGGGTTTGGTATCTTTCTCAATATAAAATATAAAATAAAGAAAGGTAGAGCGTG
NZ_CP035245.1 745261-753757    6363   ACTGCGCCCAACTCCTATTTTCCCTTCGCTTTTTGATGGGTTTGGTATCTTTCTCAATATAAAATATAAAATAAAGAAAGATAGAGCGTG
NZ_CP035259.1:745700-753548    5715   ACTGCGCCCAACTCCTATTTTCCCTTCGCTTTTTGATGGGTTTGGTATCTTTCTCAATATAAAATATAAAATAAAGAAAGGTAGAGCGTG
NZ_CP035258.1:750874-758722    5715   ACTGCGCCCAACTCCTATTTTCCCTTCGCTTTTTGATGGGTTTGGTATCTTTCTCAATATAAAATATAAAATAAAGAAAGGTAGAGCGTG
NZ_CP035260.1:726156-733994    5705   ACTGCGCCCAACTCCTATTTTCCCTTCGCTTTTTGATGGGTTTGGTATCTTTCTCAATATAAAATATAAAATAAAGAAAGGTAGAGCGTG
NZ_CP035241.1:708296-716144    5715   ACTGCGCCCAACTCCTATTTTCCCTTCGCTTTTTGATGGGTTTGGTATCTTTCTCAATATAAAATATAAAATAAAGAAAGGTAGAGCGTG
NZ_CP035261.1:779086-786938    5719   ACTGCGCCCAACTCCTATTTTCCCTTCGCTTTTTGACGGGTTTGGTATCTTTCTCAATATAAAATATAAAATAAAGAAAGGTAGAGCGTG
NZ_CP035242.1:c1532236-1524385 5718   ACTGCGCCCAACTCCTATTTTCCCTTCGCTTTTTGACGGGTTTGGTATCTTTCTCAATATAAAATATAAAATAAAGAAAGGTAGAGCGTG
NZ_CP035243.1:777729-785580    5718   ACTGCGCCCAACTCCTATTTTCCCTTCGCTTTTTGACGGGTTTGGTATCTTTCTCAATATAAAATATAAAATAAAGAAAGGTAGAGCGTG
NZ_CP035237.1:c1311322-1303456 5733   ACTGCGCCCAACTCCTATTTTCCCTTCGCTTTTTGATGGGTTTGGTATCTTTCTCAATATAAAATATAAAATAAAGAAAGGTAGAGCGTG
NZ_CP035246.1:c1302892-1295026 5733   ACTGCGCCCAACTCCTATTTTCCCTTCGCTTTTTGATGGGTTTGGTATCTTTCTCAATATAAAATATAAAATAAAGAAAGGTAGAGCGTG
NZ_CP035247.1:c1301115-1293263 5719   ACTGCGCCCAACTCCTATTTTCCCTTCGCTTTTTGATGGGTTTGGTATCTTTCTCAATATAAAATATAAAATAAAGAAAGGTAGAGCGTG
NZ_CP035238.1:738547-746413    5733   ACTGCGCCCAACTCCTATTTTCCCTTCGCTTTTTGATGGGTTTGGTATCTTTCTCAATATAAAATATAAAATAAAGAAAGGTAGAGCGTG
NZ_CP035255.1:776866-784718    5719   ACTGCGCCCAACTCCTATTTTCCCTTCGCTTTTTGATGGGTTTGGTATCTTTCTCAATATAAAATATAAAATAAAGAAAGGTAGAGCGTG
NZ_CP035256.1:807236-815102    5733   ACTGCGCCCAACTCCTATTTTCCCTTCGCTTTTTGATGGGTTTGGTATCTTTCTCAATATAAAATATAAAATAAAGAAAGGTAGAGCGTG
NZ_CP038252.1:1151812-1159673  5658   ACTGCGCCCAACTCCTATTTTCCCTTCGCTTTTTGATGGGTTTGGTATCTTTCTCAATATAAAATATAAAATAAAGAAAGGTAGAGCGTG
NZ_CP035254.1:793365-801213    5715   ACTGCGCCCAACTCCTATTTTCCCTTCGCTTTTTGATGGGTTTGGTATCTTTCTCAATATAAAATATAAAATAAAGAAAGGTAGAGCGTG
NZ_CP035251.1:748733-756585    5719   ACTGCGCCCAACTCCTATTTTCCCTTCGCTTTTTGATGGGTTTGGTATCTTTCTCAATATAAAATATAAAATAAAGAAAGGTAGAGCGCG
NZ_CP035252.1:745670-753536    5733   ACTGCGCCCAACTCCTATTTTCCCTTCGCTTTTTGATGGGTTTGGTATCTTTCTCAATATAAAATATAAAATAAAGAAAGGTAGAGCGCG
NZ_CP035257.1:776317-784169    5719   ACTGCGCCCAACTCCTATTTTCCCTTCGCTTTTTGATGGGTTTGGTATCTTTCTCAATATAAAATATAAAATAAAGAAAGGTAGAGCGCG
NZ_CP038253.1:758105-765957    5719   ACTGCGCCCAACTCCTATTTTCCCTTCGCTTTTTGATGGGTTTGGTATCTTTCTCAATATAAAATATAAAATAAAGAAAGGTAGAGCGCG
NZ_CP035263.1:743050-750901    5718   ACTGCGCCCAACTCCTATTTTCCCTTCGCTTTTTGATGGGTTTGGTATCTTTCTCAATATAAAATATAAAATAAAGAAAGGTAGAGCGTG
NZ_CP035262.1:741595-749460    5732   ACTGCGCCCAACTCCTATTTTCCCTTCGCTTTTTGATGGGTTTGGTATCTTTCTCAATATAAAATATAAAATAAAGAAAGGTAGAGCGTG
NZ_CP035236.1:764915-772766    5718   ACTGCGCCCAACTCCTATTTTCCCTTCGCTTTTTGATGGGTTTGGTATCTTTCTCAATATAAAATATAAAATAAAGAAAGGTAGAGCGTG
NZ_CP035253.1 767286-775134    5715   ACTGCGCCCAACTCCTATTTTCCCTTCGCTTTTTGATGGGTTTGGTATCTTTCTCAATATAAAATATAAAATAAAGAAAGGTAGAGCGTG
NZ_CP035249.1:763080-770921    5715   ACTGCGCCCAACTCCTATTTTCCCTTCGCTTTTTGATGGGTTTGGTATCTTTCTC-------AATATAAAATAAAGAAAGGTAGAGCGTG

NC_003028.3:777000-787000      6534   TGTTTTGATTTGAACACGAGCGGAAAACTCGGAAAATAGATAATCTGACTGAAAAATCAGGATTTCTCGTCAGGTTCCTAATTTTCAGTC
NZ_CP035248.1:765099-775111    6547   TGTTTTGATTTGAACACGAGCGGAAAACTCGGAAAATAGATAATCTGACTGAAAAATCAGGATTTCTCGTCGGCTTCCTAATTTTCAGTC
NZ_CP035235.1 c1326834-1317894 5822   TGTTTTGATTTGAACACGAGCGGAAAACTCGGAAAATAGATAATCTGACTGAAAAATCAGGATTTCTCGTCGGCTTCCTAATTTTCAGTC
NZ_CP035234.1:756284-765224    5822   TGTTTTGATTTGAACACGAGCGGAAAACTCGGAAAATAGATAATCTGACTGAAAAATCAGGATTTCTCGTCGGCTTCCTAATTTTCAGTC
NZ_CP038251.1:750745-759686    5822   TGTTTTGATTTGAACACGAGCGGAAAACTCGGAAAATAGATAATCTGACTGAAAAATCAGGATTTCTCGTCGGCTTCCTAATTTTCAGTC
NZ_CP035244.1:747023-755950    5808   TGTTTTGATTTGAACACGAGCGGAAAACTCGGAAAATAGATAATCTGACTGAAAAATCAGGATTTCTCGTCGGCTTCCTAATTTTCAGTC
NZ_CP035264.1:800836-809336    6457   TGTTTTGATTTGAACACGAGCGGAAAACTCGGAAAATAGATAATCTGACTGAAAAATCAGGATTTCTCGTCAGGTTCCTAATTTTCAGTC
NZ_CP035265.1:766128-774627    6456   TGTTTTGATTTGAACACGAGCGGAAAACTCGGAAAATAGATAATCTGACTGAAAAATCAGGATTTCTCGTCAGGTTCCTAATTTTCAGTC
NZ_CP035245.1 745261-753757    6453   TGTTTTGATTTGAACACGAGCGGAAAACTCGGAAAATAGATAATCTGACTGAAAAATCAGGATTTCTCGTCAGGTTCCTAATTTTCAGTC
NZ_CP035259.1:745700-753548    5805   TGTTTTGATTTGAACACGAGCGGAAAACTCGGAAAATAGATAATCTGACTGAAAAATCAGGATTTCTCGTCAGGTTCCTAATTTTCAGTC
NZ_CP035258.1:750874-758722    5805   TGTTTTGATTTGAACACGAGCGGAAAACTCGGAAAATAGATAATCTGACTGAAAAATCAGGATTTCTCGTCAGGTTCCTAATTTTCAGTC
NZ_CP035260.1:726156-733994    5795   TGTTTTGATTTGAACACGAGCGGAAAACTCGGAAAATAGATAATCTGACTGAAAAATCAGGATTTCTCGTCAGGTTCCTAATTTTCAGTC
NZ_CP035241.1:708296-716144    5805   TGTTTTGATTTGAACACGAGCGGAAAACTCGGAAAATAGATAATCTGACTGAAAAATCAGGATTTCTCGTCAGGTTCCTAATTTTCAGTC
NZ_CP035261.1:779086-786938    5809   TGTTTTGATTTGAACACGAGCGGAAAACTCGGAAAATAGATAATCTGACTGAAAAATCAGGATTTGTCGTCAGGTTCCTAATTTTCAGTC
NZ_CP035242.1:c1532236-1524385 5808   TGTTTTGATTTGAACACGAGCGGAAAACTCGGAAAATAGATAATCTGACTGAAAAATCAGGATTTCTCGTCGGCTTCCTAATTTTCAGTC
NZ_CP035243.1:777729-785580    5808   TGTTTTGATTTGAACACGAGCGGAAAACTCGGAAAATAGATAATCTGACTGAAAAATCAGGATTTCTCGTCGGCTTCCTAATTTTCAGTC
NZ_CP035237.1:c1311322-1303456 5823   TGTTTTGATTTGAACACGAGCGGAAAACTCGGAAAATAGATAATCTGACTGAAAAATCAGGATTTCTCGTCAGGTTCCTAATTTTCAGTC
NZ_CP035246.1:c1302892-1295026 5823   TGTTTTGATTTGAACACGAGCGGAAAACTCGGAAAATAGATAATCTGACTGAAAAATCAGGATTTCTCGTCAGGTTCCTAATTTTCAGTC
NZ_CP035247.1:c1301115-1293263 5809   TGTTTTGATTTGAACACGAGCGGAAAACTCGGAAAATAGATAATCTGACTGAAAAATCAGGATTTCTCGTCAGGTTCCTAATTTTCAGTC
NZ_CP035238.1:738547-746413    5823   TGTTTTGATTTGAACACGAGCGGAAAACTCGGAAAATAGATAATCTGACTGAAAAATCAGGATTTCTCGTCAGGTTCCTAATTTTCAGTC
NZ_CP035255.1:776866-784718    5809   TGTTTTGATTTGAACACGAGCGGAAAACTCGGAAAATAGATAATCTGACTGAAAAATCAGGATTTCTCGTCAGGTTCCTAATTTTCAGTC
NZ_CP035256.1:807236-815102    5823   TGTTTTGATTTGAACACGAGCGGAAAACTCGGAAAATAGATAATCTGACTGAAAAATCAGGATTTCTCGTCAGGTTCCTAATTTTCAGTC
NZ_CP038252.1:1151812-1159673  5748   TGTTTTGATTTGAACACGATCGGAAAACTCGGAAAATAGATAATCTGACTGAAAAATCAGGATTTCTCGTCAGGTTCCTAATTTTCAGTC
NZ_CP035254.1:793365-801213    5805   TGTTTTGATTTGAACACGAGCGGAAAACTCGGAAAATAGATAATCTGACTGAAAAATCAGGATTTCTCGTCAGGTTCCTAATTTTCAGTC
NZ_CP035251.1:748733-756585    5809   TGTTCTAATTTGAACACGAGCGGAAAACTCGGAAAATAGATAATCTGACTGAAAAATCAGGATTTCTCGTCAGGTTCCTAATTTTCAGTC
NZ_CP035252.1:745670-753536    5823   TGTTCTAATTTGAACACGAGCGGAAAACTCGGAAAATAGATAATCTGACTGAAAAATCAGGATTTCTCGTCAGGTTCCTAATTTTCAGTC
NZ_CP035257.1:776317-784169    5809   TGTTCTAATTTGAACACGAGCGGAAAACTCGGAAAATAGATAATCTGACTGAAAAATCAGGATTTCTCGTCAGGTTCCTAATTTTCAGTC
NZ_CP038253.1:758105-765957    5809   TGTTCTAATTTGAACACGAGCGGAAAACTCGGAAAATAGATAATCTGACTGAAAAATCAGGATTTCTCGTCAGGTTCCTAATTTTCAGTC
NZ_CP035263.1:743050-750901    5808   TGTTTTGATTTGAACACGAGCGGAAAACTCGGAAAATAGATAATCTGACTGAAAAATCAGGATTTCTCGTCAGGTTCCTAATTTTCAGTC
NZ_CP035262.1:741595-749460    5822   TGTTTTGATTTGAACACGAGCGGAAAACTCGGAAAATAGATAATCTGACTGAAAAATCAGGATTTCTCGTCAGGTTCCTAATTTTCAGTC
NZ_CP035236.1:764915-772766    5808   TGTTTTGATTTGAACACGAGCGGAAAACTCGGAAAATAGATAATCTGACTGAAAAATCAGGATTTCTCGTCAGGTTCCTAATTTTCAGTC
NZ_CP035253.1 767286-775134    5805   TGTTTTGATTTGAACACGAGCGGAAAACTCGGAAAATAGATAATCTGACTGAAAAATCAGGATTTCTCGTCAGGTTCCTAATTTTCAGTC
NZ_CP035249.1:763080-770921    5798   TGTTTTGATTTGAACACGAGCGGAAAACTCGGAAAATAGATAATCTGACTGAAAAATCAGGATTTCTCGTCAGGTTCCTAATTTTCAGTC

NC_003028.3:777000-787000      6624   GTTTTCTTCTCGCTCTTTGTATCATAAATTATGTCTATCCATATTGCTGCTCAGCAGGGTGAAATTGCTGATAAAATTCTTCTTCCTGGG
```

```
NZ_CP035248.1:765099-775111    6637   GTTTTCTTCTCGCTCTTTGTATCATAAATTATGTCTATCCATATTGCTGCTCAGCAGGGTGAAATTGCTGATAAAATTCTTCTTCCTGGG
NZ_CP035235.1 c1326834-1317894 5912   GTTTTCTTCTCGCTCTTTGTATCATAAATTATGTCTATCCATATTGCTGCTCAGCAGGGTGAAATTGCTGATAAAATTCTTCTTCCTGGG
NZ_CP035234.1:756284-765224    5912   GTTTTCTTCTCGCTCTTTGTATCATAAATTATGTCTATCCATATTGCTGCTCAGCAGGGTGAAATTGCTGATAAAATTCTTCTTCCTGGG
NZ_CP038251.1:750745-759686    5912   GTTTTCTTCTCGCTCTTTGTATCATAAATTATGTCTATCCATATTGCTGCTCAGCAGGGTGAAATTGCTGATAAAATTCTTCTTCCTGGG
NZ_CP035244.1:747023-755950    5898   GTTTTCTTCTCGCTCTTTGTATCATAAATTATGTCTATCCATATTGCTGCTCAGCAGGGTGAAATTGCTGATAAAATTCTTCTTCCTGGG
NZ_CP035264.1:800836-809336    6547   GTTTTCTTCTCGCTCTTTGTATCATAAATTATGTCTATCCATATTGCTGCTCAGCAGGGTGAAATTGCTGATAAAATTCTTCTTCCTGGG
NZ_CP035265.1:766128-774627    6546   GTTTTCTTCTCGCTCTTTGTATCATAAATTATGTCTATCCATATTGCTGCTCAGCAGGGTGAAATTGCTGATAAAATTCTTCTTCCTGGG
NZ_CP035245.1 745261-753757    6543   GTTTTCTTCTCGCTCTTTGTATCATAAATTATGTCTATCCATATTGCTGCTCAGCAGGGTGAAATTGCTGATAAAATTCTTCTTCCTGGG
NZ_CP035259.1:745700-753548    5895   GTTTTCTTCTCGCTCTTTGTATCATAAATTATGTCTATCCATATTGCTGCTCAGCAGGGTGAAATTGCTGATAAAATTCTTCTTCCTGGG
NZ_CP035258.1:750874-758722    5895   GTTTTCTTCTCGCTCTTTGTATCATAAATTATGTCTATCCATATTGCTGCTCAGCAGGGTGAAATTGCTGATAAAATTCTTCTTCCTGGG
NZ_CP035260.1:726156-733994    5885   GTTTTCTTCTCGCTCTTTGTATCATAAATTATGTCTATCCATATTGCTGCTCAGCAGGGTGAAATTGCTGATAAAATTCTTCTTCCTGGG
NZ_CP035241.1:708296-716144    5895   GTTTTCTTCTCGCTCTTTGTATCATAAATTATGTCTATCCATATTGCTGCTCAGCAGGGTGAAATTGCTGATAAAATTCTTCTTCCTGGG
NZ_CP035261.1:779086-786938    5899   GTTTTCTTCTCGCTCTTTGTATCATAAATTATGTCTATCCATATTGCTGCTCAGCAGGGTGAAATTGCTGATAAAATTCTTCTTCCTGGG
NZ_CP035242.1:c1532236-1524385 5898   GTTTTCTTCTCGCTCTTTGTATCATAAATTATGTCTATCCATATTGCTGCTCAGCAGGGTGAAATTGCTGATAAAATTCTTCTTCCTGGG
NZ_CP035243.1:777729-785580    5898   GTTTTCTTCTCGCTCTTTGTATCATAAATTATGTCTATCCATATTGCTGCTCAGCAGGGTGAAATTGCTGATAAAATTCTTCTTCCTGGG
NZ_CP035237.1:c1311322-1303456 5913   GTTTTCTTCTCGCTCTTTGTATCATAAATTATGTCTATCCATATTGCTGCTCAGCAGGGTGAAATTGCTGATAAAATTCTTCTTCCTGGG
NZ_CP035246.1:c1302892-1295026 5913   GTTTTCTTCTCGCTCTTTGTATCATAAATTATGTCTATCCATATTGCTGCTCAGCAGGGTGAAATTGCTGATAAAATTCTTCTTCCTGGG
NZ_CP035247.1:c1301115-1293263 5899   GTTTTCTTCTCGCTCTTTGTATCATAAATTATGTCTATCCATATTGCTGCTCAGCAGGGTGAAATTGCTGATAAAATTCTTCTTCCTGGG
NZ_CP035238.1:738547-746413    5913   GTTTTCTTCTCGCTCTTTGTATCATAAATTATGTCTATCCATATTGCTGCTCAGCAGGGTGAAATTGCTGATAAAATTCTTCTTCCTGGG
NZ_CP035255.1:776866-784718    5899   GTTTTCTTCTCGCTCTTTGTATCATAAATTATGTCTATCCATATTGCTGCTCAGCAGGGTGAAATTGCTGATAAAATTCTTCTTCCTGGG
NZ_CP035256.1:807236-815102    5913   GTTTTCTTCTCGCTCTTTGTATCATAAATTATGTCTATCCATATTGCTGCTCAGCAGGGTGAAATTGCTGATAAAATTCTTCTTCCTGGG
NZ_CP038252.1:1151812-1159673  5838   GTTTTCTTCTCGCTCTTTGTATCATAAATTATGTCTATCCATATTGCTGCTCAGCAGGGTGAAATTGCTGATAAAATTCTTCTTCCTGGG
NZ_CP035254.1:793365-801213    5895   GTTTTCTTCTCGCTCTTTGTATCATAAATTATGTCTATCCATATTGCTGCTCAGCAGGGTGAAATTGCTGATAAAATTCTTCTTCCTGGG
NZ_CP035251.1:748733-756585    5899   GTTTTCTTCTCGCTCTTTGTATCATAAATTATGTCTATCCATATTGCTGCTCAGCAGGGTGAAATTGCTGATAAAATTCTTCTTCCTGGG
NZ_CP035252.1:745670-753536    5913   GTTTTCTTCTCGCTCTTTGTATCATAAATTATGTCTATCCATATTGCTGCTCAGCAGGGTGAAATTGCTGATAAAATTCTTCTTCCTGGG
NZ_CP035257.1:776317-784169    5899   GTTTTCTTCTCGCTCTTTGTATCATAAATTATGTCTATCCATATTGCTGCTCAGCAGGGTGAAATTGCTGATAAAATTCTTCTTCCTGGG
NZ_CP038253.1:758105-765957    5899   GTTTTCTTCTCGCTCTTTGTATCATAAATTATGTCTATCCATATTGCTGCTCAGCAGGGTGAAATTGCTGATAAAATTCTTCTTCCTGGG
NZ_CP035263.1:743050-750901    5898   GTTTTCTTCTCGCTCTTTGTATCATAAATTATGTCTATCCATATTGCTGCTCAGCAGGGTGAAATTGCTGATAAAATTCTTCTTCCTGGG
NZ_CP035262.1:741595-749460    5912   GTTTTCTTCTCGCTCTTTGTATCATAAATTATGTCTATCCATATTGCTGCTCAGCAGGGTGAAATTGCTGATAAAATTCTTCTTCCTGGG
NZ_CP035236.1:764915-772766    5898   GTTTTCTTCTCGCTCTTTGTATCATAAATTATGTCTATCCATATTGCTGCTCAGCAGGGTGAAATTGCTGATAAAATTCTTCTTCCTGGG
NZ_CP035253.1 767286-775134    5895   GTTTTCTTCTCGCTCTTTGTATCATAAATTATGTCTATCCATATTGCTGCTCAGCAGGGTGAAATTGCTGATAAAATTCTTCTTCCTGGG
NZ_CP035249.1:763080-770921    5888   GTTTTCTTCTCGCTCTTTGTATCATAAATTATGTCTATCCATATTGCTGCTCAGCAGGGTGAAATTGCTGATAAAATTCTTCTTCCTGGG

NC_003028.3:777000-787000      6714   GATCCTCTTCGTGCTAAGTTTATTGCGGAGAATTTCCTTGATGATGCTGTTTGTTTTAACGAAGTGCGTAACATGTTTGGTTACACTGGT
NZ_CP035248.1:765099-775111    6727   GATCCTCTTCGTGCTAAGTTTATTGCGGAGAATTTCCTTGGTGATGCTGTTTGTTTTAACGAAGTGCGTAACATGTTTGGTTACACTGGT
NZ_CP035235.1 c1326834-1317894 6002   GATCCTCTTCGTGCTAAGTTTATTGCGGAGAATTTCCTTGGTGATGCTGTTTGTTTTAACGAAGTGCGTAACATGTTTGGTTACACTGGT
NZ_CP035234.1:756284-765224    6002   GATCCTCTTCGTGCTAAGTTTATTGCGGAGAATTTCCTTGGTGATGCTGTTTGTTTTAACGAAGTGCGTAACATGTTTGGTTACACTGGT
NZ_CP038251.1:750745-759686    6002   GATCCTCTTCGTGCTAAGTTTATTGCGGAGAATTTCCTTGGTGATGCTGTTTGTTTTAACGAAGTGCGTAACATGTTTGGTTACACTGGT
NZ_CP035244.1:747023-755950    5988   GATCCTCTTCGTGCTAAGTTTATTGCGGAGAATTTCCTTGGTGATGCTGTTTGTTTTAACGAAGTGCGTAACATGTTTGGTTACACTGGT
NZ_CP035264.1:800836-809336    6637   GATCCTCTTCGTGCTAAGTTTATTGCGGAGAATTTCCTTGATGATGCTGTTTGTTTTAACGAAGTGCGTAACATGTTTGGTTACACTGGT
NZ_CP035265.1:766128-774627    6636   GATCCTCTTCGTGCTAAGTTTATTGCGGAGAATTTCCTTGATGATGCTGTTTGTTTTAACGAAGTGCGTAACATGTTTGGTTACACTGGT
NZ_CP035245.1 745261-753757    6633   GATCCTCTTCGTGCTAAGTTTATTGCGGAGAATTTCCTTGATGATGCTGTTTGTTTTAACGAAGTGCGTAACATGTTTGGTTACACTGGT
NZ_CP035259.1:745700-753548    5985   GATCCTCTTCGTGCTAAGTTTATTGCGGAGAATTTCCTTGATGATGCTGTTTGTTTTAACGAAGTGCGTAACATGTTTGGTTACACTGGT
NZ_CP035258.1:750874-758722    5985   GATCCTCTTCGTGCTAAGTTTATTGCGGAGAATTTCCTTGATGATGCTGTTTGTTTTAACGAAGTGCGTAACATGTTTGGTTACACTGGT
NZ_CP035260.1:726156-733994    5975   GATCCTCTTCGTGCTAAGTTTATTGCGGAGAATTTCCTTGGTGATGCTGTTTGTTTTAACGAAGTGCGTAACATGTTTGGTTACACTGGT
NZ_CP035241.1:708296-716144    5985   GATCCTCTTCGTGCTAAGTTTATTGCGGAGAATTTCCTTGGTGATGCTGTTTGTTTTAACGAAGTGCGTAACATGTTTGGTTACACTGGT
NZ_CP035261.1:779086-786938    5989   GATCCTCTTCGTGCTAAGTTTATTGCGGAGAATTTCCTTGGTGATGCTGTTTGTTTTAACGAAGTGCGTAACATGTTTGGTTACACTGGT
NZ_CP035242.1:c1532236-1524385 5988   GATCCTCTTCGTGCTAAGTTTATTGCGGAGAATTTCCTTGGTGATGCTGTTTGTTTTAACGAAGTGCGTAACATGTTTGGTTACACTGGT
NZ_CP035243.1:777729-785580    5988   GATCCTCTTCGTGCTAAGTTTATTGCGGAGAATTTCCTTGGTGATGCTGTTTGTTTTAACGAAGTGCGTAACATGTTTGGTTACACTGGT
NZ_CP035237.1:c1311322-1303456 6003   GATCCTCTTCGTGCTAAGTTTATTGCGGAGAATTTCCTTGATGATGCTGTTTGTTTTAACGAAGTGCGTAACATGTTTGGTTACACTGGT
NZ_CP035246.1:c1302892-1295026 6003   GATCCTCTTCGTGCTAAGTTTATTGCGGAGAATTTCCTTGGTGATGCTGTTTGTTTTAACGAAGTGCGTAACATGTTTGGTTACACTGGT
NZ_CP035247.1:c1301115-1293263 5989   GATCCTCTTCGTGCTAAGTTTATTGCGGAGAATTTCCTTGGTGATGCTGTTTGTTTTAACGAAGTGCGTAACATGTTTGGTTACACTGGT
NZ_CP035238.1:738547-746413    6003   GATCCTCTTCGTGCTAAGTTTATTGCGGAGAATTTCCTTGGTGATGCTGTTTGTTTTAACGAAGTGCGTAACATGTTTGGTTACACTGGT
NZ_CP035255.1:776866-784718    5989   GATCCTCTTCGTGCTAAGTTTATTGCGGAGAATTTCCTTGGTGATGCTGTTTGTTTTAACGAAGTGCGTAACATGTTTGGTTACACTGGT
NZ_CP035256.1:807236-815102    6003   GATCCTCTTCGTGCTAAGTTTATTGCGGAGAATTTCCTTGGTGATGCTGTTTGTTTTAACGAAGTGCGTAACATGTTTGGTTACACTGGT
NZ_CP038252.1:1151812-1159673  5928   GATCCTCTTCGTGCTAAGTTTATTGCGGAGAATTTCCTTGATGATGCTGTTTGTTTTAACGAAGTGCGTAACATGTTTGGTTACACTGGT
NZ_CP035254.1:793365-801213    5985   GATCCTCTTCGTGCTAAGTTTATTGCGGAGAATTTCCTTGATGATGCTGTTTGTTTTAACGAAGTGCGTAACATGTTTGGTTACACTGGT
NZ_CP035251.1:748733-756585    5989   GATCCTCTTCGTGCTAAGTTTATTGCGGAGAATTTCCTTGATGATGCTGTTTGTTTTAACGAAGTGCGTAACATGTTTGGTTACACTGGT
NZ_CP035252.1:745670-753536    6003   GATCCTCTTCGTGCTAAGTTTATTGCGGAGAATTTCCTTGATGATGCTGTTTGTTTTAACGAAGTGCGTAACATGTTTGGTTACACTGGT
NZ_CP035257.1:776317-784169    5989   GATCCTCTTCGTGCTAAGTTTATTGCGGAGAATTTCCTTGATGATGCTGTTTGTTTTAACGAAGTGCGTAACATGTTTGGTTACACTGGT
NZ_CP038253.1:758105-765957    5989   GATCCTCTTCGTGCTAAGTTTATTGCGGAGAATTTCCTTGATGATGCTGTTTGTTTTAACGAAGTGCGTAACATGTTTGGTTACACTGGT
NZ_CP035263.1:743050-750901    5988   GATCCTCTTCGTGCTAAGTTTATTGCGGAGAATTTCCTTGATGATGCTGTTTGTTTTAACGAAGTGCGTAACATGTTTGGTTACACTGGT
NZ_CP035262.1:741595-749460    6002   GATCCTCTTCGTGCTAAGTTTATTGCGGAGAATTTCCTTGATGATGCTGTTTGTTTTAACGAAGTGCGTAACATGTTTGGTTACACTGGT
NZ_CP035236.1:764915-772766    5988   GATCCTCTTCGTGCTAAGTTTATTGCGGAGAATTTCCTTGATGATGCTGTTTGTTTTAACGAAGTGCGTAACATGTTTGGTTACACTGGT
NZ_CP035253.1 767286-775134    5985   GATCCTCTTCGTGCTAAGTTTATTGCGGAGAATTTCCTTGATGATGCTGTTTGTTTTAACGAAGTGCGTAACATGTTTGGTTACACTGGT
NZ_CP035249.1:763080-770921    5978   GATCCTCTTCGTGCTAAGTTTATTGCGGAGAATTTCCTTGATGATGCTGTTTGTTTTAACGAAGTGCGTAACATGTTTGGTTACACTGGT

NC_003028.3:777000-787000      6804   ACTTACAAGGGTCACTGTGTATCTGTCATGGGAACTGGGATGGGAATGCCATCTATTTCGATTTATGCGCGTGAGTTAATCGTAGACTAC
NZ_CP035248.1:765099-775111    6817   ACTTACAAGGGTCACCGTGTATCTGTCATGGGAACTGGGATGGGAATGCCATCTATTTCGATTTATGCGCGTGAGTTAATCGTAGACTAC
NZ_CP035235.1 c1326834-1317894 6092   ACTTACAAGGGTCACCGTGTATCTGTCATGGGAACTGGGATGGGAATGCCATCTATTTCGATTTATGCGCGTGAGTTAATCGTAGACTAC
NZ_CP035234.1:756284-765224    6092   ACTTACAAGGGTCACCGTGTATCTGTCATGGGAACTGGGATGGGAATGCCATCTATTTCGATTTATGCGCGTGAGTTAATCGTAGACTAC
NZ_CP038251.1:750745-759686    6092   ACTTACAAGGGTCACCGTGTATCTGTCATGGGAACTGGGATGGGAATGCCATCTATTTCGATTTATGCGCGTGAGTTAATCGTAGACTAC
NZ_CP035244.1:747023-755950    6078   ACTTACAAGGGTCACCGTGTATCTGTCATGGGAACTGGGATGGGAATGCCATCTATTTCGATTTATGCGCGTGAGTTAATCGTAGACTAC
NZ_CP035264.1:800836-809336    6727   ACTTACAAGGGTCACTGTGTATCTGTCATGGGAACTGGGATGGGAATGCCATCTATTTCGATTTATGCGCGTGAGTTAATCGTAGACTAC
NZ_CP035265.1:766128-774627    6726   ACTTACAAGGGTCACTGTGTATCTGTCATGGGAACTGGGATGGGAATGCCATCTATTTCGATTTATGCGCGTGAGTTAATCGTAGACTAC
NZ_CP035245.1 745261-753757    6723   ACTTACAAGGGTCACTGTGTATCTGTCATGGGAACTGGGATGGGAATGCCATCTATTTCGATTTATGCGCGTGAGTTAATCGTAGACTAC
NZ_CP035259.1:745700-753548    6075   ACTTACAAGGGTCACTGTGTATCTGTCATGGGAACTGGGATGGGAATGCCATCTATTTCGATTTATGCGCGTGAGTTAATCGTAGACTAC
NZ_CP035258.1:750874-758722    6075   ACTTACAAGGGTCACTGTGTATCTGTCATGGGAACTGGGATGGGAATGCCATCTATTTCGATTTATGCGCGTGAGTTAATCGTAGACTAC
NZ_CP035260.1:726156-733994    6065   ACTTACAAGGGTCACCGTGTATCTGTCATGGGAACTGGGATGGGAATGCCATCTATTTCGATTTATGCGCGTGAGTTAATCGTAGACTAC
NZ_CP035241.1:708296-716144    6075   ACTTACAAGGGTCACCGTGTATCTGTCATGGGAACTGGGATGGGAATGCCATCTATTTCGATTTATGCGCGTGAGTTAATCGTAGACTAC
NZ_CP035261.1:779086-786938    6079   ACTTACAAGGGTCACCGTGTATCTGTCATGGGAACTGGGATGGGAATGCCATCTATTTCGATTTATGCGCGTGAGTTAATCGTAGACTAC
NZ_CP035242.1:c1532236-1524385 6078   ACTTACAAGGGTCACCGTGTATCTGTCATGGGAACTGGGATGGGAATGCCATCTATTTCGATTTATGCGCGTGAGTTAATCGTAGACTAC
NZ_CP035243.1:777729-785580    6078   ACTTACAAGGGTCACCGTGTATCTGTCATGGGAACTGGGATGGGAATGCCATCTATTTCGATTTATGCGCGTGAGTTAATCGTAGACTAC
NZ_CP035237.1:c1311322-1303456 6093   ACTTACAAGGGTCACTGTGTATCTGTCATGGGAACTGGGATGGGAATGCCATCTATTTCGATTTATGCGCGTGAGTTAATCGTAGACTAC
NZ_CP035246.1:c1302892-1295026 6093   ACTTACAAGGGTCACCGTGTATCTGTCATGGGAACTGGGATGGGAATGCCATCTATTTCGATTTATGCGCGTGAGTTAATCGTAGAATAC
NZ_CP035247.1:c1301115-1293263 6079   ACTTACAAGGGTCACCGTGTATCTGTCATGGGAACTGGGATGGGAATGCCATCTATTTCGATTTATGCGCGTGAGTTAATCGTAGAATAC
NZ_CP035238.1:738547-746413    6093   ACTTACAAGGGTCACCGTGTATCTGTCATGGGAACTGGGATGGGAATGCCATCTATTTCGATTTATGCGCGTGAGTTAATCGTAGACTAC
NZ_CP035255.1:776866-784718    6079   ACTTACAAGGGTCACCGTGTATCTGTCATGGGAACTGGGATGGGAATGCCATCTATTTCGATTTATGCGCGTGAGTTAATCGTAGACTAC
NZ_CP035256.1:807236-815102    6093   ACTTACAAGGGTCACCGTGTATCTGTCATGGGAACTGGGATGGGAATGCCATCTATTTCGATTTATGCGCGTGAGTTAATCGTAGACTAC
NZ_CP038252.1:1151812-1159673  6018   ACTTACAAGGGTCACTGTGTATCTGTCATGGGAACTGGGATGGGAATGCCATCTATTTCGATTTATGCGCGTGAGTTAATCGTAGACTAC
NZ_CP035254.1:793365-801213    6075   ACTTACAAGGGTCACTGTGTATCTGTCATGGGAACTGGGATGGGAATGCCATCTATTTCGATTTATGCGCGTGAGTTAATCGTAGACTAC
NZ_CP035251.1:748733-756585    6079   ACTTACAAGGGTCACTGTGTATCTGTCATGGGAACTGGGATGGGAATGCCATCTATTTCGATTTATGCGCGTGAGTTAATCGTAGACTAC
NZ_CP035252.1:745670-753536    6093   ACTTACAAGGGTCACTGTGTATCTGTCATGGGAACTGGGATGGGAATGCCATCTATTTCGATTTATGCGCGTGAGTTAATCGTAGACTAC
NZ_CP035257.1:776317-784169    6079   ACTTACAAGGGTCACTGTGTATCTGTCATGGGAACTGGGATGGGAATGCCATCTATTTCGATTTATGCGCGTGAGTTAATCGTAGACTAC
NZ_CP038253.1:758105-765957    6079   ACTTACAAGGGTCACTGTGTATCTGTCATGGGAACTGGGATGGGAATGCCATCTATTTCGATTTATGCGCGTGAGTTAATCGTAGACTAC
```

```
NZ_CP035263.1:743050-750901    6078   ACTTACAAGGGTCACTGTGTATCTGTCATGGGAACTGGGATGGGAATGCCATCTATTTCGATTTATGCGCGTGAGTTAATCGTAGACTAC
NZ_CP035262.1:741595-749460    6092   ACTTACAAGGGTCACTGTGTATCTGTCATGGGAACTGGGATGGGAATGCCATCTATTTCGATTTATGCGCGTGAGTTAATCGTAGACTAC
NZ_CP035236.1:764915-772766    6078   ACTTACAAGGGTCACTGTGTATCTGTCATGGGAACTGGGATGGGAATGCCATCTATTTCGATTTATGCGCGTGAGTTAATCGTAGACTAC
NZ_CP035253.1 767286-775134    6075   ACTTACAAGGGTCACTGTGTATCTGTCATGGGAACTGGGATGGGAATGCCATCTATTTCGATTTATGCGCGTGAGTTAATCGTAGACTAC
NZ_CP035249.1:763080-770921    6068   ACTTACAAGGGTCACTGTGTATCTGTCATGGGAACTGGGATGGGAATGCCATCTATTTCGATTTATGCGCGTGAGTTAATCGTAGACTAC

NC_003028.3:777000-787000      6894   GGTGTGAAGAAATTGATTCGTGTGGGAACTGCAGGTTCTTTGAATGAAGAGGTTCATGTTCGTGAATTAGTTTTGGCGCAGGCGGCTGCA
NZ_CP035248.1:765099-775111    6907   GGTGTGAAGAAATTGATTCGTGTGGGAACTGCAGGTTCTTTGAATGAAGAGGTTCATGTTCGTGAATTAGTTTTGGCGCAGGCGGCTGCA
NZ_CP035235.1 c1326834-1317894 6182   GGTGTGAAGAAATTGATTCGTGTGGGAACTGCAGGTTCTTTGAATGAAGAGGTTCATGTTCGTGAATTAGTTTTGGCGCAGGCGGCTGCA
NZ_CP035234.1:756284-765224    6182   GGTGTGAAGAAATTGATTCGTGTGGGAACTGCAGGTTCTTTGAATGAAGAGGTTCATGTTCGTGAATTAGTTTTGGCGCAGGCGGCTGCA
NZ_CP038251.1:750745-759686    6182   GGTGTGAAGAAATTGATTCGTGTGGGAACTGCAGGTTCTTTGAATGAAGAGGTTCATGTTCGTGAATTAGTTTTGGCGCAGGCGGCTGCA
NZ_CP035244.1:747023-755950    6168   GGTGTGAAGAAATTGATTCGTGTGGGAACTGCAGGTTCTTTGAATGAAGAGGTTCATGTTCGTGAATTAGTTTTGGCGCAGGCGGCTGCA
NZ_CP035264.1:800836-809336    6817   GGTGTGAAGAAATTGATTCGTGTGGGAACTGCAGGTTCTTTGAATGAAGAGGTTCATGTTCGTGAATTAGTTTTGGCGCAGGCGGCTGCA
NZ_CP035265.1:766128-774627    6816   GGTGTGAAGAAATTGATTCGTGTGGGAACTGCAGGTTCTTTGAATGAAGAGGTTCATGTTCGTGAATTAGTTTTGGCGCAGGCGGCTGCA
NZ_CP035245.1 745261-753757    6813   GGTGTGAAGAAATTGATTCGTGTGGGAACTGCAGGTTCTTTGAATGAAGAGGTTCATGTTCGTGAATTAGTTTTGGCGCAGGCGGCTGCA
NZ_CP035259.1:745700-753548    6165   GGTGTGAAGAAATTGATTCGTGTGGGAACTGCAGGTTCTTTGAATGAAGAGGTTTATGTTCGTGAATTAGTTTTGGCGCAGGCGGCTGCA
NZ_CP035258.1:750874-758722    6165   GGTGTGAAGAAATTGATTCGTGTGGGAACTGCAGGTTCTTTGAATGAAGAGGTTTATGTTCGTGAATTAGTTTTGGCGCAGGCGGCTGCA
NZ_CP035260.1:726156-733994    6155   GGTGTGAAGAAATTGATTCGTGTGGGAACTGCAGGTTCTTTGAATGAAGAGGTTCATGTTCGTGAATTAGTTTTGGCGCAGGCGGCTGCA
NZ_CP035241.1:708296-716144    6165   GGTGTGAAGAAATTGATTCGTGTGGGAACTGCAGGTTCTTTGAATGAAGAGGTTCATGTTCGTGAATTAGTTTTGGCGCAGGCGGCTGCA
NZ_CP035261.1:779086-786938    6169   GGTGTGAAGAAATTGATTCGTGTGGGAACTGCAGGTTCTTTGAATGAAGAGGTTCATGTTCGTGAATTAGTTTTGGCGCAGGCGGCTGCA
NZ_CP035242.1:c1532236-1524385 6168   GGTGTGAAGAAATTGATTCGTGTGGGAACTGCAGGTTCTTTGAATGAAGAGGTTCATGTTCGTGAATTAGTTTTGGCGCAGGCGGCTGCA
NZ_CP035243.1:777729-785580    6168   GGTGTGAAGAAATTGATTCGTGTGGGAACTGCAGGTTCTTTGAATGAAGAGGTTCATGTTCGTGAATTAGTTTTGGCGCAGGCGGCTGCA
NZ_CP035237.1:c1311322-1303456 6183   GGTGTGAAGAAATTGATTCGTGTGGGAACTGCAGGTTCTTTGAATGAAGAGGTTCATGTTCGTGAATTAGTTTTGGCGCAGGCGGCTGCA
NZ_CP035246.1:c1302892-1295026 6183   GGTGTGAAGAAATTGATTCGTGTGGGAACTGCAGGTTCTTTGAATGAAGAGGTTCATGTTCGTGAATTAGTTTTGGCGCAGGCGGCTGCA
NZ_CP035247.1:c1301115-1293263 6169   GGTGTGAAGAAATTGATTCGTGTGGGAACTGCAGGTTCTTTGAATGAAGAGGTTCATGTTCGTGAATTAGTTTTGGCGCAGGCGGCTGCA
NZ_CP035238.1:738547-746413    6183   GGTGTGAAGAAATTGATTCGTGTGGGAACTGCAGGTTCTTTGAATGAAGAGGTTCATGTTCGTGAATTAGTTTTGGCGCAGGCGGCTGCA
NZ_CP035255.1:776866-784718    6169   GGTGTGAAGAAATTGATTCGTGTGGGAACTGCAGGTTCTTTGAATGAAGAGGTTCATGTTCGTGAATTAGTTTTGGCGCAGGCGGCTGCA
NZ_CP035256.1:807236-815102    6183   GGTGTGAAGAAATTGATTCGTGTGGGAACTGCAGGTTCTTTGAATGAAGAGGTTCATGTTCGTGAATTAGTTTTGGCGCAGGCGGCTGCA
NZ_CP038252.1:1151812-1159673  6108   GGTGTGAAGAAATTGATTCGTGTGGGAACTGCAGGTTCTTTGAATGAAGAGGTTCATGTTCGTGAATTAGTTTTGGCGCAGGCGGCTGCA
NZ_CP035254.1:793365-801213    6165   GGTGTGAAGAAATTGATTCGTGTGGGAACTGCAGGTTCTTTGAATGAAGAGGTTCATGTTCGTGAATTAGTTTTGGCGCAGGCGGCTGCA
NZ_CP035251.1:748733-756585    6169   GGTGTGAAGAAATTGATTCGTGTGGGAACTGCAGGTTCTTTGAATGAAGAGGTTCATGTTCGTGAATTAGTTTTGGCGCAGGCGGCTGCA
NZ_CP035252.1:745670-753536    6183   GGTGTGAAGAAATTGATTCGTGTGGGAACTGCAGGTTCTTTGAATGAAGAGGTTCATGTTCGTGAATTAGTTTTGGCGCAGGCGGCTGCA
NZ_CP035257.1:776317-784169    6169   GGTGTGAAGAAATTGATTCGTGTGGGAACTGCAGGTTCTTTGAATGAAGAGGTTCATGTTCGTGAATTAGTTTTGGCGCAGGCGGCTGCA
NZ_CP038253.1:758105-765957    6169   GGTGTGAAGAAATTGATTCGTGTGGGAACTGCAGGTTCTTTGAATGAAGAGGTTCATGTTCGTGAATTAGTTTTGGCGCAGGCGGCTGCA
NZ_CP035263.1:743050-750901    6168   GGTGTGAAGAAATTGATTCGTGTGGGAACTGCAGGTTCTTTGAATGAAGAGGTTCATGTTCGTGAATTAGTTTTGGCGCAGGCGGCTGCA
NZ_CP035262.1:741595-749460    6182   GGTGTGAAGAAATTGATTCGTGTGGGAACTGCAGGTTCTTTGAATGAAGAGGTTCATGTTCGTGAATTAGTTTTGGCGCAGGCGGCTGCA
NZ_CP035236.1:764915-772766    6168   GGTGTGAAGAAATTGATTCGTGTGGGAACTGCAGGTTCTTTGAATGAAGAGGTTCATGTTCGTGAATTAGTTTTGGCGCAGGCGGCTGCA
NZ_CP035253.1 767286-775134    6165   GGTGTGAAGAAATTGATTCGTGTGGGAACTGCAGGTTCTTTGAATGAAGAGGTTCATGTTCGTGAATTAGTTTTGGCGCAGGCGGCTGCA
NZ_CP035249.1:763080-770921    6158   GGTGTGAAGAAATTGATTCGTGTGGGAACTGCAGGTTCTTTGAATGAAGAGGTTCATGTTCGTGAATTAGTTTTGGCGCAGGCGGCTGCA

NC_003028.3:777000-787000      6984   ACCAACTCAAACATCGTTCGTAATGACTGGCCACAGTACGATTTTCCACAAATTGCTAGCTTTGATTTGCTTGATAAAGCCTACCATATC
NZ_CP035248.1:765099-775111    6997   ACCAACTCAAACATCGTTCGTAATGACTGGCCACAGTACGATTTTCCACAAATTGCTAGCTTTGATTTGCTTGATAAAGCCTACCATATC
NZ_CP035235.1 c1326834-1317894 6272   ACCAACTCAAACATCGTTCGTAATGACTGGCCACAGTACGATTTTCCACAAATTGCTAGCTTTGATTTGCTTGATAAAGCCTACCATATC
NZ_CP035234.1:756284-765224    6272   ACCAACTCAAACATCGTTCGTAATGACTGGCCACAGTACGATTTTCCACAAATTGCTAGCTTTGATTTGCTTGATAAAGCCTACCATATC
NZ_CP038251.1:750745-759686    6272   ACCAACTCAAACATCGTTCGTAATGACTGGCCACAGTACGATTTTCCACAAATTGCTAGCTTTGATTTGCTTGATAAAGCCTACCATATC
NZ_CP035244.1:747023-755950    6258   ACCAACTCAAACATCGTTCGTAATGACTGGCCACAGTACGATTTTCCACAAATTGCTAGCTTTGATTTGCTTGATAAAGCCTACCATATC
NZ_CP035264.1:800836-809336    6907   ACCAACTCAAACATCGTTCGTAATGACTGGCCACAGTACGATTTTCCACAAATTGCTAGCTTTGATTTGCTTGATAAAGCCTACCATATC
NZ_CP035265.1:766128-774627    6906   ACCAACTCAAACATCGTTCGTAATGACTGGCCACAGTACGATTTTCCACAAATTGCTAGCTTTGATTTGCTTGATAAAGCCTACCATATC
NZ_CP035245.1 745261-753757    6903   ACCAACTCAAACATCGTTCGTAATGACTGGCCACAGTACGATTTTCCACAAATTGCTAGCTTTGATTTGCTTGATAAAGCTTACCATATC
NZ_CP035259.1:745700-753548    6255   ACCAACTCAAACATCGTTCGTAATGACTGGCCACAGTACGATTTTCCACAAATTGCTAGCTTTGATTTGCTTGATAAAGCCTACCATATC
NZ_CP035258.1:750874-758722    6255   ACCAACTCAAACATCGTTCGTAATGACTGGCCACAGTACGATTTTCCACAAATTGCTAGCTTTGATTTGCTTGATAAAGCCTACCATATC
NZ_CP035260.1:726156-733994    6245   ACCAACTCAAACATCGTTCGTAATGACTGGCCACAGTACGATTTTCCACAAATTGCTAGCTTTGATTTGCTTGATAAAGCCTACCATATC
NZ_CP035241.1:708296-716144    6255   ACCAACTCAAACATCGTTCGTAATGACTGGCCACAGTACGATTTTCCACAAATTGCTAGCTTTGATTTGCTTGATAAAGCCTACCATATC
NZ_CP035261.1:779086-786938    6259   ACCAACTCAAACATCGTTCGTAATGACTGGCCACAGTACGATTTTCCACAAATTGCTAGCTTTGATTTGCTTGATAAAGCCTACCATATC
NZ_CP035242.1:c1532236-1524385 6258   ACCAACTCAAACATCGTTCGTAATGACTGGCCACAGTACGATTTTCCACAAATTGCTAGCTTTGATTTGCTTGATAAAGCCTACCATATC
NZ_CP035243.1:777729-785580    6258   ACCAACTCAAACATCGTTCGTAATGACTGGCCACAGTACGATTTTCCACAAATTGCTAGCTTTGATTTGCTTGATAAAGCCTACCATATC
NZ_CP035237.1:c1311322-1303456 6273   ACCAACTCAAACATCGTTCGTAATGACTGGCCACAGTACGATTTTCCACAAATTGCTAGCTTTGATTTGCTTGATAAAGCCTACCATATC
NZ_CP035246.1:c1302892-1295026 6273   ACCAACTCAAACATCGTTCGTAATGACTGGCCACAGTACGATTTTCCACAAATTGCTAGCTTTGATTTGCTTGATAAAGCCTACCATATC
NZ_CP035247.1:c1301115-1293263 6259   ACCAACTCAAACATCGTTCGTAATGACTGGCCACAGTACGATTTTCCACAAATTGCTAGCTTTGATTTGCTTGATAAAGCCTACCATATC
NZ_CP035238.1:738547-746413    6273   ACCAACTCAAACATCGTTCGTAATGACTGGCCACAGTACGATTTTCCACAAATTGCTAGCTTTGATTTGCTTGATAAAGCCTACCATATC
NZ_CP035255.1:776866-784718    6259   ACCAACTCAAACATCGTTCGTAATGACTGGCCACAGTACGATTTTCCACAAATTGCTAGCTTTGATTTGCTTGATAAAGCCTACCATATC
NZ_CP035256.1:807236-815102    6273   ACCAACTCAAACATCGTTCGTAATGACTGGCCACAGTACGATTTTCCACAAATTGCTAGCTTTGATTTGCTTGATAAAGCCTACCATATC
NZ_CP038252.1:1151812-1159673  6198   ACCAACTCAAACATCGTTCGTAATGACTGGCCACAGTACGATTTTCCACAAATTGCTAGCTTTGATTTGCTTGATAAAGCCTACCATATC
NZ_CP035254.1:793365-801213    6255   ACCAACTCAAACATCGTTCGTAATGACTGGCCACAGTACGATTTTCCACAAATTGCTAGCTTTGATTTGCTTGATAAAGCCTACCATATC
NZ_CP035251.1:748733-756585    6259   ACCAACTCAAACATCGTTCGTAATGACTGGCCACAGTACGATTTTCCACAAATTGCTAGCTTTGATTTGCTTGATAAAGCCTACCATATC
NZ_CP035252.1:745670-753536    6273   ACCAACTCAAACATCGTTCGTAATGACTGGCCACAGTACGATTTTCCACAAATTGCTAGCTTTGATTTGCTTGATAAAGCCTACCATATC
NZ_CP035257.1:776317-784169    6259   ACCAACTCAAACATCGTTCGTAATGACTGGCCACAGTACGATTTTCCACAAATTGCTAGCTTTGATTTGCTTGATAAAGCCTACCATATC
NZ_CP038253.1:758105-765957    6259   ACCAACTCAAACATCGTTCGTAATGACTGGCCACAGTACGATTTTCCACAAATTGCTAGCTTTGATTTGCTTGATAAAGCCTACCATATC
NZ_CP035263.1:743050-750901    6258   ACCAACTCAAACATCGTTCGTAATGACTGGCCACAGTACGATTTTCCACAAATTGCTAGCTTTGATTTGCTTGATAAAGCCTACCATATC
NZ_CP035262.1:741595-749460    6272   ACCAACTCAAACATCGTTCGTAATGACTGGCCACAGTACGATTTTCCACAAATTGCTAGCTTTGATTTGCTTGATAAAGCCTACCATATC
NZ_CP035236.1:764915-772766    6258   ACCAACTCAAACATCGTTCGTAATGACTGGCCACAGTACGATTTTCCACAAATTGCTAGCTTTGATTTGCTTGATAAAGCCTACCATATC
NZ_CP035253.1 767286-775134    6255   ACCAACTCAAACATCGTTCGTAATGACTGGCCACAGTACGATTTTCCACACATTGCTAGCTTTGATTTGCTTGATAAAGCCTACCATATC
NZ_CP035249.1:763080-770921    6248   ACCAACTCAAACATCGTTCGTAATGACTGGCCACAGTACGATTTTCCACAAATTGCTAGCTTTGATTTGCTTGATAAAGCCTACCATATC

NC_003028.3:777000-787000      7074   GCCAAAAAACTTGGTATGACTACTCACGTTGGGAACGTTTTGTCATCTGATGTCTTTTACTCAAATTACTTTGAAAAGAATATCGAGCTT
NZ_CP035248.1:765099-775111    7087   GCCAAAGAACTTGGTATGACTACTCACGTTGGGAACGTTTTGTCATCTGATGTCTTTTACTCAAATTACTTTGAAAAGAATATCGAGCTT
NZ_CP035235.1 c1326834-1317894 6362   GCCAAAGAACTTGGTATGACTACTCACGTTGGGAACGTTTTGTCATCTGATGTCTTTTACTCAAATTACTTTGAAAAGAATATCGAGCTT
NZ_CP035234.1:756284-765224    6362   GCCAAAGAACTTGGTATGACTACTCACGTTGGGAACGTTTTGTCATCTGATGTCTTTTACTCAAATTACTTTGAAAAGAATATCGAGCTT
NZ_CP038251.1:750745-759686    6362   GCCAAAGAACTTGGTATGACTACTCACGTTGGGAACGTTTTGTCATCTGATGTCTTTTACTCAAATTACTTTGAAAAGAATATCGAGCTT
NZ_CP035244.1:747023-755950    6348   GCCAAAGAACTTGGTATGACTACTCACGTTGGGAACGTTTTGTCATCTGATGTCTTTTACTCAAATTACTTTGAAAAGAATATCGAGCTT
NZ_CP035264.1:800836-809336    6997   GCCAAAGAACTTGGTATGACTACTCACGTTGGGAACGTTTTGTCATCTGATGTCTTTTACTCAAATTACTTTGAAAAGAATATCGAGCTT
NZ_CP035265.1:766128-774627    6996   GCCAAAGAACTTGGTATGACTACTCACGTTGGGAACGTTTTGTCATCTGATGTCTTTTACTCAAATTACTTTGAAAAGAATATCGAGCTT
NZ_CP035245.1 745261-753757    6993   GCCAAAAAACTTGGTATGACTACTCACGTTGGGAACGTTTTGTCATCTGATGTCTTTTACTCAAATTACTTTGAAAAGAATATCGAGCTT
NZ_CP035259.1:745700-753548    6345   GCCAAAAAACTTGGTATGACTACTCACGTTGGGAACGTTTTGTCATCTGATGTCTTTTACTCAAATTACTTTGAAAAGAATATCGAGCTT
NZ_CP035258.1:750874-758722    6345   GCCAAAAAACTTGGTATGACTACTCACGTTGGGAACGTTTTGTCATCTGATGTCTTTTACTCAAATTACTTTGAAAAGAATATCGAGCTT
NZ_CP035260.1:726156-733994    6335   GCCAAAGAACTTGGTATGACTACTCACGTTGGGAACGTTTTGTCATCTGATGTCTTTTACTCAAATTACTTTGAAAAGAATATCGAGCTT
NZ_CP035241.1:708296-716144    6345   GCCAAAGAACTTGGTATGACTACTCACGTTGGGAACGTTTTGTCATCTGATGTCTTTTACTCAAATTACTTTGAAAAGAATATCGAGCTT
NZ_CP035261.1:779086-786938    6349   GCCAAAGAACTTGGTATGACTACTCACGTTGGGAACGTTTTGTCATCTGATGTCTTTTACTCAAATTACTTTGAAAAGAATATCGAGCTT
NZ_CP035242.1:c1532236-1524385 6348   GCCAAAGAACTTGGTATGACTACTCACGTTGGGAACGTTTTGTCATCTGATGTCTTTTACTCAAATTACTTTGAAAAGAATATCGAGCTT
NZ_CP035243.1:777729-785580    6348   GCCAAAGAACTTGGTATGACTACTCACGTTGGGAACGTTTTGTCATCTGATGTCTTTTACTCAAATTACTTTGAAAAGAATATCGAGCTT
NZ_CP035237.1:c1311322-1303456 6363   GCCAAAAAACTTGGTATGACTACTCACGTTGGGAACGTTTTGTCATCTGATGTCTTTTACTCAAATTACTTTGAAAAGAATATCGAGCTT
NZ_CP035246.1:c1302892-1295026 6363   GCCAAAGAACTTGGTATGACTACTCACGTTGGGAACGTTTTGTCATCTGATGTCTTTTACTCAAATTACTTTGAAAAGAATATCGAGCTT
NZ_CP035247.1:c1301115-1293263 6349   GCCAAAGAACTTGGTATGACTACTCACGTTGGGAACGTTTTGTCATCTGATGTCTTTTACTCAAATTACTTTGAAAAGAATATCGAGCTT
NZ_CP035238.1:738547-746413    6363   GCCAAAGAACTTGGTATGACTACTCACGTTGGGAACGTTTTGTCATCTGATGTCTTTTACTCAAATTACTTTGAAAAGAATATCGAGCTT
NZ_CP035255.1:776866-784718    6349   GCCAAAGAACTTGGTATGACTACTCACGTTGGGAACGTTTTGTCATCTGATGTCTTTTACTCAAATTACTTTGAAAAGAATATCGAGCTT
```

### Ozkan et al. Supplementary Figures

```
NZ_CP035256.1:807236-815102    6363   GCCAAAGAACTTGGTATGACTACTCACGTTGGGAACGTTTTGTCATCTGATGTCTTTTACTCAAATTACTTTGAAAAGAATATCGAGCTT
NZ_CP038252.1:1151812-1159673  6288   GCCAAAGAACTTGGTATGACTACTCACGTTGGGAACGTTTTGTCATCTGATGTCTTTTACTCAAATTACTTTGAAAAGAATATCGAGCTT
NZ_CP035254.1:793365-801213    6345   GCCAAAGAACTTGGTATGACTACTCACGTTGGGAACGTTTTGTCATCTGATGTCTTTTACTCAAATTACTTTGAAAAGAATATCGAGCTT
NZ_CP035251.1:748733-756585    6349   GCCAAAAAATTTGGTATGACTACTCACGTTGGGAACGTTTTGTCATCTGATGTCTTTTACTCAAATTACTTTGAAAAGAATATCGAGCTT
NZ_CP035252.1:745670-753536    6363   GCCAAAAAATTTGGTATGACTACTCACGTTGGGAACGTTTTGTCATCTGATGTCTTTTACTCAAATTACTTTGAAAAGAATATCGAGCTT
NZ_CP035257.1:776317-784169    6349   GCCAAAAAATTTGGTATGACTACTCACGTTGGGAACGTTTTGTCATCTGATGTCTTTTACTCAAATTACTTTGAAAAGAATATCGAGCTT
NZ_CP038253.1:758105-765957    6349   GCCAAAAAATTTGGTATGACTACTCACGTTGGGAACGTTTTGTCATCTGATGTCTTTTACTCAAATTACTTTGAAAAGAATATCGAGCTT
NZ_CP035263.1:743050-750901    6348   GCCAAAAAACTTGGTATGACTACTCACGTTGGGAACGTTTTGTCATCTGATGTCTTTTACTCAAATTACTTTGAAAAGAATATCGAGCTT
NZ_CP035262.1:741595-749460    6362   GCCAAAAAACTTGGTATGACTACTCACGTTGGGAACGTTTTGTCATCTGATGTCTTTTACTCAAATTACTTTGAAAAGAATATCGAGCTT
NZ_CP035236.1:764915-772766    6348   GCCAAAAAACTTGGTATGACTACTCACGTTGGGAACGTTTTGTCATCTGATGTCTTTTACTCAAATTACTTTGAAAAGAATATCGAGCTT
NZ_CP035253.1 767286-775134    6345   GCCAAAGAACTTGGTATGACTACTCACGTTGGGAACGTTTTGTCATCTGATGTCTTTTACTCAAATTACTTTGAAAAGAATATCGAGCTT
NZ_CP035249.1:763080-770921    6338   GCCAAAAAACTTGGTATGACTACTCACGTTGGGAACGTTTTGTCATCTGATGTCTTTTACTCAAATTACTTTGAAAAGAATATCGAGCTT

NC_003028.3:777000-787000      7164   GGTAAATGGGGAGTCAAGGCTGTGGAAATGGAAGCAGCAGCTCTTTACTATCTTGCTGCCCAATACCATGTTGATGCGCTAGCTATCATG
NZ_CP035248.1:765099-775111    7177   GGTAAATGGGGAGTCAAGGCTGTGGAAATGGAAGCAGCAGCTCTTTACTATCTTGCTGCCCAATACCATGTTGATGCGCTAGCTATCATG
NZ_CP035235.1 c1326834-1317894 6452   GGTAAATGGGGAGTCAAGGCTGTGGAAATGGAAGCAGCAGCTCTTTACTATCTTGCTGCCCAATACCATGTTGATGCGCTAGCTATCATG
NZ_CP035234.1:756284-765224    6452   GGTAAATGGGGAGTCAAGGCTGTGGAAATGGAAGCAGCAGCTCTTTACTATCTTGCTGCCCAATACCATGTTGATGCGCTAGCTATCATG
NZ_CP038251.1:750745-759686    6452   GGTAAATGGGGAGTCAAGGCTGTGGAAATGGAAGCAGCAGCTCTTTACTATCTTGCTGCCCAATACCATGTTGATGCGCTAGCTATCATG
NZ_CP035244.1:747023-755950    6438   GGTAAATGGGGAGTCAAGGCTGTGGAAATGGAAGCAGCAGCTCTTTACTATCTTGCTGCCCAATACCATGTTGATGCGCTAGCTATCATG
NZ_CP035264.1:800836-809336    7087   GGTAAATGGGGAGTCAAGGCTGTGGAAATGGAAGCAGCAGCTCTTTACTATCTTGCTGCCCAATACCATGTTGATGCGCTAGCTATCATG
NZ_CP035265.1:766128-774627    7086   GGTAAATGGGGAGTCAAGGCTGTGGAAATGGAAGCAGCAGCTCTTTACTATCTTGCTGCCCAATACCATGTTGATGCGCTAGCTATCATG
NZ_CP035245.1 745261-753757    7083   GGTAAATGGGGAGTCAAGGCTGTGGAAATGGAAGCAGCAGCTCTTTACTATCTTGCTGCCCAATACCATGTTGATGCGCTAGCTATCATG
NZ_CP035259.1:745700-753548    6435   GGTAAATGGGGAGTCAAGGCTGTGGAAATGGAAGCAGCAGCTCTTTACTATCTTGCTGCCCAATACCATGTTGATGCGCTAGCTATCATG
NZ_CP035258.1:750874-758722    6435   GGTAAATGGGGAGTCAAGGCTGTGGAAATGGAAGCAGCAGCTCTTTACTATCTTGCTGCCCAATACCATGTTGATGCGCTAGCTATCATG
NZ_CP035260.1:726156-733994    6425   GGTAAATGGGGAGTCAAGGCTGTGGAAATGGAAGCAGCAGCTCTTTACTATCTTGCTGCCCAATACCATGTTGATGCGCTAGCTATCATG
NZ_CP035241.1:708296-716144    6435   GGTAAATGGGGAGTCAAGGCTGTGGAAATGGAAGCAGCAGCTCTTTACTATCTTGCTGCCCAATACCATGTTGATGCGCTAGCTATCATG
NZ_CP035261.1:779086-786938    6439   GGTAAATGGGGAGTCAAGGCTGTGGAAATGGAAGCAGCAGCTCTTTACTATCTTGCTGCCCAATACCATGTTGATGCGCTAGCTATCATG
NZ_CP035242.1:c1532236-1524385 6438   GGTAAATGGGGAGTCAAGGCTGTGGAAATGGAAGCAGCAGCTCTTTACTATCTTGCTGCCCAATACCATGTTGATGCGCTAGCTATCATG
NZ_CP035243.1:777729-785580    6438   GGTAAATGGGGAGTCAAGGCTGTGGAAATGGAAGCAGCAGCTCTTTACTATCTTGCTGCCCAATACCATGTTGATGCGCTAGCTATCATG
NZ_CP035237.1:c1311322-1303456 6453   GGTAAATGGGGAGTCAAGGCTGTGGAAATGGAAGCAGCAGCTCTTTACTATCTTGCTGCCCAATACCATGTTGATGCGCTAGCTATCATG
NZ_CP035246.1:c1302892-1295026 6453   GGTAAATGGGGAGTCAAGGCTGTGGAAATGGAAGCAGCAGCTCTTTACTATCTTGCTGCCCAATACCATGTTGATGCGCTAGCTATCATG
NZ_CP035247.1:c1301115-1293263 6439   GGTAAATGGGGAGTCAAGGCTGTGGAAATGGAAGCAGCAGCTCTTTACTATCTTGCTGCCCAATACCATGTTGATGCGCTAGCTATCATG
NZ_CP035238.1:738547-746413    6453   GGTAAATGGGGAGTCAAGGCTGTGGAAATGGAAGCAGCAGCTCTTTACTATCTTGCTGCCCAATACCATGTTGATGCGCTAGCTATCATG
NZ_CP035255.1:776866-784718    6439   GGTAAATGGGGAGTCAAGGCTGTGGAAATGGAAGCAGCAGCTCTTTACTATCTTGCTGCCCAATACCATGTTGATGCGCTAGCTATCATG
NZ_CP035256.1:807236-815102    6453   GGTAAATGGGGAGTCAAGGCTGTGGAAATGGAAGCAGCAGCTCTTTACTATCTTGCTGCCCAATACCATGTTGATGCGCTAGCTATCATG
NZ_CP038252.1:1151812-1159673  6378   GGTAAATGGGGAGTCAAGGCTGTGGAAATGGAAGCAGCAGCTCTTTACTATCTTGCTGCCCAATACCATGTTGATGCGCTAGCTATCATG
NZ_CP035254.1:793365-801213    6435   GGTAAATGGGGAGTCAAGGCTGTGGAAATGGAAGCAGCAGCTCTTTACTATCTTGCTGCCCAATACCATGTTGATGCGCTAGCTATCATG
NZ_CP035251.1:748733-756585    6439   GGTAAATGGGGAGTCAAGGCTGTGGAAATGGAAGCAGCAGCTCTTTACTATCTTGCTGCCCAATACCATGTTGATGCGCTAGCTATCATG
NZ_CP035252.1:745670-753536    6453   GGTAAATGGGGAGTCAAGGCTGTGGAAATGGAAGCAGCAGCTCTTTACTATCTTGCTGCCCAATACCATGTTGATGCGCTAGCTATCATG
NZ_CP035257.1:776317-784169    6439   GGTAAATGGGGAGTCAAGGCTGTGGAAATGGAAGCAGCAGCTCTTTACTATCTTGCTGCCCAATACCATGTTGATGCGCTAGCTATCATG
NZ_CP038253.1:758105-765957    6439   GGTAAATGGGGAGTCAAGGCTGTGGAAATGGAAGCAGCAGCTCTTTACTATCTTGCTGCCCAATACCATGTTGATGCGCTAGCTATCATG
NZ_CP035263.1:743050-750901    6438   GGTAAATGGGGAGTCAAGGCTGTGGAAATGGAAGCAGCAGCTCTTTACTATCTTGCTGCCCAATACCATGTTGATGCGCTAGCTATCATG
NZ_CP035262.1:741595-749460    6452   GGTAAATGGGGAGTCAAGGCTGTGGAAATGGAAGCAGCAGCTCTTTACTATCTTGCTGCCCAATACCATGTTGATGCGCTAGCTATCATG
NZ_CP035236.1:764915-772766    6438   GGTAAATGGGGAGTCAAGGCTGTGGAAATGGAAGCAGCAGCTCTTTACTATCTTGCTGCCCAATACCATGTTGATGCGCTAGCTATCATG
NZ_CP035253.1 767286-775134    6435   GGTAAATGGGGAGTCAAGGCTGTGGAAATGGAAGCAGCAGCTCTTTACTATCTTGCTGCCCAATACCATGTTGATGCGCTAGCTATCATG
NZ_CP035249.1:763080-770921    6428   GGTAAATGGGGAGTCAAGGCTGTGGAAATGGAAGCAGCAGCTCTTTACTATCTTGCTGCCCAATACCATGTTGATGCGCTAGCTATCATG

NC_003028.3:777000-787000      7254   ACCATCTCTGATAGCTTGGTCAATCCAGACGAAGACACAACTGCAGAAGAACGTCAAAATACCTTCACTGATATGATGAAGGTTGGTTTG
NZ_CP035248.1:765099-775111    7267   ACCATCTCTGATAGCTTGGTCAATCCAGACGAAGACACAACTGCAGAAGAACGTCAAAATACCTTCACTGATATGATGAAGGTTGGTTTG
NZ_CP035235.1 c1326834-1317894 6542   ACCATCTCTGATAGCTTGGTCAATCCAGACGAAGACACAACTGCAGAAGAACGTCAAAATACCTTCACTGATATGATGAAGGTTGGTTTG
NZ_CP035234.1:756284-765224    6542   ACCATCTCTGATAGCTTGGTCAATCCAGACGAAGACACAACTGCAGAAGAACGTCAAAATACCTTCACTGATATGATGAAGGTTGGTTTG
NZ_CP038251.1:750745-759686    6542   ACCATCTCTGATAGCTTGGTCAATCCAGACGAAGACACAACTGCAGAAGAACGTCAAAATACCTTCACTGATATGATGAAGGTTGGTTTG
NZ_CP035244.1:747023-755950    6528   ACCATCTCTGATAGCTTGGTCAATCCAGACGAAGACACAACTGCAGAAGAACGTCAAAATACCTTCACTGATATGATGAAGGTTGGTTTG
NZ_CP035264.1:800836-809336    7177   ACCATCTCTGATAGCTTGGTCAATCCAGACGAAGACACAACTGCAGAAGAACGTCAAAATACCTTCACTGATATGATGAAGGTTGGTTTG
NZ_CP035265.1:766128-774627    7176   ACCATCTCTGATAGCTTGGTCAATCCAGACGAAGACACAACTGCAGAAGAACGTCAAAATACCTTCACTGATATGATGAAGGTTGGTTTG
NZ_CP035245.1 745261-753757    7173   ACCATCTCTGATAGCTTGGTCAATCCAGACGAAGACACAACTGCAGAAGAACGTCAAAATACCTTCACTGATATGATGAAGGTTGGTTTG
NZ_CP035259.1:745700-753548    6525   ACCATCTCTGATAGCTTGGTCAATCCAGACGAAGACACAACTGCAGAAGAACGTCAAAATACCTTCACTGATATGATGAAGGTTGGTTTG
NZ_CP035258.1:750874-758722    6525   ACCATCTCTGATAGCTTGGTCAATCCAGACGAAGACACAACTGCAGAAGAACGTCAAAATACCTTCACTGATATGATGAAGGTTGGTTTG
NZ_CP035260.1:726156-733994    6515   ACCATCTCTGATAGCTTGGTCAATCCAGACGAAGACACAACTGCAGAAGAACGTCAAAATACCTTCACTGATATGATGAAGGTTGGTTTG
NZ_CP035241.1:708296-716144    6525   ACCATCTCTGATAGCTTGGTCAATCCAGACGAAGACACAACTGCAGAAGAACGTCAAAATACCTTCACTGATATGATGAAGGTTGGTTTG
NZ_CP035261.1:779086-786938    6529   ACCATCTCTGATAGCTTGGTCAATCCAGACGAAGACACAACTGCAGAAGAACGTCAAAATACCTTCACTGATATGATGAAGGTTGGTTTG
NZ_CP035242.1:c1532236-1524385 6528   ACCATCTCTGATAGCTTGGTCAATCCAGACGAAGACACAACTGCAGAAGAACGTCAAAATACCTTCACTGATATGATGAAGGTTGGTTTG
NZ_CP035243.1:777729-785580    6528   ACCATCTCTGATAGCTTGGTCAATCCAGACGAAGACACAACTGCAGAAGAACGTCAAAATACCTTCACTGATATGATGAAGGTTGGTTTG
NZ_CP035237.1:c1311322-1303456 6543   ACCATCTCTGATAGCTTGGTCAATCCAGACGAAGACACAACTGCAGAAGAACGTCAAAATACCTTCACTGATATGATGAAGGTTGGTTTG
NZ_CP035246.1:c1302892-1295026 6543   ACCATCTCTGATAGCTTGGTCAATCCAGACGAAGACACAACTGCAGAAGAACGTCAAAATACCTTCACTGATATGATGAAGGTTGGTTTG
NZ_CP035247.1:c1301115-1293263 6529   ACCATCTCTGATAGCTTGGTCAATCCAGACGAAGACACAACTGCAGAAGAACGTCAAAATACCTTCACTGATATGATGAAGGTTGGTTTG
NZ_CP035238.1:738547-746413    6543   ACCATCTCTGATAGCTTGGTCAATCCAGACGAAGACACAACTGCAGAAGAACGTCAAAATACCTTCACTGATATGATGAAGGTTGGTTTG
NZ_CP035255.1:776866-784718    6529   ACCATCTCTGATAGCTTGGTCAATCCAGACGAAGACACAACTGCAGAAGAACGTCAAAATACCTTCACTGATATGATGAAGGTTGGTTTG
NZ_CP035256.1:807236-815102    6543   ACCATCTCTGATAGCTTGGTCAATCCAGACGAAGACACAACTGCAGAAGAACGTCAAAATACCTTCACTGATATGATGAAGGTTGGTTTG
NZ_CP038252.1:1151812-1159673  6468   ACCATCTCTGATAGCTTGGTCAATCCAGACGAAGACACAACTGCAGAAGAACGTCAAAATACCTTCACTGATATGATGAAGGTTGGTTTG
NZ_CP035254.1:793365-801213    6525   ACCATCTCTGATAGCTTGGTCAATCCAGACGAAGACACAACTGCAGAAGAACGTCAAAATACCTTCACTGATATGATGAAGGTTGGTTTG
NZ_CP035251.1:748733-756585    6529   ACCATCTCTGATAGCTTGGTCAATCCAGACGAAGACACAACTGCAGAAGAACGTCAAAATACCTTCACTGATATGATGAAGGTTGGTTTG
NZ_CP035252.1:745670-753536    6543   ACCATCTCTGATAGCTTGGTCAATCCAGACGAAGACACAACTGCAGAAGAACGTCAAAATACCTTCACTGATATGATGAAGGTTGGTTTG
NZ_CP035257.1:776317-784169    6529   ACCATCTCTGATAGCTTGGTCAATCCAGACGAAGACACAACTGCAGAAGAACGTCAAAATACCTTCACTGATATGATGAAGGTTGGTTTG
NZ_CP038253.1:758105-765957    6529   ACCATCTCTGATAGCTTGGTCAATCCAGACGAAGACACAACTGCAGAAGAACGTCAAAATACCTTCACTGATATGATGAAGGTTGGTTTG
NZ_CP035263.1:743050-750901    6528   ACCATCTCTGATAGCTTGGTCAATCCAGACGAAGACACAACTGCAGAAGAACGTCAAAATACCTTCACTGATATGATGAAGGTTGGTTTG
NZ_CP035262.1:741595-749460    6542   ACCATCTCTGATAGCTTGGTCAATCCAGACGAAGACACAACTGCAGAAGAACGTCAAAATACCTTCACTGATATGATGAAGGTTGGTTTG
NZ_CP035236.1:764915-772766    6528   ACCATCTCTGATAGCTTGGTCAATCCAGACGAAGACACAACTGCAGAAGAACGTCAAAATACCTTCACTGATATGATGAAGGTTGGTTTG
NZ_CP035253.1 767286-775134    6525   ACCATCTCTGATAGCTTGGTCAATCCAGACGAAGACACAACTGCAGAAGAACGTCAAAATACCTTCACTGATATGATGAAGGTTGGTTTG
NZ_CP035249.1:763080-770921    6518   ACCATCTCTGATAGCTTGGTCAATCCAGACGAAGACACAACTGCAGAAGAACGTCAAAATACCTTCACTGATATGATGAAGGTTGGTTTG

NC_003028.3:777000-787000      7344   GAAACCTTGATTGCAGAATAATTATAGCCAAAAAGGGGCTCTTTGTCAACTGTAGTGGGTTGAAAAAAAGCTAAGCTTGAGAAAGGACAA
NZ_CP035248.1:765099-775111    7357   GAAACCTTGATTGCAGAATAATTATAGCCAAAAAGGGGCTCTTTGTCAACTGTAGTGGGTTGAAAAAAAGCTAAGCTTGAGAAAGGACAA
NZ_CP035235.1 c1326834-1317894 6632   GAAACCTTGATTGCAGAATAATTATAGCCAAAAAGG------------------------------------------------------
NZ_CP035234.1:756284-765224    6632   GAAACCTTGATTGCAGAATAATTATAGCCAAAAAGG------------------------------------------------------
NZ_CP038251.1:750745-759686    6632   GAAACCTTGATTGCAGAATAATTATAGCCAAAAAGG------------------------------------------------------
NZ_CP035244.1:747023-755950    6618   GAAACCTTGATTGCAGAATAATTATAGCCAAAAAGG------------------------------------------------------
NZ_CP035264.1:800836-809336    7267   GAAACCTTGATTGCAGAATAATTATAGCCAAAAAGG------------------------------------------------------
NZ_CP035265.1:766128-774627    7266   GAAACCTTGATTGCAGAATAATTATAGCCAAAAAGG------------------------------------------------------
NZ_CP035245.1 745261-753757    7263   GAAACCTTGATTGCAGAATAATTATAGCCAAAAAGG------------------------------------------------------
NZ_CP035259.1:745700-753548    6615   GAAACCTTGATTGCAGAATAATTATAGCCAAAAAGG------------------------------------------------------
NZ_CP035258.1:750874-758722    6615   GAAACCTTGATTGCAGAATAATTATAGCCAAAAAGG------------------------------------------------------
NZ_CP035260.1:726156-733994    6605   GAAACCTTGATTGCAGAATAATTATAGCCAAAAAGG------------------------------------------------------
NZ_CP035241.1:708296-716144    6615   GAAACCTTGATTGCAGAATAATTATAGCCAAAAAGG------------------------------------------------------
NZ_CP035261.1:779086-786938    6619   GAAACCTTGATTGCAGAATAATTATAGCCAAAAAGG------------------------------------------------------
```

```
NZ_CP035242.1:c1532236-1524385 6618    GAAACCTTGATTGCAGAATAATTATAGCCAAAAAGG------------------------------------------------------
NZ_CP035243.1:777729-785580    6618    GAAACCTTGATTGCAGAATAATTATAGCCAAAAAGG------------------------------------------------------
NZ_CP035237.1:c1311322-1303456 6633    GAAACCTTGATTGCAGAATAATTATAGCCAAAAAGG------------------------------------------------------
NZ_CP035246.1:c1302892-1295026 6633    GAAACCTTGATTGCAGAATAATTATAGCCAAAAAGG------------------------------------------------------
NZ_CP035247.1:c1301115-1293263 6619    GAAACCTTGATTGCAGAATAATTATAGCCAAAAAGG------------------------------------------------------
NZ_CP035238.1:738547-746413    6633    GAAACCTTGATTGCAGAATAATTATAGCCAAAAAGG------------------------------------------------------
NZ_CP035255.1:776866-784718    6619    GAAACCTTGATTGCAGAATAATTATAGCCAAAAAGG------------------------------------------------------
NZ_CP035256.1:807236-815102    6633    GAAACCTTGATTGCAGAATAATTATAGCCAAAAAGG------------------------------------------------------
NZ_CP038252.1:1151812-1159673  6558    GAAACCTTGATTGCAGAATAATTATAGCCAAAAAGG------------------------------------------------------
NZ_CP035254.1:793365-801213    6615    GAAACCTTGATTGCAGAATAATTATAGCCAAAAAGG------------------------------------------------------
NZ_CP035251.1:748733-756585    6619    GAAACCTTGATTGCAGAATAATTATAGCCAAAAAGG------------------------------------------------------
NZ_CP035252.1:745670-753536    6633    GAAACCTTGATTGCAGAATAATTATAGCCAAAAAGG------------------------------------------------------
NZ_CP035257.1:776317-784169    6619    GAAACCTTGATTGCAGAATAATTATAGCCAAAAAGG------------------------------------------------------
NZ_CP038253.1:758105-765957    6619    GAAACCTTGATTGCAGAATAATTATAGCCAAAAAGG------------------------------------------------------
NZ_CP035263.1:743050-750901    6618    GAAACCTTGATTGCAGAATAATTATAGCCAAAAAGG------------------------------------------------------
NZ_CP035262.1:741595-749460    6632    GAAACCTTGATTGCAGAATAATTATAGCCAAAAAGG------------------------------------------------------
NZ_CP035236.1:764915-772766    6618    GAAACCTTGATTGCAGAATAATTATAGCCAAAAAGG------------------------------------------------------
NZ_CP035253.1 767286-775134    6615    GAAACCTTGATTGCAGAATAATTATAGCCAAAAAGG------------------------------------------------------
NZ_CP035249.1:763080-770921    6608    GAAACCTTGATTGCAGAATAATTATAGCCAAAAAGG------------------------------------------------------

NC_003028.3:777000-787000      7434    ATTTCGTCCTTTCTTTTTTGATATTCAGGGCGATAAAAATCCGTTTTTTGAAGTTTTCAAAGTTCCGAAAACCAAAGGCATTGCGCTTGA
NZ_CP035248.1:765099-775111    7447    ATTTCGTCCTTTCTTTTTTGATATTCAGGGCGATAAAAATCCGTTTTTTGAAGTTTTCAAAGTTCCGAAAACCAAAGGCATTGCGCTTGA
NZ_CP035235.1 c1326834-1317894 6667    ------------------------------------------------------------------------------------------
NZ_CP035234.1:756284-765224    6667    ------------------------------------------------------------------------------------------
NZ_CP038251.1:750745-759686    6667    ------------------------------------------------------------------------------------------
NZ_CP035244.1:747023-755950    6653    ------------------------------------------------------------------------------------------
NZ_CP035264.1:800836-809336    7302    ------------------------------------------------------------------------------------------
NZ_CP035265.1:766128-774627    7301    ------------------------------------------------------------------------------------------
NZ_CP035245.1 745261-753757    7298    ------------------------------------------------------------------------------------------
NZ_CP035259.1:745700-753548    6650    ------------------------------------------------------------------------------------------
NZ_CP035258.1:750874-758722    6650    ------------------------------------------------------------------------------------------
NZ_CP035260.1:726156-733994    6640    ------------------------------------------------------------------------------------------
NZ_CP035241.1:708296-716144    6650    ------------------------------------------------------------------------------------------
NZ_CP035261.1:779086-786938    6654    ------------------------------------------------------------------------------------------
NZ_CP035242.1:c1532236-1524385 6653    ------------------------------------------------------------------------------------------
NZ_CP035243.1:777729-785580    6653    ------------------------------------------------------------------------------------------
NZ_CP035237.1:c1311322-1303456 6668    ------------------------------------------------------------------------------------------
NZ_CP035246.1:c1302892-1295026 6668    ------------------------------------------------------------------------------------------
NZ_CP035247.1:c1301115-1293263 6654    ------------------------------------------------------------------------------------------
NZ_CP035238.1:738547-746413    6668    ------------------------------------------------------------------------------------------
NZ_CP035255.1:776866-784718    6654    ------------------------------------------------------------------------------------------
NZ_CP035256.1:807236-815102    6668    ------------------------------------------------------------------------------------------
NZ_CP038252.1:1151812-1159673  6593    ------------------------------------------------------------------------------------------
NZ_CP035254.1:793365-801213    6650    ------------------------------------------------------------------------------------------
NZ_CP035251.1:748733-756585    6654    ------------------------------------------------------------------------------------------
NZ_CP035252.1:745670-753536    6668    ------------------------------------------------------------------------------------------
NZ_CP035257.1:776317-784169    6654    ------------------------------------------------------------------------------------------
NZ_CP038253.1:758105-765957    6654    ------------------------------------------------------------------------------------------
NZ_CP035263.1:743050-750901    6653    ------------------------------------------------------------------------------------------
NZ_CP035262.1:741595-749460    6667    ------------------------------------------------------------------------------------------
NZ_CP035236.1:764915-772766    6653    ------------------------------------------------------------------------------------------
NZ_CP035253.1 767286-775134    6650    ------------------------------------------------------------------------------------------
NZ_CP035249.1:763080-770921    6643    ------------------------------------------------------------------------------------------

NC_003028.3:777000-787000      7524    TAAGTTTGATGAGATTATTGGTCGCTTCCAGTTTGGCATTAGAATAGTGTAGTTGAAGGGCGTTGACGATTTTCTCTTTGTTCTTTAGAA
NZ_CP035248.1:765099-775111    7537    TAAGTTTGATGAGATTATTGGTCGCTTCCAGTTTGGCATTAGAATAGTGTAGTTGAAGGGCGTTGACGATTTTCTCTTTGTTCTTTAGAA
NZ_CP035235.1 c1326834-1317894 6667    ------------------------------------------------------------------------------------------
NZ_CP035234.1:756284-765224    6667    ------------------------------------------------------------------------------------------
NZ_CP038251.1:750745-759686    6667    ------------------------------------------------------------------------------------------
NZ_CP035244.1:747023-755950    6653    ------------------------------------------------------------------------------------------
NZ_CP035264.1:800836-809336    7302    ------------------------------------------------------------------------------------------
NZ_CP035265.1:766128-774627    7301    ------------------------------------------------------------------------------------------
NZ_CP035245.1 745261-753757    7298    ------------------------------------------------------------------------------------------
NZ_CP035259.1:745700-753548    6650    ------------------------------------------------------------------------------------------
NZ_CP035258.1:750874-758722    6650    ------------------------------------------------------------------------------------------
NZ_CP035260.1:726156-733994    6640    ------------------------------------------------------------------------------------------
NZ_CP035241.1:708296-716144    6650    ------------------------------------------------------------------------------------------
NZ_CP035261.1:779086-786938    6654    ------------------------------------------------------------------------------------------
NZ_CP035242.1:c1532236-1524385 6653    ------------------------------------------------------------------------------------------
NZ_CP035243.1:777729-785580    6653    ------------------------------------------------------------------------------------------
NZ_CP035237.1:c1311322-1303456 6668    ------------------------------------------------------------------------------------------
NZ_CP035246.1:c1302892-1295026 6668    ------------------------------------------------------------------------------------------
NZ_CP035247.1:c1301115-1293263 6654    ------------------------------------------------------------------------------------------
NZ_CP035238.1:738547-746413    6668    ------------------------------------------------------------------------------------------
NZ_CP035255.1:776866-784718    6654    ------------------------------------------------------------------------------------------
NZ_CP035256.1:807236-815102    6668    ------------------------------------------------------------------------------------------
NZ_CP038252.1:1151812-1159673  6593    ------------------------------------------------------------------------------------------
NZ_CP035254.1:793365-801213    6650    ------------------------------------------------------------------------------------------
NZ_CP035251.1:748733-756585    6654    ------------------------------------------------------------------------------------------
NZ_CP035252.1:745670-753536    6668    ------------------------------------------------------------------------------------------
NZ_CP035257.1:776317-784169    6654    ------------------------------------------------------------------------------------------
NZ_CP038253.1:758105-765957    6654    ------------------------------------------------------------------------------------------
NZ_CP035263.1:743050-750901    6653    ------------------------------------------------------------------------------------------
NZ_CP035262.1:741595-749460    6667    ------------------------------------------------------------------------------------------
NZ_CP035236.1:764915-772766    6653    ------------------------------------------------------------------------------------------
NZ_CP035253.1 767286-775134    6650    ------------------------------------------------------------------------------------------
NZ_CP035249.1:763080-770921    6643    ------------------------------------------------------------------------------------------

NC_003028.3:777000-787000      7614    AGGTTTTAAAGACAGTCTGAAAAAGAGGATGAACCTGCTTCAGATTGTCCTCAATGAGTCCGAAAAATTTCTCAGGGTCTTTGTTCTGAA
NZ_CP035248.1:765099-775111    7627    AGGTTTTAAAGACAGTCTGAAAAAGAGGATGAACCTGCTTCAGATTGTCCTCAATGAGTCCGAAAAATTTCTCAGGGTCTTTGTTCTGAA
NZ_CP035235.1 c1326834-1317894 6667    ------------------------------------------------------------------------------------------
NZ_CP035234.1:756284-765224    6667    ------------------------------------------------------------------------------------------
NZ_CP038251.1:750745-759686    6667    ------------------------------------------------------------------------------------------
NZ_CP035244.1:747023-755950    6653    ------------------------------------------------------------------------------------------
NZ_CP035264.1:800836-809336    7302    ------------------------------------------------------------------------------------------
```

```
NZ_CP035265.1:766128-774627    7301   ------------------------------------------------------------------------------------------
NZ_CP035245.1 745261-753757    7298   ------------------------------------------------------------------------------------------
NZ_CP035259.1:745700-753548    6650   ------------------------------------------------------------------------------------------
NZ_CP035258.1:750874-758722    6650   ------------------------------------------------------------------------------------------
NZ_CP035260.1:726156-733994    6640   ------------------------------------------------------------------------------------------
NZ_CP035241.1:708296-716144    6650   ------------------------------------------------------------------------------------------
NZ_CP035261.1:779086-786938    6654   ------------------------------------------------------------------------------------------
NZ_CP035242.1:c1532236-1524385 6653   ------------------------------------------------------------------------------------------
NZ_CP035243.1:777729-785580    6653   ------------------------------------------------------------------------------------------
NZ_CP035237.1:c1311322-1303456 6668   ------------------------------------------------------------------------------------------
NZ_CP035246.1:c1302892-1295026 6668   ------------------------------------------------------------------------------------------
NZ_CP035247.1:c1301115-1293263 6654   ------------------------------------------------------------------------------------------
NZ_CP035238.1:738547-746413    6668   ------------------------------------------------------------------------------------------
NZ_CP035255.1:776866-784718    6654   ------------------------------------------------------------------------------------------
NZ_CP035256.1:807236-815102    6668   ------------------------------------------------------------------------------------------
NZ_CP038252.1:1151812-1159673  6593   ------------------------------------------------------------------------------------------
NZ_CP035254.1:793365-801213    6650   ------------------------------------------------------------------------------------------
NZ_CP035251.1:748733-756585    6654   ------------------------------------------------------------------------------------------
NZ_CP035252.1:745670-753536    6668   ------------------------------------------------------------------------------------------
NZ_CP035257.1:776317-784169    6654   ------------------------------------------------------------------------------------------
NZ_CP038253.1:758105-765957    6654   ------------------------------------------------------------------------------------------
NZ_CP035263.1:743050-750901    6653   ------------------------------------------------------------------------------------------
NZ_CP035262.1:741595-749460    6667   ------------------------------------------------------------------------------------------
NZ_CP035236.1:764915-772766    6653   ------------------------------------------------------------------------------------------
NZ_CP035253.1 767286-775134    6650   ------------------------------------------------------------------------------------------
NZ_CP035249.1:763080-770921    6643   ------------------------------------------------------------------------------------------

NC_003028.3:777000-787000      7704   AGTGAAAAAGTAAGAGTTGATAGATCTGATAGTGGTGTTTCAAGTCTTCTGAATAGCTTAAAATCTTGTCAAGAATTTCTTTATTTGTTA
NZ_CP035248.1:765099-775111    7717   AGTGAAAAAGTAAGAGTTGATAGATCTGATAGTGGTGTTTCAAGTCTTCTGAATAGCTTAAAATCTTGTCAAGAATTTCTTTATTTGTTA
NZ_CP035235.1 c1326834-1317894 6667   ------------------------------------------------------------------------------------------
NZ_CP035234.1:756284-765224    6667   ------------------------------------------------------------------------------------------
NZ_CP038251.1:750745-759686    6667   ------------------------------------------------------------------------------------------
NZ_CP035244.1:747023-755950    6653   ------------------------------------------------------------------------------------------
NZ_CP035264.1:800836-809336    7302   ------------------------------------------------------------------------------------------
NZ_CP035265.1:766128-774627    7301   ------------------------------------------------------------------------------------------
NZ_CP035245.1 745261-753757    7298   ------------------------------------------------------------------------------------------
NZ_CP035259.1:745700-753548    6650   ------------------------------------------------------------------------------------------
NZ_CP035258.1:750874-758722    6650   ------------------------------------------------------------------------------------------
NZ_CP035260.1:726156-733994    6640   ------------------------------------------------------------------------------------------
NZ_CP035241.1:708296-716144    6650   ------------------------------------------------------------------------------------------
NZ_CP035261.1:779086-786938    6654   ------------------------------------------------------------------------------------------
NZ_CP035242.1:c1532236-1524385 6653   ------------------------------------------------------------------------------------------
NZ_CP035243.1:777729-785580    6653   ------------------------------------------------------------------------------------------
NZ_CP035237.1:c1311322-1303456 6668   ------------------------------------------------------------------------------------------
NZ_CP035246.1:c1302892-1295026 6668   ------------------------------------------------------------------------------------------
NZ_CP035247.1:c1301115-1293263 6654   ------------------------------------------------------------------------------------------
NZ_CP035238.1:738547-746413    6668   ------------------------------------------------------------------------------------------
NZ_CP035255.1:776866-784718    6654   ------------------------------------------------------------------------------------------
NZ_CP035256.1:807236-815102    6668   ------------------------------------------------------------------------------------------
NZ_CP038252.1:1151812-1159673  6593   ------------------------------------------------------------------------------------------
NZ_CP035254.1:793365-801213    6650   ------------------------------------------------------------------------------------------
NZ_CP035251.1:748733-756585    6654   ------------------------------------------------------------------------------------------
NZ_CP035252.1:745670-753536    6668   ------------------------------------------------------------------------------------------
NZ_CP035257.1:776317-784169    6654   ------------------------------------------------------------------------------------------
NZ_CP038253.1:758105-765957    6654   ------------------------------------------------------------------------------------------
NZ_CP035263.1:743050-750901    6653   ------------------------------------------------------------------------------------------
NZ_CP035262.1:741595-749460    6667   ------------------------------------------------------------------------------------------
NZ_CP035236.1:764915-772766    6653   ------------------------------------------------------------------------------------------
NZ_CP035253.1 767286-775134    6650   ------------------------------------------------------------------------------------------
NZ_CP035249.1:763080-770921    6643   ------------------------------------------------------------------------------------------

NC_003028.3:777000-787000      7794   AGTGCATGCGAAAAGTAGGGCGATAAAAACGTTTATCGCTCAATTTACGACTATCCTGTTGGATGAGTTTCCAGTAACGCTTGATAGCCT
NZ_CP035248.1:765099-775111    7807   AGTGCATGCGAAAAGTAGGGCGATAAAAACGTTTATCGCTCAATTTACGACTATCCTGTTGGATGAGTTTCCAGTAACGTTTGATAGCCT
NZ_CP035235.1 c1326834-1317894 6667   ------------------------------------------------------------------------------------------
NZ_CP035234.1:756284-765224    6667   ------------------------------------------------------------------------------------------
NZ_CP038251.1:750745-759686    6667   ------------------------------------------------------------------------------------------
NZ_CP035244.1:747023-755950    6653   ------------------------------------------------------------------------------------------
NZ_CP035264.1:800836-809336    7302   ------------------------------------------------------------------------------------------
NZ_CP035265.1:766128-774627    7301   ------------------------------------------------------------------------------------------
NZ_CP035245.1 745261-753757    7298   ------------------------------------------------------------------------------------------
NZ_CP035259.1:745700-753548    6650   ------------------------------------------------------------------------------------------
NZ_CP035258.1:750874-758722    6650   ------------------------------------------------------------------------------------------
NZ_CP035260.1:726156-733994    6640   ------------------------------------------------------------------------------------------
NZ_CP035241.1:708296-716144    6650   ------------------------------------------------------------------------------------------
NZ_CP035261.1:779086-786938    6654   ------------------------------------------------------------------------------------------
NZ_CP035242.1:c1532236-1524385 6653   ------------------------------------------------------------------------------------------
NZ_CP035243.1:777729-785580    6653   ------------------------------------------------------------------------------------------
NZ_CP035237.1:c1311322-1303456 6668   ------------------------------------------------------------------------------------------
NZ_CP035246.1:c1302892-1295026 6668   ------------------------------------------------------------------------------------------
NZ_CP035247.1:c1301115-1293263 6654   ------------------------------------------------------------------------------------------
NZ_CP035238.1:738547-746413    6668   ------------------------------------------------------------------------------------------
NZ_CP035255.1:776866-784718    6654   ------------------------------------------------------------------------------------------
NZ_CP035256.1:807236-815102    6668   ------------------------------------------------------------------------------------------
NZ_CP038252.1:1151812-1159673  6593   ------------------------------------------------------------------------------------------
NZ_CP035254.1:793365-801213    6650   ------------------------------------------------------------------------------------------
NZ_CP035251.1:748733-756585    6654   ------------------------------------------------------------------------------------------
NZ_CP035252.1:745670-753536    6668   ------------------------------------------------------------------------------------------
NZ_CP035257.1:776317-784169    6654   ------------------------------------------------------------------------------------------
NZ_CP038253.1:758105-765957    6654   ------------------------------------------------------------------------------------------
NZ_CP035263.1:743050-750901    6653   ------------------------------------------------------------------------------------------
NZ_CP035262.1:741595-749460    6667   ------------------------------------------------------------------------------------------
NZ_CP035236.1:764915-772766    6653   ------------------------------------------------------------------------------------------
NZ_CP035253.1 767286-775134    6650   ------------------------------------------------------------------------------------------
NZ_CP035249.1:763080-770921    6643   ------------------------------------------------------------------------------------------
```

### Ozkan et al. Supplementary Figures

```
NC_003028.3:777000-787000      7884   TGTATTCATGAGATTTTCGTTCAAACTGATTCATAATTTGAACACGAAAACGACTCATGGCACGGCTGAGATGTTGGATAATATGGAAAC
NZ_CP035248.1:765099-775111    7897   TGTATTCATGAGATTTTCGTTCAAACTGATTCATAATTTGAACACGAAAACGACTCATGGCACGGCTGAGATGTTGGATAATATGGAAAC
NZ_CP035235.1 c1326834-1317894 6667   ------------------------------------------------------------------------------------------
NZ_CP035234.1:756284-765224    6667   ------------------------------------------------------------------------------------------
NZ_CP038251.1:750745-759686    6667   ------------------------------------------------------------------------------------------
NZ_CP035244.1:747023-755950    6653   ------------------------------------------------------------------------------------------
NZ_CP035264.1:800836-809336    7302   ------------------------------------------------------------------------------------------
NZ_CP035265.1:766128-774627    7301   ------------------------------------------------------------------------------------------
NZ_CP035245.1 745261-753757    7298   ------------------------------------------------------------------------------------------
NZ_CP035259.1:745700-753548    6650   ------------------------------------------------------------------------------------------
NZ_CP035258.1:750874-758722    6650   ------------------------------------------------------------------------------------------
NZ_CP035260.1:726156-733994    6640   ------------------------------------------------------------------------------------------
NZ_CP035241.1:708296-716144    6650   ------------------------------------------------------------------------------------------
NZ_CP035261.1:779086-786938    6654   ------------------------------------------------------------------------------------------
NZ_CP035242.1:c1532236-1524385 6653   ------------------------------------------------------------------------------------------
NZ_CP035243.1:777729-785580    6653   ------------------------------------------------------------------------------------------
NZ_CP035237.1:c1311322-1303456 6668   ------------------------------------------------------------------------------------------
NZ_CP035246.1:c1302892-1295026 6668   ------------------------------------------------------------------------------------------
NZ_CP035247.1:c1301115-1293263 6654   ------------------------------------------------------------------------------------------
NZ_CP035238.1:738547-746413    6668   ------------------------------------------------------------------------------------------
NZ_CP035255.1:776866-784718    6654   ------------------------------------------------------------------------------------------
NZ_CP035256.1:807236-815102    6668   ------------------------------------------------------------------------------------------
NZ_CP038252.1:1151812-1159673  6593   ------------------------------------------------------------------------------------------
NZ_CP035254.1:793365-801213    6650   ------------------------------------------------------------------------------------------
NZ_CP035251.1:748733-756585    6654   ------------------------------------------------------------------------------------------
NZ_CP035252.1:745670-753536    6668   ------------------------------------------------------------------------------------------
NZ_CP035257.1:776317-784169    6654   ------------------------------------------------------------------------------------------
NZ_CP038253.1:758105-765957    6654   ------------------------------------------------------------------------------------------
NZ_CP035263.1:743050-750901    6653   ------------------------------------------------------------------------------------------
NZ_CP035262.1:741595-749460    6667   ------------------------------------------------------------------------------------------
NZ_CP035236.1:764915-772766    6653   ------------------------------------------------------------------------------------------
NZ_CP035253.1 767286-775134    6650   ------------------------------------------------------------------------------------------
NZ_CP035249.1:763080-770921    6643   ------------------------------------------------------------------------------------------

NC_003028.3:777000-787000      7974   GATCTAGAACGATTTTAGCACACGGAAAAAGCTGTTTAGCCAAGTCATAGTAAGGACTAAACATATCCATCGTAATGATTTTCACTTGAC
NZ_CP035248.1:765099-775111    7987   GATCTAGAACGATTTTAGCACACGGAAAAAGCTGTTTAGCCAAGTCATAGTAAGGACTAAACATATCCATCGTAATGATTTTCACTTGAC
NZ_CP035235.1 c1326834-1317894 6667   ------------------------------------------------------------------------------------------
NZ_CP035234.1:756284-765224    6667   ------------------------------------------------------------------------------------------
NZ_CP038251.1:750745-759686    6667   ------------------------------------------------------------------------------------------
NZ_CP035244.1:747023-755950    6653   ------------------------------------------------------------------------------------------
NZ_CP035264.1:800836-809336    7302   ------------------------------------------------------------------------------------------
NZ_CP035265.1:766128-774627    7301   ------------------------------------------------------------------------------------------
NZ_CP035245.1 745261-753757    7298   ------------------------------------------------------------------------------------------
NZ_CP035259.1:745700-753548    6650   ------------------------------------------------------------------------------------------
NZ_CP035258.1:750874-758722    6650   ------------------------------------------------------------------------------------------
NZ_CP035260.1:726156-733994    6640   ------------------------------------------------------------------------------------------
NZ_CP035241.1:708296-716144    6650   ------------------------------------------------------------------------------------------
NZ_CP035261.1:779086-786938    6654   ------------------------------------------------------------------------------------------
NZ_CP035242.1:c1532236-1524385 6653   ------------------------------------------------------------------------------------------
NZ_CP035243.1:777729-785580    6653   ------------------------------------------------------------------------------------------
NZ_CP035237.1:c1311322-1303456 6668   ------------------------------------------------------------------------------------------
NZ_CP035246.1:c1302892-1295026 6668   ------------------------------------------------------------------------------------------
NZ_CP035247.1:c1301115-1293263 6654   ------------------------------------------------------------------------------------------
NZ_CP035238.1:738547-746413    6668   ------------------------------------------------------------------------------------------
NZ_CP035255.1:776866-784718    6654   ------------------------------------------------------------------------------------------
NZ_CP035256.1:807236-815102    6668   ------------------------------------------------------------------------------------------
NZ_CP038252.1:1151812-1159673  6593   ------------------------------------------------------------------------------------------
NZ_CP035254.1:793365-801213    6650   ------------------------------------------------------------------------------------------
NZ_CP035251.1:748733-756585    6654   ------------------------------------------------------------------------------------------
NZ_CP035252.1:745670-753536    6668   ------------------------------------------------------------------------------------------
NZ_CP035257.1:776317-784169    6654   ------------------------------------------------------------------------------------------
NZ_CP038253.1:758105-765957    6654   ------------------------------------------------------------------------------------------
NZ_CP035263.1:743050-750901    6653   ------------------------------------------------------------------------------------------
NZ_CP035262.1:741595-749460    6667   ------------------------------------------------------------------------------------------
NZ_CP035236.1:764915-772766    6653   ------------------------------------------------------------------------------------------
NZ_CP035253.1 767286-775134    6650   ------------------------------------------------------------------------------------------
NZ_CP035249.1:763080-770921    6643   ------------------------------------------------------------------------------------------

NC_003028.3:777000-787000      8064   AACGAACGGCTCTATCGTAGCGAAGAAAGTGATTTCGGATGACAGCTTGTGTTCTGCCTTCAAGAACAGTGATAATATTAAGATTATCAA
NZ_CP035248.1:765099-775111    8077   AACGAACGGCTCTATCGTAGCGAAGAAAGTGATTTCGGATGACAGCTTGTGTTCTGCCTTCAAGAACAGTGATAATATTAAGATTATCAA
NZ_CP035235.1 c1326834-1317894 6667   ------------------------------------------------------------------------------------------
NZ_CP035234.1:756284-765224    6667   ------------------------------------------------------------------------------------------
NZ_CP038251.1:750745-759686    6667   ------------------------------------------------------------------------------------------
NZ_CP035244.1:747023-755950    6653   ------------------------------------------------------------------------------------------
NZ_CP035264.1:800836-809336    7302   ------------------------------------------------------------------------------------------
NZ_CP035265.1:766128-774627    7301   ------------------------------------------------------------------------------------------
NZ_CP035245.1 745261-753757    7298   ------------------------------------------------------------------------------------------
NZ_CP035259.1:745700-753548    6650   ------------------------------------------------------------------------------------------
NZ_CP035258.1:750874-758722    6650   ------------------------------------------------------------------------------------------
NZ_CP035260.1:726156-733994    6640   ------------------------------------------------------------------------------------------
NZ_CP035241.1:708296-716144    6650   ------------------------------------------------------------------------------------------
NZ_CP035261.1:779086-786938    6654   ------------------------------------------------------------------------------------------
NZ_CP035242.1:c1532236-1524385 6653   ------------------------------------------------------------------------------------------
NZ_CP035243.1:777729-785580    6653   ------------------------------------------------------------------------------------------
NZ_CP035237.1:c1311322-1303456 6668   ------------------------------------------------------------------------------------------
NZ_CP035246.1:c1302892-1295026 6668   ------------------------------------------------------------------------------------------
NZ_CP035247.1:c1301115-1293263 6654   ------------------------------------------------------------------------------------------
NZ_CP035238.1:738547-746413    6668   ------------------------------------------------------------------------------------------
NZ_CP035255.1:776866-784718    6654   ------------------------------------------------------------------------------------------
NZ_CP035256.1:807236-815102    6668   ------------------------------------------------------------------------------------------
NZ_CP038252.1:1151812-1159673  6593   ------------------------------------------------------------------------------------------
NZ_CP035254.1:793365-801213    6650   ------------------------------------------------------------------------------------------
NZ_CP035251.1:748733-756585    6654   ------------------------------------------------------------------------------------------
NZ_CP035252.1:745670-753536    6668   ------------------------------------------------------------------------------------------
NZ_CP035257.1:776317-784169    6654   ------------------------------------------------------------------------------------------
```

```
NZ_CP038253.1:758105-765957    6654   ------------------------------------------------------------------------------------------
NZ_CP035263.1:743050-750901    6653   ------------------------------------------------------------------------------------------
NZ_CP035262.1:741595-749460    6667   ------------------------------------------------------------------------------------------
NZ_CP035236.1:764915-772766    6653   ------------------------------------------------------------------------------------------
NZ_CP035253.1 767286-775134    6650   ------------------------------------------------------------------------------------------
NZ_CP035249.1:763080-770921    6643   ------------------------------------------------------------------------------------------

NC_003028.3:777000-787000      8154   AATCTTGCGCAATGAAACTCATCTTTCCCTTAGTGAAGGCATACTCATCCCAAGACATAATCTTTGGAAGCCGAGAAAAATCATGCTCAA
NZ_CP035248.1:765099-775111    8167   AATCTTGCGCAATGAAACTCATCTTTCCCTTAGTGAAGGCATACTCATCCCAAGACATAATCTTTGGAAGCCGAGAAAAATCATGCTCAA
NZ_CP035235.1 c1326834-1317894 6667   ------------------------------------------------------------------------------------------
NZ_CP035234.1:756284-765224    6667   ------------------------------------------------------------------------------------------
NZ_CP038251.1:750745-759686    6667   ------------------------------------------------------------------------------------------
NZ_CP035244.1:747023-755950    6653   ------------------------------------------------------------------------------------------
NZ_CP035264.1:800836-809336    7302   ------------------------------------------------------------------------------------------
NZ_CP035265.1:766128-774627    7301   ------------------------------------------------------------------------------------------
NZ_CP035245.1 745261-753757    7298   ------------------------------------------------------------------------------------------
NZ_CP035259.1:745700-753548    6650   ------------------------------------------------------------------------------------------
NZ_CP035258.1:750874-758722    6650   ------------------------------------------------------------------------------------------
NZ_CP035260.1:726156-733994    6640   ------------------------------------------------------------------------------------------
NZ_CP035241.1:708296-716144    6650   ------------------------------------------------------------------------------------------
NZ_CP035261.1:779086-786938    6654   ------------------------------------------------------------------------------------------
NZ_CP035242.1:c1532236-1524385 6653   ------------------------------------------------------------------------------------------
NZ_CP035243.1:777729-785580    6653   ------------------------------------------------------------------------------------------
NZ_CP035237.1:c1311322-1303456 6668   ------------------------------------------------------------------------------------------
NZ_CP035246.1:c1302892-1295026 6668   ------------------------------------------------------------------------------------------
NZ_CP035247.1:c1301115-1293263 6654   ------------------------------------------------------------------------------------------
NZ_CP035238.1:738547-746413    6668   ------------------------------------------------------------------------------------------
NZ_CP035255.1:776866-784718    6654   ------------------------------------------------------------------------------------------
NZ_CP035256.1:807236-815102    6668   ------------------------------------------------------------------------------------------
NZ_CP038252.1:1151812-1159673  6593   ------------------------------------------------------------------------------------------
NZ_CP035254.1:793365-801213    6650   ------------------------------------------------------------------------------------------
NZ_CP035251.1:748733-756585    6654   ------------------------------------------------------------------------------------------
NZ_CP035252.1:745670-753536    6668   ------------------------------------------------------------------------------------------
NZ_CP035257.1:776317-784169    6654   ------------------------------------------------------------------------------------------
NZ_CP038253.1:758105-765957    6654   ------------------------------------------------------------------------------------------
NZ_CP035263.1:743050-750901    6653   ------------------------------------------------------------------------------------------
NZ_CP035262.1:741595-749460    6667   ------------------------------------------------------------------------------------------
NZ_CP035236.1:764915-772766    6653   ------------------------------------------------------------------------------------------
NZ_CP035253.1 767286-775134    6650   ------------------------------------------------------------------------------------------
NZ_CP035249.1:763080-770921    6643   ------------------------------------------------------------------------------------------

NC_003028.3:777000-787000      8244   AGTGAAAGTCATTGAGCTTGCGAATGACAGTTGAAGTTGAAATGGCCAGCTGATGGGCAATATCAGTCATAGAAATTTTTTCAATTAACT
NZ_CP035248.1:765099-775111    8257   AGTGAAAGTCATTGAGCTTGCGAATGACAGTTGAAGTTGAAATGGCCAGCTGATGGGCAATATCAGTCATAGAAATTTTTTCAATTAACT
NZ_CP035235.1 c1326834-1317894 6667   ------------------------------------------------------------------------------------------
NZ_CP035234.1:756284-765224    6667   ------------------------------------------------------------------------------------------
NZ_CP038251.1:750745-759686    6667   ------------------------------------------------------------------------------------------
NZ_CP035244.1:747023-755950    6653   ------------------------------------------------------------------------------------------
NZ_CP035264.1:800836-809336    7302   ------------------------------------------------------------------------------------------
NZ_CP035265.1:766128-774627    7301   ------------------------------------------------------------------------------------------
NZ_CP035245.1 745261-753757    7298   ------------------------------------------------------------------------------------------
NZ_CP035259.1:745700-753548    6650   ------------------------------------------------------------------------------------------
NZ_CP035258.1:750874-758722    6650   ------------------------------------------------------------------------------------------
NZ_CP035260.1:726156-733994    6640   ------------------------------------------------------------------------------------------
NZ_CP035241.1:708296-716144    6650   ------------------------------------------------------------------------------------------
NZ_CP035261.1:779086-786938    6654   ------------------------------------------------------------------------------------------
NZ_CP035242.1:c1532236-1524385 6653   ------------------------------------------------------------------------------------------
NZ_CP035243.1:777729-785580    6653   ------------------------------------------------------------------------------------------
NZ_CP035237.1:c1311322-1303456 6668   ------------------------------------------------------------------------------------------
NZ_CP035246.1:c1302892-1295026 6668   ------------------------------------------------------------------------------------------
NZ_CP035247.1:c1301115-1293263 6654   ------------------------------------------------------------------------------------------
NZ_CP035238.1:738547-746413    6668   ------------------------------------------------------------------------------------------
NZ_CP035255.1:776866-784718    6654   ------------------------------------------------------------------------------------------
NZ_CP035256.1:807236-815102    6668   ------------------------------------------------------------------------------------------
NZ_CP038252.1:1151812-1159673  6593   ------------------------------------------------------------------------------------------
NZ_CP035254.1:793365-801213    6650   ------------------------------------------------------------------------------------------
NZ_CP035251.1:748733-756585    6654   ------------------------------------------------------------------------------------------
NZ_CP035252.1:745670-753536    6668   ------------------------------------------------------------------------------------------
NZ_CP035257.1:776317-784169    6654   ------------------------------------------------------------------------------------------
NZ_CP038253.1:758105-765957    6654   ------------------------------------------------------------------------------------------
NZ_CP035263.1:743050-750901    6653   ------------------------------------------------------------------------------------------
NZ_CP035262.1:741595-749460    6667   ------------------------------------------------------------------------------------------
NZ_CP035236.1:764915-772766    6653   ------------------------------------------------------------------------------------------
NZ_CP035253.1 767286-775134    6650   ------------------------------------------------------------------------------------------
NZ_CP035249.1:763080-770921    6643   ------------------------------------------------------------------------------------------

NC_003028.3:777000-787000      8334   TTTGAGCAATTTTTTGGTTGATGATACGAGGGATTTGGTGATTTTTCTTTACCAGGGGAGTCTCAGCAACCATCATTTTTGAAGAGTGAT
NZ_CP035248.1:765099-775111    8347   TTTGAGCAATTTTTTGGTTGATGATACGAGGGATTTGGTGATTTTTCTTTACCAGGGGAGTCTCAGCAACCATCATTTTTGAAGAGTGAT
NZ_CP035235.1 c1326834-1317894 6667   ------------------------------------------------------------------------------------------
NZ_CP035234.1:756284-765224    6667   ------------------------------------------------------------------------------------------
NZ_CP038251.1:750745-759686    6667   ------------------------------------------------------------------------------------------
NZ_CP035244.1:747023-755950    6653   ------------------------------------------------------------------------------------------
NZ_CP035264.1:800836-809336    7302   ------------------------------------------------------------------------------------------
NZ_CP035265.1:766128-774627    7301   ------------------------------------------------------------------------------------------
NZ_CP035245.1 745261-753757    7298   ------------------------------------------------------------------------------------------
NZ_CP035259.1:745700-753548    6650   ------------------------------------------------------------------------------------------
NZ_CP035258.1:750874-758722    6650   ------------------------------------------------------------------------------------------
NZ_CP035260.1:726156-733994    6640   ------------------------------------------------------------------------------------------
NZ_CP035241.1:708296-716144    6650   ------------------------------------------------------------------------------------------
NZ_CP035261.1:779086-786938    6654   ------------------------------------------------------------------------------------------
NZ_CP035242.1:c1532236-1524385 6653   ------------------------------------------------------------------------------------------
NZ_CP035243.1:777729-785580    6653   ------------------------------------------------------------------------------------------
NZ_CP035237.1:c1311322-1303456 6668   ------------------------------------------------------------------------------------------
NZ_CP035246.1:c1302892-1295026 6668   ------------------------------------------------------------------------------------------
NZ_CP035247.1:c1301115-1293263 6654   ------------------------------------------------------------------------------------------
NZ_CP035238.1:738547-746413    6668   ------------------------------------------------------------------------------------------
```

```
NZ_CP035255.1:776866-784718    6654   ------------------------------------------------------------------------------------------
NZ_CP035256.1:807236-815102    6668   ------------------------------------------------------------------------------------------
NZ_CP038252.1:1151812-1159673  6593   ------------------------------------------------------------------------------------------
NZ_CP035254.1:793365-801213    6650   ------------------------------------------------------------------------------------------
NZ_CP035251.1:748733-756585    6654   ------------------------------------------------------------------------------------------
NZ_CP035252.1:745670-753536    6668   ------------------------------------------------------------------------------------------
NZ_CP035257.1:776317-784169    6654   ------------------------------------------------------------------------------------------
NZ_CP038253.1:758105-765957    6654   ------------------------------------------------------------------------------------------
NZ_CP035263.1:743050-750901    6653   ------------------------------------------------------------------------------------------
NZ_CP035262.1:741595-749460    6667   ------------------------------------------------------------------------------------------
NZ_CP035236.1:764915-772766    6653   ------------------------------------------------------------------------------------------
NZ_CP035253.1 767286-775134    6650   ------------------------------------------------------------------------------------------
NZ_CP035249.1:763080-770921    6643   ------------------------------------------------------------------------------------------

NC_003028.3:777000-787000      8424   AGCACTTGAAACGGCGTTTTCTAAGGAGAATTCTAGAAGGCATACCAGTTGTTTCGAGGTAAGGGATCTTAGACGGTTTTTGAAAGTCAT
NZ_CP035248.1:765099-775111    8437   AGCACTTGAAACGGCGTTTTCTAAGGAGAATTCTAGAAGGCATACCAGTTGTTTCGAGGTAAGGGATCTTAGACGGTTTTTGAAAGTCAT
NZ_CP035235.1 c1326834-1317894 6667   ------------------------------------------------------------------------------------------
NZ_CP035234.1:756284-765224    6667   ------------------------------------------------------------------------------------------
NZ_CP038251.1:750745-759686    6667   ------------------------------------------------------------------------------------------
NZ_CP035244.1:747023-755950    6653   ------------------------------------------------------------------------------------------
NZ_CP035264.1:800836-809336    7302   ------------------------------------------------------------------------------------------
NZ_CP035265.1:766128-774627    7301   ------------------------------------------------------------------------------------------
NZ_CP035245.1 745261-753757    7298   ------------------------------------------------------------------------------------------
NZ_CP035259.1:745700-753548    6650   ------------------------------------------------------------------------------------------
NZ_CP035258.1:750874-758722    6650   ------------------------------------------------------------------------------------------
NZ_CP035260.1:726156-733994    6640   ------------------------------------------------------------------------------------------
NZ_CP035241.1:708296-716144    6650   ------------------------------------------------------------------------------------------
NZ_CP035261.1:779086-786938    6654   ------------------------------------------------------------------------------------------
NZ_CP035242.1:c1532236-1524385 6653   ------------------------------------------------------------------------------------------
NZ_CP035243.1:777729-785580    6653   ------------------------------------------------------------------------------------------
NZ_CP035237.1:c1311322-1303456 6668   ------------------------------------------------------------------------------------------
NZ_CP035246.1:c1302892-1295026 6668   ------------------------------------------------------------------------------------------
NZ_CP035247.1:c1301115-1293263 6654   ------------------------------------------------------------------------------------------
NZ_CP035238.1:738547-746413    6668   ------------------------------------------------------------------------------------------
NZ_CP035255.1:776866-784718    6654   ------------------------------------------------------------------------------------------
NZ_CP035256.1:807236-815102    6668   ------------------------------------------------------------------------------------------
NZ_CP038252.1:1151812-1159673  6593   ------------------------------------------------------------------------------------------
NZ_CP035254.1:793365-801213    6650   ------------------------------------------------------------------------------------------
NZ_CP035251.1:748733-756585    6654   ------------------------------------------------------------------------------------------
NZ_CP035252.1:745670-753536    6668   ------------------------------------------------------------------------------------------
NZ_CP035257.1:776317-784169    6654   ------------------------------------------------------------------------------------------
NZ_CP038253.1:758105-765957    6654   ------------------------------------------------------------------------------------------
NZ_CP035263.1:743050-750901    6653   ------------------------------------------------------------------------------------------
NZ_CP035262.1:741595-749460    6667   ------------------------------------------------------------------------------------------
NZ_CP035236.1:764915-772766    6653   ------------------------------------------------------------------------------------------
NZ_CP035253.1 767286-775134    6650   ------------------------------------------------------------------------------------------
NZ_CP035249.1:763080-770921    6643   ------------------------------------------------------------------------------------------

NC_003028.3:777000-787000      8514   GTTTCTTCATTAGACTTCCACAATCAGGGCAAGATGGAGCCTCATAATCCAGCTTAGCGATAATTTCTTTGTGGGTATCCATATTGATGA
NZ_CP035248.1:765099-775111    8527   ATTTCTTCATTAGACTTCCACAATCAGGGCAAGATGGAGCCTCATAATCCAGCTTAGCGATAATTTCTTTGTGGGTATCCATATTGATGA
NZ_CP035235.1 c1326834-1317894 6667   ------------------------------------------------------------------------------------------
NZ_CP035234.1:756284-765224    6667   ------------------------------------------------------------------------------------------
NZ_CP038251.1:750745-759686    6667   ------------------------------------------------------------------------------------------
NZ_CP035244.1:747023-755950    6653   ------------------------------------------------------------------------------------------
NZ_CP035264.1:800836-809336    7302   ------------------------------------------------------------------------------------------
NZ_CP035265.1:766128-774627    7301   ------------------------------------------------------------------------------------------
NZ_CP035245.1 745261-753757    7298   ------------------------------------------------------------------------------------------
NZ_CP035259.1:745700-753548    6650   ------------------------------------------------------------------------------------------
NZ_CP035258.1:750874-758722    6650   ------------------------------------------------------------------------------------------
NZ_CP035260.1:726156-733994    6640   ------------------------------------------------------------------------------------------
NZ_CP035241.1:708296-716144    6650   ------------------------------------------------------------------------------------------
NZ_CP035261.1:779086-786938    6654   ------------------------------------------------------------------------------------------
NZ_CP035242.1:c1532236-1524385 6653   ------------------------------------------------------------------------------------------
NZ_CP035243.1:777729-785580    6653   ------------------------------------------------------------------------------------------
NZ_CP035237.1:c1311322-1303456 6668   ------------------------------------------------------------------------------------------
NZ_CP035246.1:c1302892-1295026 6668   ------------------------------------------------------------------------------------------
NZ_CP035247.1:c1301115-1293263 6654   ------------------------------------------------------------------------------------------
NZ_CP035238.1:738547-746413    6668   ------------------------------------------------------------------------------------------
NZ_CP035255.1:776866-784718    6654   ------------------------------------------------------------------------------------------
NZ_CP035256.1:807236-815102    6668   ------------------------------------------------------------------------------------------
NZ_CP038252.1:1151812-1159673  6593   ------------------------------------------------------------------------------------------
NZ_CP035254.1:793365-801213    6650   ------------------------------------------------------------------------------------------
NZ_CP035251.1:748733-756585    6654   ------------------------------------------------------------------------------------------
NZ_CP035252.1:745670-753536    6668   ------------------------------------------------------------------------------------------
NZ_CP035257.1:776317-784169    6654   ------------------------------------------------------------------------------------------
NZ_CP038253.1:758105-765957    6654   ------------------------------------------------------------------------------------------
NZ_CP035263.1:743050-750901    6653   ------------------------------------------------------------------------------------------
NZ_CP035262.1:741595-749460    6667   ------------------------------------------------------------------------------------------
NZ_CP035236.1:764915-772766    6653   ------------------------------------------------------------------------------------------
NZ_CP035253.1 767286-775134    6650   ------------------------------------------------------------------------------------------
NZ_CP035249.1:763080-770921    6643   ------------------------------------------------------------------------------------------

NC_003028.3:777000-787000      8604   TATCTAGAATCTTGATGTTTGGGTCTTTAATATCGAGCAGTTTTGTGATAAAATGTAATTGTTCCATATGATTCTTTCTAATGAGTTGTT
NZ_CP035248.1:765099-775111    8617   TATCTAGAATCTTGATGTTTGGGTCTTTAATATCGAGCAGTTTTGTGATAAAATGTAATTGTTCCATATGAATCTTTCTAATGA-TGGTT
NZ_CP035235.1 c1326834-1317894 6667   ------------------------------------------------------------------------------------------
NZ_CP035234.1:756284-765224    6667   ------------------------------------------------------------------------------------------
NZ_CP038251.1:750745-759686    6667   ------------------------------------------------------------------------------------------
NZ_CP035244.1:747023-755950    6653   ------------------------------------------------------------------------------------------
NZ_CP035264.1:800836-809336    7302   ------------------------------------------------------------------------------------------
NZ_CP035265.1:766128-774627    7301   ------------------------------------------------------------------------------------------
NZ_CP035245.1 745261-753757    7298   ------------------------------------------------------------------------------------------
NZ_CP035259.1:745700-753548    6650   ------------------------------------------------------------------------------------------
NZ_CP035258.1:750874-758722    6650   ------------------------------------------------------------------------------------------
NZ_CP035260.1:726156-733994    6640   ------------------------------------------------------------------------------------------
NZ_CP035241.1:708296-716144    6650   ------------------------------------------------------------------------------------------
```

```
NZ_CP035261.1:779086-786938    6654   ------------------------------------------------------------------------------------------
NZ_CP035242.1:c1532236-1524385 6653   ------------------------------------------------------------------------------------------
NZ_CP035243.1:777729-785580    6653   ------------------------------------------------------------------------------------------
NZ_CP035237.1:c1311322-1303456 6668   ------------------------------------------------------------------------------------------
NZ_CP035246.1:c1302892-1295026 6668   ------------------------------------------------------------------------------------------
NZ_CP035247.1:c1301115-1293263 6654   ------------------------------------------------------------------------------------------
NZ_CP035238.1:738547-746413    6668   ------------------------------------------------------------------------------------------
NZ_CP035255.1:776866-784718    6654   ------------------------------------------------------------------------------------------
NZ_CP035256.1:807236-815102    6668   ------------------------------------------------------------------------------------------
NZ_CP038252.1:1151812-1159673  6593   ------------------------------------------------------------------------------------------
NZ_CP035254.1:793365-801213    6650   ------------------------------------------------------------------------------------------
NZ_CP035251.1:748733-756585    6654   ------------------------------------------------------------------------------------------
NZ_CP035252.1:745670-753536    6668   ------------------------------------------------------------------------------------------
NZ_CP035257.1:776317-784169    6654   ------------------------------------------------------------------------------------------
NZ_CP038253.1:758105-765957    6654   ------------------------------------------------------------------------------------------
NZ_CP035263.1:743050-750901    6653   ------------------------------------------------------------------------------------------
NZ_CP035262.1:741595-749460    6667   ------------------------------------------------------------------------------------------
NZ_CP035236.1:764915-772766    6653   ------------------------------------------------------------------------------------------
NZ_CP035253.1 767286-775134    6650   ------------------------------------------------------------------------------------------
NZ_CP035249.1:763080-770921    6643   ------------------------------------------------------------------------------------------

NC_003028.3:777000-787000      8694   TTGTCGCTTTTCATTATAGGTCATATGGGACTTTTTTTCTACACAAAAATAGGCTCCATAATATCTATAGTGGATTTACCCACTACAAAT
NZ_CP035248.1:765099-775111    8706   TTGTCGCTTTTCATTATAGGTCATATGGGACTTTTTTTCTACAATAAAATAGGCTCCATAATATCTATAGTGGATTTACCCACTACAAAT
NZ_CP035235.1 c1326834-1317894 6667   ------------------------------------------------------------------------------------------
NZ_CP035234.1:756284-765224    6667   ------------------------------------------------------------------------------------------
NZ_CP038251.1:750745-759686    6667   ------------------------------------------------------------------------------------------
NZ_CP035244.1:747023-755950    6653   ------------------------------------------------------------------------------------------
NZ_CP035264.1:800836-809336    7302   ------------------------------------------------------------------------------------------
NZ_CP035265.1:766128-774627    7301   ------------------------------------------------------------------------------------------
NZ_CP035245.1 745261-753757    7298   ------------------------------------------------------------------------------------------
NZ_CP035259.1:745700-753548    6650   ------------------------------------------------------------------------------------------
NZ_CP035258.1:750874-758722    6650   ------------------------------------------------------------------------------------------
NZ_CP035260.1:726156-733994    6640   ------------------------------------------------------------------------------------------
NZ_CP035241.1:708296-716144    6650   ------------------------------------------------------------------------------------------
NZ_CP035261.1:779086-786938    6654   ------------------------------------------------------------------------------------------
NZ_CP035242.1:c1532236-1524385 6653   ------------------------------------------------------------------------------------------
NZ_CP035243.1:777729-785580    6653   ------------------------------------------------------------------------------------------
NZ_CP035237.1:c1311322-1303456 6668   ------------------------------------------------------------------------------------------
NZ_CP035246.1:c1302892-1295026 6668   ------------------------------------------------------------------------------------------
NZ_CP035247.1:c1301115-1293263 6654   ------------------------------------------------------------------------------------------
NZ_CP035238.1:738547-746413    6668   ------------------------------------------------------------------------------------------
NZ_CP035255.1:776866-784718    6654   ------------------------------------------------------------------------------------------
NZ_CP035256.1:807236-815102    6668   ------------------------------------------------------------------------------------------
NZ_CP038252.1:1151812-1159673  6593   ------------------------------------------------------------------------------------------
NZ_CP035254.1:793365-801213    6650   ------------------------------------------------------------------------------------------
NZ_CP035251.1:748733-756585    6654   ------------------------------------------------------------------------------------------
NZ_CP035252.1:745670-753536    6668   ------------------------------------------------------------------------------------------
NZ_CP035257.1:776317-784169    6654   ------------------------------------------------------------------------------------------
NZ_CP038253.1:758105-765957    6654   ------------------------------------------------------------------------------------------
NZ_CP035263.1:743050-750901    6653   ------------------------------------------------------------------------------------------
NZ_CP035262.1:741595-749460    6667   ------------------------------------------------------------------------------------------
NZ_CP035236.1:764915-772766    6653   ------------------------------------------------------------------------------------------
NZ_CP035253.1 767286-775134    6650   ------------------------------------------------------------------------------------------
NZ_CP035249.1:763080-770921    6643   ------------------------------------------------------------------------------------------

NC_003028.3:777000-787000      8784   ATTATAGAGCCCAAAAAGGAAGCCCTTTATGAATTGTAGGACTTCCTTTTCTTATCCAGAAATTGATCTAGCTCTCTCTGATTTCGAAGA
NZ_CP035248.1:765099-775111    8796   ATTATAGAGCCCAAAAAGGAAGCCCTTTATGAATTGTAGGACTTCCTTTTCTTATCCAGAAATTGATCTAGCTCTCTCTGATTTCGAAGA
NZ_CP035235.1 c1326834-1317894 6667   -------------------AAGCCCTTTATGAATTGTAGGACTTCCTTTTCTTATCCAGAAATTGATCTAGCTCTCTCTGATTTCGAAGA
NZ_CP035234.1:756284-765224    6667   -------------------AAGCCCTTTATGAATTGTAGGACTTCCTTTTCTTATCCAGAAATTGATCTAGCTCTCTCTGATTTCGAAGA
NZ_CP038251.1:750745-759686    6667   -------------------AAGCCCTTTATGAATTGTAGGACTTCCTTTTCTTATCCAGAAATTGATCTAGCTCTCTCTGATTTCGAAGA
NZ_CP035244.1:747023-755950    6653   -------------------AAGCCCTTTATGAATTGTAGGACTTCCTTTTCTTATCCAGAAATTGATCTAGCTCTCTCTGATTTCGAAGA
NZ_CP035264.1:800836-809336    7302   -------------------AAGCCCTTTATGAATTGTAGGACTTCCTTTTCTTATCCAGAAATTGATCTAGCTCTCTCTGATTTCGAAGA
NZ_CP035265.1:766128-774627    7301   -------------------AAGCCCTTTATGAATTGTAGGACTTCCTTTTCTTATCCAGAAATTGATCTAGCTCTCTCTGATTTCGAAGA
NZ_CP035245.1 745261-753757    7298   -------------------AAGCCCTTTATGAATTGTAGGACTTCCTTTTCTTATCCAGAAATTGATCTAGCTCTCTCTGATTTCGAAGA
NZ_CP035259.1:745700-753548    6650   -------------------AAGCCCTTTATGAATTGTAGGACTTCCTTTTCTTATCCAGAAATTGATCTAGCTCTCTCTGATTTCGAAGG
NZ_CP035258.1:750874-758722    6650   -------------------AAGCCCTTTATGAATTGTAGGACTTCCTTTTCTTATCCAGAAATTGATCTAGCTCTCTCTGATTTCGAAGG
NZ_CP035260.1:726156-733994    6640   -------------------AAGCCCTTTATGAATTGTAGGACTTCCTTTTCTTATCCAGAAATTGATCTAGCTTTCTCTGATTTCGAAGA
NZ_CP035241.1:708296-716144    6650   -------------------AAGCCCTTTATGAATTGTAGGACTTCCTTTTCTTATCCAGAAATTGATCTAGCTCTCTCTGATTTCGAAGA
NZ_CP035261.1:779086-786938    6654   -------------------AAGCCCTTTATGAATTGTAGGACTTCCTTTTCTTATCCAGAAATTGATCTAGCTCTCTCTGATTTCGAAGA
NZ_CP035242.1:c1532236-1524385 6653   -------------------AAGCCCTTTATGAATTGTAGGACTTCCTTTTCTTATCCAGAAATTGATCTAGCTCTCTCTGATTTCGAAGA
NZ_CP035243.1:777729-785580    6653   -------------------AAGCCCTTTATGAATTGTAGGACTTCCTTTTCTTATCCAGAAATTGATCTAGCTCTCTCTGATTTCGAAGA
NZ_CP035237.1:c1311322-1303456 6668   -------------------AAGCCCTTTATGAATTGTAGGACTTCCTTTTCTTATCCAGAAATTGATCTAGCTCTCTCTGATTTCGAAGA
NZ_CP035246.1:c1302892-1295026 6668   -------------------AAGCCCTTTATGAATTGTAGGACTTCCTTTTCTTATCCAGAAATTGATCTAGCTTTCTCTGATTTCGAAGA
NZ_CP035247.1:c1301115-1293263 6654   -------------------AAGCCCTTTATGAATTGTAGGACTTCCTTTTCTTATCCAGAAATTGATCTAGCTTTCTCTGATTTCGAAGA
NZ_CP035238.1:738547-746413    6668   -------------------AAGCCCTTTATGAATTGTAGGACTTCCTTTTCTTATCCAGAAATTGATCTAGCTCTCTCTGATTTCGAAGG
NZ_CP035255.1:776866-784718    6654   -------------------AAGCCCTTTATGAATTGTAGGACTTCCTTTTCTTATCCAGAAATTGATCTAGCTCTCTCTGATTTCGAAGA
NZ_CP035256.1:807236-815102    6668   -------------------AAGCCCTTTATGAATTGTAGGACTTCCTTTTCTTATCCAGAAATTGATCTAGCTCTCTCTGATTTCGAAGA
NZ_CP038252.1:1151812-1159673  6593   -------------------AAGCCCTTTATGAATTGTAGGACTTCCTTTTCTTATCCAGAAATTGATCTAGCTCTCTCTGATTTCGAAGA
NZ_CP035254.1:793365-801213    6650   -------------------AAGCCCTTTATGAATTGTAGGACTTCCTTTTCTTATCCAGAAATTGATCTAGCTCTCTCTGATTTCGAAGA
NZ_CP035251.1:748733-756585    6654   -------------------AAGCCCTTTATGAATTGTAGGACTTCCTTTTCTTATCCAGAAATTGATCTAGTTCTCTCTGATTTCGAAGA
NZ_CP035252.1:745670-753536    6668   -------------------AAGCCCTTTATGAATTGTAGGACTTCCTTTTCTTATCCAGAAATTGATCTAGTTCTCTCTGATTTCGAAGA
NZ_CP035257.1:776317-784169    6654   -------------------AAGCCCTTTATGAATTGTAGGACTTCCTTTTCTTATCCAGAAATTGATCTAGTTCTCTCTGATTTCGAAGA
NZ_CP038253.1:758105-765957    6654   -------------------AAGCCCTTTATGAATTGTAGGACTTCCTTTTCTTATCCAGAAATTGATCTAGTTCTCTCTGATTTCGAAGA
NZ_CP035263.1:743050-750901    6653   -------------------AAGCCCTTTATGAATTGTAGGACTTCCTTTTCTTATCCAGAAATTGATCTAGCTCTCTCTGATTTCGAAGA
NZ_CP035262.1:741595-749460    6667   -------------------AAGCCCTTTATGAATTGTAGGACTTCCTTTTCTTATCCAGAAATTGATCTAGCTCTCTCTGATTTCGAAGA
NZ_CP035236.1:764915-772766    6653   -------------------AAGCCCTTTATGAATTGTAGGACTTCCTTTTCTTATCCAGAAATTGATCTAGCTCTCTCTGATTTCGAAGA
NZ_CP035253.1 767286-775134    6650   -------------------AAGCCCTTTATGAATTGTAGGACTTCCTTTTCTTATCCAGAAATTGATCTAGCTCTCTCTGATTTCGAAGA
NZ_CP035249.1:763080-770921    6643   -------------------AAGCCCTTTATGAATTGTAGGACTTCCTTTTCTTATCCAGAAATTGATCTAGCTCTCTCTGATTTCGAAGA

NC_003028.3:777000-787000      8874   ATAGTGACTTTATGTGAATATTCTTGGCAAAGTTTTTGGTAATTTTCTTTTTGAGTTTTGCTACGCCCATCCCAAAGAATCCATCTGATA
NZ_CP035248.1:765099-775111    8886   ATAGTGACTTTATGTGAATATTCTTGGCAAAGTTTTTGGTAATTTTCTTTTTGAGTTTTGCTACGCCCATCCCAAAGAATCCATCTGATA
NZ_CP035235.1 c1326834-1317894 6739   ATAGTGACTTTATGTGAATATTCTTGGCAAAGTTTTTGGTAATTTTCTTTTTGAGTTTTGCTACGTCCATCCCAAAGAATCCATCTGATA
NZ_CP035234.1:756284-765224    6739   ATAGTGACTTTATGTGAATATTCTTGGCAAAGTTTTTGGTAATTTTCTTTTTGAGTTTTGCTACGTCCATCCCAAAGAATCCATCTGATA
NZ_CP038251.1:750745-759686    6739   ATAGTGACTTTATGTGAATATTCTTGGCAAAGTTTTTGGTAATTTTCTTTTTGAGTTTTGCTACGTCCATCCCAAAGAATCCATCTGATA
NZ_CP035244.1:747023-755950    6725   ATAGTGACTTTATGTGAATATTCTTGGCAAAGTTTTTGGTAATTTTCTTTTTGAGTTTTGCTACGTCCATCCCAAAGAATCCATCTGATA
```

### Ozkan et al. Supplementary Figures

```
NZ_CP035264.1:800836-809336    7374   ATAGTGACTTTATGTGAATATTCTTGGCAAAGTTTTTGGTAATTTTCTTTTTGAGTTTTGCTACGCCCATCCCAAAGAATCCATCTGATA
NZ_CP035265.1:766128-774627    7373   ATAGTGACTTTATGTGAATATTCTTGGCAAAGTTTTTGGTAATTTTCTTTTTGAGTTTTGCTACGCCCATCCCAAAGAATCCATCTGATA
NZ_CP035245.1 745261-753757    7370   ATAGTGACTTTATGTGAATATTCTTGGCAAAGTTTTTGGTAATTTTCTTTTTGAGTTTTGCTACGCCCATCCCAAAGAATCCATCTGATA
NZ_CP035259.1:745700-753548    6722   ATAGTGACTTTATGTGAATATTCTTGGCAAAGTTTTTGGTAATTTTCTTTTTGAGTTTTGCTACGCCCATCCCAAAGAATCCATCTGATA
NZ_CP035258.1:750874-758722    6722   ATAGTGACTTTATGTGAATATTCTTGGCAAAGTTTTTGGTAATTTTCTTTTTGAGTTTTGCTACGCCCATCCCAAAGAATCCATCTGATA
NZ_CP035260.1:726156-733994    6712   ATAGTGACTTTATGTGAATATTCTTGGCAAAGTTTTTGGTAATTTTCTTTTTGAGTTTTGCTACGCCCATCCCAAAGAATCCATCTGATA
NZ_CP035241.1:708296-716144    6722   ATAGTGACTTTATGTGAATATTCTTGGCAAAGTTTTTGGTAATTTTCTTTTTGAGTTTTGCTACGCCCATCCCAAAGAATCCATCTGATA
NZ_CP035261.1:779086-786938    6726   ATAGTGACTTTATGTGAATATTCTTGGCAAAGTTTTTGGTAATTTTCTTTTTGAGTTTTGCTACGCCCATCCCAAAGAATCCATCTGATA
NZ_CP035242.1:c1532236-1524385 6725   ATAGTGACTTTATGTGAATATTCTTGGCAAAGTTTTTGGTAATTTTCTTTTTGAGTTTTGCTACGCCCATCCCAAAGAATCCATCTGATA
NZ_CP035243.1:777729-785580    6725   ATAGTGACTTTATGTGAATATTCTTGGCAAAGTTTTTGGTAATTTTCTTTTTGAGTTTTGCTACGCCCATCCCAAAGAATCCATCTGATA
NZ_CP035237.1:c1311322-1303456 6740   ATAGTGACTTTATGTGAATATTCTTGGCAAAGTTTTTGGTAATTTTCTTTTTGAGTTTTGCTACGCCCATCCCAAAGAATCCATCTGATA
NZ_CP035246.1:c1302892-1295026 6740   ATAGTGACTTTATGTGAATATTCTTGGCAAAGTTTTTGGTAATTTTCTTTTTGAGTTTTGCTACGCCCATCCCAAAGAATCCATCTGATA
NZ_CP035247.1:c1301115-1293263 6726   ATAGTGACTTTATGTGAATATTCTTGGCAAAGTTTTTGGTAATTTTCTTTTTGAGTTTTGCTACGCCCATCCCAAAGAATCCATCTGATA
NZ_CP035238.1:738547-746413    6740   ATAGTGACTTTATGTGAATATTCTTGGCAAAGTTTTTGGTAATTTTCTTTTTGAGTTTTGCTACGCCCATCCCAAAGAATCCATCTGATA
NZ_CP035255.1:776866-784718    6726   ATAGTGACTTTATGTGAATATTCTTGGCAAAGTTTTTGGTAATTTTCTTTTTGAGTTTTGCTACGCCCATCCCAAAGAATCCATCTGATA
NZ_CP035256.1:807236-815102    6740   ATAGTGACTTTATGTGAATATTCTTGGCAAAGTTTTTGGTAATTTTCTTTTTGAGTTTTGCTACGCCCATCCCAAAGAATCCATCTGATA
NZ_CP038252.1:1151812-1159673  6665   ATAGTGACTTTATGTGAATATTCTTGGCAAAGTTTTTGGTAATTTTCTTTTTGAGTTTTGCTACGCCCATCCCAAAGAATCCATCTGATA
NZ_CP035254.1:793365-801213    6722   ATAGTGACTTTATGTGAATATTCTTGGCAAAGTTTTTGGTAATTTTCTTTTTGAGTTTTGCTACGCCCATCCCAAAGAATCCATCTGATA
NZ_CP035251.1:748733-756585    6726   ATAGTGACTTTATGTGAATATTCTTGGCAAAGTTTTTGGTAATTTTCTTTTTGAGTTTTGCTACGCCCATCCCAAAGAATCCATCTGATA
NZ_CP035252.1:745670-753536    6740   ATAGTGACTTTATGTGAATATTCTTGGCAAAGTTTTTGGTAATTTTCTTTTTGAGTTTTGCTACGCCCATCCCAAAGAATCCATCTGATA
NZ_CP035257.1:776317-784169    6726   ATAGTGACTTTATGTGAATATTCTTGGCAAAGTTTTTGGTAATTTTCTTTTTGAGTTTTGCTACGCCCATCCCAAAGAATCCATCTGATA
NZ_CP038253.1:758105-765957    6726   ATAGTGACTTTATGTGAATATTCTTGGCAAAGTTTTTGGTAATTTTCTTTTTGAGTTTTGCTACGCCCATCCCAAAGAATCCATCTGATA
NZ_CP035263.1:743050-750901    6725   ATAGTGACTTTATGTGAATATTCTTGGCAAAGTTTTTGGTAATTTTCTTTTTGAGTTTTGCTACGCCCATCCCAAAGAATCCATCTGATA
NZ_CP035262.1:741595-749460    6739   ATAGTGACTTTATGTGAATATTCTTGGCAAAGTTTTTGGTAATTTTCTTTTTGAGTTTTGCTACGCCCATCCCAAAGAATCCATCTGATA
NZ_CP035236.1:764915-772766    6725   ATAGTGACTTTATGTGAATATTCTTGGCAAAGTTTTTGGTAATTTTCTTTTTGAGTTTTGCTACGCCCATCCCAAAGAATCCATCTGATA
NZ_CP035253.1 767286-775134    6722   ATAGTGACTTTATGTGAATATTCTTGGCAAAGTTTTTGGTAATTTTCTTTTTGAGTTTTGCTACGCCCATCCCAAAGAATCCATCTGATA
NZ_CP035249.1:763080-770921    6715   ATAGTGACTTTATGTGAATATTCTTGGCAAAGTTTTTGGTAATTTTCTTTTTGAGTTTTGCTACGCCCATCCCAAAGAATCCATCTGATA

NC_003028.3:777000-787000      8964   AACTCCCACTCAAAGCGTTCAGGGCAATCTACCGCCATACTTTCTCTGACTTTTCCACGGTATTTAAGATAACGCTTAAA-GGCTCTA--
NZ_CP035248.1:765099-775111    8976   AACTCCCACTCAAAGCGTTCAGGGCAATCTACCGCCATACTTTCTCTGACTTTTCCACGGTATTTAAGATAACGCTTAAA-GGCTCTA--
NZ_CP035235.1 c1326834-1317894 6829   AACTCCCAGTCAAAGCGTTCAGGGCAATCTGCCGCCATACTTTCTCTGACTTTTCCACGGTATTTAAGATAACGCTTAAAGGGCTCTATA
NZ_CP035234.1:756284-765224    6829   AACTCCCAGTCAAAGCGTTCAGGGCAATCTGCCGCCATACTTTCTCTGACTTTTCCACGGTATTTAAGATAACGCTTAAAGGGCTCTATA
NZ_CP038251.1:750745-759686    6829   AACTCCCAGTCAAAGCGTTCAGGGCAATCTGCCGCCATACTTTCTCTGACTTTTCCACGGTATTTAAGATAACGCTTAAAGGGCTCTATA
NZ_CP035244.1:747023-755950    6815   AACTCCCAGTCAAAGCGTTCAGGGCAATCTGCCGCCATACTTTCTCTGACTTTTCCACGGTATTTAAGATAACGCTTAAAGGGCTCTATA
NZ_CP035264.1:800836-809336    7464   AACTCCCACTCAAAGCGTTCAGGGCAATCTGCCGCCATACTTTCTCTGACTTTTCCACGGTATTTAAGATAACGCTTAAA-GGCTCTA--
NZ_CP035265.1:766128-774627    7463   AACTCCCACTCAAAGCGTTCAGGGCAATCTGCCGCCATACTTTCTCTGACTTTTCCACGGTATTTAAGATAACGCTTAAA-GGCTCTA--
NZ_CP035245.1 745261-753757    7460   AACTCCCACTCAAAGCGTTCAGGGCAATCTGCCGCCATACTTTCTCTGACTTTTCCACGGTATTTAAGATAACGCTTAAA-GGCTCTA--
NZ_CP035259.1:745700-753548    6812   AACTCCCACTCAAAGCGTTCAGGGCAATCTGCCGCCATACTTTCTCTGACTTTTCCACGGTATTTAAGATAACGCTTAAA-GGCTCTA--
NZ_CP035258.1:750874-758722    6812   AACTCCCACTCAAAGCGTTCAGGGCAATCTGCCGCCATACTTTCTCTGACTTTTCCACGGTATTTAAGATAACGCTTAAA-GGCTCTA--
NZ_CP035260.1:726156-733994    6802   AACTCCCAGTCAAAGCGTTCAGGGCAATCTGCCGCCATACTTTCTCTGACTTTTCCACGGTATTTAAGATAACGCTTAAA-GGCTCTA--
NZ_CP035241.1:708296-716144    6812   AACTCCCAGTCAAAGCGTTCAGGGCAATCTGCCGCCATACTTTCTCTGACTTTTCCACGGTATTTAAGATAACGCTTAAA-GGCTCTA--
NZ_CP035261.1:779086-786938    6816   AACTCCCAGTCAAAGCGTTCAGGGCAATCTGCCGCCATACTTTCTCTGACTTTTCCACGGTATTTAAGATAACGCTTAAA-GGCTCTA--
NZ_CP035242.1:c1532236-1524385 6815   AACTCCCACTCAAAGCGTTCAGGGCAATCTACCGCCATACTTTCTCTGACTTTTCCACGGTATTTAAGATAACGCTTAAA-GGCTCTA--
NZ_CP035243.1:777729-785580    6815   AACTCCCACTCAAAGCGTTCAGGGCAATCTACCGCCATACTTTCTCTGACTTTTCCACGGTATTTAAGATAACGCTTAAA-GGCTCTA--
NZ_CP035237.1:c1311322-1303456 6830   AACTCCCAGTCAAAGCGTTCAGGGCAATCTGCCGCCATACTTTCTCTGACTTTTCCACGGTATTTAAGATAACGCTTAAA-GGCTCTA--
NZ_CP035246.1:c1302892-1295026 6830   AACTCCCACTCAAAGCGTTCAGGGCAATCTGCCGCCATACTTTCTCTGACTTTTCCACGGTATTTAAGATAACGCTTAAA-GGCTCTA--
NZ_CP035247.1:c1301115-1293263 6816   AACTCCCACTCAAAGCGTTCAGGGCAATCTGCCGCCATACTTTCTCTGACTTTTCCACGGTATTTAAGATAACGCTTAAA-GGCTCTA--
NZ_CP035238.1:738547-746413    6830   AACTCCCACTCAAAGCGTTCAGGGCAATCTGCCGCCATACTTTCTCTGACTTTTCCACGGTATTTAAGATAACGCTTAAA-GGCTCTA--
NZ_CP035255.1:776866-784718    6816   AACTCCCAGTCAAAGCGTTCAGGGCAATCTGCCGCCATACTTTCTCTGACTTTTCCACGGTATTTAAGATAACGCTTAAA-GGCTCTA--
NZ_CP035256.1:807236-815102    6830   AACTCCCAGTCAAAGCGTTCAGGGCAATCTGCCGCCATACTTTCTCTGACTTTTCCACGGTATTTAAGATAACGCTTAAA-GGCTCTA--
NZ_CP038252.1:1151812-1159673  6755   AACTCCCACTCAAAGCGTTCAGGGCAATCTGCCGCCATACTTTCTCTGACTTTTCCACGGTATTTAAGATAACGCTTAAA-GGCTCTA--
NZ_CP035254.1:793365-801213    6812   AACTCCCAGTCAAAGCGTTCAGGGCAATCTGCCGCCATACTTTCTCTGACTTTTCCACGGTATTTAAGATAACGCTTAAA-GGCTCTA--
NZ_CP035251.1:748733-756585    6816   AACTCCCACTCAAAGCGTTCAGGGCAATCTGCCGCCATACTTTCTCTGACTTTTCCACGGTATTTAAGATAACGCTTAAA-GGCTCTA--
NZ_CP035252.1:745670-753536    6830   AACTCCCACTCAAAGCGTTCAGGGCAATCTGCCGCCATACTTTCTCTGACTTTTCCACGGTATTTAAGATAACGCTTAAA-GGCTCTA--
NZ_CP035257.1:776317-784169    6816   AACTCCCACTCAAAGCGTTCAGGGCAATCTGCCGCCATACTTTCTCTGACTTTTCCACGGTATTTAAGATAACGCTTAAA-GGCTCTA--
NZ_CP038253.1:758105-765957    6816   AACTCCCACTCAAAGCGTTCAGGGCAATCTGCCGCCATACTTTCTCTGACTTTTCCACGGTATTTAAGATAACGCTTAAA-GGCTCTA--
NZ_CP035263.1:743050-750901    6815   AACTCCCACTCAAAGCGTTCAGGGCAATCTGCCGCCATACTTTCTCTGACTTTTCCACGGTATTTAAGATAACGCTTAAA-GGCTCTA--
NZ_CP035262.1:741595-749460    6829   AACTCCCACTCAAAGCGTTCAGGGCAATCTGCCGCCATACTTTCTCTGACTTTTCCACGGTATTTAAGATAACGCTTAAA-GGCTCTA--
NZ_CP035236.1:764915-772766    6815   AACTCCCAGTCAAAGCGTTCAGGGCAATCTGCCGCCATACTTTCTCTGACTTTTCCACGGTATTTAAGATAACGCTTAAA-GGTTCTA--
NZ_CP035253.1 767286-775134    6812   AACTCCCAGTCAAAGCGTTCAGGGCAATCTGCCGCCATACTTTCTCTGACTTTTCCACGGTATTTAAGATAACGCTTAAA-GGCTCTA--
NZ_CP035249.1:763080-770921    6805   AACTCCCACTCAAAGCGTTCAGGGCAATCTGCCGCCATACTTTCTCTGACTTTTCCACGGTATTTAAGATAACGCTTAAA-GGCTCTA--

NC_003028.3:777000-787000      9050   ------------------------------------------------------------------------------------------
NZ_CP035248.1:765099-775111    9062   ------------------------------------------------------------------------------------------
NZ_CP035235.1 c1326834-1317894 6919   ATATTTGTAGTGGGTAAATCCACTATAGATATTATGGAGCCTATTTTTGTGTAGAAAAAAAGTCCCATATGACCTATAATGAAAAGCGAC
NZ_CP035234.1:756284-765224    6919   ATATTTGTAGTGGGTAAATCCACTATAGATATTATGGAGCCTATTTTTGTGTAGAAAAAAAGTCCCATATGACCTATAATGAAAAGCGAC
NZ_CP038251.1:750745-759686    6919   ATATTTGTAGTGGGTAAATCCCTTATAGATATTATGGAGCCTATTTTTGTGTAGAAAAAAAGTCCCATATGACCTATAATGAAAAGCGAC
NZ_CP035244.1:747023-755950    6905   ATATTTGTAGTGGGTAAATCCCTTATAGATATTATGGAGCCTATTTTTGTGTAGAAAAAAAGTCCCATATGACCTATAATGAAAAGCGAC
NZ_CP035264.1:800836-809336    7550   ------------------------------------------------------------------------------------------
NZ_CP035265.1:766128-774627    7549   ------------------------------------------------------------------------------------------
NZ_CP035245.1 745261-753757    7546   ------------------------------------------------------------------------------------------
NZ_CP035259.1:745700-753548    6898   ------------------------------------------------------------------------------------------
NZ_CP035258.1:750874-758722    6898   ------------------------------------------------------------------------------------------
NZ_CP035260.1:726156-733994    6888   ------------------------------------------------------------------------------------------
NZ_CP035241.1:708296-716144    6898   ------------------------------------------------------------------------------------------
NZ_CP035261.1:779086-786938    6902   ------------------------------------------------------------------------------------------
NZ_CP035242.1:c1532236-1524385 6901   ------------------------------------------------------------------------------------------
NZ_CP035243.1:777729-785580    6901   ------------------------------------------------------------------------------------------
NZ_CP035237.1:c1311322-1303456 6916   ------------------------------------------------------------------------------------------
NZ_CP035246.1:c1302892-1295026 6916   ------------------------------------------------------------------------------------------
NZ_CP035247.1:c1301115-1293263 6902   ------------------------------------------------------------------------------------------
NZ_CP035238.1:738547-746413    6916   ------------------------------------------------------------------------------------------
NZ_CP035255.1:776866-784718    6902   ------------------------------------------------------------------------------------------
NZ_CP035256.1:807236-815102    6916   ------------------------------------------------------------------------------------------
NZ_CP038252.1:1151812-1159673  6841   ------------------------------------------------------------------------------------------
NZ_CP035254.1:793365-801213    6898   ------------------------------------------------------------------------------------------
NZ_CP035251.1:748733-756585    6902   ------------------------------------------------------------------------------------------
NZ_CP035252.1:745670-753536    6916   ------------------------------------------------------------------------------------------
NZ_CP035257.1:776317-784169    6902   ------------------------------------------------------------------------------------------
NZ_CP038253.1:758105-765957    6902   ------------------------------------------------------------------------------------------
NZ_CP035263.1:743050-750901    6901   ------------------------------------------------------------------------------------------
NZ_CP035262.1:741595-749460    6915   ------------------------------------------------------------------------------------------
NZ_CP035236.1:764915-772766    6901   ------------------------------------------------------------------------------------------
NZ_CP035253.1 767286-775134    6898   ------------------------------------------------------------------------------------------
NZ_CP035249.1:763080-770921    6891   ------------------------------------------------------------------------------------------
```

```
NC_003028.3:777000-787000      9050   ------------------------------------------------------------------------------------------
NZ_CP035248.1:765099-775111    9062   ------------------------------------------------------------------------------------------
NZ_CP035235.1 c1326834-1317894 7009   AAAACAACTCATTAGAAAGAATCATATGGAACAATTACATTTTATCACAAAATTACTAGACATTAAAGACCCTAATATCCAGATTTTAGA
NZ_CP035234.1:756284-765224    7009   AAAACAACTCATTAGAAAGAATCATATGGAACAATTACATTTTATCACAAAATTACTAGACATTAAAGACCCTAATATCCAGATTTTAGA
NZ_CP038251.1:750745-759686    7009   AAAACAACTCATTAGAAAGAATCATATGGAACAATTACATTTTATCACAAAATTACTAGACATTAAAGACCCTAATATCCAGATTTTAGA
NZ_CP035244.1:747023-755950    6995   AAAACAACTCATTAGAAAGAATCATATGGAACAATTACATTTTATCACAAAATTACTAGACATTAAAGACCCTAATATCCAGATTTTAGA
NZ_CP035264.1:800836-809336    7550   ------------------------------------------------------------------------------------------
NZ_CP035265.1:766128-774627    7549   ------------------------------------------------------------------------------------------
NZ_CP035245.1 745261-753757    7546   ------------------------------------------------------------------------------------------
NZ_CP035259.1:745700-753548    6898   ------------------------------------------------------------------------------------------
NZ_CP035258.1:750874-758722    6898   ------------------------------------------------------------------------------------------
NZ_CP035260.1:726156-733994    6888   ------------------------------------------------------------------------------------------
NZ_CP035241.1:708296-716144    6898   ------------------------------------------------------------------------------------------
NZ_CP035261.1:779086-786938    6902   ------------------------------------------------------------------------------------------
NZ_CP035242.1:c1532236-1524385 6901   ------------------------------------------------------------------------------------------
NZ_CP035243.1:777729-785580    6901   ------------------------------------------------------------------------------------------
NZ_CP035237.1:c1311322-1303456 6916   ------------------------------------------------------------------------------------------
NZ_CP035246.1:c1302892-1295026 6916   ------------------------------------------------------------------------------------------
NZ_CP035247.1:c1301115-1293263 6902   ------------------------------------------------------------------------------------------
NZ_CP035238.1:738547-746413    6916   ------------------------------------------------------------------------------------------
NZ_CP035255.1:776866-784718    6902   ------------------------------------------------------------------------------------------
NZ_CP035256.1:807236-815102    6916   ------------------------------------------------------------------------------------------
NZ_CP038252.1:1151812-1159673  6841   ------------------------------------------------------------------------------------------
NZ_CP035254.1:793365-801213    6898   ------------------------------------------------------------------------------------------
NZ_CP035251.1:748733-756585    6902   ------------------------------------------------------------------------------------------
NZ_CP035252.1:745670-753536    6916   ------------------------------------------------------------------------------------------
NZ_CP035257.1:776317-784169    6902   ------------------------------------------------------------------------------------------
NZ_CP038253.1:758105-765957    6902   ------------------------------------------------------------------------------------------
NZ_CP035263.1:743050-750901    6901   ------------------------------------------------------------------------------------------
NZ_CP035262.1:741595-749460    6915   ------------------------------------------------------------------------------------------
NZ_CP035236.1:764915-772766    6901   ------------------------------------------------------------------------------------------
NZ_CP035253.1 767286-775134    6898   ------------------------------------------------------------------------------------------
NZ_CP035249.1:763080-770921    6891   ------------------------------------------------------------------------------------------

NC_003028.3:777000-787000      9050   ------------------------------------------------------------------------------------------
NZ_CP035248.1:765099-775111    9062   ------------------------------------------------------------------------------------------
NZ_CP035235.1 c1326834-1317894 7099   CATCATCAATAAAGATACACACAAGGAAATCATCGCCAAACTGGACTACGACGCCCCATCTTGCCCTGATTGTGGAAGTCTAATGAAGAA
NZ_CP035234.1:756284-765224    7099   CATCATCAATAAAGATACACACAAGGAAATCATCGCCAAACTGGACTACGACGCCCCATCTTGCCCTGATTGTGGAAGTCTAATGAAGAA
NZ_CP038251.1:750745-759686    7099   CATCATCAATAAAGATACACACAAGGAAATCATCGCCAAACTGGACTACGACGCCCCATCTTGCCCTGATTGTGGAAGTCTAATGAAGAA
NZ_CP035244.1:747023-755950    7085   CATCATCAATAAAGATACACACAAGGAAATCATCGCCAAACTGGACTACGACGCCCCATCTTGCCCTGAGTGCGGAAGTCAAATGAAGAA
NZ_CP035264.1:800836-809336    7550   ------------------------------------------------------------------------------------------
NZ_CP035265.1:766128-774627    7549   ------------------------------------------------------------------------------------------
NZ_CP035245.1 745261-753757    7546   ------------------------------------------------------------------------------------------
NZ_CP035259.1:745700-753548    6898   ------------------------------------------------------------------------------------------
NZ_CP035258.1:750874-758722    6898   ------------------------------------------------------------------------------------------
NZ_CP035260.1:726156-733994    6888   ------------------------------------------------------------------------------------------
NZ_CP035241.1:708296-716144    6898   ------------------------------------------------------------------------------------------
NZ_CP035261.1:779086-786938    6902   ------------------------------------------------------------------------------------------
NZ_CP035242.1:c1532236-1524385 6901   ------------------------------------------------------------------------------------------
NZ_CP035243.1:777729-785580    6901   ------------------------------------------------------------------------------------------
NZ_CP035237.1:c1311322-1303456 6916   ------------------------------------------------------------------------------------------
NZ_CP035246.1:c1302892-1295026 6916   ------------------------------------------------------------------------------------------
NZ_CP035247.1:c1301115-1293263 6902   ------------------------------------------------------------------------------------------
NZ_CP035238.1:738547-746413    6916   ------------------------------------------------------------------------------------------
NZ_CP035255.1:776866-784718    6902   ------------------------------------------------------------------------------------------
NZ_CP035256.1:807236-815102    6916   ------------------------------------------------------------------------------------------
NZ_CP038252.1:1151812-1159673  6841   ------------------------------------------------------------------------------------------
NZ_CP035254.1:793365-801213    6898   ------------------------------------------------------------------------------------------
NZ_CP035251.1:748733-756585    6902   ------------------------------------------------------------------------------------------
NZ_CP035252.1:745670-753536    6916   ------------------------------------------------------------------------------------------
NZ_CP035257.1:776317-784169    6902   ------------------------------------------------------------------------------------------
NZ_CP038253.1:758105-765957    6902   ------------------------------------------------------------------------------------------
NZ_CP035263.1:743050-750901    6901   ------------------------------------------------------------------------------------------
NZ_CP035262.1:741595-749460    6915   ------------------------------------------------------------------------------------------
NZ_CP035236.1:764915-772766    6901   ------------------------------------------------------------------------------------------
NZ_CP035253.1 767286-775134    6898   ------------------------------------------------------------------------------------------
NZ_CP035249.1:763080-770921    6891   ------------------------------------------------------------------------------------------

NC_003028.3:777000-787000      9050   ------------------------------------------------------------------------------------------
NZ_CP035248.1:765099-775111    9062   ------------------------------------------------------------------------------------------
NZ_CP035235.1 c1326834-1317894 7189   ATATGACTTTCAAAAACCGTCTAAGATCCCTTACCTCGAAACAACTGGTATGCCTTCTAGAATTCTCCTTAGAAAACGCCGTTTCAAGTG
NZ_CP035234.1:756284-765224    7189   ATATGACTTTCAAAAACCGTCTAAGATCCCTTACCTCGAAACAACTGGTATGCCTTCTAGAATTCTCCTTAGAAAACGCCGTTTCAAGTG
NZ_CP038251.1:750745-759686    7189   ATATGACTTTCAAAAACCGTCTAAGATCCCTTACCTCGAAACAACTGGTATGCCTTCTAGAATTCTCCTTAGAAAACGCCGTTTCAAGTG
NZ_CP035244.1:747023-755950    7175   ATATGACTTTCAAAAACCTTCTAAAATTCCTTATCTTGAAACAACTGGTATGCCTTCTAGAATTCTCCTTAGAAAACGCCGTTTCAAGTG
NZ_CP035264.1:800836-809336    7550   ------------------------------------------------------------------------------------------
NZ_CP035265.1:766128-774627    7549   ------------------------------------------------------------------------------------------
NZ_CP035245.1 745261-753757    7546   ------------------------------------------------------------------------------------------
NZ_CP035259.1:745700-753548    6898   ------------------------------------------------------------------------------------------
NZ_CP035258.1:750874-758722    6898   ------------------------------------------------------------------------------------------
NZ_CP035260.1:726156-733994    6888   ------------------------------------------------------------------------------------------
NZ_CP035241.1:708296-716144    6898   ------------------------------------------------------------------------------------------
NZ_CP035261.1:779086-786938    6902   ------------------------------------------------------------------------------------------
NZ_CP035242.1:c1532236-1524385 6901   ------------------------------------------------------------------------------------------
NZ_CP035243.1:777729-785580    6901   ------------------------------------------------------------------------------------------
NZ_CP035237.1:c1311322-1303456 6916   ------------------------------------------------------------------------------------------
NZ_CP035246.1:c1302892-1295026 6916   ------------------------------------------------------------------------------------------
NZ_CP035247.1:c1301115-1293263 6902   ------------------------------------------------------------------------------------------
NZ_CP035238.1:738547-746413    6916   ------------------------------------------------------------------------------------------
NZ_CP035255.1:776866-784718    6902   ------------------------------------------------------------------------------------------
NZ_CP035256.1:807236-815102    6916   ------------------------------------------------------------------------------------------
NZ_CP038252.1:1151812-1159673  6841   ------------------------------------------------------------------------------------------
NZ_CP035254.1:793365-801213    6898   ------------------------------------------------------------------------------------------
NZ_CP035251.1:748733-756585    6902   ------------------------------------------------------------------------------------------
NZ_CP035252.1:745670-753536    6916   ------------------------------------------------------------------------------------------
```

### Ozkan et al. Supplementary Figures

```
NZ_CP035257.1:776317-784169    6902    ------------------------------------------------------------------------------------------
NZ_CP038253.1:758105-765957    6902    ------------------------------------------------------------------------------------------
NZ_CP035263.1:743050-750901    6901    ------------------------------------------------------------------------------------------
NZ_CP035262.1:741595-749460    6915    ------------------------------------------------------------------------------------------
NZ_CP035236.1:764915-772766    6901    ------------------------------------------------------------------------------------------
NZ_CP035253.1 767286-775134    6898    ------------------------------------------------------------------------------------------
NZ_CP035249.1:763080-770921    6891    ------------------------------------------------------------------------------------------

NC_003028.3:777000-787000      9050    ------------------------------------------------------------------------------------------
NZ_CP035248.1:765099-775111    9062    ------------------------------------------------------------------------------------------
NZ_CP035235.1 c1326834-1317894 7279    CTATCACTGTTCAAAAATGATGGTTGCTGAGACTCCCCTGGTAAAGAAAAATCACCAAATCCCTCGTATCATCAACCAAAAAATTGCTCA
NZ_CP035234.1:756284-765224    7279    CTATCACTGTTCAAAAATGATGGTTGCTGAGACTCCCCTGGTAAAGAAAAATCACCAAATCCCTCGTATCATCAACCAAAAAATTGCTCA
NZ_CP038251.1:750745-759686    7279    CTATCACTGTTCAAAAATGATGGTTGCTGAGACTCCCCTGGTAAAGAAAAATCACCAAATCCCTCGTATCATCAACCAAAAAATTGCTCA
NZ_CP035244.1:747023-755950    7265    CTATCACTGTTCAAAAATGATGGTTGCTGAGACTCCCCTGGTAAAGAAAAATCACCAAATCCCTCGTATCATCAACCAAAAAATTGCTCA
NZ_CP035264.1:800836-809336    7550    ------------------------------------------------------------------------------------------
NZ_CP035265.1:766128-774627    7549    ------------------------------------------------------------------------------------------
NZ_CP035245.1 745261-753757    7546    ------------------------------------------------------------------------------------------
NZ_CP035259.1:745700-753548    6898    ------------------------------------------------------------------------------------------
NZ_CP035258.1:750874-758722    6898    ------------------------------------------------------------------------------------------
NZ_CP035260.1:726156-733994    6888    ------------------------------------------------------------------------------------------
NZ_CP035241.1:708296-716144    6898    ------------------------------------------------------------------------------------------
NZ_CP035261.1:779086-786938    6902    ------------------------------------------------------------------------------------------
NZ_CP035242.1:c1532236-1524385 6901    ------------------------------------------------------------------------------------------
NZ_CP035243.1:777729-785580    6901    ------------------------------------------------------------------------------------------
NZ_CP035237.1:c1311322-1303456 6916    ------------------------------------------------------------------------------------------
NZ_CP035246.1:c1302892-1295026 6916    ------------------------------------------------------------------------------------------
NZ_CP035247.1:c1301115-1293263 6902    ------------------------------------------------------------------------------------------
NZ_CP035238.1:738547-746413    6916    ------------------------------------------------------------------------------------------
NZ_CP035255.1:776866-784718    6902    ------------------------------------------------------------------------------------------
NZ_CP035256.1:807236-815102    6916    ------------------------------------------------------------------------------------------
NZ_CP038252.1:1151812-1159673  6841    ------------------------------------------------------------------------------------------
NZ_CP035254.1:793365-801213    6898    ------------------------------------------------------------------------------------------
NZ_CP035251.1:748733-756585    6902    ------------------------------------------------------------------------------------------
NZ_CP035252.1:745670-753536    6916    ------------------------------------------------------------------------------------------
NZ_CP035257.1:776317-784169    6902    ------------------------------------------------------------------------------------------
NZ_CP038253.1:758105-765957    6902    ------------------------------------------------------------------------------------------
NZ_CP035263.1:743050-750901    6901    ------------------------------------------------------------------------------------------
NZ_CP035262.1:741595-749460    6915    ------------------------------------------------------------------------------------------
NZ_CP035236.1:764915-772766    6901    ------------------------------------------------------------------------------------------
NZ_CP035253.1 767286-775134    6898    ------------------------------------------------------------------------------------------
NZ_CP035249.1:763080-770921    6891    ------------------------------------------------------------------------------------------

NC_003028.3:777000-787000      9050    ------------------------------------------------------------------------------------------
NZ_CP035248.1:765099-775111    9062    ------------------------------------------------------------------------------------------
NZ_CP035235.1 c1326834-1317894 7369    AAAGTTAATTGAAAAAATTTCTATGACTGATATTGCCCATCAGCTGGCCATTTCAACTTCAACTGTCATTCGCAAGCTCAATGACTTTCA
NZ_CP035234.1:756284-765224    7369    AAAGTTAATTGAAAAAATTTCTATGACTGATATTGCCCATCAGCTGGCCATTTCAACTTCAACTGTCATTCGCAAGCTCAATGACTTTCA
NZ_CP038251.1:750745-759686    7369    AAAGTTAATTGAAAAAATTTCTATGACTGATATTGCCCATCAGCTGGCCATTTCAACTTCAACTGTCATTCGCAAGCTCAATGACTTTCA
NZ_CP035244.1:747023-755950    7355    AAAGTTAATTGAAAAAATTTCTATGACTGATATTGCCCATCAGCTGGCCATTTCAACTTCAACTGTCATTCGCAAGCTCAATGACTTTCA
NZ_CP035264.1:800836-809336    7550    ------------------------------------------------------------------------------------------
NZ_CP035265.1:766128-774627    7549    ------------------------------------------------------------------------------------------
NZ_CP035245.1 745261-753757    7546    ------------------------------------------------------------------------------------------
NZ_CP035259.1:745700-753548    6898    ------------------------------------------------------------------------------------------
NZ_CP035258.1:750874-758722    6898    ------------------------------------------------------------------------------------------
NZ_CP035260.1:726156-733994    6888    ------------------------------------------------------------------------------------------
NZ_CP035241.1:708296-716144    6898    ------------------------------------------------------------------------------------------
NZ_CP035261.1:779086-786938    6902    ------------------------------------------------------------------------------------------
NZ_CP035242.1:c1532236-1524385 6901    ------------------------------------------------------------------------------------------
NZ_CP035243.1:777729-785580    6901    ------------------------------------------------------------------------------------------
NZ_CP035237.1:c1311322-1303456 6916    ------------------------------------------------------------------------------------------
NZ_CP035246.1:c1302892-1295026 6916    ------------------------------------------------------------------------------------------
NZ_CP035247.1:c1301115-1293263 6902    ------------------------------------------------------------------------------------------
NZ_CP035238.1:738547-746413    6916    ------------------------------------------------------------------------------------------
NZ_CP035255.1:776866-784718    6902    ------------------------------------------------------------------------------------------
NZ_CP035256.1:807236-815102    6916    ------------------------------------------------------------------------------------------
NZ_CP038252.1:1151812-1159673  6841    ------------------------------------------------------------------------------------------
NZ_CP035254.1:793365-801213    6898    ------------------------------------------------------------------------------------------
NZ_CP035251.1:748733-756585    6902    ------------------------------------------------------------------------------------------
NZ_CP035252.1:745670-753536    6916    ------------------------------------------------------------------------------------------
NZ_CP035257.1:776317-784169    6902    ------------------------------------------------------------------------------------------
NZ_CP038253.1:758105-765957    6902    ------------------------------------------------------------------------------------------
NZ_CP035263.1:743050-750901    6901    ------------------------------------------------------------------------------------------
NZ_CP035262.1:741595-749460    6915    ------------------------------------------------------------------------------------------
NZ_CP035236.1:764915-772766    6901    ------------------------------------------------------------------------------------------
NZ_CP035253.1 767286-775134    6898    ------------------------------------------------------------------------------------------
NZ_CP035249.1:763080-770921    6891    ------------------------------------------------------------------------------------------

NC_003028.3:777000-787000      9050    ------------------------------------------------------------------------------------------
NZ_CP035248.1:765099-775111    9062    ------------------------------------------------------------------------------------------
NZ_CP035235.1 c1326834-1317894 7459    CTTTGAGCATGATTTTTCTCGGCTTCCAAAGATTATGTCTTGGGATGAGTATGCCTTCACTAAGGGAAAGATGAGTTTCATTGCGCAAGA
NZ_CP035234.1:756284-765224    7459    CTTTGAGCATGATTTTTCTCGGCTTCCAAAGATTATGTCTTGGGATGAGTATGCCTTCACTAAGGGAAAGATGAGTTTCATTGCGCAAGA
NZ_CP038251.1:750745-759686    7459    CTTTGAGCATGATTTTTCTCGGCTTCCAAAGATTATGTCTTGGGATGAGTATGCCTTCACTAAGGGAAAGATGAGTTTCATTGCGCAAGA
NZ_CP035244.1:747023-755950    7445    CTTTGAGCATGATTTTTCTCGGCTTCCAAAGATTATGTCTTGGGATGAGTATGCCTTCACTAAGGGAAAGATGAGTTTCATTGCGCAAGA
NZ_CP035264.1:800836-809336    7550    ------------------------------------------------------------------------------------------
NZ_CP035265.1:766128-774627    7549    ------------------------------------------------------------------------------------------
NZ_CP035245.1 745261-753757    7546    ------------------------------------------------------------------------------------------
NZ_CP035259.1:745700-753548    6898    ------------------------------------------------------------------------------------------
NZ_CP035258.1:750874-758722    6898    ------------------------------------------------------------------------------------------
NZ_CP035260.1:726156-733994    6888    ------------------------------------------------------------------------------------------
NZ_CP035241.1:708296-716144    6898    ------------------------------------------------------------------------------------------
NZ_CP035261.1:779086-786938    6902    ------------------------------------------------------------------------------------------
NZ_CP035242.1:c1532236-1524385 6901    ------------------------------------------------------------------------------------------
NZ_CP035243.1:777729-785580    6901    ------------------------------------------------------------------------------------------
NZ_CP035237.1:c1311322-1303456 6916    ------------------------------------------------------------------------------------------
NZ_CP035246.1:c1302892-1295026 6916    ------------------------------------------------------------------------------------------
NZ_CP035247.1:c1301115-1293263 6902    ------------------------------------------------------------------------------------------
```

### Ozkan et al. Supplementary Figures

```
NZ_CP035238.1:738547-746413    6916   ------------------------------------------------------------------------------------------
NZ_CP035255.1:776866-784718    6902   ------------------------------------------------------------------------------------------
NZ_CP035256.1:807236-815102    6916   ------------------------------------------------------------------------------------------
NZ_CP038252.1:1151812-1159673  6841   ------------------------------------------------------------------------------------------
NZ_CP035254.1:793365-801213    6898   ------------------------------------------------------------------------------------------
NZ_CP035251.1:748733-756585    6902   ------------------------------------------------------------------------------------------
NZ_CP035252.1:745670-753536    6916   ------------------------------------------------------------------------------------------
NZ_CP035257.1:776317-784169    6902   ------------------------------------------------------------------------------------------
NZ_CP038253.1:758105-765957    6902   ------------------------------------------------------------------------------------------
NZ_CP035263.1:743050-750901    6901   ------------------------------------------------------------------------------------------
NZ_CP035262.1:741595-749460    6915   ------------------------------------------------------------------------------------------
NZ_CP035236.1:764915-772766    6901   ------------------------------------------------------------------------------------------
NZ_CP035253.1 767286-775134    6898   ------------------------------------------------------------------------------------------
NZ_CP035249.1:763080-770921    6891   ------------------------------------------------------------------------------------------

NC_003028.3:777000-787000      9050   ------------------------------------------------------------------------------------------
NZ_CP035248.1:765099-775111    9062   ------------------------------------------------------------------------------------------
NZ_CP035235.1 c1326834-1317894 7549   TTTTGATAATCTTAATATTATCACTGTTCTTGAAGGCAGAACACAAGCTGTCATCCGAAATCACTTTCTTCGCTACGATAGAGCCGTTCG
NZ_CP035234.1:756284-765224    7549   TTTTGATAATCTTAATATTATCACTGTTCTTGAAGGCAGAACACAAGCTGTCATCCGAAATCACTTTCTTCGCTACGATAGAGCCGTTCG
NZ_CP038251.1:750745-759686    7549   TTTTGATAATCTTAATATTATCACTGTTCTTGAAGGCAGAACACAAGCTGTCATCCGAAATCACTTTCTTCGCTACGATAGAGCCGTTCG
NZ_CP035244.1:747023-755950    7535   TTTTGATAATCTTAATATTATCACTGTTCTTGAAGGCAGAACACAAGCTGTCATCCGAAATCACTTTCTTCGCTACGATAGAGTCGTTCG
NZ_CP035264.1:800836-809336    7550   ------------------------------------------------------------------------------------------
NZ_CP035265.1:766128-774627    7549   ------------------------------------------------------------------------------------------
NZ_CP035245.1 745261-753757    7546   ------------------------------------------------------------------------------------------
NZ_CP035259.1:745700-753548    6898   ------------------------------------------------------------------------------------------
NZ_CP035258.1:750874-758722    6898   ------------------------------------------------------------------------------------------
NZ_CP035260.1:726156-733994    6888   ------------------------------------------------------------------------------------------
NZ_CP035241.1:708296-716144    6898   ------------------------------------------------------------------------------------------
NZ_CP035261.1:779086-786938    6902   ------------------------------------------------------------------------------------------
NZ_CP035242.1:c1532236-1524385 6901   ------------------------------------------------------------------------------------------
NZ_CP035243.1:777729-785580    6901   ------------------------------------------------------------------------------------------
NZ_CP035237.1:c1311322-1303456 6916   ------------------------------------------------------------------------------------------
NZ_CP035246.1:c1302892-1295026 6916   ------------------------------------------------------------------------------------------
NZ_CP035247.1:c1301115-1293263 6902   ------------------------------------------------------------------------------------------
NZ_CP035238.1:738547-746413    6916   ------------------------------------------------------------------------------------------
NZ_CP035255.1:776866-784718    6902   ------------------------------------------------------------------------------------------
NZ_CP035256.1:807236-815102    6916   ------------------------------------------------------------------------------------------
NZ_CP038252.1:1151812-1159673  6841   ------------------------------------------------------------------------------------------
NZ_CP035254.1:793365-801213    6898   ------------------------------------------------------------------------------------------
NZ_CP035251.1:748733-756585    6902   ------------------------------------------------------------------------------------------
NZ_CP035252.1:745670-753536    6916   ------------------------------------------------------------------------------------------
NZ_CP035257.1:776317-784169    6902   ------------------------------------------------------------------------------------------
NZ_CP038253.1:758105-765957    6902   ------------------------------------------------------------------------------------------
NZ_CP035263.1:743050-750901    6901   ------------------------------------------------------------------------------------------
NZ_CP035262.1:741595-749460    6915   ------------------------------------------------------------------------------------------
NZ_CP035236.1:764915-772766    6901   ------------------------------------------------------------------------------------------
NZ_CP035253.1 767286-775134    6898   ------------------------------------------------------------------------------------------
NZ_CP035249.1:763080-770921    6891   ------------------------------------------------------------------------------------------

NC_003028.3:777000-787000      9050   ------------------------------------------------------------------------------------------
NZ_CP035248.1:765099-775111    9062   ------------------------------------------------------------------------------------------
NZ_CP035235.1 c1326834-1317894 7639   TTGTCAAGTGAAAATCATTACGATGGATATGTTTAGTCCTTACTATGACTTGGCTAAACAGCTTTTTCCGTGTGCTAAAATCGTTCTAGA
NZ_CP035234.1:756284-765224    7639   TTGTCAAGTGAAAATCATTACGATGGATATGTTTAGTCCTTACTATGACTTGGCTAAACAGCTTTTTCCGTGTGCTAAAATCGTTCTAGA
NZ_CP038251.1:750745-759686    7639   TTGTCAAGTGAAAATCATTACGATGGATATGTTTAGTCCTTACTATGACTTGGCTAAACAGCTTTTTCCGTGTGCTAAAATCGTTCTAGA
NZ_CP035244.1:747023-755950    7625   TTGTCAAGTGAAAATCATTACGATGGATATGTTTAGTCCTTACTATGACTTGGCTAAACAGCTTTTTCCGTGTGCTAAAATCGTTCTAGA
NZ_CP035264.1:800836-809336    7550   ------------------------------------------------------------------------------------------
NZ_CP035265.1:766128-774627    7549   ------------------------------------------------------------------------------------------
NZ_CP035245.1 745261-753757    7546   ------------------------------------------------------------------------------------------
NZ_CP035259.1:745700-753548    6898   ------------------------------------------------------------------------------------------
NZ_CP035258.1:750874-758722    6898   ------------------------------------------------------------------------------------------
NZ_CP035260.1:726156-733994    6888   ------------------------------------------------------------------------------------------
NZ_CP035241.1:708296-716144    6898   ------------------------------------------------------------------------------------------
NZ_CP035261.1:779086-786938    6902   ------------------------------------------------------------------------------------------
NZ_CP035242.1:c1532236-1524385 6901   ------------------------------------------------------------------------------------------
NZ_CP035243.1:777729-785580    6901   ------------------------------------------------------------------------------------------
NZ_CP035237.1:c1311322-1303456 6916   ------------------------------------------------------------------------------------------
NZ_CP035246.1:c1302892-1295026 6916   ------------------------------------------------------------------------------------------
NZ_CP035247.1:c1301115-1293263 6902   ------------------------------------------------------------------------------------------
NZ_CP035238.1:738547-746413    6916   ------------------------------------------------------------------------------------------
NZ_CP035255.1:776866-784718    6902   ------------------------------------------------------------------------------------------
NZ_CP035256.1:807236-815102    6916   ------------------------------------------------------------------------------------------
NZ_CP038252.1:1151812-1159673  6841   ------------------------------------------------------------------------------------------
NZ_CP035254.1:793365-801213    6898   ------------------------------------------------------------------------------------------
NZ_CP035251.1:748733-756585    6902   ------------------------------------------------------------------------------------------
NZ_CP035252.1:745670-753536    6916   ------------------------------------------------------------------------------------------
NZ_CP035257.1:776317-784169    6902   ------------------------------------------------------------------------------------------
NZ_CP038253.1:758105-765957    6902   ------------------------------------------------------------------------------------------
NZ_CP035263.1:743050-750901    6901   ------------------------------------------------------------------------------------------
NZ_CP035262.1:741595-749460    6915   ------------------------------------------------------------------------------------------
NZ_CP035236.1:764915-772766    6901   ------------------------------------------------------------------------------------------
NZ_CP035253.1 767286-775134    6898   ------------------------------------------------------------------------------------------
NZ_CP035249.1:763080-770921    6891   ------------------------------------------------------------------------------------------

NC_003028.3:777000-787000      9050   ------------------------------------------------------------------------------------------
NZ_CP035248.1:765099-775111    9062   ------------------------------------------------------------------------------------------
NZ_CP035235.1 c1326834-1317894 7729   TCGTTTCCATATTATCCAACATCTCAGCCGTGCCATGAGTCGTTTTCGTGTTCAAATTATGAATCAGTTTGAACGAAAATCTCATGAATA
NZ_CP035234.1:756284-765224    7729   TCGTTTCCATATTATCCAACATCTCAGCCGTGCCATGAGTCGTTTTCGTGTTCAAATTATGAATCAGTTTGAACGAAAATCTCATGAATA
NZ_CP038251.1:750745-759686    7729   TCGTTTCCATATTATCCAACATCTCAGCCGTGCCATGAGTCGTTTTCGTGTTCAAATTATGAATCAGTTTGAACGAAAATCTCATGAATA
NZ_CP035244.1:747023-755950    7715   TCGTTTCCATATTATCCAACATCTCAGCCGTGCCATGAGTCGTTTTCGTGTTCAAATTATGAATCAGTTTGAACGAAAATCTCATGAATA
NZ_CP035264.1:800836-809336    7550   ------------------------------------------------------------------------------------------
NZ_CP035265.1:766128-774627    7549   ------------------------------------------------------------------------------------------
NZ_CP035245.1 745261-753757    7546   ------------------------------------------------------------------------------------------
NZ_CP035259.1:745700-753548    6898   ------------------------------------------------------------------------------------------
NZ_CP035258.1:750874-758722    6898   ------------------------------------------------------------------------------------------
NZ_CP035260.1:726156-733994    6888   ------------------------------------------------------------------------------------------
```

```
NZ_CP035241.1:708296-716144    6898   ------------------------------------------------------------------------------------------
NZ_CP035261.1:779086-786938    6902   ------------------------------------------------------------------------------------------
NZ_CP035242.1:c1532236-1524385 6901   ------------------------------------------------------------------------------------------
NZ_CP035243.1:777729-785580    6901   ------------------------------------------------------------------------------------------
NZ_CP035237.1:c1311322-1303456 6916   ------------------------------------------------------------------------------------------
NZ_CP035246.1:c1302892-1295026 6916   ------------------------------------------------------------------------------------------
NZ_CP035247.1:c1301115-1293263 6902   ------------------------------------------------------------------------------------------
NZ_CP035238.1:738547-746413    6916   ------------------------------------------------------------------------------------------
NZ_CP035255.1:776866-784718    6902   ------------------------------------------------------------------------------------------
NZ_CP035256.1:807236-815102    6916   ------------------------------------------------------------------------------------------
NZ_CP038252.1:1151812-1159673  6841   ------------------------------------------------------------------------------------------
NZ_CP035254.1:793365-801213    6898   ------------------------------------------------------------------------------------------
NZ_CP035251.1:748733-756585    6902   ------------------------------------------------------------------------------------------
NZ_CP035252.1:745670-753536    6916   ------------------------------------------------------------------------------------------
NZ_CP035257.1:776317-784169    6902   ------------------------------------------------------------------------------------------
NZ_CP038253.1:758105-765957    6902   ------------------------------------------------------------------------------------------
NZ_CP035263.1:743050-750901    6901   ------------------------------------------------------------------------------------------
NZ_CP035262.1:741595-749460    6915   ------------------------------------------------------------------------------------------
NZ_CP035236.1:764915-772766    6901   ------------------------------------------------------------------------------------------
NZ_CP035253.1 767286-775134    6898   ------------------------------------------------------------------------------------------
NZ_CP035249.1:763080-770921    6891   ------------------------------------------------------------------------------------------

NC_003028.3:777000-787000      9050   ------------------------------------------------------------------------------------------
NZ_CP035248.1:765099-775111    9062   ------------------------------------------------------------------------------------------
NZ_CP035235.1 c1326834-1317894 7819   CAAGGCTATCAAGCGTTACTGGAAACTCATCCAACAGGATAGTCGTAAACTCAGCGATAAACGTTTTTATCGCCCTACTTTTCGCATACA
NZ_CP035234.1:756284-765224    7819   CAAGGCTATCAAGCGTTACTGGAAACTCATCCAACAGGATAGTCGTAAACTCAGCGATAAACGTTTTTATCGCCCTACTTTTCGCATACA
NZ_CP038251.1:750745-759686    7819   CAAGGCTATCAAGCGTTACTGGAAACTCATCCAACAGGATAGTCGTAAACTCAGCGATAAACGTTTTTATCGCCCTACTTTTCGCATACA
NZ_CP035244.1:747023-755950    7805   CAAGGCTATCAAGCGTTACTGGAAACTCATCCAACAGGATAGTCGTAAACTCAGCGATAAACGTTTTTATCGCCCTACTTTTCGCATACA
NZ_CP035264.1:800836-809336    7550   ------------------------------------------------------------------------------------------
NZ_CP035265.1:766128-774627    7549   ------------------------------------------------------------------------------------------
NZ_CP035245.1 745261-753757    7546   ------------------------------------------------------------------------------------------
NZ_CP035259.1:745700-753548    6898   ------------------------------------------------------------------------------------------
NZ_CP035258.1:750874-758722    6898   ------------------------------------------------------------------------------------------
NZ_CP035260.1:726156-733994    6888   ------------------------------------------------------------------------------------------
NZ_CP035241.1:708296-716144    6898   ------------------------------------------------------------------------------------------
NZ_CP035261.1:779086-786938    6902   ------------------------------------------------------------------------------------------
NZ_CP035242.1:c1532236-1524385 6901   ------------------------------------------------------------------------------------------
NZ_CP035243.1:777729-785580    6901   ------------------------------------------------------------------------------------------
NZ_CP035237.1:c1311322-1303456 6916   ------------------------------------------------------------------------------------------
NZ_CP035246.1:c1302892-1295026 6916   ------------------------------------------------------------------------------------------
NZ_CP035247.1:c1301115-1293263 6902   ------------------------------------------------------------------------------------------
NZ_CP035238.1:738547-746413    6916   ------------------------------------------------------------------------------------------
NZ_CP035255.1:776866-784718    6902   ------------------------------------------------------------------------------------------
NZ_CP035256.1:807236-815102    6916   ------------------------------------------------------------------------------------------
NZ_CP038252.1:1151812-1159673  6841   ------------------------------------------------------------------------------------------
NZ_CP035254.1:793365-801213    6898   ------------------------------------------------------------------------------------------
NZ_CP035251.1:748733-756585    6902   ------------------------------------------------------------------------------------------
NZ_CP035252.1:745670-753536    6916   ------------------------------------------------------------------------------------------
NZ_CP035257.1:776317-784169    6902   ------------------------------------------------------------------------------------------
NZ_CP038253.1:758105-765957    6902   ------------------------------------------------------------------------------------------
NZ_CP035263.1:743050-750901    6901   ------------------------------------------------------------------------------------------
NZ_CP035262.1:741595-749460    6915   ------------------------------------------------------------------------------------------
NZ_CP035236.1:764915-772766    6901   ------------------------------------------------------------------------------------------
NZ_CP035253.1 767286-775134    6898   ------------------------------------------------------------------------------------------
NZ_CP035249.1:763080-770921    6891   ------------------------------------------------------------------------------------------

NC_003028.3:777000-787000      9050   ------------------------------------------------------------------------------------------
NZ_CP035248.1:765099-775111    9062   ------------------------------------------------------------------------------------------
NZ_CP035235.1 c1326834-1317894 7909   CTTAACAAATAAAGAAATTCTTGACAAGATTTTAAGCTATTCAGAAGACTTGAAACACCACTATCAGATCTATCAACTCTTACTTTTTCA
NZ_CP035234.1:756284-765224    7909   CTTAACAAATAAAGAAATTCTTGACAAGATTTTAAGCTATTCAGAAGACTTGAAACACCACTATCAGATCTATCAACTCTTACTTTTTCA
NZ_CP038251.1:750745-759686    7909   CTTAACAAATAAAGAAATTCTTGACAAGATTTTAAGCTATTCAGAAGACTTGAAACACCACTATCAGATCTATCAACTCTTACTTTTTCA
NZ_CP035244.1:747023-755950    7895   CTTAACAAATAAAGAAATTCTTGACAAGATTTTAAGCTATTCAGAAGACTTGAAACACCACTATCAGATCTATCAACTCTTACTTTTTCA
NZ_CP035264.1:800836-809336    7550   ------------------------------------------------------------------------------------------
NZ_CP035265.1:766128-774627    7549   ------------------------------------------------------------------------------------------
NZ_CP035245.1 745261-753757    7546   ------------------------------------------------------------------------------------------
NZ_CP035259.1:745700-753548    6898   ------------------------------------------------------------------------------------------
NZ_CP035258.1:750874-758722    6898   ------------------------------------------------------------------------------------------
NZ_CP035260.1:726156-733994    6888   ------------------------------------------------------------------------------------------
NZ_CP035241.1:708296-716144    6898   ------------------------------------------------------------------------------------------
NZ_CP035261.1:779086-786938    6902   ------------------------------------------------------------------------------------------
NZ_CP035242.1:c1532236-1524385 6901   ------------------------------------------------------------------------------------------
NZ_CP035243.1:777729-785580    6901   ------------------------------------------------------------------------------------------
NZ_CP035237.1:c1311322-1303456 6916   ------------------------------------------------------------------------------------------
NZ_CP035246.1:c1302892-1295026 6916   ------------------------------------------------------------------------------------------
NZ_CP035247.1:c1301115-1293263 6902   ------------------------------------------------------------------------------------------
NZ_CP035238.1:738547-746413    6916   ------------------------------------------------------------------------------------------
NZ_CP035255.1:776866-784718    6902   ------------------------------------------------------------------------------------------
NZ_CP035256.1:807236-815102    6916   ------------------------------------------------------------------------------------------
NZ_CP038252.1:1151812-1159673  6841   ------------------------------------------------------------------------------------------
NZ_CP035254.1:793365-801213    6898   ------------------------------------------------------------------------------------------
NZ_CP035251.1:748733-756585    6902   ------------------------------------------------------------------------------------------
NZ_CP035252.1:745670-753536    6916   ------------------------------------------------------------------------------------------
NZ_CP035257.1:776317-784169    6902   ------------------------------------------------------------------------------------------
NZ_CP038253.1:758105-765957    6902   ------------------------------------------------------------------------------------------
NZ_CP035263.1:743050-750901    6901   ------------------------------------------------------------------------------------------
NZ_CP035262.1:741595-749460    6915   ------------------------------------------------------------------------------------------
NZ_CP035236.1:764915-772766    6901   ------------------------------------------------------------------------------------------
NZ_CP035253.1 767286-775134    6898   ------------------------------------------------------------------------------------------
NZ_CP035249.1:763080-770921    6891   ------------------------------------------------------------------------------------------

NC_003028.3:777000-787000      9050   ------------------------------------------------------------------------------------------
NZ_CP035248.1:765099-775111    9062   ------------------------------------------------------------------------------------------
NZ_CP035235.1 c1326834-1317894 7999   CTTTCAGAACAAAGACCCTGAGAAATTTTTCGGACTCATTGAGGACAATCTGAAGCAGGTTCATCCTATTTTTCAGACTGTCTTTAAAAC
NZ_CP035234.1:756284-765224    7999   CTTTCAGAACAAAGACCCTGAGAAATTTTTCGGACTCATTGAGGACAATCTGAAGCAGGTTCATCCTATTTTTCAGACTGTCTTTAAAAC
NZ_CP038251.1:750745-759686    7999   CTTTCAGAACAAAGACCCTGAGAAATTTTTCGGACTCATTGAGGACAATCTGAAGCAGGTTCATCCTATTTTTCAGACTGTCTTTAAAAC
```

### Ozkan et al. Supplementary Figures

```
NZ_CP035244.1:747023-755950    7985   CTTTCAGAACAAAGACCCTGAGAAATTTTTCGGACTCATTGAGGACAATCTGAAGCAGGTTCATCCTATTTTTCAGACTGTCTTTAAAAC
NZ_CP035264.1:800836-809336    7550   ------------------------------------------------------------------------------------------
NZ_CP035265.1:766128-774627    7549   ------------------------------------------------------------------------------------------
NZ_CP035245.1 745261-753757    7546   ------------------------------------------------------------------------------------------
NZ_CP035259.1:745700-753548    6898   ------------------------------------------------------------------------------------------
NZ_CP035258.1:750874-758722    6898   ------------------------------------------------------------------------------------------
NZ_CP035260.1:726156-733994    6888   ------------------------------------------------------------------------------------------
NZ_CP035241.1:708296-716144    6898   ------------------------------------------------------------------------------------------
NZ_CP035261.1:779086-786938    6902   ------------------------------------------------------------------------------------------
NZ_CP035242.1:c1532236-1524385 6901   ------------------------------------------------------------------------------------------
NZ_CP035243.1:777729-785580    6901   ------------------------------------------------------------------------------------------
NZ_CP035237.1:c1311322-1303456 6916   ------------------------------------------------------------------------------------------
NZ_CP035246.1:c1302892-1295026 6916   ------------------------------------------------------------------------------------------
NZ_CP035247.1:c1301115-1293263 6902   ------------------------------------------------------------------------------------------
NZ_CP035238.1:738547-746413    6916   ------------------------------------------------------------------------------------------
NZ_CP035255.1:776866-784718    6902   ------------------------------------------------------------------------------------------
NZ_CP035256.1:807236-815102    6916   ------------------------------------------------------------------------------------------
NZ_CP038252.1:1151812-1159673  6841   ------------------------------------------------------------------------------------------
NZ_CP035254.1:793365-801213    6898   ------------------------------------------------------------------------------------------
NZ_CP035251.1:748733-756585    6902   ------------------------------------------------------------------------------------------
NZ_CP035252.1:745670-753536    6916   ------------------------------------------------------------------------------------------
NZ_CP035257.1:776317-784169    6902   ------------------------------------------------------------------------------------------
NZ_CP038253.1:758105-765957    6902   ------------------------------------------------------------------------------------------
NZ_CP035263.1:743050-750901    6901   ------------------------------------------------------------------------------------------
NZ_CP035262.1:741595-749460    6915   ------------------------------------------------------------------------------------------
NZ_CP035236.1:764915-772766    6901   ------------------------------------------------------------------------------------------
NZ_CP035253.1 767286-775134    6898   ------------------------------------------------------------------------------------------
NZ_CP035249.1:763080-770921    6891   ------------------------------------------------------------------------------------------

NC_003028.3:777000-787000      9050   ------------------------------------------------------------------------------------------
NZ_CP035248.1:765099-775111    9062   ------------------------------------------------------------------------------------------
NZ_CP035235.1 c1326834-1317894 8089   CTTTCTAAAGAACAAAGAAAAAATCGTCAATGCTCTTCAATTACCTTATTCCAATGCAAAATTGGAAGCGACCAATAATCTCATCAAACT
NZ_CP035234.1:756284-765224    8089   CTTTCTAAAGAACAAAGAAAAAATCGTCAATGCTCTTCAATTACCTTATTCCAATGCAAAATTGGAAGCGACCAATAATCTCATCAAACT
NZ_CP038251.1:750745-759686    8089   CTTTCTAAAGAACAAAGAAAAAATCGTCAATGCTCTTCAATTACCTTATTCCAATGCAAAATTGGAAGCGACCAATAATCTCATCAAACT
NZ_CP035244.1:747023-755950    8075   CTTTCTAAAGAACAAAGAAAAAATCGTCAATGCTCTTCAATTACCTTATTCCAATGCAAAATTGGAAGCGACCAATAATCTCATCAAACT
NZ_CP035264.1:800836-809336    7550   ------------------------------------------------------------------------------------------
NZ_CP035265.1:766128-774627    7549   ------------------------------------------------------------------------------------------
NZ_CP035245.1 745261-753757    7546   ------------------------------------------------------------------------------------------
NZ_CP035259.1:745700-753548    6898   ------------------------------------------------------------------------------------------
NZ_CP035258.1:750874-758722    6898   ------------------------------------------------------------------------------------------
NZ_CP035260.1:726156-733994    6888   ------------------------------------------------------------------------------------------
NZ_CP035241.1:708296-716144    6898   ------------------------------------------------------------------------------------------
NZ_CP035261.1:779086-786938    6902   ------------------------------------------------------------------------------------------
NZ_CP035242.1:c1532236-1524385 6901   ------------------------------------------------------------------------------------------
NZ_CP035243.1:777729-785580    6901   ------------------------------------------------------------------------------------------
NZ_CP035237.1:c1311322-1303456 6916   ------------------------------------------------------------------------------------------
NZ_CP035246.1:c1302892-1295026 6916   ------------------------------------------------------------------------------------------
NZ_CP035247.1:c1301115-1293263 6902   ------------------------------------------------------------------------------------------
NZ_CP035238.1:738547-746413    6916   ------------------------------------------------------------------------------------------
NZ_CP035255.1:776866-784718    6902   ------------------------------------------------------------------------------------------
NZ_CP035256.1:807236-815102    6916   ------------------------------------------------------------------------------------------
NZ_CP038252.1:1151812-1159673  6841   ------------------------------------------------------------------------------------------
NZ_CP035254.1:793365-801213    6898   ------------------------------------------------------------------------------------------
NZ_CP035251.1:748733-756585    6902   ------------------------------------------------------------------------------------------
NZ_CP035252.1:745670-753536    6916   ------------------------------------------------------------------------------------------
NZ_CP035257.1:776317-784169    6902   ------------------------------------------------------------------------------------------
NZ_CP038253.1:758105-765957    6902   ------------------------------------------------------------------------------------------
NZ_CP035263.1:743050-750901    6901   ------------------------------------------------------------------------------------------
NZ_CP035262.1:741595-749460    6915   ------------------------------------------------------------------------------------------
NZ_CP035236.1:764915-772766    6901   ------------------------------------------------------------------------------------------
NZ_CP035253.1 767286-775134    6898   ------------------------------------------------------------------------------------------
NZ_CP035249.1:763080-770921    6891   ------------------------------------------------------------------------------------------

NC_003028.3:777000-787000      9050   ------------------------------------------------------------------------------------------
NZ_CP035248.1:765099-775111    9062   ------------------------------------------------------------------------------------------
NZ_CP035235.1 c1326834-1317894 8179   TATCAAACGAAACGCCTTTGGATTTCGGAACTTTGAAAACTTCAAAAAAAGGATTTTTATCGCTCTGAACATCAAAAAAGAAAGGACGAA
NZ_CP035234.1:756284-765224    8179   TATCAAACGAAACGCCTTTGGATTTCGGAACTTTGAAAACTTCAAAAAAAGGATTTTTATCGCTCTGAACATCAAAAAAGAAAGGACGAA
NZ_CP038251.1:750745-759686    8179   TATCAAACGAAACGCCTTTGGATTTCGGAACTTTGAAAACTTCAAAAAAAGGATTTTTATCGCTCTGAACATCAAAAAAGAAAGGACGAA
NZ_CP035244.1:747023-755950    8165   TATCAAACGAAACGCCTTTGGATTTCGGAACTTTGAAAACTTCAAAAAAAGGATTTTTATCGCTCTGAACATCAAAAAAGAAAGGACGAA
NZ_CP035264.1:800836-809336    7550   ------------------------------------------------------------------------------------------
NZ_CP035265.1:766128-774627    7549   ------------------------------------------------------------------------------------------
NZ_CP035245.1 745261-753757    7546   ------------------------------------------------------------------------------------------
NZ_CP035259.1:745700-753548    6898   ------------------------------------------------------------------------------------------
NZ_CP035258.1:750874-758722    6898   ------------------------------------------------------------------------------------------
NZ_CP035260.1:726156-733994    6888   ------------------------------------------------------------------------------------------
NZ_CP035241.1:708296-716144    6898   ------------------------------------------------------------------------------------------
NZ_CP035261.1:779086-786938    6902   ------------------------------------------------------------------------------------------
NZ_CP035242.1:c1532236-1524385 6901   ------------------------------------------------------------------------------------------
NZ_CP035243.1:777729-785580    6901   ------------------------------------------------------------------------------------------
NZ_CP035237.1:c1311322-1303456 6916   ------------------------------------------------------------------------------------------
NZ_CP035246.1:c1302892-1295026 6916   ------------------------------------------------------------------------------------------
NZ_CP035247.1:c1301115-1293263 6902   ------------------------------------------------------------------------------------------
NZ_CP035238.1:738547-746413    6916   ------------------------------------------------------------------------------------------
NZ_CP035255.1:776866-784718    6902   ------------------------------------------------------------------------------------------
NZ_CP035256.1:807236-815102    6916   ------------------------------------------------------------------------------------------
NZ_CP038252.1:1151812-1159673  6841   ------------------------------------------------------------------------------------------
NZ_CP035254.1:793365-801213    6898   ------------------------------------------------------------------------------------------
NZ_CP035251.1:748733-756585    6902   ------------------------------------------------------------------------------------------
NZ_CP035252.1:745670-753536    6916   ------------------------------------------------------------------------------------------
NZ_CP035257.1:776317-784169    6902   ------------------------------------------------------------------------------------------
NZ_CP038253.1:758105-765957    6902   ------------------------------------------------------------------------------------------
NZ_CP035263.1:743050-750901    6901   ------------------------------------------------------------------------------------------
NZ_CP035262.1:741595-749460    6915   ------------------------------------------------------------------------------------------
NZ_CP035236.1:764915-772766    6901   ------------------------------------------------------------------------------------------
NZ_CP035253.1 767286-775134    6898   ------------------------------------------------------------------------------------------
```

```
NZ_CP035249.1:763080-770921    6891    ------------------------------------------------------------------------------------------

NC_003028.3:777000-787000      9050    --------------------------------------------------------AAGAGACAGGTCAATGGCGAAAAATTGAGAAAGA
NZ_CP035248.1:765099-775111    9062    --------------------------------------------------------AAGAGACAGGTCAATGGCGAAAAATTGAGAAAGA
NZ_CP035235.1 c1326834-1317894 8269   ATTTGTCCTTTCTCAAGCTTAGC-TGACTTCAACCCACTACAGTTGACAAAGAGCC----------------------------------
NZ_CP035234.1:756284-765224    8269    ATTTGTCCTTTCTCAAGCTTAGC-TGACTTCAACCCACTACAGTTGACAAAGAGCC----------------------------------
NZ_CP038251.1:750745-759686    8269    ATTTGTCCTTTCTCAAGCTTAGCTTTTCTTCAACCCACTACAGTTGACAAAGAGCC----------------------------------
NZ_CP035244.1:747023-755950    8255    ATTTGTCCTTTCTCAAGCTTAGCTTTTCTTCAACCCACTACAGTTGACAAAGAGCC----------------------------------
NZ_CP035264.1:800836-809336    7550    --------------------------------------------------------AAGAGACAGGTCAATGGCGAAAAATTGAGAAAGA
NZ_CP035265.1:766128-774627    7549    --------------------------------------------------------AAGAGACAGGTCAATGGCGAAAAATTGAGAAAGA
NZ_CP035245.1 745261-753757    7546    --------------------------------------------------------AAGAGACAGGTCAATGGCGAAAAATTGAGAAAGA
NZ_CP035259.1:745700-753548    6898    --------------------------------------------------------AAGAGACAGGTCAATGGCAAAAAATTGAGAAAGA
NZ_CP035258.1:750874-758722    6898    --------------------------------------------------------AAGAGACAGGTCAATGGCAAAAAATTGAGAAAGA
NZ_CP035260.1:726156-733994    6888    --------------------------------------------------------AAGAGACAGGTCAATGGCAAAAAATTGAGAAAGA
NZ_CP035241.1:708296-716144    6898    --------------------------------------------------------AAGAGACAGGTCAATGGCAAAAAATTGAGAAAGA
NZ_CP035261.1:779086-786938    6902    --------------------------------------------------------AAGAGACAGGTCAATGGCAAAAAATTGAGAAAGA
NZ_CP035242.1:c1532236-1524385 6901    --------------------------------------------------------AAGAGACAGGTCAATGGCGAAAAATTGAGAAAGA
NZ_CP035243.1:777729-785580    6901    --------------------------------------------------------AAGAGACAGGTCAATGGCGAAAAATTGAGAAAGA
NZ_CP035237.1:c1311322-1303456 6916    --------------------------------------------------------AAGAGACAGGTCAATGGCAAAAAATTGAGAAAGA
NZ_CP035246.1:c1302892-1295026 6916    --------------------------------------------------------AAGAGACAGGTTAATGGCGAAAAATTGAGAAAGA
NZ_CP035247.1:c1301115-1293263 6902    --------------------------------------------------------AAGAGACAGGTTAATGGCGAAAAATTGAGAAAGA
NZ_CP035238.1:738547-746413    6916    --------------------------------------------------------AAGAGACAGGTCAATGGCAAAAAATTGAGAAAGA
NZ_CP035255.1:776866-784718    6902    --------------------------------------------------------AAGAGACAGGTCAATGGCAAAAAATTGAGAAAGA
NZ_CP035256.1:807236-815102    6916    --------------------------------------------------------AAGAGACAGGTCAATGGCAAAAAATTGAGAAAGA
NZ_CP038252.1:1151812-1159673  6841    --------------------------------------------------------AAGAGACAGGTCAATGGCAAAAAATTGAGAAAGA
NZ_CP035254.1:793365-801213    6898    --------------------------------------------------------AAGAGACAGGTCAATGGCAAAAAATTGAGAAAGA
NZ_CP035251.1:748733-756585    6902    --------------------------------------------------------AAGAGACAGGTCAATGGCAAAAAATTGAGAAAGA
NZ_CP035252.1:745670-753536    6916    --------------------------------------------------------AAGAGACAGGTCAATGGCAAAAAATTGAGAAAGA
NZ_CP035257.1:776317-784169    6902    --------------------------------------------------------AAGAGACAGGTCAATGGCAAAAAATTGAGAAAGA
NZ_CP038253.1:758105-765957    6902    --------------------------------------------------------AAGAGACAGGTCAATGGCAAAAAATTGAGAAAGA
NZ_CP035263.1:743050-750901    6901    --------------------------------------------------------AAGAGACAGGTCAATGGCGAAAAATTGAGAAAGA
NZ_CP035262.1:741595-749460    6915    --------------------------------------------------------AAGAGACAGGTCAATGGCGAAAAATTGAGAAAGA
NZ_CP035236.1:764915-772766    6901    --------------------------------------------------------AAGAGACAGGTCAATGGCGAAAAATTGAGAAAGA
NZ_CP035253.1 767286-775134    6898    --------------------------------------------------------AAGAGACAGGTCAATGGCAAAAAATTGAGAAAGA
NZ_CP035249.1:763080-770921    6891    --------------------------------------------------------AAGAGACAGGTCAATGGCGAAAAATTGAGAAAGA

NC_003028.3:777000-787000      9085    TGATTTGGTCAGCTTCTTGCATTCGTTCTTGGTAGTAGCACCAAGAATAATTACCATCGATGACCCAAGCTTTATGCTTGGTGAGAAAGT
NZ_CP035248.1:765099-775111    9097    TGATTTGGTCAGCTTCTTGCATTCGTTCTTGGTAGTAGCACCAAGAATAATTACCATCGATGACCCAAGCTTTATGCTTGGTGAGAAAGT
NZ_CP035235.1 c1326834-1317894 8323   ------------------------------------------------------------------------------------------
NZ_CP035234.1:756284-765224    8323    ------------------------------------------------------------------------------------------
NZ_CP038251.1:750745-759686    8324    ------------------------------------------------------------------------------------------
NZ_CP035244.1:747023-755950    8310    ------------------------------------------------------------------------------------------
NZ_CP035264.1:800836-809336    7585    TGATTTGGTCAGCTTCTTGCATTCGTTCTTGGTAGTAGCACCAAGAATAATTACCATCGATGACCCAAGCTTTATGCTTGGTGAGAAAGT
NZ_CP035265.1:766128-774627    7584    TGATTTGGTCAGCTTCTTGCATTCGTTCTTGGTAGTAGCACCAAGAATAATTACCATCGATGACCCAAGCTTTATGCTTGGTGAGAAAGT
NZ_CP035245.1 745261-753757    7581    TGATTTGGTCAGCTTCTTGCATTCGTTCTTGGTAGTAGCACCAAGAATAATTACCATCGATGACCCAAGCTTTATGCTTGGTGAGAAAGT
NZ_CP035259.1:745700-753548    6933    TGATTTGGTCAGCTTCTTGCATTCGTTCTTGGTAGTAGCACCAAGAATAATTACCATCGATGACCCAAGCTTTATGCTTGGTGAGAAAGT
NZ_CP035258.1:750874-758722    6933    TGATTTGGTCAGCTTCTTGCATTCGTTCTTGGTAGTAGCACCAAGAATAATTACCATCGATGACCCAAGCTTTATGCTTGGTGAGAAAGT
NZ_CP035260.1:726156-733994    6923    TGATTTGGTCAGCTTCTTGCATTCGTTCTTGGTAGTAGCACCAAGAATAATTACCATCGATGACCCAAGCTTTATGCTTGGTGAGAAAGT
NZ_CP035241.1:708296-716144    6933    TGATTTGGTCAGCTTCTTGCATTCGTTCTTGGTAGTAGCACCAAGAATAATTACCATCGATGACCCAAGCTTTATGCTTGGTGAGAAAGT
NZ_CP035261.1:779086-786938    6937    TGATTTGGTCAGCTTCTTGCATTCGTTCTTGGTAGTAGCACCAAGAATAATTACCATCGATGACCCAAGCTTTATGCTTGGTGAGAAAGT
NZ_CP035242.1:c1532236-1524385 6936    TGATTTGGTCAGCTTCTTGCATTCGTTCTTGGTAGTAGCACCAAGAATAATTACCATCGATGACCCAAGCTTTATGCTTGGTGAGAAAGT
NZ_CP035243.1:777729-785580    6936    TGATTTGGTCAGCTTCTTGCATTCGTTCTTGGTAGTAGCACCAAGAATAATTACCATCGATGACCCAAGCTTTATGCTTGGTGAGAAAGT
NZ_CP035237.1:c1311322-1303456 6951    TGATTTGGTCAGCTTCTTGCATTCGTTCTTGGTAGTAGCACCAAGAATAATTACCATCGATGACCCAAGCTTTATGCTTGGTGAGAAAGT
NZ_CP035246.1:c1302892-1295026 6951    TGATTTGGTCAGCTTCTTGCATTCGTTCTTGGTAGTAGCACCAAGAATAATTACCATCGATGACCCAAGCTTTATGCTTGGTGAGAAAGT
NZ_CP035247.1:c1301115-1293263 6937    TGATTTGGTCAGCTTCTTGCATTCGTTCTTGGTAGTAGCACCAAGAATAATTACCATCGATGACCCAAGCTTTATGCTTGGTGAGAAAGT
NZ_CP035238.1:738547-746413    6951    TGATTTGGTCAGCTTCTTGCATTCGTTCTTGGTAGTAGCACCAAGAATAATTACCATCGATGACCCAAGCTTTATGCTTGGTGAGAAAGT
NZ_CP035255.1:776866-784718    6937    TGATTTGGTCAGCTTCTTGCATTCGTTCTTGGTAGTAGCACCAAGAATAATTACCATCGATGCCCCAAGCTTTATGCTTGGTGAGAAAGT
NZ_CP035256.1:807236-815102    6951    TGATTTGGTCAGCTTCTTGCATTCGTTCTTGGTAGTAGCACCAAGAATAATTACCATCGATGCCCCAAGCTTTATGCTTGGTGAGAAAGT
NZ_CP038252.1:1151812-1159673  6876    TGATTTGGTCAGCTTCTTGCATTCGTTCTTGGTAGTAGCACCAAGAATAATTACCATCGATGACCCAAGCTTTATGCTTGGTGAGAAAGT
NZ_CP035254.1:793365-801213    6933    TGATTTGGTCAGCTTCTTGCATTCGTTCTTGGTAGTAGCACCAAGAATAATTACCATCGATGACCCAAGCTTTATGCTTGGTGAGAAAGT
NZ_CP035251.1:748733-756585    6937    TGATTTGGTCAGCTTCTTGCATTCGTTCTTGGTAGTAGCACCAAGAATAATTACCATCGATGACCCAAGCTTTATGCTTGGTGAGAAAGT
NZ_CP035252.1:745670-753536    6951    TGATTTGGTCAGCTTCTTGCATTCGTTCTTGGTAGTAGCACCAAGAATAATTACCATCGATGACCCAAGCTTTATGCTTGGTGAGAAAGT
NZ_CP035257.1:776317-784169    6937    TGATTTGGTCAGCTTCTTGCATTCGTTCTTGGTAGTAGCACCAAGAATAATTACCATCGATGACCCAAGCTTTATGCTTGGTGAGAAAGT
NZ_CP038253.1:758105-765957    6937    TGATTTGGTCAGCTTCTTGCATTCGTTCTTGGTAGTAGCACCAAGAATAATTACCATCGATGACCCAAGCTTTATGCTTGGTGAGAAAGT
NZ_CP035263.1:743050-750901    6936    TGATTTGGTCAGCTTCTTGCATTCGTTCTTGGTAGTAGCACCAAGAATAATTACCATCGATGACCCAAGCTTTATGCTTGGTGAGAAAGT
NZ_CP035262.1:741595-749460    6950    TGATTTGGTCAGCTTCTTGCATTCGTTCTTGGTAGTAGCACCAAGAATAATTACCATCGATGACCCAAGCTTTATGCTTGGTGAGAAAGT
NZ_CP035236.1:764915-772766    6936    TGATTTGGTCAGCTTCTTGCATTCGTTCTTGGTAGCAGCACCAAGAATAATTACCATCGATGACCCAAGCTTTATGCTTGGTGAGAAAGT
NZ_CP035253.1 767286-775134    6933    TGATTTGGTCAGCTTCTTGCATTCGTTCTTGGTAGTAGCACCAAGAATAATTACCATCGATGACCCAAGCTTTATGCTTGGTGAGAAAGT
NZ_CP035249.1:763080-770921    6926    TGATTTGGTCAGCTTCTTGCATTCGTTCTTGGTAGTAGCACCAAGAATAATTACCATCGATGACCCAAGCTTTATGCTTGGTGAGAAAGT

NC_003028.3:777000-787000      9175    TTTTTATCTCGGTTAACATCCATTCGCAGTCACTGTCTTGCCAACCAGGTTGAAATTGGAGTGTGTCCATGTGCAGTTTTGGAATGGAGT
NZ_CP035248.1:765099-775111    9187    TTTTTATCTCGGTTAACATCCATTCGCAGTCACTGTCTTGCCAACCAGGTTGAAATTGGAGTGTGTCCATGTGCAGTTTTGGAATGGAGT
NZ_CP035235.1 c1326834-1317894 8323   ------------------------------------------------------------------------------------------
NZ_CP035234.1:756284-765224    8323    ------------------------------------------------------------------------------------------
NZ_CP038251.1:750745-759686    8324    ------------------------------------------------------------------------------------------
NZ_CP035244.1:747023-755950    8310    ------------------------------------------------------------------------------------------
NZ_CP035264.1:800836-809336    7675    TTTTTATCTCGGTTAACATCCATTCGTGGTCACTGTCTTGCCAACCAGGTTGAAATTGGAGTGTGTCCATGTGCAGTTTTGGAATGGAGT
NZ_CP035265.1:766128-774627    7674    TTTTTATCTCGGTTAACATCCATTCGTGGTCACTGTCTTGCCAACCAGGTTGAAATTGGAGTGTGTCCATGTGCAGTTTTGGAATGGAGT
NZ_CP035245.1 745261-753757    7671    TTTTTATCTCGGTTAACATCCATTCGCAGTCACTGTCTTGCCAACCAGGTTGAAATTGGAGTGTGTCCATGTGCAGTTTTGGAATGGAGT
NZ_CP035259.1:745700-753548    7023    TTTTTATCTCGGTTAACATCCATTCGCAGTCACTGTCTTGCCAACCAGGTTGAAATTGGAGTGTGTCCATGTGCAGTTTTGGAATGGAGT
NZ_CP035258.1:750874-758722    7023    TTTTTATCTCGGTTAACATCCATTCGCAGTCACTGTCTTGCCAACCAGGTTGAAATTGGAGTGTGTCCATGTGCAGTTTTGGAATGGAGT
NZ_CP035260.1:726156-733994    7013    TTTTTATCTCGGTTAACATCCATTCGCAGTCACTGTCTTGCCAACCAGGTTGAAATTGGAGTGTGTCCATGTGCAGTTTTGGAATGGAGT
NZ_CP035241.1:708296-716144    7023    TTTTTATCTCGGTTAACATCCATTCGCAGTCACTGTCTTGCCAACCAGGTTGAAATTGGAGTGTGTCCATGTGCAGTTTTGGAATGGAGT
NZ_CP035261.1:779086-786938    7027    TTTTTATCTCGGTTAACATCCATTCGCAGTCACTGTCTTGCCAACCAGGTTGAAATTGGAGTGTGTCCATGTGCAGTTTTGGAATGGAGT
NZ_CP035242.1:c1532236-1524385 7026    TTTTTATCTCGGTTAACATCCATTCGCAGTCACTGTCTTGCCAATCAGGTTGAAATTGGAGTGTGTCCATGTGCAGTTTTGGAATGGAGT
NZ_CP035243.1:777729-785580    7026    TTTTTATCTCGGTTAACATCCATTCGCAGTCACTGTCTTGCCAACCAGGTTGAAATTGGAGTGTGTCCATGTGCAGTTTTGGAATGGAGT
NZ_CP035237.1:c1311322-1303456 7041    TTTTTATCTCGGTTAACATCCATTCGCAGTCACTGTCTTGCCAACCAGGTTGAAATTGGAGTGTGTCCATGTGCAGTTTTGGAATGGAGT
NZ_CP035246.1:c1302892-1295026 7041    TTTTTATCTCGGTTAACATCCATTCGCAGTCACTGTCTTGCCAACCAGGTTGAAATTGGAGTGTGTCCATGTGCAGTTTTGGAATGGAGT
NZ_CP035247.1:c1301115-1293263 7027    TTTTTATCTCGGTTAACATCCATTCGCAGTCACTGTCTTGCCAACCAGGTTGAAATTGGAGTGTGTCCATGTGCAGTTTTGGAATGGAGT
NZ_CP035238.1:738547-746413    7041    TTTTTATCTCGGTTAACATCCATTCGCAGTCACTGTCTTGCCAACCAGGTTGAAATTGGAGTGTGTCCATGTGCAGTTTTGGAATGGAGT
NZ_CP035255.1:776866-784718    7027    TTTTTATCTCGGTTAACATCCATTCGCAGTCACTGTCTTGCCAACCAGGTTGAAATTGGAGTGTGTCCATGTGCAGTTTTGGAATGGAGT
NZ_CP035256.1:807236-815102    7041    TTTTTATCTCGGTTAACATCCATTCGCAGTCACTGTCTTGCCAACCAGGTTGAAATTGGAGTGTGTCCATGTGCAGTTTTGGAATGGAGT
NZ_CP038252.1:1151812-1159673  6966    TTTTTATCTCGGTTAACATCCATTCGCAGTCACTGTCTTGCCAACCAGGTTGAAATTGGAGTGTGTCCATGTGCAGTTTTGGAATGGAGT
NZ_CP035254.1:793365-801213    7023    TTTTTATCTCGGTTAACATCCATTCGCAGTCACTGTCTTGCCAACCAGGTTGAAATTGGAGTGTGTCCATGTGCAGTTTTGGAATGGAGT
NZ_CP035251.1:748733-756585    7027    TTTTTATCTCGGTTAACATCCATTCGCAGTCACTGTCTTGCCAACCAGGTTGAAATTGGAGTGTGTCCATGTGCAGTTTTGGAATGGAGT
```

### Ozkan et al. Supplementary Figures

```
NZ_CP035252.1:745670-753536    7041   TTTTTATCTCGGTTAACATCCATTCGCAGTCACTGTCTTGCCAACCAGGTTGAAATTGGAGTGTGTCCATGTGCAGTTTTGGAATGGAGT
NZ_CP035257.1:776317-784169    7027   TTTTTATCTCGGTTAACATCCATTCGCAGTCACTGTCTTGCCAACCAGGTTGAAATTGGAGTGTGTCCATGTGCAGTTTTGGAATGGAGT
NZ_CP038253.1:758105-765957    7027   TTTTTATCTCGGTTAACATCCATTCGCAGTCACTGTCTTGCCAACCAGGTTGAAATTGGAGTGTGTCCATGTGCAGTTTTGGAATGGAGT
NZ_CP035263.1:743050-750901    7026   TTTTTATCTCGGTTAACATCCATTCGCAGTCACTGTCTTGCCAACCAGGTTGAAATTGGAGTGTGTCCATGTGCAGTTTTGGAATGGAGT
NZ_CP035262.1:741595-749460    7040   TTTTTATCTCGGTTAACATCCATTCGCAGTCACTGTCTTGCCAACCAGGTTGAAATTGGAGTGTGTCCATGTGCAGTTTTGGAATGGAGT
NZ_CP035236.1:764915-772766    7026   TTTTTATATCGGTTAACATCCATTCGTGGTCACTGTCTTGCCAACCAGGTTGAAATTGGAGTGTGTCCATGTGCAGTTTTGGAATGGAGT
NZ_CP035253.1 767286-775134    7023   TTTTTATCTCGGTTAACATCCATTCGCAGTCACTGTCTTGCCAACCAGGTTGAAATTGGAGTGTGTCCATGTGCAGTTTTGGAATGGAGT
NZ_CP035249.1:763080-770921    7016   TTTTTATCTCGGTTAACATCCATTCGCAGTCACTGTCTTGCCAACCAGGTTGAAATTGGAGTGTGTCCATGTGCAGTTTTGGAATGGAGT

NC_003028.3:777000-787000      9265   AGTAGTTAGATAACTTTTCTGCTATAGTTGACTTACCAGAACCAGAATATCCGATAATTGCGATTTTCATTTTCTACCTTTTCCTATTTG
NZ_CP035248.1:765099-775111    9277   AGTAGTTAGATAACTTTTCTGCTATAGTTGACTTACCAGAACCAGAATATCCGATAATTGCGATTTTCATTTTCTACCTTTTCCTATTTG
NZ_CP035235.1 c1326834-1317894 8323   ------------------------------------------------------------------------------------------
NZ_CP035234.1:756284-765224    8323   ------------------------------------------------------------------------------------------
NZ_CP038251.1:750745-759686    8324   ------------------------------------------------------------------------------------------
NZ_CP035244.1:747023-755950    8310   ------------------------------------------------------------------------------------------
NZ_CP035264.1:800836-809336    7765   AGTAGTTAGATAACTTTTCTGCTAGAGTTGACTTACCAGAACCAGAATATCCGATAATTGCGATTTTCATTTTCTACCTTTTCCTATTTG
NZ_CP035265.1:766128-774627    7764   AGTAGTTAGATAACTTTTCTGCTAGAGTTGACTTACCAGAACCAGAATATCCGATAATTGCGATTTTCATTTTCTACCTTTTCCTATTTG
NZ_CP035245.1 745261-753757    7761   AGTAGTTAGATAACTTTTCTGCTAGAGTTGACTTACCAGAACCAGAATATCCGATAATTGCGATTTTCATTTTCTACCTTTTCCTATTTG
NZ_CP035259.1:745700-753548    7113   AGTAGTTAGATAACTTTTCTGCTAGAGTTGACTTACCAGAACCAGAATATCCGATAATTGTGATTTTCATTTTCTACCTTTTCCTATTTG
NZ_CP035258.1:750874-758722    7113   AGTAGTTAGATAACTTTTCTGCTAGAGTTGACTTACCAGAACCAGAATATCCGATAATTGTGATTTTCATTTTCTACCTTTTCCTATTTG
NZ_CP035260.1:726156-733994    7103   AGTAGTTAGATAACTTTTCTGCTAGAGTTGACTTACCAGAACCAGAATATCCGATAATTGTGATTTTCATTTTCTACCTTTTCCTATTTG
NZ_CP035241.1:708296-716144    7113   AGTAGTTAGATAACTTTTCTGCTAGAGTTGACTTACCAGAACCAGAATATCCGATAATTGTGATTTTCATTTTCTACCTTTTCCTATTTG
NZ_CP035261.1:779086-786938    7117   AGTAGTTAGATAACTTTTCTGCTAGAGTTGACTTACCAGAACCAGAATATCCGATAATTGTGATTTTCATTTTCTACCTTTTCCTATTTG
NZ_CP035242.1:c1532236-1524385 7116   AGTAGTTAGATAACTTTTCTGCTATAGTTGACTTACCAGAACCAGAATATCCGATAATTGCGATTTTCATTTTCTACCTTTTCCTATTTG
NZ_CP035243.1:777729-785580    7116   AGTAGTTAGATAACTTTTCTGCTATAGTTGACTTACCAGAACCAGAATATCCGATAATTGCGATTTTCATTTTCTACCTTTTCCTATTTG
NZ_CP035237.1:c1311322-1303456 7131   AGTAGTTAGATAACTTTTCTGCTAGAGTTGACTTACCAGAACCAGAATATCCGATAATTGTGATTTTCATTTTCTACCTTTTCCTATTTG
NZ_CP035246.1:c1302892-1295026 7131   AGTAGTTAGATAACTTTTCTGCTAGAGTTGACTTACCAGAACCAGAATATCCGATAATTGTGATTTTCATTTTCTACCTTTTCCTATTTG
NZ_CP035247.1:c1301115-1293263 7117   AGTAGTTAGATAACTTTTCTGCTAGAGTTGACTTACCAGAACCAGAATATCCGATAATTGTGATTTTCATTTTCTACCTTTTCCTATTTG
NZ_CP035238.1:738547-746413    7131   AGTAGTTAGATAACTTTTCTGCTAGAGTTGACTTACCAGAACCAGAATATCCGATAATTGTGATTTTCATTTTCTACCTTTTCCTATTTG
NZ_CP035255.1:776866-784718    7117   AGTAGTTAGATAACTTTTCTGCTAGAGTTGACTTACCAGAACCAGAATATCCGATAATTGTGATTTTCATTTTCTACCTTTTCCTATTTG
NZ_CP035256.1:807236-815102    7131   AGTAGTTAGATAACTTTTCTGCTAGAGTTGACTTACCAGAACCAGAATATCCGATAATTGTGATTTTCATTTTCTACCTTTTCCTATTTG
NZ_CP038252.1:1151812-1159673  7056   AGTAGTTAGATAACTTTTCTGCTAGAGTTGACTTACCAGAACCAGAATATCCGATAATTGTGATTTTCATTTTCTACCTTTTCCTATTTG
NZ_CP035254.1:793365-801213    7113   AGTAGTTAGATAACTTTTCTGCTAGAGTTGACTTACCAGAACCAGAATATCCGATAATTGTGATTTTCATTTTCTACCTTTTCCTATTTG
NZ_CP035251.1:748733-756585    7117   AGTAGTTAGATAACTTTTCTGCTAGAGTTGACTTACCAGAACCAGAATATCCGATAATTGTGATTTTCATTTTCTACCTTTTCCTATTTG
NZ_CP035252.1:745670-753536    7131   AGTAGTTAGATAACTTTTCTGCTAGAGTTGACTTACCAGAACCAGAATATCCGATAATTGTGATTTTCATTTTCTACCTTTTCCTATTTG
NZ_CP035257.1:776317-784169    7117   AGTAGTTAGATAACTTTTCTGCTAGAGTTGACTTACCAGAACCAGAATATCCGATAATTGTGATTTTCATTTTCTACCTTTTCCTATTTG
NZ_CP038253.1:758105-765957    7117   AGTAGTTAGATAACTTTTCTGCTAGAGTTGACTTACCAGAACCAGAATATCCGATAATTGTGATTTTCATTTTCTACCTTTTCCTATTTG
NZ_CP035263.1:743050-750901    7116   AGTAGTTAGATAACTTTTCTGCTATAGTTGACTTACCAGAACCAGAATATCCGATAATTGCGATTTTCATTTTCTACTTTTTCCTATTTG
NZ_CP035262.1:741595-749460    7130   AGTAGTTAGATAACTTTTCTGCTATAGTTGACTTACCAGAACCAGAATATCCGATAATTGCGATTTTCATTTTCTACTTTTTCCTATTTG
NZ_CP035236.1:764915-772766    7116   AGTAGTTAGATAACTTTTCTGCTAGAGTTGACTTACCAGAACCAGAATATCCGATAATTGCGATTTTCATTTTCTACCTTTTCCTATTTG
NZ_CP035253.1 767286-775134    7113   AGTAGTTAGATAACTTTTCTGCTAGAGTTGACTTACCAGAACCAGAATATCCGATAATTGTGATTTTCATTTTCTACCTTTTCCTATTTG
NZ_CP035249.1:763080-770921    7106   AGTAGTTAGATAACTTTTCTGCTATAGTTGACTTACCAGAACCAGAATATCCGATAATTGCGATTTTCATTTTCTACTTTTTCCTATTTG

NC_003028.3:777000-787000      9355   GAGACAAAAAAACAGCCTCTATGGACTGTTTCTTATTTAACAAGTTTAGCTGAAAGACGAGCTTTATCGCGGCTTGCTTTGTTTTTGTGA
NZ_CP035248.1:765099-775111    9367   GAGACAAAAAAACAGCCTCTATGGACTGTTTCTTATTTAACAAGTTTAGCTGAAAGACGAGCTTTATCGCGGCTTGCTTTGTTTTTGTGA
NZ_CP035235.1 c1326834-1317894 8323   -----------------------------TTCTTATTTAGCAAGTTTAGCTGAAAGACGAGCTTTATCGCGGCTTGCTTTGTTTTTGTGA
NZ_CP035234.1:756284-765224    8323   -----------------------------TTCTTATTTAGCAAGTTTAGCTGAAAGACGAGCTTTATCGCGGCTTGCTTTGTTTTTGTGA
NZ_CP038251.1:750745-759686    8324   -----------------------------TTCTTATTTAGCAAGTTTAGCTGAAAGACGAGCTTTATCGCGGCTTGCTTTGTTTTTGTGA
NZ_CP035244.1:747023-755950    8310   -----------------------------TTCTTATTTAGCAAGTTTAGCTGAAAGACGAGCTTTATCGCGGCTTGCTTTGTTTTTGTGA
NZ_CP035264.1:800836-809336    7855   GAGACAAAAAAACAGCCTCTATGGACTGTTTCTTATTTAGCAAGTTTAGCTGAAAGACGAGCTTTATCGCGGCTTGCTTTGTTTTTGTGA
NZ_CP035265.1:766128-774627    7854   GAGACAAAAAAACAGCCTCTATGGACTGTTTCTTATTTAGCAAGTTTAGCTGAAAGACGAGCTTTATCGCGGCTTGCTTTGTTTTTGTGA
NZ_CP035245.1 745261-753757    7851   GAGACAAAAAAACAGCCTCTATGGACTGTTTCTTATTTAGCAAGTTTAGCTGAAAGACGAGCTTTATCGCGGCTTGCTTTGTTTTTGTGA
NZ_CP035259.1:745700-753548    7203   GAGACAAAAAAACAGCCTCTATGGACTGTTTCTTATTTAGCAAGTTTAGCTGAAAGACGAGCTTTATCGCGGCTTGCTTTGTTTTTGTGA
NZ_CP035258.1:750874-758722    7203   GAGACAAAAAAACAGCCTCTATGGACTGTTTCTTATTTAGCAAGTTTAGCTGAAAGACGAGCTTTATCGCGGCTTGCTTTGTTTTTGTGA
NZ_CP035260.1:726156-733994    7193   GAGACAAAAAAACAGCCTCTATGGACTGTTTCTTATTTAGCAAGTTTAGTTGAAAGACGAGCTTTATCGCGGCTTGCTTTGTTTTTGTGA
NZ_CP035241.1:708296-716144    7203   GAGACAAAAAAACAGCCTCTATGGACTGTTTCTTATTTAGCAAGTTTAGTTGAAAGACGAGCTTTATCGCGGCTTGCTTTGTTTTTGTGA
NZ_CP035261.1:779086-786938    7207   GAGACAAAAAAACAGCCTCTATGGACTGTTTCTTATTTAGCAAGTTTAGCTGAAAGACGAGCTTTATCGCGGCTTGCTTTGTTTTTGTGA
NZ_CP035242.1:c1532236-1524385 7206   GAGACAAAAAAACAGCCTCTATGGACTGTTTCTTATTTAACAAGTTTAGCTGAAAGACGAGCTTTATCGCGGCTTGCTTTGTTTTTGTGA
NZ_CP035243.1:777729-785580    7206   GAGACAAAAAAACAGCCTCTATGGACTGTTTCTTATTTAACAAGTTTAGCTGAAAGACGAGCTTTATCGCGGCTTGCTTTGTTTTTGTGA
NZ_CP035237.1:c1311322-1303456 7221   GAGACAAAAAAACAGCCTCTATGGACTGTTTCTTATTTAGCAAGTTTAGTTGAAAGACGAGCTTTATCGCGGCTTGCTTTGTTTTTGTGA
NZ_CP035246.1:c1302892-1295026 7221   GAGACAAAAAAACAGCCTCTATGGACTGTTTCTTATTTAGCAAGTTTAGCTGAAAGACGAGCTTTATCGCGGCTTGCTTTGTTTTTGTGA
NZ_CP035247.1:c1301115-1293263 7207   GAGACAAAAAAACAGCCTCTATGGACTGTTTCTTATTTAGCAAGTTTAGCTGAAAGACGAGCTTTATCGCGGCTTGCTTTGTTTTTGTGA
NZ_CP035238.1:738547-746413    7221   GAGACAAAAAAACAGCCTCTATGGACTGTTTCTTATTTAGCAAGTTTAGCTGAAAGACGAGCTTTATCGCGGCTTGCTTTGTTTTTGTGA
NZ_CP035255.1:776866-784718    7207   GAGACAAAAAAACAGCCTCTATGGACTGTTTCTTATTTAGCAAGTTTAGCTGAAAGACGAGCTTTATCGCGGCTTGCTTTGTTTTTGTGA
NZ_CP035256.1:807236-815102    7221   GAGACAAAAAAACAGCCTCTATGGACTGTTTCTTATTTAGCAAGTTTAGCTGAAAGACGAGCTTTATCGCGGCTTGCTTTGTTTTTGTGA
NZ_CP038252.1:1151812-1159673  7146   GAGACAAAAAAACAGCCTCTATGGACTGTTTCTTATTTAGCAAGTTTAGCTGAAAGACGAGCTTTATCGCGGCTTGCTTTGTTTTTGTGA
NZ_CP035254.1:793365-801213    7203   GAGACAAAAAAACAGCCTCTATGGACTGTTTCTTATTTAGCAAGTTTAGCTGAAAGACGAGCTTTATCGCGGCTTGCTTTGTTTTTGTGA
NZ_CP035251.1:748733-756585    7207   GAGACAAAAAAACAGCCTCTATGGACTGTTTCTTATTTAGCAAGTTTAGCTGAAAGACGAGCTTTATCGCGGCTTGCTTTGTTTTTGTGA
NZ_CP035252.1:745670-753536    7221   GAGACAAAAAAACAGCCTCTATGGACTGTTTCTTATTTAGCAAGTTTAGCTGAAAGACGAGCTTTATCGCGGCTTGCTTTGTTTTTGTGA
NZ_CP035257.1:776317-784169    7207   GAGACAAAAAAACAGCCTCTATGGACTGTTTCTTATTTAGCAAGTTTAGCTGAAAGACGAGCTTTATCGCGGCTTGCTTTGTTTTTGTGA
NZ_CP038253.1:758105-765957    7207   GAGACAAAAAAACAGCCTCTATGGACTGTTTCTTATTTAGCAAGTTTAGCTGAAAGACGAGCTTTATCGCGGCTTGCTTTGTTTTTGTGA
NZ_CP035263.1:743050-750901    7206   GAGACAAAAAAACAGCCTCTATGGACTGTTTCTTATTTAGCAAGTTTAGCTGAAAGACGAGCTTTATCGCGGCTTGCTTTGTTTTTGTGA
NZ_CP035262.1:741595-749460    7220   GAGACAAAAAAACAGCCTCTATGGACTGTTTCTTATTTAGCAAGTTTAGCTGAAAGACGAGCTTTATCGCGGCTTGCTTTGTTTTTGTGA
NZ_CP035236.1:764915-772766    7206   GAGACAAAAAAACAGCCTCTATGGACTGTTTCTTATTTAGCAAGTTTAGCTGAAAGACGAGCTTTATCGCGGCTTGCTTTGTTTTTGTGA
NZ_CP035253.1 767286-775134    7203   GAGACAAAAAAACAGCCTCTATGGACTGTTTCTTATTTAGCAAGTTTAGCTGAAAGACGAGCTTTATCGCGGCTTGCTTTGTTTTTGTGA
NZ_CP035249.1:763080-770921    7196   GAGACAAAAAAACAGCCTCTATGGACTGTTTCTTATTTAGCAAGTTTAGCTGAAAGACGAGCTTTATCGCGGCTTGCTTTGTTTTTGTGA

NC_003028.3:777000-787000      9445   ATCAAACCTTTAGTTTCTGCTTTATCGATAGCTGAGCTAGCAGCACGGAAAAGTTCTTCAGATGGGTTTGCTTCGAAAGCTTTTATAGCA
NZ_CP035248.1:765099-775111    9457   ATCAAACCTTTAGTTTCTGCTTTATCGATAGCTGAGCTAGCAGCACGGAAAAGTTCTTCAGATGGGTTTGCTTCGAAAGCTTTTATAGCA
NZ_CP035235.1 c1326834-1317894 8385   ATCAAACCTTTAGTTTCTGCTTTATCGATAGCTGAGCTAGCAGCACGGAAAAGTTCTTCAGATGGGTTTGCTTCGAAAGCTTTTATAGCA
NZ_CP035234.1:756284-765224    8385   ATCAAACCTTTAGTTTCTGCTTTATCGATAGCTGAGCTAGCAGCACGGAAAAGTTCTTCAGATGGGTTTGCTTCGAAAGCTTTTATAGCA
NZ_CP038251.1:750745-759686    8386   ATCAAACCTTTAGTTTCTGCTTTATCGATAGCTGAGCTAGCAGCACGGAAAAGTTCTTCAGATGGGTTTGCTTCGAAAGCTTTTATAGCA
NZ_CP035244.1:747023-755950    8372   ATCAAACCTTTAGTTTCTGCTTTATCGATAGCTGAGCTAGCAGCACGGAAAAGTTCTTCAGATGGGTTTGCTTCGAAAGCTTTTATAGCA
NZ_CP035264.1:800836-809336    7945   ATCAAACCTTTAGTTTCTGCTTTATCGATAGCTGAGCTAGCAGCACGGAAAAGTTCTTCAGATGGGTTTGCTTCGAAAGCTTTTATAGCA
NZ_CP035265.1:766128-774627    7944   ATCAAACCTTTAGTTTCTGCTTTATCGATAGCTGAGCTAGCAGCACGGAAAAGTTCTTCAGATGGGTTTGCTTCGAAAGCTTTTATAGCA
NZ_CP035245.1 745261-753757    7941   ATCAAACCTTTAGTTTCTGCTTTATCGATAGCTGAGCTAGCAGCACGGAAAAGTTCTTCAGATGGGTTTGCTTCGAAAGCTTTTATAGCA
NZ_CP035259.1:745700-753548    7293   ATCAAACCTTTAGTTTCTGCTTTATCGATAGCTGAGCTAGCAGCACGGAAAAGTTCTTCAGATGGGTTTGCTTCGAAAGCTTTTATAGCA
NZ_CP035258.1:750874-758722    7293   ATCAAACCTTTAGTTTCTGCTTTATCGATAGCTGAGCTAGCAGCACGGAAAAGTTCTTCAGATGGGTTTGCTTCGAAAGCTTTTATAGCA
NZ_CP035260.1:726156-733994    7283   ATCAAACCTTTAGTTTCTGCTTTATCGATAGCTGAGCTAGCAGCACGGAAAAGTTCTTCAGATGGGTTTGCTTCGAAAGCTTTTATAGCA
NZ_CP035241.1:708296-716144    7293   ATCAAACCTTTAGTTTCTGCTTTATCGATAGCTGAGCTAGCAGCACGGAAAAGTTCTTCAGATGGGTTTGCTTCGAAAGCTTTTATAGCA
NZ_CP035261.1:779086-786938    7297   ATCAAACCTTTAGTTTCTGCTTTATCGATAGCTGAGCTAGCAGCACGGAAAAGTTCTTCAGATGGGTTTGCTTCGAAAGCTTTTATAGCA
NZ_CP035242.1:c1532236-1524385 7296   ATCAAACCTTTAGTTTCTGCTTTATCGATAGCTGAGCTAGCAGCACGGAAAAGTTCTTCAGATGGGTTTGCTTCGAAAGCTTTTATAGCA
NZ_CP035243.1:777729-785580    7296   ATCAAACCTTTAGTTTCTGCTTTATCGATAGCTGAGCTAGCAGCACGGAAAAGTTCTTCAGATGGGTTTGCTTCGAAAGCTTTTATAGCA
NZ_CP035237.1:c1311322-1303456 7311   ATCAAACCTTTAGTTTCTGCTTTATCGATAGCTGAGCTAGCAGCACGGAAAAGTTCTTCAGATGGGTTTGCTTCGAAAGCTTTTATAGCA
NZ_CP035246.1:c1302892-1295026 7311   ATCAAACCTTTAGTTTCTGCTTTATCGATAGCTGAGCTAGCAGCACGGAAAAGTTCTTCAGATGGGTTTGCTTCGAAAGCTTTTATAGCA
```

```
NZ_CP035247.1:c1301115-1293263 7297   ATCAAACCTTTAGTTTCTGCTTTATCGATAGCTGAGCTAGCAGCACGGAAAAGTTCTTCAGATGGGTTTGCTTCGAAAGCTTTTATAGCA
NZ_CP035238.1:738547-746413    7311   ATCAAACCTTTAGTTTCTGCTTTATCGATAGCTGAGCTAGCAGCACGGAAAAGTTCTTCAGATGGGTTTGCTTCGAAAGCTTTTATAGCA
NZ_CP035255.1:776866-784718    7297   ATCAAACCTTTAGTTTCTGCTTTATCGATAGCTGAGCTAGCAGCACGGAAAAGTTCTTCAGATGGGTTTGCTTCGAAAGCTTTTATAGCA
NZ_CP035256.1:807236-815102    7311   ATCAAACCTTTAGTTTCTGCTTTATCGATAGCTGAGCTAGCAGCACGGAAAAGTTCTTCAGATGGGTTTGCTTCGAAAGCTTTTATAGCA
NZ_CP038252.1:1151812-1159673  7236   ATCAAACCTTTAGTTTCTGCTTTATCGATAGCTGAGCTAGCAGCACGGAAAAGTTCTTCAGATGGGTTTGCTTCGAAAGCTTTTATAGCA
NZ_CP035254.1:793365-801213    7293   ATCAAACCTTTAGTTTCTGCTTTATCGATAGCTGAGCTAGCAGCACGGAAAAGTTCTTCAGATGGGTTTGCTTCGAAAGCTTTTATAGCA
NZ_CP035251.1:748733-756585    7297   ATCAAACCTTTAGTTTCTGCTTTATCGATAGCTGAGCTAGCAGCACGGAAAAGTTCTTCAGATGGGTTTGCTTCAAAAGCTTTTATAGCA
NZ_CP035252.1:745670-753536    7311   ATCAAACCTTTAGTTTCTGCTTTATCGATAGCTGAGCTAGCAGCACGGAAAAGTTCTTCAGATGGGTTTGCTTCAAAAGCTTTTATAGCA
NZ_CP035257.1:776317-784169    7297   ATCAAACCTTTAGTTTCTGCTTTATCGATAGCTGAGCTAGCAGCACGGAAAAGTTCTTCAGATGGGTTTGCTTCAAAAGCTTTTATAGCA
NZ_CP038253.1:758105-765957    7297   ATCAAACCTTTAGTTTCTGCTTTATCGATAGCTGAGCTAGCAGCACGGAAAAGTTCTTCAGATGGGTTTGCTTCAAAAGCTTTTATAGCA
NZ_CP035263.1:743050-750901    7296   ATCAAACCTTTAGTTTCTGCTTTATCGATAGCTGAGCTAGCAGCACGGAAAAGTTCTTCAGATGGGTTTGCTTCGAAAGCTTTTATAGCA
NZ_CP035262.1:741595-749460    7310   ATCAAACCTTTAGTTTCTGCTTTATCGATAGCTGAGCTAGCAGCACGGAAAAGTTCTTCAGATGGGTTTGCTTCGAAAGCTTTTATAGCA
NZ_CP035236.1:764915-772766    7296   ATCAAACCTTTAGTTTCTGCTTTATCGATAGCTGAGCTAGCAGCACGGAAAAGTTCTTCAGATGGGTTTGCTTCGAAAGCTTTTATAGCA
NZ_CP035253.1 767286-775134    7293   ATCAAACCTTTAGTTTCTGCTTTATCGATAGCTGAGCTAGCAGCACGGAAAAGTTCTTCAGATGGGTTTGCTTCGAAAGCTTTTATAGCA
NZ_CP035249.1:763080-770921    7286   ATCAAACCTTTAGTTTCTGCTTTATCGATAGCTGAGCTAGCAGCACGGAAAAGTTCTTCAGATGGGTTTGCTTCGAAAGCTTTTATAGCA

NC_003028.3:777000-787000      9535   GTACGCATAGCTGATTTTTGAGCTGAGTTCTTTTCGTTTTGTTTAACGTTCAATTCAGCGCGTTTGATAGCTGATTTAATGTTTGCCAAT
NZ_CP035248.1:765099-775111    9547   GTACGCATAGCTGATTTTTGAGCTGAGTTCTTTTCGTTTTGTTTAACGTTCAATTCAGCGCGTTTGATAGCTGATTTAATGTTTGCCAAT
NZ_CP035235.1 c1326834-1317894 8475   GTACGCATAGCTGATTTTTGAGCTGAGTTCTTTTCGTTTTGTTTAACGTTCAATTCAGCGCGTTTGATAGCTGATTTAATGTTTGCCAAT
NZ_CP035234.1:756284-765224    8475   GTACGCATAGCTGATTTTTGAGCTGAGTTCTTTTCGTTTTGTTTAACGTTCAATTCAGCGCGTTTGATAGCTGATTTAATGTTTGCCAAT
NZ_CP038251.1:750745-759686    8476   GTACGCATAGCTGATTTTTGAGCTGAGTTCTTTTCGTTTTGTTTAACGTTCAATTCAGCGCGTTTGATAGCTGATTTAATGTTTGCCAAT
NZ_CP035244.1:747023-755950    8462   GTACGCATAGCTGATTTTTGAGCTGAGTTCTTTTCGTTTTGTTTAACGTTCAATTCAGCGCGTTTGATAGCTGATTTAATGTTTGCCAAT
NZ_CP035264.1:800836-809336    8035   GTACGCATAGCTGATTTTTGAGCTGAGTTCTTTTCGTTTTGTTTAACGTTCAATTCAGCGCGTTTGATAGCTGATTTAATGTTTGCCAAT
NZ_CP035265.1:766128-774627    8034   GTACGCATAGCTGATTTTTGAGCTGAGTTCTTTTCGTTTTGTTTAACGTTCAATTCAGCGCGTTTGATAGCTGATTTAATGTTTGCCAAT
NZ_CP035245.1 745261-753757    8031   GTACGCATAGCTGATTTTTGAGCTGAGTTCTTTTCGTTTTGTTTAACGTTCAATTCAGCGCGTTTGATAGCTGATTTAATGTTTGCCAAT
NZ_CP035259.1:745700-753548    7383   GTACGCATAGCTGATTTTTGAGCTGAGTTCTTTTCGTTTTGTTTAACGTTCAATTCAGCGCGTTTGATAGCTGATTTAATGTTTGCCAAT
NZ_CP035258.1:750874-758722    7383   GTACGCATAGCTGATTTTTGAGCTGAGTTCTTTTCGTTTTGTTTAACGTTCAATTCAGCGCGTTTGATAGCTGATTTAATGTTTGCCAAT
NZ_CP035260.1:726156-733994    7373   GTACGCATAGCTGATTTTTGAGCTGAGTTCTTTTCGTTTTGTTTAACGTTCAATTCAGCGCGTTTGATAGCTGATTTAATGTTTGCCAAT
NZ_CP035241.1:708296-716144    7383   GTACGCATAGCTGATTTTTGAGCTGAGTTCTTTTCGTTTTGTTTAACGTTCAATTCAGCGCGTTTGATAGCTGATTTAATGTTTGCCAAT
NZ_CP035261.1:779086-786938    7387   GTACGCATAGCTGATTTTTGAGCTGAGTTCTTTTCGTTTTGTTTAACGTTCAATTCAGCGCGTTTGATAGCTGATTTAATGTTTGCCAAT
NZ_CP035242.1:c1532236-1524385 7386   GTACGCATAGCTGATTTTTGAGCTGAGTTCTTTTCGTTTTGTTTAACGTTCAATTCAGCGCGTTTGATAGCTGATTTAATGTTTGCCAAT
NZ_CP035243.1:777729-785580    7386   GTACGCATAGCTGATTTTTGAGCTGAGTTCTTTTCGTTTTGTTTAACGTTCAATTCAGCGCGTTTGATAGCTGATTTAATGTTTGCCAAT
NZ_CP035237.1:c1311322-1303456 7401   GTACGCATAGCTGATTTTTGAGCTGAGTTCTTTTCGTTTTGTTTAACGTTCAATTCAGCGCGTTTGATAGCTGATTTAATGTTTGCCAAT
NZ_CP035246.1:c1302892-1295026 7401   GTACGCATAGCTGATTTTTGAGCTGAGTTCTTTTCGTTTTGTTTAACGTTCAATTCAGCGCGTTTGATAGCTGATTTAATGTTTGCCAAT
NZ_CP035247.1:c1301115-1293263 7387   GTACGCATAGCTGATTTTTGAGCTGAGTTCTTTTCGTTTTGTTTAACGTTCAATTCAGCGCGTTTGATAGCTGATTTAATGTTTGCCAAT
NZ_CP035238.1:738547-746413    7401   GTACGCATAGCTGATTTTTGAGCTGAGTTCTTTTCGTTTTGTTTAACGTTCAATTCAGCGCGTTTGATAGCTGATTTAATGTTTGCCAAT
NZ_CP035255.1:776866-784718    7387   GTACGCATAGCTGATTTTTGAGCTGAGTTCTTTTCGTTTTGTTTAACGTTCAATTCAGCGCGTTTGATAGCTGATTTAATGTTTGCCAAT
NZ_CP035256.1:807236-815102    7401   GTACGCATAGCTGATTTTTGAGCTGAGTTCTTTTCGTTTTGTTTAACGTTCAATTCAGCGCGTTTGATAGCTGATTTAATGTTTGCCAAT
NZ_CP038252.1:1151812-1159673  7326   GTACGCATAGCTGATTTTTGAGCTGAGTTCTTTTCGTTTTGTTTAACGTTCAATTCAGCGCGTTTGATAGCTGATTTAATGTTTGCCAAT
NZ_CP035254.1:793365-801213    7383   GTACGCATAGCTGATTTTTGAGCTGAGTTCTTTTCGTTTTGTTTAACGTTCAATTCAGCGCGTTTGATAGCTGATTTAATGTTTGCCAAT
NZ_CP035251.1:748733-756585    7387   GTACGCATAGCTGATTTTTGAGCTGAGTTCTTTTCGTTTTGTTTAACGTTCAATTCAGCGCGTTTGATAGCTGATTTAATGTTTGCCAAT
NZ_CP035252.1:745670-753536    7401   GTACGCATAGCTGATTTTTGAGCTGAGTTCTTTTCGTTTTGTTTAACGTTCAATTCAGCGCGTTTGATAGCTGATTTAATGTTTGCCAAT
NZ_CP035257.1:776317-784169    7387   GTACGCATAGCTGATTTTTGAGCTGAGTTCTTTTCGTTTTGTTTAACGTTCAATTCAGCGCGTTTGATAGCTGATTTAATGTTTGCCAAT
NZ_CP038253.1:758105-765957    7387   GTACGCATAGCTGATTTTTGAGCTGAGTTCTTTTCGTTTTGTTTAACGTTCAATTCAGCGCGTTTGATAGCTGATTTAATGTTTGCCAAT
NZ_CP035263.1:743050-750901    7386   GTACGCATAGCTGATTTTTGAGCTGAGTTCTTTTCGTTTTGTTTAACGTTCAATTCAGCGCGTTTGATAGCTGATTTAATGTTTGCCAAT
NZ_CP035262.1:741595-749460    7400   GTACGCATAGCTGATTTTTGAGCTGAGTTCTTTTCGTTTTGTTTAACGTTCAATTCAGCGCGTTTGATAGCTGATTTAATGTTTGCCAAT
NZ_CP035236.1:764915-772766    7386   GTACGCATAGCTGATTTTTGAGCTGAGTTCTTTTCGTTTTGTTTAACGTTCAATTCAGCGCGTTTGATAGCTGATTTAATGTTTGCCAAT
NZ_CP035253.1 767286-775134    7383   GTACGCATAGCTGATTTTTGAGCTGAGTTCTTTTCGTTTTGTTTAACGTTCAATTCAGCGCGTTTGATAGCTGATTTAATGTTTGCCAAT
NZ_CP035249.1:763080-770921    7376   GTACGCATAGCTGATTTTTGAGCTGAGTTCTTTTCGTTTTGTTTAACGTTCAATTCAGCGCGTTTGATAGCTGATTTAATGTTTGCCAAT

NC_003028.3:777000-787000      9625   GTTCTTACCTCCATATTTACTAACTATACCATTATATATGAAAACTTACGTTTTGACAAGGGAAAATTACTTTTTTAAGTAAATTTCATC
NZ_CP035248.1:765099-775111    9637   GTTCTTACCTCCATATTTACTAACTATACCATTATATATGAAAACTTACGTTTTGACAAGGGAAAATTACTTTTTTAAGTAAATTTCATC
NZ_CP035235.1 c1326834-1317894 8565   GTTCTTACCTCCATATTTACTAACTATACCATTATATATGAAAACTTACGTTTTGACAAGGGAAAATTATTTTTTTAAGTAAATTTCATC
NZ_CP035234.1:756284-765224    8565   GTTCTTACCTCCATATTTACTAACTATACCATTATATATGAAAACTTACGTTTTGACAAGGGAAAATTATTTTTTTAAGTAAATTTCATC
NZ_CP038251.1:750745-759686    8566   GTTCTTACCTCCATATTTACTAACTATACCATTATATATGAAAACTTACGTTTTGACAAGGGAAAATTATTTTTTTAAGTAAATTTCATC
NZ_CP035244.1:747023-755950    8552   GTTCTTACCTCCATATTTACTAACTATACCATTATATATGAAAACTTACGTTTTGACAAGGGAAAATTATTTTTTTAAGTAAATTTCATC
NZ_CP035264.1:800836-809336    8125   GTTCTTACCTCCATATTTACTAACTATACCATTATATATGAAAACTTACGTTTTGACAAGGGAAAATTACTTTTTTAAGTAAATTTCATC
NZ_CP035265.1:766128-774627    8124   GTTCTTACCTCCATATTTACTAACTATACCATTATATATGAAAACTTACGTTTTGACAAGGGAAAATTACTTTTTTAAGTAAATTTCATC
NZ_CP035245.1 745261-753757    8121   GTTCTTACCTCCATATTTACTAACTATACCATTATATATGAAAACTTACGTTTTGACAAGGGAAAATTACTTTTTTAAGTAAATTTCATC
NZ_CP035259.1:745700-753548    7473   GTTCTTACCTCCATATTTACTAACTATACCATTATATATGAAAACTTACGTTTTGACAAGGGAAAATTACTTTTTTAAGTAAATTTCATC
NZ_CP035258.1:750874-758722    7473   GTTCTTACCTCCATATTTACTAACTATACCATTATATATGAAAACTTACGTTTTGACAAGGGAAAATTACTTTTTTAAGTAAATTTCATC
NZ_CP035260.1:726156-733994    7463   GTTCTTACCTCCATATTTACTAACTATACCATTATATATGAAAACTTACGTTTTGACAAGGGAAAATTATTTTTTTAAGTAAATTTCATC
NZ_CP035241.1:708296-716144    7473   GTTCTTACCTCCATATTTACTAACTATACCATTATATATGAAAACTTACGTTTTGACAAGGGAAAATTACTTTTTTAAGTAAATTTCATC
NZ_CP035261.1:779086-786938    7477   GTTCTTACCTCCATATTTACTAACTATACCATTATATATGAAAACTTACGTTTTGACAAGGGAAAATTACTTTTTTAAGTAAATTTCATC
NZ_CP035242.1:c1532236-1524385 7476   GTTCTTACCTCCATATTTACTAACTATACCATTATATATGAAAACTTACGTTTTGACAAGGGAAAATTACTTTTTTAAGTAAATTTCATC
NZ_CP035243.1:777729-785580    7476   GTTCTTACCTCCATATTTACTAACTATACCATTATATATGAAAACTTACGTTTTGACAAGGGAAAATTACTTTTTTAAGTAAATTTCATC
NZ_CP035237.1:c1311322-1303456 7491   GTTCTTACCTCCATATTTACTAACTATACCATTATATATGAAAACTTACGTTTTGACAAGGGAAAATTACTTTTTTAAGTAAATTTCATC
NZ_CP035246.1:c1302892-1295026 7491   GTTCTTACCTCCATATTTACTAACTATACCATTATATATGAAAACTTACGTTTTGACAAGGGAAAATTATTTTTTTAAGTAAATTTCATC
NZ_CP035247.1:c1301115-1293263 7477   GTTCTTACCTCCATATTTACTAACTATACCATTATATATGAAAACTTACGTTTTGACAAGGGAAAATTATTTTTTTAAGTAAATTTCATC
NZ_CP035238.1:738547-746413    7491   GTTCTTACCTCCATATTTACTAACTATACCATTATATATGAAAACTTACGTTTTGACAAGGGAAAATTATTTTTTTAAGTAAATTTCATC
NZ_CP035255.1:776866-784718    7477   GTTCTTACCTCCATATTTACTAACTATACCATTATATATGAAAACTTACGTTTTGACAAGGGAAAATTATTTTTTTAAGTAAATTTCATC
NZ_CP035256.1:807236-815102    7491   GTTCTTACCTCCATATTTACTAACTATACCATTATATATGAAAACTTACGTTTTGACAAGGGAAAATTATTTTTTTAAGTAAATTTCATC
NZ_CP038252.1:1151812-1159673  7416   GTTCTTACCTCCATATTTACTAACTATACCATTATATATGAAAACTTACGTTTTGACAAGGGAAAATTATTTTTTTAAGTAAATTTCATC
NZ_CP035254.1:793365-801213    7473   GTTCTTACCTCCATATTTACTAACTATACCATTATATATGAAAACTTACGTTTTGACAAGGGAAAATTATTTTTTTAAGTAAATTTCATC
NZ_CP035251.1:748733-756585    7477   GTTCTTACCTCCATATTTACTAACTATACCATTATATATGAAAACTTACGTTTTGACAAGGGAAAATTATTTTTTTAAGTAAATTTCATC
NZ_CP035252.1:745670-753536    7491   GTTCTTACCTCCATATTTACTAACTATACCATTATATATGAAAACTTACGTTTTGACAAGGGAAAATTATTTTTTTAAGTAAATTTCATC
NZ_CP035257.1:776317-784169    7477   GTTCTTACCTCCATATTTACTAACTATACCATTATATATGAAAACTTACGTTTTGACAAGGGAAAATTATTTTTTTAAGTAAATTTCATC
NZ_CP038253.1:758105-765957    7477   GTTCTTACCTCCATATTTACTAACTATACCATTATATATGAAAACTTACGTTTTGACAAGGGAAAATTATTTTTTTAAGTAAATTTCATC
NZ_CP035263.1:743050-750901    7476   GTTCTTACCTCCATATTTACTAACTATACCATTATATATGAAAACTTACGTTTTGACAAGGGAAAATTACTTTTTTAAGTAAATTTCATC
NZ_CP035262.1:741595-749460    7490   GTTCTTACCTCCATATTTACTAACTATACCATTATATATGAAAACTTACGTTTTGACAAGGGAAAATTACTTTTTTAAGTAAATTTCATC
NZ_CP035236.1:764915-772766    7476   ATTCTTACCTCCATATTTACTAACTATACCATTATATATGAAAACTTACGTTTTGACAAGGGAAAATTACTTTTTTAAGTAAATTTCATC
NZ_CP035253.1 767286-775134    7473   GTTCTTACCTCCATATTTACTAACTATACCATTATATATGAAAACTTACGTTTTGACAAGGGAAAATTATTTTTTTAAGTAAATTTCATC
NZ_CP035249.1:763080-770921    7466   GTTCTTACCTCCATATTTACTAACTATACCATTATATATGAAAACTTACGTTTTGACAAGGGAAAATTACTTTTTTAAGTAAATTTCATC

NC_003028.3:777000-787000      9715   GATTTCATGGTTCTTGCTTTTATGAAGAATCACTTCTGCACGATTTCTGGTTGGTTCAATATAATTTTGCAGATTTGTGAGATTGATACT
NZ_CP035248.1:765099-775111    9727   GATTTCATGGTTCTTGCTTTTATGAAGAATCACTTCTGCACGATTTCTGGTTGGTTCAATATAATTTTGCAGATTTGTGAGATTGATACT
NZ_CP035235.1 c1326834-1317894 8655   GATTTCATGGTTCTTGCTTTTATGAAGAATCACTTCTGCACGATTTCTGGTTGGTTCAATATAATTTTGTAGATTTGTGAGATTGATACT
NZ_CP035234.1:756284-765224    8655   GATTTCATGGTTCTTGCTTTTATGAAGAATCACTTCTGCACGATTTCTGGTTGGTTCAATATAATTTTGTAGATTTGTGAGATTGATACT
NZ_CP038251.1:750745-759686    8656   GATTTCATGGTTCTTGCTTTTATGAAGAATCACTTCTGCACGATTTCTGGTTGGTTCAATATAATTTTGTAGATTTGTGAGATTGATACT
NZ_CP035244.1:747023-755950    8642   GATTTCATGGTTCTTGCTTTTATGAAGAATCACTTCTGCACGATTTCTGGTTGGTTCAATATAATTTTGTAGATTTGTGAGATTGATACT
NZ_CP035264.1:800836-809336    8215   GATTTCATGGTTCTTGCTTTTATGAAGAATCACTTCTGCACGATTTCTGGTTGGTTCAATATAATTTTGCAGATTTGTGAGATTGATACT
NZ_CP035265.1:766128-774627    8214   GATTTCATGGTTCTTGCTTTTATGAAGAATCACTTCTGCACGATTTCTGGTTGGTTCAATATAATTTTGCAGATTTGTGAGATTGATACT
NZ_CP035245.1 745261-753757    8211   GATTTCATGGTTCTTGCTTTTATGAAGAATCACTTCTGCACGATTTCTGGTTGGTTCAATATAATTTTGCAGATTTGTGAGATTGATACT
NZ_CP035259.1:745700-753548    7563   GATTTCATGGTTCTTGCTTTTATGAAGAATCACTTCTGCACGATTTCTGGTTGGTTCAATATAATTTTGCAGATTTGTGAGATTGATACT
NZ_CP035258.1:750874-758722    7563   GATTTCATGGTTCTTGCTTTTATGAAGAATCACTTCTGCACGATTTCTGGTTGGTTCAATATAATTTTGCAGATTTGTGAGATTGATACT
```

### Ozkan et al. Supplementary Figures

```
NZ_CP035260.1:726156-733994    7553   GATTTCATGGTTCTTGCTTTTATGAAGAATCACTTCTGCACGATTTCTGGTTGGTTCAATATAATTTTGTAGATTTGTGAGATTGATACT
NZ_CP035241.1:708296-716144    7563   GATTTCATGGTTCTTGCTTTTATGAAGAATCACTTCTGCACGATTTCTGGTTGGTTCAATATAATTTTGCAGATTTGTGAGATTGATACT
NZ_CP035261.1:779086-786938    7567   GATTTCATGGTTCTTGCTTTTATGAAGAATCACTTCTGCACGATTTCTGGTTGGTTCAATATAATTTTGCAGATTTGTGAGATTGATACT
NZ_CP035242.1:c1532236-1524385 7566   GATTTCATGGTTCTTGCTTTTATGAAGAATCACTTCTGCACGATTTCTGGTTGGTTCAATATAATTTTGCAGATTTGTGAGATTGATACT
NZ_CP035243.1:777729-785580    7566   GATTTCATGGTTCTTGCTTTTATGAAGAATCACTTCTGCACGATTTCTGGTTGGTTCAATATAATTTTGCAGATTTGTGAGATTGATACT
NZ_CP035237.1:c1311322-1303456 7581   GATTTCATGGTTCTTGCTTTTATGAAGAATCACTTCTGCACGATTTCTGGTTGGTTCAATATAATTTTGCAGATTTGTGAGATTGATACT
NZ_CP035246.1:c1302892-1295026 7581   GATTTCATGGTTCTTGCTTTTATGAAGAATCACTTCTGCACGATTTCTGGTTGGTTCAATATAATTTTGTAGATTTGTGAGATTGATACT
NZ_CP035247.1:c1301115-1293263 7567   GATTTCATGGTTCTTGCTTTTATGAAGAATCACTTCTGCACGATTTCTGGTTGGTTCAATATAATTTTGTAGATTTGTGAGATTGATACT
NZ_CP035238.1:738547-746413    7581   GATTTCATGGTTCTTGCTTTTATGAAGAATCACTTCTGCACGATTTCTGGTTGGTTCAATATAATTTTGTAGATTTGTGAGATTGATACT
NZ_CP035255.1:776866-784718    7567   GATTTCATGGTTCTTGCTTTTATGAAGAATCACTTCTGCACGATTTCTGGTTGGTTCAATATAATTTTGTAGATTTGTGAGATTGATACT
NZ_CP035256.1:807236-815102    7581   GATTTCATGGTTCTTGCTTTTATGAAGAATCACTTCTGCACGATTTCTGGTTGGTTCAATATAATTTTGTAGATTTGTGAGATTGATACT
NZ_CP038252.1:1151812-1159673  7506   GATTTCATGGTTCTTGCTTTTATGAAGAATCACTTCTGCACGATTTCTGGTTGGTTCAATATAATTTTGTAGATTTGTGAGATTGATACT
NZ_CP035254.1:793365-801213    7563   GATTTCATGGTTCTTGCTTTTATGAAGAATCACTTCTGCACGATTTCTGGTTGGTTCAATATAATTTTGTAGATTTGTGAGATTGATACT
NZ_CP035251.1:748733-756585    7567   GATTTCATGGTTCTTGCTTTTATGAAGAATCACTTCTGCACGATTTCTGGTTGGTTCAATATAATTTTGTAGATTTGTGAGATTGATACT
NZ_CP035252.1:745670-753536    7581   GATTTCATGGTTCTTGCTTTTATGAAGAATCACTTCTGCACGATTTCTGGTTGGTTCAATATAATTTTGTAGATTTGTGAGATTGATACT
NZ_CP035257.1:776317-784169    7567   GATTTCATGGTTCTTGCTTTTATGAAGAATCACTTCTGCACGATTTCTGGTTGGTTCAATATAATTTTGTAGATTTGTGAGATTGATACT
NZ_CP038253.1:758105-765957    7567   GATTTCATGGTTCTTGCTTTTATGAAGAATCACTTCTGCACGATTTCTGGTTGGTTCAATATAATTTTGTAGATTTGTGAGATTGATACT
NZ_CP035263.1:743050-750901    7566   GATTTCATGGTTCTTGCTTTTATGAAGAATCACTTCTGCACGATTTCTGGTTGGTTCAATATAATTTTGCAGATTTGTGAGATTGATACT
NZ_CP035262.1:741595-749460    7580   GATTTCATGGTTCTTGCTTTTATGAAGAATCACTTCTGCACGATTTCTGGTTGGTTCAATATAATTTTGCAGATTTGTGAGATTGATACT
NZ_CP035236.1:764915-772766    7566   GATTTCATGGTTCTTGCTTTTATGAAGAATCACTTCTGCACGATTTCTGGTTGGTTCAATATAATTTTGCAGATTTGTGAGATTGATACT
NZ_CP035253.1 767286-775134    7563   GATTTCATGGTTCTTGCTTTTATGAAGAATCACTTCTGCACGATTTCTGGTTGGTTCAATATAATTTTGTAGATTTGTGAGATTGATACT
NZ_CP035249.1:763080-770921    7556   GATTTCATGGTTCTTGCTTTTATGAAGAATCACTTCTGCACGATTTCTGGTTGGTTCAATATAATTTTGCAGATTTGTGAGATTGATACT

NC_003028.3:777000-787000      9805   GATCCAGACCTGATGGGCAAAGGACTCCACTTCCCCAATCGGCATCTGAGTAAAACGATAATAGTAGCTATCAGGGTCGTTTTGGGCTAG
NZ_CP035248.1:765099-775111    9817   GATCCAGACCTGATGGGCAAAGGACTCCACTTCCCCAATCGGCATCTGAGTAAAACGATAATAGTAGCTATCAGGGTCGTTTTGGGCTAG
NZ_CP035235.1 c1326834-1317894 8745   GGTCCAGACCTGATGGGCAAAGGCTTCCACTTCCCCAATCGGCATCTGAGTAAAACGATAATAGTAGCTATCAGAGTCGTTTTGGGCTAG
NZ_CP035234.1:756284-765224    8745   GGTCCAGACCTGATGGGCAAAGGCTTCCACTTCCCCAATCGGCATCTGAGTAAAACGATAATAGTAGCTATCAGAGTCGTTTTGGGCTAG
NZ_CP038251.1:750745-759686    8746   GGTCCAGACCTGATGGGCAAAGGCTTCCACTTCCCCAATCGGCATCTGAGTAAAACGATAATAGTAGCTATCAGAGTCGTTTTGGGCTAG
NZ_CP035244.1:747023-755950    8732   GGTCCAGACCTGATGGGCAAAGGCTTCCACTTCCCCAATCGGCATCTGAGTAAAACGATAATAGTAGCTATCAGAGTCGTTTTGGGCTAG
NZ_CP035264.1:800836-809336    8305   GATCCAGACCTGATGGGCAAAGGCTTCCACTTCCCCAATCGGCATCTGAGTAAAACGATAATAGTAGCTATCAGGGTCGTTTTGGGCTAG
NZ_CP035265.1:766128-774627    8304   GATCCAGACCTGATGGGCAAAGGCTTCCACTTCCCCAATCGGCATCTGAGTAAAACGATAATAGTAGCTATCAGGGTCGTTTTGGGCTAG
NZ_CP035245.1 745261-753757    8301   GGTCCAGACCTGATGGGCAAAGGACTCCACTTCCCCAATCGGCATCTGAGTAAAACGATAATAGTAGCTATCAGGGTCGTTTTGGGCTAG
NZ_CP035259.1:745700-753548    7653   GATCCAGACCTGATGGGCAAAGGACTCCACTTCCCCAATCGGCATCTGAGTAAAACGATAATAGTAGCTATCAGGGTCGTTTTGGGCTAG
NZ_CP035258.1:750874-758722    7653   GATCCAGACCTGATGGGCAAAGGACTCCACTTCCCCAATCGGCATCTGAGTAAAACGATAATAGTAGCTATCAGGGTCGTTTTGGGCTAG
NZ_CP035260.1:726156-733994    7643   GGTCCAGACCTGATGGGCAAAGGCTTCCACTTCCCCAATCGGCATCTGAGTAAAACGATAATAGTAGCTATCAGGGTCGTTTTGGGCTAG
NZ_CP035241.1:708296-716144    7653   GATCCAGACCTGATGGGCAAAGGACTCCACTTCCCCAATCGGCATCTGAGTAAAACGATAATAGTAGCTATCAGGGTCGTTTTGGGCTAG
NZ_CP035261.1:779086-786938    7657   GATCCAGACCTGATGGGCAAAGGACTCCACTTCCCCAATCGGCATCTGAGTAAAACGATAATAGTAGCTATCAGGGTCGTTTTGGGCTAG
NZ_CP035242.1:c1532236-1524385 7656   GATCCAGACCTGATGGGCAAAGGACTCCACTTCCCCAATCGGCATCTGAGTAAAACGATAATAGTAGCTATCAGGGTCGTTTTGGGCTAG
NZ_CP035243.1:777729-785580    7656   GATCCAGACCTGATGGGCAAAGGACTCCACTTCCCCAATCGGCATCTGAGTAAAACGATAATAGTAGCTATCAGGGTCGTTTTGGGCTAG
NZ_CP035237.1:c1311322-1303456 7671   GATCCAGACCTGATGGGCAAAGGACTCCACTTCCCCAATCGGCATCTGAGTAAAACGATAATAGTAGCTATCAGGGTCGTTTTGGGCTAG
NZ_CP035246.1:c1302892-1295026 7671   GGTCCAGACCTGATGGGCAAAGGCTTCCACTTCCCCAATCGGCATCTGAGTAAAACGATAATAGTAGCTATCAGGGTCGTTTTGGGCTAG
NZ_CP035247.1:c1301115-1293263 7657   GGTCCAGACCTGATGGGCAAAGGCTTCCACTTCCCCAATCGGCATCTGAGTAAAACGATAATAGTAGCTATCAGGGTCGTTTTGGGCTAG
NZ_CP035238.1:738547-746413    7671   GGTCCAGACCTGATGGGCAAAGGCTTCCACTTCCCCAATCGGCATCTGAGTAAAACGATAATAGTAGCTATCAGGGTCGTTTTGGGCTAG
NZ_CP035255.1:776866-784718    7657   GGTCCAGACCTGATGGGCAAAGGCTTCCACTTCCCCAATCGGCATCTGAGTAAAACGATAATAGTAGCTATCAGGGTCGTTTTGGGCTAG
NZ_CP035256.1:807236-815102    7671   GGTCCAGACCTGATGGGCAAAGGCTTCCACTTCCCCAATCGGCATCTGAGTAAAACGATAATAGTAGCTATCAGGGTCGTTTTGGGCTAG
NZ_CP038252.1:1151812-1159673  7596   GGTCCAGACCTGATGGGCAAAGGCTTCCACTTCCCCAATCGGCATCTGAGTAAAACGATAATAGTAGCTATCAGGGTCGTTTTGGGCTAG
NZ_CP035254.1:793365-801213    7653   GGTCCAGACCTGATGGGCAAAGGCTTCCACTTCCCCAATCGGCATCTGAGTAAAACGATAATAGTAGCTATCAGGGTCGTTTTGGGCTAG
NZ_CP035251.1:748733-756585    7657   GGTCCAGACCTGATGGGCAAAGGCTTCCACTTCCCCAATCGGCATCTGAGTAAAACGATAATAGTAGCTATCAGGGTCGTTTTGGGCTAG
NZ_CP035252.1:745670-753536    7671   GGTCCAGACCTGATGGGCAAAGGCTTCCACTTCCCCAATCGGCATCTGAGTAAAACGATAATAGTAGCTATCAGGGTCGTTTTGGGCTAG
NZ_CP035257.1:776317-784169    7657   GGTCCAGACCTGATGGGCAAAGGCTTCCACTTCCCCAATCGGCATCTGAGTAAAACGATAATAGTAGCTATCAGGGTCGTTTTGGGCTAG
NZ_CP038253.1:758105-765957    7657   GGTCCAGACCTGATGGGCAAAGGCTTCCACTTCCCCAATCGGCATCTGAGTAAAACGATAATAGTAGCTATCAGGGTCGTTTTGGGCTAG
NZ_CP035263.1:743050-750901    7656   GATCCAGACCTGATGGGCAAAGGACTCCACTTCCCCAATCGGCATCTGAGTAAAACGATAATAGTAGCTATCAGGGTCGTTTTGGGCTAG
NZ_CP035262.1:741595-749460    7670   GATCCAGACCTGATGGGCAAAGGACTCCACTTCCCCAATCGGCATCTGAGTAAAACGATAATAGTAGCTATCAGGGTCGTTTTGGGCTAG
NZ_CP035236.1:764915-772766    7656   GATCCAGACCTGATGGGCAAAGGACTCCACTTCCCCAATCGGCATCTGAGTAAAACGATAATAGTAGCTATCAGGGTCGTTTTGGGCTAG
NZ_CP035253.1 767286-775134    7653   GGTCCAGACCTGATGGGCAAAGGCTTCCACTTCCCCAATCGGCATCTGAGTAAAACGATAATAGTAGCTATCAGGGTCGTTTTGGGCTAG
NZ_CP035249.1:763080-770921    7646   GATCCAGACCTGATGGGCAAAGGACTCCACTTCCCCAATCGGCATCTGAGTAAAACGATAATAGTAGCTATCAGGGTCGTTTTGGGCTAG

NC_003028.3:777000-787000      9895   ACTCAGCATTTTCAAGAAACGGTCCAGATACCAACTTTCAATATCATCCACTCCAGCATCTACATAGATGGAAAAGTCAAAGAAGTCAGT
NZ_CP035248.1:765099-775111    9907   ACTCAGCATTTTCAAGAAACGGTCCAGATACCAACTTTCAATATCATCCACTCCAGCATCTACATAGATGGAAAAGTCAAAGAAGTCAGT
NZ_CP035235.1 c1326834-1317894 8835   ACTCAGCATTTTCAAGAAACGGTCCAGATACCAACTTTCAATATCATCCACTCCAGCATCTACATAGATGGAAAAGTCAAAGAAGTCAGT
NZ_CP035234.1:756284-765224    8835   ACTCAGCATTTTCAAGAAACGGTCCAGATACCAACTTTCAATATCATCCACTCCAGCATCTACATAGATGGAAAAGTCAAAGAAGTCAGT
NZ_CP038251.1:750745-759686    8836   ACTCAGCATTTTCAAGAAACGGTCCAGATACCAACTTTCAATATCATCCACTCCAGCATCTACATAGATGGAAAAGTCAAAGAAGTCAGT
NZ_CP035244.1:747023-755950    8822   ACTCAGCATTTTCAAGAAACGGTCCAGATACCAACTTTCAATATCATCCACTCCAGCATCTACATAGATGGAAAAGTCAAAGAAGTCAGT
NZ_CP035264.1:800836-809336    8395   ACTCAGCATTTTCAAGAAACGGTCCAGATACCAACTTTCAATATCATCCACTCCAGCATCTACATAGATGGAAAAGTCAAAGAAGTCAGT
NZ_CP035265.1:766128-774627    8394   ACTCAGCATTTTCAAGAAACGGTCCAGATACCAACTTTCAATATCATCCACTCCAGCATCTACATAGATGGAAAAGTCAAAGAAGTCAGT
NZ_CP035245.1 745261-753757    8391   GCTCAGCATTTTCAAGAAACGGTCCAGATACCAACTTTCAATATCATCCACTCCAGCATCTACATAGATGGAAAAGTCAAAGAAGTCAGT
NZ_CP035259.1:745700-753548    7743   GCTCAGCATTTTCAAGAAACGGTCCAGATACCAACTTTCAATATCATCCACTCCAGCATCTACATAGATGGAAAAGTCAAAGAAGTCAGT
NZ_CP035258.1:750874-758722    7743   GCTCAGCATTTTCAAGAAACGGTCCAGATACCAACTTTCAATATCATCCACTCCAGCATCTACATAGATGGAAAAGTCAAAGAAGTCAGT
NZ_CP035260.1:726156-733994    7733   GCTCAGCATTTTCAAGAAACGGTCCAGATACCAACTTTCAATATCATCCACTCCAGCATCTACATAGATGGAAAAGTCAAAGAAGTCAGT
NZ_CP035241.1:708296-716144    7743   GCTCAGCATCTTCAAGAAACGGTCCAGATACCAACTTTCAATATCATCCACTCCAGCATCTACATAGATGGAAAAGTCAAAGAAGTCAGT
NZ_CP035261.1:779086-786938    7747   GCTCAGCATTTTCAAGAAACGGTCCAGATACCAACTTTCAATATCATCCACTCCAGCATCTACATAGATGGAAAAGTCAAAGAAGTCAGT
NZ_CP035242.1:c1532236-1524385 7746   GCTCAGCATTTTCAAGAAACGGTCCAGATACCAACTTTCAATATCATCCACTCCAGCATCTACATAGATGGAAAAGTCAAAGAAGTCAGT
NZ_CP035243.1:777729-785580    7746   GCTCAGCATTTTCAAGAAACGGTCCAGATACCAACTTTCAATATCATCCACTCCAGCATCTACATAGATGGAAAAGTCAAAGAAGTCAGT
NZ_CP035237.1:c1311322-1303456 7761   GCTCAGCATTTTCAAGAAACGGTCCAGATACCAACTTTCAATATCATCCACTCCAGCATCTACATAGATGGAAAAGTCAAAGAAGTCAGT
NZ_CP035246.1:c1302892-1295026 7761   ACTCAGCATTTTCAAGAAACGGTCCAGATACCAACTTTCAATATCATCCACTCCAGCATCTACATAGATGGAAAAGTCAAAGAAGTCAGT
NZ_CP035247.1:c1301115-1293263 7747   ACTCAGCATTTTCAAGAAACGGTCCAGATACCAACTTTCAATATCATCCACTCCAGCATCTACATAGATGGAAAAGTCAAAGAAGTCAGT
NZ_CP035238.1:738547-746413    7761   ACTCAGCATTTTCAAGAAACGGTCCAGATACCAACTTTCAATATCATCCACTCCAGCATCTACATAGATGGAAAAGTCAAAGAAGTCAGT
NZ_CP035255.1:776866-784718    7747   ACTCAGCATTTTCAAGAAACGGTCCAGATACCAACTTTCAATATCATCCACTCCAGCATCTACATAGATGGAAAAGTCAAAGAAGTCAGT
NZ_CP035256.1:807236-815102    7761   ACTCAGCATTTTCAAGAAACGGTCCAGATACCAACTTTCAATATCATCCACTCCAGCATCTACATAGATGGAAAAGTCAAAGAAGTCAGT
NZ_CP038252.1:1151812-1159673  7686   ACTCAGCATTTTCAAGAAACGGTCCAGATACCAACTTTCAATATCATCCACTCCAGCATCTACATAGATGGAAAAGTCAAAGAAGTCAGT
NZ_CP035254.1:793365-801213    7743   ACTCAGCATTTTCAAGAAACGGTCCAGATACCAACTTTCAATATCATCCACTCCAGCATCTACATAGATGGAAAAGTCAAAGAAGTCAGT
NZ_CP035251.1:748733-756585    7747   ACTCAGCATTTTCAAGAAACGGTCCAGATACCAACTTTCAATATCATCCACTCCAGCATCTACATAGATGGAAAAGTCAAAGAAGTCAGT
NZ_CP035252.1:745670-753536    7761   ACTCAGCATTTTCAAGAAACGGTCCAGATACCAACTTTCAATATCATCCACTCCAGCATCTACATAGATGGAAAAGTCAAAGAAGTCAGT
NZ_CP035257.1:776317-784169    7747   ACTCAGCATTTTCAAGAAACGGTCCAGATACCAACTTTCAATATCATCCACTCCAGCATCTACATAGATGGAAAAGTCAAAGAAGTCAGT
NZ_CP038253.1:758105-765957    7747   ACTCAGCATTTTCAAGAAACGGTCCAGATACCAACTTTCAATATCATCCACTCCAGCATCTACATAGATGGAAAAGTCAAAGAAGTCAGT
NZ_CP035263.1:743050-750901    7746   GCTCAGCATTTTCAAGAAACGGTCCAGATACCAACTTTCAATATCATCCACTCCAGCATCTACATAGATGGAAAAGTCAAAGAAGTCAGT
NZ_CP035262.1:741595-749460    7760   GCTCAGCATTTTCAAGAAACGGTCCAGATACCAACTTTCAATATCATCCACTCCAGCATCTACATAGATGGAAAAGTCAAAGAAGTCAGT
NZ_CP035236.1:764915-772766    7746   GCTCAGCATTTTCAAGAAACGGTCCAGATACCAACTTTCAATATCATCCACTCCAGCATCTACATAGATGGAAAAGTCAAAGAAGTCAGT
NZ_CP035253.1 767286-775134    7743   ACTCAGCATTTTCAAGAAACGGTCCAGATACCAACTTTCAATATCATCCACTCCAGCATCTACATAGATGGAAAAGTCAAAGAAGTCAGT
NZ_CP035249.1:763080-770921    7736   GCTCAGCATTTTCAAGAAACGGTCCAGATACCAACTTTCAATATCATCCACTCCAGCATCTACATAGATGGAAAAGTCAAAGAAGTCAGT

NC_003028.3:777000-787000      9985   GATATAGAGACGATCGT
NZ_CP035248.1:765099-775111    9997   GATATAGAGACGATCGT
NZ_CP035235.1 c1326834-1317894 8925   GATATAGAGACGATCGT
NZ_CP035234.1:756284-765224    8925   GATATAGAGACGATCGT
```

```
NZ_CP038251.1:750745-759686    8926   GATATAGAGACGATCGT
NZ_CP035244.1:747023-755950    8912   GATATAGAGACGATCGT
NZ_CP035264.1:800836-809336    8485   GATATAGAGACGATCGT
NZ_CP035265.1:766128-774627    8484   GATATAGAGACGATCGT
NZ_CP035245.1 745261-753757    8481   GATATAGAGACGATCGT
NZ_CP035259.1:745700-753548    7833   GATATAGAGACGATCGT
NZ_CP035258.1:750874-758722    7833   GATATAGAGACGATCGT
NZ_CP035260.1:726156-733994    7823   GATATAGAGACGATCGT
NZ_CP035241.1:708296-716144    7833   GATATAGAGACGATCGT
NZ_CP035261.1:779086-786938    7837   GATATAGAGACGATCGT
NZ_CP035242.1:c1532236-1524385 7836   GATATAGAGACGATCGT
NZ_CP035243.1:777729-785580    7836   GATATAGAGACGATCGT
NZ_CP035237.1:c1311322-1303456 7851   GATATAGAGACGATCGT
NZ_CP035246.1:c1302892-1295026 7851   GATATAGAGACGATCGT
NZ_CP035247.1:c1301115-1293263 7837   GATATAGAGACGATCGT
NZ_CP035238.1:738547-746413    7851   GATATAGAGACGATCGT
NZ_CP035255.1:776866-784718    7837   GATATAGAGACGATCGT
NZ_CP035256.1:807236-815102    7851   GATATAGAGACGATCGT
NZ_CP038252.1:1151812-1159673  7776   GATATAGAGACGATCGT
NZ_CP035254.1:793365-801213    7833   GATATAGAGACGATCGT
NZ_CP035251.1:748733-756585    7837   GATATAGAGACGATCGT
NZ_CP035252.1:745670-753536    7851   GATATAGAGACGATCGT
NZ_CP035257.1:776317-784169    7837   GATATAGAGACGATCGT
NZ_CP038253.1:758105-765957    7837   GATATAGAGACGATCGT
NZ_CP035263.1:743050-750901    7836   GATATAGAGACGATCGT
NZ_CP035262.1:741595-749460    7850   GATATAGAGACGATCGT
NZ_CP035236.1:764915-772766    7836   GATATAGAGACGATCGT
NZ_CP035253.1 767286-775134    7833   GATATAGAGACGATCGT
NZ_CP035249.1:763080-770921    7826   GATATAGAGACGATCGT
```

**Supplementary Figure S9:** Alignment of 33 *S. pneumoniae* genomes segments to illustrate genomic arrangements pictured in Figure 4. In addition to the insertion sequence that creates the promoter for SP_0835as (underlined), there are several other insertions/rearrangements in this locus across the 33 genomes assessed.
